# Discovery of an AKT Degrader with Prolonged Inhibition of Downstream Signaling

**DOI:** 10.1101/848887

**Authors:** Inchul You, Emily C. Erickson, Katherine A. Donovan, Nicholas A. Eleuteri, Eric S. Fischer, Nathanael S. Gray, Alex Toker

## Abstract

The PI3K/AKT signaling cascade is one of the most commonly dysregulated pathways in cancer, with over half of tumors exhibiting aberrant AKT activation. Although potent small molecule AKT inhibitors have entered clinical trials, robust and durable therapeutic responses have not been observed. As an alternative strategy to target AKT, we report the development of INY-03-041, a pan-AKT degrader consisting of the ATP-competitive AKT inhibitor GDC-0068 conjugated to lenalidomide, a recruiter of the E3 ubiquitin ligase substrate adaptor Cereblon (CRBN). INY-03-041 induced potent degradation of all three AKT isoforms and displayed enhanced anti-proliferative effects relative to GDC-0068. Notably, INY-03-041 promoted sustained AKT degradation and inhibition of downstream signaling effects for up to 96 hours, even after compound washout. Our findings indicate that AKT degradation may confer prolonged pharmacological effects compared to inhibition, and highlight the potential advantages of AKT-targeted degradation.

## INTRODUCTION

The serine/threonine kinase AKT is a central component of the phosphoinositide 3-kinase (PI3K) signaling cascade and is a key regulator of critical cellular processes, including proliferation, survival and metabolism (Manning and Cantley, 2007). The AKT protein kinase family is comprised of three highly homologous isoforms, AKT1, AKT2, and AKT3 that possess both redundant functions and isoform-specific activities (Toker, 2012). Hyperactivation of AKT, due to gain-of-function mutations or amplification of oncogenes (receptor tyrosine kinases, PI3K) or inactivation of tumor suppressor genes (PTEN, INPP4B, PHLPP), is one of the most common molecular perturbations in cancer and promotes malignant phenotypes associated with tumor initiation and progression (Cantley and Neel, 1999; Fruman et al., 2017; Shayesteh et al., 1999). Thus, AKT is an attractive therapeutic target and significant efforts have been made to develop AKT targeted therapies (Brown and Banerji, 2017).

Current strategies to target AKT have focused on ATP-competitive, allosteric, and covalent inhibitors. Several ATP-competitive inhibitors, such as ipatasertib (GDC-0068) and capivasertib (AZD5363), are currently under clinical investigation in phase II and III studies (Oliveira et al., 2019; Turner et al., 2019). However, these inhibitors suffer from a lack of selectivity among the AGC kinase family, and this may limit their clinical efficacy or tolerability (Huck and Mochalkin, 2017). By contrast, allosteric inhibitors which target the pleckstrin homology (PH) domain, such as MK-2206 and miransertib (ARQ-092), exhibit a high degree of specificity towards AKT, but either lack significant efficacy in the clinic or require further clinical evaluation (Do et al., 2015; Keppler-Noreuil et al., 2019). Covalent allosteric inhibitors which target AKT at Cys296 and Cys310 have been reported, but have not advanced to clinical trials (Uhlenbrock et al., 2019).

An alternative pharmacological approach to inhibiting AKT activity is to directly reduce cellular AKT protein levels via targeted protein degradation. Heterobifunctional degraders, also known as PROTACs (proteolysis targeting chimeras), consist of a moiety that binds to an E3 ubiquitin ligase chemically linked to a second moiety that engages a target protein, thereby recruiting the E3 ligase into close proximity to the target protein to induce its ubiquitination and subsequent proteasomal degradation (Winter et al., 2015). Several advantages of degraders over inhibitors have been reported, which include enhancing the selectivity of multi-targeted inhibitors (CDK9) (Olson et al., 2018), abrogating non-kinase dependent functions (FAK) (Cromm et al., 2018), and overcoming resistance mutations (BTK) (Dobrovolsky et al., 2018). Because the pharmacological effects of degraders depend on the re-synthesis rate of the targeted protein rather than sustained target occupancy, small molecule degraders may also have significantly prolonged effects in comparison to reversible inhibitors. However, while the potential for degraders to achieve an extended pharmacological duration of action has been noted, there have been no reported degraders to date that highlight such a feature (Churcher, 2018). Given the long half-life of AKT (Basso et al., 2002) and its importance in cancer etiology and progression, AKT is an attractive protein to target for degradation.

Here, we report the development of INY-03-041, a pan-AKT degrader that induces potent degradation of all three AKT isoforms for an extended period of time. Through cell-based assays, we demonstrate that INY-03-041 exhibits more potent and prolonged effects on downstream signaling than GDC-0068, which may explain its enhanced anti-proliferative effects in comparison to the parent catalytic inhibitor. Our data demonstrate that AKT degradation has more durable pharmacological effects than AKT inhibition, which highlights a potentially novel aspect of degrader pharmacology.

## RESULTS

### Design and Development of the Pan-AKT Degrader INY-03-041

To develop an AKT-targeting heterobifunctional degrader, we designed compounds based on GDC-0068, the most advanced AKT inhibitor in clinical trials (Oliveira et al., 2019). The cocrystal structure of GDC-0068 bound to AKT1 (PDB ID: 4EKL) revealed that the isopropylamine is solvent-exposed, suggesting that the amine could serve as a suitable attachment site for linkers without adversely affecting affinity to AKT (**Figure 1A**). A ten hydrocarbon linker was used to conjugate GDC-0068 with lenalidomide to generate INY-03-041 (**Figure 1B**).

**Figure 1.**
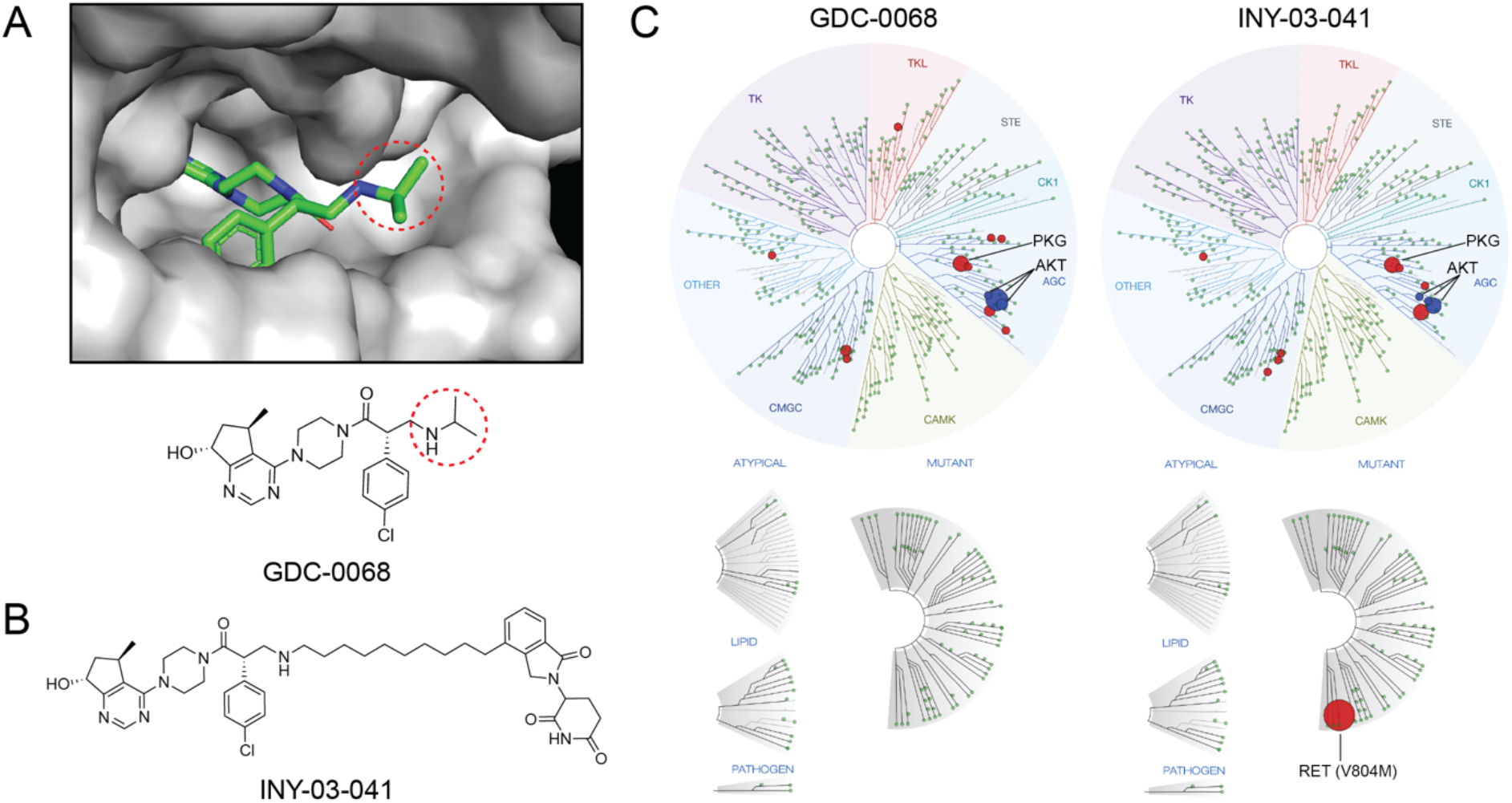
Design and Development of INY-03-041. (A) Co-crystal structure of GDC-0068 (green) bound to AKT1 (gray, PDB: 4EKL) revealing solvent-exposed isopropylamine (circled) where linker was attached. (B) Chemical structure of INY-03-041. (C) TREE*spot* visualization of the biochemical kinome selectivity profile of GDC-0068 and INY-03-041 (1 μM). AKT isoforms are highlighted in blue, while all other inhibited kinases are highlighted in red (Table S2).

To verify that conjugation of the linker and lenalidomide did not affect the ability of INY-03-041 to bind to AKT, INY-03-041 was tested in a commercially-available fluorescence resonance energy transfer (FRET)-based assay (Invitrogen, Z’-Lyte) for AKT1, AKT2, and AKT3 inhibition (**Table S1**). INY-03-041 had similar inhibitory activity against AKT1 (IC_50_ = 2.0 nM), AKT2 (IC_50_ = 6.8 nM), and AKT3 (IC_50_ = 3.5 nM) as GDC-0068 (IC_50_s for AKT1, 2, and 3 = 5, 18, and 8 nM, respectively) (Blake et al., 2012), demonstrating that INY-03-041 retained comparable biochemical affinity to all three AKT isoforms as its parent inhibitor. In addition, we evaluated the biochemical selectivity of INY-03-041 against a panel of 468 kinases at 1 μM (KINOMEscan) and observed that INY-03-041 had a similar selectivity profile as GDC-0068 (**Figure 1C**). Although INY-03-041 scored as a strong binder of RET (V804M) in the KINOMEscan assay, this was confirmed to be a false positive, as its biochemical IC_50_ was determined to be >10000 nM (Invitrogen, LanthaScreen) (**Tables S1 and S2**).

### INY-03-041 is a potent and highly selective pan-AKT degrader

After verifying that INY-03-041 engaged AKT biochemically, we sought to characterize its degradation activity in cells. We first chose to evaluate the AKT degrader in the triple negative breast cancer cell line MDA-MB-468 due to their high expression of all three AKT isoforms. We found that INY-03-041 induced potent degradation of all three AKT isoforms in a dosedependent manner after 12 hour treatment, with maximal degradation observed between 100 to 250 nM (**Figure 2A**). At concentrations of 500 nM and greater, we observed diminished AKT degradation, consistent with the hook effect, in which independent engagement of AKT and CRBN by INY-03-041 prevents formation of a productive ternary complex (An and Fu, 2018). Treatment of MDA-MB-468 cells with 250 nM of INY-03-041 over time revealed partial degradation of all AKT isoforms within 4 hours and progressive loss of AKT abundance out to 24 hours (**Figure 2B**).

**Figure 2.**
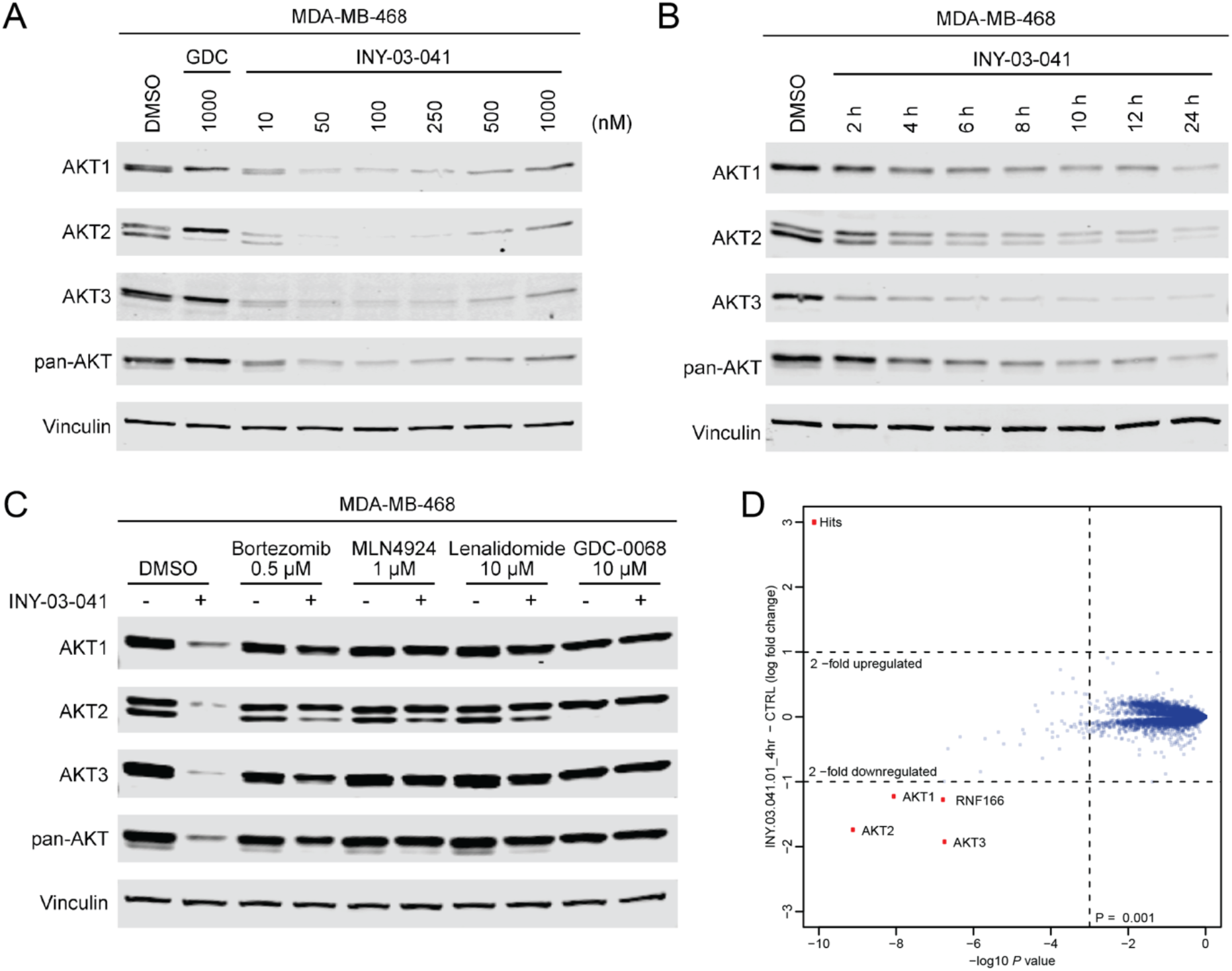
INY-03-041 induces potent degradation of AKT isoforms dependent on CRBN, neddylation, and the proteasome. (A) Immunoblots for AKT1, AKT2, AKT3, pan-AKT, and Vinculin in MDA-MB-468 cells after 12 hour treatment with DMSO, GDC-0068 (GDC), or INY-03-041 at the concentrations indicated (n=4). (B) Immunoblots for AKT1, AKT2, AKT3, pan-AKT, and Vinculin in MDA-MB-468 cells after treatment with INY-03-041 (250 nM) at indicated times or DMSO (24 h) (n=4). (C) Immunoblots for AKT1, AKT2, AKT3, pan-AKT, and Vinculin after 12 hour co-treatment of MDA-MB-468 cells with DMSO, bortezomib (0.5 μM), MLN-4924 (1 μM), lenalidomide (Len, 10 μM), or GDC-0068 (GDC, 10 μM) and either INY-03-041 (250 nM) or DMSO (n=4). (D) Scatterplot depicts the change in relative protein abundance of INY-03-041 (250 nM, 4 hour) treated MOLT4 cells compared to DMSO vehicle control treated cells. Protein abundance measurements were made using TMT quantitative mass spectrometry and significant changes were assessed by moderated t-test as implemented in the limma package (Ritchie et al., 2015). The log2 fold change (log_2_ FC) is shown on the y-axis and negative log10 p value (−log_10_ p value) on the x-axis for three independent biological replicates of each treatment. See also Table S3.

To ensure that INY-03-041-induced AKT degradation was dependent on CRBN, we synthesized INY-03-112, a negative control compound with an N-methylated glutarimide that substantially weakens CRBN binding (**Figure S1A**) (Brand et al., 2018). INY-03-112 did not induce potent degradation of any AKT isoform (**Figure S1B**), demonstrating that INY-03-041-induced AKT degradation was CRBN-dependent. Furthermore, co-treatment of INY-03-041 with bortezomib, a proteasome inhibitor, or MLN-4924, a NEDD8-activating enzyme inhibitor that prevents neddylation required for the function of cullin RING ligases such as CRL4^CRBN^ (Soucy et al., 2009), prevented AKT destabilization, indicating that degradation was dependent on the ubiquitin-proteasome system (**Figure 2C**). Finally, we co-treated INY-03-041 with excess quantities of either GDC-0068 or lenalidomide to compete for binding to AKT or CRBN, respectively, both of which prevented AKT degradation, demonstrating that engagement to both AKT and CRBN are required for INY-03-041-induced AKT degradation (**Figure 2C**).

To broadly assess degrader selectivity, MOLT4 cells, a cell line that is amenable to proteomics and expresses all three AKT isoforms, were treated with 250 nM of INY-03-041 for 4 hours and an unbiased, multiplexed mass spectrometry (MS)-based proteomic analysis was performed (Donovan et al., 2018). This analysis identified significant downregulation of all three AKT isoforms, as well as RNF166, a ring-finger protein known to be degraded by lenalidomide (**Figure 2D**) (Kronke et al., 2015). Although INY-03-041 exhibited potent *in vitro* inhibition of S6K1 (IC_50_ = 37.3 nM) and PKG1 (IC_50_ = 33.2 nM), both of which are known off-targets of GDC-0068, no downregulation of either kinases was observed in the proteomics (**Table S3**). Further immunoblot analysis confirmed that INY-03-041 did not induce S6K1 degradation (**Figure S2**).

As CRBN-targeting degraders often destabilize zinc finger proteins, and because we observed IMiD-induced protein degradation of RNF166, we examined whether INY-03-041 affected protein abundance levels of Ikaros (IKZF1) and Aiolos (IKZF3), well-established targets of lenalidomide (Kronke et al., 2014). Immunoblot analysis revealed weak IKZF1 and IKZF3 degradation after 24 hours of drug treatment, albeit at relatively high concentrations of 500 nM or greater, indicating that INY-03-041 is primarily a selective degrader for AKT (**Figure S2**).

### INY-03-041 exhibits enhanced anti-proliferative effects compared to GDC-0068

As AKT has well-characterized functions in regulating cell proliferation, we next compared the anti-proliferative effects of AKT degradation and inhibition using growth rate inhibition (GR) to account for variation in division rates among cells, as this can confound other drug response metrics, such as IC_50_ values (Hafner et al., 2017). In a panel of cell lines with PI3K pathway mutations that have been reported to be sensitive (ZR-75-1, T47D, LNCaP, and MCF-7) and insensitive (MDA-MB-468 and HCC1937) to AKT inhibition (**Table S4**) (Lin et al., 2013), we found that INY-03-041 was most potent in ZR-75-1 cells (GR_50_ = 16 nM), with a 14-fold increased potency compared to GDC-0068 (GR_50_ = 229 nM). The anti-proliferative effect of INY-03-041 was degradation-dependent, as INY-03-112 was significantly less potent (GR_50_ = 413 nM) than INY-03-041 and had a comparable GR_50_ value to GDC-0068 (**Figure 3A**; **Table S5**). Similar trends were seen in the other cell lines sensitive to AKT inhibition, with 8-to 14-fold lower GR_50_ values for INY-03-041 in comparison to GDC-0068 (**Figures 3A-D**; **Table S5**). In addition, lenalidomide, used as a control for RNF166, IKZF1, and IKZF3 degradation, did not have strong anti-proliferative effects, suggesting that the enhanced anti-proliferative effects were due to AKT degradation (**Figures 3A-D**).

**Figure 3.**
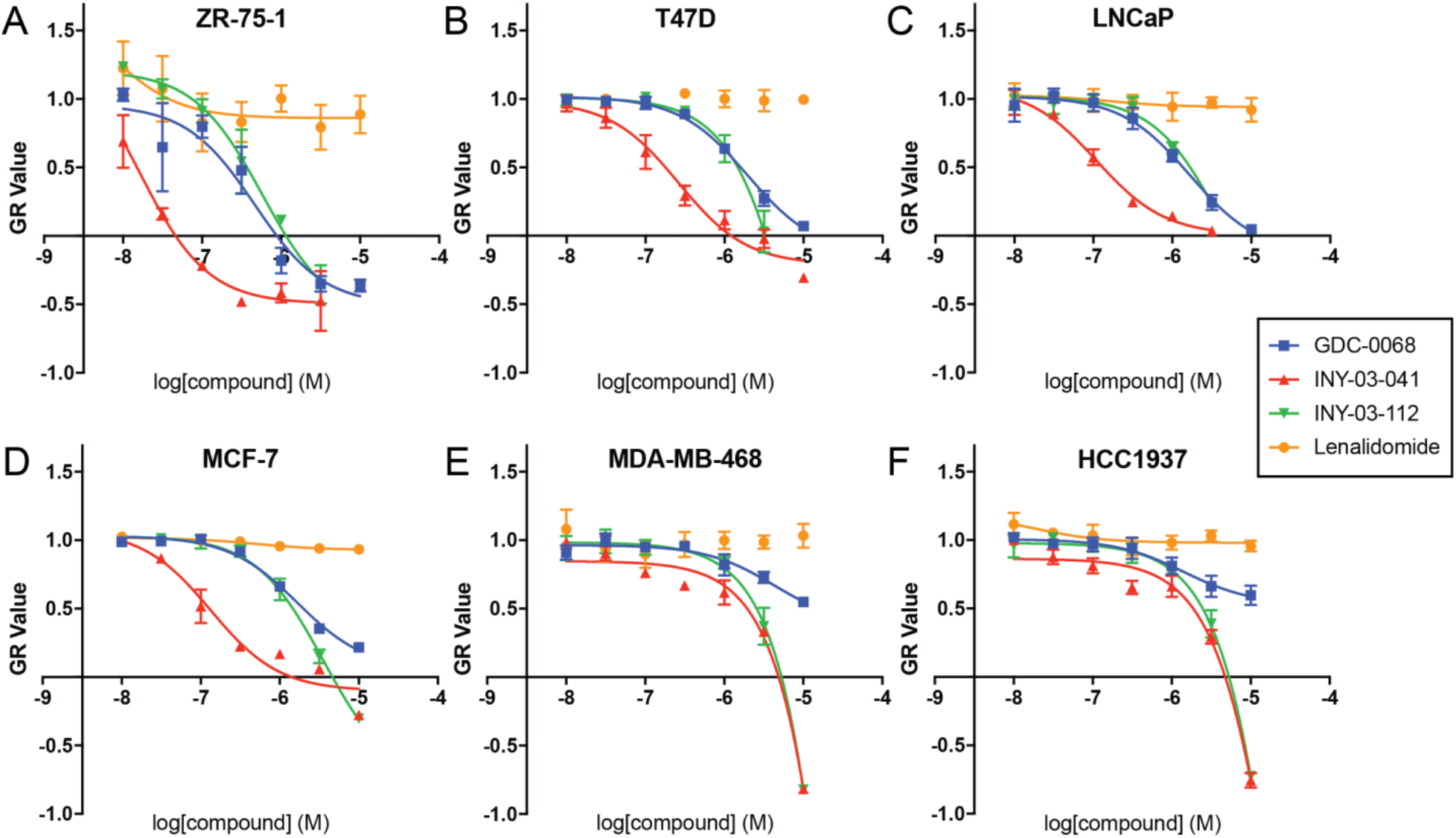
INY-03-041 induces enhanced anti-proliferative effects compared to GDC-0068. GR values across concentrations in (A) ZR-75-1, (B) T47D, (C) LNCaP, (D) MCF-7, (E) MDA-MB-468, and (F) HCC1937 cells after 72 hour treatment with GDC-0068 (blue), INY-03-041 (red), INY-03-112 (green), and lenalidomide (orange). Error bars represent standard deviation of three technical replicates.

While INY-03-041 displayed enhanced anti-proliferative effects compared to GDC-0068 in MDA-MB-468 and HCC1937 cells, there were no apparent differences in GR_50_ values between INY-03-041 and INY-03-112, its non-CRBN binding control (**Figures 3E** and **3F**; **Table S5**). Thus, the anti-proliferative effects of INY-03-041 in these cell lines were likely due to off-target effects unrelated to AKT degradation that manifest at elevated concentrations of INY-03-041 and INY-03-112. This is consistent with previous studies reporting resistance of MDA-MB-468 and HCC1937 to AKT inhibition (Lin et al., 2013), and suggest that AKT degradation has similar phenotypic effects as AKT inhibition in these cell lines. Overall, the data show that INY-03-041 suppresses proliferation more potently than GDC-0068, and highlight the potential therapeutic value of targeted AKT degradation.

### INY-03-041 suppresses downstream signaling more potently than GDC-0068

Given the enhanced anti-proliferative effects of INY-03-041 compared to GDC-0068, we sought to compare their effects on downstream AKT signaling. In T47D cells, which were highly sensitive to INY-03-041 in terms of anti-proliferation, we confirmed that INY-03-041 induced potent AKT degradation, with no detectable levels of all three AKT isoforms observed after a 24 hour treatment with 250 nM of INY-03-041 (**Figure S3A**). Moreover, INY-03-041 treatment resulted in robust and dose-dependent downregulation of PRAS40 phosphorylation (pPRAS40), a well-established substrate of AKT (Wang et al., 2012) (**Figure 4A**). While 250 nM of INY-03-041 significantly reduced pPRAS40 levels, up to 1 μM of GDC-0068 was needed to observe comparable effects (**Figure 4A**). To test whether these effects were cell-type specific, we also compared the effects of INY-03-041 and GDC-0068 in MDA-MB-468 cells. Similar to what was observed in T47D cells, INY-03-041 significantly reduced levels of pPRAS40 at 250 nM, while an equivalent dose of GDC-0068 did not noticeably affect pPRAS40 (**Figure 4A**). Thus, INY-03-041 suppresses downstream AKT signaling more potently than GDC-0068.

**Figure 4.**
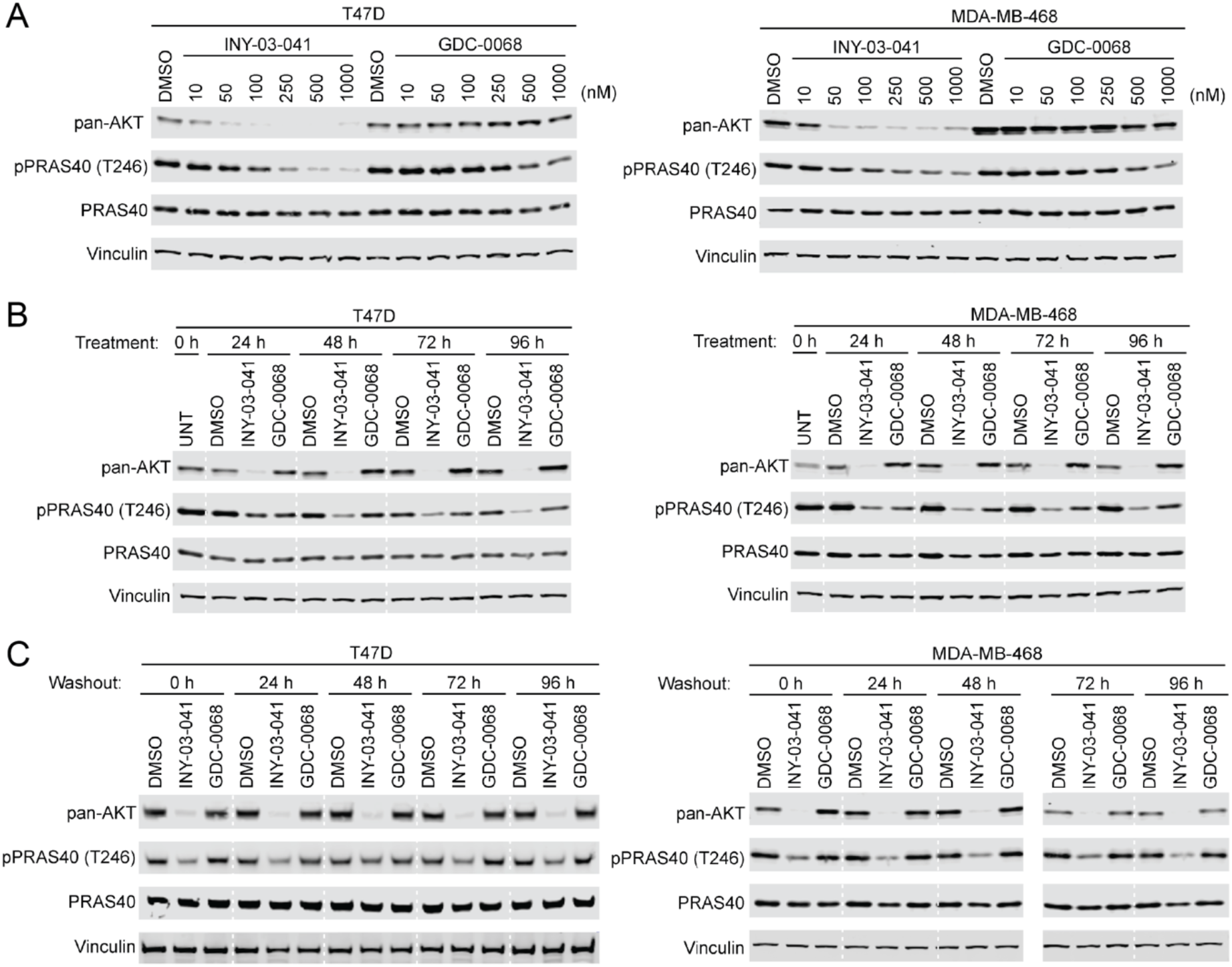
INY-03-041 exhibits more potent and durable downstream signaling effects than GDC-0068. (A) Immunoblots of pan-AKT, phospho-PRAS4O (T246), total PRAS40, and Vinculin after treating T47D or MDA-MB-468 cells for 24 hours with DMSO, INY-03-041, or GDC-0068 at the concentrations indicated (n=3). (B) Immunoblots of pan-AKT, phospho-PRAS40 (T246), total PRAS40, and Vinculin after treatment of T47D or MDA-MB-468 cells with 250 nM of INY-03-041 or GDC-0068 at time points indicated (n=3). (C) Immunoblots of pan-AKT, phospho-PRAS40 (T246), total PRAS40, and Vinculin in T47D or MDA-MB-468 cells treated for 12 hours with INY-03-041 or GDC-0068 (250 nM), followed by washout for indicated times (n=4). Solid vertical white line indicates samples run on separate gels.

Notably, we also found that INY-03-041 promoted sustained destabilization of all three AKT isoforms for at least 96 hours after treatment with 250 nM of INY-03-041 in both T47D and MDA-MB-468 cells (**Figure 4B**). This durable AKT degradation resulted in sustained inhibition of downstream signaling, as pPRAS40 levels were also significantly reduced for up to 96 hours (**Figure 4B**). In contrast, treatment with equivalent dose of GDC-0068 not only resulted in less pronounced downregulation of pPRAS40 levels, but the duration of this effect was also shorter (**Figure 4B**).

To further characterize the mechanism underlying the extended duration of AKT degradation induced by INY-03-041, we performed compound washout experiments after 12 hours of treatment with either 250 nM of INY-03-041 or GDC-0068. We observed no detectable rebound of AKT levels for up to 96 hours after washout in INY-03-041 treated cells (**Figure 4C**), suggesting that the re-synthesis rate of AKT is slow. Consistently, INY-03-041 potently suppressed levels of pPRAS40 for up to 96 hours after washout, while washout in GDC-0068 treated cells resulted in rebound of pPRAS40, as would be expected of a reversible inhibitor (**Figure 4C**). Taken together, our data suggest that INY-03-041-mediated AKT degradation resulted in more potent and durable pharmacological effects than AKT inhibition.

## DISCUSSION

Heterobifunctional degraders have garnered attention recently due to potential advantages over traditional small molecule inhibitors, including their ability to target proteins deemed undruggable through the use of non-functional ligands (Farnaby et al., 2019; Silva et al., 2019) and to achieve selectivity with promiscuous ligands (Nowak et al., 2018). However, one advantage of degraders that has not been fully investigated is their potential to exert long-lasting pharmacological effects. Given that degraders deplete abundance of a targeted protein, shortterm drug exposure may deliver sustained pharmacological effects, uncoupling pharmacodynamics from pharmacokinetics. This distinction may be more apparent for proteins with slow re-synthesis rates, as longer periods of time may be required for protein levels to recover and re-establish physiological signaling.

Consistent with the long reported half-life for AKT (Basso et al., 2002), INY-03-041 destabilized all three AKT isoforms and diminished downstream signaling effects for up to 96 hours, even after compound washout. This durable and prolonged pharmacodynamic effect, which was significantly greater than that of the parent AKT inhibitor GDC-0068, may explain, at least in part, the more potent anti-proliferative effects of INY-03-041. To the best of our knowledge, this is the first example of a small molecule degrader that sustains knockdown and downstream signaling effects for such an extended period of time, and exemplifies a distinguishing feature of degrader pharmacology.

Although several ATP-competitive and allosteric AKT inhibitors have entered clinical trials, dose-limiting toxicities or lack of clinical efficacy have hindered their progress. Moreover, AKT has been reported to have kinase-independent functions (Vivanco et al., 2014) that would reduce efficacy of small molecule inhibitors, and resistance to AKT inhibitors can arise in some contexts due to upregulation of AKT3 (Stottrup et al., 2016). Degraders have been reported to abrogate kinase-independent functions (Cromm et al., 2018) and overcome inhibitor-induced compensatory upregulation of target proteins (Cai et al., 2012; Han et al., 2019). Targeted AKT degradation may be a promising alternative approach for therapeutic intervention in cancers with PI3K/AKT pathway alterations. Although our data indicate that AKT degraders can induce more potent and durable downstream effects than AKT inhibitors, further optimization, particularly of the pharmacokinetic properties of the large heterobifunctional degrader molecules will be necessary to investigate the therapeutic viability of targeted AKT degradation.

Our data support the value of INY-03-041 as a chemical probe for studying the acute biological effects of pan-AKT depletion. Existing methods for inducing AKT1, AKT2 and AKT3 depletion in cells have relied on CRISPR knockout or repression, doxycycline-inducible shRNAs, or transient transfection with siRNA, all of which require relatively long incubation periods (Degtyarev et al., 2008; Koseoglu et al., 2007). This not only prevents assessment of acute biological changes resulting from loss of AKT, but also may allow reprogramming and compensation of cellular networks. By contrast, INY-03-041 induces rapid degradation of all three AKT isoforms, thereby permitting the investigation of the phenotypes of acute pan-AKT depletion. Therefore, INY-03-041 will be a useful chemical tool to explore AKT biology inaccessible with current strategies.

## SIGNIFICANCE

While many small molecule AKT inhibitors have been developed, here we demonstrate an alternative approach to targeting AKT by inducing its degradation, which resulted in distinct pharmacological effects. INY-03-041, a heterobifunctional degrader, promoted rapid and highly selective degradation of all three AKT isoforms and had more potent anti-proliferative effects than GDC-0068, the most clinically advanced AKT inhibitor. More importantly, INY-03-041-induced degradation of AKT and the ensuing suppression of downstream signaling were sustained for several days. Our work showcases the extended pharmacodynamic effects of degraders, and presents INY-03-041 as a tool to study acute changes to the AKT signaling network after rapid AKT degradation.

## ACKNOWLEDGEMENTS

We would like to thank Eric Wang and Milka Kostic for helpful comments on the manuscript. We would also like to thank Caitlin Mills and Kartik Subramanian (Laboratory for Systems Pharmacology at Harvard Medical School) for their help with proliferation data collection and analysis. This work was supported by the Training Grant in Chemical Biology (NIH 7 T32 GM095450-07, I.Y.), the Training Grant in Pharmacological Sciences (NIH 1 T32 GM132089-01, I.Y.), the National Science Foundation Graduate Research Fellowship Program (DGE1745303, E.C.E.), the NIH 1R01 CA218278-01A1 (E.S.F., N.S.G.) and the Ludwig Center at Harvard (E.C.E., A.T.).

## AUTHOR CONTRIBUTIONS

A.T. and N.S.G. conceived the study. I.Y. designed and synthesized the compounds. E.C.E. and I.Y. designed and conducted the biological profiling of the degraders. K.A.D. and N.A.E. performed the proteomics. I.Y and E.C.E wrote the manuscript, with guidance from E.S.F., N.S.G., and A.T. All authors gave feedback on the manuscript.

