## Supplemental Figures S1-S5 and Methods for "Discovery of an AKT Degrader with Prolonged Inhibition of Downstream Signaling"

### Supplemental Information

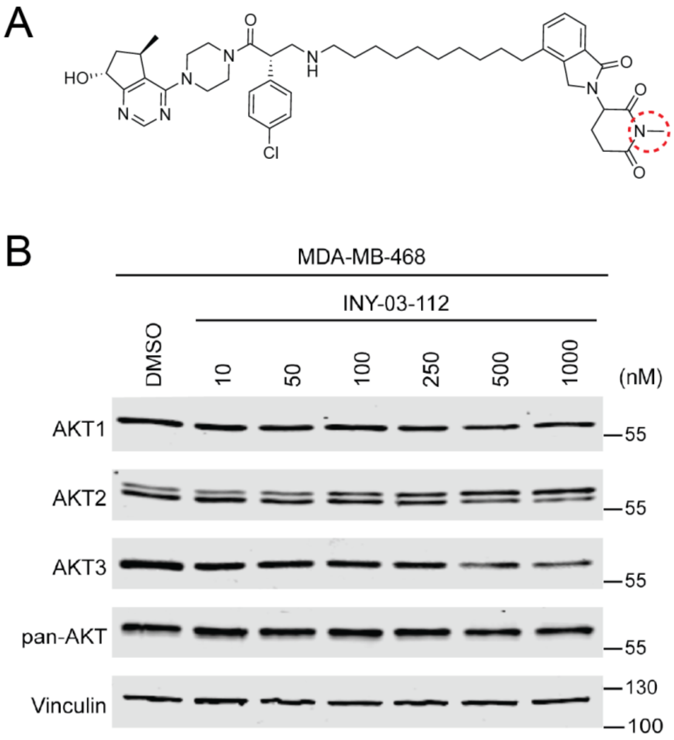


**Figure S1. Negative control INY-03-112 does not degrade AKT isoforms.** *Related to Table S1*. (A) Chemical structure of negative control compound INY-03-112, with N-methylated glutarimide circled. (B) Immunoblots for AKT1, AKT2, AKT3, pan-AKT, and Vinculin in MDA-MB-468 cells after 12 hour treatment with DMSO or INY-03-112 at the concentrations indicated (representative of n=2).

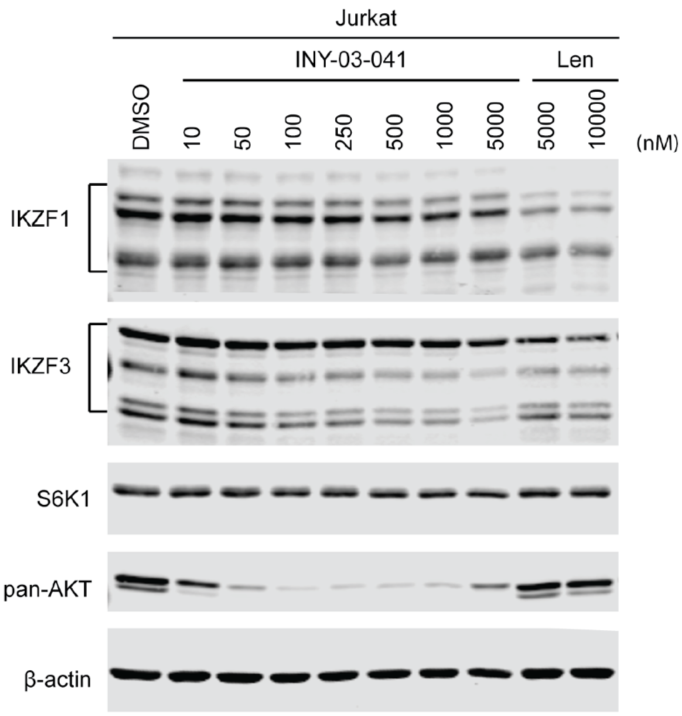


**Figure S2. INY-03-041-mediated degradation of IKZF1 and IKZF3.** (A) Immunoblots for IKZF1, IKZF3, S6K1, pan-AKT, and β-actin in Jurkat cells after 24 hour treatment with DMSO, INY-03-041, or lenalidomide (Len) at the concentrations indicated (n=3).

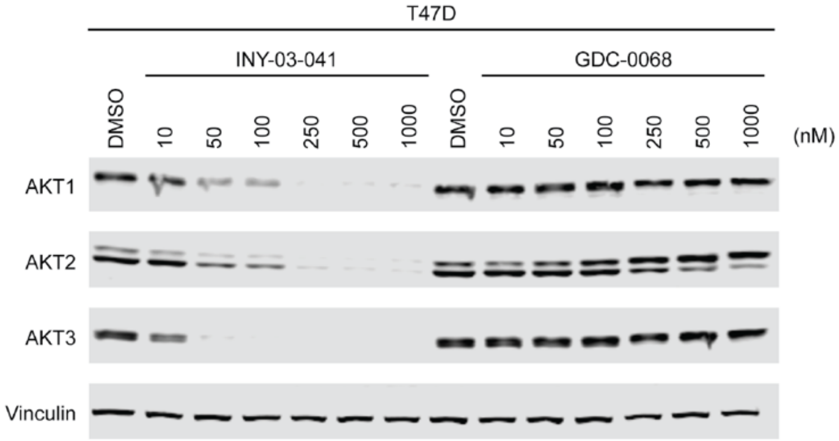


**Figure S3. INY-03-041 induces degradation of AKT isoforms in T47D cells.** *Related to Figure 4.* (A) Immunoblots of AKT1, AKT2, AKT3, and Vinculin after treating T47D cells for 24 hours with DMSO, INY-03-041, or GDC-0068 at the concentrations indicated (n=3).

**Table S1. Biochemical $IC_{50}$ (nM) values for INY-03-041 and INY-03-112.** *Related to Figure 1 and Figure S1.* Z'lyte kinase assays (Invitrogen) were conducted to assess $IC_{50}$ values for all kinases listed in the table with the exception of RET (V804M) which was assayed using LanthaScreen assays (Invitrogen).

| | **Compound** | |
|---|---|---|
| **Kinase** | **INY-03-041 $IC_{50}$ (nM)** | **INY-03-112 $IC_{50}$ (nM)** |
| AKT1 | 1.96 | 1.5 |
| AKT2 | 6.78 | 13.7 |
| AKT3 | 3.51 | 4.45 |
| PKG1 | 33.2 | N/A |
| S6K1 | 37.3 | N/A |
| PKN1 | 51.7 | N/A |
| ßMSK2 | 3.31 | N/A |
| Haspin | 4020 | N/A |
| RET (V804M) | >10000 | N/A |

**Table S2. KINOMEscan hits.** *Related to Figure 1*.

**Table S3. Proteomics output from limma processing of MOLT4 cells treated for 4 hours with INY-03-041 vs DMSO control.** *Related to Figure 2*.

**Table S4**. **Hormone receptor and mutational status of PIK3CA and PTEN in cancer cell line panel.** The tissue, cancer subtype, and status of each gene (Meric-Bernstam et al., 2012; Vlietstra et al., 1998) as wild-type (wt), mutated (mut) or deleted (del) is indicated. Triple negative breast cancer (TNBC).

| **Cell Line** | **Tissue** | **Cancer Subtype** | **PIK3CA** | **PTEN** |
|---|---|---|---|---|
| ZR-75-1 | Breast | Luminal A | wt | mut (L108R) |
| T47D | Breast | Luminal A | mut (H1047R) | wt |
| LNCaP | Prostate | Androgen-dependent | wt | del/mut |
| MCF-7 | Breast | Luminal A | mut (E545K) | wt |
| MDA-MB-468 | Breast | TNBC – Basal A | wt | del |
| HCC1937 | Breast | TNBC – Basal A | wt | del |

**Table S5**. **GR values indicate anti-proliferative advantage of INY-03-041.** *Related to Figure 3.* (A) GR values were calculated after 72 hours treatment with the compounds indicated over a range of concentrations. $GR_{50}$ values represent compound potency, $GR_{max}$ values measure the efficacy of the drug at high concentrations. $GR_{AOC}$ captures changes in potency and efficacy and is calculated by integrating GR curve over a range of concentrations.

$GR_{50}$ (μM)

| Cell Line | INY-03-041 | INY-03-112 | GDC-0068 | Lenalidomide |
|---|---|---|---|---|
| ZR-75-1 | 0.01599132 | 0.41332816 | 0.2289724 | inf |
| T47D | 0.17831981 | 1.33985401 | 1.52509853 | inf |
| LNCaP | 0.13009989 | 1.38277274 | 1.31522936 | inf |
| MCF-7 | 0.14798964 | 1.49338161 | 1.71566724 | inf |
| MDA-MB-468 | 1.68557896 | 2.69522214 | 12.9176626 | inf |
| HCC1937 | 1.64527129 | 2.41978757 | inf | inf |

$GR_{max}$

| Cell Line | INY-03-041 | INY-03-112 | GDC-0068 | Lenalidomide |
|---|---|---|---|---|
| ZR-75-1 | -0.6878133 | -0.5171039 | -0.3841938 | 0.74474387 |
| T47D | -0.3208748 | -0.1634936 | 0.05440203 | 0.99850631 |
| LNCaP | 0.01067184 | 0.21216665 | 0.01683696 | 0.82853147 |
| MCF-7 | -0.3054814 | -0.3317977 | 0.21842009 | 0.90960366 |
| MDA-MB-468 | -0.8194262 | -0.8318163 | 0.51778759 | 0.93954195 |
| HCC1937 | -0.8131698 | -0.7255681 | 0.53817904 | 0.90453 |

$GR_{AOC}$

| Cell Line | INY-03-041 | INY-03-112 | GDC-0068 | Lenalidomide |
|---|---|---|---|---|
| ZR-75-1 | 1.16640547 | 0.32890379 | 0.7053818 | 0.08476616 |
| T47D | 0.63026903 | 0.19803381 | 0.28472386 | -0.0100805 |
| LNCaP | 0.54925615 | 0.186322 | 0.32963774 | 0.04177353 |
| MCF-7 | 0.62747921 | 0.31979881 | 0.24329199 | 0.02059167 |
| MDA-MB-468 | 0.43042352 | 0.31894929 | 0.14038282 | 0.03254079 |
| HCC1937 | 0.41471365 | 0.31312878 | 0.15607112 | -0.0084259 |

**KEY RESOURCES TABLE**

**Antibodies**

| Antibodies | Source | CAT# |
|---|---|---|
| **AKT1** | Cell Signaling Technology | CST2938 |
| **AKT2** | Cell Signaling Technology | CST3063 |
| **AKT3** | Cell Signaling Technology | CST8018 |
| **Pan-AKT** | Cell Signaling Technology | CST4691<br>CST2920 |
| **Phospho-PRAS40 (T246)** | Cell Signaling Technology | CST2997 |
| **Total PRAS40** | Cell Signaling Technology | CST2691 |
| **Vinculin** | Cell Signaling Technology | CST13901 |
| **S6K1** | Cell Signaling Technology | CST2708 |
| **IKZF1 (Ikaros)** | Cell Signaling Technology | CST14859 |
| **IKZF3 (Aiolos)** | Cell Signaling Technology | CST15103 |
| **$\beta$-actin** | Cell Signaling Technology | CST4970 |

**Chemicals**

| Chemicals | Source | CAT# |
|---|---|---|
| **DMSO** | Fisher | BP231-100 |
| **Bortezomib (PS-341)** | Cayman Chemical | 10008822 |
| **Pevonedistat (MLN4924)** | Selleckchem | S7109 |
| **GDC-0068** | Selleckchem | S2808 |
| **Lenalidomide** | Sigma Aldrich | 901558 |
| **INY-03-041** | This study | N/A |
| **INY-03-112** | This study | N/A |

**Experimental Models: Cell Lines**

| Cell Line | Source | Identifier |
|---|---|---|
| **MOLT4** | ATCC | CRL-1582 |
| **Jurkat** | ATCC | TIB-152 |
| **ZR-75-1** | ATCC | CRL-1500 |
| **LNCaP** | Balk Lab | |
| **T47D** | ATCC | HTB-133 |
| **MCF7** | ATCC | HTB-22 |
| **MDA-MB-468** | ATCC | HTB-132 |
| **HCC1937** | ATCC | CRL-2336 |

**Critical Commercial Assays**

| Assay | Source | Identifier |
|---|---|---|
| **Z' Lyte (AKT1)** | Invitrogen | Assay ID 247 |
| **Z' Lyte (AKT2)** | Invitrogen | Assay ID 250 |
| **Z' Lyte (AKT3)** | Invitrogen | Assay ID 253 |
| **Z' Lyte (PKG1)** | Invitrogen | Assay ID 757 |
| **Z' Lyte (S6K1)** | Invitrogen | Assay ID 810 |
| **Z' Lyte (PKN1)** | Invitrogen | Assay ID 703 |
| **Z' Lyte (βMSK2)** | Invitrogen | Assay ID 801 |
| **Z' Lyte (Haspin)** | Invitrogen | Assay ID 1115 |
| **Lanthascreen (RET (V804M))** | Invitrogen | Assay ID 1847 |
| **Deposited Data** | | Pride: PXD015207 |

**Software and Algorithms**

| Software | Source | CAT# |
|---|---|---|
| **GraphPad Prism** | GraphPad Software, Inc. | www.graphpad.com/ |
| **Columbus image storage and analysis** | PerkinElmer Informatics | http://www.perkinelmer.com/product/image-data-storage-and-analysis-system-columbus |
| **Adobe Illustrator** | Adobe Creative Cloud | https://www.adobe.com/creativecloud.html |
| **Proteome Discoverer 2.2** | Thermo Fisher Scientific | https://www.thermofisher.com/order/catalog/product/OPTON-30795 |
| **R Framework** | Team RCR: A Language and Environment for Statistical Computing | http://www.R-project.org/ |

#### METHODS DETAILS

##### Cell Culture and Reagents

MOLT4, Jurkat, ZR-75-1, LNCaP, T47D, MCF-7, MDA-MB-468, and HCC1937 cells were cultured in RPMI media (Wisent Bioproducts, Cat #350000CL) supplemented with 10% heat inactivated fetal bovine serum (Thermo Fisher Scientific) at 37°C in the presence of 5% $CO_2$.

##### Drug Treatment Experiments

Cells were plated at 250,000 cells per mL (MDA-MB-468) or 200,000 cells per mL (T47D) in 2 mL per well RPMI media with 10% serum in 6-well treated tissue culture plates (Greiner, Cat # TCG-657160) or 60 mm treated tissue culture plates (Corning, Cat # 430166) and incubated overnight. The next day, cells were treated with the indicated compounds at the appropriate concentration and protein lysates were harvested at the times specified.

##### Immunoblotting

Cells were washed once in 1x PBS then lysed in RIPA buffer (150 mM Tris-HCl, 150 mM NaCl, 0.5% (w/v) sodium deoxycholate, 1% (v/v) NP-40, pH 7.5) containing 0.1% (w/v) sodium dodecyl sulfate, 1 mM sodium pyrophosphate, 20 mM sodium fluoride, 50 nM calyculin, and 0.5% (v/v) protease inhibitor cocktail (Sigma-Aldrich®) for 15 minutes. Cell extracts were precleared by centrifugation at 14,000 rpm for 10 minutes at 4°C. The Bio-Rad DC protein assay was used to assess protein concentration, and sample concentration was normalized using SDS sample buffer. Lysates were resolved on acrylamide gels by SDS-polyacrylamide gel electrophoresis and electrophoretically transferred to nitrocellulose membrane (BioRad) at 100 volts for 90 minutes. Membranes were blocked in 5% (w/v) nonfat dry milk in tris-buffered saline (TBS) buffer for 1 hour then incubated with specific primary antibodies diluted in 5% (w/v) nonfat dry milk in TBS-T (TBS with 0.05% Tween-20) at 4°C overnight, shaking. The next day, membranes were washed with TBS-T then incubated for 1 hour at room temperature with fluorophore-conjugated secondary antibodies (LI-COR Biosciences). The membrane was washed again with TBS-T then imaged with a LI-COR Odyssey CLx Imaging System (LI-COR Biosciences).

##### Growth Rate Assay and Analysis

Cell lines were plated at densities ranging from 500 to 2000 cells per well in a 384-well plate using a Matrix WellMate Reagent Dispenser (Thermo Fisher) and allowed 24 hours to adhere to plate prior to treatment. A D300 Digital Dispenser (Hewlett-Packard) was used to treat cells with dilution series of compounds as indicated. At the time of treatment and after 72 hours of treatment, cells were stained and fixed for subsequent analysis. Cells were stained with LIVE/DEAD Far Red Dead Cell Stain (LDR) (Thermo Fisher Scientific) at 1:2000 for one hour at 37°C. Cells were fixed for 30 minutes at room temperature in 4% formaldehyde (Sigma Aldrich) then permeabilized with 0.5% Triton X-100 in PBS. Cells were blocked for one hour using Odyssey Blocking Buffer (LI-COR Biosciences) and stained overnight at 4°C with 2 µg/ml Hoechst 33342 (Sigma Aldrich). An Operetta microscope was used to image fixed cells, and data was stored and analyzed using Columbus software (PerkinElmer). Nuclei were segmented by Hoechst signal using the Columbus system (Perkin Elmer). The average LDR and Hoechst intensities were determined within the nuclear area. Dead cells were classified by LDR signal. Experiments were performed in technical triplicate.

**In Vitro Kinase Assays**

Z'-LYTE assays were conducted for AKT1, AKT2, AKT3, PKG1, S6K1, PKN1, ßMSK2, and Haspin at Life Technologies in a 10-point dose response using Km ATP concentrations. LanthaScreen assays were conducted for RET (V804M) in a 10-point dose response at Life Technologies.

**TMT LC-MS Sample Preparation**

MOLT4 cells were treated with DMSO, 250 nM INY-03-041-01 for 4 hours in biological triplicates. Cells were harvested by centrifugation. Lysis buffer (8 M Urea, 50 mM NaCl, 50 mM 4-(2hydroxyethyl)-1-piperazineethanesulfonic acid (EPPS) pH 8.5, 1x Roche protease inhibitor and 1x Roche PhosphoStop was added to the cell pellets and cells were homogenized by 20 passes through a 21 gauge (1.25 in. long) needle to achieve a cell lysate with a protein concentration between 0.5 – 4 mg $mL^{-1}$. The homogenized sample was clarified by centrifugation at 20,000 x g for 10 minutes at 4 °C. A Bradford assay was used to determine the final protein concentration in the cell lysate. 200 µg protein for each sample were reduced and alkylated as previously described(An et al., 2017). Proteins were precipitated using methanol/chloroform. In brief, four volumes of methanol were added to the cell lysate, followed by one volume of chloroform, and finally three volumes of water. The mixture was vortexed and centrifuged at 14,000 x g for 5 minutes to separate the chloroform phase from the aqueous phase. The precipitated protein was washed with three volumes of methanol, centrifuged at 14,000 x g for 5 min, and the resulting washed precipitated protein was allowed to air dry. Precipitated protein was resuspended in 4 M Urea, 50 mM HEPES pH 7.4, followed by dilution to 1 M urea with the addition of 200 mM EPPS pH 8 for digestion with LysC (1:50; enzyme:protein) for 12 hours at RT. The LysC digestion was diluted to 0.5 M Urea, 200 mM EPPS pH 8 and then digested with trypsin (1:50; enzyme:protein) for 6 hours at 37°C. Tandem mass tag (TMT) reagents (Thermo Fisher Scientific) were dissolved in anhydrous acetonitrile (ACN) according to manufacturer's instructions. Anhydrous ACN was added to each peptide sample to a final concentration of 30% v/v, and labeling was induced with the addition of TMT reagent to each sample at a ratio of 1:4 peptide:TMT label. The 11-plex labeling reactions were performed for 1.5 hours at RT and the reaction quenched by the addition of 0.3% hydroxylamine for 15 minutes at RT. The sample channels were combined at a 1:1:1:1:1:1:1:1:1:1:1 ratio, desalted using $C_{18}$ solid phase extraction cartridges (Waters) and analyzed by LC-MS for channel ratio comparison. Samples were then combined using the adjusted volumes determined in the channel ratio analysis and dried down in a speed vacuum. The combined sample was then resuspended in 1% formic acid and acidified (pH 2−3) before being subjected to desalting with C18 SPE (Sep-Pak, Waters). Samples were then offline fractionated into 96 fractions by high pH reverse-phase HPLC (Agilent LC1260) through an aeris peptide xb-c18 column (phenomenex) with mobile phase A containing 5% acetonitrile and 10 mM $NH_4HCO_3$ in LC-MS grade $H_2O$, and mobile phase B containing 90% acetonitrile and 10 mM $NH_4HCO_3$ in LC-MS grade $H_2O$ (both pH 8.0). The 96 resulting fractions were then pooled in a non-continuous manner into 24 fractions or 48 fractions and every fraction was used for subsequent mass spectrometry analysis.

Data were collected using an Orbitrap Fusion Lumos mass spectrometer (Thermo Fisher Scientific, San Jose, CA, USA) coupled with a Proxeon EASY-nLC 1200 LC pump (Thermo Fisher Scientific). Peptides were separated on a 50 cm and 75 µm inner diameter Easyspray column (ES803a, Thermo Fisher Scientific). Peptides were separated using a 190 minute gradient of 6 – 27% acetonitrile in 1.0% formic acid with a flow rate of 300 nL/min.

Each analysis used an $MS^3$-based TMT method as described previously (McAlister et al., 2014). The data were acquired using a mass range of *m/z* 340 – 1350, resolution 120,000, AGC target 5 x $10^5$, maximum injection time 100 ms, dynamic exclusion of 120 seconds for the peptide measurements in the Orbitrap. Data dependent $MS^2$ spectra were acquired in the ion trap with a normalized collision energy (NCE) set at 35%, AGC target set to 1.8 x $10^4$ and a maximum injection time of 120 ms. $MS^3$ scans were acquired in the Orbitrap with a HCD collision energy set to 55%, AGC target set to 2 x $10^5$, maximum injection time of 150 ms, resolution at 50,000 and with a maximum synchronous precursor selection (SPS) precursors set to 10.

**LC-MS Data analysis**

Proteome Discoverer 2.2 (Thermo Fisher) was used for .RAW file processing and controlling peptide and protein level false discovery rates, assembling proteins from peptides, and protein quantification from peptides. MS/MS spectra were searched against a Uniprot human database (September 2016) with both the forward and reverse sequences. Database search criteria are as follows: tryptic with two missed cleavages, a precursor mass tolerance of 10 ppm, fragment ion mass tolerance of 0.6 Da, static alkylation of cysteine (57.02146 Da), static TMT labeling of lysine residues and N-termini of peptides (229.16293 Da), variable phosphorylation of serine, threonine and tyrosine (79.966 Da), and variable oxidation of methionine (15.99491 Da). TMT reporter ion intensities were measured using a 0.003 Da window around the theoretical *m/z* for each reporter ion in the $MS^3$ scan. Peptide spectral matches with poor quality $MS^3$ spectra were excluded from quantitation (summed signal-to-noise across 10 channels < 200 and precursor isolation specificity < 0.5). Only proteins containing at least two unique peptides identified in the experiment were included in final quantitation. Reporter ion intensities were normalized and scaled using in house scripts and the R framework(Team, 2013). Statistical analysis was carried out using a moderated t-test in the limma package within the R framework (Ritchie et al., 2015).

#### Chemistry Synthetic Scheme

**Scheme 1**

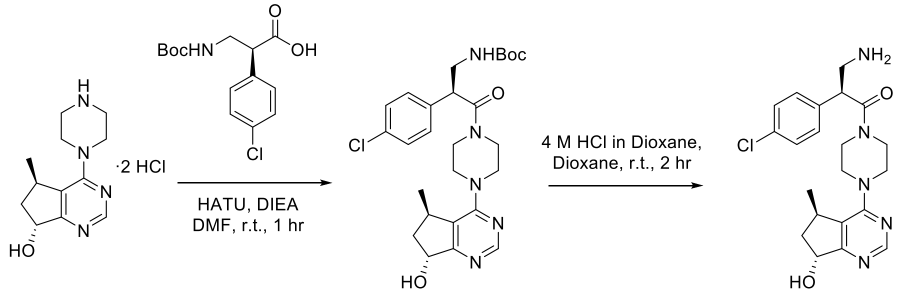


1 2 3

**Scheme 2**

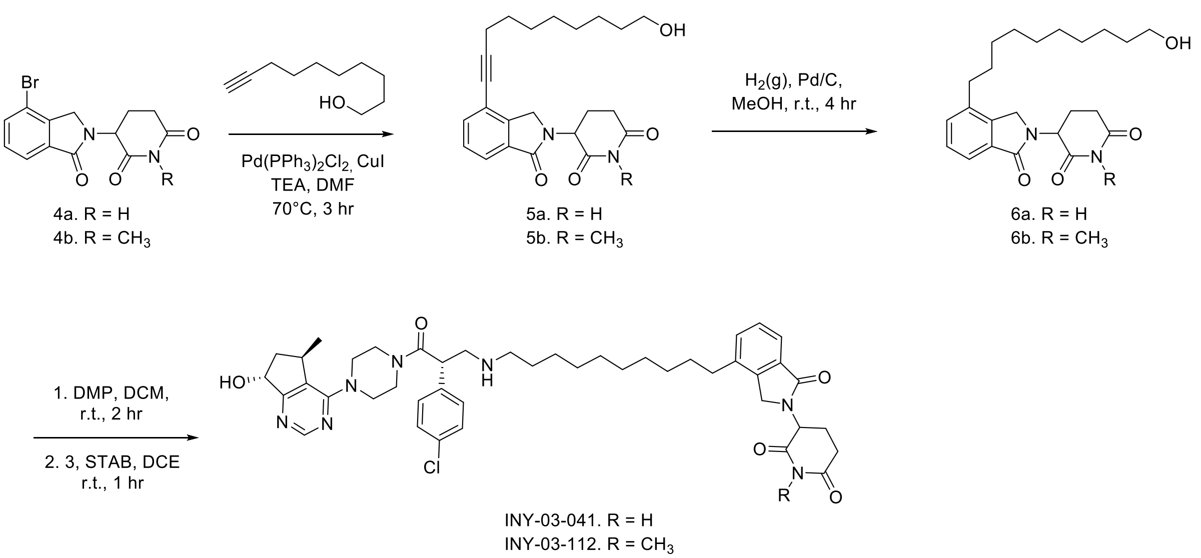


#### General Chemistry Methods

Reagents and solvents were purchased from commercial suppliers and were used without further purification unless otherwise noted. Reactions were monitored using a Waters Acquity UPLC/MS system (Waters PDA eλ Detector, QDa Detector, Sample manager - FL, Binary Solvent Manager) using Acquity UPLC® BEH C18 column (2.1 x 50 mm, 1.7 µm particle size): solvent gradient = 85% A at 0 min, 1% A at 1.7 min; solvent A = 0.1% formic acid in Water; solvent B = 0.1% formic acid in Acetonitrile; flow rate: 0.6 mL/min. Products were purified by flash column chromatography using CombiFlash®Rf with Teledyne Isco RediSep® normal-phase silica flash columns (4 g, 12 g, 24 g, 40 g or 80 g) and preparative HPLC using Waters SunFireTM Prep C18 column (19 x 100 mm, 5 µm particle size) using a gradient of 15-75% methanol in water containing 0.05% trifluoroacetic acid (TFA) over 48 min (60 min run time) at a flow of 40 mL/min. $^{1}$H NMR spectra were recorded on 500 MHz Bruker Avance III spectrometer, and chemical shifts are reported in million (ppm, δ) downfield from tetramethylsilane (TMS).

Coupling constants (J) are reported in Hz. Spin multiplicities are described as s (singlet), br (broad singlet), d (doublet), t (triplet), q (quartet) and m (multiplet). Purities of assayed compounds were in all cases greater than 95%, as determined by reverse-phase HPLC analysis.

**Abbreviations Used**
DCM: Dichloromethane
DIEA: N,N-Diisopropylethylamine
DMF: Dimethylformamide
TEA: Triethylamine
DMP: Dess–Martin periodinane
STAB: Sodium triacetoxyborohydride
DCE: 1,2-Dichloroethane
MeOH: Methanol

**Chemical Synthesis of INY-03-041 and INY-03-112**
**Synthesis of 3**
Tert-butyl((S)-2-(4-chlorophenyl)-3-(4-((5R,7R)-7-hydroxy-5-methyl-6,7-dihydro-5H-cyclopenta[d]pyrimidin-4-yl)piperazin-1-yl)-3-oxopropyl)carbamate (**2**):
To a solution of known intermediate **1** (Blake et al., 2012) (727 mg, 2.37 mmol), (S)-3-((tert-butoxycarbonyl)amino)-2-(4-chlorophenyl)propanoic acid (711 mg, 2.37 mmol), and HATU (902 mg, 2.37 mmol) in DMF (12 mL) was added DIEA (2.1 mL) and stirred for 1 hour at r.t. The reaction mixture was diluted ethyl acetate (30 mL) and washed with water (2 x 15 mL) and brine (2 x 15 mL). The organic layer was dried over anhydrous sodium sulfate, filtered, and concentrated in vacuo to afford **2** as a yellow oil (1.22 g, 100% yield). The crude was used directly for the next reaction. LC-MS: m/z 516.27 [M+1].

(S)-3-amino-2-(4-chlorophenyl)-1-(4-((5R,7R)-7-hydroxy-5-methyl-6,7-dihydro-5H-cyclopenta[d]pyrimidin-4-yl)piperazin-1-yl)propan-1-one (**3**):
To a solution of intermediate **2** (1.22 g, 2.37 mmol) in dioxane (7 mL) was added 4 M HCl in dioxane (5 mL). The reaction was stirred at r.t. for 4 hours. The reaction mixture was concentrated in vacuo and purified by reverse-phase HPLC (95-35% methanol in water) to obtain **3** as a TFA salt. The resultant TFA salt was dissolved in a mixture of 4:1 chloroform: isopropanol (50 mL) and washed with sat. aq sodium bicarbonate (50 mL) to give **3** as a white foam (868 mg, 88% yield). LC-MS: m/z 416.25 [M+1]. $^{1}$H NMR (500 MHz, DMSO-d6) δ 8.43 (s, 1H), 7.43 – 7.38 (m, 2H), 7.35 – 7.28 (m, 2H), 4.84 (t, J = 7.3, 6.2 Hz, 1H), 4.07 (dd, J = 8.3, 5.3 Hz, 1H), 3.73 – 3.66 (m, 1H), 3.64 – 3.56 (m, 3H), 3.52 – 3.35 (m, 4H), 3.22 – 3.16 (m, 1H), 3.09 (dd, J = 12.6, 8.3 Hz, 1H), 2.72 – 2.67 (m, 1H), 2.01 – 1.87 (m, 2H), 1.04 (dd, J = 6.5, 3.9 Hz, 3H).

**Synthesis of INY-03-041 and INY-03-112**
3-(4-(10-hydroxydec-1-yn-1-yl)-1-oxoisoindolin-2-yl)piperidine-2,6-dione (**5a**)
Known intermediate **4a** (Zhou et al., 2018) (500 mg, 1.55 mmol), dec-9-yn-1-ol (478 mg, 3.10 mmol), $Pd(PPh_3)_2Cl_2$ (113 mg, 0.16 mmol), and CuI (61 mg, 0.32 mmol) were dissolved in TEA

(4 mL) and DMF (8 mL). The reaction mixture was flushed with nitrogen (x3) and stirred at 70°C for 3 hours. The reaction mixture was cooled to r.t., diluted with ethyl acetate (50 mL), and filtered through celite. The organic layer was washed with brine (3 x 20 mL), dried over anhydrous sodium sulfate, filtered, and concentrated in vacuo. The crude material was purified by column chromatography on silica gel (9:1 DCM:MeOH) to afford **5a** as a pale yellow solid (503 mg, 82% yield). LC-MS: m/z 397.24 [M+1]. $^1$H NMR (500 MHz, DMSO-d6) δ 11.00 (s, 1H), 7.71 (dd, J = 7.6, 1.0 Hz, 1H), 7.64 (dd, J = 7.7, 1.0 Hz, 1H), 7.52 (t, J = 7.6 Hz, 1H), 5.76 (s, 2H), 5.15 (dd, J = 13.3, 5.2 Hz, 1H), 4.45 (d, J = 17.6 Hz, 1H), 4.31 (d, J = 17.6 Hz, 1H), 3.37 (t, J = 6.5 Hz, 2H), 2.92 (ddd, J = 17.3, 13.6, 5.5 Hz, 1H), 2.64 – 2.57 (m, 1H), 2.50 – 2.40 (m, 2H), 2.06 – 1.99 (m, 1H), 1.62 – 1.54 (m, 2H), 1.47 – 1.37 (m, 3H), 1.36 – 1.23 (m, 5H), 1.18 (t, J = 7.3 Hz, 1H).

3-(4-(10-hydroxydecyl)-1-oxoisoindolin-2-yl)piperidine-2,6-dione (**6a**)
To a solution of intermediate **5a** (100 mg, 0.25 mmol) in MeOH (10 mL) was added Pd/C (10 mg). $H_2$ (g) was introduced to the reaction mixture, and the reaction was stirred at r.t. for 4 hours. The reaction mixture was filtered over celite and concentrated in vacuo to afford **6a** as a white solid (91 mg, 91% yield). LC-MS: m/z 401.30 [M+1]. $^1$H NMR (500 MHz, DMSO-d6) δ 10.98 (s, 1H), 7.60 – 7.54 (m, 1H), 7.50 – 7.41 (m, 2H), 5.13 (dd, J = 13.3, 5.1 Hz, 1H), 4.45 (d, J = 17.2 Hz, 1H), 4.34 – 4.28 (m, 2H), 3.36 (td, J = 6.5, 4.9 Hz, 2H), 2.92 (ddd, J = 17.3, 13.7, 5.4 Hz, 1H), 2.66 – 2.58 (m, 3H), 2.43 (qd, J = 13.3, 4.5 Hz, 1H), 2.01 (dtd, J = 12.7, 5.2, 2.2 Hz, 1H), 1.66 – 1.54 (m, 2H), 1.43 – 1.35 (m, 2H), 1.33 – 1.21 (m, 12H).

3-(4-(10-(((S)-2-(4-chlorophenyl)-3-(4-((5R,7R)-7-hydroxy-5-methyl-6,7-dihydro-5H-cyclopenta[d]pyrimidin-4-yl)piperazin-1-yl)-3-oxopropyl)amino)decyl)-1-oxoisoindolin-2-yl)piperidine-2,6-dione (**INY-03-41**)
To a solution of intermediate **6a** (83 mg, 0.21 mmol) in DCM (4 mL) was added DMP (132 mg, 0.31 mmol). The reaction was stirred at r.t. for 2 hours. The reaction mixture was filtered and concentrated in vacuo to obtain 10-(2-(2,6-dioxopiperidin-3-yl)-1-oxoisoindolin-4-yl)decanal (84 mg, quantitative yield). LC-MS: m/z 399.27 [M+1]. The crude material was then dissolved in DCE (2 mL), followed by addition of intermediate **3** (131 mg, 0.31 mmol). The reaction was stirred at r.t. for 30 minutes, after which STAB (89 mg, 0.42 mmol) was added. The reaction mixture was stirred for an additional 1 hour. The reaction was quenched by sat. aq sodium bicarbonate (10 mL) and extracted with DCM (3 x 20 mL). The organic layers were combined, dried over anhydrous sodium sulfate, filtered, and concentrated in vacuo. The crude residue was purified by reverse-phase HPLC (75-15% methanol in water) to obtain **INY-03-041** (TFA salt). The resultant TFA salt was dissolved in 4:1 chloroform:isopropanol (10 mL) and washed with sat. aq sodium bicarbonate. The organic layer was dried over anhydrous sodium sulfate, filtered, and concentrated in vacuo to afford title compound as an off-white solid (42 mg, 25% yield). LC-MS: m/z 798.42 [M+1]. $^1$H NMR (500 MHz, DMSO-d6) δ 8.43 (s, 1H), 7.56 (dd, J = 5.4, 3.2 Hz, 1H), 7.45 – 7.43 (m, 2H), 7.41 (d, J = 8.2 Hz, 2H), 7.34 (d, J = 8.2 Hz, 2H), 5.38 (d, 1H), 5.13 (dd, J = 13.3, 5.1 Hz, 1H), 4.87 – 4.80 (m, 1H), 4.45 (d, J = 17.1 Hz, 1H), 4.29 (dd, J = 15.4, 7.6 Hz, 2H), 3.70 – 3.56 (m, 5H), 3.51 – 3.42 (m, 3H), 3.41 – 3.36 (m, 1H), 3.24 – 3.16 (m, 2H), 2.92 (ddd, J = 17.9, 13.8, 5.4 Hz, 1H), 2.77 – 2.71 (m, 1H), 2.63 (t, J = 7.5 Hz, 3H), 2.60 –

2.55 (m, 2H), 2.42 (tt, J = 13.3, 6.6 Hz, 1H), 2.04 – 1.93 (m, 2H), 1.93 – 1.87 (m, 1H), 1.62 – 1.55 (m, 2H), 1.41 – 1.35 (m, 2H), 1.32 – 1.26 (m, 4H), 1.22 (s, 8H), 1.03 (d, J = 6.8 Hz, 3H).

3-(4-(10-hydroxydec-1-yn-1-yl)-1-oxoisoindolin-2-yl)-1-methylpiperidine-2,6-dione (**5b**)
**5b** was synthesized with similar procedures as **5a** using intermediate **4b** (90 mg, 0.27 mmol) as the starting material. **5b** was obtained as a light orange solid (49 mg, 44% yield). LC-MS: m/z 411.51 [M+1]. $^1$H NMR (500 MHz, DMSO-d6) δ 7.71 (dd, J = 7.6, 1.0 Hz, 1H), 7.63 (dd, J = 6.8, 1.2 Hz, 1H), 7.52 (t, J = 7.6 Hz, 1H), 5.21 (dd, J = 13.5, 5.1 Hz, 1H), 4.44 (d, J = 17.6 Hz, 1H), 4.30 (d, J = 17.6 Hz, 1H), 3.36 (t, J = 6.5 Hz, 2H), 3.04 – 2.95 (m, 4H), 2.79 – 2.72 (m, 1H), 2.47 (t, J = 7.1 Hz, 2H), 2.45 – 2.39 (m, 1H), 2.03 (dtd, J = 12.7, 5.2, 2.2 Hz, 1H), 1.61 – 1.52 (m, 2H), 1.47 – 1.34 (m, 4H), 1.34 – 1.24 (m, 6H).

3-(4-(10-hydroxydecyl)-1-oxoisoindolin-2-yl)-1-methylpiperidine-2,6-dione (**6b**)
**6b** was synthesized with similar procedures as **6a** using intermediate **5b** (42 mg, 0.1 mmol) as the starting material. **6b** was obtained as an off-white solid (33 mg, 79% yield). LC-MS: m/z 415.57 [M+1]. $^1$H NMR (500 MHz, DMSO-d6) δ 7.60 – 7.53 (m, 1H), 7.49 – 7.43 (m, 2H), 5.20 (dd, J = 13.4, 5.1 Hz, 1H), 4.45 (d, J = 17.1 Hz, 1H), 4.32 – 4.27 (m, 2H), 3.36 (td, J = 6.6, 5.1 Hz, 2H), 3.01 (s, 3H), 3.00 – 2.95 (m, 1H), 2.79 – 2.74 (m, 1H), 2.66 – 2.61 (m, 2H), 2.48 – 2.39 (m, 1H), 2.08 – 1.98 (m, 1H), 1.63 – 1.56 (m, 2H), 1.43 – 1.34 (m, 2H), 1.32 – 1.22 (m, 12H).

3-(4-(10-(((S)-2-(4-chlorophenyl)-3-(4-((5R,7R)-7-hydroxy-5-methyl-6,7-dihydro-5H-cyclopenta[d]pyrimidin-4-yl)piperazin-1-yl)-3-oxopropyl)amino)decyl)-1-oxoisoindolin-2-yl)-1-methylpiperidine-2,6-dione (**INY-03-112**)
**INY-03-112** was synthesized with similar procedures as **INY-03-041** using intermediate **6b** (27 mg, 0.07 mmol) as the starting material. **INY-03-112** was obtained as an off-white solid (9 mg, 16% yield). LC-MS: m/z 812.47 [M+1]. $^1$H NMR (500 MHz, DMSO-d6) δ 8.43 (s, 1H), 7.57 (dd, J = 5.8, 2.8 Hz, 1H), 7.49 – 7.43 (m, 4H), 7.37 – 7.32 (m, 2H), 5.40 (d, J = 5.5 Hz, 1H), 5.21 (dd, J = 13.4, 5.1 Hz, 1H), 4.83 (q, J = 6.3 Hz, 1H), 4.45 (d, J = 17.1 Hz, 1H), 4.38 (dd, J = 8.9, 5.1 Hz, 1H), 4.29 (d, J = 17.1 Hz, 1H), 3.76 – 3.41 (m, 8H), 3.41 – 3.34 (m, 3H), 3.10 (t, J = 10.0 Hz, 1H), 3.01 (s, 3H), 2.99 – 2.95 (m, 1H), 2.83 – 2.73 (m, 3H), 2.63 (t, J = 7.7 Hz, 2H), 2.43 (qd, J = 13.2, 4.5 Hz, 1H), 2.05 – 1.87 (m, 3H), 1.59 (t, J = 7.5 Hz, 2H), 1.50 (s, 2H), 1.33 – 1.27 (m, 4H), 1.23 (s, 8H), 1.02 (d, J = 6.9 Hz, 3H).

**Table S2: KINOMEscan kinase selectivity prolfiles of INY-03-041 and GDC-0068. Related to Figur**

Both compounds were profiled at 1 µM against a diverse panel of 468 kinases (390 non-mutants kinases). Scores reported as percent of DMSO control (% control). The lower the score, the lower the Kd value is likely to be, such that scores of zero represent strong binding. Scores below 1 are highlighted in navy blue, between 1 and 10 in aqua blue, and between 10 and 35 in gray.

| | INY-03-041 | GDC-0068 |
|---|---|---|
| AAK1 | 95 | 100 |
| ABL1(E255K)-phosphorylated | 90 | 100 |
| ABL1(F317I)-nonphosphorylated | 70 | 100 |
| ABL1(F317I)-phosphorylated | 86 | 100 |
| ABL1(F317L)-nonphosphorylated | 87 | 100 |
| ABL1(F317L)-phosphorylated | 93 | 100 |
| ABL1(H396P)-nonphosphorylated | 74 | 95 |
| ABL1(H396P)-phosphorylated | 56 | 100 |
| ABL1(M351T)-phosphorylated | 100 | 100 |
| ABL1(Q252H)-nonphosphorylated | 61 | 86 |
| ABL1(Q252H)-phosphorylated | 65 | 100 |
| ABL1(T315I)-nonphosphorylated | 70 | 97 |
| ABL1(T315I)-phosphorylated | 87 | 98 |
| ABL1(Y253F)-phosphorylated | 40 | 100 |
| ABL1-nonphosphorylated | 68 | 74 |
| ABL1-phosphorylated | 60 | 88 |
| ABL2 | 86 | 100 |
| ACVR1 | 91 | 100 |
| ACVR1B | 82 | 75 |
| ACVR2A | 78 | 100 |
| ACVR2B | 98 | 100 |
| ACVRL1 | 96 | 93 |
| ADCK3 | 100 | 94 |
| ADCK4 | 84 | 100 |
| AKT1 | 11 | 0.7 |
| AKT2 | 1.2 | 6 |
| AKT3 | 26 | 20 |
| ALK | 75 | 75 |
| ALK(C1156Y) | 98 | 100 |
| ALK(L1196M) | 75 | 99 |
| AMPK-alpha1 | 98 | 100 |
| AMPK-alpha2 | 89 | 77 |
| ANKK1 | 65 | 100 |
| ARK5 | 88 | 98 |
| ASK1 | 100 | 100 |
| ASK2 | 74 | 100 |
| AURKA | 100 | 93 |

| | | |
|---|---|---|
| AURKB | 100 | 94 |
| AURKC | 100 | 94 |
| AXL | 87 | 87 |
| BIKE | 100 | 94 |
| BLK | 100 | 100 |
| BMPR1A | 77 | 78 |
| BMPR1B | 73 | 13 |
| BMPR2 | 83 | 100 |
| BMX | 100 | 94 |
| BRAF | 75 | 77 |
| BRAF(V600E) | 91 | 97 |
| BRK | 88 | 99 |
| BRSK1 | 98 | 100 |
| BRSK2 | 91 | 85 |
| BTK | 93 | 100 |
| BUB1 | 93 | 100 |
| CAMK1 | 100 | 100 |
| CAMK1B | 87 | 94 |
| CAMK1D | 96 | 100 |
| CAMK1G | 100 | 100 |
| CAMK2A | 96 | 100 |
| CAMK2B | 91 | 100 |
| CAMK2D | 96 | 100 |
| CAMK2G | 90 | 100 |
| CAMK4 | 100 | 87 |
| CAMKK1 | 96 | 100 |
| CAMKK2 | 87 | 100 |
| CASK | 85 | 99 |
| CDC2L1 | 83 | 100 |
| CDC2L2 | 100 | 100 |
| CDC2L5 | 41 | 93 |
| CDK11 | 84 | 78 |
| CDK2 | 86 | 100 |
| CDK3 | 70 | 98 |
| CDK4 | 83 | 100 |
| CDK4-cyclinD1 | 82 | 100 |
| CDK4-cyclinD3 | 78 | 100 |
| CDK5 | 100 | 100 |
| CDK7 | 89 | 100 |
| CDK8 | 93 | 100 |
| CDK9 | 82 | 100 |
| CDKL1 | 88 | 58 |
| CDKL2 | 100 | 97 |
| CDKL3 | 94 | 83 |
| CDKL5 | 82 | 100 |
| CHEK1 | 92 | 100 |
| CHEK2 | 93 | 100 |
| CIT | 100 | 80 |

| | | |
|---|---|---|
| CLK1 | 31 | 36 |
| CLK2 | 86 | 5.5 |
| CLK3 | 78 | 72 |
| CLK4 | 32 | 18 |
| CSF1R | 100 | 70 |
| CSF1R-autoinhibited | 86 | 98 |
| CSK | 100 | 100 |
| CSNK1A1 | 90 | 93 |
| CSNK1A1L | 90 | 100 |
| CSNK1D | 92 | 100 |
| CSNK1E | 82 | 94 |
| CSNK1G1 | 94 | 98 |
| CSNK1G2 | 88 | 100 |
| CSNK1G3 | 91 | 100 |
| CSNK2A1 | 100 | 100 |
| CSNK2A2 | 99 | 99 |
| CTK | 100 | 100 |
| DAPK1 | 85 | 99 |
| DAPK2 | 89 | 100 |
| DAPK3 | 71 | 100 |
| DCAMKL1 | 77 | 71 |
| DCAMKL2 | 100 | 94 |
| DCAMKL3 | 100 | 99 |
| DDR1 | 96 | 100 |
| DDR2 | 63 | 92 |
| DLK | 97 | 100 |
| DMPK | 99 | 85 |
| DMPK2 | 98 | 100 |
| DRAK1 | 98 | 100 |
| DRAK2 | 100 | 100 |
| DYRK1A | 70 | 77 |
| DYRK1B | 73 | 91 |
| DYRK2 | 76 | 100 |
| EGFR | 79 | 100 |
| EGFR(E746-A750del) | 76 | 100 |
| EGFR(G719C) | 100 | 76 |
| EGFR(G719S) | 100 | 78 |
| EGFR(L747-E749del, A750P) | 87 | 100 |
| EGFR(L747-S752del, P753S) | 97 | 92 |
| EGFR(L747-T751del,Sins) | 100 | 75 |
| EGFR(L858R) | 88 | 99 |
| EGFR(L858R,T790M) | 98 | 100 |
| EGFR(L861Q) | 100 | 73 |
| EGFR(S752-I759del) | 90 | 82 |
| EGFR(T790M) | 89 | 97 |
| EIF2AK1 | 77 | 100 |
| EPHA1 | 93 | 99 |
| EPHA2 | 100 | 100 |

| | | |
|---|---|---|
| EPHA3 | 76 | 100 |
| EPHA4 | 100 | 96 |
| EPHA5 | 100 | 100 |
| EPHA6 | 100 | 99 |
| EPHA7 | 96 | 100 |
| EPHA8 | 87 | 100 |
| EPHB1 | 90 | 100 |
| EPHB2 | 100 | 100 |
| EPHB3 | 100 | 100 |
| EPHB4 | 100 | 100 |
| EPHB6 | 63 | 100 |
| ERBB2 | 84 | 100 |
| ERBB3 | 100 | 100 |
| ERBB4 | 100 | 100 |
| ERK1 | 86 | 100 |
| ERK2 | 100 | 100 |
| ERK3 | 100 | 100 |
| ERK4 | 100 | 100 |
| ERK5 | 89 | 100 |
| ERK8 | 96 | 100 |
| ERN1 | 84 | 98 |
| FAK | 98 | 100 |
| FER | 100 | 82 |
| FES | 100 | 100 |
| FGFR1 | 98 | 94 |
| FGFR2 | 100 | 95 |
| FGFR3 | 100 | 97 |
| FGFR3(G697C) | 86 | 100 |
| FGFR4 | 96 | 96 |
| FGR | 100 | 98 |
| FLT1 | 100 | 100 |
| FLT3 | 91 | 95 |
| FLT3(D835H) | 100 | 95 |
| FLT3(D835V) | 91 | 100 |
| FLT3(D835Y) | 82 | 91 |
| FLT3(ITD) | 97 | 100 |
| FLT3(ITD,D835V) | 88 | 100 |
| FLT3(ITD,F691L) | 82 | 100 |
| FLT3(K663Q) | 98 | 91 |
| FLT3(N841I) | 83 | 100 |
| FLT3(R834Q) | 75 | 100 |
| FLT3-autoinhibited | 87 | 100 |
| FLT4 | 100 | 100 |
| FRK | 89 | 91 |
| FYN | 82 | 87 |
| GAK | 100 | 100 |
| GCN2(Kin.Dom.2,S808G) | 91 | 100 |
| GRK1 | 64 | 47 |

| | | |
|---|---|---|
| GRK2 | 88 | 96 |
| GRK3 | 39 | 97 |
| GRK4 | 88 | 87 |
| GRK7 | 81 | 97 |
| GSK3A | 87 | 95 |
| GSK3B | 67 | 100 |
| HASPIN | 19 | 35 |
| HCK | 96 | 100 |
| HIPK1 | 29 | 41 |
| HIPK2 | 67 | 49 |
| HIPK3 | 52 | 62 |
| HIPK4 | 76 | 86 |
| HPK1 | 100 | 100 |
| HUNK | 100 | 100 |
| ICK | 64 | 100 |
| IGF1R | 69 | 100 |
| IKK-alpha | 94 | 100 |
| IKK-beta | 84 | 99 |
| IKK-epsilon | 84 | 80 |
| INSR | 82 | 72 |
| INSRR | 88 | 100 |
| IRAK1 | 80 | 71 |
| IRAK3 | 97 | 98 |
| IRAK4 | 69 | 73 |
| ITK | 92 | 100 |
| JAK1(JH1domain-catalytic) | 80 | 98 |
| JAK1(JH2domain-pseudokinase) | 96 | 84 |
| JAK2(JH1domain-catalytic) | 73 | 99 |
| JAK3(JH1domain-catalytic) | 78 | 100 |
| JNK1 | 77 | 81 |
| JNK2 | 54 | 99 |
| JNK3 | 63 | 98 |
| KIT | 100 | 82 |
| KIT(A829P) | 80 | 100 |
| KIT(D816H) | 84 | 100 |
| KIT(D816V) | 100 | 100 |
| KIT(L576P) | 100 | 57 |
| KIT(V559D) | 100 | 91 |
| KIT(V559D,T670I) | 100 | 91 |
| KIT(V559D,V654A) | 81 | 100 |
| KIT-autoinhibited | 100 | 100 |
| LATS1 | 91 | 95 |
| LATS2 | 88 | 100 |
| LCK | 94 | 99 |
| LIMK1 | 93 | 100 |
| LIMK2 | 99 | 100 |
| LKB1 | 86 | 100 |
| LOK | 98 | 96 |

| | | |
|---|---|---|
| LRRK2 | 67 | 94 |
| LRRK2(G2019S) | 93 | 89 |
| LTK | 100 | 94 |
| LYN | 100 | 100 |
| LZK | 65 | 100 |
| MAK | 100 | 88 |
| MAP3K1 | 69 | 95 |
| MAP3K15 | 96 | 100 |
| MAP3K2 | 69 | 100 |
| MAP3K3 | 95 | 100 |
| MAP3K4 | 90 | 99 |
| MAP4K2 | 56 | 72 |
| MAP4K3 | 99 | 88 |
| MAP4K4 | 92 | 100 |
| MAP4K5 | 83 | 100 |
| MAPKAPK2 | 98 | 76 |
| MAPKAPK5 | 81 | 70 |
| MARK1 | 97 | 87 |
| MARK2 | 74 | 100 |
| MARK3 | 59 | 87 |
| MARK4 | 100 | 94 |
| MAST1 | 100 | 98 |
| MEK1 | 97 | 76 |
| MEK2 | 78 | 64 |
| MEK3 | 95 | 97 |
| MEK4 | 78 | 100 |
| MEK5 | 47 | 86 |
| MEK6 | 100 | 99 |
| MELK | 100 | 100 |
| MERTK | 100 | 100 |
| MET | 88 | 93 |
| MET(M1250T) | 56 | 98 |
| MET(Y1235D) | 92 | 100 |
| MINK | 80 | 91 |
| MKK7 | 96 | 100 |
| MKNK1 | 90 | 100 |
| MKNK2 | 57 | 100 |
| MLCK | 79 | 100 |
| MLK1 | 85 | 69 |
| MLK2 | 99 | 93 |
| MLK3 | 100 | 94 |
| MRCKA | 98 | 100 |
| MRCKB | 100 | 100 |
| MST1 | 92 | 92 |
| MST1R | 100 | 91 |
| MST2 | 100 | 84 |
| MST3 | 85 | 100 |
| MST4 | 85 | 100 |

| | | |
|---|---|---|
| MTOR | 100 | 100 |
| MUSK | 72 | 100 |
| MYLK | 84 | 95 |
| MYLK2 | 80 | 90 |
| MYLK4 | 100 | 88 |
| MYO3A | 98 | 100 |
| MYO3B | 100 | 88 |
| NDR1 | 73 | 72 |
| NDR2 | 98 | 100 |
| NEK1 | 100 | 95 |
| NEK10 | 84 | 82 |
| NEK11 | 98 | 71 |
| NEK2 | 85 | 100 |
| NEK3 | 58 | 100 |
| NEK4 | 89 | 100 |
| NEK5 | 100 | 82 |
| NEK6 | 100 | 100 |
| NEK7 | 96 | 100 |
| NEK9 | 95 | 100 |
| NIK | 100 | 99 |
| NIM1 | 86 | 100 |
| NLK | 93 | 100 |
| OSR1 | 74 | 100 |
| p38-alpha | 94 | 100 |
| p38-beta | 100 | 73 |
| p38-delta | 100 | 97 |
| p38-gamma | 100 | 100 |
| PAK1 | 98 | 95 |
| PAK2 | 90 | 86 |
| PAK3 | 99 | 91 |
| PAK4 | 96 | 100 |
| PAK6 | 80 | 97 |
| PAK7 | 65 | 95 |
| PCTK1 | 88 | 100 |
| PCTK2 | 100 | 77 |
| PCTK3 | 100 | 100 |
| PDGFRA | 100 | 100 |
| PDGFRB | 100 | 94 |
| PDPK1 | 100 | 75 |
| PFCDPK1(P.falciparum) | 99 | 100 |
| PFPK5(P.falciparum) | 85 | 100 |
| PFTAIRE2 | 96 | 100 |
| PFTK1 | 100 | 98 |
| PHKG1 | 98 | 100 |
| PHKG2 | 77 | 100 |
| PIK3C2B | 99 | 98 |
| PIK3C2G | 93 | 99 |
| PIK3CA | 100 | 86 |

| | | |
|---|---|---|
| PIK3CA(C420R) | 91 | 91 |
| PIK3CA(E542K) | 77 | 77 |
| PIK3CA(E545A) | 100 | 68 |
| PIK3CA(E545K) | 91 | 54 |
| PIK3CA(H1047L) | 81 | 96 |
| PIK3CA(H1047Y) | 96 | 100 |
| PIK3CA(I800L) | 89 | 62 |
| PIK3CA(M1043I) | 100 | 95 |
| PIK3CA(Q546K) | 84 | 100 |
| PIK3CB | 76 | 100 |
| PIK3CD | 100 | 80 |
| PIK3CG | 97 | 89 |
| PIK4CB | 53 | 100 |
| PIKFYVE | 81 | 100 |
| PIM1 | 98 | 98 |
| PIM2 | 100 | 76 |
| PIM3 | 100 | 93 |
| PIP5K1A | 87 | 88 |
| PIP5K1C | 100 | 100 |
| PIP5K2B | 81 | 100 |
| PIP5K2C | 100 | 100 |
| PKAC-alpha | 98 | 43 |
| PKAC-beta | 66 | 56 |
| PKMYT1 | 94 | 100 |
| PKN1 | 4.9 | 6.8 |
| PKN2 | 84 | 70 |
| PKNB(M.tuberculosis) | 100 | 96 |
| PLK1 | 52 | 100 |
| PLK2 | 81 | 92 |
| PLK3 | 82 | 67 |
| PLK4 | 91 | 97 |
| PRKCD | 94 | 68 |
| PRKCE | 97 | 100 |
| PRKCH | 61 | 33 |
| PRKCI | 37 | 81 |
| PRKCQ | 62 | 79 |
| PRKD1 | 89 | 93 |
| PRKD2 | 84 | 89 |
| PRKD3 | 95 | 100 |
| PRKG1 | 2.6 | 4.1 |
| PRKG2 | 33 | 12 |
| PRKR | 100 | 80 |
| PRKX | 100 | 100 |
| PRP4 | 86 | 87 |
| PYK2 | 90 | 100 |
| QSK | 91 | 100 |
| RAF1 | 96 | 93 |
| RET | 90 | 92 |

| | | |
|---|---|---|
| RET(M918T) | 98 | 98 |
| RET(V804L) | 100 | 100 |
| RET(V804M) | 0 | 92 |
| RIOK1 | 99 | 100 |
| RIOK2 | 82 | 100 |
| RIOK3 | 100 | 97 |
| RIPK1 | 100 | 100 |
| RIPK2 | 93 | 100 |
| RIPK4 | 51 | 100 |
| RIPK5 | 69 | 96 |
| ROCK1 | 72 | 22 |
| ROCK2 | 92 | 34 |
| ROS1 | 100 | 92 |
| RPS6KA4(Kin.Dom.1-N-terminal) | 27 | 74 |
| RPS6KA4(Kin.Dom.2-C-terminal) | 69 | 65 |
| RPS6KA5(Kin.Dom.1-N-terminal) | 75 | 65 |
| RPS6KA5(Kin.Dom.2-C-terminal) | 100 | 100 |
| RSK1(Kin.Dom.1-N-terminal) | 100 | 77 |
| RSK1(Kin.Dom.2-C-terminal) | 72 | 100 |
| RSK2(Kin.Dom.1-N-terminal) | 78 | 97 |
| RSK2(Kin.Dom.2-C-terminal) | 39 | 87 |
| RSK3(Kin.Dom.1-N-terminal) | 91 | 100 |
| RSK3(Kin.Dom.2-C-terminal) | 88 | 100 |
| RSK4(Kin.Dom.1-N-terminal) | 72 | 100 |
| RSK4(Kin.Dom.2-C-terminal) | 77 | 100 |
| S6K1 | 46 | 59 |
| SBK1 | 77 | 100 |
| SGK | 81 | 100 |
| SgK110 | 99 | 73 |
| SGK2 | 100 | 100 |
| SGK3 | 52 | 96 |
| SIK | 85 | 94 |
| SIK2 | 88 | 100 |
| SLK | 95 | 96 |
| SNARK | 80 | 100 |
| SNRK | 58 | 98 |
| SRC | 100 | 82 |
| SRMS | 74 | 84 |
| SRPK1 | 86 | 99 |
| SRPK2 | 100 | 100 |
| SRPK3 | 83 | 95 |
| STK16 | 100 | 95 |
| STK33 | 81 | 92 |
| STK35 | 99 | 100 |
| STK36 | 100 | 97 |
| STK39 | 55 | 95 |
| SYK | 89 | 94 |
| TAK1 | 100 | 100 |

| | | |
|---|---|---|
| TAOK1 | 79 | 85 |
| TAOK2 | 84 | 72 |
| TAOK3 | 99 | 71 |
| TBK1 | 92 | 65 |
| TEC | 95 | 100 |
| TESK1 | 94 | 90 |
| TGFBR1 | 77 | 91 |
| TGFBR2 | 99 | 87 |
| TIE1 | 82 | 100 |
| TIE2 | 95 | 91 |
| TLK1 | 79 | 100 |
| TLK2 | 100 | 100 |
| TNIK | 61 | 100 |
| TNK1 | 90 | 100 |
| TNK2 | 95 | 100 |
| TNNI3K | 100 | 93 |
| TRKA | 67 | 87 |
| TRKB | 66 | 100 |
| TRKC | 75 | 100 |
| TRPM6 | 96 | 100 |
| TSSK1B | 84 | 92 |
| TSSK3 | 100 | 94 |
| TTK | 100 | 100 |
| TXK | 89 | 98 |
| TYK2(JH1domain-catalytic) | 67 | 100 |
| TYK2(JH2domain-pseudokinase) | 91 | 74 |
| TYRO3 | 90 | 91 |
| ULK1 | 73 | 100 |
| ULK2 | 92 | 100 |
| ULK3 | 66 | 80 |
| VEGFR2 | 94 | 100 |
| VPS34 | 79 | 77 |
| VRK2 | 93 | 100 |
| WEE1 | 84 | 100 |
| WEE2 | 92 | 100 |
| WNK1 | 96 | 100 |
| WNK2 | 89 | 100 |
| WNK3 | 99 | 100 |
| WNK4 | 81 | 100 |
| YANK1 | 82 | 100 |
| YANK2 | 88 | 76 |
| YANK3 | 81 | 87 |
| YES | 95 | 100 |
| YSK1 | 100 | 96 |
| YSK4 | 92 | 93 |
| ZAK | 95 | 97 |
| ZAP70 | 79 | 100 |

e 1

Table S3

| | ID | logFC | AveExpr | t | P.Value | adj.P.Val | B |
|---|---|---|---|---|---|---|---|
| **P31751** | AKT2 | -1.739007527 | 3.057717968 | -53.86658945 | 7.89E-10 | 6.27E-06 | 9.871288575 |
| **P31749** | AKT1 | -1.224858439 | 3.221868404 | -36.95370339 | 8.98E-09 | 3.57E-05 | 8.316332427 |
| **P63261** | ACTG1 | -0.491837683 | 3.281277158 | -22.49288943 | 2.18E-07 | 0.000288758 | 7.6556987 |
| **Q9UL63-1** | MKLN1 | -0.30779296 | 3.268969323 | -20.07382366 | 4.51E-07 | 0.000512121 | 7.359684989 |
| **P11926** | ODC1 | -0.995963375 | 3.236907714 | -23.34691379 | 1.72E-07 | 0.000288758 | 6.606081366 |
| **Q9UKT9-1** | IKZF3 | -0.336087445 | 3.415095519 | -15.89604323 | 1.99E-06 | 0.00175563 | 5.81858533 |
| **Q9UKK3** | PARP4 | -0.234616308 | 3.243909607 | -14.02124859 | 4.39E-06 | 0.00348796 | 5.101121477 |
| **Q96A37-1** | RNF166 | -1.275614815 | 3.365716489 | -23.42000549 | 1.68E-07 | 0.000288758 | 4.935810597 |
| **Q8WTP8-2** | AEN | -0.84518177 | 3.228747127 | -16.6031387 | 1.51E-06 | 0.001499184 | 4.694294716 |
| **Q16667** | CDKN3 | -0.418484228 | 3.244680693 | -13.33002301 | 6.03E-06 | 0.004355745 | 4.550406547 |
| **Q9Y243-1** | AKT3 | -1.92474433 | 3.004713126 | -23.09367149 | 1.84E-07 | 0.000288758 | 4.439609398 |
| **P01106** | MYC | -0.429429793 | 3.269956478 | -11.38547393 | 1.61E-05 | 0.010675004 | 3.773199027 |
| **Q13772-4** | NCOA4 | -0.257271495 | 3.22860648 | -10.96887275 | 2.03E-05 | 0.011524641 | 3.57975291 |
| **Q86VP1-2** | TAX1BP1 | -0.197682393 | 3.226524517 | -11.10055293 | 1.89E-05 | 0.011524641 | 3.473678073 |
| **Q14534** | SQLE | 0.179517544 | 3.029794204 | 9.837533545 | 3.97E-05 | 0.021003283 | 2.806107929 |
| **Q9H871-1** | RMND5A | -0.195025676 | 3.232891317 | -8.764537349 | 7.99E-05 | 0.035288477 | 2.058315788 |
| **P05455** | SSB | 0.085183538 | 3.169349897 | 9.611936234 | 4.57E-05 | 0.022686612 | 1.967369317 |
| **Q92817** | EVPL | -0.235122241 | 3.163442651 | -8.304869974 | 0.000110481 | 0.039898643 | 1.925947664 |
| **P17707-1** | AMD 1.00 | -0.316189787 | 3.287985689 | -8.316947207 | 0.000109525 | 0.039898643 | 1.902216492 |
| **Q96E35** | ZMYND19 | -0.637804459 | 3.252820549 | -9.176114743 | 6.06E-05 | 0.028312178 | 1.767284735 |
| **Q9BXS4** | TMEM59 | 0.369134311 | 3.032175777 | 8.311319997 | 0.000109969 | 0.039898643 | 1.714974976 |
| **Q9Y2T2** | AP3M1 | -0.107021476 | 3.221073387 | -8.353624329 | 0.000106678 | 0.039898643 | 1.631770387 |
| **P47974** | ZFP36L2 | -0.299738585 | 3.197644853 | -7.888475596 | 0.000150094 | 0.051847637 | 1.625542807 |
| **Q5VV67-1** | PPRC1 | -0.495081167 | 3.324651728 | -7.745047953 | 0.000167327 | 0.055392221 | 1.230107304 |
| **Q9H8N7** | ZNF395 | 0.36134627 | 2.987345507 | 7.471638694 | 0.000206818 | 0.060858116 | 1.198104238 |
| **Q14CB8-2** | ARHGAP19 | -0.153591224 | 3.157092854 | -6.816568651 | 0.000352966 | 0.090461836 | 0.675156484 |
| **O60381-1** | HBP1 | 0.726589387 | 2.900367785 | 7.671835437 | 0.000176987 | 0.056246475 | 0.474821696 |
| **P52789** | HK2 | -0.106158875 | 3.213539073 | -7.368036682 | 0.000224482 | 0.063696677 | 0.468281263 |
| **P62753** | RPS6 | -0.21291348 | 3.163233373 | -7.154483877 | 0.000266593 | 0.07303725 | 0.453704645 |
| **Q14582** | MXD4 | 0.593304079 | 2.99751674 | 6.750457613 | 0.000373367 | 0.092700049 | 0.442823975 |
| **Q2TB10** | ZNF800 | -0.158609055 | 3.232979088 | -6.552581107 | 0.000442902 | 0.100538825 | 0.394494337 |
| **P46976-1** | GYG1 | 0.378850052 | 3.152566259 | 6.552693679 | 0.000442859 | 0.100538825 | 0.375847678 |
| **Q99608** | NDN | -0.228674581 | 3.309395032 | -6.331758308 | 0.000538453 | 0.118833639 | 0.329489332 |
| **P23396-1** | RPS3 | -0.063992732 | 3.202419092 | -7.569842778 | 0.000191523 | 0.058525035 | 0.200595007 |
| **Q8IZT6-1** | ASPM | -0.114191034 | 3.151055164 | -6.643689187 | 0.000409208 | 0.098519882 | 0.19452402 |
| **P17612** | PRKACA | 0.155215046 | 3.152714848 | 6.176216026 | 0.000619819 | 0.120108864 | 0.158143353 |
| **P62269** | RPS18 | -0.148062208 | 3.193088407 | -6.917719819 | 0.000324156 | 0.085847388 | 0.113838171 |

| | | | | | | | |
|---|---|---|---|---|---|---|---|
| **Q9BPZ3** | PAIP2 | 0.351575192 | 3.188588143 | 5.979059673 | 0.000743714 | 0.130801336 | 0.050207293 |
| **O94868** | FCHSD2 | -0.100199677 | 3.218779143 | -6.280161602 | 0.000564021 | 0.119580152 | -0.045085734 |
| **Q96G74-1** | OTUD5 | -0.251225656 | 3.291778667 | -6.059421391 | 0.00069011 | 0.127509823 | -0.080001931 |
| **Q96RQ3** | MCCC1 | 0.348191696 | 3.116828235 | 5.792825581 | 0.000887033 | 0.137578741 | -0.081735371 |
| **Q9UEE5** | STK17A | -0.222787163 | 3.28487835 | -5.777154221 | 0.000900452 | 0.137578741 | -0.13289687 |
| **Q15004-1** | KIAA0101; PCLAF | -0.184687353 | 3.191964757 | -6.139133873 | 0.000641218 | 0.12129708 | -0.24800493 |
| **Q58FF6** | HSP90AB4P | -0.256370918 | 3.205484077 | -5.643153459 | 0.001025042 | 0.145427831 | -0.252408808 |
| **Q6PI26** | SHQ1 | -0.173281345 | 3.249326584 | -5.869087192 | 0.000824873 | 0.13653365 | -0.287501605 |
| **Q9BWT1** | CDCA7 | -0.121351121 | 3.239615172 | -5.80045109 | 0.000880585 | 0.137578741 | -0.342931171 |
| **Q9NZM5** | GLTSCR2; NOP53 | -0.322655177 | 3.260737747 | -6.01193703 | 0.000721234 | 0.1302319 | -0.349689769 |
| **Q92187-1** | ST8SIA4 | 0.227384365 | 3.126474574 | 5.694855694 | 0.000974799 | 0.145427831 | -0.357274177 |
| **Q5VWQ8-1** | DAB2IP | -0.069336993 | 3.212691306 | -6.264720827 | 0.000571938 | 0.119580152 | -0.401017362 |
| **Q15014** | MORF4L2 | -0.198334262 | 3.220317749 | -5.600577706 | 0.001068616 | 0.146662208 | -0.406326423 |
| **Q92871** | PMM1 | -0.21941506 | 3.228860107 | -5.461622959 | 0.001226048 | 0.158829039 | -0.419228988 |
| **Q9NS23** | RASSF1 | -0.235532149 | 3.263574671 | -5.426280831 | 0.001270152 | 0.160180326 | -0.466057068 |
| **Q6PIF6-1** | MYO7B | -0.118470763 | 3.150483434 | -5.886129198 | 0.000811665 | 0.13653365 | -0.558694035 |
| **Q9H3R5** | CENPH | -0.090910405 | 3.190813962 | -5.959710942 | 0.000757314 | 0.130801336 | -0.561368493 |
| **Q7L590-1** | MCM10 | -0.170700135 | 3.200957276 | -5.450737162 | 0.001239446 | 0.158829039 | -0.568959934 |
| **Q01581** | HMGCS1 | -0.111286415 | 3.058054724 | -6.215786667 | 0.000597871 | 0.120108864 | -0.603323258 |
| **Q9H0U9** | TSPYL1 | -0.188336964 | 3.273871526 | -5.189146731 | 0.001616734 | 0.175016668 | -0.710755468 |
| **P55008-1** | AIF1 | -0.184543473 | 3.191967887 | -5.115025594 | 0.001746009 | 0.17784671 | -0.818476819 |
| **Q96QE3-1** | ATAD5 | -0.111018345 | 3.185540434 | -5.67776847 | 0.00099109 | 0.145427831 | -0.825285578 |
| **Q53EL6-1** | PDCD4 | 0.199650084 | 3.029318109 | 5.598627379 | 0.001070662 | 0.146662208 | -0.828479251 |
| **P41134** | ID1 | -0.296657013 | 3.277092086 | -5.014527125 | 0.001940232 | 0.185724587 | -0.841039598 |
| **Q8WWV3-1** | RTN4IP1 | 0.245549879 | 3.172033395 | 5.212120515 | 0.001578876 | 0.175016668 | -0.856358004 |
| **Q8TD26** | CHD6 | -0.089483462 | 3.193053966 | -5.782139085 | 0.000896159 | 0.137578741 | -0.866002792 |
| **P45844** | ABCG1 | -0.134490717 | 3.260538676 | -5.291421227 | 0.001455654 | 0.170076071 | -0.887608334 |
| **Q9NRM2** | ZNF277 | -0.286345426 | 3.325072223 | -4.928816321 | 0.002125157 | 0.191867897 | -0.909494848 |
| **Q14201-2** | BTG3 | -0.242712884 | 3.262173759 | -4.929985837 | 0.002122505 | 0.191867897 | -0.909627955 |
| **Q9H977** | WDR54 | -0.169206103 | 3.182359349 | -5.099050067 | 0.001775369 | 0.178211211 | -0.942342931 |
| **Q99755-3** | PIP5K1A | -0.123612874 | 3.237377878 | -5.076962429 | 0.00181688 | 0.178211211 | -1.009764904 |
| **Q9H1K1** | ISCU | 0.163530901 | 3.155083175 | 5.320198235 | 0.001413644 | 0.167632854 | -1.010869529 |
| **O60243-1** | HS6ST1 | -0.144462827 | 3.264444012 | -4.980832645 | 0.002010687 | 0.186559092 | -1.019599647 |
| **Q8N0T1** | C8orf59 | -0.235056098 | 3.253474172 | -4.99365481 | 0.001983543 | 0.186559092 | -1.036782541 |
| **Q96JP5-1** | ZFP91 | -0.119155502 | 3.241515069 | -5.184829966 | 0.001623961 | 0.175016668 | -1.06963306 |
| **O43561** | LAT | -0.235517227 | 3.253129027 | -4.818325412 | 0.002393362 | 0.198075676 | -1.070007 |
| **Q8IW45** | CARKD; NAXD | 0.136999617 | 3.15699127 | 5.142832439 | 0.001696197 | 0.175016668 | -1.074479854 |
| **Q6TFL4-1** | KLHL24 | 0.377114281 | 3.054338781 | 4.757292918 | 0.002557632 | 0.205256465 | -1.090491265 |
| **P01130** | LDLR | 0.122221389 | 3.040828934 | 5.327860945 | 0.001402689 | 0.167632854 | -1.117127593 |

| | | | | | | | |
|---|---|---|---|---|---|---|---|
| **Q9P2G9** | KLHL8 | -0.204866453 | 3.23083465 | -4.840633732 | 0.002336294 | 0.195387921 | -1.131611814 |
| **Q13137-4** | CALCOCO2 | -0.214518977 | 3.198957675 | -5.328606731 | 0.001401628 | 0.167632854 | -1.138023533 |
| **Q96CS2-1** | HAUS1 | -0.084824131 | 3.20113148 | -5.251706711 | 0.001515958 | 0.174554833 | -1.139840411 |
| **L0R8F8** | SMCR7L; MIEF1 | 0.183227447 | 3.168350825 | 4.684860476 | 0.002769168 | 0.21154847 | -1.19383184 |
| **Q02224** | CENPE | -0.104894222 | 3.177354998 | -5.474260233 | 0.001210699 | 0.158829039 | -1.214215708 |
| **P14866** | HNRNPL | 0.03792677 | 3.176788661 | 6.19762954 | 0.000607831 | 0.120108864 | -1.214785287 |
| **Q96CX6** | LRRC58 | -0.127934699 | 3.196163917 | -5.087090104 | 0.001797712 | 0.178211211 | -1.218124904 |
| **Q8NG31-1** | CASC5; KNL1 | -0.088132145 | 3.188682878 | -5.652174031 | 0.001016069 | 0.145427831 | -1.22345276 |
| **Q969M1** | TOMM40L | 0.276492747 | 3.182799479 | 4.634015577 | 0.002929334 | 0.211577775 | -1.228223673 |
| **Q14517** | FAT1 | -0.097545028 | 3.242048595 | -5.480526954 | 0.001203168 | 0.158829039 | -1.235325795 |
| **Q13422** | IKZF1 | -0.292176583 | 3.430748378 | -4.568783019 | 0.003150174 | 0.220369654 | -1.294289085 |
| **P41208** | CETN2 | -0.155951181 | 3.192845898 | -4.847736458 | 0.002318444 | 0.195387921 | -1.30989364 |
| **P49815** | TSC2 | -0.070332427 | 3.223260065 | -5.234621206 | 0.001542761 | 0.175016668 | -1.327562401 |
| **P46531** | NOTCH1 | -0.090228079 | 3.208543829 | -5.40169949 | 0.001301882 | 0.161616392 | -1.403396943 |
| **Q9UPQ4** | TRIM35 | -0.169130552 | 3.255318367 | -4.516876024 | 0.003339206 | 0.222941074 | -1.411511593 |
| **P14921-1** | ETS1 | 0.090271699 | 3.15249388 | 4.873741494 | 0.002254385 | 0.195387921 | -1.487365702 |
| **Q8NCA9** | ZNF784 | 0.907504871 | 3.123535768 | 4.640494572 | 0.00290836 | 0.211577775 | -1.495410311 |
| **Q9H3L0** | MMADHC | -0.155673473 | 3.250464232 | -4.544494332 | 0.003237099 | 0.221713378 | -1.534363621 |
| **Q13564-4** | NAE1 | -0.064775029 | 3.196133913 | -5.146836551 | 0.001689156 | 0.175016668 | -1.560824059 |
| **Q9UGP4** | LIMD1 | -0.096186785 | 3.21138841 | -4.894472303 | 0.002204736 | 0.195387921 | -1.565907741 |
| **Q69YH5-1** | CDCA2 | -0.095520998 | 3.179317229 | -4.976760911 | 0.002019394 | 0.186559092 | -1.575198632 |
| **Q3B7T1** | C10orf137; EDRF1 | -0.161490263 | 3.268206216 | -4.845317434 | 0.002324506 | 0.195387921 | -1.582252961 |
| **Q6ZRP7** | QSOX2 | -0.091205549 | 3.204678466 | -4.759660373 | 0.00255103 | 0.205256465 | -1.597596386 |
| **O95864** | FADS2 | -0.10583589 | 3.145885045 | -4.878062352 | 0.002243934 | 0.195387921 | -1.676593937 |
| **Q9UHH9** | IP6K2 | -0.325130595 | 3.264489209 | -4.206705771 | 0.004769361 | 0.25603089 | -1.696019233 |
| **Q8TAT5** | NEIL3 | -0.130538304 | 3.174241642 | -4.577231238 | 0.003120552 | 0.220369654 | -1.712088628 |
| **P62280** | RPS11 | -0.126948816 | 3.174995109 | -5.158966516 | 0.001668028 | 0.175016668 | -1.733172768 |
| **P18887** | XRCC1 | 0.074018099 | 3.172852258 | 5.032384123 | 0.001904018 | 0.184480797 | -1.789716533 |
| **Q9Y6A5** | TACC3 | -0.104562939 | 3.166060437 | -4.858605367 | 0.002291426 | 0.195387921 | -1.795744577 |
| **P05091** | ALDH2 | 0.240845808 | 3.171088468 | 4.353946841 | 0.004019715 | 0.238753388 | -1.815763969 |
| **O76080** | ZFAND5 | 0.129171491 | 3.205660678 | 4.565433189 | 0.003162006 | 0.220369654 | -1.824522227 |
| **Q9UBF8** | PI4KB | -0.102830568 | 3.213217751 | -4.686968357 | 0.002762743 | 0.21154847 | -1.843250252 |
| **Q15834** | CCDC85B | -0.213610606 | 3.291922717 | -4.054640475 | 0.005710082 | 0.262198515 | -1.847586442 |
| **Q15438-1** | CYTH1 | -0.151675065 | 3.172243661 | -4.462370813 | 0.00355142 | 0.22754864 | -1.848684912 |
| **Q9H8K7** | C10orf88 | -0.12806851 | 3.219786097 | -4.322425973 | 0.004168459 | 0.238753388 | -1.849502428 |
| **P50542-2** | PEX5 | -0.25986799 | 3.196572999 | -4.028232822 | 0.005893514 | 0.262198515 | -1.850342813 |
| **Q9UID6** | ZNF639 | -0.102838758 | 3.23391593 | -4.466155856 | 0.003536207 | 0.22754864 | -1.895199738 |
| **Q9UPR3** | SMG5 | -0.081003437 | 3.199473932 | -4.526915983 | 0.003301677 | 0.222303624 | -1.898779589 |
| **Q8NI60-1** | ADCK3; COQ8A | 0.241899726 | 3.13577697 | 4.501591548 | 0.003397253 | 0.223067579 | -1.901964403 |

| | | | | | | | |
|---|---|---|---|---|---|---|---|
| **P68402-1** | PAFAH1B2 | 0.26098333 | 3.106668958 | 4.410152115 | 0.00376891 | 0.236436902 | -1.915409487 |
| **Q9Y4E5-4** | ZNF451 | -0.183284342 | 3.236829378 | -4.003970976 | 0.006067795 | 0.267825749 | -1.930044909 |
| **Q14157-5** | UBAP2L | 0.209185898 | 3.165449621 | 4.039941726 | 0.00581139 | 0.262198515 | -1.938075429 |
| **Q9NQR1-2** | SETD8; KMT5A | -0.192216335 | 3.195543326 | -3.990983387 | 0.006163425 | 0.269057205 | -1.943703979 |
| **Q8TBE9** | NANP | -0.087595305 | 3.193472174 | -4.528366752 | 0.003296294 | 0.222303624 | -1.946080167 |
| **Q96GJ1** | TRMT2B | 0.207041274 | 3.186780333 | 4.30876526 | 0.004234815 | 0.238753388 | -1.94634322 |
| **Q9UBI9** | HECA | 0.140078431 | 3.160179476 | 4.300419615 | 0.00427593 | 0.238753388 | -1.961899039 |
| **P49366-1** | DHPS | -0.094329101 | 3.226209458 | -4.645910766 | 0.002890956 | 0.211577775 | -1.976907996 |
| **Q9H6S3-3** | EPS8L2 | -0.330168052 | 3.2279315 | -3.89204517 | 0.006949006 | 0.277900297 | -1.99126093 |
| **P03915** | ND5 | 0.323652339 | 3.182519145 | 3.891363241 | 0.00695479 | 0.277900297 | -1.993916839 |
| **Q9HA77** | CARS2 | 0.17183476 | 3.216469659 | 4.645231473 | 0.002893132 | 0.211577775 | -1.998725583 |
| **O43683** | BUB1 | -0.132260874 | 3.141940057 | -4.73673659 | 0.00261577 | 0.207822936 | -2.010233951 |
| **Q9H009** | NACA2 | -0.342333802 | 3.249332134 | -3.921860875 | 0.006701216 | 0.277900297 | -2.014979247 |
| **Q96DP5** | MTFMT | 0.257841189 | 3.200767286 | 3.889939819 | 0.006966881 | 0.277900297 | -2.015686954 |
| **P16403** | HIST1H1C | -0.156522808 | 3.179869443 | -4.506909008 | 0.003376932 | 0.223067579 | -2.039349001 |
| **Q06945** | SOX4 | 0.12746851 | 3.200368177 | 4.268592836 | 0.004436848 | 0.243996877 | -2.051584405 |
| **Q96R06** | SPAG5 | -0.079743018 | 3.177493359 | -4.704196845 | 0.002710845 | 0.211153546 | -2.054578196 |
| **Q9NSP4-1** | CENPM | -0.158500171 | 3.237491698 | -4.298426402 | 0.004285815 | 0.238753388 | -2.060788618 |
| **O00767** | SCD | -0.237276273 | 3.132262221 | -4.027331233 | 0.00589989 | 0.262198515 | -2.06426682 |
| **Q99767-1** | APBA2 | -0.113068873 | 3.210057451 | -4.115215501 | 0.005312719 | 0.262198515 | -2.077300246 |
| **O75964** | ATP5L | 0.227472605 | 3.186113529 | 4.070495856 | 0.005602987 | 0.262198515 | -2.07933897 |
| **P51530** | DNA2 | -0.106670074 | 3.20758842 | -4.607468377 | 0.003017052 | 0.215950234 | -2.105032899 |
| **Q13823** | GNL2 | -0.113895807 | 3.236374075 | -4.766206588 | 0.002532873 | 0.205256465 | -2.130973168 |
| **Q9BV79** | MECR | 0.18854764 | 3.197064618 | 4.086041047 | 0.005500142 | 0.262198515 | -2.15037027 |
| **Q16589** | CCNG2 | 0.825375293 | 2.810338284 | 4.347065056 | 0.004051679 | 0.238753388 | -2.164192681 |
| **Q92506** | HSD17B8 | 0.188953264 | 3.169061935 | 4.265457957 | 0.004453058 | 0.243996877 | -2.170088181 |
| **P61247** | RPS3A | -0.134479232 | 3.179938195 | -5.1466257 | 0.001689526 | 0.175016668 | -2.1767923 |
| **Q15120-2** | PDK3 | 0.166278924 | 3.194530665 | 4.194600487 | 0.004837583 | 0.25795034 | -2.190726746 |
| **Q9BQ61** | C19orf43; TRIR | 0.100383092 | 3.158906291 | 4.407714306 | 0.003779419 | 0.236436902 | -2.194905171 |
| **Q15119-1** | PDK2 | 0.289670863 | 3.129281586 | 4.05307937 | 0.005720748 | 0.262198515 | -2.201084573 |
| **P68543** | UBXN2A | -0.185413827 | 3.252597725 | -3.817594702 | 0.007612991 | 0.28115912 | -2.20458463 |
| **Q9Y512** | SAMM50 | 0.177724533 | 3.1830045 | 4.37626491 | 0.003917967 | 0.238753388 | -2.239309038 |
| **P52815** | MRPL12 | 0.2772029 | 3.153935142 | 4.323533842 | 0.004163128 | 0.238753388 | -2.252066067 |
| **Q9H3H5-1** | DPAGT1 | -0.223464793 | 3.193916182 | -3.689232569 | 0.008927891 | 0.286465393 | -2.263183052 |
| **P35222** | CTNNB1 | -0.10244951 | 3.181178811 | -4.257058084 | 0.004496819 | 0.24470704 | -2.278703185 |
| **O43678** | NDUFA2 | 0.234135689 | 3.190971852 | 3.883438637 | 0.007022398 | 0.277900297 | -2.29926193 |
| **P52429** | DGKE | -0.081719682 | 3.206308228 | -4.101752136 | 0.005398317 | 0.262198515 | -2.30996022 |
| **Q8TCF1-1** | ZFAND1 | 0.274572599 | 3.209826539 | 3.594007466 | 0.010064292 | 0.293973515 | -2.312888981 |
| **Q9UNF1** | MAGED2 | -0.081087873 | 3.203774392 | -4.4541062 | 0.003584891 | 0.227855671 | -2.320265226 |

| | | | | | | | |
|---|---|---|---|---|---|---|---|
| **Q96C34-1** | RUNDC1 | -0.180294542 | 3.142140979 | -3.828586981 | 0.007510703 | 0.28115912 | -2.323749632 |
| **Q9Y421-1** | FAM32A | -0.120930359 | 3.170179909 | -3.900831755 | 0.006874949 | 0.277900297 | -2.337474891 |
| **Q9NYK6-1** | C21orf91 | -0.218980382 | 3.247607396 | -3.765315533 | 0.008120911 | 0.282982452 | -2.338025765 |
| **Q9UBU8-1** | MORF4L1 | -0.093376406 | 3.22847529 | -4.318125041 | 0.004189225 | 0.238753388 | -2.348772914 |
| **Q9H6K4** | OPA3 | 0.182329151 | 3.175982956 | 3.752941207 | 0.008246515 | 0.282982452 | -2.354562352 |
| **Q9UHQ4-2** | BCAP29 | -0.113447922 | 3.18110618 | -3.867816572 | 0.007157809 | 0.277900297 | -2.355650327 |
| **P51798-1** | CLCN7 | -0.144732322 | 3.190407656 | -3.682571244 | 0.009002631 | 0.286465393 | -2.367417473 |
| **Q13309-1** | SKP2 | -0.095604619 | 3.192623713 | -4.05336882 | 0.005718769 | 0.262198515 | -2.374771936 |
| **O94880** | PHF14 | -0.137033673 | 3.14892975 | -3.812105099 | 0.007664647 | 0.28115912 | -2.374860571 |
| **Q0D2I5-5** | IFFO1 | -0.127000375 | 3.210889878 | -3.718909147 | 0.008603098 | 0.283479686 | -2.382660584 |
| **O96020** | CCNE2 | -0.145590401 | 3.246979799 | -3.734146683 | 0.00844139 | 0.282982452 | -2.383750119 |
| **Q9NVR0** | KLHL11 | -0.091454537 | 3.224296662 | -4.09990504 | 0.005410179 | 0.262198515 | -2.387229611 |
| **Q9HCJ0-2** | TNRC6C | -0.057541828 | 3.199233992 | -4.387797015 | 0.003866518 | 0.238135529 | -2.394729135 |
| **Q5RI15-2** | COX20 | 0.297265395 | 3.201291573 | 3.510026408 | 0.011198703 | 0.299378139 | -2.406574107 |
| **O15091** | KIAA0391 | 0.231323479 | 3.165727002 | 3.919987398 | 0.006716496 | 0.277900297 | -2.413205787 |
| **P41002** | CCNF | -0.356963919 | 3.131672203 | -3.583731872 | 0.010196094 | 0.295122294 | -2.440253902 |
| **Q6P4A7-1** | SFXN4 | 0.168803514 | 3.197413198 | 3.913179846 | 0.006772341 | 0.277900297 | -2.454307862 |
| **P15880** | RPS2 | -0.14428683 | 3.199757164 | -4.658132757 | 0.002852108 | 0.211577775 | -2.462356904 |
| **Q9H4H8** | FAM83D | -0.126732121 | 3.21416164 | -3.699339855 | 0.008815786 | 0.285883345 | -2.489451788 |
| **P29084** | GTF2E2 | -0.100828519 | 3.176721399 | -4.07218045 | 0.00559174 | 0.262198515 | -2.489882124 |
| **Q8NB46** | ANKRD52 | -0.092183689 | 3.188440887 | -3.950609176 | 0.006471499 | 0.277900297 | -2.495740557 |
| **Q9BSK4** | FEM1A | -0.138841312 | 3.267665218 | -3.70625221 | 0.008740007 | 0.284587526 | -2.498465272 |
| **Q9UIF9-1** | BAZ2A | -0.060051573 | 3.178043018 | -4.155983681 | 0.005062555 | 0.262198515 | -2.49863369 |
| **Q9UH92** | MLX | -0.125670267 | 3.241223581 | -3.654640077 | 0.009323572 | 0.28710988 | -2.512035255 |
| **Q9NW81-4** | ATP5SL; DMAC2 | 0.221642185 | 3.19687883 | 3.449950451 | 0.012095706 | 0.299378139 | -2.515028 |
| **O95478** | NSA2 | -0.234489709 | 3.257997095 | -3.718986573 | 0.008602268 | 0.283479686 | -2.517231873 |
| **P11387** | TOP 1.00 | -0.134280395 | 3.196650833 | -4.715360621 | 0.002677798 | 0.210644595 | -2.517796397 |
| **P04920-1** | SLC4A2 | -0.107962723 | 3.211615334 | -3.926700561 | 0.006661922 | 0.277900297 | -2.521528043 |
| **Q9BZ67** | FRMD8 | -0.106094268 | 3.200266758 | -3.805860252 | 0.007723879 | 0.28115912 | -2.522120583 |
| **Q96S59-1** | RANBP9 | -0.080588768 | 3.221960084 | -4.072113183 | 0.005592189 | 0.262198515 | -2.527251291 |
| **Q9BZ23** | PANK2 | -0.115221533 | 3.185088094 | -3.831202241 | 0.007486591 | 0.28115912 | -2.532165102 |
| **Q9HB19** | PLEKHA2 | -0.068514967 | 3.188481767 | -4.139439011 | 0.005162469 | 0.262198515 | -2.536342574 |
| **Q9P2B7** | KIAA1430; CFAP97 | -0.117484242 | 3.231677465 | -3.709233043 | 0.00870755 | 0.284587526 | -2.537961795 |
| **Q15417** | CNN3 | -0.206425101 | 3.218710968 | -3.393878801 | 0.013003942 | 0.304707969 | -2.556367433 |
| **Q9NP64** | ZCCHC17 | -0.11042586 | 3.18773386 | -3.807848039 | 0.00770497 | 0.28115912 | -2.578213498 |
| **P23588** | EIF4B | -0.119597558 | 3.186350551 | -4.300514499 | 0.00427546 | 0.238753388 | -2.578639959 |
| **O60930** | RNASEH1 | 0.085943743 | 3.183084087 | 3.814656336 | 0.007640592 | 0.28115912 | -2.58637091 |
| **Q7Z4G4** | TRMT11 | 0.158042269 | 3.195584031 | 3.458867622 | 0.011957747 | 0.299378139 | -2.596239297 |
| **Q9NX20** | MRPL16 | 0.193052666 | 3.19902078 | 3.875865266 | 0.007087688 | 0.277900297 | -2.596610659 |

| | | | | | | | |
|---|---|---|---|---|---|---|---|
| **Q9BR61** | ACBD6 | -0.082860737 | 3.196159366 | -4.113107851 | 0.005326019 | 0.262198515 | -2.597089044 |
| **Q16533** | SNAPC1 | -0.150514795 | 3.17718398 | -3.480669058 | 0.011627626 | 0.299378139 | -2.601581158 |
| **Q9NXV2** | KCTD5 | -0.155666209 | 3.256923475 | -3.798411455 | 0.00779519 | 0.281512655 | -2.605469674 |
| **P98170** | XIAP | -0.084335786 | 3.221476174 | -3.807203716 | 0.007711094 | 0.28115912 | -2.611834815 |
| **Q15782-4** | CHI3L2 | 0.252478549 | 2.98063582 | 3.461222656 | 0.011921599 | 0.299378139 | -2.620301086 |
| **O00330-1** | PDHX | 0.191475433 | 3.201888829 | 3.905097719 | 0.006839308 | 0.277900297 | -2.625157139 |
| **P31689-1** | DNAJA1 | -0.056091965 | 3.234559882 | -4.462964129 | 0.00354903 | 0.22754864 | -2.648498644 |
| **O60678** | PRMT3 | -0.060920975 | 3.210380635 | -4.156115023 | 0.005061771 | 0.262198515 | -2.664033984 |
| **P37268** | FDFT1 | 0.057885669 | 3.09143083 | 4.358270736 | 0.003999775 | 0.238753388 | -2.674606981 |
| **Q9BY43-2** | CHMP4A | -0.0875898 | 3.204423641 | -4.115423213 | 0.00531141 | 0.262198515 | -2.679846819 |
| **Q9NZJ0** | DTL | -0.074348556 | 3.229599679 | -4.053395969 | 0.005718583 | 0.262198515 | -2.690784787 |
| **Q8N4T8** | CBR4 | 0.242365048 | 3.172355363 | 3.50035216 | 0.011338108 | 0.299378139 | -2.691305676 |
| **Q969V5** | MUL1 | 0.198708659 | 3.16215449 | 3.303283108 | 0.014632037 | 0.305857884 | -2.702717165 |
| **Q9H147** | DNTTIP1 | 0.206517947 | 3.178467624 | 3.44482504 | 0.012175786 | 0.300424289 | -2.704069735 |
| **Q16763** | UBE2S | -0.080043323 | 3.203145374 | -4.026284258 | 0.005907304 | 0.262198515 | -2.709042109 |
| **Q96EV2** | RBM33 | 0.098819819 | 3.156529674 | 4.057038865 | 0.005693738 | 0.262198515 | -2.715536494 |
| **O15217** | GSTA4 | -0.32199236 | 3.1463332 | -3.230546558 | 0.016099449 | 0.305857884 | -2.722872144 |
| **Q6PGN9-1** | PSRC1 | -0.165393318 | 3.174125707 | -3.374507252 | 0.013334759 | 0.305857884 | -2.729785686 |
| **O75170-5** | PPP6R2 | 0.057655143 | 3.194493885 | 4.050459824 | 0.005738696 | 0.262198515 | -2.747222853 |
| **P10515** | DLAT | 0.215412611 | 3.212612418 | 3.871431027 | 0.007126225 | 0.277900297 | -2.751739361 |
| **Q13946-1** | PDE7A | 0.047240845 | 3.157698086 | 4.151900474 | 0.005087013 | 0.262198515 | -2.752086046 |
| **Q8NCC3** | PLA2G15 | 0.133515076 | 3.190367652 | 3.472989115 | 0.011742769 | 0.299378139 | -2.757303317 |
| **Q9NYP9** | MIS18A | -0.120141438 | 3.223248461 | -3.750357106 | 0.008273013 | 0.282982452 | -2.758959482 |
| **Q9ULD5-3** | ZNF777 | 0.116365985 | 3.177421886 | 3.43130603 | 0.012389795 | 0.302793718 | -2.762288651 |
| **Q6P2H3-1** | CEP85 | -0.125756903 | 3.209318137 | -3.629288129 | 0.009625759 | 0.289956215 | -2.762831134 |
| **P51553-1** | IDH3G | 0.197851682 | 3.206377959 | 3.756512527 | 0.008210047 | 0.282982452 | -2.774410184 |
| **Q15629** | TRAM1 | 0.071985261 | 3.216960665 | 4.054458632 | 0.005711323 | 0.262198515 | -2.778532716 |
| **Q9Y5P8-1** | PPP2R3B | -0.159246397 | 3.192469739 | -3.518629706 | 0.0110763 | 0.299378139 | -2.787022115 |
| **Q9Y3S2** | ZNF330 | -0.096621541 | 3.218421817 | -3.747758183 | 0.008299758 | 0.282982452 | -2.81203248 |
| **Q9HBU6-1** | ETNK1 | -0.126213603 | 3.185607418 | -3.933868112 | 0.006604194 | 0.277900297 | -2.821687511 |
| **Q14469** | HES1 | -0.20311389 | 3.16453717 | -3.234441015 | 0.016016968 | 0.305857884 | -2.825899103 |
| **Q9NRG7-1** | SDR39U1 | 0.208028158 | 3.167607544 | 3.327982204 | 0.014167273 | 0.305857884 | -2.826449542 |
| **P26599-3** | PTBP1 | 0.060103301 | 3.191537865 | 4.549035764 | 0.003220645 | 0.221713378 | -2.831563066 |
| **Q9Y2Y0** | ARL2BP | -0.092810857 | 3.176875591 | -3.89737147 | 0.006904009 | 0.277900297 | -2.832133819 |
| **O95872** | GPANK1 | 0.212552502 | 3.189991106 | 3.389490744 | 0.013078089 | 0.304707969 | -2.833862783 |
| **Q71RC2-4** | LARP4 | -0.108881392 | 3.267330495 | -3.931696509 | 0.006621626 | 0.277900297 | -2.839070466 |
| **Q6ZTW0-1** | TPGS1 | -0.154497857 | 3.273035056 | -3.298518024 | 0.014723592 | 0.305857884 | -2.856032925 |
| **O14880** | MGST3 | 0.291263075 | 3.181446403 | 3.424700505 | 0.012495849 | 0.302793718 | -2.864564891 |
| **Q9H9B4** | SFXN1 | 0.119897105 | 3.185222378 | 3.934238278 | 0.006601228 | 0.277900297 | -2.866345736 |

| | | | | | | | |
|---|---|---|---|---|---|---|---|
| **Q8NHZ8** | CDC26 | -0.383325255 | 3.172307046 | -3.379595327 | 0.01324699 | 0.305857884 | -2.86970387 |
| **Q8IUE6** | HIST2H2AB | 0.220766146 | 3.192356997 | 3.360781266 | 0.013574701 | 0.305857884 | -2.871284346 |
| **Q86YD1-1** | PTOV1 | -0.125768334 | 3.271841678 | -3.289566085 | 0.014897276 | 0.305857884 | -2.883131806 |
| **Q96CK0** | ZNF653 | -0.186602206 | 3.296300527 | -3.148342947 | 0.017952065 | 0.305857884 | -2.887791089 |
| **O75208-1** | COQ9 | 0.165963928 | 3.180475164 | 3.364666381 | 0.013506313 | 0.305857884 | -2.897451465 |
| **Q9UKT4** | FBXO5 | -0.165085753 | 3.18610469 | -3.66982319 | 0.009147583 | 0.28710988 | -2.905088522 |
| **P27816-1** | MAP4 | -0.150647689 | 3.195193866 | -4.296125047 | 0.00429726 | 0.238753388 | -2.905416782 |
| **Q9UHQ7** | WBP5; TCEAL9 | -0.50337405 | 2.968480389 | -3.085161251 | 0.019532185 | 0.305857884 | -2.91039315 |
| **Q5T890-1** | ERCC6L2 | 0.337427975 | 3.191580999 | 3.070562233 | 0.0199182 | 0.305857884 | -2.911537823 |
| **O94906-1** | PRPF6 | -0.048517023 | 3.184366678 | -4.332240844 | 0.004121497 | 0.238753388 | -2.91943901 |
| **Q9HCM4** | EPB41L5 | -0.094663613 | 3.225878643 | -3.486839681 | 0.011536004 | 0.299378139 | -2.924432294 |
| **Q8N490-2** | PNKD | 0.211109804 | 3.175075107 | 3.310735337 | 0.014490089 | 0.305857884 | -2.92792866 |
| **Q9GZR7** | DDX24 | -0.103560894 | 3.170622938 | -4.227048189 | 0.004657106 | 0.251705502 | -2.931927861 |
| **P62829** | RPL23 | -0.134644271 | 3.173849699 | -4.032106372 | 0.005866204 | 0.262198515 | -2.932013307 |
| **O43615** | TIMM44 | 0.21209971 | 3.193275992 | 3.916098676 | 0.006748334 | 0.277900297 | -2.941611018 |
| **Q16611** | BAK1 | 0.129125422 | 3.16651876 | 3.394799735 | 0.012988439 | 0.304707969 | -2.942506428 |
| **O15165** | LDLRAD4 | -0.142797302 | 3.203700926 | -3.250647059 | 0.01567862 | 0.305857884 | -2.946866376 |
| **P79522-2** | PRR3 | 0.184415712 | 3.125787399 | 3.256984296 | 0.015548423 | 0.305857884 | -2.948895363 |
| **P83876** | TXNL4A | -0.068887762 | 3.182985679 | -3.662779707 | 0.009228767 | 0.28710988 | -2.955122965 |
| **P04179** | SOD2 | 0.202607513 | 3.181174447 | 3.85518448 | 0.007269408 | 0.2802201 | -2.964065227 |
| **Q9UHV7** | MED13 | 0.072448535 | 3.166188029 | 3.715975172 | 0.008634624 | 0.283479686 | -2.970652509 |
| **Q03013-1** | GSTM4 | 0.109054495 | 3.145465623 | 3.341983129 | 0.013910953 | 0.305857884 | -2.981515868 |
| **Q8TEL6-3** | TRPC4AP | -0.112709355 | 3.206611826 | -3.45868039 | 0.011960626 | 0.299378139 | -2.988780634 |
| **Q14761** | PTPRCAP | -0.112721676 | 3.235574845 | -3.419654477 | 0.012577531 | 0.302793718 | -3.000516984 |
| **Q9UJZ1** | STOML2 | 0.215497874 | 3.189618873 | 3.650391771 | 0.009373477 | 0.28710988 | -3.005096146 |
| **Q5T5X7** | BEND3 | -0.060499728 | 3.18263983 | -3.831754744 | 0.007481508 | 0.28115912 | -3.007046604 |
| **Q5ST30-4** | VARS2 | 0.21382065 | 3.188782719 | 3.750773787 | 0.008268734 | 0.282982452 | -3.012947997 |
| **Q9Y5Z0-1** | BACE2 | -0.198785868 | 3.191107025 | -3.144958374 | 0.018033121 | 0.305857884 | -3.015018012 |
| **Q9H000** | MKRN2 | -0.096965558 | 3.20261306 | -3.630640168 | 0.009609375 | 0.289956215 | -3.017872274 |
| **Q9H5V7** | IKZF5 | 0.102138119 | 3.231730667 | 3.39039605 | 0.013062754 | 0.304707969 | -3.0208574 |
| **P24723** | PRKCH | -0.094644189 | 3.185356952 | -3.83106086 | 0.007487892 | 0.28115912 | -3.021932941 |
| **O75794** | CDC123 | -0.078993212 | 3.259843684 | -3.769802106 | 0.008075891 | 0.282982452 | -3.025094212 |
| **Q7Z4L5** | TTC21B | -0.172046258 | 3.243788959 | -3.133141378 | 0.018319231 | 0.305857884 | -3.026812963 |
| **Q96CP6-1** | GRAMD1A | -0.111426981 | 3.191840243 | -3.223871885 | 0.016241883 | 0.305857884 | -3.030138324 |
| **Q8ND25-2** | ZNRF1 | -0.086379247 | 3.222970094 | -3.500923172 | 0.011329828 | 0.299378139 | -3.034801237 |
| **Q8N6I1-1** | EID2 | -0.152215381 | 3.261241203 | -3.110436118 | 0.018882794 | 0.305857884 | -3.034893002 |
| **Q9BVW5** | TIPIN | -0.180188987 | 3.211818466 | -3.534371446 | 0.010856103 | 0.299378139 | -3.0350499 |
| **Q9H6Y7** | RNF167 | -0.121850123 | 3.191148571 | -3.219502339 | 0.016335865 | 0.305857884 | -3.03567567 |
| **P35241-5** | RDX | -0.083906993 | 3.177907303 | -3.779106893 | 0.007983396 | 0.282982452 | -3.038140066 |

| | | | | | | | |
|---|---|---|---|---|---|---|---|
| **Q8IVH4** | MMAA | 0.216829879 | 3.172328412 | 3.412701255 | 0.012691039 | 0.302793718 | -3.04194027 |
| **P20700** | LMNB1 | 0.053518692 | 3.175832038 | 4.636376354 | 0.002921672 | 0.211577775 | -3.050678098 |
| **P26583** | HMGB2 | 0.123095768 | 3.172890627 | 4.39959402 | 0.003814663 | 0.236777323 | -3.05089277 |
| **Q9NYR9-1** | NKIRAS2 | -0.127164446 | 3.190469414 | -3.480380207 | 0.011631934 | 0.299378139 | -3.053846753 |
| **P53370** | NUDT6 | 0.184653298 | 3.199828332 | 3.040394239 | 0.020742118 | 0.306882921 | -3.057368266 |
| **O75251** | NDUFS7 | 0.147278862 | 3.187986222 | 3.535987499 | 0.010833769 | 0.299378139 | -3.058121874 |
| **P26367-2** | PAX6 | 0.207220009 | 3.195247753 | 2.991834542 | 0.022146184 | 0.312352362 | -3.068552197 |
| **Q7Z6M4** | MTERFD2; MTERF4 | 0.220773004 | 3.179611635 | 3.089628205 | 0.019415688 | 0.305857884 | -3.075726944 |
| **O00444** | PLK4 | 0.189527002 | 3.201330103 | 2.949182354 | 0.023463641 | 0.320063886 | -3.077576803 |
| **Q86YA3-1** | C4orf21; ZGRF1 | -0.094080016 | 3.175093568 | -3.600625319 | 0.009980394 | 0.293973515 | -3.089027598 |
| **P35268** | RPL22 | -0.070903748 | 3.206723856 | -3.556598007 | 0.010553298 | 0.298384176 | -3.089179162 |
| **P53597** | SUCLG1 | 0.223372677 | 3.187555272 | 3.851659701 | 0.007300889 | 0.2802201 | -3.093660575 |
| **Q9H6L5** | FAM134B; RETREG | -0.124684231 | 3.251154285 | -3.45977082 | 0.01194387 | 0.299378139 | -3.09693833 |
| **Q6IAA8** | LAMTOR1 | -0.101563964 | 3.189021291 | -3.624315679 | 0.009686278 | 0.290405592 | -3.09702077 |
| **Q96I99-1** | SUCLG2 | 0.176825974 | 3.178585247 | 3.720292089 | 0.008588282 | 0.283479686 | -3.10142163 |
| **Q9NQ50** | MRPL40 | 0.187872386 | 3.21149263 | 3.450211969 | 0.012091635 | 0.299378139 | -3.10206702 |
| **O95229** | ZWINT | -0.201513174 | 3.203104203 | -3.373345329 | 0.013354891 | 0.305857884 | -3.103545209 |
| **Q96CQ1** | SLC25A36 | 0.276771792 | 3.192971717 | 2.924887019 | 0.024251378 | 0.322917088 | -3.114893074 |
| **Q8N0X4** | CLYBL | 0.188498499 | 3.208523507 | 3.256575572 | 0.015556785 | 0.305857884 | -3.131554332 |
| **Q14106** | TOB2 | -0.247984613 | 3.216474695 | -2.870434723 | 0.026121395 | 0.328952544 | -3.137508001 |
| **Q9HC21** | SLC25A19 | 0.288035555 | 3.209953519 | 3.249253394 | 0.01570741 | 0.305857884 | -3.140060907 |
| **Q07864** | POLE | -0.053331772 | 3.184032277 | -4.078147333 | 0.005552102 | 0.262198515 | -3.141280082 |
| **Q9UMX1** | SUFU | -0.174976296 | 3.200376824 | -2.910493539 | 0.024731361 | 0.32394477 | -3.148413245 |
| **Q99611** | SEPHS2 | 0.100242836 | 3.193083098 | 3.317066268 | 0.014370672 | 0.305857884 | -3.150478817 |
| **Q9BX63** | BRIP1 | -0.068665299 | 3.165300952 | -3.866370062 | 0.007170492 | 0.277900297 | -3.153036097 |
| **Q9HAW4** | CLSPN | -0.07328171 | 3.192975422 | -3.803121736 | 0.007750012 | 0.28115912 | -3.156114128 |
| **P08069** | IGF1R | -0.587049536 | 3.192351203 | -3.091275265 | 0.019372922 | 0.305857884 | -3.160533249 |
| **Q9H7P6** | FAM125B; MVB12E | -0.11166342 | 3.192717785 | -3.129379234 | 0.018411343 | 0.305857884 | -3.162583163 |
| **Q9Y2I1-1** | NISCH | -0.055561862 | 3.182155762 | -3.736360032 | 0.00841818 | 0.282982452 | -3.172464411 |
| **Q13625-3** | TP53BP2 | -0.061168561 | 3.200772061 | -3.736360245 | 0.008418177 | 0.282982452 | -3.186346039 |
| **O95139** | NDUFB6 | 0.276314211 | 3.211421128 | 3.021212105 | 0.021284994 | 0.309578361 | -3.196765486 |
| **Q96NH3-4** | C6orf170; TBC1D32 | -0.255375009 | 3.110339713 | -2.799638873 | 0.028785637 | 0.331452005 | -3.198817569 |
| **Q4LDG9-1** | DNAL1 | -0.094213228 | 3.19961029 | -3.536569236 | 0.010825742 | 0.299378139 | -3.199045108 |
| **Q8IYB3** | SRRM1 | 0.053821888 | 3.175108766 | 3.747545662 | 0.008301949 | 0.282982452 | -3.201352957 |
| **Q9HB20** | PLEKHA3 | 0.137213291 | 3.180740052 | 3.20730516 | 0.016601333 | 0.305857884 | -3.201381371 |
| **P60228** | EIF3E | -0.076755063 | 3.205636866 | -3.993971161 | 0.006141279 | 0.269057205 | -3.202045298 |
| **Q96RS0** | TGS1 | -0.08903047 | 3.201502456 | -3.752957887 | 0.008246345 | 0.282982452 | -3.205805187 |
| **P31937** | HIBADH | 0.212566265 | 3.169664458 | 3.306530679 | 0.014569994 | 0.305857884 | -3.209254091 |
| **Q9ULJ3** | ZBTB21; ZNF295 | -0.085170552 | 3.194292567 | -3.578281086 | 0.010266775 | 0.295122294 | -3.218960249 |

| | | | | | | | |
|---|---|---|---|---|---|---|---|
| **Q9UJ04** | TSPYL4 | -0.271289913 | 3.223968175 | -2.779848918 | 0.029581005 | 0.334311646 | -3.220275125 |
| **P30038** | ALDH4A1 | 0.195031388 | 3.175886501 | 3.601921646 | 0.00996405 | 0.293973515 | -3.222795824 |
| **Q9BSH5** | HDHD3 | 0.217091358 | 3.195005271 | 3.119248655 | 0.018661872 | 0.305857884 | -3.224157922 |
| **Q96Q45-3** | TMEM237 | -0.151316331 | 3.200812176 | -2.900306669 | 0.025077192 | 0.32636083 | -3.232056585 |
| **Q8N163-1** | CCAR2; KIAA1967 | 0.067128171 | 3.148231876 | 4.177629138 | 0.00493506 | 0.261393654 | -3.233756338 |
| **Q9Y399** | MRPS2 | 0.2247428 | 3.19664367 | 3.108094164 | 0.018941978 | 0.305857884 | -3.239651145 |
| **Q9Y2S7** | POLDIP2 | 0.170959176 | 3.184889084 | 3.32986488 | 0.01413251 | 0.305857884 | -3.244284992 |
| **Q8NBN7** | RDH13 | 0.175979949 | 3.183740831 | 3.178870878 | 0.017238481 | 0.305857884 | -3.245329813 |
| **Q9ULA0** | DNPEP | 0.057684641 | 3.186314095 | 3.667223764 | 0.009177453 | 0.28710988 | -3.250868371 |
| **Q5VZL5** | ZMYM4 | -0.039997482 | 3.188119947 | -3.868286976 | 0.00715369 | 0.277900297 | -3.25133811 |
| **O75600-2** | GCAT | 0.182213248 | 3.146753181 | 3.286401094 | 0.014959213 | 0.305857884 | -3.261253969 |
| **P20023-3** | CR2 | 0.078625633 | 3.130757952 | 3.586549534 | 0.010159765 | 0.295122294 | -3.262161339 |
| **Q5THK1-1** | PRR14L | -0.052257152 | 3.21662669 | -3.571839025 | 0.010351003 | 0.295122294 | -3.266345516 |
| **Q9NUT2-1** | ABCB8 | 0.188381141 | 3.171079362 | 3.187158345 | 0.017050092 | 0.305857884 | -3.269710819 |
| **Q9BW92** | TARS2 | 0.218032591 | 3.170502328 | 3.573474045 | 0.010329554 | 0.295122294 | -3.278971344 |
| **A4D1E9-1** | GTPBP10 | 0.151067749 | 3.190729729 | 3.478298802 | 0.01166303 | 0.299378139 | -3.279144401 |
| **Q9BZE4** | GTPBP4 | -0.085214486 | 3.180093371 | -3.832637949 | 0.00747339 | 0.28115912 | -3.279172858 |
| **Q9Y6G9** | DYNC1LI1 | 0.055846023 | 3.167744026 | 4.048996802 | 0.005748747 | 0.262198515 | -3.283609129 |
| **O43736-1** | ITM2A | -0.067472519 | 3.225672965 | -3.523003056 | 0.01101464 | 0.299378139 | -3.287393202 |
| **Q9Y2Z9-1** | COQ6 | 0.23136433 | 3.187845475 | 3.232704915 | 0.016053681 | 0.305857884 | -3.289526078 |
| **Q6P1Q0-7** | LETMD1 | 0.273764948 | 3.173572206 | 2.749376919 | 0.030851614 | 0.33673269 | -3.292764441 |
| **Q9NWU2** | GID8 | -0.129406841 | 3.228367396 | -3.066626843 | 0.020023655 | 0.305857884 | -3.293771385 |
| **Q9Y2C4** | EXOG | 0.18786352 | 3.217966347 | 3.056550642 | 0.020296409 | 0.305857884 | -3.294795268 |
| **P16219** | ACADS | 0.197447169 | 3.173471507 | 3.520453527 | 0.01105054 | 0.299378139 | -3.294868477 |
| **P41732** | TSPAN7 | -0.146044071 | 3.176755531 | -3.482789908 | 0.011596046 | 0.299378139 | -3.296910768 |
| **Q7KZN9** | COX15 | 0.227630578 | 3.197926375 | 3.164036965 | 0.017581319 | 0.305857884 | -3.300468662 |
| **Q5W0B1** | RNF219 | -0.111913076 | 3.229251925 | -3.568058332 | 0.010400786 | 0.295122294 | -3.304758676 |
| **O95182** | NDUFA7 | 0.232460325 | 3.185022207 | 3.385869002 | 0.013139634 | 0.305246758 | -3.307264025 |
| **O75521** | ECI2 | 0.084907257 | 3.174595837 | 3.512474849 | 0.011163718 | 0.299378139 | -3.318249781 |
| **Q562F6** | SGOL2; SGO2 | -0.161618499 | 3.185471605 | -3.226900131 | 0.016177094 | 0.305857884 | -3.321316966 |
| **Q9UNZ5** | C19orf53 | -0.12428641 | 3.202084484 | -3.079304737 | 0.019686063 | 0.305857884 | -3.325098399 |
| **P15104** | GLUL | -0.204627128 | 3.231865104 | -3.283780906 | 0.0150107 | 0.305857884 | -3.32921703 |
| **P23786** | CPT2 | 0.181509444 | 3.173434025 | 3.397069979 | 0.012950306 | 0.304707969 | -3.332882159 |
| **O95613** | PCNT | -0.044868173 | 3.188112512 | -4.073280022 | 0.005584412 | 0.262198515 | -3.335707857 |
| **P61086** | UBE2K | -0.048716001 | 3.151655119 | -3.669608113 | 0.00915005 | 0.28710988 | -3.340496565 |
| **P30679** | GNA15 | -0.09546892 | 3.180055051 | -3.504816314 | 0.011273546 | 0.299378139 | -3.341809179 |
| **Q9HBT8** | ZNF286A | -0.274341358 | 3.236719232 | -2.672651838 | 0.034313669 | 0.345097821 | -3.345974906 |
| **Q8WVM0** | TFB1M | 0.160687103 | 3.190066596 | 2.989620105 | 0.022212597 | 0.312352362 | -3.349129419 |
| **P51587** | BRCA2 | -0.091852315 | 3.1974209 | -3.656783097 | 0.009298508 | 0.28710988 | -3.359909107 |

| | | | | | | | |
|---|---|---|---|---|---|---|---|
| **Q9HC77** | CENPJ | -0.102369485 | 3.189717349 | -3.091984431 | 0.01935454 | 0.305857884 | -3.361610983 |
| **P04183** | TK1 | -0.17304221 | 3.146365531 | -3.119565319 | 0.018653985 | 0.305857884 | -3.362375051 |
| **Q96GA3** | LTV1 | -0.066554358 | 3.213025729 | -3.685829803 | 0.008965985 | 0.286465393 | -3.363423259 |
| **Q02535** | ID3 | -0.170321381 | 3.160067426 | -2.700470856 | 0.033013187 | 0.339814312 | -3.370400948 |
| **Q9Y5B8** | NME7 | -0.072588102 | 3.188136392 | -3.470594633 | 0.011778922 | 0.299378139 | -3.372510849 |
| **P34932** | HSPA4 | 0.027853192 | 3.170628054 | 4.345393697 | 0.004059484 | 0.238753388 | -3.373689276 |
| **Q9UJS0-2** | SLC25A13 | 0.152189782 | 3.157998217 | 2.92542234 | 0.02423372 | 0.322917088 | -3.374091795 |
| **P85037-1** | FOXK1 | 0.059174255 | 3.169915009 | 3.607883186 | 0.009889262 | 0.293973515 | -3.379115211 |
| **Q9NY65** | TUBA8; EHHADH | -0.167356937 | 3.224685763 | -3.132041618 | 0.018346106 | 0.305857884 | -3.381054413 |
| **Q9NZJ6** | COQ3 | 0.251991285 | 3.191424305 | 3.177011516 | 0.017281055 | 0.305857884 | -3.383512625 |
| **Q9BT67** | NDFIP1 | 0.189051627 | 3.097446774 | 2.730441266 | 0.031670219 | 0.33673269 | -3.384646317 |
| **Q6P087** | RPUSD3 | 0.191643166 | 3.163624262 | 3.328169285 | 0.014163814 | 0.305857884 | -3.390934077 |
| **O00287** | RFXAP | -0.088323342 | 3.175984933 | -3.220654281 | 0.016311031 | 0.305857884 | -3.396975008 |
| **Q96I59** | NARS2 | 0.221781551 | 3.193231616 | 3.425446619 | 0.01248382 | 0.302793718 | -3.401186396 |
| **Q8N9N8** | EIF1AD | -0.101969676 | 3.20780923 | -3.157793114 | 0.017727821 | 0.305857884 | -3.403214478 |
| **Q9GZT6** | CCDC90B | 0.16532879 | 3.199718093 | 3.213481173 | 0.016466335 | 0.305857884 | -3.405753111 |
| **Q86VE9-4** | SERINC5 | 0.20983091 | 3.115482237 | 2.760815947 | 0.030367979 | 0.336035644 | -3.405987955 |
| **Q96AQ6** | PBXIP1 | 0.129001845 | 2.982038272 | 3.24739108 | 0.015745971 | 0.305857884 | -3.414031071 |
| **Q9Y253-1** | POLH | -0.116210058 | 3.243504085 | -3.04290414 | 0.020672193 | 0.306620093 | -3.418518476 |
| **Q8WYP5-2** | AHCTF1 | -0.068675174 | 3.198669123 | -3.904052339 | 0.006848023 | 0.277900297 | -3.420034803 |
| **O95707** | POP4 | -0.136598101 | 3.198494339 | -3.339529198 | 0.013955512 | 0.305857884 | -3.420683532 |
| **Q3ZCQ8-2** | TIMM50 | 0.210925646 | 3.17296084 | 3.154709641 | 0.017800655 | 0.305857884 | -3.421125953 |
| **O14530** | TXNDC9 | -0.130466959 | 3.18206755 | -2.887347247 | 0.025524626 | 0.328144258 | -3.42128833 |
| **Q96AT1** | KIAA1143 | -0.059632571 | 3.185554405 | -3.483947311 | 0.011578851 | 0.299378139 | -3.423927185 |
| **Q9P1Q0** | VPS54 | -0.223948395 | 3.162590879 | -2.582295291 | 0.038923231 | 0.352501702 | -3.438026133 |
| **Q99798** | ACO2 | 0.189538735 | 3.183530075 | 3.759175679 | 0.008182968 | 0.282982452 | -3.443815586 |
| **Q16822-1** | PCK2 | 0.182983593 | 3.201884268 | 3.315840941 | 0.014393701 | 0.305857884 | -3.451017302 |
| **Q9H4A6** | GOLPH3 | 0.079877424 | 3.154550971 | 3.308991228 | 0.014523176 | 0.305857884 | -3.452054886 |
| **Q9H300** | PARL | 0.206453434 | 3.17832669 | 3.156401826 | 0.017760644 | 0.305857884 | -3.453919582 |
| **Q9Y375** | NDUFAF1 | 0.217068828 | 3.199864968 | 3.172771794 | 0.017378559 | 0.305857884 | -3.456208946 |
| **O94986-4** | CEP152 | -0.059747604 | 3.19643945 | -3.628543566 | 0.009634794 | 0.289956215 | -3.460165653 |
| **Q9BRT2** | MNF1; UQCC2 | 0.18918246 | 3.188660162 | 3.095937286 | 0.019252421 | 0.305857884 | -3.460876142 |
| **Q99643** | SDHC | 0.206002707 | 3.184560704 | 3.122621163 | 0.018578064 | 0.305857884 | -3.460993426 |
| **Q9UGM6-1** | WARS2 | 0.200353827 | 3.176471333 | 2.998032025 | 0.021961445 | 0.311022609 | -3.461219872 |
| **O95997** | PTTG1 | -0.106461031 | 3.143090176 | -2.976250295 | 0.022618131 | 0.315819071 | -3.468777558 |
| **O75880** | SCO1 | 0.266032308 | 3.213423389 | 3.068489583 | 0.019973666 | 0.305857884 | -3.469147516 |
| **Q6IN84-1** | MRM1 | 0.199378716 | 3.208422689 | 3.289365955 | 0.014901184 | 0.305857884 | -3.469745265 |
| **P24534** | EEF1B2 | -0.061299899 | 3.198376844 | -3.902772441 | 0.00685871 | 0.277900297 | -3.470548358 |
| **P78344-1** | EIF4G2 | 0.202643532 | 3.20176015 | 3.036109832 | 0.020862069 | 0.306942846 | -3.475633822 |

| | | | | | | | |
|---|---|---|---|---|---|---|---|
| **Q8N4Q0** | ZADH2 | 0.240600994 | 3.198297076 | 2.744539889 | 0.031058569 | 0.33673269 | -3.478252413 |
| **P56589** | PEX3 | -0.079938898 | 3.185246174 | -3.228909652 | 0.016134254 | 0.305857884 | -3.478608129 |
| **P62701** | RPS4X | -0.12472854 | 3.167352948 | -3.743257318 | 0.0083463 | 0.282982452 | -3.485805567 |
| **Q9Y2Q9** | MRPS28 | 0.106918233 | 3.209390975 | 2.912979987 | 0.024647725 | 0.32394477 | -3.489689847 |
| **P13804-1** | ETFA | 0.180430362 | 3.187365502 | 3.576356631 | 0.010291858 | 0.295122294 | -3.489867569 |
| **P50213-1** | IDH3A | 0.151019815 | 3.176196212 | 3.491848663 | 0.011462209 | 0.299378139 | -3.494057101 |
| **P04062** | GBA | -0.062589459 | 3.187605544 | -3.417204673 | 0.012617396 | 0.302793718 | -3.49499302 |
| **Q13287** | NMI | -0.057653465 | 3.200417498 | -3.681561913 | 0.009014015 | 0.286465393 | -3.496320747 |
| **Q9UPU7-1** | TBC1D2B | -0.066437017 | 3.207621171 | -3.348409332 | 0.013794997 | 0.305857884 | -3.500567352 |
| **Q99618** | CDCA3 | -0.241300091 | 3.140145072 | -2.540858266 | 0.041250526 | 0.355540374 | -3.507607907 |
| **P21912** | SDHB | 0.30586249 | 3.180713288 | 3.482146004 | 0.011605624 | 0.299378139 | -3.513107622 |
| **O75616-1** | ERAL1 | 0.258083851 | 3.147714695 | 3.257760397 | 0.015532559 | 0.305857884 | -3.517125066 |
| **P57086** | SCAND1 | -0.342629724 | 3.165187762 | -2.511331706 | 0.042997439 | 0.357204257 | -3.517254037 |
| **Q9H0H5** | RACGAP1 | -0.09005673 | 3.18207422 | -3.594764815 | 0.010054651 | 0.293973515 | -3.518739534 |
| **Q9BX70-1** | BTBD2 | -0.096912675 | 3.165709244 | -2.925164962 | 0.024242208 | 0.322917088 | -3.520984202 |
| **Q9H496** | TOR1AIP2 | -0.110484726 | 3.237295286 | -2.812432039 | 0.028283584 | 0.330820062 | -3.52163224 |
| **P55196-1** | MLLT4; AFDN | -0.283449203 | 3.234773707 | -2.492275825 | 0.04416575 | 0.357204257 | -3.532079817 |
| **Q9H6F5** | CCDC86 | -0.088370712 | 3.222323938 | -3.482398134 | 0.011601872 | 0.299378139 | -3.533056815 |
| **Q9BPW8** | NIPSNAP1 | 0.217308494 | 3.190575533 | 3.368640197 | 0.013436751 | 0.305857884 | -3.536579125 |
| **P19823** | ITIH2 | 0.105921852 | 3.254967214 | 2.828357734 | 0.027671569 | 0.32997678 | -3.539040072 |
| **Q9Y385** | UBE2J1 | 0.093606906 | 3.171401789 | 3.066509371 | 0.020026812 | 0.305857884 | -3.540164389 |
| **P07237** | P4HB | 0.020886273 | 3.194993594 | 4.14498157 | 0.005128756 | 0.262198515 | -3.540653102 |
| **Q8WVD3-1** | RNF138 | -0.07805787 | 3.207871193 | -3.032376489 | 0.020967199 | 0.307351282 | -3.541205165 |
| **Q9ULM6** | CNOT6 | 0.10763534 | 3.173412927 | 3.119456084 | 0.018656705 | 0.305857884 | -3.54506139 |
| **Q00987-11** | MDM2 | -0.263486105 | 3.178114466 | -2.819045065 | 0.02802772 | 0.330092368 | -3.549125175 |
| **P11117** | ACP2 | -0.096518988 | 3.191924958 | -3.122680631 | 0.01857659 | 0.305857884 | -3.550740605 |
| **O95989** | NUDT3 | 0.170047775 | 3.142999216 | 3.122228685 | 0.018587796 | 0.305857884 | -3.552222099 |
| **O60239** | SH3BP5 | -0.215049054 | 3.176697098 | -2.776286609 | 0.029726637 | 0.335004444 | -3.556266777 |
| **P27635** | RPL10 | -0.121265666 | 3.167491055 | -3.635407487 | 0.009551848 | 0.289956215 | -3.558137092 |
| **Q7Z5L2-2** | R3HCC1L | -0.079995727 | 3.198984269 | -3.06020703 | 0.020196975 | 0.305857884 | -3.55829782 |
| **Q9NRN9** | METTL5 | -0.053067216 | 3.211897244 | -3.290158869 | 0.014885706 | 0.305857884 | -3.562806224 |
| **P27695** | APEX1 | 0.104937257 | 3.170750496 | 3.618120832 | 0.00976226 | 0.291583285 | -3.563431319 |
| **Q567V2-1** | MPV17L2 | 0.184347789 | 3.20990378 | 2.597242318 | 0.038117887 | 0.352231442 | -3.567424729 |
| **P31040** | SDHA | 0.214935314 | 3.18934904 | 3.737797369 | 0.008403144 | 0.282982452 | -3.568092087 |
| **Q15032-2** | R3HDM1 | -0.146060161 | 3.134446155 | -2.719840819 | 0.032138497 | 0.338199147 | -3.570146583 |
| **Q9NRX2** | MRPL17 | 0.174660147 | 3.183015745 | 3.11307104 | 0.018816445 | 0.305857884 | -3.570682844 |
| **Q9BUE6-2** | ISCA1 | 0.208267684 | 3.160987684 | 2.621308746 | 0.036857759 | 0.348943902 | -3.575367607 |
| **Q9H1U9** | SLC25A51 | 0.144362932 | 3.228113362 | 2.698696274 | 0.03309457 | 0.339814312 | -3.579428118 |
| **Q969S8-1** | HDAC10 | 0.191000388 | 3.149546681 | 2.471553202 | 0.045473902 | 0.357414656 | -3.580268126 |

| | | | | | | | |
|---|---|---|---|---|---|---|---|
| **Q9NPL8** | TIMMDC1 | 0.242831589 | 3.181861288 | 3.037321309 | 0.020828076 | 0.306942846 | -3.582813897 |
| **Q9UID3-1** | VPS51 | -0.062102998 | 3.196644082 | -3.425155995 | 0.012488504 | 0.302793718 | -3.587678435 |
| **Q6P1K8** | GTF2H2D; GTF2H2 | -0.097847659 | 3.236605676 | -3.130337075 | 0.018387844 | 0.305857884 | -3.594095233 |
| **Q3SXM5** | HSDL1 | 0.088379924 | 3.155465783 | 3.05524035 | 0.02033217 | 0.305857884 | -3.595080092 |
| **Q9Y333** | LSM 2.00 | -0.06545035 | 3.19671609 | -3.457436183 | 0.011979777 | 0.299378139 | -3.597363573 |
| **Q14865** | ARID5B | 0.147049609 | 3.170521107 | 2.929220163 | 0.024108837 | 0.322917088 | -3.597537073 |
| **Q96HP4** | OXNAD1 | 0.180019738 | 3.179566801 | 2.849684508 | 0.02687397 | 0.328952544 | -3.600294346 |
| **Q8NFF5** | FLAD1 | 0.070637314 | 3.191718754 | 3.123382321 | 0.018559205 | 0.305857884 | -3.601611976 |
| **Q53R41** | FASTKD1 | 0.172809266 | 3.204807129 | 3.068103231 | 0.019984023 | 0.305857884 | -3.602121697 |
| **Q5T3I0-3** | GPATCH4 | -0.084868765 | 3.195256448 | -3.056575405 | 0.020295734 | 0.305857884 | -3.603368652 |
| **Q9Y676** | MRPS18B | 0.199830806 | 3.186249955 | 2.947870102 | 0.023505482 | 0.320063886 | -3.605690529 |
| **Q5TC12-1** | ATPAF1 | 0.144065347 | 3.185568727 | 2.874704076 | 0.025969358 | 0.328952544 | -3.605813053 |
| **Q8WWZ3-1** | EDARADD | -0.295090909 | 3.200209448 | -2.450081775 | 0.046871958 | 0.359457245 | -3.60974128 |
| **Q96AG3** | SLC25A46 | 0.098445496 | 3.187050869 | 3.131866355 | 0.018350393 | 0.305857884 | -3.61497697 |
| **Q9Y2R9** | MRPS7 | 0.229252479 | 3.178996706 | 3.144471418 | 0.018044815 | 0.305857884 | -3.616122547 |
| **P42765** | ACAA2 | 0.179134476 | 3.175045046 | 3.500185365 | 0.011340528 | 0.299378139 | -3.617522777 |
| **Q8NEZ2** | VPS37A | 0.075055282 | 3.149753814 | 3.140678017 | 0.018136195 | 0.305857884 | -3.619975141 |
| **Q13114-1** | TRAF3 | -0.091347994 | 3.191192822 | -2.936437161 | 0.023873414 | 0.322027627 | -3.621522972 |
| **Q5JTW2-5** | CEP78 | -0.063928029 | 3.179768626 | -3.203120589 | 0.01669348 | 0.305857884 | -3.622720575 |
| **Q9Y5N6** | ORC6 | -0.105637845 | 3.196331812 | -3.125085357 | 0.018517085 | 0.305857884 | -3.623676448 |
| **Q5TC82-1** | RC3H1 | 0.075212125 | 3.192473369 | 3.137520223 | 0.018212644 | 0.305857884 | -3.625285465 |
| **Q9BXW7** | CECR5; HDHD5 | 0.152909814 | 3.18354331 | 3.048745545 | 0.020510431 | 0.305857884 | -3.627894707 |
| **O14933** | UBE2L6 | 0.32627618 | 3.118721979 | 2.781194984 | 0.029526174 | 0.334311646 | -3.628900426 |
| **Q9BTT4** | MED10 | -0.123395108 | 3.195404851 | -3.152804459 | 0.017845818 | 0.305857884 | -3.635395399 |
| **Q68CL5-5** | TPGS2 | -0.281306088 | 3.191309442 | -2.402977166 | 0.050097808 | 0.36851891 | -3.637328646 |
| **P98177-1** | FOXO4 | -0.25408894 | 3.242955773 | -2.409698349 | 0.04962378 | 0.367474614 | -3.639126945 |
| **Q86US8-1** | SMG6 | -0.067783887 | 3.193797036 | -3.412919924 | 0.012687453 | 0.302793718 | -3.641640239 |
| **Q9C0H2-4** | TTYH3 | -0.185853833 | 3.201120711 | -2.501796556 | 0.04357795 | 0.357204257 | -3.641909594 |
| **P09564** | CD7 | -0.157855234 | 3.208087643 | -2.943770186 | 0.023636722 | 0.320063886 | -3.642632315 |
| **Q86TI0** | TBC1D1 | -0.045653793 | 3.173341933 | -3.417201159 | 0.012617454 | 0.302793718 | -3.642737329 |
| **O96008** | TOMM40 | 0.228226132 | 3.169326371 | 3.371558303 | 0.013385918 | 0.305857884 | -3.643628356 |
| **Q9BXL8** | CDCA4 | -0.371027132 | 3.153207776 | -2.547256646 | 0.040881854 | 0.355540374 | -3.647193266 |
| **O60573** | EIF4E2 | 0.175005001 | 3.159995159 | 2.899688018 | 0.02509836 | 0.32636083 | -3.651827878 |
| **Q68CZ1** | RPGRIP1L | -0.111951697 | 3.217794349 | -2.864534421 | 0.026333077 | 0.328952544 | -3.652347363 |
| **Q9BRT6** | LLPH | -0.208054103 | 3.174113765 | -2.822800773 | 0.027883504 | 0.32997678 | -3.656894145 |
| **Q6ZN18-1** | AEBP2 | -0.070716168 | 3.203030568 | -3.056631639 | 0.020294201 | 0.305857884 | -3.661644242 |
| **P12694** | BCKDHA | 0.250046323 | 3.141373532 | 3.154602537 | 0.01780319 | 0.305857884 | -3.662111802 |
| **Q9GZT3-1** | SLIRP | 0.246404561 | 3.187671167 | 3.317122069 | 0.014369625 | 0.305857884 | -3.664227582 |
| **P49069** | CAMLG | -0.128981366 | 3.16731008 | -2.620038025 | 0.036923187 | 0.348943902 | -3.666670221 |

| | | | | | | | |
|---|---|---|---|---|---|---|---|
| **Q6PML9** | SLC30A9 | 0.184563154 | 3.187829782 | 2.832339477 | 0.027520765 | 0.32997678 | -3.666925838 |
| **P60903** | S100A10 | -0.155352286 | 3.190423441 | -2.708695741 | 0.03263874 | 0.339814312 | -3.670878722 |
| **P18583-3** | SON | 0.160862603 | 3.152814334 | 2.561508678 | 0.040073003 | 0.355476279 | -3.671171635 |
| **P23378** | GLDC | 0.108083872 | 3.180537505 | 3.183724296 | 0.017127883 | 0.305857884 | -3.674246502 |
| **P23368-1** | ME2 | 0.197617684 | 3.184047888 | 3.488176748 | 0.011516255 | 0.299378139 | -3.674791979 |
| **Q86Y79** | PTRH1 | 0.109016523 | 3.165104735 | 2.84317225 | 0.027114894 | 0.328952544 | -3.681782115 |
| **Q9H0R6-1** | QRSL1 | 0.198460283 | 3.171426239 | 3.174327014 | 0.017342724 | 0.305857884 | -3.684500679 |
| **Q8N488** | RYBP | -0.092389997 | 3.2016809 | -2.990777254 | 0.022177867 | 0.312352362 | -3.684707798 |
| **P06702** | S100A9 | -1.000717012 | -0.414926557 | -2.518949565 | 0.04253946 | 0.355540374 | -3.685076036 |
| **Q53GL0-1** | PLEKHO1 | -0.067061856 | 3.224050277 | -3.103545844 | 0.01905749 | 0.305857884 | -3.686214355 |
| **Q96FX2-1** | DPH3 | -0.321743487 | 3.259820205 | -2.615085847 | 0.037179337 | 0.349573768 | -3.6876672 |
| **Q9Y2K6** | USP20 | -0.09982333 | 3.244040701 | -3.30461093 | 0.014606635 | 0.305857884 | -3.690217388 |
| **Q6IQ26** | DENND5A | 0.144307014 | 3.178153531 | 2.5369983 | 0.041474617 | 0.355540374 | -3.694982471 |
| **Q8IX90** | SKA3 | -0.083896927 | 3.172875527 | -3.212934947 | 0.016478227 | 0.305857884 | -3.69587819 |
| **Q8IZ69** | TRMT2A | -0.046449339 | 3.192207659 | -3.362699662 | 0.013540885 | 0.305857884 | -3.7039977 |
| **Q9H204** | MED28 | -0.113164306 | 3.21357022 | -2.735335811 | 0.031456443 | 0.33673269 | -3.705565605 |
| **Q9NP97** | DYNLRB1 | -0.210641351 | 3.208271297 | -3.30098876 | 0.014676043 | 0.305857884 | -3.706200083 |
| **Q15649** | ZNHIT3 | -0.125106769 | 3.236033818 | -2.660181809 | 0.034914063 | 0.345787303 | -3.707680621 |
| **Q9Y234** | LIPT1 | 0.230583921 | 3.174743594 | 2.384612238 | 0.051417108 | 0.371709669 | -3.708464041 |
| **O60830-2** | TIMM17B | 0.18437624 | 3.206198193 | 2.599922634 | 0.037975331 | 0.352231442 | -3.711263072 |
| **Q8NCU4** | KIAA1407; CCDC1 | 0.321776696 | 3.124598551 | 2.363534986 | 0.052975668 | 0.375126274 | -3.714160475 |
| **P29218-3** | IMPA1 | 0.040044436 | 3.179196838 | 3.471925409 | 0.011758814 | 0.299378139 | -3.715371667 |
| **Q86VP3-2** | PACS2 | -0.177324103 | 3.221573147 | -2.4415445 | 0.047440197 | 0.361027168 | -3.716984869 |
| **P62750** | RPL23A | -0.099178145 | 3.195732672 | -3.761650082 | 0.008157896 | 0.282982452 | -3.717702098 |
| **Q6KCM7-3** | SLC25A25 | 0.251227911 | 3.240112559 | 2.469819766 | 0.045585142 | 0.357414656 | -3.720339761 |
| **Q4L235-1** | AASDH | 0.675426634 | 3.077194149 | 2.31094556 | 0.057080699 | 0.377614679 | -3.721813518 |
| **Q0ZGT2** | NEXN | -0.110619186 | 3.201523129 | -2.971302824 | 0.022770203 | 0.316828828 | -3.722259569 |
| **Q96G46** | DUS3L | 0.058999045 | 3.172122603 | 3.469351928 | 0.011797733 | 0.299378139 | -3.724423403 |
| **Q5JTZ9** | AARS2 | 0.164028342 | 3.195527743 | 3.200001404 | 0.016762527 | 0.305857884 | -3.725512031 |
| **Q5T440** | IBA57 | 0.194378741 | 3.193984433 | 2.946053024 | 0.023563551 | 0.320063886 | -3.726962134 |
| **Q96D53-1** | ADCK4; COQ8B | 0.194574038 | 3.203262084 | 2.56042883 | 0.040133697 | 0.355476279 | -3.732079357 |
| **O75940** | SMNDC1 | 0.096624967 | 3.160104759 | 3.068724858 | 0.019967361 | 0.305857884 | -3.732633341 |
| **Q9BQ75** | CMSS1 | 0.070990027 | 3.146227654 | 3.200865555 | 0.016743367 | 0.305857884 | -3.73415902 |
| **Q03933-2** | HSF2 | -0.080133083 | 3.229100441 | -3.099076444 | 0.019171737 | 0.305857884 | -3.734281996 |
| **P61353** | RPL27 | -0.136090244 | 3.179679951 | -3.450464473 | 0.012087706 | 0.299378139 | -3.738170569 |
| **Q8TB03** | CXorf38 | -0.071906445 | 3.189732091 | -3.027014115 | 0.021119199 | 0.307875291 | -3.739929402 |
| **Q96QV1-1** | HHIP | 0.132235585 | 3.162718426 | 2.557862246 | 0.040278342 | 0.355540374 | -3.741530609 |
| **P20810-6** | CAST | -0.145693946 | 3.188366062 | -2.894205529 | 0.025286786 | 0.327738203 | -3.742424309 |
| **Q9NQ55-3** | PPAN-P2RY11; PP | -0.128541974 | 3.194348241 | -3.042416411 | 0.020685761 | 0.306620093 | -3.746862063 |

| | | | | | | | |
|---|---|---|---|---|---|---|---|
| **Q8TCZ2-5** | CD99L2 | -0.211756373 | 3.215121832 | -2.46829557 | 0.045683189 | 0.357414656 | -3.751039166 |
| **Q9H6E5** | TUT1 | -0.076531671 | 3.205332119 | -3.168140733 | 0.017485743 | 0.305857884 | -3.751374786 |
| **P0CG08** | GPR89A; GPR89B | 0.145599712 | 3.211373868 | 2.854700932 | 0.02668994 | 0.328952544 | -3.75279892 |
| **Q96A35** | MRPL24 | 0.145608479 | 3.201896391 | 3.16818743 | 0.017484658 | 0.305857884 | -3.753471804 |
| **Q9P227** | ARHGAP23 | -0.139881738 | 3.144570857 | -2.81861333 | 0.028044348 | 0.330092368 | -3.759782431 |
| **O15498** | YKT6 | -0.077192546 | 3.17959207 | -3.327770131 | 0.014171195 | 0.305857884 | -3.761233002 |
| **Q8NE00** | TMEM104 | 0.338949149 | 3.134254512 | 2.277037919 | 0.059899824 | 0.382023723 | -3.764334844 |
| **P62487** | POLR2G | -0.113772908 | 3.203269262 | -3.179349232 | 0.017227546 | 0.305857884 | -3.770473467 |
| **O75027-2** | ABCB7 | 0.206416622 | 3.206137497 | 3.002568034 | 0.02182728 | 0.310784487 | -3.770855507 |
| **Q9UIJ7-1** | AK3 | 0.2169701 | 3.182065452 | 3.190023127 | 0.016985488 | 0.305857884 | -3.771082592 |
| **P84103** | SRSF3 | -0.054077982 | 3.176243928 | -3.651947155 | 0.009355172 | 0.28710988 | -3.771090748 |
| **Q9HA38-1** | ZMAT3 | 0.193257281 | 3.245471744 | 2.308766711 | 0.057257683 | 0.377614679 | -3.771917235 |
| **Q9BVJ6-1** | UTP14A | -0.069865555 | 3.195700162 | -3.375956718 | 0.013309692 | 0.305857884 | -3.772503491 |
| **Q9GZN2-1** | TGIF2 | -0.178392434 | 3.277228627 | -2.380359703 | 0.051727699 | 0.372598883 | -3.776405427 |
| **Q96KF7** | SMIM8 | 0.237817251 | 3.200476945 | 2.518043994 | 0.042593634 | 0.355540374 | -3.779131543 |
| **P14406** | COX7A2 | 0.154259811 | 3.180422928 | 2.692059936 | 0.033400801 | 0.341970827 | -3.779996235 |
| **Q9BRT9** | GINS4 | -0.052777298 | 3.18197495 | -3.231303246 | 0.016083387 | 0.305857884 | -3.787415086 |
| **Q2M2Z5** | PLK1S1; KIZ | -0.55059296 | 2.88014488 | -2.400716259 | 0.050258316 | 0.368557155 | -3.78921891 |
| **P35790** | CHKA | -0.102789691 | 3.253652618 | -2.585525718 | 0.038747675 | 0.352231442 | -3.791287399 |
| **Q71UM5** | RPS27L | 0.114237478 | 3.178065444 | 2.930300464 | 0.024073439 | 0.322917088 | -3.791638948 |
| **P62263** | RPS14 | -0.054150808 | 3.201146631 | -3.326913458 | 0.014187048 | 0.305857884 | -3.792675565 |
| **O75410-1** | TACC1 | -0.048561203 | 3.208270835 | -3.452306562 | 0.012059086 | 0.299378139 | -3.793902708 |
| **Q9UIG0** | BAZ1B | -0.09102158 | 3.202383603 | -3.648510755 | 0.009395666 | 0.28710988 | -3.79466135 |
| **Q8N5G0-2** | C4orf52; SMIM20 | 0.221299048 | 3.199146382 | 2.725846765 | 0.03187229 | 0.33673269 | -3.796932533 |
| **P29374-1** | ARID4A | 0.054228025 | 3.161086498 | 3.353683796 | 0.013700608 | 0.305857884 | -3.801064219 |
| **Q8TCY9** | URGCP | -0.084335823 | 3.19695473 | -3.05571527 | 0.0203192 | 0.305857884 | -3.802676617 |
| **Q5SVS4** | SLC25A30 | 0.224060736 | 3.191440517 | 2.396539695 | 0.050556226 | 0.368977419 | -3.804842475 |
| **A5YKK6** | CNOT1 | 0.114198314 | 3.194359716 | 2.592575555 | 0.038367436 | 0.352231442 | -3.805432193 |
| **Q9H7X0** | NAA60 | 0.18587147 | 3.23589498 | 2.353332442 | 0.053747554 | 0.376021618 | -3.807107302 |
| **O96000** | NDUFB10 | 0.149957829 | 3.177883627 | 3.257070982 | 0.01554665 | 0.305857884 | -3.808685401 |
| **Q96KC2** | ARL5B | -0.088277724 | 3.20529721 | -2.793373835 | 0.029034946 | 0.333102237 | -3.811078477 |
| **Q9BXC1** | GPR174 | 0.119233134 | 3.187931467 | 2.501493032 | 0.043596563 | 0.357204257 | -3.82317536 |
| **Q9NYA4** | MTMR4 | -0.076677098 | 3.251458456 | -2.839717219 | 0.027243647 | 0.328952544 | -3.824102183 |
| **Q8TAE8** | GADD45GIP1 | 0.157279505 | 3.191922459 | 2.856702279 | 0.026616894 | 0.328952544 | -3.826240883 |
| **Q8N9Q2** | SREK1IP1 | -0.218645797 | 3.211038843 | -2.312073781 | 0.056989277 | 0.377614679 | -3.827872691 |
| **Q9BQI6** | ANKRD32; SLF1 | -0.132603834 | 3.212975754 | -2.452199288 | 0.046732115 | 0.359078004 | -3.830162798 |
| **Q9NUL7** | DDX28 | 0.217775453 | 3.18905977 | 2.868789257 | 0.026180246 | 0.328952544 | -3.830396802 |
| **P11498** | PC | 0.218763357 | 3.19445612 | 3.225062649 | 0.016216373 | 0.305857884 | -3.83205766 |
| **Q3KRA6** | C2orf76 | 0.130192747 | 3.171806726 | 2.736852343 | 0.031390517 | 0.33673269 | -3.832840966 |

| | | | | | | | |
|---|---|---|---|---|---|---|---|
| **Q8WV74** | NUDT8 | 0.147752473 | 3.168940363 | 2.475132956 | 0.045245071 | 0.357414656 | -3.833900413 |
| **P62699** | YPEL5 | 0.225320067 | 3.136050014 | 2.342402645 | 0.054587353 | 0.376021618 | -3.836008676 |
| **P21953** | BCKDHB | 0.158978652 | 3.148759381 | 2.784151001 | 0.029406139 | 0.333759678 | -3.837997581 |
| **Q9BWG6-1** | SCNM1; TNFAIP8L: | -0.085165378 | 3.221722811 | -2.907924572 | 0.024818091 | 0.324308771 | -3.840195658 |
| **Q9UPR0-1** | PLCL2 | -0.079901806 | 3.186585806 | -3.011858149 | 0.021555236 | 0.309585734 | -3.843753436 |
| **Q86SF2** | GALNT7 | -0.059439838 | 3.160993912 | -3.225674692 | 0.016203278 | 0.305857884 | -3.845897052 |
| **P32969** | RPL9 | -0.081150467 | 3.201031584 | -3.598023572 | 0.010013286 | 0.293973515 | -3.846913244 |
| **Q96AP0** | ACD | -0.126985264 | 3.199698755 | -2.751465891 | 0.030762688 | 0.33673269 | -3.847958773 |
| **O95905-3** | ECD | -0.097996442 | 3.225252948 | -3.160734692 | 0.017658638 | 0.305857884 | -3.848361576 |
| **Q07817-1** | BCL2L1 | -0.141967892 | 3.170271694 | -2.442449714 | 0.047379608 | 0.360911779 | -3.852238261 |
| **Q15050** | RRS1 | -0.075931059 | 3.211256692 | -3.113926469 | 0.018794959 | 0.305857884 | -3.853781791 |
| **Q8ND83-1** | SLAIN1 | -0.200033354 | 3.193588277 | -2.809430687 | 0.028400527 | 0.330820062 | -3.855864461 |
| **P82650** | MRPS22 | 0.17741495 | 3.196522522 | 3.069700831 | 0.019941232 | 0.305857884 | -3.861003104 |
| **Q9C086** | INO80B | -0.221071026 | 3.21340189 | -2.598887372 | 0.038030326 | 0.352231442 | -3.862776459 |
| **P30304** | CDC25A | -0.142262768 | 3.238286503 | -2.381997315 | 0.051607864 | 0.372073031 | -3.866877703 |
| **Q96CM8-2** | ACSF2 | 0.21613181 | 3.195474874 | 3.099919296 | 0.019150136 | 0.305857884 | -3.872595064 |
| **Q9BRZ2-1** | TRIM56 | -0.059224101 | 3.187327476 | -3.144685601 | 0.01803967 | 0.305857884 | -3.87468736 |
| **Q12834** | CDC20 | -0.092453776 | 3.201533876 | -3.032516357 | 0.02096325 | 0.307351282 | -3.874862775 |
| **P62195-1** | PSMC5 | -0.038743935 | 3.168662108 | -3.54851492 | 0.010662337 | 0.299378139 | -3.87487126 |
| **Q9UMS0** | NFU1 | 0.117074932 | 3.19464484 | 2.87792642 | 0.02585523 | 0.328952544 | -3.874875681 |
| **P45880-2** | VDAC2 | 0.183679943 | 3.180588774 | 3.39446664 | 0.012994044 | 0.304707969 | -3.876462495 |
| **O94760-1** | DDAH1 | 0.063313635 | 3.186709141 | 2.959895536 | 0.02312502 | 0.318891564 | -3.876601763 |
| **Q9H845** | ACAD9 | 0.173734122 | 3.199408179 | 3.264164311 | 0.015402325 | 0.305857884 | -3.876687221 |
| **Q8N587** | ZNF561 | -0.185579234 | 3.216903473 | -2.21961886 | 0.065003546 | 0.393401641 | -3.876756166 |
| **Q01081** | U2AF1; LOC102724 | 0.057998933 | 3.194763251 | 3.390181765 | 0.013066382 | 0.304707969 | -3.880082599 |
| **P62851** | RPS25 | -0.158278792 | 3.191205301 | -3.011951111 | 0.021552532 | 0.309585734 | -3.881606325 |
| **Q8IYR2** | SMYD4 | 0.068099127 | 3.169226939 | 2.966858718 | 0.022907736 | 0.317241514 | -3.883518554 |
| **Q8N3I7-1** | BBS5 | -0.139550102 | 3.167787444 | -2.339795147 | 0.054789695 | 0.376021618 | -3.884864424 |
| **Q15819** | UBE2V2 | -0.066535076 | 3.173453629 | -3.241721062 | 0.015864008 | 0.305857884 | -3.885299175 |
| **O15014** | ZNF609 | 0.055351767 | 3.163892203 | 3.181881856 | 0.017169777 | 0.305857884 | -3.885370455 |
| **Q96TA2-2** | YME1L1 | 0.184289396 | 3.198726432 | 3.219990146 | 0.016325344 | 0.305857884 | -3.885614734 |
| **P48634-1** | PRRC2A | -0.03488499 | 3.18892217 | -3.502006533 | 0.011314135 | 0.299378139 | -3.885808518 |
| **P80404** | ABAT | 0.221117104 | 3.169928344 | 2.622948483 | 0.036773511 | 0.348943902 | -3.886112121 |
| **Q9H5Q4** | TFB2M | 0.187529758 | 3.183830332 | 3.117002643 | 0.018717912 | 0.305857884 | -3.887178305 |
| **Q02543** | RPL18A | -0.208657863 | 3.173171435 | -3.529603486 | 0.010922288 | 0.299378139 | -3.890668104 |
| **O60925** | PFDN1 | -0.11182677 | 3.177531422 | -2.920441386 | 0.024398556 | 0.323077552 | -3.893880072 |
| **Q7Z7L8** | C11orf96 | -0.250470511 | 3.102709526 | -2.142521449 | 0.072562267 | 0.404366176 | -3.895814396 |
| **Q6DKK2** | TTC19 | 0.100418797 | 3.166759679 | 2.834259708 | 0.027448351 | 0.32997678 | -3.899529354 |
| **Q9ULI2-1** | RIMKLB | 0.15735661 | 3.144797537 | 2.486355332 | 0.044535439 | 0.357204257 | -3.903506315 |

| | | | | | | | |
|---|---|---|---|---|---|---|---|
| **Q330K2-1** | NDUFAF6 | 0.2134082 | 3.221768482 | 2.660155039 | 0.034915363 | 0.345787303 | -3.90783079 |
| **Q9Y697** | NFS1 | 0.162413281 | 3.184813614 | 3.132795564 | 0.018327677 | 0.305857884 | -3.909258786 |
| **O43324-1** | EEF1E1 | -0.050041573 | 3.188832181 | -3.227713123 | 0.016159747 | 0.305857884 | -3.910337807 |
| **Q15070-1** | OXA1L | 0.189798683 | 3.219608516 | 2.820702226 | 0.027963989 | 0.330092368 | -3.912714194 |
| **Q9NP81-1** | SARS2 | 0.217444424 | 3.164822853 | 3.018835971 | 0.021353296 | 0.309578361 | -3.91569552 |
| **Q13740-1** | ALCAM | 0.075292314 | 3.162609746 | 2.896300591 | 0.025214604 | 0.327336648 | -3.920397729 |
| **Q9NQ92** | COPRS | -0.332785088 | 3.150603262 | -2.29892109 | 0.058064489 | 0.379089796 | -3.921461741 |
| **Q9UBT7** | CTNNAL1 | -0.138217374 | 3.236918315 | -2.51942589 | 0.042510994 | 0.355540374 | -3.923657018 |
| **Q9NQX7** | ITM2C | 0.099698363 | 3.137574795 | 2.594743803 | 0.038251279 | 0.352231442 | -3.925424128 |
| **P62899** | RPL31 | -0.182551347 | 3.204179965 | -3.239393702 | 0.015912735 | 0.305857884 | -3.925529954 |
| **P43155** | CRAT | 0.284746315 | 3.148584899 | 2.134855087 | 0.073361098 | 0.405447658 | -3.925578337 |
| **Q9Y6N1** | COX11 | 0.198650057 | 3.185316434 | 2.64605108 | 0.035607822 | 0.347548094 | -3.929586386 |
| **P84090** | ERH | -0.098037604 | 3.177384391 | -3.413278564 | 0.012681573 | 0.302793718 | -3.929900487 |
| **Q7L0Q8-1** | RHOU | -0.078302156 | 3.148189671 | -2.732491913 | 0.031580468 | 0.33673269 | -3.931219904 |
| **Q9Y448** | C15orf23; KNSTRN | -0.089133782 | 3.201458817 | -2.802910559 | 0.028656348 | 0.331404204 | -3.931706154 |
| **A0PJW6** | TMEM223 | 0.183201855 | 3.184962375 | 2.185083477 | 0.068285088 | 0.398319894 | -3.933002415 |
| **Q9UHQ9** | CYB5R1 | 0.079217655 | 3.195806129 | 3.017497125 | 0.021391884 | 0.309578361 | -3.933274994 |
| **Q99417** | MYCBP | -0.235476718 | 3.144533222 | -2.922031805 | 0.024345794 | 0.322917088 | -3.934831655 |
| **Q9NP92** | MRPS30 | 0.224747312 | 3.200390473 | 3.04917678 | 0.020498543 | 0.305857884 | -3.935735244 |
| **Q9H5L6** | THAP9 | 0.227022474 | 3.20911061 | 2.110201721 | 0.075990808 | 0.411304347 | -3.937624044 |
| **Q7Z4V0-1** | ZNF438 | -0.128009454 | 3.261990584 | -2.465379835 | 0.045871362 | 0.357577769 | -3.937771918 |
| **Q7RTV3-1** | ZNF367 | -0.168108963 | 3.250764891 | -2.34535164 | 0.05435944 | 0.376021618 | -3.940603208 |
| **Q9UDX5** | MTFP1 | 0.293205425 | 3.168616054 | 2.429616222 | 0.048246155 | 0.363332416 | -3.940707675 |
| **Q9Y287** | ITM2B | 0.136681862 | 3.102641591 | 2.672674115 | 0.034312606 | 0.345097821 | -3.943501215 |
| **Q66K64** | DCAF15 | -0.116794344 | 3.197525139 | -2.660165964 | 0.034914833 | 0.345787303 | -3.943505564 |
| **Q9HCE1** | MOV10 | -0.086351763 | 3.191477184 | -3.332310613 | 0.014087488 | 0.305857884 | -3.943848428 |
| **Q7L5Y9-1** | MAEA | -0.169269236 | 3.24365822 | -3.152136839 | 0.017861674 | 0.305857884 | -3.943880106 |
| **Q15544** | TAF11 | -0.106288392 | 3.188228143 | -2.736314201 | 0.031413894 | 0.33673269 | -3.944674092 |
| **Q16740** | CLPP | 0.178867249 | 3.196673612 | 2.903358652 | 0.024973042 | 0.325797735 | -3.946934774 |
| **Q9H9J2** | MRPL44 | 0.178766506 | 3.195522004 | 2.950167904 | 0.023432269 | 0.320063886 | -3.949113691 |
| **O43676** | NDUFB3 | 0.19917717 | 3.210417641 | 2.943774754 | 0.023636575 | 0.320063886 | -3.950341297 |
| **Q9NUJ1** | ABHD10 | 0.201347437 | 3.19422313 | 3.113132279 | 0.018814906 | 0.305857884 | -3.950343366 |
| **O95169** | NDUFB8 | 0.216898855 | 3.176794037 | 2.570792034 | 0.039555172 | 0.35510265 | -3.951969506 |
| **A6NCE7** | MAP1LC3B2 | -0.167155603 | 3.208878354 | -2.720972972 | 0.032088136 | 0.338117029 | -3.95225993 |
| **Q9Y371-2** | SH3GLB1 | -0.06830999 | 3.186280172 | -2.84134387 | 0.027182948 | 0.328952544 | -3.953535622 |
| **Q15329** | E2F5 | -0.245404061 | 3.223307818 | -2.089909545 | 0.078226644 | 0.415568143 | -3.95397357 |
| **Q5TF39** | KIAA1919; MFSD4E | -0.220651674 | 3.208062702 | -2.164667443 | 0.07030384 | 0.401998301 | -3.959183848 |
| **B0YJ81** | PTPLA; HACD1 | -0.119837792 | 3.240256685 | -2.4136066 | 0.049350282 | 0.366780161 | -3.960210774 |
| **Q9NP80-1** | PNPLA8 | 0.170399948 | 3.202130024 | 2.436881877 | 0.047753558 | 0.362024828 | -3.960576719 |

| | | | | | | | |
|---|---|---|---|---|---|---|---|
| **P12429** | ANXA3 | 0.070627594 | 3.199866489 | 3.065413472 | 0.02005629 | 0.305857884 | -3.962429038 |
| **Q96BQ5** | CCDC127 | 0.199157627 | 3.131908686 | 2.281282868 | 0.059539202 | 0.381175634 | -3.962535736 |
| **P49748-3** | ACADVL | 0.206626765 | 3.198609002 | 3.211959072 | 0.016499495 | 0.305857884 | -3.96444053 |
| **Q9H649** | NSUN3 | 0.243232554 | 3.186686578 | 2.465648951 | 0.04585396 | 0.357577769 | -3.969448878 |
| **Q8NDD1** | C1orf131 | -0.102829716 | 3.219544924 | -2.727952908 | 0.031779491 | 0.33673269 | -3.971122541 |
| **O75935** | DCTN3 | -0.056380897 | 3.154825371 | -3.175185354 | 0.01732298 | 0.305857884 | -3.972169123 |
| **P61601** | NCALD | 0.274388507 | 3.242128088 | 2.057569351 | 0.081927781 | 0.423947857 | -3.973206604 |
| **P53007** | SLC25A1 | 0.241709492 | 3.190539736 | 3.007000759 | 0.021697019 | 0.310090823 | -3.97938806 |
| **Q9Y3A0-1** | COQ4 | 0.362480605 | 3.218206836 | 2.060938247 | 0.081534153 | 0.423686898 | -3.980244395 |
| **Q9BRP1** | PDCD2L | -0.096352728 | 3.233914299 | -2.74159717 | 0.031185194 | 0.33673269 | -3.984650817 |
| **O94817-1** | ATG12 | -0.112361546 | 3.196499806 | -2.875453728 | 0.025942759 | 0.328952544 | -3.984933694 |
| **Q9NRY5** | FAM114A2 | -0.052512903 | 3.179449316 | -3.20852719 | 0.016574526 | 0.305857884 | -3.985095872 |
| **Q8N4L2** | TMEM55A; PIP4P2 | -0.196108233 | 3.163596358 | -2.136255532 | 0.07321451 | 0.405447658 | -3.987660555 |
| **Q08117-2** | AES | 0.114984984 | 3.198077223 | 2.700253124 | 0.033023161 | 0.339814312 | -3.988056985 |
| **P63104-1** | YWHAZ | 0.060971019 | 3.16535737 | 3.657872265 | 0.009285799 | 0.28710988 | -3.988221474 |
| **O95900** | TRUB2 | 0.157952531 | 3.183133372 | 2.659469874 | 0.034948673 | 0.345787303 | -3.990584728 |
| **P46977** | STT3A | -0.050644945 | 3.207519939 | -3.090270012 | 0.019399012 | 0.305857884 | -3.990610294 |
| **Q86YH6-1** | PDSS2 | 0.194862943 | 3.180703595 | 2.652273517 | 0.035300554 | 0.346250492 | -3.994693002 |
| **Q00325-1** | SLC25A3 | 0.211593042 | 3.187707953 | 3.310360534 | 0.014497192 | 0.305857884 | -3.994899507 |
| **P09661** | SNRPA1 | 0.06659021 | 3.150073289 | 2.987116607 | 0.022287936 | 0.312858048 | -3.998045853 |
| **Q9Y2W6** | TDRKH | 0.160756364 | 3.189690549 | 2.809843767 | 0.028384401 | 0.330820062 | -4.000259399 |
| **Q8TEW0** | PARD3 | 0.183185467 | 3.11289763 | 2.263022975 | 0.061106426 | 0.38369455 | -4.001847225 |
| **Q8NCW5** | APOA1BP; NAXE | 0.074757411 | 3.177797816 | 3.242390098 | 0.015850031 | 0.305857884 | -4.002919234 |
| **Q16798** | ME3 | 0.184684575 | 3.21218697 | 2.27092949 | 0.060422691 | 0.383145812 | -4.009356829 |
| **Q16629** | SRSF7 | -0.120792515 | 3.177014948 | -2.954412169 | 0.023297679 | 0.319689228 | -4.012319005 |
| **P46199** | MTIF2 | 0.179397278 | 3.17869018 | 2.799764628 | 0.028780656 | 0.331452005 | -4.014228525 |
| **Q14005-1** | IL16 | 0.074790013 | 3.176768018 | 2.943441954 | 0.023647263 | 0.320063886 | -4.014958152 |
| **Q9H3K2** | GHITM | 0.186143254 | 3.147995041 | 2.710683985 | 0.032548898 | 0.339814312 | -4.017541895 |
| **Q52LW3** | ARHGAP29 | -0.096923222 | 3.214102786 | -2.91150382 | 0.024697342 | 0.32394477 | -4.018281801 |
| **Q9UGI0** | ZRANB1 | -0.327513396 | 3.031086142 | -2.04929959 | 0.082902148 | 0.425319102 | -4.018731231 |
| **Q69YU5** | C12orf73 | 0.189081725 | 3.214476641 | 2.397056278 | 0.05051928 | 0.368977419 | -4.018788775 |
| **Q5T5P2-1** | KIAA1217 | 0.163202061 | 3.153937507 | 2.115935466 | 0.075370821 | 0.409678928 | -4.019392663 |
| **Q7Z3D4** | LYSMD3 | -0.112875845 | 3.208915196 | -2.59111814 | 0.038445721 | 0.352231442 | -4.02687539 |
| **Q9BVX2** | TMEM106C | -0.183472342 | 3.221153261 | -2.244281714 | 0.062758961 | 0.387695915 | -4.027782857 |
| **Q5TA31** | RNF187 | -0.166699556 | 3.212016893 | -2.139587281 | 0.07286696 | 0.404796855 | -4.02805174 |
| **O95236-1** | APOL3 | -0.08290042 | 3.171888839 | -2.667317288 | 0.034569167 | 0.345474259 | -4.02868408 |
| **O15533-3** | TAPBP | -0.133619164 | 3.157327037 | -2.761470167 | 0.030340563 | 0.336035644 | -4.030869003 |
| **Q9Y576** | ASB1 | -0.112153639 | 3.231875365 | -2.402370339 | 0.050140836 | 0.36851891 | -4.032368645 |
| **Q6ICC9** | LDOC1L; RTL6 | -0.147476676 | 3.232688426 | -2.22353189 | 0.064642018 | 0.392645894 | -4.033531488 |

| | | | | | | | |
|---|---|---|---|---|---|---|---|
| **P45954** | ACADSB | 0.215891914 | 3.176544673 | 3.075835767 | 0.019777823 | 0.305857884 | -4.033767772 |
| **Q14094** | CCNI | 0.231114084 | 3.081949526 | 2.152259139 | 0.071560298 | 0.403638325 | -4.034915443 |
| **P46087-4** | NOP2 | -0.095686391 | 3.182381624 | -3.303787491 | 0.014622382 | 0.305857884 | -4.036293487 |
| **Q13442** | PDAP1 | 0.092560138 | 3.144907658 | 3.247714837 | 0.01573926 | 0.305857884 | -4.037368754 |
| **Q9Y619** | SLC25A15 | 0.249221352 | 3.181549626 | 2.4033104 | 0.050074195 | 0.36851891 | -4.042066595 |
| **P62241** | RPS8 | -0.144373798 | 3.166671908 | -3.453025462 | 0.012047937 | 0.299378139 | -4.042220026 |
| **Q16527** | CSRP2 | -0.106828847 | 3.149295576 | -2.487120829 | 0.044487459 | 0.357204257 | -4.042478638 |
| **P40939** | HADHA | 0.219259909 | 3.176542161 | 3.33977275 | 0.013951082 | 0.305857884 | -4.04615667 |
| **Q96EH3** | MALSU1 | 0.199555194 | 3.234599488 | 2.290382979 | 0.058773609 | 0.37966232 | -4.049268286 |
| **P83111-1** | LACTB | 0.153210038 | 3.183759134 | 2.773630848 | 0.029835703 | 0.335282407 | -4.050422541 |
| **Q9C0D4** | ZNF518B | -0.180758785 | 3.20630859 | -2.252496241 | 0.062029074 | 0.385317429 | -4.050508543 |
| **P80217-2** | IFI35 | -0.095789309 | 3.193329709 | -2.827251374 | 0.027713627 | 0.32997678 | -4.051380249 |
| **Q9UIL8** | PHF11 | -0.159636069 | 3.247013001 | -2.48037157 | 0.044912359 | 0.357204257 | -4.05242993 |
| **Q9Y6H3** | XRCC6BP1; ATP23 | 0.198724704 | 3.198495538 | 2.365583257 | 0.052822088 | 0.374840509 | -4.054626302 |
| **P16152** | CBR1; SETD4 | -0.879296423 | 2.57177853 | -1.973275737 | 0.092421096 | 0.446659583 | -4.056302131 |
| **P51784** | USP11 | -0.037711588 | 3.196228926 | -3.278458345 | 0.015115881 | 0.305857884 | -4.057083163 |
| **Q9H857-2** | NT5DC2 | 0.176330692 | 3.199711334 | 2.706838365 | 0.032722905 | 0.339814312 | -4.057514874 |
| **Q14687** | GSE1 | -0.055598877 | 3.227141962 | -3.056046249 | 0.020310167 | 0.305857884 | -4.062067348 |
| **Q86SQ9-2** | DHDDS | 0.112667669 | 3.223681246 | 2.266415266 | 0.060812103 | 0.383378328 | -4.062101689 |
| **P28799** | GRN | 0.078689651 | 3.054287932 | 2.588990298 | 0.038560318 | 0.352231442 | -4.062259474 |
| **P55809** | OXCT1 | 0.175718838 | 3.178079783 | 3.246352557 | 0.01576752 | 0.305857884 | -4.062656877 |
| **Q96EY7-1** | PTCD3 | 0.183006898 | 3.196358261 | 3.071342216 | 0.019897371 | 0.305857884 | -4.06267222 |
| **Q9BSH4** | TACO1 | 0.184106024 | 3.175277149 | 2.834112648 | 0.027453889 | 0.32997678 | -4.064041507 |
| **Q8NDY6** | BHLHE23 | -0.175719393 | 3.144794276 | -2.211606472 | 0.065750271 | 0.394427892 | -4.064459922 |
| **P17096-3** | HMGA1 | -0.102507262 | 3.187760158 | -2.945366032 | 0.023585545 | 0.320063886 | -4.066483328 |
| **Q8IY95** | TMEM192 | -0.125560904 | 3.199566329 | -2.459029987 | 0.046283959 | 0.358190865 | -4.066667136 |
| **Q96HY7** | DHTKD1 | 0.236753819 | 3.17500541 | 2.764211643 | 0.030225963 | 0.336035644 | -4.06724362 |
| **P48047** | ATP5O | 0.131688085 | 3.183979312 | 3.017860845 | 0.021381394 | 0.309578361 | -4.070285716 |
| **P04004** | VTN | 0.104450682 | 3.277752257 | 2.649255204 | 0.03544925 | 0.346852575 | -4.07134856 |
| **P36508** | ZNF76 | 0.140070178 | 3.172544516 | 2.175280112 | 0.069246992 | 0.399872475 | -4.072273448 |
| **Q15031** | LARS2 | 0.180343452 | 3.180400389 | 3.010759265 | 0.021587224 | 0.309585734 | -4.076472162 |
| **P18124** | RPL7 | -0.156205955 | 3.17017696 | -3.466334725 | 0.011843542 | 0.299378139 | -4.078070921 |
| **Q9Y6D0** | SELK; SELENOK | -0.113907384 | 3.197937562 | -2.303533092 | 0.05768511 | 0.377966845 | -4.078341343 |
| **Q9GZU8** | FAM192A | 0.043492018 | 3.163988258 | 3.044417059 | 0.020630165 | 0.306620093 | -4.078581263 |
| **Q9BVV7** | TIMM21 | 0.245482483 | 3.19160658 | 2.549319175 | 0.040763748 | 0.355540374 | -4.079045758 |
| **Q9Y6N5** | SQRDL; SQOR | 0.192666343 | 3.190274565 | 3.081441325 | 0.019629774 | 0.305857884 | -4.079363328 |
| **Q6JQN1-5** | ACAD10 | 0.203496727 | 3.202065602 | 2.38275878 | 0.051552241 | 0.372073031 | -4.080650968 |
| **Q9UKV3-1** | ACIN1 | 0.191094792 | 3.108306097 | 2.29366187 | 0.058500239 | 0.379210276 | -4.080995166 |
| **Q9NWR8** | CCDC109B; MCUB | 0.275791988 | 3.18293771 | 2.562717222 | 0.040005189 | 0.355476279 | -4.081137982 |

| | | | | | | | |
|---|---|---|---|---|---|---|---|
| **Q9HBL7** | PLGRKT | 0.198462945 | 3.174378147 | 2.755807002 | 0.030578758 | 0.336493392 | -4.081523008 |
| **Q9UK39** | CCRN4L; NOCT | -0.100825687 | 3.237130047 | -2.579316244 | 0.039085864 | 0.352883173 | -4.086683168 |
| **Q9Y2E5** | MAN2B2 | 0.102840286 | 3.18818234 | 2.586759281 | 0.038680858 | 0.352231442 | -4.087032127 |
| **Q9UK41-1** | VPS28 | -0.05565274 | 3.189270965 | -2.767504696 | 0.030088912 | 0.336035644 | -4.087333094 |
| **Q9BPX3** | NCAPG | -0.072702807 | 3.175518141 | -3.320104195 | 0.014313749 | 0.305857884 | -4.0909305 |
| **Q9UBS3** | DNAJB9 | 0.169058846 | 3.170766018 | 2.336689222 | 0.055031724 | 0.376021618 | -4.092955391 |
| **Q02241-1** | KIF23 | -0.11422498 | 3.200198486 | -3.06047563 | 0.020189691 | 0.305857884 | -4.093408511 |
| **Q96EY8** | MMAB | 0.171987713 | 3.18202415 | 2.759716412 | 0.030414116 | 0.336078092 | -4.095573155 |
| **Q5T8I3-1** | FAM102B | 0.11724705 | 3.142805914 | 2.459030993 | 0.046283894 | 0.358190865 | -4.096797804 |
| **Q9HAV4** | XPO5 | -0.083912071 | 3.190730736 | -3.420722797 | 0.012560189 | 0.302793718 | -4.096931582 |
| **Q93073-1** | SECISBP2L | 0.139736969 | 3.147196626 | 2.235436019 | 0.06355476 | 0.389084368 | -4.09701249 |
| **Q9UBX3-2** | SLC25A10 | 0.237489397 | 3.228643822 | 2.866050148 | 0.026278524 | 0.328952544 | -4.099468378 |
| **Q70CQ3** | USP30 | 0.256912812 | 3.173515221 | 2.306726664 | 0.057423904 | 0.377614679 | -4.100527585 |
| **Q99496** | RNF2 | 0.074658736 | 3.182125111 | 2.855315105 | 0.026667501 | 0.328952544 | -4.101768029 |
| **A6NJ78** | METTL15 | 0.233154042 | 3.211188677 | 2.181213394 | 0.068663186 | 0.398897914 | -4.103162233 |
| **P22033** | MUT | 0.209530876 | 3.153982247 | 3.085120906 | 0.019533241 | 0.305857884 | -4.103341093 |
| **Q9BUR5-1** | APOO | 0.168204954 | 3.217556337 | 2.732828608 | 0.031565757 | 0.33673269 | -4.109258697 |
| **Q9NP72** | RAB18 | -0.061332431 | 3.177723292 | -2.874305356 | 0.025983516 | 0.328952544 | -4.109635431 |
| **O43426** | SYNJ1 | -0.06003245 | 3.227301241 | -3.048440341 | 0.020518849 | 0.305857884 | -4.110172758 |
| **O94782** | USP1 | -0.073310079 | 3.232235813 | -3.071364833 | 0.019896767 | 0.305857884 | -4.110396786 |
| **Q92567** | FAM168A | -0.149463722 | 3.200470338 | -2.309966899 | 0.057160124 | 0.377614679 | -4.111422777 |
| **O76021** | RSL1D1 | -0.160923993 | 3.19871206 | -3.278519529 | 0.015114667 | 0.305857884 | -4.113307319 |
| **O60942-1** | RNGTT | 0.107757598 | 3.170152808 | 3.066230409 | 0.020034312 | 0.305857884 | -4.11352291 |
| **P24941-1** | CDK2 | -0.053778956 | 3.189492856 | -3.166731839 | 0.017518492 | 0.305857884 | -4.121596898 |
| **Q96EU6** | RRP36 | -0.095046366 | 3.180141782 | -2.780725021 | 0.029545305 | 0.334311646 | -4.12462604 |
| **P49753-1** | ACOT2 | 0.196051363 | 3.186540206 | 2.593349192 | 0.038325949 | 0.352231442 | -4.125470131 |
| **Q8TCG2** | PI4K2B | -0.112253665 | 3.185547909 | -2.82629986 | 0.027749853 | 0.32997678 | -4.127071988 |
| **Q6ZT12-4** | UBR3 | -0.095574868 | 3.188418225 | -2.677127848 | 0.034100827 | 0.345097821 | -4.127779082 |
| **Q9BPX6-3** | MICU1 | 0.159989691 | 3.155391933 | 2.874761498 | 0.025967319 | 0.328952544 | -4.128041902 |
| **Q86Y91-4** | KIF18B | -0.061372931 | 3.169423251 | -2.812188031 | 0.028293072 | 0.330820062 | -4.132432283 |
| **P09429** | HMGB1 | 0.09447188 | 3.174743266 | 3.683297142 | 0.008994453 | 0.286465393 | -4.135943125 |
| **Q8N448** | LNX2 | -0.128502853 | 3.194045808 | -2.218091418 | 0.065145227 | 0.393496543 | -4.136064327 |
| **Q9HC36** | RNMTL1; MRM3 | 0.100989788 | 3.123913216 | 2.487869657 | 0.044440575 | 0.357204257 | -4.136377787 |
| **P53680** | AP2S1 | -0.083241252 | 3.19004903 | -2.740747759 | 0.031221846 | 0.33673269 | -4.139359158 |
| **P17275** | JUNB | -0.127571699 | 3.275612938 | -2.377426242 | 0.051943079 | 0.373408128 | -4.141416349 |
| **Q8IXM2-3** | C17orf49 | -0.082495384 | 3.167507284 | -2.661300063 | 0.034859773 | 0.345787303 | -4.14346872 |
| **Q9NRY2-1** | INIP | -0.223667256 | 3.146168169 | -2.258246373 | 0.061523331 | 0.384217614 | -4.144069258 |
| **Q9C0B9** | ZCCHC2 | -0.250786128 | 3.304771822 | -1.870269444 | 0.107072986 | 0.475555333 | -4.144844912 |
| **O75306-1** | NDUFS2 | 0.223847148 | 3.186966604 | 2.998394286 | 0.021950698 | 0.311022609 | -4.147527525 |

| | | | | | | | |
|---|---|---|---|---|---|---|---|
| **Q9H7H0-3** | METTL17 | 0.161446523 | 3.160752211 | 2.619208704 | 0.036965954 | 0.348943902 | -4.149992264 |
| **Q13283** | G3BP1 | 0.037073627 | 3.187339968 | 3.405401661 | 0.012811403 | 0.304707969 | -4.151253323 |
| **A8MPS7-1** | YDJC | 0.238819795 | 3.202609352 | 1.871163989 | 0.106936373 | 0.475471224 | -4.152009503 |
| **P50440-1** | GATM | 0.199646948 | 3.189811911 | 2.669646689 | 0.034457355 | 0.345097821 | -4.155110454 |
| **Q86UT6** | NLRX1 | 0.211536913 | 3.190265708 | 2.046921318 | 0.083184511 | 0.425585663 | -4.155721766 |
| **Q9BTE3-1** | MCMBP | -0.055682637 | 3.193028958 | -3.281958246 | 0.015046628 | 0.305857884 | -4.157296662 |
| **Q96AV8** | E2F7 | -0.109792325 | 3.217108295 | -2.455803437 | 0.046495091 | 0.358626496 | -4.157461009 |
| **Q96AQ8-1** | CCDC90A; MCUR1 | 0.170904093 | 3.192168183 | 2.807490983 | 0.028476378 | 0.330820062 | -4.161014026 |
| **Q53HL2** | CDCA8 | -0.089750652 | 3.180764376 | -2.756007505 | 0.030570291 | 0.336493392 | -4.161204599 |
| **Q6P3S1** | DENND1B | 0.081627204 | 3.206171459 | 2.646262057 | 0.035597358 | 0.347548094 | -4.161619982 |
| **Q5XUX0-1** | FBXO31 | -0.171440238 | 3.205543527 | -2.004269162 | 0.088415214 | 0.437773138 | -4.162215118 |
| **Q99684** | GFI1 | -0.139662212 | 3.198031703 | -2.154466587 | 0.071335118 | 0.403638325 | -4.164301375 |
| **P08574** | CYC1 | 0.206489441 | 3.191988556 | 2.910863363 | 0.024718902 | 0.32394477 | -4.164488767 |
| **Q9UEE9-1** | CFDP1 | -0.073512447 | 3.190033609 | -2.655650208 | 0.035134981 | 0.346250492 | -4.169504 |
| **Q9H816** | DCLRE1B | 0.080066514 | 3.194256918 | 2.489787483 | 0.044320736 | 0.357204257 | -4.169564647 |
| **Q9NVA1** | UQCC; UQCC1 | 0.144225162 | 3.175379196 | 2.365666595 | 0.052815849 | 0.374840509 | -4.171496064 |
| **Q96E14** | RMI2 | 0.075088801 | 3.177875588 | 2.478090949 | 0.04505689 | 0.357262465 | -4.175379629 |
| **Q13950-1** | RUNX2 | 0.12036617 | 3.226752984 | 2.231543575 | 0.063908203 | 0.390577439 | -4.17632924 |
| **P18583-5** | SON | 0.05732064 | 3.194756234 | 2.867190634 | 0.026237556 | 0.328952544 | -4.180328376 |
| **Q13011** | ECH1 | 0.120834791 | 3.181716667 | 2.940547913 | 0.023740418 | 0.320778271 | -4.182149835 |
| **Q6P1A2** | LPCAT3 | -0.098227004 | 3.207652388 | -2.434149298 | 0.047938206 | 0.362534913 | -4.186337092 |
| **Q86WA6** | BPHL | 0.191170036 | 3.211057048 | 2.676467081 | 0.03413216 | 0.345097821 | -4.186743439 |
| **Q9UQ35** | SRRM2 | -0.030136949 | 3.192953864 | -3.570347098 | 0.010370617 | 0.295122294 | -4.187264531 |
| **Q9NR96-4** | TLR9 | -0.18281147 | 3.287135518 | -2.014000292 | 0.087193569 | 0.435693653 | -4.188398592 |
| **Q9UBQ6** | EXTL2 | 0.112869746 | 3.163339669 | 2.350854237 | 0.053936792 | 0.376021618 | -4.18923608 |
| **Q96EX1** | C1orf212; SMIM12 | 0.224661745 | 3.13011294 | 2.344839903 | 0.054398919 | 0.376021618 | -4.191065886 |
| **Q8N442** | GUF1 | 0.22355251 | 3.173046893 | 2.59074765 | 0.038465648 | 0.352231442 | -4.191841794 |
| **Q12981** | BNIP1 | 0.108463042 | 3.202929186 | 2.761736629 | 0.030329404 | 0.336035644 | -4.192117602 |
| **Q9Y2D5-6** | PALM2-AKAP2; PAI | -0.054032413 | 3.17007567 | -3.209344399 | 0.016556626 | 0.305857884 | -4.194424247 |
| **Q96BX8** | MOB3A | -0.099475229 | 3.241452668 | -2.803970729 | 0.028614586 | 0.331403622 | -4.196524593 |
| **Q9UKV3-5** | ACIN1 | 0.124269274 | 3.153690218 | 2.138202699 | 0.073011189 | 0.404796855 | -4.197474661 |
| **Q9C0D2-3** | KIAA1731; CEP295 | -0.347498834 | 2.944076047 | -1.81412217 | 0.115998593 | 0.492508572 | -4.198911755 |
| **Q5BKZ1** | ZNF326 | 0.042772306 | 3.185427526 | 3.014686603 | 0.021473134 | 0.309585734 | -4.199115198 |
| **Q7L1V2** | MON1B | 0.109108028 | 3.174389512 | 2.499170376 | 0.043739268 | 0.357204257 | -4.200469721 |
| **Q9Y2Y1** | POLR3K | -0.169052006 | 3.239965027 | -2.258844164 | 0.061470997 | 0.384217614 | -4.201298294 |
| **Q8IVH8** | MAP4K3 | -0.093718115 | 3.174201514 | -2.307970093 | 0.057322532 | 0.377614679 | -4.202499185 |
| **Q8NBL1** | POGLUT1 | 0.115226994 | 3.227148697 | 2.735157277 | 0.031464214 | 0.33673269 | -4.205523091 |
| **Q8NFV4** | ABHD11 | 0.145801405 | 3.192665872 | 2.269179152 | 0.060573377 | 0.383378328 | -4.205669543 |
| **Q8TEY7-1** | USP33 | -0.102504571 | 3.207906993 | -2.282446624 | 0.059440727 | 0.381159465 | -4.20801414 |

| | | | | | | | |
|---|---|---|---|---|---|---|---|
| **Q9H6Z4** | RANBP3 | -0.050420121 | 3.171661683 | -3.092251853 | 0.019347613 | 0.305857884 | -4.213072033 |
| **P54105** | CLNS1A | -0.050597335 | 3.211720952 | -3.038497416 | 0.020795132 | 0.306942846 | -4.213644307 |
| **Q8TB37-1** | NUBPL | 0.182686274 | 3.137930554 | 2.161757961 | 0.070596428 | 0.402227373 | -4.215312926 |
| **Q9NVH0** | EXD2 | 0.184221671 | 3.170995263 | 1.864378533 | 0.107976921 | 0.477393788 | -4.218662929 |
| **Q96CB9** | NSUN4 | 0.154015246 | 3.159878465 | 2.542954172 | 0.041129379 | 0.355540374 | -4.219540881 |
| **Q9BS16** | CENPK | -0.087766592 | 3.208178963 | -2.433860945 | 0.047957734 | 0.362534913 | -4.220268312 |
| **Q96CM3** | RPUSD4 | 0.172619459 | 3.209522911 | 2.523193564 | 0.042286528 | 0.355540374 | -4.221187592 |
| **O43462** | MBTPS2 | -0.193953106 | 3.190942054 | -1.872665893 | 0.106707391 | 0.475441804 | -4.221314092 |
| **P13010** | XRCC5 | 0.031346055 | 3.177185613 | 3.720998546 | 0.008580724 | 0.283479686 | -4.22181286 |
| **O15382** | BCAT2 | 0.159250256 | 3.185470289 | 2.92864451 | 0.024127722 | 0.322917088 | -4.22313449 |
| **Q9UL42** | PNMA2 | -0.105028215 | 3.190456952 | -2.624251753 | 0.036706694 | 0.348943902 | -4.225600641 |
| **Q9NWU1-1** | OXSM | 0.18995325 | 3.188886803 | 2.571129131 | 0.039536501 | 0.35510265 | -4.226050017 |
| **Q99952** | PTPN18 | -0.274544596 | 3.245493482 | -2.011334818 | 0.087526506 | 0.436259778 | -4.226799907 |
| **Q96A49** | SYAP1 | 0.073006909 | 3.186073529 | 2.841375788 | 0.027181759 | 0.328952544 | -4.227743749 |
| **Q8NBE8** | PHOSPHO2-KLHL2 | -0.195474577 | 3.262208298 | -1.873643775 | 0.106558561 | 0.475403904 | -4.228881801 |
| **O43808** | SLC25A17 | -0.074942047 | 3.188875336 | -2.464832509 | 0.045906775 | 0.357577769 | -4.228925216 |
| **Q9H9P8-1** | L2HGDH | 0.181214548 | 3.195365014 | 2.891006137 | 0.025397443 | 0.328144258 | -4.229056025 |
| **Q9NVX7-2** | KBTBD4 | 0.076772421 | 3.170035541 | 2.534103722 | 0.041643498 | 0.355540374 | -4.231703739 |
| **Q9Y3C1** | NOP16 | -0.071899929 | 3.207390329 | -2.982466591 | 0.0224286 | 0.314277294 | -4.232753271 |
| **Q8NBT0** | POC1A | -0.073140099 | 3.191643933 | -2.739545697 | 0.031273793 | 0.33673269 | -4.233437963 |
| **Q5JRX3** | PITRM1 | 0.199000874 | 3.181514285 | 3.171185286 | 0.017415198 | 0.305857884 | -4.23470273 |
| **Q16626** | MEA1 | -0.54527535 | 3.164074801 | -2.147082295 | 0.072091216 | 0.404094866 | -4.237750759 |
| **O14757** | CHEK1 | -0.047503121 | 3.215201789 | -3.138338995 | 0.018192789 | 0.305857884 | -4.237784035 |
| **P82664** | MRPS10 | 0.305229614 | 3.164660471 | 2.524963283 | 0.042181522 | 0.355540374 | -4.239898481 |
| **Q8N5M1** | ATPAF2 | 0.197875569 | 3.185227585 | 2.194660848 | 0.067358486 | 0.395595275 | -4.240111629 |
| **O75879** | PET112; GATB | 0.192631947 | 3.161659481 | 2.455150494 | 0.046537938 | 0.358626496 | -4.240177854 |
| **O43824** | GTPBP6 | 0.136353353 | 3.191228212 | 2.307706065 | 0.057344042 | 0.377614679 | -4.241113547 |
| **P46781** | RPS9 | -0.120488197 | 3.148829172 | -3.129280735 | 0.018413761 | 0.305857884 | -4.245297176 |
| **P18065** | IGFBP2 | -0.298865938 | 3.237443608 | -2.858885478 | 0.026537454 | 0.328952544 | -4.247036926 |
| **Q99741** | CDC6 | -0.081926461 | 3.223211268 | -2.636236879 | 0.036098193 | 0.348481345 | -4.248141744 |
| **Q9NZN3** | EHD3 | 0.100049838 | 3.233254462 | 2.200956017 | 0.066756418 | 0.394427892 | -4.24899271 |
| **Q9UK53-1** | ING1 | 0.12907624 | 3.148922564 | 2.201503547 | 0.066704312 | 0.394427892 | -4.249241652 |
| **Q9NYK5-1** | MRPL39 | 0.166561668 | 3.218900392 | 3.028186753 | 0.021085859 | 0.307875291 | -4.251417697 |
| **Q5T160** | RARS2 | 0.14976749 | 3.1646796 | 2.478293421 | 0.045044039 | 0.357262465 | -4.253014521 |
| **Q13813-2** | SPTAN1 | 0.194670281 | 3.125032386 | 2.191445926 | 0.067668091 | 0.397062767 | -4.257182464 |
| **Q9Y605** | MRFAP1 | -0.132470679 | 3.289633704 | -2.236198036 | 0.063485801 | 0.389084368 | -4.25808065 |
| **Q9P0J1-2** | PDP1 | 0.182131627 | 3.183776757 | 2.706189221 | 0.032752374 | 0.339814312 | -4.260021229 |
| **P40818** | USP8 | 0.151947309 | 3.207228432 | 2.48201993 | 0.044808197 | 0.357204257 | -4.260660499 |
| **Q96M94** | KLHL15 | -0.108678081 | 3.187544389 | -2.306615064 | 0.057433011 | 0.377614679 | -4.26109013 |

| | | | | | | | |
|---|---|---|---|---|---|---|---|
| **P30084** | ECHS1 | 0.177716376 | 3.168407258 | 3.054346032 | 0.020356617 | 0.305857884 | -4.261542841 |
| **Q86VU5** | COMTD1 | 0.223855838 | 3.177586027 | 1.821831957 | 0.114731233 | 0.489548681 | -4.262096957 |
| **Q53GL7** | PARP10 | -0.05453818 | 3.187520925 | -2.670182144 | 0.034431706 | 0.345097821 | -4.262974843 |
| **Q02338** | BDH1 | 0.154999391 | 3.193693243 | 2.483888675 | 0.044690414 | 0.357204257 | -4.266054211 |
| **P09493-4** | TPM1 | -0.116168514 | 3.196406554 | -2.10576706 | 0.076473857 | 0.412201353 | -4.269023337 |
| **Q13526** | PIN1 | -0.048365067 | 3.182208504 | -2.82268454 | 0.027887956 | 0.32997678 | -4.270177174 |
| **Q86SQ7** | SDCCAG8 | -0.140752001 | 3.225991515 | -2.188387418 | 0.067963978 | 0.397403835 | -4.271408875 |
| **Q9P2R7** | SUCLA2 | 0.141557127 | 3.177358182 | 2.962979605 | 0.023028512 | 0.318193956 | -4.27475304 |
| **P17568** | NDUFB7 | 0.145402228 | 3.184965747 | 2.371440171 | 0.052385471 | 0.374522685 | -4.277100062 |
| **Q9Y4H4** | GPSM3 | -0.319753408 | 3.230961759 | -1.73834011 | 0.12920195 | 0.514283314 | -4.28239487 |
| **Q03154-1** | ACY1 | 0.132768542 | 3.150149905 | 2.029489062 | 0.085283827 | 0.431338184 | -4.28246998 |
| **P02774-3** | GC | 0.226401894 | 3.231013154 | 1.919329047 | 0.0998286 | 0.461022182 | -4.282572053 |
| **O95167** | NDUFA3 | 0.135854182 | 3.196419451 | 2.316540751 | 0.056628786 | 0.377614679 | -4.284083962 |
| **P62820** | RAB1A | 0.135307492 | 3.1526323 | 2.822709996 | 0.027886981 | 0.32997678 | -4.284865026 |
| **Q15797** | SMAD1 | 0.084959894 | 3.154020951 | 2.523782313 | 0.042251564 | 0.355540374 | -4.286666078 |
| **Q16134** | ETFDH | 0.158643586 | 3.204772884 | 2.701940241 | 0.03294596 | 0.339814312 | -4.287004508 |
| **O60333-3** | KIF1B | -0.106856052 | 3.253695594 | -2.185581806 | 0.068236557 | 0.398319894 | -4.28802129 |
| **Q7Z7H8-2** | MRPL10 | 0.145634921 | 3.182886896 | 2.435916844 | 0.047818683 | 0.362172964 | -4.288969074 |
| **Q96L50** | LRR1 | -0.071255236 | 3.218414165 | -2.455848751 | 0.046492118 | 0.358626496 | -4.289507978 |
| **O75182-1** | SIN3B | -0.071585289 | 3.208600094 | -2.724893732 | 0.031914376 | 0.33673269 | -4.289843408 |
| **Q7Z3T8** | ZFYVE16 | -0.044399757 | 3.198771948 | -2.859011766 | 0.026532867 | 0.328952544 | -4.290111316 |
| **A2RU30** | TESPA1 | 0.19576902 | 3.200651897 | 1.809044398 | 0.116840735 | 0.492826136 | -4.295546062 |
| **Q7Z6J2** | GRASP | -0.272955418 | 3.092471867 | -1.704880053 | 0.13548297 | 0.524095099 | -4.296093403 |
| **Q9NQX6** | ZNF331 | -0.133345478 | 3.237793348 | -2.09803032 | 0.077324021 | 0.414399115 | -4.297248044 |
| **Q13501-1** | SQSTM1 | -0.082012365 | 3.19035884 | -2.578170957 | 0.039148578 | 0.353048187 | -4.298410134 |
| **Q14108-1** | SCARB2 | -0.093239193 | 3.187618669 | -2.584228805 | 0.038818055 | 0.35246794 | -4.30091989 |
| **O75531** | BANF1 | -0.189476071 | 3.194942046 | -2.538883891 | 0.04136499 | 0.355540374 | -4.302356717 |
| **Q15928** | ZNF141 | -0.618667912 | 2.747675261 | -2.042734247 | 0.083683971 | 0.426895123 | -4.302846454 |
| **Q86W74** | ANKRD46 | -0.111174485 | 3.248380053 | -2.175614324 | 0.069213973 | 0.399872475 | -4.303269656 |
| **P62424** | RPL7A | -0.181953971 | 3.133840774 | -3.245711601 | 0.015780835 | 0.305857884 | -4.30384364 |
| **Q9NUL3** | STAU2 | 0.097189747 | 3.197500564 | 2.198240066 | 0.067015496 | 0.394739983 | -4.30612494 |
| **P0C7P0** | CISD3 | 0.209689579 | 3.149550852 | 2.337661655 | 0.054955829 | 0.376021618 | -4.306808872 |
| **Q9HCK8-1** | CHD8 | -0.030172652 | 3.184891815 | -3.254512147 | 0.015599073 | 0.305857884 | -4.308098055 |
| **O60566-3** | BUB1B | -0.085875952 | 3.180875163 | -2.977316501 | 0.022585502 | 0.315819071 | -4.310744138 |
| **Q8NHH9** | ATL2 | -0.125414963 | 3.171954887 | -2.48771963 | 0.044449964 | 0.357204257 | -4.312112656 |
| **Q07020** | RPL18 | -0.151798225 | 3.151779674 | -2.695549525 | 0.033239404 | 0.340757503 | -4.312438423 |
| **Q96EL2** | MRPS24 | 0.216628738 | 3.179962771 | 2.350103516 | 0.053994254 | 0.376021618 | -4.31303344 |
| **Q5TAP6** | UTP14C | 0.095749085 | 3.19780846 | 2.197111025 | 0.067123497 | 0.394900782 | -4.31398726 |
| **Q8TDN6** | BRIX1 | -0.115030712 | 3.201772568 | -2.909956585 | 0.024749462 | 0.32394477 | -4.314202447 |

| | | | | | | | |
|---|---|---|---|---|---|---|---|
| **Q9BX40** | LSM14B | -0.076220931 | 3.175294653 | -2.689386266 | 0.03352502 | 0.342800882 | -4.318316038 |
| **Q9NVS9** | PNPO | 0.071510115 | 3.191104596 | 2.757660528 | 0.03050058 | 0.336493392 | -4.320918581 |
| **Q96BW9-3** | TAMM41 | 0.168607312 | 3.174929364 | 2.483678064 | 0.044703672 | 0.357204257 | -4.324517352 |
| **Q96BD8** | SKA1 | -0.095044642 | 3.178296078 | -2.728423781 | 0.031758783 | 0.33673269 | -4.326243678 |
| **Q2NKX8** | ERCC6L | -0.045409895 | 3.187528262 | -3.193727929 | 0.016902334 | 0.305857884 | -4.328086438 |
| **Q69YI7-1** | NAIF1 | 0.103037798 | 3.164117548 | 2.318932779 | 0.056436712 | 0.377614679 | -4.328098683 |
| **Q9ULF5** | SLC39A10 | -0.069434722 | 3.229608699 | -2.362502632 | 0.053053248 | 0.375341102 | -4.331334686 |
| **P40429** | RPL13A | -0.182016133 | 3.167733794 | -3.258876101 | 0.015509784 | 0.305857884 | -4.331564132 |
| **Q5SSJ5-1** | HP1BP3 | -0.097763814 | 3.168105658 | -2.922669335 | 0.024324678 | 0.322917088 | -4.334135206 |
| **Q7Z3D6-2** | C14orf159; DGLUC | 0.189576847 | 3.162414057 | 2.517901014 | 0.042602195 | 0.355540374 | -4.339654825 |
| **Q9Y2Z4** | YARS2 | 0.182987219 | 3.20279315 | 2.357934563 | 0.053397947 | 0.376021618 | -4.339723548 |
| **Q8TF47-1** | ZFP90 | -0.096331202 | 3.177015202 | -2.133818024 | 0.073469842 | 0.405641342 | -4.34020778 |
| **P82663** | MRPS25 | 0.162647326 | 3.192230542 | 2.322848361 | 0.056123746 | 0.37738805 | -4.34073803 |
| **Q96AP4-1** | ZUFSP | -0.159801601 | 3.247322588 | -1.995314391 | 0.089554491 | 0.440291106 | -4.340765964 |
| **O14772** | FPGT | 0.08737691 | 3.195732064 | 2.348975099 | 0.054080744 | 0.376021618 | -4.341024983 |
| **O43156** | TTI1 | -0.06268617 | 3.194467395 | -3.08108362 | 0.019639186 | 0.305857884 | -4.341826911 |
| **Q14153** | FAM53B | -0.14120585 | 3.238340269 | -2.07750065 | 0.079626445 | 0.418407477 | -4.342158876 |
| **Q5EBL8-2** | PDZD11 | -0.132173329 | 3.229978921 | -2.404474188 | 0.049991822 | 0.36851891 | -4.342296902 |
| **Q92552-2** | MRPS27 | 0.178072354 | 3.220191778 | 2.85166472 | 0.026801164 | 0.328952544 | -4.3444993 |
| **Q96N46** | TTC14 | -0.058657716 | 3.176062713 | -2.505928909 | 0.04332537 | 0.357204257 | -4.348553891 |
| **O00142-6** | TK2 | 0.138890805 | 3.180181093 | 1.844993694 | 0.111004597 | 0.482457068 | -4.349737073 |
| **Q6NUK1** | SLC25A24 | 0.195324339 | 3.216060374 | 2.652302848 | 0.035299112 | 0.346250492 | -4.35175354 |
| **Q13547** | HDAC1 | 0.050767982 | 3.183272801 | 3.082923168 | 0.019590837 | 0.305857884 | -4.352656794 |
| **Q9H1X3-3** | DNAJC25-GNG10; | 0.136909227 | 3.248705579 | 2.163049731 | 0.070466371 | 0.401998301 | -4.352989495 |
| **Q96MW1** | CCDC43 | -0.088465013 | 3.180867037 | -2.726293111 | 0.031852599 | 0.33673269 | -4.356142994 |
| **Q96G25-2** | MED8 | 0.080326443 | 3.165590427 | 2.615694744 | 0.037147742 | 0.349573768 | -4.356584149 |
| **P62081** | RPS7 | -0.094218204 | 3.213138337 | -2.857651689 | 0.026582317 | 0.328952544 | -4.356649065 |
| **Q9P0B6** | CCDC167 | -0.122801631 | 3.200188773 | -2.026116649 | 0.085696044 | 0.431546517 | -4.356795359 |
| **P06576** | ATP5B | 0.194543323 | 3.193325772 | 3.344583708 | 0.013863901 | 0.305857884 | -4.357770097 |
| **Q5T1M5** | FKBP15 | -0.035061389 | 3.189643826 | -3.051196759 | 0.020442956 | 0.305857884 | -4.358408318 |
| **P04049** | RAF1 | -0.060824959 | 3.213387493 | -2.857604378 | 0.026584039 | 0.328952544 | -4.360933557 |
| **Q8TBP6** | SLC25A40 | 0.220072116 | 3.185271428 | 2.353747357 | 0.053715937 | 0.376021618 | -4.362719728 |
| **O94889** | KLHL18 | -0.099798212 | 3.216099964 | -2.079898521 | 0.079353997 | 0.417804844 | -4.36377677 |
| **P42695** | NCAPD3 | -0.092744414 | 3.191422101 | -3.131850195 | 0.018350788 | 0.305857884 | -4.364135764 |
| **Q8IWT6** | LRRC8A | 0.075151032 | 3.161022686 | 2.322253474 | 0.056171179 | 0.37738805 | -4.364460411 |
| **Q8IXM3** | MRPL41 | 0.159230402 | 3.198857918 | 2.452670756 | 0.046701038 | 0.359078004 | -4.365305985 |
| **P31350-2** | RRM2 | -0.125183753 | 3.207176931 | -2.849195826 | 0.02689197 | 0.328952544 | -4.369113843 |
| **O94985-2** | CLSTN1 | 0.098084551 | 3.155131122 | 2.555102625 | 0.040434474 | 0.355540374 | -4.369983779 |
| **Q07021** | C1QBP | 0.193573461 | 3.186251679 | 2.844186069 | 0.027077236 | 0.328952544 | -4.370463493 |

| | | | | | | | |
|---|---|---|---|---|---|---|---|
| **Q9NRD5** | PICK1 | -0.250616098 | 3.232353148 | -1.939563293 | 0.096983643 | 0.455254009 | -4.37263542 |
| **Q9NUS5** | AP5S1 | -0.117620257 | 3.195957362 | -1.974874647 | 0.092210084 | 0.446440655 | -4.372977543 |
| **Q9NZ72** | STMN3 | -0.138556742 | 3.300375605 | -2.168011402 | 0.06996908 | 0.401998301 | -4.373058742 |
| **P36551** | CPOX | 0.133512672 | 3.178594945 | 2.958804211 | 0.023159274 | 0.318891564 | -4.376506465 |
| **Q8N7B6** | PACRGL | 0.115582963 | 3.175931982 | 2.060267156 | 0.081612413 | 0.423686898 | -4.377939974 |
| **Q5RHP9** | C1orf173; ERICH3 | 0.253405356 | 3.121496945 | 1.599978495 | 0.157119467 | 0.555457754 | -4.378013601 |
| **Q8NBJ4** | GOLM1 | -0.130966716 | 3.221907238 | -2.184064195 | 0.068384464 | 0.398319894 | -4.37869931 |
| **Q9Y5X0** | SNX10 | -0.2144051 | 3.135271387 | -2.078027932 | 0.079566454 | 0.418381741 | -4.381447964 |
| **P86452** | ZBED6 | 0.126692401 | 3.216326981 | 2.007732012 | 0.087978541 | 0.43768911 | -4.38180267 |
| **Q13185** | CBX3; C15orf57; C( | 0.092634521 | 3.161263843 | 3.049557624 | 0.02048805 | 0.305857884 | -4.381893643 |
| **Q4G176** | ACSF3 | 0.171962415 | 3.193013297 | 2.653847146 | 0.03522329 | 0.346250492 | -4.384087826 |
| **O00584** | RNASET2 | -0.101351835 | 3.179270161 | -2.328787993 | 0.055652409 | 0.376021618 | -4.387472186 |
| **Q02218** | OGDH | 0.202204934 | 3.186088304 | 2.971304652 | 0.022770146 | 0.316828828 | -4.387576437 |
| **P62277** | RPS13 | -0.212934421 | 3.150359533 | -2.809306378 | 0.028405382 | 0.330820062 | -4.388433902 |
| **P56181-2** | NDUFV3 | 0.199328083 | 3.194718133 | 2.585872313 | 0.038728889 | 0.352231442 | -4.390279575 |
| **P62330** | ARF6 | -0.065152431 | 3.142624169 | -2.544900717 | 0.041017199 | 0.355540374 | -4.394144643 |
| **Q9UKA4** | AKAP11 | -0.051566945 | 3.202311165 | -3.012956659 | 0.021523309 | 0.309585734 | -4.396830341 |
| **Q96JG8-4** | MAGED4B; MAGED | -0.148248582 | 3.156331744 | -1.952102182 | 0.095261121 | 0.451948632 | -4.396959687 |
| **Q5SRD1** | TIMM23B; LINC008 | 0.264522582 | 3.232330426 | 2.36646213 | 0.052756332 | 0.374840509 | -4.398915269 |
| **Q96SL8** | FIZ1 | -0.153734197 | 3.181298201 | -1.899299701 | 0.102725951 | 0.468248815 | -4.399045511 |
| **P02786** | TFRC | -0.101872079 | 3.177478461 | -3.248706513 | 0.015718723 | 0.305857884 | -4.400712152 |
| **Q71F23** | MLF1IP; CENPU | -0.068878764 | 3.174997026 | -2.559456208 | 0.040188447 | 0.355540374 | -4.401046167 |
| **P12259** | F5 | -0.063107327 | 3.233608072 | -2.84368824 | 0.027095721 | 0.328952544 | -4.40240516 |
| **Q8N8Q8-1** | COX18 | 0.154614266 | 3.169636586 | 2.014866854 | 0.087085603 | 0.435428013 | -4.403801863 |
| **O75064** | DENND4B | 0.109350377 | 3.156567897 | 2.414398223 | 0.049295075 | 0.366780161 | -4.404869214 |
| **Q5T7W7** | TSTD2 | -0.149533058 | 3.236866265 | -2.253552556 | 0.061935849 | 0.385039377 | -4.407350425 |
| **Q17R31-3** | TATDN3 | 0.187003394 | 3.125057319 | 2.08336275 | 0.078962046 | 0.416709711 | -4.410148029 |
| **P32322** | PYCR1 | 0.167941387 | 3.212650882 | 2.790243033 | 0.029160391 | 0.333307048 | -4.411178657 |
| **Q9P015** | MRPL15 | 0.167298939 | 3.192937879 | 2.841183503 | 0.027188926 | 0.328952544 | -4.412518638 |
| **O75127** | PTCD1 | 0.156273487 | 3.229017239 | 2.542608503 | 0.041149334 | 0.355540374 | -4.414582181 |
| **Q15526** | SURF1 | 0.183194449 | 3.157710357 | 2.230689736 | 0.063986002 | 0.390752332 | -4.416570555 |
| **O15131** | KPNA5 | -0.105471306 | 3.202398461 | -1.971765247 | 0.092620881 | 0.44675374 | -4.416963339 |
| **Q9BZE2** | PUS3 | -0.086819518 | 3.185503556 | -2.396280801 | 0.050574753 | 0.368977419 | -4.417230562 |
| **Q15555** | MAPRE2 | -0.068322175 | 3.180905699 | -2.828394282 | 0.027670181 | 0.32997678 | -4.422979708 |
| **Q8WV99** | ZFAND2B | -0.126026747 | 3.220769124 | -1.989355286 | 0.090320745 | 0.442415735 | -4.423319509 |
| **Q8WZ42-12** | TTN | 0.214479025 | 3.210259726 | 1.929112334 | 0.098442894 | 0.457316603 | -4.423898277 |
| **Q13505-1** | MTX1 | 0.099173889 | 3.14595134 | 2.562866558 | 0.039996818 | 0.355476279 | -4.425821905 |
| **Q16649** | NFIL3 | -0.09946899 | 3.176829905 | -2.205300195 | 0.066344142 | 0.394427892 | -4.427503527 |
| **Q00059** | TFAM | 0.184737509 | 3.197385665 | 2.739960812 | 0.031255843 | 0.33673269 | -4.427726177 |

| | | | | | | | |
|---|---|---|---|---|---|---|---|
| **P78406** | RAE1 | 0.038462157 | 3.179470195 | 3.113373853 | 0.018808836 | 0.305857884 | -4.427782932 |
| **Q96HE7** | ERO1L; ERO1A | 0.06635976 | 3.169635926 | 3.005550915 | 0.021739533 | 0.310090823 | -4.429875197 |
| **Q9Y5J9** | TIMM8B | 0.161390074 | 3.241579121 | 2.331074854 | 0.055472025 | 0.376021618 | -4.4306857 |
| **Q96IR7** | HPDL | 0.134401873 | 3.162603114 | 2.160647105 | 0.070708466 | 0.402227373 | -4.431524498 |
| **Q93088** | BHMT | -0.285382224 | 3.167301827 | -1.527178834 | 0.174006309 | 0.574454961 | -4.432942668 |
| **P47914** | RPL29 | -0.16346733 | 3.168266405 | -2.839810442 | 0.027240164 | 0.328952544 | -4.433362591 |
| **Q96BR5** | SELRC1; COA7 | 0.182837908 | 3.173419591 | 2.791610545 | 0.029105527 | 0.333203768 | -4.437778062 |
| **Q16891-2** | IMMT | 0.238330996 | 3.173603069 | 2.320168516 | 0.056337748 | 0.377614679 | -4.438639968 |
| **Q8IWL3** | HSCB | 0.150288638 | 3.200904301 | 2.068458544 | 0.08066231 | 0.42108109 | -4.441591044 |
| **Q02252** | ALDH6A1 | 0.188280133 | 3.178125426 | 2.78818966 | 0.029242978 | 0.333336389 | -4.446248935 |
| **Q96GN5-1** | CDCA7L | -0.05403278 | 3.201209302 | -2.771572909 | 0.02992051 | 0.335760523 | -4.446550108 |
| **Q9BU70** | C9orf156; TRMO | -0.163614865 | 3.25972394 | -1.643624345 | 0.147746385 | 0.546231571 | -4.446629171 |
| **Q9BRP7** | FDXACB1 | 0.112018516 | 3.210403198 | 1.977569659 | 0.0918555 | 0.445393971 | -4.447517607 |
| **Q96D70** | R3HDM4 | 0.164892095 | 3.219852687 | 1.614295285 | 0.153984712 | 0.552139704 | -4.44815391 |
| **Q13433-1** | SLC39A6 | -0.073837093 | 3.190847448 | -2.303278926 | 0.057705951 | 0.377966845 | -4.44935793 |
| **Q8IXJ9** | ASXL1 | -0.054989839 | 3.183304073 | -2.599573977 | 0.037993843 | 0.352231442 | -4.45083027 |
| **Q14011** | CIRBP | 0.11278819 | 3.14316882 | 1.903305574 | 0.102139918 | 0.467185751 | -4.452705635 |
| **O60701** | UGDH | 0.050798451 | 3.189672867 | 2.778784696 | 0.029624433 | 0.334326872 | -4.456743203 |
| **P53350** | PLK1 | -0.087141359 | 3.162037932 | -2.679254794 | 0.034000176 | 0.345097821 | -4.460955011 |
| **Q92543** | SNX19 | -0.112731671 | 3.197642011 | -1.878958527 | 0.105753234 | 0.475403904 | -4.461643434 |
| **P50914** | RPL14 | -0.200548022 | 3.124507271 | -2.519947567 | 0.04247984 | 0.355540374 | -4.462582538 |
| **Q9H190** | SDCBP2 | -0.189564901 | 3.120730465 | -1.566725347 | 0.164633408 | 0.563893388 | -4.465176984 |
| **Q96I51** | WBSCR16; RCC1L | 0.189232584 | 3.18151563 | 2.154862921 | 0.071294765 | 0.403638325 | -4.469467561 |
| **P82909** | MRPS36 | 0.209574615 | 3.171577594 | 2.149587604 | 0.071833784 | 0.403638325 | -4.470010324 |
| **Q9H7P9-1** | PLEKHG2 | -0.074228016 | 3.172957081 | -2.272642533 | 0.060275588 | 0.383111637 | -4.470816987 |
| **Q9BVS5** | TRMT61B | 0.136964011 | 3.21487157 | 2.165001216 | 0.070270354 | 0.401998301 | -4.47208036 |
| **Q969V6** | MKL1 | -0.08001161 | 3.228342422 | -2.378597157 | 0.051856997 | 0.373191888 | -4.474247118 |
| **Q8IYU8** | EFHA1; MICU2 | 0.160310304 | 3.175193505 | 2.317746845 | 0.056531856 | 0.377614679 | -4.475157314 |
| **Q5T2D3** | OTUD3 | 0.133001303 | 3.197911248 | 1.719421144 | 0.132718133 | 0.519961998 | -4.476711057 |
| **Q5W0Q7** | USPL1 | -0.121602456 | 3.233473482 | -2.149089674 | 0.071884874 | 0.403638325 | -4.476888151 |
| **Q8IZ83** | ALDH16A1 | 0.050437864 | 3.18260422 | 2.652674537 | 0.035280847 | 0.346250492 | -4.477148079 |
| **O75554** | WBP4 | -0.04708608 | 3.186064499 | -2.632707536 | 0.036276272 | 0.348943902 | -4.477856886 |
| **O76031** | CLPX | 0.171505274 | 3.161953503 | 2.859480181 | 0.026515859 | 0.328952544 | -4.478169681 |
| **O14526-1** | FCHO1 | -0.055634405 | 3.201838307 | -2.516288877 | 0.042698837 | 0.355972987 | -4.47947175 |
| **Q13043-1** | STK4 | -0.048659544 | 3.188461986 | -2.862435698 | 0.026408812 | 0.328952544 | -4.483801269 |
| **Q8WXI2-1** | CNKSR2 | -0.146363821 | 3.202254835 | -1.837972731 | 0.112121566 | 0.484388026 | -4.486221526 |
| **Q96LI5-1** | CNOT6L | 0.067554466 | 3.149682349 | 2.412379723 | 0.04943597 | 0.367073627 | -4.486245982 |
| **O00506** | STK25 | -0.110419766 | 3.163863552 | -2.097334566 | 0.077400941 | 0.414399115 | -4.486394797 |
| **Q96E52-1** | OMA1 | 0.192316081 | 3.222389046 | 2.276431453 | 0.059951528 | 0.382023723 | -4.486534917 |

| | | | | | | | |
|---|---|---|---|---|---|---|---|
| **Q86SZ2** | TRAPPC6B | -0.070746348 | 3.183328426 | -2.551768514 | 0.040623957 | 0.355540374 | -4.486928826 |
| **P09001** | MRPL3 | 0.191221814 | 3.22002439 | 2.710574813 | 0.032553825 | 0.339814312 | -4.487465958 |
| **O77932** | DOM3Z; DXO | 0.088453927 | 3.191502847 | 2.084436115 | 0.078840999 | 0.416709711 | -4.489660325 |
| **P63208** | SKP1 | -0.095154417 | 3.191852553 | -2.685519929 | 0.033705515 | 0.343761638 | -4.491311087 |
| **O43491-4** | EPB41L2 | -0.055316085 | 3.189441537 | -3.192175463 | 0.016937125 | 0.305857884 | -4.492138804 |
| **P62256** | UBE2H | -0.063453732 | 3.187254675 | -2.344957194 | 0.054389868 | 0.376021618 | -4.49331126 |
| **Q96QD8** | SLC38A2 | 0.147309933 | 3.211142984 | 2.002514836 | 0.088637264 | 0.438495059 | -4.494562003 |
| **Q9BTL3** | FAM103A1 | -0.083342513 | 3.177141649 | -2.432676068 | 0.048038063 | 0.362796969 | -4.496582538 |
| **Q9Y4C2-1** | FAM115A; TCAF1 | -0.068908179 | 3.208720993 | -2.846520581 | 0.026990735 | 0.328952544 | -4.497298343 |
| **P49662** | CASP4 | -0.08324567 | 3.208166246 | -2.481471023 | 0.044842855 | 0.357204257 | -4.498237795 |
| **P24752** | ACAT1 | 0.176775952 | 3.177915931 | 3.005720699 | 0.02173455 | 0.310090823 | -4.498499001 |
| **P49888** | SULT1E1 | 0.193119054 | 3.231366788 | 1.643546531 | 0.147762614 | 0.546231571 | -4.500658397 |
| **Q5JS37** | NHLRC3 | 0.113883351 | 3.168261731 | 2.103970885 | 0.076670388 | 0.412980496 | -4.501197073 |
| **P53677** | AP3M2 | -0.096967533 | 3.190577673 | -2.321067781 | 0.056265843 | 0.377560911 | -4.501736857 |
| **P82912-1** | MRPS11 | 0.179896818 | 3.206891734 | 2.505323819 | 0.043362259 | 0.357204257 | -4.501926683 |
| **Q9NTX5** | ECHDC1 | -0.061034807 | 3.165467589 | -2.425998437 | 0.048493398 | 0.364158835 | -4.503073818 |
| **P08708** | RPS17; RPS17L | -0.055245441 | 3.199801859 | -3.170929257 | 0.017421119 | 0.305857884 | -4.50324404 |
| **O14874** | BCKDK | 0.189360761 | 3.216230508 | 2.0041029 | 0.088436235 | 0.437773138 | -4.504232286 |
| **Q92665** | MRPS31 | 0.175492742 | 3.174170737 | 2.652803393 | 0.035274517 | 0.346250492 | -4.50479598 |
| **P23193** | TCEA1 | 0.047881781 | 3.171036854 | 3.056261663 | 0.02030429 | 0.305857884 | -4.508713418 |
| **P12235** | SLC25A4 | 0.181345396 | 3.210030793 | 2.389178769 | 0.051085733 | 0.370663149 | -4.509105153 |
| **Q8NFG4-1** | FLCN | -0.093751772 | 3.19729525 | -2.403642808 | 0.050050653 | 0.36851891 | -4.509771091 |
| **Q15800** | MSMO1 | 0.065377782 | 3.103708073 | 2.350893598 | 0.053933781 | 0.376021618 | -4.512083589 |
| **Q6ZRI6** | C15orf39 | -0.058891756 | 3.265058126 | -2.483242786 | 0.044731087 | 0.357204257 | -4.51420149 |
| **P55273** | CDKN2D | -0.151678051 | 3.272648756 | -1.747048722 | 0.127613726 | 0.511807701 | -4.515766913 |
| **Q9NYY8** | FASTKD2 | 0.179471784 | 3.212149421 | 2.57574544 | 0.039281743 | 0.353847451 | -4.516321793 |
| **Q96DV4** | MRPL38 | 0.159019994 | 3.200623841 | 2.591155471 | 0.038443713 | 0.352231442 | -4.516730949 |
| **Q9BUN8** | DERL1 | 0.087812677 | 3.179100709 | 2.237541142 | 0.063364442 | 0.389084368 | -4.516768871 |
| **Q96PU8** | QKI | 0.071271728 | 3.183700585 | 2.714528745 | 0.032375909 | 0.339797349 | -4.518675334 |
| **O95835** | LATS1 | -0.055524412 | 3.243124317 | -2.495429289 | 0.043970148 | 0.357204257 | -4.521336778 |
| **Q96CU9** | FOXRED1 | 0.19611518 | 3.18936989 | 2.279841883 | 0.059661368 | 0.381343177 | -4.522913123 |
| **Q8WUM9** | SLC20A1 | -0.093502546 | 3.216671609 | -2.245568269 | 0.062644069 | 0.387695915 | -4.528931186 |
| **P16401** | HIST1H1B | -0.115549205 | 3.173892329 | -2.837999611 | 0.027307896 | 0.329227974 | -4.528986062 |
| **Q6PKC3-1** | TXNDC11 | -0.22017004 | 3.246070552 | -1.482620639 | 0.185158528 | 0.588198522 | -4.52944943 |
| **Q9H1E3** | NUCKS1 | 0.162627003 | 3.123626937 | 2.542986231 | 0.041127529 | 0.355540374 | -4.529910332 |
| **Q99797** | MIPEP | 0.117551883 | 3.182626427 | 2.641615049 | 0.035828596 | 0.347992906 | -4.530889455 |
| **Q9NVV4** | MTPAP | 0.216615296 | 3.189918172 | 2.743400296 | 0.031107541 | 0.33673269 | -4.531330816 |
| **O00716** | E2F3 | 0.101440005 | 3.214241756 | 2.254365149 | 0.061864232 | 0.384904994 | -4.531809705 |
| **Q8TCC3-2** | MRPL30 | 0.162675952 | 3.18176018 | 2.178272358 | 0.068951941 | 0.399844221 | -4.533747629 |

| | | | | | | | |
|---|---|---|---|---|---|---|---|
| **Q96ND0** | FAM210A | 0.207988065 | 3.185605364 | 2.439647778 | 0.047567411 | 0.361292298 | -4.533850575 |
| **Q15532** | SS18 | -0.115440537 | 3.195736636 | -2.262637074 | 0.06114 | 0.38369455 | -4.539364932 |
| **Q9BRX8-1** | FAM213A | -0.063815076 | 3.183840643 | -2.482646332 | 0.04476868 | 0.357204257 | -4.539811495 |
| **Q3KQU3** | MAP7D1 | -0.055821755 | 3.195588091 | -2.445005463 | 0.047208978 | 0.360649353 | -4.542673434 |
| **Q5W0U4** | BSPRY | 0.422500409 | 3.09247833 | 1.708889926 | 0.134715066 | 0.523635615 | -4.544023078 |
| **Q9NWU5** | MRPL22 | 0.164857633 | 3.222260204 | 2.215606531 | 0.065376392 | 0.393496543 | -4.545149648 |
| **Q8IVU3-1** | HERC6 | 0.15607537 | 3.124149567 | 1.645135163 | 0.147431602 | 0.546231571 | -4.549419021 |
| **O60313-2** | OPA1 | 0.144031468 | 3.16173858 | 3.058720494 | 0.020237337 | 0.305857884 | -4.55024337 |
| **Q6ICB0** | DESI1 | -0.085206376 | 3.164340717 | -1.980536969 | 0.09146666 | 0.444209209 | -4.550840488 |
| **O60513** | B4GALT4 | 0.169982972 | 3.200429698 | 1.509978311 | 0.178235672 | 0.57989621 | -4.551331923 |
| **O95801** | TTC4 | -0.067598256 | 3.203570101 | -2.787099568 | 0.029286923 | 0.333359029 | -4.556185579 |
| **Q9H790** | EXO5 | -0.165620386 | 3.231224032 | -1.488745325 | 0.183587508 | 0.586019588 | -4.556514166 |
| **Q6ZW76-1** | ANKS3 | 0.229274449 | 3.184721075 | 1.429937359 | 0.199189268 | 0.604614147 | -4.556591697 |
| **Q53QV2** | LBH | -0.441385457 | 3.173378588 | -1.386312805 | 0.211530927 | 0.61590617 | -4.556670297 |
| **P42574** | CASP3 | 0.057886963 | 3.17510076 | 2.765559989 | 0.030169767 | 0.336035644 | -4.556775423 |
| **Q8WZA0-2** | LZIC | 0.055276425 | 3.17367873 | 2.606781703 | 0.037613068 | 0.351571562 | -4.557302495 |
| **P53999** | SUB1 | 0.105596632 | 3.171014853 | 3.062475889 | 0.020135538 | 0.305857884 | -4.558380717 |
| **P46778** | RPL21 | -0.148714859 | 3.175177354 | -2.917603977 | 0.024492991 | 0.323253998 | -4.562534395 |
| **Q9NSV4-3** | DIAPH3 | -0.065390289 | 3.176834866 | -2.726392749 | 0.031848206 | 0.33673269 | -4.563850716 |
| **Q8N2A8** | PLD6 | -0.183320478 | 3.263321321 | -1.523379421 | 0.174932452 | 0.575026202 | -4.563929721 |
| **Q7Z7G8-2** | VPS13B | -0.044440708 | 3.220100575 | -2.704465205 | 0.032830776 | 0.339814312 | -4.565815731 |
| **Q9NYF5-1** | FAM13B | -0.107696772 | 3.233448079 | -1.7879108 | 0.120410072 | 0.499295152 | -4.566070436 |
| **Q9UJY4** | GGA2 | -0.079982207 | 3.22550615 | -2.460685438 | 0.046176021 | 0.358190865 | -4.567536912 |
| **Q02127** | DHODH | 0.186397222 | 3.202687946 | 2.663216955 | 0.034766917 | 0.345787303 | -4.569964613 |
| **Q99759-2** | MAP3K3 | -0.133642135 | 3.177561069 | -1.747570066 | 0.127519246 | 0.511687077 | -4.572516392 |
| **Q7Z7F0-1** | KIAA0907; KHDC4 | 0.055510519 | 3.171615497 | 2.652521955 | 0.035288344 | 0.346250492 | -4.572573211 |
| **Q9H582-1** | ZNF644 | -0.040165511 | 3.201005054 | -2.698477264 | 0.033104629 | 0.339814312 | -4.57307913 |
| **P52292** | KPNA2 | -0.084635392 | 3.182170718 | -2.88929552 | 0.02545682 | 0.328144258 | -4.573352555 |
| **Q8IZ81** | ELMOD2 | -0.049755176 | 3.207944274 | -2.645123941 | 0.035653846 | 0.347570313 | -4.573676375 |
| **P50336** | PPOX | 0.174620935 | 3.154508957 | 2.364701684 | 0.052888132 | 0.374840509 | -4.574250633 |
| **Q6P3W7** | SCYL2 | -0.03519987 | 3.196690244 | -2.966475051 | 0.022919651 | 0.317241514 | -4.575290394 |
| **P05165-1** | PCCA | 0.202906783 | 3.207492933 | 2.29821559 | 0.058122749 | 0.379089796 | -4.576520699 |
| **P42025** | ACTR1B | 0.060343479 | 3.183575198 | 2.294991886 | 0.058389725 | 0.379161351 | -4.576948576 |
| **Q9NRP4** | ACN9; SDHAF3 | 0.198869076 | 3.229891739 | 1.809727866 | 0.116727037 | 0.492826136 | -4.578555095 |
| **Q9NSE4** | IARS2 | 0.192905768 | 3.177827916 | 3.065266312 | 0.020060252 | 0.305857884 | -4.579977843 |
| **Q9BSL1** | UBAC1 | -0.092891477 | 3.195194181 | -2.346096278 | 0.054302046 | 0.376021618 | -4.580465416 |
| **O00411** | POLRMT | 0.156852073 | 3.192400261 | 2.629209703 | 0.036453674 | 0.348943902 | -4.580670532 |
| **O15347** | HMGB3 | 0.065581595 | 3.138628281 | 2.804778087 | 0.028582826 | 0.331403622 | -4.584513961 |
| **Q9Y4D8-5** | HECTD4 | 0.079880445 | 3.178131849 | 2.12678243 | 0.074211894 | 0.406910627 | -4.584673791 |

| | | | | | | | |
|---|---|---|---|---|---|---|---|
| **Q9UHY8-2** | FEZ2 | -0.109540267 | 3.217280846 | -1.863986017 | 0.108037416 | 0.477395592 | -4.587209294 |
| **Q99714-1** | HSD17B10 | 0.148916006 | 3.172975235 | 2.747202086 | 0.030944484 | 0.33673269 | -4.587596124 |
| **Q9BTE7** | DCUN1D5 | -0.058434381 | 3.188544862 | -2.446293669 | 0.047123215 | 0.360340657 | -4.589018188 |
| **Q14197** | ICT1; MRPL58 | 0.119486355 | 3.196571247 | 2.596449779 | 0.038160146 | 0.352231442 | -4.590001277 |
| **P62266** | RPS23 | -0.158763946 | 3.194809234 | -2.57086382 | 0.039551196 | 0.35510265 | -4.593907595 |
| **O00217** | NDUFS8 | 0.160086091 | 3.184712727 | 2.568627263 | 0.039675294 | 0.355476279 | -4.595188717 |
| **O43395** | PRPF3 | -0.049925707 | 3.19207615 | -2.884530691 | 0.025622992 | 0.328538932 | -4.595470177 |
| **P05387** | RPLP2 | -0.109863253 | 3.201684181 | -3.07554657 | 0.019785494 | 0.305857884 | -4.596377524 |
| **A3KMH1** | VWA8 | 0.168633372 | 3.189880596 | 2.917596085 | 0.024493254 | 0.323253998 | -4.596522286 |
| **Q16181** | 7-Sep | 0.037500302 | 3.173630764 | 3.164304211 | 0.017575078 | 0.305857884 | -4.596659841 |
| **Q08881** | ITK | 0.050864201 | 3.156080242 | 2.619962975 | 0.036927055 | 0.348943902 | -4.599304904 |
| **Q15468-2** | STIL | -0.068817435 | 3.179233619 | -2.241383035 | 0.063018611 | 0.387825609 | -4.599456056 |
| **Q9BZ95** | WHSC1L1; NSD3 | -0.073471116 | 3.222554377 | -2.458767896 | 0.046301072 | 0.358190865 | -4.601052188 |
| **O43819** | SCO2 | 0.17627245 | 3.221247875 | 2.209465893 | 0.065951242 | 0.394427892 | -4.601481089 |
| **Q9NRL2** | BAZ1A | -0.051851504 | 3.19193761 | -3.139727655 | 0.018159167 | 0.305857884 | -4.603237574 |
| **Q8WWC4** | C2orf47; MAIP1 | 0.184900373 | 3.210319762 | 2.165369601 | 0.070233413 | 0.401998301 | -4.604437349 |
| **Q9H3K6-1** | BOLA2B; BOLA2; L | -0.064898553 | 3.183312419 | -2.586532325 | 0.038693142 | 0.352231442 | -4.604568922 |
| **O95995** | GAS8 | 0.149244374 | 3.232458281 | 1.82616329 | 0.11402517 | 0.48833912 | -4.606772708 |
| **Q8N1S5-1** | SLC39A11 | -0.351468913 | 2.865145828 | -1.284362105 | 0.243077669 | 0.649360189 | -4.607465765 |
| **Q9UDY4** | DNAJB4 | 0.081311308 | 3.198113747 | 2.111326145 | 0.07586882 | 0.411304347 | -4.607490855 |
| **Q56VL3** | OCIAD2 | 0.212677964 | 3.181704185 | 2.440469903 | 0.047512227 | 0.361229326 | -4.609632721 |
| **O75312** | ZNF259; ZPR1 | -0.063519763 | 3.21246807 | -2.740652527 | 0.031225958 | 0.33673269 | -4.613184453 |
| **O75477** | ERLIN1 | -0.063087542 | 3.188947391 | -2.481784711 | 0.044823046 | 0.357204257 | -4.613190249 |
| **Q8N9M5** | TMEM102 | 0.08584202 | 3.188335236 | 2.265143496 | 0.060922274 | 0.383378328 | -4.613865998 |
| **P55061-2** | TMBIM6 | 0.103715395 | 3.208413314 | 2.176740073 | 0.069102872 | 0.399872475 | -4.613877076 |
| **Q9H2G9-1** | BLZF1 | -0.119006129 | 3.205002276 | -1.835249851 | 0.112557703 | 0.485489116 | -4.614407824 |
| **Q4G0I0** | C16orf91 | 0.236840396 | 3.211138153 | 1.961067241 | 0.094048198 | 0.448519793 | -4.615236442 |
| **Q8IWT0** | ZBTB8OS | -0.115112232 | 3.186550913 | -1.799674905 | 0.118410306 | 0.496152504 | -4.616010742 |
| **O15235** | MRPS12 | 0.165925759 | 3.185726422 | 2.093152736 | 0.077864896 | 0.414913883 | -4.616872834 |
| **Q9H467** | CUEDC2 | -0.122884129 | 3.192426698 | -2.348161068 | 0.054143226 | 0.376021618 | -4.617945903 |
| **O15118** | NPC1 | -0.105758049 | 3.185038209 | -1.936514941 | 0.097407052 | 0.455770924 | -4.621843366 |
| **Q9HCU5** | PREB | -0.062996566 | 3.195749276 | -2.528421805 | 0.041977097 | 0.355540374 | -4.623519098 |
| **Q9BTT6** | LRRC1 | 0.076493724 | 3.176274065 | 2.015391357 | 0.087020319 | 0.435375587 | -4.625759263 |
| **Q9BWN1** | PRR14 | 0.128324311 | 3.232478762 | 1.536021514 | 0.171868409 | 0.571776747 | -4.627408485 |
| **Q9HBI0-1** | PARVG | -0.051503278 | 3.19906937 | -2.638686526 | 0.035975133 | 0.348430364 | -4.628043964 |
| **Q9NXE8** | CWC25 | -0.075616154 | 3.176649141 | -2.408081074 | 0.049737416 | 0.367936472 | -4.628923359 |
| **O00463** | TRAF5 | -0.06142581 | 3.201849015 | -2.488277697 | 0.04441505 | 0.357204257 | -4.629000825 |
| **Q96DX7** | TRIM44 | -0.129710131 | 3.267553241 | -2.149061586 | 0.071887757 | 0.403638325 | -4.630175484 |
| **Q8WWB7** | C1orf85; GLMP | 0.142509714 | 3.197997048 | 1.490459216 | 0.183150085 | 0.585730065 | -4.630615536 |

| | | | | | | | |
|---|---|---|---|---|---|---|---|
| **Q969H8** | C19orf10; MYDGF | -0.054950952 | 3.186734546 | -2.485053589 | 0.044617155 | 0.357204257 | -4.632736428 |
| **P46939** | UTRN | -0.029176813 | 3.167786252 | -3.08158337 | 0.019626038 | 0.305857884 | -4.632752501 |
| **P20592** | MX2 | -0.047953416 | 3.155833025 | -2.639948497 | 0.03591191 | 0.348376219 | -4.634072233 |
| **P61513** | RPL37A | -0.075400594 | 3.194840965 | -2.888196191 | 0.025495056 | 0.328144258 | -4.636948463 |
| **O95772-1** | STARD3NL | -0.1196878 | 3.146138939 | -1.674210994 | 0.141496302 | 0.537118068 | -4.637415144 |
| **Q9BV90** | SNRNP25 | 0.107572443 | 3.213563325 | 1.797486788 | 0.118779801 | 0.496164837 | -4.638360507 |
| **Q53T94** | TAF1B | -0.111274339 | 3.175620877 | -1.969756374 | 0.09288725 | 0.44685719 | -4.638668141 |
| **Q5BJF6** | ODF2 | -0.15020509 | 3.203249655 | -1.665644872 | 0.143220872 | 0.539313633 | -4.64000661 |
| **Q9BYD6** | MRPL1 | 0.173379357 | 3.200028654 | 2.506031052 | 0.043319146 | 0.357204257 | -4.641838556 |
| **Q92966** | SNAPC3 | -0.099613974 | 3.237349818 | -2.10814303 | 0.076214668 | 0.411642107 | -4.642778148 |
| **P14927** | UQCRB | 0.148449423 | 3.22564627 | 2.459787194 | 0.046234556 | 0.358190865 | -4.642788094 |
| **Q5T2S8** | ARMC4 | -0.234747712 | 3.244713724 | -1.278772409 | 0.244921282 | 0.650150213 | -4.644151181 |
| **P83731** | RPL24 | -0.139418122 | 3.141853076 | -2.686638361 | 0.033653197 | 0.343669218 | -4.644854399 |
| **P43250-2** | GRK6 | -0.063072533 | 3.174934402 | -2.565118236 | 0.039870821 | 0.355476279 | -4.644957611 |
| **Q9UNW1** | MINPP1 | 0.080479487 | 3.192339805 | 2.315357016 | 0.056724086 | 0.377614679 | -4.645177773 |
| **O43772** | SLC25A20 | 0.185770315 | 3.201005998 | 2.38230909 | 0.051585082 | 0.372073031 | -4.64686645 |
| **Q7Z589-5** | C11orf30; EMSY | 0.042533626 | 3.173120327 | 2.532640169 | 0.04172916 | 0.355540374 | -4.647312808 |
| **Q9NUB1** | ACSS1 | 0.141947061 | 3.190605651 | 2.672380411 | 0.034326621 | 0.345097821 | -4.647312961 |
| **Q9NS91** | RAD18 | -0.049873253 | 3.164402008 | -2.743173123 | 0.031117313 | 0.33673269 | -4.649814765 |
| **Q6P9G4** | TMEM154 | -0.343056719 | 3.062830117 | -1.447095936 | 0.194516494 | 0.600401532 | -4.65006303 |
| **P10109** | FDX1 | 0.176989523 | 3.204096275 | 1.902660861 | 0.102234012 | 0.467205335 | -4.650127711 |
| **O43516-1** | WIPF1 | 0.10254069 | 3.157666577 | 2.467357986 | 0.04574361 | 0.357414656 | -4.650508709 |
| **Q96AD5-1** | PNPLA2 | 0.078551514 | 3.16777618 | 2.147151365 | 0.072084106 | 0.404094866 | -4.650775269 |
| **Q9NQH7-1** | XPNPEP3 | 0.165475862 | 3.173510567 | 2.158116455 | 0.070964374 | 0.403010689 | -4.651005094 |
| **Q9UHD8-1** | 9-Sep | -0.043534222 | 3.185474833 | -3.15768969 | 0.017730258 | 0.305857884 | -4.651121471 |
| **Q16706** | MAN2A1 | -0.050146873 | 3.188457708 | -2.560760484 | 0.040115046 | 0.355476279 | -4.651297398 |
| **P11182** | DBT | 0.156861034 | 3.206304656 | 2.438992282 | 0.047611458 | 0.361292298 | -4.651418639 |
| **Q9NPJ3-1** | ACOT13 | 0.09587553 | 3.177526803 | 2.564957215 | 0.039879817 | 0.355476279 | -4.654440023 |
| **Q9P0M9** | MRPL27 | 0.196558945 | 3.188261592 | 2.551276267 | 0.04065201 | 0.355540374 | -4.654761466 |
| **O43709-3** | WBSCR22; BUD23 | -0.072095822 | 3.189681524 | -2.228497554 | 0.064186189 | 0.390938548 | -4.655504179 |
| **P08962** | CD63 | -0.106730188 | 3.213127595 | -1.830894136 | 0.113258828 | 0.487243547 | -4.655965421 |
| **Q12851** | MAP4K2 | 0.09625867 | 3.181529147 | 2.211999337 | 0.065713454 | 0.394427892 | -4.656419676 |
| **Q9BZE1** | MRPL37 | 0.213075232 | 3.189954363 | 2.66315682 | 0.034769826 | 0.345787303 | -4.657397859 |
| **Q9Y277** | VDAC3 | 0.223059321 | 3.177087296 | 2.887835565 | 0.025507613 | 0.328144258 | -4.658369088 |
| **Q16875** | PFKFB3 | -0.231455236 | 3.155648129 | -2.467245335 | 0.045750875 | 0.357414656 | -4.66127049 |
| **P49821** | NDUFV1 | 0.163097483 | 3.193758903 | 2.603509871 | 0.037785416 | 0.352231442 | -4.661430944 |
| **Q3KQV9** | UAP1L1 | -0.078624836 | 3.206548221 | -2.121366726 | 0.074788257 | 0.407645581 | -4.663081968 |
| **P49336** | CDK8 | -0.093146736 | 3.177492012 | -2.129760138 | 0.07389691 | 0.406305157 | -4.663654868 |
| **Q7Z6M2** | FBXO33 | 0.105477161 | 3.176169328 | 2.018031837 | 0.086692405 | 0.434648757 | -4.663759792 |

| | | | | | | | |
|---|---|---|---|---|---|---|---|
| **Q9BQP7** | C20orf72; MGME1 | 0.207764085 | 3.176912371 | 2.676160817 | 0.034146693 | 0.345097821 | -4.666444511 |
| **Q9HB09-1** | BCL2L12 | -0.109005945 | 3.200520571 | -1.782082061 | 0.121413004 | 0.499487166 | -4.667896588 |
| **P78348-1** | ASIC1 | -0.264072935 | 3.1946136 | -1.193723049 | 0.27450338 | 0.674170362 | -4.668251521 |
| **P19419-1** | ELK1 | 0.11816475 | 3.168034621 | 1.842450322 | 0.11140796 | 0.483014501 | -4.668325652 |
| **O00488** | ZNF593 | -0.151828644 | 3.23612221 | -1.838772977 | 0.111993702 | 0.484371236 | -4.669443834 |
| **Q9HBH1** | PDF | 0.14321732 | 3.202573226 | 2.162889189 | 0.070482521 | 0.401998301 | -4.670543316 |
| **A6NK58** | LIPT2 | 0.168285698 | 3.213313961 | 1.865151413 | 0.1078579 | 0.477133083 | -4.672860131 |
| **P04181-1** | OAT | 0.186147135 | 3.190193632 | 2.470228782 | 0.045558869 | 0.357414656 | -4.673654204 |
| **P55084** | HADHB | 0.169921395 | 3.18663916 | 2.86774929 | 0.026217513 | 0.328952544 | -4.674529068 |
| **Q5HYK3-1** | COQ5 | 0.228961893 | 3.160705089 | 2.339999611 | 0.054773801 | 0.376021618 | -4.674659309 |
| **Q96P47-4** | AGAP3 | -0.102136688 | 3.173241686 | -1.770894739 | 0.123360713 | 0.50442659 | -4.677878461 |
| **P19404** | NDUFV2 | 0.160818301 | 3.187818214 | 2.488585024 | 0.044395835 | 0.357204257 | -4.678412261 |
| **O95831-1** | AIFM1 | 0.161582332 | 3.17903648 | 2.670307227 | 0.034425718 | 0.345097821 | -4.679386737 |
| **P82933** | MRPS9 | 0.18847142 | 3.202050485 | 2.624970334 | 0.036669909 | 0.348943902 | -4.679496312 |
| **P06276** | BCHE | -0.16160867 | 3.210193829 | -1.383857942 | 0.212245472 | 0.617236559 | -4.680423201 |
| **Q6UX07** | DHRS13 | 0.087278426 | 3.204071428 | 1.977369585 | 0.091881777 | 0.445393971 | -4.682750935 |
| **P13051** | UNG | -0.129787158 | 3.210254637 | -1.798250082 | 0.118650781 | 0.496152504 | -4.685312121 |
| **Q9NVC3-1** | SLC38A7 | -0.172255494 | 3.151849414 | -1.46969949 | 0.188513454 | 0.592460202 | -4.685523696 |
| **O75911** | DHRS3 | 0.137357007 | 3.158383375 | 1.758057711 | 0.125632917 | 0.509142907 | -4.685588942 |
| **Q8NE86** | MCU | 0.217972145 | 3.212953551 | 2.430662629 | 0.048174886 | 0.363139916 | -4.686126129 |
| **Q8NHG7** | SVIP | -0.134654317 | 3.166790077 | -1.728373248 | 0.131043018 | 0.517978495 | -4.68687938 |
| **Q969P6** | TOP1MT | 0.186830317 | 3.226623821 | 1.720564182 | 0.132503108 | 0.519961998 | -4.688639093 |
| **Q9P2E3-1** | ZNFX1 | -0.037149842 | 3.19412917 | -2.702433008 | 0.032923447 | 0.339814312 | -4.688708007 |
| **Q9Y2U9-1** | KLHDC2 | 0.078015456 | 3.149522914 | 2.310061924 | 0.057152407 | 0.377614679 | -4.689639342 |
| **Q969Z0-1** | TBRG4 | 0.171516823 | 3.187659125 | 2.732897613 | 0.031562743 | 0.33673269 | -4.690689385 |
| **P05166-2** | PCCB | 0.124691679 | 3.167335301 | 2.241721428 | 0.062988242 | 0.387825609 | -4.691219062 |
| **O15514** | POLR2D | -0.125404047 | 3.207450447 | -1.988251827 | 0.09046335 | 0.442736078 | -4.691422789 |
| **Q9Y2Z2-6** | MTO1 | 0.143172312 | 3.176524358 | 2.140320854 | 0.072790662 | 0.404796855 | -4.693243107 |
| **P17931** | LGALS3 | 0.229255468 | 3.065745243 | 1.598735023 | 0.157394551 | 0.555457754 | -4.693404664 |
| **Q8N5I9** | C12orf45 | -0.064361688 | 3.246223517 | -2.355984794 | 0.053545776 | 0.376021618 | -4.693848924 |
| **Q9NR30-1** | DDX21 | -0.096141199 | 3.173926429 | -3.136973166 | 0.018225924 | 0.305857884 | -4.693980753 |
| **Q8TCS8** | PNPT1 | 0.174171921 | 3.192734604 | 2.87970035 | 0.02579263 | 0.328952544 | -4.697877518 |
| **P50542-3** | PEX5 | -0.135099922 | 3.206356502 | -1.82189352 | 0.114721168 | 0.489548681 | -4.700783573 |
| **Q16795** | NDUFA9 | 0.138386303 | 3.19699282 | 2.701089748 | 0.032984854 | 0.339814312 | -4.704622921 |
| **O75909-4** | CCNK | -0.083841415 | 3.200109848 | -2.607176437 | 0.037592331 | 0.351571562 | -4.70588865 |
| **Q9P032** | NDUFAF4 | 0.143733521 | 3.197965809 | 2.360142656 | 0.05323104 | 0.376021618 | -4.706165711 |
| **Q96II8-2** | LRCH3 | -0.059188304 | 3.178632056 | -2.357688158 | 0.053416605 | 0.376021618 | -4.706715543 |
| **Q96GY0** | ZC2HC1A | 0.147089743 | 3.221722868 | 1.377144083 | 0.214210774 | 0.619843797 | -4.706903408 |
| **Q9BXK1** | KLF16 | -0.069072163 | 3.241597276 | -2.235312389 | 0.063565955 | 0.389084368 | -4.706998559 |

| | | | | | | | |
|---|---|---|---|---|---|---|---|
| **Q9UMY1-1** | NOL7 | -0.122326524 | 3.175226837 | -2.28162291 | 0.059510411 | 0.381175634 | -4.711778641 |
| **Q8TE82** | SH3TC1 | -0.067009115 | 3.203249028 | -2.149559447 | 0.071836672 | 0.403638325 | -4.711885665 |
| **Q9UI43** | FTSJ2; MRM2 | 0.159517609 | 3.170326058 | 1.551466262 | 0.168192786 | 0.566515005 | -4.714630258 |
| **O15234** | CASC3 | -0.08629332 | 3.215278066 | -2.229582448 | 0.064087038 | 0.390938548 | -4.714774871 |
| **Q10588** | BST1 | -0.132072952 | 3.130551473 | -1.820446459 | 0.114957986 | 0.489727184 | -4.71503864 |
| **P28331-2** | NDUFS1 | 0.164650896 | 3.179823943 | 2.82789959 | 0.027688977 | 0.32997678 | -4.715179244 |
| **O43542** | XRCC3 | -0.049351622 | 3.222740446 | -2.309085795 | 0.057231729 | 0.377614679 | -4.715333858 |
| **P49411** | TUFM | 0.153910107 | 3.186306642 | 3.00013141 | 0.021899241 | 0.311022609 | -4.715439823 |
| **A0MZ66-3** | KIAA1598; SHTN1 | 0.059521355 | 3.168166129 | 2.205476453 | 0.066327469 | 0.394427892 | -4.715779233 |
| **Q9UNK9** | ANGEL1 | 0.095923535 | 3.176356215 | 2.202780696 | 0.066582933 | 0.394427892 | -4.717052899 |
| **Q96BP2** | CHCHD1 | 0.193670123 | 3.167084509 | 2.096631965 | 0.077478696 | 0.414504124 | -4.717060301 |
| **Q8NDF8-5** | PAPD5 | -0.233060719 | 3.176788049 | -1.220375918 | 0.264919418 | 0.66823064 | -4.71715861 |
| **Q6ZN54** | DEF8 | -0.084350675 | 3.196434979 | -1.919081151 | 0.099863962 | 0.461022182 | -4.718322634 |
| **Q9Y2I8-2** | WDR37 | 0.053984814 | 3.162603958 | 2.409627824 | 0.04962873 | 0.367474614 | -4.719167858 |
| **Q969J3-1** | LOH12CR1; BORC[illegible] | -0.067828282 | 3.185655135 | -2.14507632 | 0.072298013 | 0.404094866 | -4.719209766 |
| **Q16656-4** | NRF1 | -0.065625222 | 3.177856245 | -2.372289562 | 0.052322462 | 0.374506268 | -4.719437857 |
| **Q6P1M3-1** | LLGL2 | -0.174311945 | 3.143150934 | -1.362903324 | 0.21843341 | 0.625163344 | -4.719509236 |
| **Q9H981-1** | ACTR8 | -0.058048585 | 3.195851307 | -2.60169209 | 0.037881528 | 0.352231442 | -4.722035705 |
| **Q14061** | COX17 | -0.229125083 | 3.19279593 | -2.101769674 | 0.076911932 | 0.41316112 | -4.722367562 |
| **O14730** | RIOK3 | -0.076886318 | 3.212142064 | -2.201753725 | 0.066680518 | 0.394427892 | -4.723950051 |
| **Q9Y4W6** | AFG3L2 | 0.221536049 | 3.183815113 | 2.822111035 | 0.02790993 | 0.32997678 | -4.724082882 |
| **Q15573-1** | TAF1A | -0.150763037 | 3.179847849 | -1.722700785 | 0.132102072 | 0.519814431 | -4.724469912 |
| **O43296** | ZNF264 | -0.12675706 | 3.183983932 | -1.662558425 | 0.143847141 | 0.539341922 | -4.724542611 |
| **Q16625-1** | OCLN | 0.214821942 | 2.72284128 | 1.127767117 | 0.299488747 | 0.690882899 | -4.72530656 |
| **Q96C92** | SDCCAG3 | -0.172364665 | 3.164832759 | -1.523834593 | 0.174821259 | 0.574976204 | -4.725726871 |
| **Q96MH2** | HEXIM2 | -0.150955334 | 3.15701823 | -1.478310767 | 0.186271431 | 0.588831734 | -4.726320004 |
| **O94768** | STK17B | -0.116405435 | 3.154163797 | -1.969548939 | 0.092914799 | 0.44685719 | -4.729108507 |
| **P32519-1** | ELF1 | 0.04551999 | 3.1611362 | 2.581714591 | 0.038954877 | 0.352501702 | -4.729146349 |
| **Q8N8K9-2** | KIAA1958 | -0.135206771 | 3.205937705 | -1.743475891 | 0.128263027 | 0.512600477 | -4.729449922 |
| **P34897-1** | SHMT2 | 0.140279615 | 3.206430569 | 2.924089837 | 0.0242777 | 0.322917088 | -4.730794745 |
| **O00458** | IFRD1 | -0.157157522 | 3.27533629 | -1.620733365 | 0.152594359 | 0.552031202 | -4.731337155 |
| **Q96C36** | PYCR2 | 0.185530974 | 3.188201727 | 2.627858337 | 0.036522456 | 0.348943902 | -4.731579686 |
| **Q712K3** | UBE2R2 | -0.089899008 | 3.192113405 | -2.372950315 | 0.0522735 | 0.374493198 | -4.732111009 |
| **Q15583** | TGIF1 | 0.148047924 | 3.145304217 | 1.532556542 | 0.172703215 | 0.57291317 | -4.732434798 |
| **Q13576-1** | IQGAP2 | 0.036372635 | 3.173897777 | 2.993827081 | 0.022086608 | 0.312238617 | -4.736977081 |
| **Q9H6X2** | ANTXR1 | 0.106090177 | 3.145982719 | 2.054995391 | 0.082229814 | 0.424506738 | -4.737137334 |
| **Q15040** | JOSD1 | 0.098406135 | 3.20434279 | 1.782158575 | 0.121399786 | 0.499487166 | -4.737158809 |
| **Q96FC9-2** | DDX11 | -0.06791124 | 3.227396133 | -2.421236402 | 0.048820847 | 0.364893351 | -4.737565652 |
| **Q96EA4** | SPDL1 | -0.089090763 | 3.186493936 | -2.337804232 | 0.05494471 | 0.376021618 | -4.739921024 |

| | | | | | | | |
|---|---|---|---|---|---|---|---|
| **P11279** | LAMP1 | -0.123415367 | 3.182720297 | -2.443788162 | 0.047290169 | 0.360768714 | -4.740237819 |
| **Q3MII6-1** | TBC1D25 | 0.084988491 | 3.157371796 | 1.973010779 | 0.09245611 | 0.446659583 | -4.74216446 |
| **O75439** | PMPCB | 0.169983399 | 3.208271334 | 2.784294644 | 0.029400319 | 0.333759678 | -4.743076501 |
| **P13073** | COX4I1 | 0.197552508 | 3.199612139 | 2.530886851 | 0.041832026 | 0.355540374 | -4.743245746 |
| **Q9Y3D3** | MRPS16 | 0.181656347 | 3.189179366 | 2.138290688 | 0.073002014 | 0.404796855 | -4.744855227 |
| **Q9NVH6-8** | TMLHE | 0.212383971 | 3.174226631 | 1.865207353 | 0.107849291 | 0.477133083 | -4.745295772 |
| **Q8N0Z8** | PUSL1 | 0.120271453 | 3.240950987 | 1.863402067 | 0.108127478 | 0.477527965 | -4.745506433 |
| **Q9HD40-1** | SEPSECS | -0.055404894 | 3.164816232 | -2.271082017 | 0.060409578 | 0.383145812 | -4.746011247 |
| **Q9GZS1-2** | POLR1E | -0.087570775 | 3.168621122 | -2.520174824 | 0.042466276 | 0.355540374 | -4.746338513 |
| **Q99828-2** | CIB1 | -0.132492419 | 3.234724281 | -1.898653278 | 0.102820829 | 0.468365275 | -4.747622099 |
| **O75323-1** | GBAS; NIPSNAP2 | 0.158740065 | 3.200526139 | 2.169230286 | 0.069847462 | 0.401998301 | -4.74874851 |
| **Q8NI37** | PPTC7 | 0.16056674 | 3.198302638 | 1.942857155 | 0.096528182 | 0.454581358 | -4.749210637 |
| **Q96E29** | MTERFD1; MTERF3 | 0.184113554 | 3.211547344 | 2.26574663 | 0.06087 | 0.383378328 | -4.749842124 |
| **O95822** | MLYCD | 0.152338012 | 3.192912672 | 2.34308712 | 0.054534366 | 0.376021618 | -4.749928974 |
| **Q96G28-2** | CCDC104; CFAP36 | -0.070881031 | 3.213514025 | -2.067973568 | 0.08071825 | 0.42108109 | -4.750174687 |
| **O00418** | EEF2K; LOC10193( | 0.060473401 | 3.182011781 | 2.16487632 | 0.070282882 | 0.401998301 | -4.75273625 |
| **P82673-1** | MRPS35 | 0.178387182 | 3.176241779 | 2.499849596 | 0.043697486 | 0.357204257 | -4.754643022 |
| **Q1ZZU3** | SWI5 | -0.098707077 | 3.205572415 | -1.876328078 | 0.106151067 | 0.475403904 | -4.754885101 |
| **Q56P03** | EAPP | 0.058202786 | 3.164484016 | 2.27633009 | 0.059960174 | 0.382023723 | -4.755530498 |
| **P61586** | RHOA | 0.231182272 | 3.12997914 | 2.331550053 | 0.055434618 | 0.376021618 | -4.755833245 |
| **O75575-1** | CRCP | -0.16713867 | 3.221047763 | -2.015525051 | 0.087003686 | 0.435375587 | -4.755835432 |
| **O95363** | FARS2 | 0.158692054 | 3.200386143 | 2.219284712 | 0.065034514 | 0.393401641 | -4.7562358 |
| **Q01850** | CDR2; LOC101060 | -0.075641866 | 3.238010797 | -2.086258777 | 0.07863588 | 0.416709711 | -4.758875048 |
| **Q13428-4** | TCOF1 | -0.073528634 | 3.184587279 | -2.175207292 | 0.069254188 | 0.399872475 | -4.758880637 |
| **P46779** | RPL28 | -0.072729195 | 3.171635971 | -2.675075991 | 0.034198223 | 0.345097821 | -4.759171847 |
| **P27986** | PIK3R1 | -0.053559821 | 3.201005233 | -2.416878954 | 0.049122487 | 0.366458362 | -4.760228684 |
| **Q9BYC8** | MRPL32 | 0.202438487 | 3.184492376 | 2.410345442 | 0.049578389 | 0.367474614 | -4.76173673 |
| **P27540** | ARNT | -0.061180075 | 3.18334123 | -2.352074005 | 0.053843564 | 0.376021618 | -4.76494961 |
| **Q9H999** | PANK3 | -0.139769318 | 3.128620279 | -1.790049284 | 0.12004413 | 0.499295152 | -4.76751733 |
| **Q96GC5-1** | MRPL48 | 0.166680522 | 3.215296513 | 2.316460926 | 0.056635208 | 0.377614679 | -4.769228012 |
| **P05141** | SLC25A5 | 0.208493601 | 3.193917795 | 2.504477715 | 0.043413896 | 0.357204257 | -4.769447328 |
| **Q13371** | PDCL | -0.084939167 | 3.216524771 | -2.12322776 | 0.074589689 | 0.407645581 | -4.769654844 |
| **Q96EB1-1** | ELP4 | -0.046335033 | 3.206294541 | -2.632315771 | 0.036296096 | 0.348943902 | -4.771037709 |
| **Q2NKX9-1** | C2orf68 | -0.118423062 | 3.173159497 | -1.637884013 | 0.148948215 | 0.546231571 | -4.772583933 |
| **Q86Z02** | HIPK1 | 0.06680014 | 3.170092344 | 2.012861714 | 0.08733563 | 0.435855266 | -4.772619845 |
| **Q96B97-1** | SH3KBP1 | -0.128476412 | 3.205609579 | -1.565823854 | 0.164841722 | 0.563893388 | -4.772752942 |
| **Q969X5** | ERGIC1 | -0.07213184 | 3.200793383 | -2.221255099 | 0.064852122 | 0.393401641 | -4.773170686 |
| **Q9P275-2** | USP36 | -0.072417795 | 3.196792953 | -2.762743083 | 0.030287295 | 0.336035644 | -4.773508246 |
| **Q9H4I3-1** | TRABD | 0.139813926 | 3.194650091 | 2.073987084 | 0.08002736 | 0.419060309 | -4.773860119 |

| | | | | | | | |
|---|---|---|---|---|---|---|---|
| **Q9H9Y2** | RPF1 | -0.080609781 | 3.217770312 | -2.06523034 | 0.081035408 | 0.422181192 | -4.773864262 |
| **Q9NVI7-2** | ATAD3A | 0.207686349 | 3.185435488 | 2.286101909 | 0.059132509 | 0.379928744 | -4.774352078 |
| **Q96E09** | FAM122A | -0.141585932 | 3.210326691 | -2.259555405 | 0.061408789 | 0.384167582 | -4.774461625 |
| **Q92932-3** | PTPRN2 | 0.279636352 | 3.225896805 | 2.107228825 | 0.076314292 | 0.411900167 | -4.775084255 |
| **Q86U38** | NOP9 | -0.06203196 | 3.182360716 | -2.481934757 | 0.044813573 | 0.357204257 | -4.775429473 |
| **Q9BV44** | THUMPD3 | -0.046209677 | 3.181845926 | -2.592918099 | 0.038349061 | 0.352231442 | -4.776314576 |
| **Q03989** | ARID5A | 0.09775671 | 3.166031203 | 1.675885701 | 0.141161463 | 0.536483353 | -4.776577748 |
| **O60711-2** | LPXN | -0.100224924 | 3.197721089 | -2.244927196 | 0.062701291 | 0.387695915 | -4.777114292 |
| **Q8NC60** | NOA1 | 0.118928152 | 3.200503651 | 2.329874768 | 0.055566611 | 0.376021618 | -4.777704025 |
| **P82930** | MRPS34 | 0.126946402 | 3.183754999 | 2.305677298 | 0.057509599 | 0.377614679 | -4.778241362 |
| **O95571** | ETHE1 | 0.150381852 | 3.219460274 | 2.20135196 | 0.066718734 | 0.394427892 | -4.779023013 |
| **O43464** | HTRA2 | 0.122216754 | 3.179786969 | 2.058723149 | 0.081792755 | 0.423686898 | -4.779310316 |
| **Q9UPT9-1** | USP22 | -0.059140922 | 3.19336238 | -2.545291049 | 0.040994743 | 0.355540374 | -4.779739499 |
| **Q9H2U1** | DHX36 | -0.050742237 | 3.186874406 | -2.807374963 | 0.028480922 | 0.330820062 | -4.781071999 |
| **Q13049** | TRIM32 | -0.044720715 | 3.211965766 | -2.637213936 | 0.036049057 | 0.348430364 | -4.781677313 |
| **Q969K3-2** | RNF34 | -0.085756794 | 3.174809689 | -2.066131817 | 0.080931046 | 0.421914148 | -4.782862511 |
| **Q13398-8** | ZNF211 | 0.081905338 | 3.181103354 | 1.916601987 | 0.100218287 | 0.461621569 | -4.78339892 |
| **O43791** | SPOP | -0.099472973 | 3.196199913 | -1.961841869 | 0.093944117 | 0.448519793 | -4.784881728 |
| **P16333-1** | NCK1 | -0.058750437 | 3.191135714 | -2.633193349 | 0.036251705 | 0.348943902 | -4.787477523 |
| **Q9Y5R4** | HEMK1 | 0.18720137 | 3.133840974 | 1.075227374 | 0.320718721 | 0.708595727 | -4.788846079 |
| **P42766** | RPL35 | -0.171113635 | 3.119085563 | -2.095477933 | 0.077606581 | 0.414504124 | -4.788853373 |
| **Q969T9** | WBP2 | -0.082148652 | 3.190942938 | -2.33463136 | 0.05519269 | 0.376021618 | -4.788962336 |
| **Q9NX14-2** | NDUFB11 | 0.194398061 | 3.186203997 | 2.422579059 | 0.048728291 | 0.364544514 | -4.789379946 |
| **P54619-3** | PRKAG1 | -0.066584556 | 3.179423484 | -2.520150757 | 0.042467712 | 0.355540374 | -4.792368619 |
| **Q9Y250** | LZTS1 | -0.066887962 | 3.206748195 | -2.255406677 | 0.061772562 | 0.384904994 | -4.792380885 |
| **Q96PK6-5** | RBM14-RBM4; RBI | 0.104192864 | 3.18225484 | 1.661013763 | 0.144161544 | 0.539756583 | -4.793205057 |
| **Q96K58-2** | ZNF668 | -0.091457884 | 3.16117981 | -2.215799538 | 0.065358407 | 0.393496543 | -4.793631916 |
| **P21359-1** | NF1 | 0.042367386 | 3.175618757 | 2.882840031 | 0.025682231 | 0.328575403 | -4.795661075 |
| **Q8TBZ8** | ZNF564 | -0.113044087 | 3.131966991 | -1.524562574 | 0.174643558 | 0.5748429 | -4.795925982 |
| **Q7KYR7** | BTN2A1 | 0.098825923 | 3.150828193 | 1.721477951 | 0.132331452 | 0.519961998 | -4.796552416 |
| **P55197** | MLLT10 | 0.095360729 | 3.188123846 | 2.150990909 | 0.071689995 | 0.403638325 | -4.797049311 |
| **Q9H7F4** | TMEM185B | -0.102036199 | 3.229424782 | -1.721218052 | 0.132380254 | 0.519961998 | -4.797178749 |
| **P22695** | UQCRC2 | 0.206426043 | 3.196180695 | 2.727721352 | 0.031789679 | 0.33673269 | -4.797218491 |
| **Q8IVV7** | GID4 | -0.063323932 | 3.232314869 | -2.326924949 | 0.05579981 | 0.376021618 | -4.797967969 |
| **Q8N344** | MIER2 | -0.091421375 | 3.238214573 | -1.996504458 | 0.089402244 | 0.440291106 | -4.798894117 |
| **Q15742-1** | NAB2 | -0.075674216 | 3.216485366 | -2.189140833 | 0.067890969 | 0.397403835 | -4.800301598 |
| **P31930** | UQCRC1 | 0.216032739 | 3.188524845 | 2.768376467 | 0.030052741 | 0.336035644 | -4.801333533 |
| **Q96G21** | IMP4 | -0.082527835 | 3.16966237 | -2.386889751 | 0.051251561 | 0.37144139 | -4.80347407 |
| **Q9P2K6** | KLHDC5; KLHL42 | -0.127704502 | 3.220524922 | -1.798634376 | 0.118585875 | 0.496152504 | -4.804003091 |

| | | | | | | | |
|---|---|---|---|---|---|---|---|
| **Q9NWS8** | RMND1 | 0.175177454 | 3.181614974 | 2.127959579 | 0.074087212 | 0.406910627 | -4.804271971 |
| **Q7L3S4** | ZNF771 | 0.133128277 | 3.201354343 | 1.192249678 | 0.275041682 | 0.674446347 | -4.8047575 |
| **P07203** | GPX1 | 0.082159372 | 3.170392153 | 2.392165698 | 0.05087018 | 0.369774545 | -4.805022096 |
| **Q8N0U4-1** | FAM185A | 0.185134933 | 3.182583097 | 1.981261611 | 0.091371951 | 0.444209209 | -4.80571254 |
| **P15529-2** | CD46 | -0.082734824 | 3.170606528 | -2.075879698 | 0.079811152 | 0.418547591 | -4.808575783 |
| **Q9UQ88-1** | CDK11A | -0.13108024 | 3.168697708 | -1.281329412 | 0.244076413 | 0.650150213 | -4.808810955 |
| **Q8N183** | NDUFAF2 | 0.171499144 | 3.210309649 | 2.499661949 | 0.043709025 | 0.357204257 | -4.810349068 |
| **Q6YN16** | HSDL2 | 0.109396116 | 3.184213491 | 2.619233665 | 0.036964666 | 0.348943902 | -4.81118961 |
| **P78417** | GSTO1 | -0.050215577 | 3.15609853 | -2.808727093 | 0.028428016 | 0.330820062 | -4.811633525 |
| **Q9NYP3-1** | DONSON | -0.079234921 | 3.17058097 | -1.997880995 | 0.089226463 | 0.440291106 | -4.812326391 |
| **Q04756** | HGFAC | 0.180190181 | 3.147220162 | 1.568509505 | 0.164221856 | 0.563603735 | -4.81282588 |
| **Q92947-1** | GCDH | 0.141866618 | 3.18986707 | 2.629983812 | 0.036414334 | 0.348943902 | -4.812837347 |
| **Q99538** | LGMN | -0.133630368 | 3.17864138 | -2.042864565 | 0.083668381 | 0.426895123 | -4.813505433 |
| **Q53EQ6-1** | TIGD5 | -0.160129384 | 3.248655995 | -1.122127017 | 0.301710723 | 0.691344519 | -4.815065424 |
| **P53814-6** | SMTN | -0.131033201 | 3.090984587 | -1.270764712 | 0.247583653 | 0.652280071 | -4.817423968 |
| **Q9H6Y2** | WDR55 | 0.064850468 | 3.169042872 | 2.518294475 | 0.042578643 | 0.355540374 | -4.817588131 |
| **Q9UK80-1** | USP21 | -0.181637986 | 3.060509538 | -1.023080726 | 0.342983793 | 0.724735701 | -4.8180271 |
| **Q9HAW0** | BRF2 | -0.096231165 | 3.214670258 | -1.757499856 | 0.125732572 | 0.509142907 | -4.818402698 |
| **O60353-1** | FZD6 | -0.097474258 | 3.233033187 | -1.856659514 | 0.109172714 | 0.479873319 | -4.819993816 |
| **Q5T9A4** | ATAD3B | 0.214574656 | 3.214242386 | 1.496013083 | 0.181739198 | 0.584582158 | -4.821107928 |
| **O94887-1** | FARP2 | -0.081498469 | 3.252560578 | -1.865742821 | 0.107766913 | 0.477133083 | -4.821614827 |
| **Q5PSV4** | BRMS1L | -0.060148706 | 3.185177226 | -2.166398221 | 0.070130371 | 0.401998301 | -4.821877628 |
| **Q13423** | NNT | 0.206656166 | 3.193931559 | 2.928285001 | 0.024139524 | 0.322917088 | -4.824548162 |
| **P46776** | RPL27A | -0.152330875 | 3.173404057 | -2.637854867 | 0.036016863 | 0.348430364 | -4.825231823 |
| **Q96K83** | ZNF521 | 0.147415561 | 3.160132185 | 1.219683463 | 0.265164748 | 0.66823064 | -4.825537893 |
| **Q10469** | MGAT2 | -0.056377732 | 3.16970933 | -2.236233338 | 0.063482608 | 0.389084368 | -4.827100257 |
| **Q9H0R3** | TMEM222 | -0.138746607 | 3.168756494 | -1.479705023 | 0.185910733 | 0.588813889 | -4.828443078 |
| **Q3ZAQ7** | VMA21 | -0.173022408 | 3.208183905 | -1.944819426 | 0.096257857 | 0.454408005 | -4.828543731 |
| **Q96LL9** | DNAJC30 | 0.171578496 | 3.22401851 | 1.413921151 | 0.203642655 | 0.608995611 | -4.829062381 |
| **Q9NV06** | DCAF13 | -0.068936313 | 3.192225458 | -2.443410038 | 0.047315419 | 0.360768714 | -4.829919784 |
| **Q969Z4** | RELT | 0.156485832 | 3.159600842 | 0.993289427 | 0.356246555 | 0.734600528 | -4.830156281 |
| **P33993-1** | MCM7 | 0.043670362 | 3.176379052 | 3.078014321 | 0.019720143 | 0.305857884 | -4.830189038 |
| **P30793-1** | GCH1 | -0.05271963 | 3.238971574 | -2.273814824 | 0.060175132 | 0.382779365 | -4.830766501 |
| **Q06609** | RAD51 | -0.069379977 | 3.203962045 | -2.328110803 | 0.055705941 | 0.376021618 | -4.831594473 |
| **Q6P161** | MRPL54 | 0.142905992 | 3.192990176 | 1.997021006 | 0.089336241 | 0.440291106 | -4.831614054 |
| **Q9UI09** | NDUFA12 | 0.127554765 | 3.170181603 | 2.189950711 | 0.067812578 | 0.397403835 | -4.832911036 |
| **Q7Z340** | ZNF551 | 0.141112148 | 3.17348806 | 1.603903783 | 0.156254071 | 0.554834464 | -4.833693492 |
| **Q6UWD8** | C16orf54; LOC101 | -0.154347128 | 3.149153863 | -1.509641611 | 0.178319403 | 0.57989621 | -4.833819374 |
| **Q96RN5** | MED15 | -0.052616717 | 3.204236079 | -2.468350736 | 0.045679637 | 0.357414656 | -4.834091584 |

| | | | | | | | |
|---|---|---|---|---|---|---|---|
| **Q9BT73** | PSMG3 | -0.077382094 | 3.181541926 | -1.995502669 | 0.089530387 | 0.440291106 | -4.834931816 |
| **Q8WVZ9** | KBTBD7 | 0.077117757 | 3.176932874 | 2.155024061 | 0.071278365 | 0.403638325 | -4.834954012 |
| **P49207** | RPL34 | -0.162512294 | 3.176446007 | -2.295537929 | 0.058344415 | 0.379161351 | -4.837458861 |
| **Q8WUK0** | PTPMT1 | 0.158248081 | 3.185679019 | 1.966308855 | 0.093346156 | 0.447307121 | -4.838735292 |
| **Q9C035** | TRIM5 | -0.091521579 | 3.15090656 | -2.160919639 | 0.070680962 | 0.402227373 | -4.839782236 |
| **Q8IZQ1-1** | WDFY3 | 0.08421702 | 3.162922811 | 1.680819816 | 0.140179331 | 0.535444609 | -4.840512472 |
| **O75683** | SURF6 | -0.075576047 | 3.210647087 | -2.347455145 | 0.05419747 | 0.376021618 | -4.84060013 |
| **P24863-1** | CCNC | -0.100644415 | 3.177893991 | -1.87932961 | 0.10569723 | 0.475403904 | -4.840731595 |
| **Q96DA6** | DNAJC19 | 0.143791427 | 3.202050826 | 2.455897593 | 0.046488915 | 0.358626496 | -4.841003524 |
| **P39023** | RPL3 | -0.118254984 | 3.173678648 | -3.136820552 | 0.01822963 | 0.305857884 | -4.842369099 |
| **Q2VPK5-1** | CTU2 | -0.089488158 | 3.190881385 | -2.159538857 | 0.070820421 | 0.40248086 | -4.842645397 |
| **P17535** | JUND | -0.109469273 | 3.204243897 | -1.715676306 | 0.133424953 | 0.521170723 | -4.842791075 |
| **P37173-2** | TGFBR2 | 0.244989996 | 3.127626557 | 1.282137956 | 0.243809789 | 0.650023079 | -4.84307559 |
| **Q9Y5Q9** | GTF3C3 | -0.057908058 | 3.214550864 | -2.607167258 | 0.037592813 | 0.351571562 | -4.843444411 |
| **Q9BVK2-1** | ALG8 | -0.116528621 | 3.2077 | -1.984079092 | 0.091004641 | 0.443577837 | -4.84467914 |
| **P54652** | HSPA2 | -0.404120094 | 2.93688541 | -1.472101409 | 0.187885617 | 0.591197824 | -4.845688759 |
| **O15321-1** | TM9SF1 | -0.065721351 | 3.143276025 | -2.094313562 | 0.077735827 | 0.414504124 | -4.846689434 |
| **O95168-1** | NDUFB4 | 0.178387756 | 3.211812255 | 2.073178588 | 0.0801199 | 0.419060309 | -4.848932737 |
| **O60664-1** | PLIN3 | -0.031475051 | 3.185202471 | -2.669006124 | 0.034488064 | 0.345097821 | -4.849148814 |
| **Q9UBT6** | POLK | -0.048732774 | 3.193476673 | -2.42420698 | 0.048616315 | 0.364544514 | -4.850595064 |
| **Q969E8** | TSR2 | -0.156498491 | 3.225425732 | -1.843755771 | 0.111200745 | 0.482781376 | -4.85083958 |
| **Q5SW79-1** | CEP170 | -0.137679948 | 3.127929668 | -1.462592355 | 0.190382454 | 0.594337366 | -4.851719189 |
| **Q8TDG4** | HELQ | 0.083835909 | 3.205565955 | 1.94425016 | 0.096336202 | 0.454507796 | -4.852002648 |
| **P56385** | ATP5I | 0.151102004 | 3.212587912 | 2.254347498 | 0.061865787 | 0.384904994 | -4.853018468 |
| **Q9Y305-4** | ACOT9 | 0.178565304 | 3.180940777 | 2.400338234 | 0.050285205 | 0.368557155 | -4.855081895 |
| **A6NFQ2** | FAM115C; TCAF2 | 0.142324398 | 3.195908122 | 0.977787277 | 0.363305592 | 0.738441741 | -4.855168612 |
| **Q53HC0** | CCDC92 | -0.191750788 | 3.192057378 | -1.44210391 | 0.19586557 | 0.601527622 | -4.855928734 |
| **P58317** | ZNF121 | -0.078319974 | 3.218794018 | -1.765657138 | 0.124282946 | 0.50674745 | -4.856840281 |
| **Q6P9H5-1** | GIMAP6 | 0.070919461 | 3.167738752 | 2.224810378 | 0.064524342 | 0.392319975 | -4.860747953 |
| **Q8NE01** | CNNM3 | 0.056680154 | 3.175195428 | 2.309327924 | 0.057212043 | 0.377614679 | -4.861008105 |
| **O95299** | NDUFA10 | 0.233614838 | 3.176669861 | 2.470122265 | 0.045565709 | 0.357414656 | -4.861316042 |
| **Q96EN9-1** | C19orf60; REX1BD | -0.129903804 | 3.185538753 | -1.410599935 | 0.204577342 | 0.610527836 | -4.863429605 |
| **P36542-1** | ATP5C1 | 0.167530183 | 3.202598318 | 2.563727222 | 0.039948609 | 0.355476279 | -4.863507864 |
| **Q96JY6-5** | PDLIM2 | 0.048418802 | 3.2013513 | 2.327724091 | 0.055736534 | 0.376021618 | -4.864071208 |
| **P57075** | UBASH3A | -0.050395009 | 3.184309707 | -2.605309136 | 0.037690535 | 0.351881669 | -4.865026424 |
| **Q9NRA8-3** | EIF4ENIF1 | -0.071461539 | 3.191326703 | -2.398304953 | 0.050430089 | 0.368977419 | -4.865580931 |
| **Q16829** | DUSP7 | -0.082476415 | 3.237928882 | -1.792457744 | 0.119633282 | 0.49841973 | -4.865689043 |
| **Q969S9** | GFM2 | 0.194524021 | 3.184200982 | 2.200718488 | 0.066779035 | 0.394427892 | -4.866638912 |
| **Q02447-1** | SP3 | -0.112766424 | 3.149491898 | -1.633903451 | 0.149787042 | 0.546701296 | -4.867366172 |

| | | | | | | | |
|---|---|---|---|---|---|---|---|
| **Q9Y6H1** | CHCHD2 | 0.113611723 | 3.192750409 | 1.788498097 | 0.120309464 | 0.499295152 | -4.867660169 |
| **Q5T013** | HYI | 0.147709528 | 3.175463899 | 1.574097927 | 0.162938994 | 0.561580888 | -4.868303098 |
| **P58557** | YBEY | 0.20299571 | 3.192213916 | 2.027331091 | 0.08554737 | 0.431538958 | -4.868825187 |
| **Q9UN86** | G3BP2 | -0.114275577 | 3.208506007 | -1.860471848 | 0.108580511 | 0.478730389 | -4.869095608 |
| **Q969W9** | PMEPA1 | -0.194453461 | 3.178708639 | -1.202546712 | 0.271298325 | 0.671838876 | -4.869469803 |
| **O75937** | DNAJC8 | 0.065921507 | 3.179502577 | 2.762837097 | 0.030283365 | 0.336035644 | -4.871487357 |
| **Q9UQ90** | SPG7 | 0.163394091 | 3.167378907 | 2.188145106 | 0.067987476 | 0.397403835 | -4.871513574 |
| **P86790** | CCZ1; CCZ1B | -0.051019025 | 3.216775237 | -2.539695738 | 0.041317883 | 0.355540374 | -4.872772275 |
| **Q96FZ2** | C3orf37; HMCES | -0.063017551 | 3.158048298 | -2.346290975 | 0.054287049 | 0.376021618 | -4.873768731 |
| **Q2QGD7** | ZXDC | -0.071385522 | 3.217728176 | -1.791894611 | 0.119729222 | 0.498558001 | -4.873849406 |
| **Q8NFP9-1** | NBEA | 0.098429002 | 3.189089941 | 1.479285335 | 0.18601924 | 0.588813889 | -4.873867381 |
| **Q9Y6M9** | NDUFB9 | 0.17409427 | 3.201876334 | 2.174335849 | 0.069340367 | 0.400056839 | -4.874226003 |
| **Q9HAV7** | GRPEL1 | 0.118105156 | 3.179075315 | 2.555527967 | 0.040410368 | 0.355540374 | -4.874955342 |
| **P35914** | HMGCL | 0.133159217 | 3.186848005 | 2.153387684 | 0.071445086 | 0.403638325 | -4.876052675 |
| **P49796-3** | RGS3 | -0.048492162 | 3.195890367 | -2.393362075 | 0.050784107 | 0.369486931 | -4.876523908 |
| **Q8IYQ7** | THNSL1 | 0.17635724 | 3.184147613 | 2.393523924 | 0.050772474 | 0.369486931 | -4.876961327 |
| **Q9Y3D9** | MRPS23 | 0.16156885 | 3.203596059 | 2.325595072 | 0.055905273 | 0.376413045 | -4.877233458 |
| **O94905-1** | ERLIN2 | 0.046865215 | 3.183520245 | 2.540494261 | 0.041271605 | 0.355540374 | -4.878297295 |
| **Q8N823-1** | ZNF611 | -0.107100297 | 3.219738046 | -1.574639803 | 0.162815102 | 0.561580888 | -4.881023846 |
| **Q8NC96** | NECAP1 | -0.068378116 | 3.205645862 | -2.245723553 | 0.062630216 | 0.387695915 | -4.881121454 |
| **O15047** | SETD1A | 0.046116674 | 3.174332461 | 2.579524856 | 0.039074452 | 0.352883173 | -4.881147314 |
| **Q8N4H5** | TOMM5 | 0.232271486 | 3.169794134 | 1.939520455 | 0.09698958 | 0.455254009 | -4.882786147 |
| **O00161** | SNAP23 | -0.099603532 | 3.165181712 | -2.344878086 | 0.054395972 | 0.376021618 | -4.883005192 |
| **Q6P1X6** | C8orf82 | 0.128670881 | 3.182733239 | 2.039892067 | 0.084024717 | 0.427385647 | -4.885624837 |
| **Q8N9A8-2** | CNEP1R1 | -0.146109005 | 3.188664691 | -1.313721087 | 0.233592526 | 0.640812038 | -4.886647944 |
| **P0C0L4-1** | C4A; C4B | 0.096709276 | 3.230371887 | 2.068339626 | 0.080676023 | 0.42108109 | -4.887010117 |
| **P42696** | RBM34 | -0.154453168 | 3.133070345 | -2.240261103 | 0.063119402 | 0.388145242 | -4.887055628 |
| **Q2TB18** | ASTE1 | -0.067043274 | 3.197272999 | -2.201627051 | 0.066692565 | 0.394427892 | -4.887138346 |
| **Q49A88-1** | CCDC14 | -0.069325125 | 3.215650093 | -1.931340573 | 0.098129956 | 0.456822639 | -4.888309764 |
| **Q8TDD1-2** | DDX54 | -0.108768754 | 3.199869997 | -2.613999583 | 0.037235773 | 0.349690566 | -4.889999284 |
| **O00635** | TRIM38 | 0.053819103 | 3.160815609 | 2.417189886 | 0.049100899 | 0.366458362 | -4.891975433 |
| **Q6PK04** | CCDC137 | -0.128760099 | 3.170223863 | -2.054118399 | 0.082332977 | 0.424508448 | -4.892007577 |
| **O95999** | BCL10 | -0.08492337 | 3.196879417 | -2.263568166 | 0.061059026 | 0.38369455 | -4.894563655 |
| **Q13029** | PRDM2 | -0.085482721 | 3.197436274 | -2.132655056 | 0.073591981 | 0.405653949 | -4.894828902 |
| **Q92535** | PIGC | 0.114779329 | 3.203887533 | 1.529512618 | 0.173439684 | 0.574167672 | -4.89557949 |
| **Q9C0C9** | UBE2O | -0.030052105 | 3.18332514 | -2.957118575 | 0.023212289 | 0.319068571 | -4.895908455 |
| **Q96MC6** | HIAT1; MFSD14A | 0.188641998 | 3.057122837 | 1.347040013 | 0.223224517 | 0.629408553 | -4.896078119 |
| **O43924** | PDE6D | -0.079380236 | 3.189321859 | -1.898322562 | 0.102869403 | 0.468365275 | -4.89613933 |
| **Q9NPQ8** | RIC8A | -0.050019018 | 3.182674596 | -2.504367536 | 0.043420625 | 0.357204257 | -4.896540142 |

| | | | | | | | |
|---|---|---|---|---|---|---|---|
| **P0DI81-3** | TRAPPC2 | -0.121503927 | 3.148311408 | -1.643548728 | 0.147762156 | 0.546231571 | -4.896587693 |
| **P17542-1** | TAL1 | -0.074089259 | 3.229357688 | -1.877594148 | 0.105959402 | 0.475403904 | -4.897060411 |
| **O43181** | NDUFS4 | 0.135846162 | 3.191564326 | 2.134643852 | 0.073383235 | 0.405447658 | -4.899161866 |
| **Q7Z7L1** | SLFN11 | -0.038913168 | 3.218194313 | -2.511191712 | 0.043005903 | 0.357204257 | -4.89934776 |
| **Q9Y2G7** | ZFP30 | -0.128534964 | 3.137728338 | -1.234830737 | 0.259842505 | 0.66338326 | -4.900403387 |
| **Q4KMQ1** | TPRN | -0.119001101 | 3.196582261 | -1.388577181 | 0.210873744 | 0.61590617 | -4.901722645 |
| **Q13769** | THOC5 | 0.032938121 | 3.194392075 | 2.673777745 | 0.034259998 | 0.345097821 | -4.902184225 |
| **Q9NS69** | TOMM22 | 0.165309821 | 3.15576082 | 1.669878474 | 0.142366056 | 0.538432214 | -4.90441116 |
| **P47985** | UQCRFS1 | 0.149621996 | 3.211583448 | 2.082662654 | 0.079041099 | 0.416709711 | -4.90495125 |
| **Q9Y240** | CLEC11A | 0.086422131 | 3.192206924 | 1.797505376 | 0.118776658 | 0.496164837 | -4.907088658 |
| **Q8TF32** | ZNF431 | 0.130373456 | 3.176634054 | 1.605169561 | 0.155975967 | 0.554834464 | -4.907102928 |
| **O43741** | LOC101060511; PF | -0.089754197 | 3.180200467 | -1.575258479 | 0.162673759 | 0.561580888 | -4.907897025 |
| **Q9UKV5** | AMFR | 0.106651117 | 3.147777562 | 2.046435523 | 0.083242306 | 0.425585663 | -4.908440079 |
| **Q96ER9** | CCDC51 | 0.164030323 | 3.195126237 | 2.135530045 | 0.073290412 | 0.405447658 | -4.911307106 |
| **Q9BWW8** | APOL6 | -0.290731072 | 3.237810118 | -1.377922606 | 0.213982049 | 0.619843797 | -4.911678874 |
| **Q9Y2R0** | COA3 | 0.167194967 | 3.173689518 | 1.632336194 | 0.150118537 | 0.547357402 | -4.911824833 |
| **O75179-1** | ANKRD17 | -0.065222334 | 3.198159527 | -2.665922356 | 0.034636308 | 0.345710389 | -4.912807959 |
| **Q96CD0** | FBXL8 | 0.155225105 | 3.116884168 | 1.408485 | 0.205174565 | 0.611215566 | -4.912811613 |
| **Q9UBC1-1** | NFKBIL1 | -0.143646057 | 3.192568656 | -1.493886548 | 0.18227823 | 0.584895208 | -4.913821119 |
| **Q9HC62** | SENP2 | 0.064493446 | 3.179019552 | 2.042488705 | 0.083713354 | 0.426895123 | -4.914788765 |
| **Q99807-1** | COQ7 | -0.159203052 | 3.027655463 | -1.605685374 | 0.155862771 | 0.554834464 | -4.914814006 |
| **Q9Y4F3-1** | KIAA0430; MARF1 | 0.040572451 | 3.148648311 | 2.491230436 | 0.044230793 | 0.357204257 | -4.915626476 |
| **P11177** | PDHB | 0.224686009 | 3.188348932 | 2.631293767 | 0.036347866 | 0.348943902 | -4.915786723 |
| **Q9H4G0-1** | EPB41L1 | 0.120233835 | 3.224903023 | 1.24312541 | 0.256967165 | 0.659782078 | -4.915878898 |
| **Q9BQE4** | VIMP; SELENOS | -0.07805434 | 3.209118498 | -2.110075859 | 0.076004475 | 0.411304347 | -4.916825237 |
| **Q86XN8-3** | MEX3D | 0.129816527 | 3.138167597 | 1.31984947 | 0.231654175 | 0.639059869 | -4.917030677 |
| **Q5SVZ6** | ZMYM1 | -0.045860015 | 3.202184411 | -2.450705037 | 0.046830752 | 0.359457245 | -4.923844307 |
| **Q658Y4** | FAM91A1 | 0.040734987 | 3.175750255 | 2.531975151 | 0.041768145 | 0.355540374 | -4.924515494 |
| **Q5JTC6-1** | AMER1; FAM123B | -0.162242486 | 3.269221008 | -1.021024006 | 0.343886666 | 0.725100733 | -4.92467149 |
| **Q9Y5Y2** | NUBP2 | 0.076508323 | 3.145934707 | 2.167063363 | 0.070063822 | 0.401998301 | -4.924981771 |
| **Q96RR1-1** | C10orf2; TWNK | 0.160142533 | 3.199743248 | 1.909174815 | 0.101287237 | 0.464354932 | -4.9250804 |
| **P31151** | S100A7 | -0.453256264 | 0.515420116 | -1.23173855 | 0.260921476 | 0.664371245 | -4.926283784 |
| **Q9H175** | CSRNP2 | -0.106088152 | 3.141414037 | -1.582201055 | 0.161095521 | 0.561414897 | -4.926519909 |
| **P56182** | RRP1 | -0.118410073 | 3.202581586 | -2.522753257 | 0.042312696 | 0.355540374 | -4.926745247 |
| **P07954** | FH | 0.091295769 | 3.179550758 | 2.715473802 | 0.032333536 | 0.339797349 | -4.927469715 |
| **Q8WUZ0-2** | BCL7C | -0.088823582 | 3.176939309 | -1.966417482 | 0.093331662 | 0.447307121 | -4.929680211 |
| **Q8WVF1-1** | OSCP1 | 0.142384149 | 3.15688288 | 1.235115819 | 0.259743224 | 0.663342949 | -4.931024889 |
| **Q9Y4C5-1** | CHST2 | -0.18974234 | 3.13515005 | -0.956345894 | 0.373247926 | 0.745838726 | -4.931037364 |
| **Q8N594-1** | MPND | -0.107961025 | 3.173261174 | -1.407345514 | 0.205496989 | 0.611717339 | -4.931451083 |

| | | | | | | | |
|---|---|---|---|---|---|---|---|
| **Q9UPG8** | PLAGL2 | -0.116407869 | 3.214184844 | -1.556784957 | 0.166944057 | 0.566424749 | -4.9331603 |
| **Q99871-2** | HAUS7 | -0.071784611 | 3.198901348 | -2.039996422 | 0.084012182 | 0.427385647 | -4.935511268 |
| **Q96K49-1** | TMEM87B | -0.085809825 | 3.149104749 | -1.82724307 | 0.113849814 | 0.48833912 | -4.936035142 |
| **Q8N8A6** | DDX51 | -0.041225034 | 3.182041413 | -2.585703977 | 0.038738012 | 0.352231442 | -4.937229916 |
| **Q8N3F8** | MICALL1 | -0.046426016 | 3.185154 | -2.149972057 | 0.071794362 | 0.403638325 | -4.938643852 |
| **Q68CQ7-1** | GLT8D1 | -0.108739942 | 3.180847288 | -1.848507728 | 0.110449643 | 0.481625912 | -4.938974338 |
| **P13747** | HLA-E | 0.058288398 | 3.163113985 | 2.297143488 | 0.058211397 | 0.379089796 | -4.939729283 |
| **Q7L8L6** | FASTKD5 | 0.16822885 | 3.189092969 | 2.469140031 | 0.04562884 | 0.357414656 | -4.939868325 |
| **P22626** | HNRNPA2B1 | 0.043776337 | 3.166614603 | 3.194574787 | 0.016883388 | 0.305857884 | -4.94010478 |
| **Q6RI45** | BRWD3 | -0.112339739 | 3.172495388 | -1.604015569 | 0.156229491 | 0.554834464 | -4.940272381 |
| **Q86X40** | LRRC28 | -0.137105854 | 3.134220026 | -1.098514782 | 0.311161435 | 0.69914525 | -4.94086206 |
| **Q53H12** | AGK | 0.209376303 | 3.186291416 | 2.565596381 | 0.039844119 | 0.355476279 | -4.942327837 |
| **Q6PI48** | DARS2 | 0.148505969 | 3.174977724 | 2.294708214 | 0.058413278 | 0.379161351 | -4.942856855 |
| **P00403** | COX2 | 0.216427296 | 3.139742942 | 2.149249664 | 0.071868454 | 0.403638325 | -4.943086725 |
| **Q16698** | DECR1 | 0.144349942 | 3.183154562 | 2.586831118 | 0.03867697 | 0.352231442 | -4.944663543 |
| **Q96EW2-1** | HSPBAP1 | 0.063661837 | 3.199184814 | 2.046458352 | 0.083239589 | 0.425585663 | -4.944945945 |
| **P78559-2** | MAP1A | -0.077354375 | 3.18094284 | -2.968272196 | 0.022863896 | 0.317241514 | -4.945350117 |
| **P61619** | SEC61A1 | -0.086557835 | 3.198701841 | -2.076758969 | 0.079710906 | 0.418547591 | -4.945730541 |
| **Q96NL6-1** | SCLT1 | -0.06785286 | 3.199450292 | -1.868549332 | 0.107336159 | 0.475884923 | -4.946505436 |
| **Q6UB35-1** | MTHFD1L | 0.165623654 | 3.18542477 | 2.704540146 | 0.032827364 | 0.339814312 | -4.946725274 |
| **O60885-1** | BRD4 | 0.023899544 | 3.180523288 | 2.72556776 | 0.031884605 | 0.33673269 | -4.948094769 |
| **Q92526-1** | CCT6B | 0.125123324 | 3.17305317 | 1.07006272 | 0.322870395 | 0.710272801 | -4.949768516 |
| **Q9NRK6** | ABCB10 | 0.130329543 | 3.166637048 | 2.005411028 | 0.088270984 | 0.437773138 | -4.951481216 |
| **O14647** | CHD2 | 0.055943367 | 3.166640975 | 2.497907589 | 0.043817059 | 0.357204257 | -4.951735793 |
| **Q86XL3-1** | ANKLE2 | -0.037902499 | 3.206106705 | -2.535623648 | 0.04155473 | 0.355540374 | -4.953623689 |
| **O75676** | RPS6KA4 | -0.046994901 | 3.218416822 | -2.166119629 | 0.070158264 | 0.401998301 | -4.954069387 |
| **Q5JWR5** | DOPEY1 | -0.112809424 | 3.195711719 | -1.506825391 | 0.179021168 | 0.580540072 | -4.954188819 |
| **Q9NZM4** | GLTSCR1; BICRA | 0.070496116 | 3.184045141 | 1.968327897 | 0.093077125 | 0.447295866 | -4.954741399 |
| **Q8TD16-2** | BICD2 | -0.067534845 | 3.203871222 | -2.517957209 | 0.04259883 | 0.355540374 | -4.955154005 |
| **Q9BYD3** | MRPL4 | 0.147830902 | 3.193359035 | 2.259875517 | 0.061380812 | 0.384167582 | -4.955166014 |
| **Q9BUB4** | ADAT1 | 0.078943812 | 3.168384401 | 1.995350213 | 0.089549904 | 0.440291106 | -4.956112843 |
| **Q8WUY9** | DEPDC1B | -0.096118549 | 3.149279577 | -2.10587892 | 0.076461634 | 0.412201353 | -4.957457374 |
| **P61764-2** | STXBP1 | -0.05684657 | 3.195403029 | -2.36739068 | 0.05268695 | 0.374840509 | -4.957766386 |
| **Q9Y450-4** | HBS1L | -0.047986032 | 3.186487357 | -2.79287907 | 0.029054733 | 0.333102237 | -4.95781145 |
| **Q96PN7** | TRERF1 | 0.092857454 | 3.196342856 | 1.603720971 | 0.156294275 | 0.554834464 | -4.958763036 |
| **Q02413** | DSG1 | -0.279357274 | 1.737194288 | -0.767350601 | 0.469936895 | 0.80622946 | -4.960487934 |
| **O60281-1** | ZNF292 | -0.056382979 | 3.197397121 | -2.305709765 | 0.057506945 | 0.377614679 | -4.960625918 |
| **P26440** | IVD | 0.122230688 | 3.187126901 | 2.540817348 | 0.041252895 | 0.355540374 | -4.96117587 |
| **Q9BPU9** | B9D2 | -0.124993416 | 3.161317534 | -1.148678459 | 0.291368853 | 0.685264687 | -4.962403153 |

| | | | | | | | |
|---|---|---|---|---|---|---|---|
| **Q12931** | TRAP1 | 0.161832181 | 3.187405473 | 2.670454951 | 0.034418647 | 0.345097821 | -4.963311446 |
| **Q13105-2** | ZBTB17 | 0.096733812 | 3.181277026 | 1.749937995 | 0.127090971 | 0.510226256 | -4.964731165 |
| **Q9NVH1-1** | DNAJC11 | 0.174920066 | 3.175871714 | 2.359493444 | 0.053280057 | 0.376021618 | -4.96537747 |
| **O75052** | NOS1AP | -0.151437247 | 3.243979587 | -1.275594693 | 0.245974797 | 0.651206185 | -4.967309592 |
| **Q5TB80** | KIAA1009; CEP162 | 0.094804205 | 3.158460435 | 1.403871992 | 0.206482666 | 0.612058044 | -4.969870663 |
| **Q8TF20-2** | ZNF721 | -0.169124399 | 3.241762773 | -1.165128192 | 0.285111441 | 0.681061455 | -4.969903749 |
| **Q16659** | MAPK6 | -0.127105404 | 3.216616939 | -0.951801355 | 0.375381958 | 0.748219118 | -4.970281616 |
| **P08559-4** | PDHA1 | 0.176891682 | 3.1704064 | 2.483448778 | 0.044718111 | 0.357204257 | -4.971298208 |
| **Q8WVD5-1** | RNF141 | 0.118143049 | 3.187681269 | 1.589623613 | 0.159424057 | 0.558776357 | -4.973220136 |
| **Q5SXM8** | DNLZ | 0.18165545 | 3.119494769 | 1.543736026 | 0.170023225 | 0.569040056 | -4.973612644 |
| **Q9P1F3** | ABRACL | -0.072755384 | 3.163390786 | -1.906549523 | 0.101667767 | 0.465562193 | -4.974215348 |
| **P59998-3** | ARPC4 | 0.082878881 | 3.193137742 | 2.243077384 | 0.062866706 | 0.387791908 | -4.977136024 |
| **Q9NVN8** | GNL3L | -0.072680167 | 3.240153137 | -2.547561721 | 0.040864362 | 0.355540374 | -4.97851627 |
| **Q9UL54-2** | TAOK2 | -0.231777152 | 3.174026823 | -1.356834822 | 0.220255313 | 0.625644783 | -4.978750799 |
| **Q9H974-4** | QTRTD1; QTRT2 | 0.099455019 | 3.174739344 | 2.134639128 | 0.07338373 | 0.405447658 | -4.98045497 |
| **Q71F56** | MED13L | 0.050039315 | 3.181845956 | 2.090025415 | 0.07821369 | 0.415568143 | -4.980661415 |
| **Q9HCN8** | SDF2L1 | 0.093537552 | 3.175246241 | 1.976610721 | 0.091981512 | 0.445605557 | -4.983118307 |
| **Q86XA9** | HEATR5A | 0.137330498 | 3.160602485 | 1.173464641 | 0.281983655 | 0.679261937 | -4.984291364 |
| **Q6IQ55** | TTBK2 | -0.116609791 | 3.200438057 | -0.839112889 | 0.431306846 | 0.782716512 | -4.984520978 |
| **P13498** | CYBA | -0.126008392 | 3.247206772 | -1.37347713 | 0.215291049 | 0.620866563 | -4.986164732 |
| **Q9BX95** | SGPP1 | 0.228464594 | 3.253668055 | 1.694973311 | 0.137398164 | 0.528633612 | -4.986768658 |
| **Q9NXC2-1** | GFOD1 | -0.400378396 | 3.166664964 | -0.90627127 | 0.39728021 | 0.762571436 | -4.98752409 |
| **Q9H9L4-1** | KANSL2 | 0.054561364 | 3.194014031 | 2.352571838 | 0.053805561 | 0.376021618 | -4.988319282 |
| **Q96BS2-1** | TESC | -0.062962643 | 3.204947889 | -2.205668841 | 0.066309276 | 0.394427892 | -4.990274264 |
| **P05386** | RPLP1 | -0.189541125 | 3.200530496 | -1.682154923 | 0.139914702 | 0.534948175 | -4.991167195 |
| **Q13356** | PPIL2 | -0.036188104 | 3.187339927 | -2.540320138 | 0.041281691 | 0.355540374 | -4.991839967 |
| **Q9NRC8** | SIRT7 | -0.137930847 | 3.189772408 | -0.902966965 | 0.398906211 | 0.762953742 | -4.991866218 |
| **Q9UPM8-1** | AP4E1 | -0.038103494 | 3.18657169 | -2.296535829 | 0.058261704 | 0.379106666 | -4.993320365 |
| **Q02086-1** | SP2 | 0.065929393 | 3.171237339 | 2.029152511 | 0.085324875 | 0.431338184 | -4.993597899 |
| **Q969U6** | FBXW5 | 0.106376504 | 3.209209911 | 1.345384718 | 0.223729806 | 0.629881158 | -4.994026401 |
| **Q9UHQ1-2** | NARF | -0.119180315 | 3.172826524 | -2.469929697 | 0.045578079 | 0.357414656 | -4.994118217 |
| **Q9NRZ9** | HELLS | -0.061060885 | 3.182489175 | -2.884101152 | 0.025638029 | 0.328538932 | -4.995897225 |
| **Q6PJP8** | DCLRE1A | 0.135430552 | 3.143810615 | 1.229257538 | 0.26178997 | 0.665372759 | -4.996641764 |
| **Q9Y2H0** | DLGAP4 | -0.060438735 | 3.177191145 | -2.292543376 | 0.058593344 | 0.379210276 | -4.996765201 |
| **Q8N584-1** | TTC39C | -0.067877782 | 3.195229772 | -1.807222855 | 0.117144284 | 0.493122332 | -4.996873606 |
| **Q9P0T4** | ZNF581 | -0.148404873 | 3.191459045 | -1.14865618 | 0.291377405 | 0.685264687 | -4.996956066 |
| **Q99707** | MTR | -0.062354921 | 3.182608864 | -2.612485717 | 0.037314576 | 0.350016887 | -4.998189895 |
| **O43251-8** | RBFOX2 | 0.098906177 | 3.154134325 | 1.663785833 | 0.143597777 | 0.539320158 | -4.999232232 |
| **Q9HA92** | RSAD1 | 0.162144996 | 3.156497545 | 1.762957206 | 0.124760951 | 0.507540068 | -5.000484559 |

| | | | | | | | |
|---|---|---|---|---|---|---|---|
| **O96005-1** | CLPTM1 | -0.048688588 | 3.169452066 | -2.329733547 | 0.055577752 | 0.376021618 | -5.000887889 |
| **Q96GM8** | TOE1 | -0.05924321 | 3.179030082 | -2.306248607 | 0.057462928 | 0.377614679 | -5.001179389 |
| **Q9BX79-4** | STRA6 | 0.107329733 | 3.198294578 | 1.231337386 | 0.261061738 | 0.664371245 | -5.002017865 |
| **Q9P2W9** | STX18 | -0.063721481 | 3.210797197 | -2.189479736 | 0.067858154 | 0.397403835 | -5.002814761 |
| **P62841** | RPS15 | -0.0881127 | 3.19745273 | -2.286650544 | 0.059086389 | 0.379928744 | -5.003051947 |
| **P35240-2** | NF2 | -0.063446222 | 3.186525616 | -2.315055131 | 0.056748417 | 0.377614679 | -5.005052313 |
| **Q8IY47** | KBTBD2 | 0.068802149 | 3.192873252 | 1.885059271 | 0.104836186 | 0.473790384 | -5.005078623 |
| **P35219** | CA8 | -0.076235318 | 3.195839038 | -1.89365625 | 0.10355718 | 0.469881093 | -5.005793007 |
| **Q86Y97** | SUV420H2; KMT5C | -0.125406576 | 3.180524618 | -1.149965856 | 0.290875018 | 0.685147353 | -5.006013358 |
| **Q9UIS9-9** | MBD1 | -0.114108553 | 3.219900155 | -1.985971893 | 0.090758706 | 0.443191118 | -5.006584128 |
| **P62993** | GRB2 | -0.032422552 | 3.197328447 | -2.707414915 | 0.032696755 | 0.339814312 | -5.006671206 |
| **P82921** | MRPS21 | 0.134591894 | 3.158074944 | 1.297496147 | 0.23879343 | 0.644491512 | -5.010088658 |
| **Q92624** | APPBP2 | -0.080636978 | 3.189085546 | -2.049339547 | 0.082897413 | 0.425319102 | -5.010750328 |
| **Q5VYK3** | KIAA0368 | 0.104543375 | 3.195986359 | 1.450790301 | 0.193523564 | 0.598629202 | -5.011363195 |
| **Q8IWU2** | LMTK2 | 0.044174291 | 3.199088878 | 2.353215107 | 0.053756498 | 0.376021618 | -5.011374102 |
| **P50452** | SERPINB8 | 0.085769315 | 3.141520172 | 1.858671423 | 0.108859793 | 0.479429633 | -5.012254656 |
| **O60266** | ADCY3 | 0.13233627 | 3.207282829 | 0.825496207 | 0.438456908 | 0.786897076 | -5.012545106 |
| **Q6IQ23-2** | PLEKHA7 | -0.085265767 | 3.191364032 | -2.266434954 | 0.060810399 | 0.383378328 | -5.01265614 |
| **Q8N720-2** | ZNF655 | 0.12504657 | 3.139690102 | 1.14763931 | 0.291767973 | 0.685264687 | -5.013144256 |
| **Q15051-1** | IQCB1 | -0.08009471 | 3.176206849 | -2.121184672 | 0.07480771 | 0.407645581 | -5.01339336 |
| **P34949** | MPI | 0.063592758 | 3.165622042 | 2.336943513 | 0.055011867 | 0.376021618 | -5.014910745 |
| **Q52LA3** | LIN52 | -0.128177965 | 3.190622867 | -1.364789097 | 0.217870004 | 0.624306497 | -5.01567035 |
| **Q5SWA1** | PPP1R15B | -0.10531828 | 3.258152922 | -1.24364415 | 0.25678826 | 0.659782078 | -5.017125555 |
| **Q9H2G4** | TSPYL2 | 0.108637511 | 3.198644964 | 1.204660742 | 0.270535185 | 0.671731232 | -5.01796966 |
| **Q14847** | LASP1 | 0.064621209 | 3.19604134 | 2.499085727 | 0.043744478 | 0.357204257 | -5.018872042 |
| **Q7Z4V5** | HDGFRP2; HDGFL2 | 0.04107368 | 3.157964606 | 2.52377081 | 0.042252247 | 0.355540374 | -5.019067708 |
| **Q8WUJ0** | STYX | -0.065482655 | 3.215204259 | -1.825342978 | 0.114158565 | 0.48833912 | -5.019216372 |
| **P57105** | SYNJ2BP | 0.125478992 | 3.165922131 | 2.025987908 | 0.085711819 | 0.431546517 | -5.020337063 |
| **Q6PL24** | TMED8 | -0.078086885 | 3.199447948 | -2.076301744 | 0.079763018 | 0.418547591 | -5.021209181 |
| **P35568** | IRS1 | 0.155815021 | 2.898530377 | 0.9185721 | 0.391270921 | 0.759806823 | -5.022067921 |
| **P60983** | GMFB | -0.075507995 | 3.179421783 | -1.996214565 | 0.089439306 | 0.440291106 | -5.022073286 |
| **Q9UBT2** | UBA2 | 0.075149248 | 3.170707976 | 2.789295947 | 0.029198453 | 0.333307048 | -5.023762827 |
| **Q15652** | JMJD1C | 0.047098435 | 3.149814951 | 2.582690381 | 0.038901715 | 0.352501702 | -5.024369263 |
| **Q86WC4** | OSTM1 | -0.069789302 | 3.211632016 | -1.784320584 | 0.121026873 | 0.499487166 | -5.024818955 |
| **P61077-3** | UBE2D3 | -0.057361686 | 3.187809381 | -2.237017542 | 0.063411725 | 0.389084368 | -5.024892409 |
| **P30281-1** | CCND3 | -0.100499428 | 3.124023451 | -1.481146523 | 0.185538488 | 0.588699395 | -5.025700986 |
| **Q9Y584** | TIMM22 | 0.153789169 | 3.217012616 | 1.028303308 | 0.340699622 | 0.722995142 | -5.02586148 |
| **O43837-1** | IDH3B | 0.179863678 | 3.194135295 | 2.555458184 | 0.040414322 | 0.355540374 | -5.02649581 |
| **Q9ULD0** | OGDHL | 0.172286587 | 3.166590433 | 2.05005226 | 0.082812987 | 0.425319102 | -5.026518178 |

| | | | | | | | |
|---|---|---|---|---|---|---|---|
| **Q4V328** | GRIPAP1 | 0.038178333 | 3.174613582 | 2.423057808 | 0.048695333 | 0.364544514 | -5.026699247 |
| **Q9NUJ7** | PLCXD1 | 0.119667661 | 3.213480536 | 1.34697624 | 0.223243965 | 0.629408553 | -5.02794181 |
| **P25325-2** | MPST | 0.091173859 | 3.173611613 | 1.669786106 | 0.142384655 | 0.538432214 | -5.028196558 |
| **Q9HAP2-1** | MLXIP | 0.064287005 | 3.199758832 | 1.873757755 | 0.106541227 | 0.475403904 | -5.028557383 |
| **P30048** | PRDX3 | 0.181139832 | 3.169697941 | 2.616598625 | 0.037100891 | 0.349573768 | -5.028936982 |
| **P16220-1** | CREB1 | -0.060780575 | 3.174434703 | -1.855567396 | 0.109342945 | 0.479873319 | -5.029002578 |
| **Q9NV96** | TMEM30A | -0.073173148 | 3.166770089 | -2.160481433 | 0.070725191 | 0.402227373 | -5.030049695 |
| **Q8NDZ4** | C3orf58 | -0.078593095 | 3.190089399 | -1.882880854 | 0.105162738 | 0.474456534 | -5.031332053 |
| **P40926** | MDH2 | 0.174313009 | 3.170136487 | 2.83177364 | 0.027542142 | 0.32997678 | -5.032129176 |
| **P0CK96** | SLC35E2B | -0.122963015 | 3.155398894 | -1.479628884 | 0.185930414 | 0.588813889 | -5.032597049 |
| **Q86UY8** | NT5DC3 | 0.128806261 | 3.19095602 | 2.049262143 | 0.082906587 | 0.425319102 | -5.033877141 |
| **Q96NC0** | ZMAT2 | 0.096213212 | 3.200976824 | 2.389981878 | 0.051027683 | 0.370580387 | -5.036057799 |
| **O94876** | TMCC1 | -0.144179337 | 3.177347256 | -1.146243371 | 0.292304849 | 0.685264687 | -5.036338131 |
| **Q9NUE0** | ZDHHC18 | -0.097999893 | 3.189956138 | -1.150797196 | 0.290556495 | 0.684803131 | -5.036427046 |
| **O75467** | ZNF324 | -0.069631764 | 3.184061213 | -1.683849835 | 0.139579444 | 0.533923294 | -5.03651363 |
| **Q6UVJ0** | SASS6 | -0.060673513 | 3.179543759 | -2.241394652 | 0.063017568 | 0.387825609 | -5.038159956 |
| **Q8WU39-1** | MZB1 | -0.073644242 | 3.191994093 | -2.482944148 | 0.044749905 | 0.357204257 | -5.03950916 |
| **P56211** | ARPP19 | -0.196923408 | 3.17095129 | -2.004481212 | 0.088388412 | 0.437773138 | -5.041562724 |
| **Q03936** | ZNF92 | -0.090492351 | 3.147308749 | -1.274351915 | 0.246387891 | 0.6518654 | -5.042412828 |
| **O14949** | UQCRQ | 0.175544449 | 3.206388248 | 2.196828723 | 0.06715053 | 0.394900782 | -5.044979799 |
| **Q9Y262** | EIF3L | -0.042151672 | 3.198710527 | -2.849211453 | 0.026891394 | 0.328952544 | -5.046537592 |
| **P55160** | NCKAP1L | -0.0454014 | 3.177840313 | -2.77432494 | 0.029807157 | 0.335282407 | -5.046989829 |
| **Q9BQB6-1** | VKORC1 | -0.085258767 | 3.213964317 | -1.618297262 | 0.153119055 | 0.552031202 | -5.048486674 |
| **Q86Y39-2** | NDUFA11 | 0.122617695 | 3.212958126 | 1.573299301 | 0.163121748 | 0.561580888 | -5.048531527 |
| **O14548** | COX7A2L | 0.101057719 | 3.195629896 | 1.972403601 | 0.092536397 | 0.446659583 | -5.048856581 |
| **Q9H5H4** | ZNF768 | 0.113883067 | 3.193547087 | 0.937845224 | 0.381994092 | 0.753274524 | -5.04900578 |
| **Q99766-1** | ATP5S | 0.156795351 | 3.190605287 | 1.921761541 | 0.09948227 | 0.459999988 | -5.049402537 |
| **Q6V9R5-1** | ZNF562 | -0.095354539 | 3.233969926 | -1.638558753 | 0.148806469 | 0.546231571 | -5.04963916 |
| **Q13325** | IFIT5 | -0.055374018 | 3.199523343 | -2.090161146 | 0.078198519 | 0.415568143 | -5.05189101 |
| **O60674** | JAK2 | 0.102347379 | 3.185648184 | 1.327524278 | 0.229246737 | 0.637286678 | -5.053501213 |
| **Q8WV16-1** | DCAF4 | -0.082895262 | 3.238397729 | -1.963864034 | 0.093672955 | 0.448493945 | -5.054111093 |
| **Q8N3Z3-1** | GTPBP8 | 0.087044458 | 3.252774898 | 1.508945932 | 0.17849252 | 0.57989621 | -5.055334555 |
| **Q9NXS2** | QPCTL | -0.126191123 | 3.139462187 | -1.010098854 | 0.348714328 | 0.730176717 | -5.055609898 |
| **Q13098-7** | GPS1 | 0.056049446 | 3.205559965 | 2.527512406 | 0.042030749 | 0.355540374 | -5.056104543 |
| **P17026** | ZNF22 | -0.074182551 | 3.195608934 | -1.901580031 | 0.102391948 | 0.467531052 | -5.056653136 |
| **P38646** | HSPA9 | 0.162679426 | 3.187919455 | 3.027269943 | 0.02111192 | 0.307875291 | -5.057378702 |
| **Q9BYG3** | MKI67IP; NIFK | -0.114468208 | 3.198961273 | -2.343162292 | 0.05452855 | 0.376021618 | -5.058338652 |
| **Q93063-3** | EXT2 | 0.074034934 | 3.208982531 | 1.579190155 | 0.161778211 | 0.561414897 | -5.058518455 |
| **Q6DCA0** | AMMECR1L | -0.095003719 | 3.109551573 | -0.514441801 | 0.624037462 | 0.885078428 | -5.059138086 |

| | | | | | | | |
|---|---|---|---|---|---|---|---|
| **Q8NDZ2-1** | SIMC1 | -0.085284973 | 3.238740522 | -1.648056282 | 0.146824784 | 0.545761036 | -5.059459539 |
| **Q8NBU5** | ATAD1 | 0.07945271 | 3.168649212 | 2.288039924 | 0.058969759 | 0.379928744 | -5.061398194 |
| **Q9H2U2** | PPA2 | 0.141383208 | 3.168789597 | 2.644108513 | 0.035704324 | 0.347635849 | -5.061964563 |
| **Q9NRP2** | CMC2 | 0.101092674 | 3.15319125 | 1.104274374 | 0.308834002 | 0.696978291 | -5.0619739 |
| **Q96BW5-1** | PTER | 0.049808791 | 3.167831636 | 2.287436915 | 0.059020349 | 0.379928744 | -5.063026737 |
| **Q5SR56** | HIATL1; MFSD14B | 0.12500127 | 3.085624322 | 1.15605978 | 0.288546936 | 0.683513836 | -5.063701338 |
| **P27815-1** | PDE4A | 0.108067271 | 3.147057181 | 0.985239692 | 0.359898557 | 0.73661838 | -5.06447691 |
| **O00764-1** | PDXK | -0.060911225 | 3.202237176 | -2.013202302 | 0.087293111 | 0.435855266 | -5.065122885 |
| **Q16539-2** | MAPK14 | -0.121646001 | 3.179955067 | -1.363259503 | 0.218326896 | 0.625083672 | -5.065137125 |
| **Q92546** | RGP1 | -0.121371914 | 3.201445024 | -0.736499019 | 0.487259323 | 0.816334529 | -5.067051081 |
| **P17024** | ZNF20 | -0.094215992 | 3.205825606 | -1.17566307 | 0.281163654 | 0.678447235 | -5.067090949 |
| **Q6P9F7** | LRRC8B | -0.11142292 | 3.15674116 | -1.036715685 | 0.337045868 | 0.721128781 | -5.067280692 |
| **Q9H1J1-1** | UPF3A | 0.108182145 | 3.178784257 | 0.896963096 | 0.401873379 | 0.765129163 | -5.067656322 |
| **O75420** | GIGYF1 | 0.101824115 | 3.02299639 | 1.95608385 | 0.094720518 | 0.450361767 | -5.069204554 |
| **P22830-2** | FECH | 0.213837614 | 3.210348734 | 2.270624885 | 0.060448887 | 0.383145812 | -5.070551354 |
| **Q9H9S5** | FKRP | -0.106394228 | 3.232320215 | -0.62141819 | 0.55557103 | 0.84999265 | -5.07092718 |
| **O15523** | DDX3Y | -0.060039844 | 3.164877179 | -1.936954835 | 0.097345839 | 0.45575291 | -5.070978868 |
| **O75081** | CBFA2T3 | -0.090809333 | 3.250929238 | -1.777309191 | 0.122240286 | 0.501393428 | -5.071426595 |
| **Q92685-1** | ALG3 | 0.091071507 | 3.178842527 | 1.653429984 | 0.145714666 | 0.543674934 | -5.071538396 |
| **Q8IWZ3-6** | ANKHD1-EIF4EBP3 | -0.09677004 | 3.199658321 | -1.605595035 | 0.15588259 | 0.554834464 | -5.072236534 |
| **Q15633** | TARBP2 | -0.052989298 | 3.170899521 | -1.923026755 | 0.099302603 | 0.459766423 | -5.073025725 |
| **Q5VYS8-1** | ZCCHC6 | -0.042664058 | 3.174639476 | -2.376910481 | 0.051981043 | 0.373408128 | -5.07475137 |
| **Q8IY92** | SLX4 | -0.120318395 | 3.175229095 | -1.375127127 | 0.214804362 | 0.619913059 | -5.075370828 |
| **Q5R372-5** | RABGAP1L | 0.086694776 | 3.196094074 | 1.291472807 | 0.240749927 | 0.646357456 | -5.076309382 |
| **Q9BRJ2** | MRPL45 | 0.152003505 | 3.202310738 | 2.264883188 | 0.060944849 | 0.383378328 | -5.076545646 |
| **Q8WTV1** | THAP3 | -0.102599486 | 3.189267278 | -1.116881308 | 0.30378956 | 0.692399426 | -5.076595475 |
| **P23771-2** | GATA3 | 0.061537546 | 3.201903766 | 1.781803158 | 0.121461196 | 0.499487166 | -5.076733107 |
| **Q16549** | PCSK7 | 0.110847537 | 3.199870058 | 1.074013306 | 0.321223469 | 0.709119329 | -5.078057562 |
| **I3L1I5** | | 0.126695222 | 3.213850915 | 1.618703302 | 0.153031482 | 0.552031202 | -5.079735904 |
| **Q9Y3U8** | RPL36 | -0.110075892 | 3.179213623 | -2.34039815 | 0.054742833 | 0.376021618 | -5.080008814 |
| **P20963-1** | CD247 | -0.114981205 | 3.223380906 | -2.097314295 | 0.077403183 | 0.414399115 | -5.080461532 |
| **Q9NWW7** | C2orf42 | 0.107594921 | 3.241080525 | 1.326223426 | 0.229653226 | 0.637461728 | -5.081908645 |
| **P35443** | THBS4 | 0.122630941 | 3.225042302 | 0.829414913 | 0.436390552 | 0.785483221 | -5.082005818 |
| **Q9ULW3** | ABT1 | -0.05746589 | 3.173239379 | -2.089707765 | 0.078249206 | 0.415568143 | -5.083336266 |
| **Q9NSG2-1** | C1orf112 | -0.071324696 | 3.194835473 | -1.881456017 | 0.105376868 | 0.475110045 | -5.084685833 |
| **Q6P1L8** | MRPL14 | 0.163705426 | 3.192786978 | 1.75416569 | 0.126329778 | 0.509218658 | -5.084764215 |
| **A9UHW6-3** | MIF4GD | -0.075081676 | 3.197158115 | -2.078006537 | 0.079568887 | 0.418381741 | -5.085529037 |
| **Q9Y6X8** | ZHX2 | 0.054490572 | 3.184708796 | 2.132480829 | 0.073610297 | 0.405653949 | -5.085672565 |
| **Q49AG3** | ZBED5 | 0.067394247 | 3.159037141 | 1.636387467 | 0.149263061 | 0.546464774 | -5.085884366 |

| | | | | | | | |
|---|---|---|---|---|---|---|---|
| **Q9NQL2-1** | RRAGD | 0.095984764 | 3.239855608 | 1.301577428 | 0.237475689 | 0.644090219 | -5.086118939 |
| **Q14520-1** | HABP2 | 0.078187388 | 3.239076949 | 1.871116494 | 0.106943622 | 0.475471224 | -5.086239584 |
| **P51178-2** | PLCD1 | 0.113993175 | 3.161809847 | 0.947106894 | 0.377596227 | 0.749625692 | -5.086593544 |
| **P36776** | LONP1 | 0.146599099 | 3.180740291 | 2.678922027 | 0.034015902 | 0.345097821 | -5.088156846 |
| **Q9NW82** | WDR70 | -0.037955515 | 3.174970913 | -2.484916755 | 0.044625754 | 0.357204257 | -5.088608791 |
| **Q9UBQ5** | EIF3K | -0.061210454 | 3.193046704 | -2.336068552 | 0.055080222 | 0.376021618 | -5.08957346 |
| **Q13546** | RIPK1 | -0.038162816 | 3.196978999 | -2.428103043 | 0.048349407 | 0.363765191 | -5.089884174 |
| **P67936** | TPM4 | -0.0567083 | 3.187130403 | -2.329025237 | 0.055633668 | 0.376021618 | -5.089928307 |
| **Q53H54** | TRMT12 | -0.078309718 | 3.201504229 | -1.553476869 | 0.167719713 | 0.566424749 | -5.09013166 |
| **Q9H0U4** | RAB1B | 0.144111647 | 3.189353049 | 2.05954122 | 0.081697153 | 0.423686898 | -5.090737283 |
| **Q9Y3A4** | RRP7A | -0.063150111 | 3.194211939 | -2.042757837 | 0.083681149 | 0.426895123 | -5.091107302 |
| **Q6PD74** | AAGAB | -0.065234 | 3.214404149 | -1.718474145 | 0.132896534 | 0.519961998 | -5.091135652 |
| **Q9NS86** | LANCL2 | 0.072008943 | 3.160456381 | 1.962769523 | 0.093819627 | 0.448493945 | -5.091565676 |
| **Q96NL8** | C8orf37 | -0.087997995 | 3.166727215 | -1.386680317 | 0.21142414 | 0.61590617 | -5.094136207 |
| **Q16739** | UGCG | -0.112357388 | 3.187206869 | -1.240111951 | 0.25800859 | 0.660717666 | -5.094576551 |
| **O75489** | NDUFS3 | 0.182735385 | 3.201161715 | 2.109193449 | 0.076100364 | 0.411304347 | -5.094795643 |
| **Q9ULI3** | HEG1 | 0.104809886 | 3.2239091 | 0.705017728 | 0.505372701 | 0.829070021 | -5.096745092 |
| **Q9Y3P8** | SIT 1.00 | -0.099124322 | 3.208139542 | -1.26912535 | 0.248131799 | 0.652351802 | -5.096876879 |
| **Q9Y2K2** | SIK3 | -0.123111382 | 3.228133434 | -1.467427883 | 0.189108996 | 0.593158693 | -5.097139199 |
| **P51149** | RAB7A | -0.03893745 | 3.170462916 | -2.622828887 | 0.036779649 | 0.348943902 | -5.097614767 |
| **Q9BYV9** | BACH2 | -0.067897869 | 3.109973574 | -1.895042126 | 0.103352441 | 0.469881093 | -5.098479107 |
| **Q9BTK6** | PAGR1 | -0.219692769 | 3.171382482 | -1.688666626 | 0.138630853 | 0.532345155 | -5.100892732 |
| **Q0P651-1** | C4orf29; ABHD18 | 0.102889436 | 3.185035653 | 1.247742251 | 0.255378664 | 0.659782078 | -5.100935988 |
| **P42701** | IL12RB1 | -0.205498353 | 3.218831785 | -0.983021548 | 0.360910013 | 0.736749757 | -5.101333834 |
| **Q9H7Z7** | PTGES2 | 0.10886169 | 3.182831351 | 2.352661013 | 0.053798757 | 0.376021618 | -5.101400181 |
| **Q9ULC8-1** | ZDHHC8 | 0.094280602 | 3.247136545 | 0.529850936 | 0.613897125 | 0.879837049 | -5.101764321 |
| **Q8N0Z3** | SPICE1 | -0.068468677 | 3.202626932 | -2.017882463 | 0.086710923 | 0.434648757 | -5.102305835 |
| **Q9BQQ3-1** | GORASP1 | 0.071703155 | 3.209989574 | 1.678379092 | 0.140664338 | 0.53619047 | -5.102487979 |
| **Q8N8R3-1** | SLC25A29 | 0.159817848 | 3.228185893 | 1.460947097 | 0.190817526 | 0.595227814 | -5.102489317 |
| **Q68CJ6** | NUGGC | 0.074675484 | 3.167623551 | 1.668450539 | 0.142653829 | 0.53919347 | -5.104230081 |
| **Q96QD9-1** | FYTTD1 | -0.105759323 | 3.121964011 | -1.782616688 | 0.121320676 | 0.499487166 | -5.105629716 |
| **Q9UDR5** | AASS | 0.18807536 | 3.184353882 | 2.405422003 | 0.049924839 | 0.36851891 | -5.106078512 |
| **Q6UWW8** | CES3 | 0.086421329 | 3.196480411 | 1.78262913 | 0.121318528 | 0.499487166 | -5.106089592 |
| **P29017** | CD1C | 0.055423689 | 3.20004512 | 1.926195335 | 0.098854055 | 0.458056247 | -5.107617255 |
| **Q9GZY4** | COA1 | 0.118651869 | 3.186591218 | 1.394087845 | 0.209282061 | 0.614302382 | -5.108843412 |
| **A7KAX9** | ARHGAP32 | -0.087810279 | 3.186447708 | -1.93120258 | 0.098149307 | 0.456822639 | -5.110997479 |
| **Q99501** | GAS2L1 | -0.094955659 | 3.144093476 | -0.738791039 | 0.485957759 | 0.81473183 | -5.111172012 |
| **O43291** | SPINT2 | 0.054199964 | 3.144731934 | 2.166772231 | 0.070092943 | 0.401998301 | -5.112020975 |
| **Q10713** | PMPCA | 0.180344038 | 3.178295413 | 2.405105754 | 0.049947178 | 0.36851891 | -5.112554671 |

| | | | | | | | |
|---|---|---|---|---|---|---|---|
| **Q13308-6** | PTK7 | -0.039125338 | 3.160408819 | -2.481211413 | 0.044859257 | 0.357204257 | -5.113199915 |
| **Q8N6N3** | C1orf52 | -0.266935668 | 3.187942716 | -1.811843307 | 0.116375805 | 0.492686484 | -5.113262943 |
| **P62917** | RPL8 | -0.088766302 | 3.173192525 | -2.397107832 | 0.050515594 | 0.368977419 | -5.113447855 |
| **Q6RW13** | AGTRAP | 0.094788033 | 3.210129018 | 1.357593332 | 0.220026852 | 0.625644783 | -5.114111743 |
| **Q96A26** | FAM162A | 0.208477611 | 3.207934902 | 2.287502386 | 0.059014854 | 0.379928744 | -5.115645253 |
| **Q9NXG0-2** | CNTLN | 0.052352955 | 3.245888949 | 0.309402612 | 0.76673769 | 0.940879515 | -5.11606684 |
| **P17098** | ZNF8 | -0.051906412 | 3.173018708 | -1.999062573 | 0.089075854 | 0.440116703 | -5.119023565 |
| **Q96KN1** | FAM84B | 0.071837906 | 3.193157473 | 1.463910712 | 0.190034483 | 0.593951206 | -5.120829669 |
| **P52746** | ZNF142 | -0.079190969 | 3.224417637 | -1.567158318 | 0.164533446 | 0.563893388 | -5.12210682 |
| **P52435** | POLR2J | -0.057318852 | 3.206516977 | -2.144023872 | 0.07240675 | 0.404094866 | -5.122375069 |
| **Q9Y2B1** | TMEM5 | -0.07970101 | 3.19601113 | -1.466396181 | 0.189380043 | 0.59346782 | -5.123864453 |
| **Q13895** | BYSL | 0.053298901 | 3.189486157 | 2.487500816 | 0.044463662 | 0.357204257 | -5.124324623 |
| **P36897-2** | TGFBR1 | -0.088396806 | 3.164501445 | -1.213086561 | 0.26751173 | 0.67010832 | -5.124539235 |
| **P61604** | HSPE1 | 0.161221168 | 3.199275599 | 2.768842545 | 0.030033422 | 0.336035644 | -5.124865176 |
| **P46777** | RPL5 | -0.102203803 | 3.172706938 | -2.845390499 | 0.027032571 | 0.328952544 | -5.125428577 |
| **Q08357** | SLC20A2 | 0.070925484 | 3.188080775 | 1.665049984 | 0.143341378 | 0.539313633 | -5.125743542 |
| **Q9NY12-1** | GAR1 | -0.094440358 | 3.136556962 | -2.145788925 | 0.072224481 | 0.404094866 | -5.126882066 |
| **P13598** | ICAM2 | -0.071111581 | 3.126581401 | -1.574803066 | 0.162777792 | 0.561580888 | -5.127317675 |
| **Q8N1E6** | FBXL14 | 0.117627765 | 3.199829222 | 1.158063308 | 0.287784941 | 0.682522792 | -5.127491357 |
| **Q96FH0-1** | MEF2BNB; BORCS | 0.095360711 | 3.199957102 | 1.096021479 | 0.312173427 | 0.699835744 | -5.127835326 |
| **Q6UWB1** | IL27RA | 0.089769011 | 3.234675267 | 0.917089511 | 0.39199155 | 0.759806823 | -5.127939193 |
| **Q5BKX6-1** | SLC45A4 | -0.100093107 | 3.235480676 | -1.122850296 | 0.301425019 | 0.691344519 | -5.127958958 |
| **P25686-3** | DNAJB2 | -0.077614665 | 3.240112154 | -0.420566113 | 0.687683122 | 0.907431059 | -5.128907724 |
| **Q8IYB8** | SUPV3L1 | 0.142702524 | 3.184502409 | 2.368009321 | 0.052640777 | 0.374840509 | -5.12912505 |
| **Q6GPH6-2** | ITPRIPL1 | -0.080707473 | 3.220268303 | -1.595670534 | 0.15807442 | 0.556829942 | -5.129595928 |
| **Q96QE5** | TEFM | 0.105582259 | 3.197153109 | 1.794055256 | 0.119361524 | 0.498071065 | -5.131075455 |
| **Q96HY6-1** | DDRGK1 | 0.126302702 | 3.162251423 | 1.619399798 | 0.152881376 | 0.552031202 | -5.131209394 |
| **Q9Y6C9** | MTCH2 | 0.130165178 | 3.195270477 | 2.467979209 | 0.045703567 | 0.357414656 | -5.13124313 |
| **Q9NP73-1** | ALG13 | 0.102673619 | 3.185230195 | 0.52408699 | 0.617679633 | 0.881750816 | -5.131686584 |
| **O95948** | ONECUT2 | 0.098377816 | 3.182006816 | 0.933610072 | 0.384018126 | 0.754456977 | -5.13392556 |
| **P01730** | CD4 | -0.074134206 | 3.213814731 | -1.723237327 | 0.132001548 | 0.519814431 | -5.135248922 |
| **Q9UJ68-1** | MSRA | 0.093842218 | 3.214627822 | 1.35897786 | 0.219610382 | 0.625602181 | -5.137920507 |
| **P09012** | SNRPA | 0.088020239 | 3.167408386 | 2.199747871 | 0.066871538 | 0.394427892 | -5.141048663 |
| **P30050-1** | RPL12 | -0.051478194 | 3.187668836 | -2.74394155 | 0.031084271 | 0.33673269 | -5.141766072 |
| **Q9NUQ2** | AGPAT5 | 0.086717564 | 3.242275705 | 1.832512568 | 0.112997818 | 0.486858819 | -5.142229974 |
| **P05026** | ATP1B1 | -0.081061296 | 3.197905766 | -1.59274274 | 0.15872654 | 0.557345662 | -5.142251492 |
| **O60673** | REV3L | -0.074488976 | 3.205398088 | -1.499638111 | 0.180823715 | 0.583290466 | -5.144877192 |
| **Q02818** | NUCB1 | -0.25809051 | 3.154633793 | -2.01919974 | 0.086547761 | 0.434379005 | -5.145165075 |
| **O75369-8** | FLNB | 0.089521205 | 3.173692095 | 1.180291689 | 0.279443787 | 0.676725357 | -5.148721375 |

| | | | | | | | |
|---|---|---|---|---|---|---|---|
| **Q8TEK3-2** | DOT1L | -0.05445067 | 3.206581009 | -2.304731467 | 0.057586951 | 0.377810342 | -5.149318871 |
| **Q16718** | NDUFA5 | 0.226324886 | 3.183265789 | 1.817747257 | 0.115400998 | 0.491192304 | -5.149520192 |
| **O43529** | CHST10 | -0.080909443 | 3.214978225 | -1.449791306 | 0.193791606 | 0.598629202 | -5.150621039 |
| **P63098** | PPP3R1 | -0.157760709 | 3.202717481 | -1.861761412 | 0.108380906 | 0.478381277 | -5.152059871 |
| **O96028-1** | WHSC1; NSD2 | -0.0782892 | 3.179732273 | -2.527938812 | 0.042005583 | 0.355540374 | -5.152424646 |
| **Q96KS0-1** | EGLN2; RAB4B-EG | -0.088523724 | 3.193456884 | -0.983417878 | 0.360729127 | 0.736666658 | -5.152696321 |
| **Q6UXV4** | APOOL | 0.204979206 | 3.199497852 | 1.95728394 | 0.094558176 | 0.450024955 | -5.154204738 |
| **O14709** | ZNF197; ZNF660-Z | -0.046564842 | 3.122095359 | -0.312755567 | 0.76430227 | 0.940369694 | -5.154237223 |
| **Q9Y5W8** | SNX13 | 0.081593301 | 3.172339294 | 1.397528262 | 0.208293832 | 0.614231901 | -5.154918485 |
| **Q9H939-2** | PSTPIP2 | -0.059289888 | 3.097777432 | -0.216571742 | 0.83520957 | 0.960912428 | -5.155293369 |
| **P17010-3** | ZFX | 0.110322823 | 3.176169671 | 1.082336085 | 0.317776206 | 0.705887314 | -5.155738168 |
| **Q9NYL9** | TMOD3 | -0.029937069 | 3.186856975 | -2.469210353 | 0.045624318 | 0.357414656 | -5.156203206 |
| **Q9C0H5-2** | ARHGAP39 | 0.093930196 | 3.275280792 | 0.936748415 | 0.382517488 | 0.754089677 | -5.156317017 |
| **Q16520** | BATF | -0.112876295 | 3.190483994 | -0.680196132 | 0.519962132 | 0.834396917 | -5.156529018 |
| **Q8NHY2-1** | RFWD2 | 0.04719838 | 3.16168704 | 1.971199703 | 0.092695793 | 0.44675374 | -5.157349548 |
| **O00541-1** | PES 1.00 | -0.073598148 | 3.196681363 | -2.532633318 | 0.041729562 | 0.355540374 | -5.15747527 |
| **Q96J84-2** | KIRREL; KIRREL1 | -0.070982935 | 3.170478826 | -1.490255944 | 0.183201915 | 0.585730065 | -5.158463449 |
| **Q14493-1** | SLBP | -0.05644649 | 3.204035846 | -1.916546675 | 0.100226206 | 0.461621569 | -5.159131812 |
| **Q16637** | SMN1; SMN2 | 0.113057088 | 3.16730372 | 2.270334573 | 0.060473864 | 0.383145812 | -5.159805007 |
| **Q7Z460-3** | CLASP1 | 0.105726229 | 3.162479804 | 1.033136954 | 0.338596376 | 0.721920542 | -5.159806207 |
| **Q5TA50** | GLTPD1; CPTP | 0.091932822 | 3.190662904 | 1.505114318 | 0.179448789 | 0.580990067 | -5.159976301 |
| **Q9NSA3** | CTNNBIP1 | -0.435230266 | 3.174680896 | -1.130703666 | 0.298337248 | 0.689670498 | -5.16128712 |
| **O75164-1** | KDM4A | -0.042950414 | 3.141074 | -2.121564955 | 0.074767081 | 0.407645581 | -5.162308594 |
| **Q9BUW7** | C9orf16 | -0.149870791 | 3.245754209 | -0.680404105 | 0.519838773 | 0.834367486 | -5.163627977 |
| **P31944** | CASP14 | -0.143735164 | 1.709210016 | -0.631582655 | 0.549308093 | 0.846638465 | -5.165281563 |
| **Q562R1** | ACTBL2 | 0.127759096 | 3.220760598 | 1.356062433 | 0.220488171 | 0.625865114 | -5.165657073 |
| **Q9H078-2** | CLPB | 0.122497799 | 3.168036588 | 2.312287137 | 0.056972006 | 0.377614679 | -5.167666717 |
| **O43169** | CYB5B | 0.065443896 | 3.18626082 | 2.280396975 | 0.059614277 | 0.381343177 | -5.170298745 |
| **Q86V81** | ALYREF | -0.058119208 | 3.171344758 | -2.446737769 | 0.047093686 | 0.360340657 | -5.170824025 |
| **Q6NYC1-3** | JMJD6 | -0.053135929 | 3.210318801 | -2.153303667 | 0.071453657 | 0.403638325 | -5.170862899 |
| **P52757-9** | CHN2 | 0.084271767 | 3.188463316 | 1.633381872 | 0.149897286 | 0.546801624 | -5.171517219 |
| **P26368** | U2AF2 | 0.038726126 | 3.185779173 | 2.525437005 | 0.04215346 | 0.355540374 | -5.172371981 |
| **Q8NDN9** | RCBTB1 | -0.053946569 | 3.186603205 | -1.820505758 | 0.114948272 | 0.489727184 | -5.172731301 |
| **Q96A19** | CCDC102A | -0.088873375 | 3.27085024 | -1.592640196 | 0.158749426 | 0.557345662 | -5.173006186 |
| **Q86XZ4** | SPATS2 | -0.072669953 | 3.187362826 | -1.973996544 | 0.09232591 | 0.446659583 | -5.174920543 |
| **Q9Y2R5** | MRPS17 | 0.195734539 | 3.170814125 | 1.986297939 | 0.09071641 | 0.443191118 | -5.175043847 |
| **Q8TDM6-1** | DLG5 | -0.100133993 | 3.179664658 | -1.152954607 | 0.289731258 | 0.684482558 | -5.175516824 |
| **Q99661-1** | KIF2C | -0.054191458 | 3.157446084 | -2.480335044 | 0.04491467 | 0.357204257 | -5.175896232 |
| **Q92782-2** | DPF1 | 0.096300993 | 3.172790701 | 1.072338122 | 0.321920987 | 0.709476351 | -5.176046476 |

| | | | | | | | |
|---|---|---|---|---|---|---|---|
| **P06753** | TPM3 | -0.128852504 | 3.146001124 | -1.51606875 | 0.176727359 | 0.577967785 | -5.176150578 |
| **Q9BQ83-1** | SLX1B; SLX1A | -0.00204067 | 3.108741959 | -0.011790826 | 0.990947681 | 0.996844685 | -5.176167963 |
| **O75746** | SLC25A12 | 0.182914518 | 3.182511115 | 2.209969466 | 0.065903907 | 0.394427892 | -5.17669643 |
| **Q6PJI9-1** | WDR59 | 0.067113789 | 3.169662286 | 1.967726739 | 0.093157146 | 0.447295866 | -5.177967798 |
| **Q9BVT8** | TMUB1 | 0.076544279 | 3.188057061 | 1.204915088 | 0.270443493 | 0.671731232 | -5.178518476 |
| **Q96FX7** | TRMT61A | 0.052351089 | 3.180466646 | 2.055714617 | 0.082145306 | 0.424346201 | -5.178681941 |
| **Q9H040-1** | SPRTN | 0.147966145 | 3.15041325 | 1.601785683 | 0.156720483 | 0.555124493 | -5.178862225 |
| **Q86WG5** | SBF2 | 0.083915235 | 3.216828964 | 1.582358585 | 0.161059877 | 0.561414897 | -5.178867296 |
| **O95620** | DUS4L | 0.101503996 | 3.117390668 | 1.104134732 | 0.308890261 | 0.696978291 | -5.179810302 |
| **Q9NPI8** | FANCF | 0.076601359 | 3.183980926 | 1.700253978 | 0.13637409 | 0.526478203 | -5.181978787 |
| **Q8N5U6-2** | RNF10 | 0.053794581 | 3.186060389 | 1.809388328 | 0.116783507 | 0.492826136 | -5.182630235 |
| **P09913** | IFIT2 | 0.059288276 | 3.1496886 | 1.739427325 | 0.129002634 | 0.51400498 | -5.182713637 |
| **Q99551** | MTERF; MTERF1 | 0.155267896 | 3.095940533 | 1.126032805 | 0.300170548 | 0.691062013 | -5.183464658 |
| **Q16854** | DGUOK | 0.121900111 | 3.208663827 | 1.765707711 | 0.124274009 | 0.50674745 | -5.183619364 |
| **Q15493-1** | RGN | 0.082842498 | 3.221968124 | 1.278821634 | 0.244904994 | 0.650150213 | -5.18381928 |
| **Q9BT17-1** | MTG1 | 0.122007211 | 3.18338658 | 1.932323734 | 0.097992192 | 0.456626373 | -5.183950189 |
| **Q9UMR5-3** | PPT2 | 0.085524374 | 3.171415142 | 1.704559874 | 0.135544466 | 0.524095099 | -5.185722985 |
| **Q15937** | ZNF79 | 0.091515796 | 3.268710603 | 0.72544412 | 0.493569906 | 0.821410327 | -5.186534827 |
| **P40222** | TXLNA | -0.072548601 | 3.200840715 | -2.339316495 | 0.054826922 | 0.376021618 | -5.187375098 |
| **Q9NRN7** | AASDHPPT | -0.039170999 | 3.188892786 | -2.292322677 | 0.058611733 | 0.379210276 | -5.188402456 |
| **Q14CZ0** | C16orf72 | -0.097321388 | 3.195718672 | -1.35949863 | 0.219453914 | 0.625602181 | -5.189132407 |
| **Q6IQ20** | NAPEPLD | 0.095016973 | 3.212374739 | 1.089958374 | 0.314645601 | 0.701837878 | -5.189928038 |
| **A2A3K4-2** | PTPDC1 | 0.087584284 | 3.164036667 | 1.113749718 | 0.30503622 | 0.693599527 | -5.190015031 |
| **P36404** | ARL 2.00 | -0.071049188 | 3.142722752 | -1.930665513 | 0.098224658 | 0.456905685 | -5.190373441 |
| **O15040** | TECPR2 | 0.100683729 | 3.184786475 | 1.255810052 | 0.252623176 | 0.656572825 | -5.190423921 |
| **Q15907** | RAB11B | -0.11168634 | 3.125848749 | -1.18772249 | 0.276701316 | 0.675408632 | -5.190515585 |
| **Q7Z591** | AKNA | -0.072248366 | 3.210215241 | -2.210123302 | 0.065889454 | 0.394427892 | -5.191268581 |
| **Q7L0Y3** | TRMT10C | 0.143302773 | 3.184048517 | 2.476088402 | 0.045184198 | 0.357414656 | -5.191357687 |
| **P32314** | FOXN2 | -0.08653199 | 3.187460971 | -1.15969323 | 0.287166283 | 0.682522792 | -5.191685777 |
| **P50748** | KNTC1 | -0.033712347 | 3.16338272 | -2.651380233 | 0.035344492 | 0.346253998 | -5.191930351 |
| **P52569-3** | SLC7A2 | 0.103356228 | 3.212323061 | 0.770877914 | 0.467983623 | 0.80548741 | -5.192160528 |
| **Q9UJT1** | TUBD1 | 0.094494112 | 3.171296859 | 1.435611624 | 0.197632827 | 0.603708078 | -5.192293976 |
| **Q9H330-2** | TMEM245 | -0.045659391 | 3.201646067 | -1.991547028 | 0.090038162 | 0.441848793 | -5.192524249 |
| **Q96GK7** | FAHD2A | 0.062626067 | 3.187745399 | 1.722382413 | 0.132161756 | 0.519814431 | -5.192857663 |
| **Q5U5Q3** | MEX3C | 0.137762212 | 3.110647656 | 1.16917908 | 0.283587942 | 0.679787821 | -5.192960889 |
| **Q9UBS9-2** | SUCO | -0.090832206 | 3.183052017 | -0.990948692 | 0.357305492 | 0.734600528 | -5.193121238 |
| **O60524** | NEMF | -0.067804066 | 3.186883198 | -2.229066262 | 0.064134194 | 0.390938548 | -5.195400129 |
| **Q7L592** | NDUFAF7 | 0.174027994 | 3.182584998 | 2.09789439 | 0.077339042 | 0.414399115 | -5.19624575 |
| **Q9BUY5** | ZNF426 | 0.075085376 | 3.255580557 | 0.592672532 | 0.573515935 | 0.858598851 | -5.196648071 |

| | | | | | | | |
|---|---|---|---|---|---|---|---|
| **A2RU67** | KIAA1467; FAM234 | 0.072886617 | 3.169082419 | 1.742098363 | 0.128514219 | 0.512665171 | -5.196704992 |
| **Q6DD87** | ZNF787 | -0.082144177 | 3.160590473 | -1.306191141 | 0.235993732 | 0.642113083 | -5.196734963 |
| **Q96C90** | PPP1R14B | -0.091154146 | 3.147454901 | -1.11499685 | 0.304539242 | 0.693086302 | -5.20007826 |
| **Q8N8R7** | ARL14EP | -0.038603809 | 3.207950575 | -2.095181533 | 0.077639461 | 0.414504124 | -5.201161536 |
| **Q08999** | RBL2 | 0.075708727 | 3.158815005 | 2.110351538 | 0.075974543 | 0.411304347 | -5.201577987 |
| **P10809** | HSPD1 | 0.18264198 | 3.199496936 | 3.105117478 | 0.01901749 | 0.305857884 | -5.201921137 |
| **Q6R327-3** | RICTOR | 0.035983202 | 3.170352201 | 2.454176156 | 0.046601952 | 0.358771814 | -5.202178963 |
| **Q6P1R3-1** | MSANTD2 | 0.087664283 | 3.188730439 | 1.847908732 | 0.110544047 | 0.481773149 | -5.203048243 |
| **Q9NUD5** | ZCCHC3 | -0.060556751 | 3.163683243 | -2.05894866 | 0.08176639 | 0.423686898 | -5.20341754 |
| **Q9NS93** | TM7SF3 | -0.094296526 | 3.208659045 | -1.638363868 | 0.148847396 | 0.546231571 | -5.203660507 |
| **Q96LA8-1** | PRMT6 | 0.075261186 | 3.193708462 | 1.625100644 | 0.151657969 | 0.550444296 | -5.203981688 |
| **O15054-1** | KDM6B | 0.082902529 | 3.179660883 | 0.901416038 | 0.399671117 | 0.763210165 | -5.204316121 |
| **Q9BWE0-4** | REPIN1 | -0.096394395 | 3.200682064 | -1.066314151 | 0.32443943 | 0.710703247 | -5.205786934 |
| **O15211** | RGL2 | -0.019187581 | 3.219622315 | -0.133347613 | 0.897969552 | 0.978757548 | -5.205861434 |
| **Q9Y2K2-4** | SIK3 | 0.07266953 | 3.153558772 | 0.634723313 | 0.547381742 | 0.846115385 | -5.206031793 |
| **P11362-21** | FGFR1 | 0.12965397 | 3.159297918 | 0.983313881 | 0.360776585 | 0.736666658 | -5.206372237 |
| **Q7Z417** | NUFIP2 | -0.044413876 | 3.208203263 | -2.55817667 | 0.040260592 | 0.355540374 | -5.206398558 |
| **O75648-1** | TRMU | 0.154040325 | 3.210140464 | 2.020827629 | 0.086346551 | 0.433917362 | -5.208090113 |
| **O15062** | ZBTB5 | 0.0887165 | 3.191154898 | 1.653231535 | 0.14575552 | 0.543674934 | -5.210349546 |
| **O76075** | DFFB | -0.082271002 | 3.154800627 | -1.027667466 | 0.340977067 | 0.722995142 | -5.210983294 |
| **Q6PKG0-1** | LARP1 | -0.029397007 | 3.184491864 | -2.708744893 | 0.032636516 | 0.339814312 | -5.211634977 |
| **Q8N5B7** | CERS5 | -0.010524091 | 3.140741312 | -0.062188776 | 0.952289251 | 0.989472196 | -5.212361185 |
| **Q3SYG4** | BBS9 | 0.069402765 | 3.177446604 | 1.840510029 | 0.111716643 | 0.48369958 | -5.212368374 |
| **Q969X0** | RILPL2 | -0.094655585 | 3.180401575 | -1.5723043 | 0.163349708 | 0.561580888 | -5.212795723 |
| **Q96SB4-2** | SRPK1 | -0.051670059 | 3.189150472 | -2.328473309 | 0.055677278 | 0.376021618 | -5.213177507 |
| **Q9HBR0** | SLC38A10 | -0.053911873 | 3.161910618 | -2.091267641 | 0.078074955 | 0.415568143 | -5.213459237 |
| **Q16587-5** | ZNF74 | -0.094235399 | 3.187040886 | -0.796803705 | 0.453799463 | 0.796605553 | -5.215115259 |
| **Q8N8D1** | PDCD7 | -0.114063009 | 3.204858647 | -1.733669316 | 0.130061612 | 0.515715456 | -5.216437722 |
| **Q9BQ24-2** | ZFYVE21 | -0.061098095 | 3.224785553 | -1.737640032 | 0.12933045 | 0.514537019 | -5.216625313 |
| **Q8N257** | HIST3H2BB | -0.095513997 | 3.181074157 | -1.206206952 | 0.269978181 | 0.671731232 | -5.217275923 |
| **Q9BZR9** | TRIM8 | 0.087658207 | 3.175874242 | 0.598321884 | 0.569962302 | 0.855535705 | -5.217431659 |
| **O75486-1** | SUPT3H | -0.117708327 | 3.055183844 | -0.993096316 | 0.356333824 | 0.734600528 | -5.217718291 |
| **Q9NTG7** | SIRT3 | 0.101608079 | 3.101493449 | 0.746102989 | 0.481821217 | 0.81235548 | -5.218682406 |
| **Q6DN12-1** | MCTP2 | -0.075488295 | 3.192263901 | -1.34827368 | 0.222848592 | 0.628740079 | -5.218835256 |
| **Q6IN85** | SMEK1; PPP4R3A | -0.034651052 | 3.171685708 | -2.415269675 | 0.049234375 | 0.366780161 | -5.218912697 |
| **Q4VC44-3** | FLYWCH1 | 0.076269359 | 3.208045791 | 1.076211838 | 0.320309903 | 0.708257023 | -5.219259818 |
| **P23528** | CFL1 | 0.081863904 | 3.158380006 | 2.813549072 | 0.028240191 | 0.330820062 | -5.219606548 |
| **P43378** | PTPN9 | 0.050337474 | 3.179716811 | 1.990993696 | 0.09010942 | 0.44185107 | -5.219701303 |
| **Q7Z6V5-1** | ADAT2 | -0.076993972 | 3.18722006 | -1.087817987 | 0.31552214 | 0.702585033 | -5.220058615 |

| | | | | | | | |
|---|---|---|---|---|---|---|---|
| **Q9NPD7** | NRN1 | -0.108981039 | 3.191251045 | -1.366058899 | 0.217491361 | 0.623815474 | -5.220195763 |
| **Q5H9S7** | DCAF17 | -0.106157724 | 3.181657525 | -0.96555723 | 0.368951163 | 0.743611616 | -5.221289692 |
| **Q08334** | IL10RB | -0.068279291 | 3.043681924 | -0.443942891 | 0.671546042 | 0.901175787 | -5.221801857 |
| **P35453** | HOXD13 | -0.147365056 | 3.119350245 | -1.63906391 | 0.148700431 | 0.546231571 | -5.223729591 |
| **Q3T8J9-1** | GON4L | -0.037092329 | 3.183231781 | -2.146171504 | 0.072185035 | 0.404094866 | -5.223876903 |
| **Q9P0J0** | NDUFA13 | 0.193241429 | 3.166712446 | 2.151794659 | 0.071607771 | 0.403638325 | -5.2250522 |
| **O00623** | PEX12 | 0.052669774 | 3.169616688 | 1.742537031 | 0.128434177 | 0.512665171 | -5.22557474 |
| **P56202** | CTSW | 0.093056038 | 3.276874943 | 1.359061924 | 0.219585118 | 0.625602181 | -5.226191026 |
| **Q13015** | MLLT11 | -0.219344758 | 3.17000012 | -1.790828026 | 0.119911138 | 0.49905395 | -5.226345491 |
| **Q9Y2Q3-1** | GSTK1 | 0.062400179 | 3.183840124 | 2.261185783 | 0.061266434 | 0.38388156 | -5.226417018 |
| **Q9NQY0-1** | BIN3 | -0.063127155 | 3.219095859 | -1.813000786 | 0.116184063 | 0.492573307 | -5.226718388 |
| **O15484** | CAPN5 | -0.082461448 | 3.162272896 | -0.662232098 | 0.530688143 | 0.839849063 | -5.226950477 |
| **Q9BSA9-1** | TMEM175 | -0.078452857 | 3.199260054 | -0.638687267 | 0.544956367 | 0.84497922 | -5.228084598 |
| **Q8WTV0-1** | SCARB1 | -0.075581952 | 3.204759988 | -1.320200928 | 0.231543444 | 0.638976263 | -5.228667516 |
| **Q86YC2** | PALB2 | 0.062287061 | 3.198490317 | 1.640424014 | 0.148415287 | 0.546231571 | -5.229982282 |
| **Q01543** | FLI1 | 0.0458119 | 3.194844298 | 2.177807828 | 0.068997662 | 0.399844221 | -5.230133391 |
| **Q99590-1** | SCAF11 | -0.042500889 | 3.179428691 | -2.502625277 | 0.043527174 | 0.357204257 | -5.230229092 |
| **P00367** | GLUD1 | 0.1477585 | 3.175835316 | 2.618925167 | 0.036980587 | 0.348943902 | -5.230264032 |
| **Q86WV5** | TEN1 | 0.105132682 | 3.185610388 | 0.986359572 | 0.359388744 | 0.73661838 | -5.230804091 |
| **P78314-4** | SH3BP2 | 0.087081725 | 3.148719043 | 0.677639847 | 0.521479915 | 0.834809173 | -5.231392181 |
| **P49406** | MRPL19 | 0.127278738 | 3.162466616 | 2.102431558 | 0.076839221 | 0.413084194 | -5.233528653 |
| **Q96DR7** | ARHGEF26 | -0.010111781 | 3.200171074 | -0.062684344 | 0.951909628 | 0.989472196 | -5.234228041 |
| **P01824** | IGHV4-39 | -0.084699004 | 3.124076932 | -0.798010835 | 0.453146447 | 0.796162876 | -5.234880326 |
| **Q86SK9-1** | SCD5 | -0.092113558 | 3.191096283 | -1.443239378 | 0.195557968 | 0.60151299 | -5.236235425 |
| **O95926** | SYF2 | -0.134986204 | 3.110087059 | -2.058651462 | 0.081801138 | 0.423686898 | -5.236450074 |
| **Q13277-1** | STX3 | -0.094066604 | 3.214339472 | -1.283731673 | 0.243284993 | 0.649360189 | -5.237236463 |
| **Q8IVD9** | NUDCD3 | -0.050478348 | 3.189422098 | -2.073618383 | 0.080069548 | 0.419060309 | -5.237281669 |
| **P51513-1** | NOVA1 | 0.069238877 | 3.171687772 | 1.479337165 | 0.186005836 | 0.588813889 | -5.237294291 |
| **Q99623** | PHB2 | 0.070854369 | 3.1822345 | 2.473945791 | 0.045320825 | 0.357414656 | -5.2384426 |
| **P08048** | ZFY | 0.122685963 | 3.103446487 | 0.697965832 | 0.509490167 | 0.829141305 | -5.239570442 |
| **O94979-7** | SEC31A | 0.073018938 | 3.243875013 | 0.661611542 | 0.531061156 | 0.839849063 | -5.240104878 |
| **O60907** | TBL1X | 0.076882317 | 3.231504665 | 1.262035546 | 0.250514551 | 0.654931921 | -5.240602692 |
| **P36578** | RPL4 | -0.127991134 | 3.172056635 | -2.859992619 | 0.026497266 | 0.328952544 | -5.241561212 |
| **Q96MH7** | C5orf34 | -0.0950198 | 3.134224924 | -0.805350383 | 0.449190235 | 0.794482729 | -5.24164003 |
| **P04350** | TUBB4A | -0.055480919 | 3.191965025 | -1.813961153 | 0.116025206 | 0.492508572 | -5.242230228 |
| **P11310-2** | ACADM | 0.166420787 | 3.17069305 | 2.586838922 | 0.038676548 | 0.352231442 | -5.242619632 |
| **Q6PHR2-1** | ULK3 | -0.053180521 | 3.177867015 | -1.911721587 | 0.100919432 | 0.463471034 | -5.24354471 |
| **Q9H4B0** | OSGEPL1 | 0.115075433 | 3.174113522 | 1.283937923 | 0.243217148 | 0.649360189 | -5.244958869 |
| **Q9BQ95** | ECSIT | 0.127055117 | 3.227286968 | 1.799759827 | 0.118395988 | 0.496152504 | -5.245043752 |

| | | | | | | | |
|---|---|---|---|---|---|---|---|
| **P48382** | RFX5 | 0.046700058 | 3.188312208 | 2.103219206 | 0.076752785 | 0.413084194 | -5.245146492 |
| **Q86VX9** | MON1A | -0.068753376 | 3.177343648 | -1.387686574 | 0.211132003 | 0.61590617 | -5.246130366 |
| **Q6P4R8-2** | NFRKB | 0.040391478 | 3.194171753 | 2.370903391 | 0.052425331 | 0.374522685 | -5.246818448 |
| **Q68D10-1** | SPTY2D1 | -0.081001725 | 3.233821121 | -2.08217383 | 0.079096343 | 0.416724434 | -5.247588655 |
| **Q96E16** | C8orf40; SMIM19 | 0.177121124 | 3.247758459 | 1.04317546 | 0.334261516 | 0.719277002 | -5.24797276 |
| **Q96RK0** | CIC | -0.067478311 | 3.182840291 | -1.574497541 | 0.16284762 | 0.561580888 | -5.24868827 |
| **Q8N684-3** | CPSF7 | -0.048881207 | 3.180293142 | -2.365173224 | 0.052852796 | 0.374840509 | -5.248863736 |
| **P01876** | IGHA1 | 0.124985663 | 2.313527299 | 0.458810171 | 0.661379698 | 0.896934512 | -5.249120293 |
| **Q96IQ9-2** | ZNF414 | 0.076039953 | 3.133250701 | 1.274416405 | 0.24636644 | 0.6518654 | -5.249592123 |
| **P25963** | NFKBIA | -0.079194531 | 3.186242192 | -1.146808075 | 0.292087565 | 0.685264687 | -5.250547193 |
| **P18433** | PTPRA | 0.054403682 | 3.161369946 | 2.036684969 | 0.084410884 | 0.42852682 | -5.251699534 |
| **Q7Z6K5-2** | C15orf38-AP3S2; C | 0.047324005 | 3.157156898 | 0.371095647 | 0.722414489 | 0.926635956 | -5.252921199 |
| **Q9NR48** | ASH1L | -0.191274217 | 3.141248269 | -0.780581736 | 0.462639022 | 0.802800142 | -5.253421069 |
| **Q6PIW4** | FIGNL1 | -0.052879925 | 3.210128267 | -2.220406076 | 0.064930649 | 0.393401641 | -5.253510475 |
| **O94925-1** | GLS | 0.166978669 | 3.20438092 | 1.57232429 | 0.163345125 | 0.561580888 | -5.253536996 |
| **Q8N612-2** | FAM160A2 | 0.087063677 | 3.188115112 | 1.45050998 | 0.193598743 | 0.598629202 | -5.25449818 |
| **Q02078-4** | MEF2A | -0.080106404 | 3.205651341 | -1.230787995 | 0.261253931 | 0.664637361 | -5.254561787 |
| **A8K8P3** | SFI1 | -0.052999904 | 3.143097602 | -0.412682704 | 0.693165864 | 0.909991705 | -5.255252903 |
| **Q9UMX0-3** | UBQLN1 | 0.076498177 | 3.16432118 | 1.513813883 | 0.177284401 | 0.578452275 | -5.255879329 |
| **Q96NE9-2** | FRMD6 | 0.07329109 | 3.10155979 | 1.705376384 | 0.135387694 | 0.524095099 | -5.256815756 |
| **Q9NP50-1** | FAM60A; SINHCAF | -0.053770446 | 3.200766472 | -1.821080403 | 0.114854179 | 0.489727184 | -5.257074515 |
| **P67870** | CSNK2B | -0.060532666 | 3.18025311 | -2.244654355 | 0.062725661 | 0.387695915 | -5.258128693 |
| **Q02978** | SLC25A11 | 0.142675096 | 3.195051068 | 2.126005763 | 0.074294274 | 0.407081381 | -5.259798817 |
| **O95164** | UBL3 | -0.09763716 | 3.099831495 | -0.685122339 | 0.517045232 | 0.833280927 | -5.262244251 |
| **Q9NVJ2** | ARL8B | -0.081693176 | 3.127281256 | -1.753658574 | 0.126420851 | 0.509218658 | -5.262320027 |
| **Q7Z4H7** | HAUS6 | -0.034528224 | 3.19326624 | -2.358930431 | 0.053322603 | 0.376021618 | -5.262405749 |
| **P56559-2** | ARL4C | -0.084615164 | 3.170071683 | -1.618305489 | 0.15311728 | 0.552031202 | -5.262549638 |
| **Q9BQ70** | TCF25 | -0.08227504 | 3.164367199 | -2.111626871 | 0.075836228 | 0.411304347 | -5.263945932 |
| **A6NDG6** | PGP | 0.066969816 | 3.189783047 | 2.208712907 | 0.066022086 | 0.394427892 | -5.264615871 |
| **Q9NP73-2** | ALG13 | -0.078265861 | 3.203608898 | -1.543700818 | 0.170031604 | 0.569040056 | -5.264949837 |
| **Q96AE7-2** | TTC17 | -0.077847684 | 3.251865324 | -1.036086566 | 0.337318026 | 0.721233457 | -5.266101406 |
| **Q8IXI1** | RHOT2 | 0.133389933 | 3.133944472 | 2.032347306 | 0.084936013 | 0.430367743 | -5.267269224 |
| **P63151-2** | PPP2R2A | -0.035358029 | 3.176731348 | -2.219593743 | 0.065005873 | 0.393401641 | -5.268061609 |
| **P13473-2** | LAMP2 | -0.100634893 | 3.178727812 | -1.496958275 | 0.181500084 | 0.58404948 | -5.268852326 |
| **Q8TDQ7** | GNPDA2 | -0.05831795 | 3.184489139 | -1.617246666 | 0.153345861 | 0.552031202 | -5.268985197 |
| **Q9Y6G3** | MRPL42 | 0.118564071 | 3.204063623 | 1.553020235 | 0.167827045 | 0.566424749 | -5.269981523 |
| **O94788-1** | ALDH1A2 | 0.04808203 | 3.180816386 | 1.841886201 | 0.11149762 | 0.483014501 | -5.270033306 |
| **Q15436** | SEC23A | -0.034354503 | 3.183952642 | -2.231976814 | 0.063868765 | 0.390577439 | -5.270645375 |
| **Q86V87** | FAM160B2 | -0.10450169 | 3.188191373 | -1.15769725 | 0.287924036 | 0.6826489 | -5.270657202 |

| | | | | | | | |
|---|---|---|---|---|---|---|---|
| **P20701-2** | ITGAL | -0.077624705 | 3.194134665 | -2.163866591 | 0.070384254 | 0.401998301 | -5.271863181 |
| **Q6P280** | ZNF529 | 0.080388491 | 3.151355237 | 0.546387755 | 0.603116182 | 0.872497827 | -5.27203927 |
| **Q8IWK6** | GPR125; ADGRA3 | 0.074076807 | 3.199779332 | 0.608740442 | 0.563443124 | 0.852578515 | -5.273537414 |
| **P52565-1** | ARHGDIA | 0.088568827 | 3.17035184 | 2.486697497 | 0.044513986 | 0.357204257 | -5.273705545 |
| **P48380** | RFX3 | 0.086977188 | 3.147974224 | 1.260884966 | 0.250903108 | 0.655139636 | -5.274032479 |
| **P20336** | RAB3A | -0.094720651 | 3.236329102 | -1.135781007 | 0.296354966 | 0.688880164 | -5.274492928 |
| **Q13424** | SNTA1 | -0.072306938 | 3.143926949 | -0.778292177 | 0.463896218 | 0.802800142 | -5.275357729 |
| **P61011-1** | SRP54 | -0.036651804 | 3.185887093 | -2.539611713 | 0.041322756 | 0.355540374 | -5.276135785 |
| **Q9BY49-1** | PECR | 0.075683222 | 3.21954627 | 0.581882156 | 0.580339667 | 0.863283777 | -5.277637163 |
| **Q9NX00** | TMEM160 | 0.135195207 | 3.166112302 | 1.662663416 | 0.143825794 | 0.539341922 | -5.277655862 |
| **Q6ZU67-1** | BEND4 | -0.09680407 | 3.243729533 | -0.907921964 | 0.396469792 | 0.762021989 | -5.278814902 |
| **Q9Y3D2** | MSRB2 | 0.095670979 | 3.264611959 | 1.481281665 | 0.185503624 | 0.588699395 | -5.278920338 |
| **Q96J88-3** | EPSTI1 | -0.091966108 | 3.166316025 | -1.594989411 | 0.158225904 | 0.55698042 | -5.279178276 |
| **Q9H0A8-1** | COMMD4 | -0.039800023 | 3.184077288 | -2.124818606 | 0.074420372 | 0.407491284 | -5.280017163 |
| **Q7Z3C6** | ATG9A | -0.08046024 | 3.136772312 | -1.661492839 | 0.144063962 | 0.539645535 | -5.28028507 |
| **O14639-1** | ABLIM1 | -0.092067974 | 3.168559533 | -1.206369393 | 0.269919721 | 0.671731232 | -5.281522799 |
| **Q8N157-1** | AHI1 | 0.035933804 | 3.213881064 | 0.290840433 | 0.780271629 | 0.94472083 | -5.283382767 |
| **Q6ZMS4** | ZNF852 | -0.044927367 | 3.154337373 | -0.280440768 | 0.787890972 | 0.947802649 | -5.283475195 |
| **Q8IVP5** | FUNDC1 | 0.071405849 | 3.245668209 | 1.307179744 | 0.235677246 | 0.641994289 | -5.285393527 |
| **Q9NRJ4-1** | TULP4 | 0.106008528 | 3.190891326 | 1.003573739 | 0.351623151 | 0.732738778 | -5.287291659 |
| **Q5T200-1** | ZC3H13 | 0.043052397 | 3.188728825 | 2.243796515 | 0.062802346 | 0.387695915 | -5.28743653 |
| **Q6N021** | TET2 | -0.063080503 | 3.168463409 | -1.349990244 | 0.222326455 | 0.627993355 | -5.287448691 |
| **Q9UM11-1** | FZR1 | 0.092087294 | 3.15591094 | 1.685311359 | 0.139290967 | 0.533333364 | -5.288091703 |
| **Q7L2R6** | ZNF765 | 0.007764307 | 3.236358566 | 0.055456523 | 0.957447705 | 0.989500907 | -5.28822772 |
| **Q6UB28** | METAP1D | 0.133407231 | 3.135915588 | 0.661882328 | 0.530898367 | 0.839849063 | -5.288833983 |
| **Q8N3X6-3** | LCORL | 0.101274547 | 3.226027172 | 0.678916514 | 0.520721544 | 0.834650661 | -5.289326305 |
| **Q96B36-3** | AKT1S1 | 0.071975953 | 3.183949396 | 1.422371982 | 0.201281743 | 0.607592494 | -5.289428936 |
| **O14773** | TPP1 | 0.059117172 | 3.168861003 | 1.784969821 | 0.120915107 | 0.499487166 | -5.29038862 |
| **O43674-1** | NDUFB5 | 0.185754543 | 3.161577155 | 1.939572272 | 0.096982398 | 0.455254009 | -5.290692916 |
| **Q8IXX5** | TMEM183A | 0.053733174 | 3.220398138 | 0.320088035 | 0.758986482 | 0.9384213 | -5.292750424 |
| **P22570-7** | FDXR | 0.145553376 | 3.18604605 | 2.150496934 | 0.071740577 | 0.403638325 | -5.293703345 |
| **O75503** | CLN5 | 0.050488211 | 3.177889175 | 1.830536503 | 0.113316584 | 0.487243547 | -5.293979739 |
| **Q9UNS1** | TIMELESS | -0.051919192 | 3.191133109 | -2.530072106 | 0.041879917 | 0.355540374 | -5.294154306 |
| **Q96ST8** | CEP89 | -0.083604364 | 3.285724813 | -1.145118027 | 0.292738256 | 0.68530312 | -5.294654645 |
| **P85299-5** | PRR5 | -0.106903245 | 3.097284424 | -0.926273079 | 0.387543885 | 0.757123654 | -5.296254439 |
| **Q96K31-1** | C8orf76; ZHX1-C8( | -0.117759793 | 3.140096095 | -1.031960569 | 0.339107295 | 0.721920542 | -5.296332695 |
| **Q8NDH2** | CCDC168 | 0.081448195 | 3.127576605 | 0.639483366 | 0.544470071 | 0.844555781 | -5.297088763 |
| **Q1KMD3** | HNRNPUL2 | -0.028318478 | 3.165794575 | -2.747715046 | 0.030922553 | 0.33673269 | -5.298667549 |
| **Q9UBV8** | PEF1 | 0.07174608 | 3.177783654 | 1.520438533 | 0.175652456 | 0.576677174 | -5.298853284 |

| | | | | | | | |
|---|---|---|---|---|---|---|---|
| **O75674** | TOM1L1 | 0.011522132 | 3.233432189 | 0.086278583 | 0.933853212 | 0.986793807 | -5.30026537 |
| **P11169** | SLC2A3 | -0.068244239 | 3.161682353 | -1.360234155 | 0.219233093 | 0.625602181 | -5.30040042 |
| **Q9P2X0-2** | DPM3 | -0.116754369 | 3.193580318 | -0.888268489 | 0.406199504 | 0.766336431 | -5.300592114 |
| **Q96HI0** | SENP5 | -0.056278806 | 3.191990939 | -1.733066809 | 0.130172904 | 0.515822305 | -5.300791605 |
| **O60683-1** | PEX10 | 0.081716499 | 3.177958051 | 0.976508286 | 0.363892829 | 0.73923 | -5.302017723 |
| **Q9NPR2-1** | SEMA4B | -0.143026366 | 3.06871527 | -0.935265056 | 0.383226217 | 0.754258979 | -5.302331442 |
| **Q9Y224** | C14orf166; RTRAF | -0.057333028 | 3.178731504 | -2.51830451 | 0.042578042 | 0.355540374 | -5.302505257 |
| **P47929** | LGALS7; LGALS7B | -0.323626436 | -0.428843765 | -1.326647507 | 0.22952064 | 0.637461728 | -5.303620363 |
| **Q8WWW8-2** | GAB3 | 0.1002334 | 3.169536877 | 0.931482811 | 0.38503785 | 0.754667212 | -5.303864928 |
| **Q14692** | BMS1 | -0.073674695 | 3.184748075 | -2.302516543 | 0.057768511 | 0.378064927 | -5.304237045 |
| **P17035-1** | ZNF28 | -0.071234773 | 3.164408454 | -1.039412353 | 0.335881275 | 0.720858943 | -5.304764094 |
| **Q9Y597** | KCTD3 | -0.061648839 | 3.12233848 | -0.453723779 | 0.664849202 | 0.898624353 | -5.307115848 |
| **Q0P6D6** | CCDC15 | -0.019055722 | 3.197689672 | -0.139022888 | 0.893658626 | 0.978276697 | -5.30746174 |
| **Q9BXJ8-1** | TMEM120A | -0.079456799 | 3.149446846 | -1.025150568 | 0.342077064 | 0.724167939 | -5.307541545 |
| **Q9Y5U9** | IER3IP1 | -0.272409998 | 3.195999577 | -1.111625155 | 0.305884393 | 0.694598791 | -5.308557661 |
| **P33981** | TTK | 0.042120499 | 3.138431957 | 1.978062136 | 0.091790851 | 0.445393971 | -5.308973387 |
| **Q9NZJ9-1** | NUDT4 | -0.070659617 | 3.181745634 | -1.318781507 | 0.231990938 | 0.639101249 | -5.309028845 |
| **Q9C0K7** | STRADB | -0.084102055 | 3.216853368 | -0.716872834 | 0.498500103 | 0.82494966 | -5.30936632 |
| **Q6P1Q9-1** | METTL2B | -0.045052193 | 3.217218222 | -2.265158751 | 0.060920952 | 0.383378328 | -5.310387946 |
| **Q9H488-1** | POFUT1 | 0.039346153 | 3.189538866 | 2.117482296 | 0.07520444 | 0.409500549 | -5.310463803 |
| **O00165-2** | HAX1 | 0.080955826 | 3.183351529 | 1.815451251 | 0.115779145 | 0.492169772 | -5.311246151 |
| **Q9P0P0** | RNF181 | -0.120859984 | 3.176039363 | -1.676997398 | 0.14093961 | 0.536483353 | -5.313216241 |
| **Q9HD47** | RANGRF | 0.143982473 | 3.212326531 | 1.455081872 | 0.19237593 | 0.596793208 | -5.313945894 |
| **Q9GZY8-2** | MFF | 0.073619142 | 3.18064749 | 1.373179099 | 0.215379062 | 0.620895009 | -5.314210431 |
| **Q96FF9** | CDCA5 | -0.049778838 | 3.121869236 | -1.705852919 | 0.135296278 | 0.524095099 | -5.314697211 |
| **P52294** | KPNA1 | -0.04552652 | 3.172206152 | -1.893684799 | 0.103552958 | 0.469881093 | -5.314799662 |
| **P48735** | IDH2 | 0.200187025 | 3.182180667 | 2.630471282 | 0.036389585 | 0.348943902 | -5.315009789 |
| **P46100** | ATRX | 0.066458463 | 3.20166131 | 1.181963742 | 0.278824689 | 0.676427621 | -5.315921344 |
| **Q14242-2** | SELPLG | -0.121523482 | 3.177082261 | -0.671774131 | 0.524973378 | 0.837910607 | -5.316639863 |
| **Q9BV86-1** | NTMT1 | 0.05298706 | 3.187882187 | 2.08380902 | 0.078911696 | 0.416709711 | -5.317137299 |
| **Q9NQ66-1** | PLCB1 | -0.103810908 | 3.16475301 | -1.031981429 | 0.33909823 | 0.721920542 | -5.317875814 |
| **Q9BVG3** | TRIM62 | -0.065293697 | 3.134205004 | -0.630542157 | 0.54994721 | 0.846638465 | -5.318225407 |
| **P60510** | PPP4C | -0.047721853 | 3.187557261 | -2.121551663 | 0.0747685 | 0.407645581 | -5.318503185 |
| **Q9P2G1** | ANKIB1 | -0.057459008 | 3.200780357 | -1.886343926 | 0.104644079 | 0.473191353 | -5.319772395 |
| **P78413-2** | IRX4 | -0.055633967 | 3.227266112 | -0.512848126 | 0.625091355 | 0.885387512 | -5.320168532 |
| **Q8WWM7-1** | ATXN2L | -0.049721515 | 3.182186444 | -2.500263654 | 0.043672036 | 0.357204257 | -5.320441595 |
| **Q5U5Z8** | AGBL2 | 0.066990384 | 3.222064263 | 0.517808718 | 0.621814081 | 0.883385606 | -5.321037191 |
| **P00973-3** | OAS1 | -0.057282384 | 3.233622503 | -0.477175006 | 0.648928529 | 0.89354197 | -5.321599605 |
| **P17022** | ZNF18 | 0.028707711 | 3.193489873 | 0.227663415 | 0.826932052 | 0.959077751 | -5.322518555 |

| | | | | | | | |
|---|---|---|---|---|---|---|---|
| **Q96MW7** | TIGD1 | 0.064736754 | 3.139907747 | 1.711226877 | 0.134269459 | 0.522414715 | -5.325545865 |
| **P98153-1** | DGCR2 | 0.088136468 | 3.207925447 | 0.612488159 | 0.561109058 | 0.852576894 | -5.325586806 |
| **P23497-4** | SP100 | -0.110033849 | 3.177503746 | -1.047927213 | 0.332225172 | 0.718239181 | -5.325830015 |
| **Q969Y0** | NXPE3 | -0.029105402 | 3.125797057 | -0.17325082 | 0.867747632 | 0.970868803 | -5.326563498 |
| **Q6PCB8** | EMB | 0.056238979 | 3.175169432 | 1.826492172 | 0.113971732 | 0.48833912 | -5.326697464 |
| **Q14691** | GINS1 | -0.043655106 | 3.174275944 | -2.19455968 | 0.067368207 | 0.395595275 | -5.327373249 |
| **Q9Y252** | RNF6 | -0.059114636 | 3.235848319 | -1.430608109 | 0.199004704 | 0.604393109 | -5.327472342 |
| **P43403-1** | ZAP70 | 0.065729326 | 3.173360795 | 2.297621871 | 0.058171824 | 0.379089796 | -5.328328574 |
| **Q8N3X6-1** | LCORL | -0.118544694 | 3.224486664 | -1.208569308 | 0.269129059 | 0.671554766 | -5.328760956 |
| **O95825-1** | CRYZL1 | -0.050769035 | 3.187606891 | -2.199917243 | 0.066855387 | 0.394427892 | -5.329373775 |
| **P40426-1** | PBX3 | 0.030042711 | 3.258929241 | 0.22697002 | 0.827448831 | 0.959077751 | -5.33062373 |
| **O94826** | TOMM70A; TOMM7 | 0.147571554 | 3.177473609 | 2.506457463 | 0.043293173 | 0.357204257 | -5.330703024 |
| **Q68DC2-4** | ANKS6 | -0.078607698 | 3.181305929 | -1.063996458 | 0.325412632 | 0.710703247 | -5.331396096 |
| **Q96LW7-2** | C9orf89; CARD19 | 0.090615235 | 3.204049744 | 1.563769649 | 0.165317322 | 0.56468019 | -5.332166395 |
| **P14923** | JUP | -0.21412054 | 2.168731631 | -1.24412769 | 0.256621591 | 0.659782078 | -5.332287626 |
| **P00451** | F8 | 0.006790066 | 3.216497149 | 0.053690598 | 0.958801189 | 0.990326778 | -5.332303065 |
| **P49642** | PRIM1 | -0.055143756 | 3.183494841 | -2.372970348 | 0.052272016 | 0.374493198 | -5.332339805 |
| **P20933** | AGA | -0.09257788 | 3.259834778 | -0.922538538 | 0.389347914 | 0.75855365 | -5.332363309 |
| **Q6NXG1-1** | ESRP1 | -0.026127058 | 3.186460347 | -0.163054674 | 0.875449356 | 0.972759481 | -5.333313661 |
| **Q8WVV4-1** | POF1B | 0.313176359 | 1.286955147 | 0.851345594 | 0.424955606 | 0.778986255 | -5.333392689 |
| **Q2T9K0-1** | TMEM44 | -0.073921821 | 3.150095424 | -0.758901211 | 0.474638572 | 0.809054593 | -5.334546001 |
| **Q5F1R6-2** | DNAJC21 | -0.040678337 | 3.1912001 | -1.986255122 | 0.090721963 | 0.443191118 | -5.33580219 |
| **Q8NC44** | FAM134A; RETREG | -0.049084355 | 3.167952055 | -0.442606894 | 0.672463328 | 0.90142081 | -5.33657081 |
| **Q9HC98-2** | NEK6 | 0.077078856 | 3.222556736 | 1.090495409 | 0.314425985 | 0.701837878 | -5.336579923 |
| **Q9Y6W3** | CAPN7 | -0.033059665 | 3.201369062 | -2.319040819 | 0.056428052 | 0.377614679 | -5.336599015 |
| **O75386-2** | TULP3 | -0.077370854 | 3.184031812 | -1.524496706 | 0.17465963 | 0.5748429 | -5.336645578 |
| **Q8N8K9-3** | KIAA1958 | 0.068294003 | 3.182826296 | 1.411218915 | 0.204402849 | 0.610519037 | -5.336718564 |
| **Q9NSI2-1** | FAM207A | -0.069670169 | 3.219942329 | -1.812524475 | 0.116262929 | 0.492644784 | -5.337084931 |
| **Q9Y227-1** | ENTPD4 | 0.095382711 | 3.169913504 | 0.932827019 | 0.384393251 | 0.754667212 | -5.337380493 |
| **Q9HBG6-6** | IFT122 | -0.054019145 | 3.170537771 | -1.948407251 | 0.095765532 | 0.452621745 | -5.337874776 |
| **O75822** | EIF3J | -0.079453066 | 3.171472629 | -2.286169065 | 0.059126861 | 0.379928744 | -5.338288713 |
| **Q02446** | SP4 | 0.062589321 | 3.196123687 | 1.743608032 | 0.128238956 | 0.512600477 | -5.338729455 |
| **Q9UBE8** | NLK | 0.079242723 | 3.147186502 | 1.406763725 | 0.205661786 | 0.611865721 | -5.338864041 |
| **O75570** | MTRF1 | 0.127319138 | 3.224987712 | 1.045042814 | 0.333460077 | 0.719098069 | -5.338992781 |
| **Q96NY9** | MUS81 | 0.077294622 | 3.168534584 | 1.169442553 | 0.283489091 | 0.679787821 | -5.33945239 |
| **Q9H6Q3-1** | SLA2 | 0.099068584 | 3.212586401 | 0.84428647 | 0.428612383 | 0.781217109 | -5.339698101 |
| **O43257** | ZNHIT1 | -0.066363115 | 3.196127875 | -1.126566806 | 0.299960482 | 0.690882899 | -5.340038253 |
| **Q9H3C7** | GGNBP2 | -0.12924573 | 3.194520681 | -1.960627747 | 0.0941073 | 0.448519793 | -5.340211974 |
| **A8MW92-2** | PHF20L1 | -0.23419897 | 2.974943023 | -1.454158299 | 0.192622382 | 0.597035716 | -5.340968972 |

| | | | | | | | |
|---|---|---|---|---|---|---|---|
| **P35555** | FBN1 | -0.084469735 | 3.134855484 | -1.187238391 | 0.276879284 | 0.675408632 | -5.341352405 |
| **P25705-1** | ATP5A1 | 0.169690641 | 3.188410633 | 2.800570209 | 0.02874877 | 0.331452005 | -5.341537953 |
| **Q5BJH7** | YIF1B | -0.105592546 | 3.176148752 | -1.330408924 | 0.228347615 | 0.635676875 | -5.341566642 |
| **O43566** | RGS14 | -0.093006431 | 3.165250349 | -0.955261026 | 0.373756509 | 0.746240828 | -5.342320976 |
| **Q5T3U5** | ABCC10 | -0.13267069 | 3.203121435 | -1.61605594 | 0.153603302 | 0.552139704 | -5.343134886 |
| **Q12968-1** | NFATC3 | -0.0623728 | 3.206376582 | -1.909683781 | 0.101213626 | 0.46428537 | -5.344958635 |
| **Q08ER8** | ZNF543 | -0.081040693 | 3.231772956 | -0.748737823 | 0.480336489 | 0.81235548 | -5.346568069 |
| **B2RTY4-4** | MYO9A | 0.047226583 | 3.18609133 | 1.967170807 | 0.093231209 | 0.447295866 | -5.346594474 |
| **Q9BRT3** | MIEN1 | 0.099789116 | 3.174965801 | 1.467864834 | 0.188994308 | 0.593033086 | -5.347812977 |
| **Q9BWU1-1** | CDK19 | 0.052342807 | 3.137404981 | 1.647995365 | 0.146837414 | 0.545761036 | -5.34805814 |
| **Q9Y2L8** | ZKSCAN5 | 0.068178917 | 3.184380132 | 0.592400225 | 0.573687555 | 0.858693976 | -5.348946244 |
| **Q9HCG7** | GBA2 | 0.078576399 | 3.153774208 | 1.319000346 | 0.231921896 | 0.639101249 | -5.349227409 |
| **Q15973-4** | ZNF124 | -0.058717562 | 3.207920334 | -1.608563221 | 0.155232644 | 0.554148197 | -5.350478526 |
| **Q9NNW5** | WDR6 | -0.189040106 | 3.247773569 | -1.995873162 | 0.089482974 | 0.440291106 | -5.350578663 |
| **Q6P1S2-1** | C3orf33 | 0.120902603 | 3.175862433 | 0.732831727 | 0.489346731 | 0.818152311 | -5.351121703 |
| **Q9BRQ3-1** | NUDT22 | -0.07047938 | 3.236151825 | -1.398039432 | 0.208147362 | 0.614231901 | -5.351519473 |
| **O96018** | APBA3 | -0.011817777 | 3.159617061 | -0.08026398 | 0.938452557 | 0.987739899 | -5.352115851 |
| **Q5T6L9-1** | C6orf70; ERMARD | -0.051257503 | 3.195130792 | -0.461886869 | 0.65928542 | 0.896852471 | -5.352405911 |
| **Q15063-1** | POSTN | 0.168288348 | 3.184392741 | 0.625346831 | 0.553145213 | 0.848243603 | -5.352834361 |
| **Q96PE7** | MCEE | 0.158284009 | 3.168772798 | 1.416162032 | 0.20301419 | 0.608695713 | -5.352957511 |
| **Q8N4P3** | HDDC3 | 0.069065084 | 3.163314035 | 1.680082372 | 0.140325703 | 0.535746138 | -5.353889476 |
| **Q676U5** | ATG16L1 | 0.035486855 | 3.206182216 | 2.021615097 | 0.086249387 | 0.433703403 | -5.354024786 |
| **Q8N465** | D2HGDH | 0.135115044 | 3.253253216 | 1.134676544 | 0.296785233 | 0.689058642 | -5.354829121 |
| **Q9NP58** | ABCB6 | 0.020988354 | 3.127310593 | 0.173330445 | 0.867687547 | 0.970868803 | -5.355298753 |
| **P28702-3** | RXRB | -0.075438763 | 3.151091946 | -1.635140116 | 0.149525961 | 0.546464774 | -5.35743636 |
| **Q86VH2-1** | KIF27 | 0.024630443 | 3.203172115 | 0.150062405 | 0.885284384 | 0.975708125 | -5.357636963 |
| **Q14CZ7** | FASTKD3 | 0.163074892 | 3.199877106 | 1.859759088 | 0.10869099 | 0.478951702 | -5.358024685 |
| **Q6EEV4-1** | POLR2M | -0.125861291 | 3.208352222 | -0.906529898 | 0.397153153 | 0.762571436 | -5.358708734 |
| **Q96FZ7** | CHMP6 | -0.045849585 | 3.160532398 | -1.803370619 | 0.117788767 | 0.495149077 | -5.360019848 |
| **P21281** | ATP6V1B2 | 0.045705357 | 3.175206668 | 2.414066034 | 0.049318234 | 0.366780161 | -5.361424147 |
| **Q01459** | CTBS | 0.056357753 | 3.226373038 | 1.489052292 | 0.183509093 | 0.58600472 | -5.3621034 |
| **Q12770** | SCAP | -0.077798372 | 3.153839528 | -1.397212423 | 0.208384378 | 0.614231901 | -5.362309124 |
| **A6ND36-1** | FAM83G | -0.03591871 | 3.234894427 | -0.209306508 | 0.840643923 | 0.961788885 | -5.3631576 |
| **A5PLL7** | TMEM189; TMEM1 | 0.084433806 | 3.21660322 | 1.036054045 | 0.337332099 | 0.721233457 | -5.363174841 |
| **Q9BV10** | ALG12 | 0.088633816 | 3.173121373 | 0.977758246 | 0.363318913 | 0.738441741 | -5.363911263 |
| **Q7Z478** | DHX29 | -0.040950821 | 3.185601716 | -2.426169884 | 0.048481652 | 0.364158835 | -5.364470455 |
| **Q9P1U0** | ZNRD1 | 0.079807001 | 3.20689045 | 0.55139064 | 0.599875612 | 0.872097298 | -5.36459948 |
| **Q96H86** | ZNF764 | -0.008830271 | 3.126674422 | -0.047019469 | 0.963915534 | 0.99136685 | -5.365251319 |
| **Q8TB36** | GDAP1 | 0.125581458 | 3.161455135 | 1.743885229 | 0.128188476 | 0.512600477 | -5.365620938 |

| | | | | | | | |
|---|---|---|---|---|---|---|---|
| **Q7Z7A4-1** | PXK | -0.053632512 | 3.198888936 | -1.753466106 | 0.126455433 | 0.509218658 | -5.366536693 |
| **Q9Y5J6** | FXC1; TIMM10B | 0.056344851 | 3.214362587 | 1.578477364 | 0.161940225 | 0.561414897 | -5.366599207 |
| **Q9Y228-1** | TRAF3IP3 | 0.073598954 | 3.197661964 | 0.999993866 | 0.353227127 | 0.732738778 | -5.36676833 |
| **Q9H8H3** | METTL7A | 0.057764587 | 3.202944421 | 0.418831099 | 0.688888047 | 0.908052188 | -5.366847645 |
| **Q04206** | RELA | -0.049076101 | 3.188061327 | -2.100458578 | 0.077056166 | 0.413656241 | -5.367769485 |
| **Q96H78** | SLC25A44 | 0.124824977 | 3.156630701 | 0.801728366 | 0.451139544 | 0.79513757 | -5.368555446 |
| **Q6ZMN8-2** | CCNI2 | 0.078714363 | 3.205496667 | 0.698437408 | 0.509214141 | 0.82913633 | -5.368691162 |
| **Q15424-3** | SAFB | 0.05242992 | 3.167852296 | 2.496604717 | 0.04389747 | 0.357204257 | -5.368757364 |
| **Q14209** | E2F2 | 0.060587757 | 3.158297358 | 0.584680754 | 0.578565311 | 0.861450787 | -5.368801507 |
| **Q96BR6-1** | ZNF669 | -0.063744373 | 3.209688829 | -1.325269921 | 0.229951581 | 0.637461728 | -5.3690873 |
| **Q5QJE6** | DNTTIP2 | -0.061549426 | 3.195573614 | -2.40149303 | 0.050203111 | 0.368557155 | -5.370238411 |
| **Q9P031** | CCDC59 | -0.059363928 | 3.152206563 | -1.63558796 | 0.14943152 | 0.546464774 | -5.370611063 |
| **Q70Z35-1** | PREX2 | -0.03626268 | 3.21422059 | -0.249748533 | 0.810522563 | 0.955498404 | -5.371148695 |
| **Q63HK3** | ZKSCAN2 | 0.079549651 | 3.110696631 | 0.670648749 | 0.525645329 | 0.838233243 | -5.371323664 |
| **Q99575** | POP1 | -0.041840812 | 3.173793868 | -2.282918068 | 0.059400882 | 0.381159465 | -5.371884063 |
| **Q9H0S4** | DDX47 | -0.060047087 | 3.179294404 | -2.25692002 | 0.061639612 | 0.384400879 | -5.372279723 |
| **Q96GC6-1** | ZNF274 | -0.072012218 | 3.232457698 | -0.700097884 | 0.508242999 | 0.82913633 | -5.372903659 |
| **P00505** | GOT2 | 0.18962745 | 3.197423785 | 2.423821393 | 0.048642813 | 0.364544514 | -5.373232285 |
| **Q9NZN8-1** | CNOT2 | -0.053047405 | 3.184403995 | -2.08906119 | 0.078321549 | 0.415674486 | -5.373979449 |
| **Q9NPJ8-3** | NXT2 | -0.070768523 | 3.21330184 | -1.08543477 | 0.31650047 | 0.703974147 | -5.374106976 |
| **Q08AG7** | MZT1 | 0.236086705 | 3.189081083 | 0.996847199 | 0.354641753 | 0.734091082 | -5.374760229 |
| **B1AJZ9-3** | FHAD1 | 0.083339062 | 3.168959724 | 1.706812738 | 0.135112332 | 0.523998833 | -5.375873963 |
| **Q8N4C8-1** | MINK1 | -0.095147822 | 3.203594726 | -1.414920696 | 0.203362111 | 0.608695713 | -5.376513186 |
| **O60220** | TIMM8A | -0.121161306 | 3.159189581 | -0.808863471 | 0.447305242 | 0.792383533 | -5.377016056 |
| **Q16836-2** | HADH | 0.124151292 | 3.173701276 | 2.290340311 | 0.058777175 | 0.37966232 | -5.377242524 |
| **Q6NYC8** | PPP1R18 | -0.046258436 | 3.170109421 | -2.139853308 | 0.072839281 | 0.404796855 | -5.377821644 |
| **Q9NX40** | OCIAD1 | 0.13811113 | 3.205578217 | 1.943604362 | 0.096425157 | 0.454510849 | -5.3804515 |
| **Q96SN8** | CDK5RAP2 | -0.066076032 | 3.204299091 | -0.913662221 | 0.393661257 | 0.760194378 | -5.380825803 |
| **Q99933** | BAG1 | 0.075121359 | 3.127588139 | 1.95195027 | 0.095281807 | 0.451948632 | -5.381212055 |
| **Q6ZQX7-4** | C17orf97 | -0.105782048 | 3.173830018 | -1.588593922 | 0.159654954 | 0.559241222 | -5.381760772 |
| **Q6NVY1** | HIBCH | 0.113196253 | 3.205178521 | 2.341464646 | 0.054660053 | 0.376021618 | -5.382289074 |
| **Q9NW68** | BSDC1 | -0.058136412 | 3.180067298 | -1.776317409 | 0.122412876 | 0.501640511 | -5.382290621 |
| **P19474** | TRIM21 | -0.053977089 | 3.201397108 | -2.211331942 | 0.06577601 | 0.394427892 | -5.382408538 |
| **P38117** | ETFB | 0.143993252 | 3.196318231 | 2.216861781 | 0.065259514 | 0.393496543 | -5.382602998 |
| **Q9UH03-1** | 3-Sep | -0.091501763 | 3.116094878 | -1.007645115 | 0.349805939 | 0.731407657 | -5.382684046 |
| **P17301** | ITGA2 | 0.076833088 | 3.205254262 | 0.894312501 | 0.403188563 | 0.766336431 | -5.383803354 |
| **Q9BYD1** | MRPL13 | 0.151215331 | 3.185674528 | 1.96333463 | 0.09374387 | 0.448493945 | -5.384769708 |
| **P05106** | ITGB3 | 0.064708655 | 3.178988024 | 0.912855266 | 0.394055168 | 0.760194378 | -5.385720144 |
| **O60499** | STX10 | 0.064988242 | 3.203177513 | 1.21027076 | 0.268518904 | 0.67097152 | -5.386140755 |

| | | | | | | | |
|---|---|---|---|---|---|---|---|
| **Q6ZMD2-1** | SPNS3 | -0.069949008 | 3.226841626 | -1.001550932 | 0.352528771 | 0.732738778 | -5.388967559 |
| **Q7LG56-6** | RRM2B | 0.076163262 | 3.18540443 | 1.340326299 | 0.225280225 | 0.630229362 | -5.389694097 |
| **O15169** | AXIN1 | -0.068079168 | 3.223538686 | -1.634636412 | 0.14963225 | 0.546587689 | -5.391253208 |
| **Q969S2-1** | NEIL2 | 0.07948552 | 3.225039411 | 1.158831555 | 0.287493204 | 0.682522792 | -5.391305613 |
| **A6NFI3** | ZNF316 | -0.045216298 | 3.157068563 | -0.356819871 | 0.732575795 | 0.928164811 | -5.391721078 |
| **O15269** | SPTLC1 | -0.049332421 | 3.193309111 | -1.857041846 | 0.109113181 | 0.479873319 | -5.392204541 |
| **P62910** | RPL32 | -0.119580264 | 3.187690728 | -2.59155919 | 0.038422012 | 0.352231442 | -5.39285629 |
| **Q92738-2** | USP6NL | 0.07578993 | 3.149893884 | 1.307088341 | 0.235706491 | 0.641994289 | -5.394360091 |
| **Q9Y5T5-2** | USP16 | 0.031435101 | 3.195729596 | 2.279011518 | 0.059731883 | 0.381486984 | -5.394753888 |
| **P29508** | SERPINB3 | -0.392233266 | -1.426421764 | -1.584058981 | 0.160675604 | 0.560635782 | -5.394986798 |
| **Q3MIX3** | ADCK5 | 0.035920186 | 3.15126272 | 0.227846871 | 0.82679534 | 0.959077751 | -5.395314804 |
| **A2AJT9-1** | CXorf23; BCLAF3 | 0.072884526 | 3.167892337 | 1.534872827 | 0.172144742 | 0.571776747 | -5.395452663 |
| **Q9BXW9-1** | FANCD2 | -0.03501697 | 3.192476548 | -2.395507072 | 0.050630163 | 0.369042796 | -5.395597906 |
| **Q4FZB7-1** | SUV420H1; KMT5B | -0.067887067 | 3.227489686 | -0.891901841 | 0.40438748 | 0.766336431 | -5.396706665 |
| **Q9UHN6-1** | TMEM2 | 0.063535828 | 3.170477658 | 0.585555409 | 0.578011418 | 0.861366689 | -5.397045687 |
| **Q9BXR0** | QTRT1 | 0.050799552 | 3.172493738 | 1.990679922 | 0.090149853 | 0.44185107 | -5.397563481 |
| **O95336** | PGLS | -0.084377025 | 3.173988466 | -2.117047206 | 0.075251202 | 0.409500549 | -5.398104896 |
| **Q9BXY0** | MAK16 | -0.043734633 | 3.203194556 | -2.102373295 | 0.076845619 | 0.413084194 | -5.398968645 |
| **Q8N511** | TMEM199 | -0.060930404 | 3.194166902 | -1.349923299 | 0.222346798 | 0.627993355 | -5.399002883 |
| **P43351** | RAD52 | -0.00311832 | 3.115818613 | -0.025122498 | 0.980714214 | 0.994229224 | -5.399707066 |
| **Q96N16-5** | JAKMIP1 | -0.135777675 | 3.160126964 | -1.001226743 | 0.352674082 | 0.732738778 | -5.40009914 |
| **P15622-1** | ZNF250 | -0.072009282 | 3.203812163 | -1.032836285 | 0.338726902 | 0.721920542 | -5.400986748 |
| **Q6PJG9** | LRFN4 | -0.028901643 | 3.140548372 | -0.256103791 | 0.805819286 | 0.954293355 | -5.401003267 |
| **Q8TF40-1** | FNIP1 | 0.047868247 | 3.177025232 | 1.753645253 | 0.126423244 | 0.509218658 | -5.402308889 |
| **Q9HBH0** | RHOF | -0.071548841 | 3.151859927 | -1.51342728 | 0.17738007 | 0.578452275 | -5.403100753 |
| **Q92753-2** | RORB | 0.078654836 | 3.191503452 | 0.83839805 | 0.431680102 | 0.783036167 | -5.404237918 |
| **Q7L3T8** | PARS2 | 0.136003443 | 3.170670557 | 1.265071141 | 0.249491921 | 0.65453932 | -5.404899621 |
| **P78317** | RNF4 | -0.03166087 | 3.260526664 | -0.234758176 | 0.821649827 | 0.959013938 | -5.405809294 |
| **Q86TB9** | PATL1 | 0.085174045 | 3.200540243 | 0.89005208 | 0.405309239 | 0.766336431 | -5.405890689 |
| **Q9BXL5** | HEMGN | -0.180626401 | 3.187207452 | -1.022814859 | 0.343100399 | 0.724789331 | -5.405954705 |
| **P02792** | FTL | 0.078397217 | 3.182796838 | 2.072167917 | 0.080235733 | 0.419390067 | -5.406192626 |
| **Q9HD33-1** | MRPL47 | 0.153384203 | 3.195202804 | 1.987987102 | 0.090497595 | 0.442736078 | -5.406260301 |
| **Q9NYV4-2** | CDK12 | -0.079066941 | 3.141840572 | -1.30108422 | 0.237634594 | 0.644090219 | -5.406633801 |
| **Q9NR31** | SAR1A | -0.072167029 | 3.127044019 | -1.544954211 | 0.169733546 | 0.569040056 | -5.40681947 |
| **Q96RP9** | GFM1 | 0.174639453 | 3.19435183 | 2.370353812 | 0.052466173 | 0.374522685 | -5.407027269 |
| **Q96SW2-1** | CRBN | 0.097833044 | 3.146061523 | 1.778972728 | 0.121951326 | 0.500466574 | -5.407063213 |
| **P53367** | ARFIP1 | 0.037358442 | 3.253699464 | 0.294802197 | 0.777375883 | 0.943803697 | -5.407311492 |
| **O60256-1** | PRPSAP2 | 0.028633495 | 3.177740536 | 2.12413439 | 0.074493146 | 0.407608849 | -5.407331504 |
| **Q9H765** | ASB8 | -0.058644391 | 3.241602118 | -1.279921251 | 0.244541374 | 0.650150213 | -5.408024202 |

| | | | | | | | |
|---|---|---|---|---|---|---|---|
| **P28324** | ELK4 | 0.066260127 | 3.21175832 | 1.374355808 | 0.215031749 | 0.620343952 | -5.408797249 |
| **Q9BWW4** | SSBP3 | 0.085768515 | 3.123480316 | 1.285271301 | 0.242778944 | 0.649360189 | -5.408818847 |
| **Q9HD23-4** | MRS2 | 0.150496567 | 3.177928299 | 1.439551953 | 0.196558513 | 0.603422483 | -5.409523766 |
| **Q9BXK5-1** | BCL2L13 | 0.149969928 | 3.165551264 | 1.881121595 | 0.105427188 | 0.475110045 | -5.409784446 |
| **Q86TJ2-1** | TADA2B | 0.049731926 | 3.198554192 | 1.570081365 | 0.163860072 | 0.562848368 | -5.410046811 |
| **P04156-1** | PRNP | -0.064246146 | 3.211071928 | -0.981611644 | 0.361554071 | 0.73711755 | -5.410262155 |
| **P23381** | WARS | 0.0265352 | 3.172995327 | 2.300896839 | 0.057901653 | 0.37862439 | -5.410718122 |
| **O43167-1** | ZBTB24 | -0.087636707 | 3.208953121 | -1.674965793 | 0.141345295 | 0.536801322 | -5.411138701 |
| **Q9NYC9-1** | DNAH9 | 0.051041786 | 3.18697891 | 0.407775937 | 0.696588583 | 0.912062672 | -5.411357329 |
| **Q08257** | CRYZ | 0.126876559 | 3.202825353 | 2.173866524 | 0.069386825 | 0.400056839 | -5.412277045 |
| **Q5VV63-1** | ATRNL1 | -0.019939696 | 3.232491319 | -0.143581931 | 0.890198415 | 0.977017047 | -5.412531142 |
| **Q14188** | TFDP2 | -0.060346099 | 3.156572526 | -1.874640053 | 0.10640714 | 0.475403904 | -5.412651859 |
| **P17017** | ZNF14 | -0.033515158 | 3.008872212 | -0.241308142 | 0.81678214 | 0.95663256 | -5.413595433 |
| **Q5T5Y3-3** | CAMSAP1 | -0.031462626 | 3.194484478 | -2.25780655 | 0.061561866 | 0.384217614 | -5.413603092 |
| **O94898** | LRIG2 | -0.073791886 | 3.164550151 | -0.996284064 | 0.354895387 | 0.734091082 | -5.413770624 |
| **Q7L3V2** | C22orf29; RTL10; G | -0.02855582 | 3.119124956 | -0.224658333 | 0.829172375 | 0.959338069 | -5.413829724 |
| **P17040-1** | ZSCAN20 | 0.083151133 | 3.150950722 | 0.556062501 | 0.596858346 | 0.871378088 | -5.414146533 |
| **Q9Y673** | ALG5 | -0.043235488 | 3.177370031 | -1.894520058 | 0.103429521 | 0.469881093 | -5.414243291 |
| **Q8NC69** | KCTD6 | -0.017556771 | 3.244463407 | -0.105275761 | 0.919345104 | 0.983742622 | -5.416120847 |
| **P36956-4** | SREBF1 | -0.084104573 | 3.124924266 | -1.284058427 | 0.243177517 | 0.649360189 | -5.41646297 |
| **P50991** | CCT4 | 0.02669545 | 3.18648084 | 2.742549263 | 0.031144166 | 0.33673269 | -5.41646519 |
| **P16885** | PLCG2 | 0.0296297 | 3.174977181 | 2.274431919 | 0.060122321 | 0.382749875 | -5.417855147 |
| **P15848** | ARSB | 0.078294509 | 3.185045422 | 1.407495636 | 0.205454485 | 0.611717339 | -5.418089763 |
| **P63172** | DYNLT1 | -0.012539632 | 3.202532735 | -0.082969283 | 0.936383494 | 0.987232823 | -5.418469694 |
| **P43251-2** | BTD | 0.064941203 | 3.15658039 | 0.869858371 | 0.415473654 | 0.77203561 | -5.41853584 |
| **Q96S97** | MYADM | 0.098536049 | 3.180907388 | 0.87954677 | 0.410573799 | 0.769906516 | -5.418900608 |
| **Q9NYJ1-2** | COA4 | -0.577711065 | 3.170845947 | -1.192887974 | 0.274808368 | 0.674207874 | -5.419331456 |
| **Q6L8Q7-1** | PDE12 | 0.070624863 | 3.179429672 | 2.216965752 | 0.065249843 | 0.393496543 | -5.41942394 |
| **Q99442** | SEC62 | -0.042973837 | 3.207946192 | -1.943414945 | 0.096451264 | 0.454510849 | -5.419569326 |
| **Q96B70** | LENG9 | -0.065966561 | 3.19764143 | -0.897403266 | 0.401655284 | 0.765047743 | -5.420217482 |
| **Q9H0E2** | TOLLIP | 0.045436263 | 3.190636934 | 1.854823882 | 0.109458987 | 0.479940204 | -5.420628443 |
| **Q96HJ3-1** | CCDC34 | 0.057269696 | 3.25467037 | 0.456637008 | 0.662860943 | 0.897330072 | -5.42074016 |
| **P02675** | FGB | -0.333124525 | 2.758450206 | -0.658289551 | 0.533060802 | 0.840132297 | -5.420796663 |
| **Q6ZS86** | GK5 | 0.105231424 | 3.19365141 | 1.270697442 | 0.247606125 | 0.652280071 | -5.421230281 |
| **P12236** | SLC25A6 | 0.155184021 | 3.224075518 | 1.723492097 | 0.131953841 | 0.519814431 | -5.421645013 |
| **Q9Y535** | POLR3H | -0.087358245 | 3.194251109 | -1.375530196 | 0.214685621 | 0.619913059 | -5.422167768 |
| **Q96CG3** | TIFA | 0.048685203 | 3.174613187 | 1.861351709 | 0.108444283 | 0.478395241 | -5.423241317 |
| **Q8TEU7** | RAPGEF6 | -0.027964003 | 3.187495475 | -2.335245539 | 0.055144598 | 0.376021618 | -5.424260059 |
| **Q9BWQ6** | YIPF2 | 0.080791733 | 3.199251159 | 1.132738578 | 0.297541464 | 0.689530645 | -5.426168712 |

| | | | | | | | |
|---|---|---|---|---|---|---|---|
| **Q9Y580** | RBM7 | 0.038122273 | 3.188655932 | 1.875093175 | 0.106338342 | 0.475403904 | -5.426417248 |
| **Q9ULX3** | NOB1 | -0.061808231 | 3.199161835 | -2.083222838 | 0.078977838 | 0.416709711 | -5.4275721 |
| **Q9UGM3-9** | DMBT1 | -0.003189523 | 2.756220174 | -0.014946766 | 0.988524917 | 0.996184213 | -5.428368079 |
| **A6NJZ7** | RIMBP3C | -0.026297879 | 3.169777148 | -0.167059834 | 0.872422227 | 0.97208641 | -5.428529649 |
| **P61163** | ACTR1A | 0.069446237 | 3.189449785 | 2.165653601 | 0.070204948 | 0.401998301 | -5.42860839 |
| **Q9NVN3-1** | RIC8B | -0.052918359 | 3.206276102 | -1.686542842 | 0.139048337 | 0.533333364 | -5.428743509 |
| **Q96C00** | ZBTB9 | -0.04809027 | 3.221210902 | -1.756801317 | 0.125857466 | 0.509142907 | -5.428943053 |
| **O15056** | SYNJ2 | -0.036484018 | 3.194733145 | -2.127114066 | 0.074176746 | 0.406910627 | -5.429295475 |
| **Q15131-1** | CDK10 | 0.072461324 | 3.181167742 | 0.873751577 | 0.413499534 | 0.771122868 | -5.430021895 |
| **Q3B7J2-1** | GFOD2 | -0.09364963 | 3.152003735 | -1.227213561 | 0.262507342 | 0.666332535 | -5.430052531 |
| **O75347** | TBCA | -0.056936112 | 3.18799573 | -2.244758903 | 0.062716322 | 0.387695915 | -5.430390761 |
| **Q9UKU7** | ACAD8 | 0.16239538 | 3.188713517 | 1.8250767 | 0.114201899 | 0.48833912 | -5.430742967 |
| **Q562E7-1** | WDR81 | 0.067404189 | 3.155573055 | 1.553859806 | 0.167629753 | 0.566424749 | -5.431780974 |
| **Q2NL68-1** | C19orf55; PROSER | 0.064191589 | 3.13074648 | 0.617843182 | 0.557784105 | 0.851117349 | -5.431905977 |
| **Q99808-2** | SLC29A1 | -0.126478594 | 3.175285521 | -1.431659053 | 0.198715838 | 0.603977556 | -5.432275633 |
| **P51114-1** | FXR1 | -0.0462217 | 3.209307305 | -1.845597435 | 0.110909057 | 0.482457068 | -5.433358976 |
| **P09669** | COX6C | 0.159988411 | 3.190668437 | 1.935956932 | 0.097484756 | 0.455866032 | -5.433654328 |
| **Q5T6S3-2** | PHF19 | 0.033548933 | 3.270251507 | 0.226020019 | 0.828157007 | 0.959077751 | -5.433758452 |
| **P32189-3** | GK | 0.147215714 | 3.166533427 | 2.132338162 | 0.073625298 | 0.405653949 | -5.433903106 |
| **P25791-3** | LMO2 | -0.070037715 | 3.136601524 | -0.512691494 | 0.625194987 | 0.885387512 | -5.434024432 |
| **Q8N6T7-1** | SIRT6 | -0.048369597 | 3.167530681 | -1.800733962 | 0.11823187 | 0.496152504 | -5.434212958 |
| **Q14C87** | TMEM132D | 0.083260596 | 3.165240349 | 0.916480353 | 0.392287928 | 0.759806823 | -5.434224834 |
| **Q8N0V3-1** | RBFA | 0.098376714 | 3.206157241 | 1.257234418 | 0.252139379 | 0.656218606 | -5.43429338 |
| **Q96F24-1** | NRBF2 | -0.05722593 | 3.196825372 | -1.799469014 | 0.118445026 | 0.496152504 | -5.435376825 |
| **P22090** | RPS4Y1 | -0.133276573 | 3.128908464 | -1.272840369 | 0.246891137 | 0.652280071 | -5.435727093 |
| **Q3ZCQ3** | FAM174B | 0.065958313 | 3.19863544 | 0.715113873 | 0.49951588 | 0.825004623 | -5.435954302 |
| **Q9UHX3** | EMR2; ADGRE2 | 0.065332511 | 3.135183725 | 1.270188064 | 0.247776344 | 0.652280071 | -5.436933453 |
| **Q9BYZ6-2** | RHOBTB2 | 0.055781757 | 3.133293261 | 0.508789938 | 0.627779336 | 0.885601354 | -5.437783492 |
| **Q13239-5** | SLA | -0.123293767 | 3.192131387 | -1.238038997 | 0.258727094 | 0.661376267 | -5.43849208 |
| **O15027-5** | SEC16A | -0.046469408 | 3.224502971 | -0.404942591 | 0.698568505 | 0.913150177 | -5.438699397 |
| **P04083** | ANXA1 | 0.039312927 | 3.158734249 | 2.551542393 | 0.040636841 | 0.355540374 | -5.438951645 |
| **Q9Y575-3** | GPR75-ASB3; ASB | -0.070499815 | 3.214249251 | -1.319485468 | 0.231768908 | 0.639101249 | -5.438959136 |
| **Q16774-2** | GUK1 | 0.051311357 | 3.179177192 | 2.058623842 | 0.081804368 | 0.423686898 | -5.439892572 |
| **Q13825** | AUH | 0.113892273 | 3.251533508 | 0.914690984 | 0.393159502 | 0.760194378 | -5.440471733 |
| **Q9P0S2** | COX16 | -0.084158839 | 3.198176719 | -1.13810936 | 0.295449608 | 0.688880164 | -5.441175814 |
| **P15153** | RAC2 | 0.07820248 | 3.191990442 | 2.005747149 | 0.088228573 | 0.437773138 | -5.441826145 |
| **Q9Y3B2** | EXOSC1 | 0.054336154 | 3.174395665 | 2.095880657 | 0.077561928 | 0.414504124 | -5.442468404 |
| **Q15018** | FAM175B; ABRAXA | 0.045536947 | 3.196722596 | 2.115784211 | 0.075387111 | 0.409678928 | -5.442812405 |
| **Q8IUR7-1** | ARMC8 | -0.060522975 | 3.212505574 | -1.762020302 | 0.124927238 | 0.507956453 | -5.443504669 |

| | | | | | | | |
|---|---|---|---|---|---|---|---|
| **Q9UDV6** | ZNF212 | -0.061081852 | 3.216920466 | -0.710500968 | 0.502186268 | 0.826806835 | -5.444381896 |
| **Q9NP90** | RAB9B | -0.07242687 | 3.192101047 | -1.35700991 | 0.220202558 | 0.625644783 | -5.444578924 |
| **P32242** | OTX1 | -0.065494608 | 3.196910283 | -1.153977062 | 0.289340842 | 0.683966971 | -5.446359615 |
| **O43865** | AHCYL1 | -0.045841307 | 3.172889172 | -1.982112158 | 0.091260911 | 0.444137581 | -5.446950922 |
| **Q8TF71** | SLC16A10 | -0.027791411 | 3.186237341 | -0.263861747 | 0.800089794 | 0.952507937 | -5.447066699 |
| **Q8NCE2-1** | MTMR14 | -0.032745044 | 3.171852528 | -1.967569122 | 0.093178139 | 0.447295866 | -5.447121479 |
| **Q92934** | BAD | -0.088203628 | 3.177213195 | -1.679125377 | 0.140515871 | 0.536062029 | -5.447182045 |
| **Q9BX68** | HINT2 | 0.073846257 | 3.192665302 | 1.87020881 | 0.107082252 | 0.475555333 | -5.447329324 |
| **Q9NVR7** | TBCCD1 | -0.074636483 | 3.195800369 | -1.386466355 | 0.211486305 | 0.61590617 | -5.447523059 |
| **Q9H0U6** | MRPL18 | 0.130925625 | 3.189975366 | 1.277218478 | 0.245435968 | 0.650646902 | -5.44838087 |
| **Q9P203-1** | BTBD7 | -0.016980685 | 3.216363184 | -0.132728864 | 0.898439782 | 0.978757548 | -5.448590823 |
| **Q6PIZ9** | TRAT1 | 0.0735624 | 3.146522305 | 1.555314027 | 0.167288536 | 0.566424749 | -5.449364795 |
| **O95980** | RECK | 0.052212852 | 3.168513141 | 1.553406514 | 0.167736246 | 0.566424749 | -5.450294618 |
| **Q96CS4** | ZNF689 | -0.067389552 | 3.172934175 | -0.534973214 | 0.610546408 | 0.877336085 | -5.450559892 |
| **A6NNY8** | USP27X | -0.021678902 | 3.165050382 | -0.128215117 | 0.901871424 | 0.97952714 | -5.450862386 |
| **P27824-2** | CANX | -0.034894095 | 3.199351936 | -2.508686329 | 0.043157679 | 0.357204257 | -5.451805675 |
| **Q9NZ09-4** | UBAP1 | -0.064929269 | 3.217305605 | -1.498227709 | 0.181179399 | 0.58362594 | -5.453067638 |
| **Q9BU64-1** | CENPO | -0.061399553 | 3.182218087 | -1.351874081 | 0.22175469 | 0.627993355 | -5.453828565 |
| **Q8N4V1** | MMGT1 | -0.056096945 | 3.191820706 | -1.707759844 | 0.134931057 | 0.523800962 | -5.454774474 |
| **Q96EL3** | MRPL53 | 0.122728157 | 3.189794472 | 1.54685955 | 0.169281385 | 0.568204733 | -5.455162423 |
| **P12757** | SKIL | -0.073957215 | 3.210655271 | -1.4750834 | 0.187108824 | 0.590381098 | -5.45591662 |
| **Q53HC5** | KLHL26 | -0.063647672 | 3.217826801 | -1.302815178 | 0.237077315 | 0.643518711 | -5.456276996 |
| **Q53EP0** | FNDC3B | 0.031113112 | 3.262464936 | 0.327072676 | 0.753936022 | 0.936379818 | -5.456583168 |
| **Q9H900-1** | ZWILCH | -0.066308346 | 3.187391693 | -1.780595507 | 0.121670084 | 0.49982876 | -5.456648681 |
| **Q32MZ4-4** | LRRFIP1 | 0.057076338 | 3.186182951 | 0.485383195 | 0.643402431 | 0.891474831 | -5.456931813 |
| **Q96T52-1** | IMMP2L | 0.072776006 | 3.211152573 | 1.04723649 | 0.332520557 | 0.718682216 | -5.457409874 |
| **Q12891** | HYAL2 | -0.009526758 | 3.164944207 | -0.067669427 | 0.94809164 | 0.989472196 | -5.4579468 |
| **Q8WVP7-3** | LMBR1 | -0.079179873 | 3.171031982 | -0.841854108 | 0.429877665 | 0.782603406 | -5.458102077 |
| **O14972** | DSCR3 | -0.051753635 | 3.192917936 | -1.469080966 | 0.18867544 | 0.592734825 | -5.458313654 |
| **O14607-4** | UTY | 0.079886539 | 3.186563697 | 0.903239653 | 0.398771836 | 0.762880385 | -5.458696032 |
| **Q8IXW5** | RPAP2 | -0.066519215 | 3.196693825 | -2.049388793 | 0.082891576 | 0.425319102 | -5.459930588 |
| **Q9BW60** | ELOVL1 | -0.057860768 | 3.1850152 | -1.33011466 | 0.228439192 | 0.635709064 | -5.460572971 |
| **Q96KQ7-2** | EHMT2 | -0.038788961 | 3.19394215 | -2.292903919 | 0.058563315 | 0.379210276 | -5.460723289 |
| **Q9NXN4-1** | GDAP2 | -0.043016525 | 3.179837564 | -1.892541765 | 0.103722115 | 0.470360848 | -5.460782268 |
| **P24588** | AKAP5 | -0.070845045 | 3.202683607 | -0.854445054 | 0.423357205 | 0.777884596 | -5.461014052 |
| **Q92622-1** | KIAA0226; RUBCN | 0.026047752 | 3.197644832 | 0.162560746 | 0.875822829 | 0.972759481 | -5.462134711 |
| **P47736-4** | RAP1GAP | -0.09164515 | 3.229189047 | -1.066270193 | 0.324457866 | 0.710703247 | -5.464072189 |
| **O95260** | ATE1 | 0.058584953 | 3.199856265 | 0.760558615 | 0.473713776 | 0.808788313 | -5.46423749 |
| **Q96AE7** | TTC17 | 0.000161281 | 3.274510252 | 0.001086806 | 0.99916559 | 0.999417174 | -5.464435544 |

| | | | | | | | |
|---|---|---|---|---|---|---|---|
| **Q5J8M3-1** | EMC4 | -0.052070816 | 3.200676182 | -1.39705647 | 0.208429101 | 0.614231901 | -5.464677205 |
| **P35221-2** | CTNNA1 | -0.072307601 | 3.179336087 | -2.05408416 | 0.082337007 | 0.424508448 | -5.465720528 |
| **Q5T9C2** | FAM102A | -0.058427363 | 3.173075657 | -0.511091533 | 0.626254092 | 0.885387512 | -5.465833747 |
| **P54259** | ATN1 | -0.064400432 | 3.171234872 | -1.199741724 | 0.272313722 | 0.672884265 | -5.466253737 |
| **Q9HAV0** | GNB4 | 0.153583139 | 3.206977742 | 1.676581657 | 0.141022537 | 0.536483353 | -5.466675689 |
| **Q8IUX1-1** | TMEM126B | 0.093182132 | 3.210158619 | 0.98742028 | 0.358906388 | 0.736063824 | -5.466738451 |
| **Q8TCX1-2** | DYNC2LI1 | -0.059222478 | 3.150892743 | -0.756372994 | 0.47605164 | 0.809713018 | -5.467348371 |
| **P31323** | PRKAR2B | 0.053803869 | 3.182593685 | 1.825132782 | 0.114192771 | 0.48833912 | -5.467941577 |
| **P22083** | FUT4 | -0.024010783 | 3.158864559 | -0.178910272 | 0.863479413 | 0.969659921 | -5.468085951 |
| **Q9BSF4** | C19orf52; TIMM29 | 0.079555701 | 3.19523294 | 1.197015641 | 0.273303653 | 0.672884265 | -5.468214913 |
| **Q8TCD5** | NT5C | 0.046319474 | 3.148692181 | 1.744654345 | 0.128048512 | 0.512600477 | -5.468441851 |
| **P08651** | NFIC | 0.068457397 | 3.192697631 | 1.558223477 | 0.166607809 | 0.566132944 | -5.468937103 |
| **O43310-2** | CTIF | -0.077472789 | 3.218270014 | -1.103504427 | 0.309144304 | 0.696978291 | -5.469498326 |
| **Q9UBL6-1** | CPNE7 | -0.058582012 | 3.237880226 | -0.658317613 | 0.533043891 | 0.840132297 | -5.469906692 |
| **Q9NNW7-1** | TXNRD2 | 0.230597938 | 3.146590068 | 1.516270434 | 0.176677613 | 0.577967785 | -5.470100342 |
| **Q9NXL9-1** | MCM9 | -0.052224915 | 3.213601455 | -1.489149676 | 0.183484223 | 0.58600472 | -5.471039367 |
| **Q9NRF8** | CTPS2 | 0.074908294 | 3.199402651 | 0.429086637 | 0.681780218 | 0.904168215 | -5.471363157 |
| **P45985-2** | MAP2K4 | 0.040799606 | 3.180425211 | 1.884078159 | 0.104983134 | 0.473915342 | -5.471625582 |
| **P33552** | CKS2 | -0.120755636 | 3.205728838 | -1.78901411 | 0.120221136 | 0.499295152 | -5.471641397 |
| **Q9BV81** | EMC6 | -0.062834827 | 3.187257976 | -0.53891205 | 0.607976727 | 0.875726951 | -5.47189073 |
| **Q96F46** | IL17RA | -0.022255165 | 3.142106542 | -0.226112699 | 0.828087911 | 0.959077751 | -5.472511435 |
| **Q8TAP9** | MPLKIP | 0.070779486 | 3.206443854 | 0.91345127 | 0.393764203 | 0.760194378 | -5.474589865 |
| **Q15181** | PPA1 | 0.053065322 | 3.195474092 | 2.703898007 | 0.032856613 | 0.339814312 | -5.474626611 |
| **Q9H5J0** | ZBTB3 | 0.05953705 | 3.188643872 | 0.484975356 | 0.64367643 | 0.891474831 | -5.474853199 |
| **Q9NZ43-1** | USE1 | -0.052336737 | 3.193545017 | -1.598379154 | 0.157473361 | 0.555457754 | -5.477182589 |
| **P16455** | MGMT | -0.068096692 | 3.186820885 | -1.955515707 | 0.09479747 | 0.450458072 | -5.478211636 |
| **Q9ULU4-13** | ZMYND8 | 0.079565992 | 3.19575031 | 1.444841137 | 0.195124793 | 0.600645673 | -5.478403792 |
| **Q5TF21** | SOGA3 | -0.070938513 | 3.090548262 | -1.761206613 | 0.125071831 | 0.508284244 | -5.478914747 |
| **Q659C4-1** | LARP1B | -0.057541387 | 3.217313298 | -1.561166352 | 0.165921892 | 0.565325887 | -5.479040792 |
| **Q96LT7-1** | C9orf72 | 0.066140689 | 3.159678959 | 0.883323798 | 0.408675202 | 0.768662396 | -5.479208848 |
| **Q5JTJ3-2** | COA6 | -0.153023153 | 3.161537193 | -1.753068806 | 0.126526847 | 0.509248125 | -5.48105929 |
| **P09622** | DLD | 0.172725585 | 3.170213237 | 2.368131297 | 0.052631678 | 0.374840509 | -5.481627151 |
| **Q9BQE5** | APOL2 | 0.04151382 | 3.176989657 | 1.826394639 | 0.113987577 | 0.48833912 | -5.481669599 |
| **P54098** | POLG | 0.171366469 | 3.197928811 | 1.8841764 | 0.104968411 | 0.473915342 | -5.481987512 |
| **Q96AX2-1** | RAB37 | 0.075232354 | 3.203013046 | 0.809587197 | 0.446917613 | 0.792204208 | -5.482376433 |
| **Q8NCW6** | GALNT11 | 0.062990492 | 3.200773378 | 1.578100921 | 0.162025851 | 0.561414897 | -5.482498234 |
| **Q9ULH7-5** | MKL2 | 0.049446216 | 3.197038116 | 1.712767651 | 0.133976439 | 0.521785691 | -5.482638687 |
| **P31785-1** | IL2RG | -0.0703746 | 3.198625357 | -1.51548875 | 0.176870487 | 0.578048547 | -5.483008825 |
| **P48651** | PTDSS1 | -0.052649976 | 3.218216924 | -1.723611514 | 0.131931485 | 0.519814431 | -5.483033132 |

| | | | | | | | |
|---|---|---|---|---|---|---|---|
| **O43286** | B4GALT5 | -0.064940456 | 3.104423862 | -0.535826038 | 0.60998952 | 0.876853037 | -5.484234368 |
| **Q9H061-1** | TMEM126A | 0.099054163 | 3.200176094 | 1.509564052 | 0.178338696 | 0.57989621 | -5.484631085 |
| **P30405** | PPIF | 0.124812298 | 3.21057151 | 1.847230865 | 0.110650976 | 0.481974785 | -5.484911296 |
| **Q9H7J1** | PPP1R3E | -0.063013022 | 3.177379187 | -0.972110145 | 0.365917835 | 0.740729864 | -5.485054215 |
| **Q9BVM4** | GGACT | -0.058021818 | 3.154032276 | -0.778914623 | 0.463554199 | 0.802800142 | -5.485166116 |
| **Q9NWS1-1** | PARPBP | -0.095413778 | 3.162852252 | -1.195091967 | 0.274004043 | 0.673773483 | -5.48540202 |
| **Q9HCX3** | ZNF304 | 0.04010797 | 3.174542915 | 0.367739184 | 0.724798188 | 0.926648409 | -5.486117812 |
| **Q9BYD2** | MRPL9 | 0.120343098 | 3.172566544 | 1.714898287 | 0.133572253 | 0.521489705 | -5.486545465 |
| **Q9UJJ9** | GNPTG | -0.207724062 | 3.200461545 | -1.289858158 | 0.241276776 | 0.647178928 | -5.487037383 |
| **O75145-1** | PPFIA3 | 0.068349469 | 3.176078057 | 1.225222438 | 0.263207787 | 0.666686026 | -5.487455784 |
| **Q9Y653** | GPR56; ADGRG1 | -0.055377874 | 3.135308242 | -1.584961362 | 0.160472024 | 0.560635782 | -5.487527569 |
| **Q9UL12** | SARDH | -0.006965728 | 3.180540258 | -0.068533664 | 0.947429884 | 0.989472196 | -5.488088552 |
| **O95278** | EPM2A | 0.086357348 | 3.179950997 | 0.970765912 | 0.366538494 | 0.741119482 | -5.489723269 |
| **Q56NI9** | ESCO2 | -0.075360686 | 3.113341329 | -2.130721111 | 0.073795547 | 0.406305157 | -5.489924304 |
| **Q8NEF3-2** | CCDC112 | -0.072035535 | 3.211825614 | -0.60178247 | 0.567791951 | 0.854214553 | -5.490219962 |
| **Q9NR81-2** | ARHGEF3 | 0.031346687 | 3.19038409 | 0.240853851 | 0.817119471 | 0.956656724 | -5.491538031 |
| **Q9UPQ9** | TNRC6B | -0.039130789 | 3.156954811 | -2.033716911 | 0.084769852 | 0.430074378 | -5.491784998 |
| **Q96SD1** | DCLRE1C | 0.056158605 | 3.198762998 | 1.561339848 | 0.165881536 | 0.565325887 | -5.492021931 |
| **Q9H869-9** | YY1AP1 | -0.065845166 | 3.188915391 | -1.234551839 | 0.259939665 | 0.663418129 | -5.4927197 |
| **Q99829** | CPNE1 | 0.051981324 | 3.161438104 | 2.265253663 | 0.060912723 | 0.383378328 | -5.492898651 |
| **O95429** | BAG4 | -0.143201769 | 3.187052744 | -1.981910829 | 0.091287183 | 0.444137581 | -5.494636149 |
| **Q5VT06** | CEP350 | 0.036574011 | 3.166648263 | 2.187744601 | 0.068026333 | 0.397403835 | -5.49492265 |
| **Q9ULE0** | WWC3 | 0.062776994 | 3.209297789 | 1.391202974 | 0.210113965 | 0.615470161 | -5.494956509 |
| **Q9GZU1** | MCOLN1 | 0.015078135 | 3.129119503 | 0.095616984 | 0.926717692 | 0.985528933 | -5.495949224 |
| **Q6IQ22** | RAB12 | -0.054392337 | 3.211293404 | -1.457688486 | 0.191681916 | 0.596505086 | -5.495989061 |
| **O95208-1** | EPN2 | 0.061171812 | 3.213392877 | 0.750055785 | 0.479594985 | 0.812101908 | -5.496132503 |
| **P06681-1** | C2 | -0.25515579 | 2.715083334 | -0.753044633 | 0.477916299 | 0.811322085 | -5.496422544 |
| **Q15173** | PPP2R5B | -0.024533758 | 3.204753027 | -0.198006148 | 0.849115263 | 0.964964904 | -5.496740815 |
| **Q8N5H7-1** | SH2D3C | -0.075922178 | 3.179817144 | -0.882609813 | 0.409033601 | 0.768994785 | -5.49674285 |
| **Q15034** | HERC3 | 0.080862239 | 3.185856944 | 1.423956945 | 0.200841721 | 0.606617388 | -5.49694775 |
| **Q6P4F2** | FDX1L | -0.007623421 | 3.202414222 | -0.040757486 | 0.968717903 | 0.992306703 | -5.498143622 |
| **Q9H8X2** | IPPK | -0.097205854 | 3.213953091 | -0.991662839 | 0.356982154 | 0.734600528 | -5.498160713 |
| **Q9Y689-1** | ARL5A | -0.07742429 | 3.186805625 | -1.030716339 | 0.33964835 | 0.722416219 | -5.498751128 |
| **E9PAV3** | NACA | -0.047676158 | 3.226915228 | -2.203507354 | 0.066513973 | 0.394427892 | -5.499265996 |
| **Q8WV60** | PTCD2 | 0.117699912 | 3.172068786 | 1.614218076 | 0.154001458 | 0.552139704 | -5.499726584 |
| **P51858** | HDGF | -0.075148438 | 3.163272965 | -2.261501558 | 0.061238901 | 0.38388156 | -5.500347576 |
| **Q5JPH6-2** | EARS2 | 0.117123259 | 3.183077627 | 1.705693844 | 0.135326788 | 0.524095099 | -5.500723466 |
| **Q8N4P2** | TTC30B | -0.020103392 | 3.169154393 | -0.15784268 | 0.879392017 | 0.97338977 | -5.501861006 |
| **Q8N5X7** | EIF4E3 | 0.060415733 | 3.198520472 | 1.100148565 | 0.31049977 | 0.698646466 | -5.501980856 |

| | | | | | | | |
|---|---|---|---|---|---|---|---|
| **Q9C0A6** | SETD5 | -0.061588067 | 3.192878224 | -1.432732937 | 0.19842106 | 0.603708078 | -5.502260031 |
| **Q1ED39** | C16orf88; KNOP1 | -0.083617758 | 3.194759888 | -2.163692192 | 0.070401778 | 0.401998301 | -5.503822038 |
| **Q9H7N4** | SCAF1 | -0.056760958 | 3.157603076 | -1.593324849 | 0.158596683 | 0.557345662 | -5.505079957 |
| **Q15434** | RBMS2 | 0.061606112 | 3.171345511 | 1.079839507 | 0.318807102 | 0.707150066 | -5.505182036 |
| **Q6B0I6** | KDM4D | 0.048266123 | 3.177881498 | 0.607194775 | 0.564407457 | 0.852578515 | -5.505673029 |
| **O00459** | PIK3R2 | -0.050061639 | 3.184380405 | -1.555793319 | 0.167176217 | 0.566424749 | -5.506224479 |
| **P31641** | SLC6A6 | 0.092385199 | 3.184397932 | 1.032571888 | 0.338841715 | 0.721920542 | -5.506294166 |
| **Q13588** | GRAP | -0.059220537 | 3.174334977 | -1.666063946 | 0.143136038 | 0.539313633 | -5.506761328 |
| **Q16595-1** | FXN | 0.120352488 | 3.210396823 | 1.599269663 | 0.157276221 | 0.555457754 | -5.507581866 |
| **Q92805** | GOLGA1 | -0.063072501 | 3.196975438 | -1.442883379 | 0.195654362 | 0.601527622 | -5.507642636 |
| **Q8TBE0** | BAHD1 | -0.133998276 | 3.106517174 | -1.345291514 | 0.223758288 | 0.629881158 | -5.508989882 |
| **Q9UNY5** | ZNF232 | 0.022855797 | 3.180773003 | 0.217797992 | 0.834293299 | 0.960780962 | -5.50912252 |
| **Q9Y5V3-2** | MAGED1 | -0.052306108 | 3.179671255 | -1.893883306 | 0.103523609 | 0.469881093 | -5.509287959 |
| **Q9NTX7** | RNF146 | -0.072968667 | 3.220540134 | -0.810814603 | 0.446260758 | 0.792020296 | -5.50929666 |
| **O75845** | SC5D; SC5DL | 0.071055556 | 3.184354919 | 0.934812438 | 0.383442671 | 0.754258979 | -5.510375686 |
| **O95183** | VAMP5 | -0.06243097 | 3.182926452 | -1.419700657 | 0.202025339 | 0.608654462 | -5.510499443 |
| **Q15906** | VPS72 | -0.076685559 | 3.192890509 | -1.475682559 | 0.186953101 | 0.59035866 | -5.51063008 |
| **Q92626** | PXDN | -0.042248796 | 3.174388661 | -1.622413143 | 0.152233553 | 0.55152557 | -5.510742837 |
| **Q8ND24** | RNF214 | -0.037395217 | 3.164005952 | -1.735709429 | 0.129685453 | 0.515290389 | -5.510819969 |
| **O14908-1** | GIPC1 | 0.075700165 | 3.169542266 | 1.741700749 | 0.128586812 | 0.512665171 | -5.511049115 |
| **O15143** | ARPC1B | 0.034626011 | 3.1645271 | 2.463735618 | 0.04597783 | 0.357780473 | -5.511337547 |
| **Q6P9B6** | KIAA1609; TLDC1 | -0.059026072 | 3.147268026 | -1.549515728 | 0.168652912 | 0.567053487 | -5.511654962 |
| **O95644-1** | NFATC1 | -0.053432105 | 3.212263768 | -2.037306331 | 0.084335927 | 0.428420039 | -5.511834976 |
| **Q15697-1** | ZNF174 | 0.076658007 | 3.191905963 | 0.500737396 | 0.633131251 | 0.887490806 | -5.512060702 |
| **Q9UKS7-1** | IKZF2 | -0.051891823 | 3.199961898 | -1.664776168 | 0.143396877 | 0.539313633 | -5.51226575 |
| **Q6ZPD9-1** | DPY19L3 | -0.078275931 | 3.182447642 | -0.801286881 | 0.451377551 | 0.79513757 | -5.512594419 |
| **Q8NHX9** | TPCN2 | -0.048559543 | 3.220534247 | -0.650742086 | 0.537621533 | 0.84335148 | -5.512941309 |
| **P20774** | OGN | 0.155625322 | 2.768462232 | 1.014224319 | 0.346885078 | 0.728715479 | -5.513878848 |
| **Q8WVN8-1** | UBE2Q2 | -0.086675582 | 3.211247654 | -1.124289681 | 0.30085711 | 0.691344519 | -5.514014344 |
| **A8MTY0** | ZNF724P; ZNF724 | -0.056617321 | 3.175557795 | -0.657437381 | 0.533574523 | 0.840132297 | -5.514480628 |
| **Q8TEH3** | DENND1A | 0.058303914 | 3.228915859 | 0.875664622 | 0.412532027 | 0.770644294 | -5.514521509 |
| **Q8WU79-1** | SMAP2 | -0.07889506 | 3.16670212 | -1.918448199 | 0.099954306 | 0.461169746 | -5.514756488 |
| **Q86VW0** | SESTD1 | -0.056099874 | 3.201089138 | -1.461142668 | 0.190765761 | 0.595227814 | -5.51551433 |
| **Q9H4L4** | SENP3 | -0.034877346 | 3.184949316 | -1.970982378 | 0.092724596 | 0.44675374 | -5.515668198 |
| **Q6NVH7** | SWSAP1 | -0.024102799 | 3.16882187 | -0.227948028 | 0.826719961 | 0.959077751 | -5.515687345 |
| **Q9NYV6** | RRN3 | -0.06942015 | 3.270825257 | -1.52874238 | 0.173626503 | 0.574454961 | -5.517047148 |
| **Q92597** | NDRG1 | 0.079544457 | 3.174419277 | 1.813842545 | 0.116044814 | 0.492508572 | -5.518325718 |
| **A8K0R7-5** | ZNF839 | -0.051831237 | 3.176460968 | -0.485014466 | 0.643650151 | 0.891474831 | -5.518406163 |
| **Q92560** | BAP1 | 0.037305282 | 3.188652974 | 1.934026783 | 0.097754009 | 0.456510811 | -5.519928385 |

| | | | | | | | |
|---|---|---|---|---|---|---|---|
| **Q16644** | MAPKAPK3 | -0.041543224 | 3.185567657 | -1.872394078 | 0.106748796 | 0.475441804 | -5.520349945 |
| **Q8N6Q8** | METTL25 | 0.052770701 | 3.184731604 | 0.424641117 | 0.68485702 | 0.905978473 | -5.521478032 |
| **Q6P4Q7** | CNNM4 | -0.086219298 | 3.173634736 | -0.944993179 | 0.378596484 | 0.750267026 | -5.522845975 |
| **Q9BU61-1** | NDUFAF3 | 0.148596581 | 3.190609124 | 1.417135743 | 0.202741655 | 0.608695713 | -5.523194983 |
| **P30626-1** | SRI | -0.044493055 | 3.174574784 | -2.206112942 | 0.066267298 | 0.394427892 | -5.523403955 |
| **Q9BS91** | SLC35A5 | -0.060889484 | 3.198795439 | -1.019374444 | 0.344612165 | 0.725897233 | -5.523474791 |
| **O95273-1** | CCNDBP1 | -0.058175894 | 3.251693099 | -1.465546202 | 0.189603615 | 0.593538505 | -5.523899252 |
| **O60512** | B4GALT3 | 0.030008739 | 3.200159685 | 0.223612877 | 0.829952183 | 0.959820974 | -5.525194717 |
| **Q9UN36-1** | NDRG2 | 0.020935123 | 3.167328108 | 0.193568526 | 0.852447957 | 0.966699831 | -5.526048918 |
| **P09884** | POLA1 | -0.03554527 | 3.184068792 | -2.431298902 | 0.048131605 | 0.363139916 | -5.526251037 |
| **P41162** | ETV3 | 0.063810669 | 3.167896719 | 1.16273431 | 0.286014989 | 0.681580409 | -5.526508258 |
| **Q9NX61** | TMEM161A | -0.062317565 | 3.215174069 | -0.854735688 | 0.423207549 | 0.777884596 | -5.526522759 |
| **Q8TAM2-4** | TTC8 | 0.052960204 | 3.166301553 | 1.474432191 | 0.187278208 | 0.590446574 | -5.526649186 |
| **P51606-1** | RENBP | -0.066504922 | 3.16888826 | -1.238256677 | 0.258651564 | 0.661376267 | -5.526710993 |
| **Q6UWY0** | ARSK | 0.052198078 | 3.199893358 | 1.358291743 | 0.21981668 | 0.625644783 | -5.5268616 |
| **Q9UBV2** | SEL1L | 0.058667702 | 3.192545967 | 1.900702323 | 0.102520381 | 0.467580037 | -5.527779437 |
| **Q93075** | TATDN2 | 0.044879907 | 3.181832866 | 1.615512805 | 0.153720866 | 0.552139704 | -5.528130677 |
| **A0A0U1RRE** | LOC550643; LINC0 | 0.105475183 | 3.144904975 | 0.471083971 | 0.653044923 | 0.895639895 | -5.528325583 |
| **Q05481-1** | ZNF91 | -0.040534488 | 3.112342049 | -0.406212844 | 0.697680544 | 0.912939342 | -5.528358294 |
| **O60858-3** | TRIM13 | -0.072319332 | 3.205688402 | -1.225692699 | 0.263042212 | 0.66662532 | -5.529706314 |
| **Q9NZZ3** | CHMP5 | 0.069850267 | 3.197963736 | 1.608122945 | 0.15532889 | 0.554148197 | -5.529722139 |
| **Q03181** | PPARD | -0.012637075 | 3.194986896 | -0.123476195 | 0.905476719 | 0.98075504 | -5.530740094 |
| **Q9NXB0-1** | MKS1 | 0.056850257 | 3.194637209 | 0.881720709 | 0.40948023 | 0.769470301 | -5.532482884 |
| **Q0VDD7** | C19orf57 | 0.070248799 | 3.255116326 | 0.859676868 | 0.420669092 | 0.776717624 | -5.532577098 |
| **Q14442** | PIGH | 0.044290523 | 3.207378858 | 0.529517942 | 0.614115301 | 0.879837049 | -5.532638772 |
| **Q9BS34** | ZNF670 | 0.077203803 | 3.211374862 | 1.197483609 | 0.273133501 | 0.672884265 | -5.533426672 |
| **P56381** | ATP5E | 0.085966692 | 3.200660739 | 1.5786673 | 0.161897039 | 0.561414897 | -5.533813852 |
| **O14561** | NDUFAB1 | 0.130126096 | 3.187860797 | 1.786736892 | 0.120611413 | 0.499295152 | -5.534283292 |
| **Q96PM9** | ZNF385A | -0.039821892 | 3.146527089 | -0.459088925 | 0.661189816 | 0.896934512 | -5.534608781 |
| **P78368** | CSNK1G2 | 0.058744149 | 3.170316954 | 1.198509655 | 0.272760749 | 0.672884265 | -5.536178369 |
| **Q92613** | PHF16; JADE3 | -0.055679107 | 3.16562952 | -1.933695511 | 0.097800294 | 0.456510811 | -5.53621285 |
| **Q8N5I4** | DHRSX | 0.063657123 | 3.194328233 | 0.867273463 | 0.416788202 | 0.773310479 | -5.536548305 |
| **Q7Z3V4-1** | UBE3B | -0.061533713 | 3.19462935 | -1.483556022 | 0.1849178 | 0.588139279 | -5.537648311 |
| **Q7Z412** | PEX26 | 0.055237846 | 3.147663759 | 0.630517579 | 0.549962312 | 0.846638465 | -5.538121058 |
| **Q9NY27** | PPP4R2 | 0.03871004 | 3.180143635 | 2.202058996 | 0.066651495 | 0.394427892 | -5.53815405 |
| **Q15723-5** | ELF2 | 0.038282854 | 3.174548811 | 1.877332728 | 0.105998949 | 0.475403904 | -5.539293032 |
| **Q9NXK8** | FBXL12 | 0.066542971 | 3.184592623 | 1.03402439 | 0.338211358 | 0.721920542 | -5.540101142 |
| **Q13724-1** | MOGS | -0.059784544 | 3.19267151 | -1.852249553 | 0.109861701 | 0.480909762 | -5.540390064 |
| **A8MVW0** | FAM171A2 | -0.045102235 | 3.105565163 | -0.320689464 | 0.758551091 | 0.9384213 | -5.541372432 |

| | | | | | | | |
|---|---|---|---|---|---|---|---|
| **P35080-2** | PFN2 | -0.066557878 | 3.210124171 | -1.134311576 | 0.296927528 | 0.689187615 | -5.541762982 |
| **Q7L4E1-1** | FAM73B; MIGA2 | 0.018072383 | 3.111700552 | 0.14101939 | 0.89214301 | 0.978205385 | -5.544010929 |
| **Q4G0N4** | NADK2; NADKD1 | 0.159388995 | 3.18976975 | 2.026228583 | 0.08568233 | 0.431546517 | -5.544096548 |
| **Q8NF37** | LPCAT1 | -0.062809025 | 3.19077679 | -1.6501949 | 0.146382022 | 0.544988363 | -5.544210043 |
| **Q6P6C2-1** | ALKBH5 | 0.054753736 | 3.189596568 | 1.78276459 | 0.121295146 | 0.499487166 | -5.54703766 |
| **Q9BQG2** | NUDT12 | 0.052077354 | 3.183981085 | 0.60910456 | 0.563216097 | 0.852578515 | -5.547598519 |
| **Q7Z398** | ZNF550 | -0.055618482 | 3.194384426 | -0.655003324 | 0.535043582 | 0.840674016 | -5.547683801 |
| **Q04637-9** | EIF4G1 | 0.01222255 | 3.219578064 | 0.104578714 | 0.919876883 | 0.983742622 | -5.548312835 |
| **O94925-3** | GLS | 0.115997427 | 3.169935609 | 1.256372859 | 0.252431918 | 0.656572825 | -5.548342872 |
| **Q9ULT8** | HECTD1 | -0.033858866 | 3.188121913 | -2.321985214 | 0.056192582 | 0.37738805 | -5.548966288 |
| **A1A4S6** | ARHGAP10 | -0.032519691 | 3.214452043 | -0.393507502 | 0.706585132 | 0.917890594 | -5.549935446 |
| **Q15036-1** | SNX17 | -0.052908911 | 3.230571731 | -1.850146163 | 0.110191822 | 0.481294133 | -5.550192106 |
| **Q4G148-1** | GXYLT1 | 0.069572759 | 3.097348226 | 0.276093497 | 0.791083585 | 0.948368382 | -5.550331785 |
| **Q5T447** | HECTD3 | 0.049502851 | 3.165517594 | 2.050417052 | 0.082769809 | 0.425319102 | -5.551608221 |
| **Q9Y5S9** | RBM8A; LOC10106 | -0.095188265 | 3.164994255 | -1.653660692 | 0.145667185 | 0.543674934 | -5.55218945 |
| **Q08554-1** | DSC1 | -0.332981693 | 1.669151073 | -0.671834964 | 0.524937071 | 0.837910607 | -5.55291556 |
| **Q9BQ90** | KLHDC3 | 0.059107716 | 3.149108483 | 1.420396946 | 0.201831279 | 0.608326826 | -5.55349374 |
| **Q9NUK0** | MBNL3 | -0.055052925 | 3.151239615 | -1.063094081 | 0.32579218 | 0.710703247 | -5.553817279 |
| **O75534-4** | CSDE1 | 0.05644566 | 3.194390226 | 1.410206625 | 0.204688287 | 0.610527836 | -5.553843952 |
| **Q96RK4-2** | BBS4 | 0.037664324 | 3.127895402 | 1.722610639 | 0.132118969 | 0.519814431 | -5.554060819 |
| **O00748** | CES2 | 0.047747427 | 3.185240765 | 0.494120849 | 0.63754677 | 0.889065271 | -5.55423809 |
| **Q03060-31** | CREM | 0.056267802 | 3.211180003 | 0.927913077 | 0.386753666 | 0.756278088 | -5.554306406 |
| **Q9H2P9-5** | DPH5 | 0.039817305 | 3.213346841 | 1.670655251 | 0.142209744 | 0.538432214 | -5.55497006 |
| **Q8WUR7** | C15orf40 | 0.096385319 | 3.242696281 | 0.63631673 | 0.546405999 | 0.845412981 | -5.555907005 |
| **Q9UHJ6** | SHPK | -0.036795533 | 3.171355177 | -1.850552948 | 0.110127903 | 0.481294133 | -5.555992463 |
| **Q9NPA5-1** | ZFP64 | -0.081854222 | 3.069597133 | -1.279045096 | 0.244831061 | 0.650150213 | -5.556357137 |
| **Q9UN81** | LOC440264; L1RE1 | 0.033007471 | 3.180286188 | 0.356813638 | 0.732580244 | 0.928164811 | -5.556492493 |
| **Q9UK73** | FEM1B | -0.052004589 | 3.239565595 | -1.623931898 | 0.151908027 | 0.551100124 | -5.5575706 |
| **P00395** | COX1 | 0.123608298 | 3.20033523 | 1.417201368 | 0.202723299 | 0.608695713 | -5.55835661 |
| **Q92750-1** | TAF4B | 0.045356347 | 3.229575856 | 0.52514633 | 0.616983502 | 0.881353777 | -5.558485482 |
| **Q96NU7** | AMDHD1 | 0.047235239 | 3.160334312 | 0.545108911 | 0.603946114 | 0.873076436 | -5.55861758 |
| **Q9UHK6-5** | AMACR | 0.007873673 | 3.149822446 | 0.05821893 | 0.955330781 | 0.989472196 | -5.558637536 |
| **Q9NV79** | PCMTD2 | 0.063413055 | 3.064374541 | 0.82680492 | 0.437766038 | 0.786862931 | -5.559206031 |
| **O15085-2** | ARHGEF11 | 0.071152708 | 3.151856965 | 0.762035368 | 0.472890819 | 0.808597546 | -5.559661635 |
| **Q6ZSC3** | RBM43 | -0.04603408 | 3.185267511 | -0.48306818 | 0.64495853 | 0.891546795 | -5.560280272 |
| **Q5T4D3-3** | TMTC4 | 0.049948487 | 3.153888936 | 0.707150672 | 0.504131625 | 0.828319674 | -5.560411953 |
| **O43566-6** | RGS14 | -0.057572866 | 3.223386889 | -0.714064121 | 0.50012275 | 0.825209342 | -5.561651792 |
| **Q99973** | TEP1 | 0.000699061 | 3.147484622 | 0.0073636 | 0.994346558 | 0.997608555 | -5.561865142 |
| **Q08211** | DHX9 | 0.020585319 | 3.190498913 | 2.730960763 | 0.031647457 | 0.33673269 | -5.562220904 |

| | | | | | | | |
|---|---|---|---|---|---|---|---|
| **Q9Y303** | AMDHD2 | 0.063929211 | 3.125951858 | 1.343715499 | 0.224240375 | 0.629881158 | -5.562346541 |
| **Q9Y5K3** | PCYT1B | -0.048320325 | 3.083231328 | -0.733645917 | 0.48888278 | 0.817619097 | -5.562835433 |
| **Q9UNQ0-1** | ABCG2 | -0.055569806 | 3.212003048 | -0.56099629 | 0.593681281 | 0.869936882 | -5.562874398 |
| **Q9H2D6** | TRIOBP | -0.017206326 | 3.191032984 | -0.196168399 | 0.850495025 | 0.965552187 | -5.56300967 |
| **Q04837** | SSBP1 | 0.112642215 | 3.146492417 | 2.046726473 | 0.083207687 | 0.425585663 | -5.563224581 |
| **Q08623-4** | HDHD1; PUDP | 0.095921528 | 3.202120167 | 1.488411528 | 0.183672813 | 0.586056424 | -5.563289437 |
| **Q96QF0-1** | RAB3IP | 0.052585704 | 3.167383233 | 1.335349169 | 0.226815028 | 0.63274066 | -5.563424796 |
| **Q9NR33** | POLE4 | -0.096416986 | 3.169963812 | -1.182154436 | 0.278754156 | 0.676427621 | -5.563623848 |
| **P36894** | BMPR1A | 0.015423278 | 3.146308391 | 0.087567179 | 0.932868179 | 0.986793807 | -5.563704783 |
| **Q86XK3-2** | SFR1 | -0.058845743 | 3.172845806 | -1.215917137 | 0.266502527 | 0.669528065 | -5.565281658 |
| **P61970** | NUTF2 | 0.037847644 | 3.187653908 | 2.138681379 | 0.072961293 | 0.404796855 | -5.565378755 |
| **A0JLT2** | MED19 | -0.088998363 | 3.203074032 | -1.12541586 | 0.300413395 | 0.691292041 | -5.565918136 |
| **Q96E11** | MRRF | 0.058344398 | 3.187757062 | 1.506910916 | 0.178999818 | 0.580540072 | -5.566523895 |
| **Q8NCN5** | PDPR | 0.111833714 | 3.17951652 | 2.01723566 | 0.08679115 | 0.434776599 | -5.569018842 |
| **Q9NQ11** | ATP13A2 | -0.042383098 | 3.201908706 | -0.434947786 | 0.677733708 | 0.903662615 | -5.571007836 |
| **Q08AM6** | VAC14 | -0.050127538 | 3.181146192 | -1.933930119 | 0.097767512 | 0.456510811 | -5.571273908 |
| **Q9H4Z3** | PCIF1 | 0.038869335 | 3.19657118 | 1.982094743 | 0.091263184 | 0.444137581 | -5.571902284 |
| **P19544-7** | WT1 | -0.04665539 | 3.238820353 | -0.448315258 | 0.668548275 | 0.90001396 | -5.572003446 |
| **P61925** | PKIA | -0.404495939 | 3.153590241 | -1.000242888 | 0.353115366 | 0.732738778 | -5.572478718 |
| **Q9NPI7** | KRCC1 | 0.00061347 | 3.156295486 | 0.004442719 | 0.996589057 | 0.998480498 | -5.572497278 |
| **O14745** | SLC9A3R1 | -0.05670962 | 3.19317149 | -2.552768204 | 0.040567045 | 0.355540374 | -5.573437742 |
| **Q8TCT0** | CERK | -0.068213247 | 3.195739587 | -0.809688985 | 0.446863115 | 0.792204208 | -5.574014325 |
| **Q8IZ40** | RCOR2 | -0.107923674 | 3.207713148 | -1.450637955 | 0.193564418 | 0.598629202 | -5.574262112 |
| **Q03518** | TAP1 | -0.033535146 | 3.193827379 | -1.888393559 | 0.104338295 | 0.472077308 | -5.574287873 |
| **Q6DKI7** | PVRIG | -0.030112331 | 3.189983377 | -0.25169319 | 0.809082482 | 0.955498404 | -5.574391722 |
| **Q9BSG0** | PRADC1 | 0.012783374 | 3.168731669 | 0.059371724 | 0.954447473 | 0.989472196 | -5.574507648 |
| **Q68DD2-3** | PLA2G4F | -0.043509153 | 3.176523002 | -0.510565557 | 0.626602476 | 0.885387512 | -5.574890681 |
| **Q9NZL9** | MAT2B | -0.08477226 | 3.195565715 | -1.355793821 | 0.220569203 | 0.625865114 | -5.575214682 |
| **P59923** | ZNF445 | -0.057866097 | 3.106199515 | -0.47963789 | 0.647267858 | 0.89271426 | -5.576070547 |
| **A2RU49** | AGPHD1; HYKK | 0.053645764 | 3.166778447 | 1.1627563 | 0.286006678 | 0.681580409 | -5.576110978 |
| **P35475** | IDUA | 0.072220151 | 3.209832429 | 0.951603469 | 0.375475094 | 0.748219118 | -5.577709541 |
| **Q9UKR5** | C14orf1; ERG28 | 0.064335515 | 3.158080868 | 1.662880531 | 0.143781661 | 0.539341922 | -5.578093415 |
| **Q9P265** | DIP2B | -0.026240036 | 3.162625388 | -2.202599338 | 0.066600156 | 0.394427892 | -5.57883751 |
| **Q8IVB5** | LIX1L; LOC101060! | -0.042224916 | 3.185659513 | -1.445951641 | 0.194824985 | 0.600645673 | -5.579231318 |
| **Q14766-4** | LTBP1 | 0.037349665 | 3.154410252 | 0.348561968 | 0.738480615 | 0.930567563 | -5.579243757 |
| **Q9C037-1** | TRIM4 | -0.028387807 | 3.191019544 | -2.084313975 | 0.078854764 | 0.416709711 | -5.580535181 |
| **P63146** | UBE2B | -0.072031345 | 3.157335237 | -1.050568787 | 0.331097455 | 0.71677637 | -5.582047722 |
| **Q13443** | ADAM9 | 0.08059445 | 3.16608695 | 1.433874714 | 0.198108081 | 0.603708078 | -5.582329225 |
| **Q7Z6J8** | UBE3D | -0.083180592 | 3.197920115 | -0.826366806 | 0.437997232 | 0.786897076 | -5.582497045 |

| | | | | | | | |
|---|---|---|---|---|---|---|---|
| **Q9HBH9-1** | MKNK2 | 0.087001063 | 3.139721434 | 0.703013374 | 0.506540782 | 0.82913633 | -5.582870625 |
| **Q9C026** | TRIM9 | -0.069100505 | 3.168781515 | -1.023804843 | 0.342666365 | 0.724485042 | -5.583184974 |
| **Q6UWP2** | DHRS11 | 0.045988787 | 3.195153369 | 1.647865569 | 0.146864329 | 0.545761036 | -5.583399423 |
| **O43684** | BUB3 | 0.024024623 | 3.171742851 | 2.327633986 | 0.055743665 | 0.376021618 | -5.58346902 |
| **O15055-1** | PER2 | -0.052484026 | 3.230995683 | -1.269294389 | 0.24807523 | 0.652351802 | -5.583745106 |
| **P49419** | ALDH7A1 | 0.060073363 | 3.189133311 | 1.842194962 | 0.111448538 | 0.483014501 | -5.584316936 |
| **P35789** | ZNF93 | -0.036770616 | 3.141572813 | -0.373747228 | 0.720533754 | 0.925644897 | -5.585646108 |
| **Q9NX55-2** | HYPK | -0.178237852 | 3.196206573 | -1.438044702 | 0.19696883 | 0.603701501 | -5.586012937 |
| **Q5SQI0-1** | ATAT1 | -0.039452518 | 3.179355977 | -1.535274495 | 0.172048068 | 0.571776747 | -5.586207882 |
| **Q9H0R5** | GBP3 | -0.045734298 | 3.107314374 | -0.652792786 | 0.536379937 | 0.842162373 | -5.586293811 |
| **Q8N5Z5-1** | KCTD17 | 0.066310163 | 3.241256092 | 0.892747221 | 0.403966737 | 0.766336431 | -5.586303837 |
| **Q9NR28-1** | DIABLO | 0.074510409 | 3.171677945 | 1.56754201 | 0.164444909 | 0.56388209 | -5.587208421 |
| **P00747** | PLG | 0.005314593 | 3.205963944 | 0.04379334 | 0.966389491 | 0.992001409 | -5.589088687 |
| **Q9NVS2-2** | MRPS18A | 0.165457263 | 3.184186334 | 1.006686877 | 0.35023297 | 0.731407657 | -5.590243368 |
| **Q8IXQ3** | C9orf40 | -0.152277923 | 3.198912178 | -1.641051477 | 0.148283915 | 0.546231571 | -5.590482092 |
| **Q8IYL2** | TRMT44 | -0.055226511 | 3.173963241 | -1.380328227 | 0.213276676 | 0.61901473 | -5.590680828 |
| **Q8N1B3-1** | FAM58A; CCNQ | -0.06114784 | 3.212038999 | -0.459037148 | 0.661225084 | 0.896934512 | -5.591085977 |
| **Q14254** | FLOT2 | -0.064401973 | 3.181359363 | -1.615840573 | 0.153649909 | 0.552139704 | -5.5914113 |
| **Q96SZ5** | ADO | 0.040321077 | 3.19894989 | 1.731267921 | 0.130505731 | 0.516883365 | -5.591759561 |
| **Q9UHL4** | DPP7 | 0.045487993 | 3.172876237 | 1.742478798 | 0.128444799 | 0.512665171 | -5.591768956 |
| **O75385** | ULK1 | 0.056740522 | 3.168575319 | 1.626298367 | 0.151402115 | 0.549766822 | -5.592148496 |
| **Q9BRP4** | PAAF1 | -0.093101612 | 3.118722025 | -1.728379804 | 0.131041798 | 0.517978495 | -5.592503193 |
| **Q9H840** | GEMIN7 | -0.050503322 | 3.166785572 | -0.824015162 | 0.439239692 | 0.787933926 | -5.592764972 |
| **Q7Z449-1** | CYP2U1 | -0.066945604 | 3.202439096 | -1.489364157 | 0.183429458 | 0.58600472 | -5.592961543 |
| **Q5JU69-1** | TOR2A | 0.070278339 | 3.151853134 | 0.950306898 | 0.376085773 | 0.748872548 | -5.593924768 |
| **Q8WZ82** | OVCA2 | 0.043229575 | 3.20157989 | 1.566486786 | 0.16468851 | 0.563893388 | -5.593943241 |
| **P54710-2** | FXYD2 | -0.056214475 | 3.186208346 | -1.076313797 | 0.320267587 | 0.708257023 | -5.594093015 |
| **P78382** | SLC35A1 | 0.104912477 | 3.154884576 | 1.002562929 | 0.352075466 | 0.732738778 | -5.594427458 |
| **P04066** | FUCA1 | 0.045951593 | 3.177065307 | 1.563366891 | 0.16541072 | 0.564756413 | -5.594679742 |
| **O60733** | PLA2G6 | -0.049522794 | 3.169264152 | -0.51290227 | 0.625055534 | 0.885387512 | -5.594765824 |
| **Q96LU5** | IMMP1L | 0.049131211 | 3.160271923 | 0.670175051 | 0.525928332 | 0.838233243 | -5.596622931 |
| **Q8NFZ8** | CADM4 | -0.052474713 | 3.187715629 | -0.837201638 | 0.432305336 | 0.783333621 | -5.597236419 |
| **P49441** | INPP1 | 0.060818103 | 3.181992178 | 0.870083498 | 0.415359311 | 0.77203561 | -5.598317531 |
| **Q9BRQ6** | CHCHD6 | 0.109847297 | 3.22713314 | 1.780862836 | 0.121623814 | 0.49982876 | -5.598412706 |
| **Q9UI14** | RABAC1 | 0.059554116 | 3.174124806 | 0.95678704 | 0.373041271 | 0.745722908 | -5.598456694 |
| **Q96LW9** | ZNF323; ZSCAN31 | 0.058537551 | 3.14787398 | 0.684203328 | 0.517588594 | 0.833280927 | -5.598767979 |
| **Q9Y281-1** | CFL2 | -0.007798845 | 3.230738607 | -0.060866791 | 0.953302008 | 0.989472196 | -5.598777556 |
| **O15061-1** | SYNM | -0.025478535 | 3.168640139 | -0.229675204 | 0.82543322 | 0.959077751 | -5.598979485 |
| **P82914** | MRPS15 | 0.111965768 | 3.18675381 | 1.640046118 | 0.14849446 | 0.546231571 | -5.599644473 |

| | | | | | | | |
|---|---|---|---|---|---|---|---|
| **Q8NDH3** | NPEPL1 | 0.070863852 | 3.178891748 | 1.092786404 | 0.313490512 | 0.701109468 | -5.600689241 |
| **Q8N531-1** | FBXL6 | -0.0537469 | 3.145885516 | -1.143095692 | 0.293518471 | 0.685934755 | -5.600849546 |
| **Q5THJ4** | VPS13D | 0.030910685 | 3.179161802 | 1.999415247 | 0.08903095 | 0.440116703 | -5.601044756 |
| **Q01469** | FABP5 | 0.039396471 | 3.172414203 | 2.447343903 | 0.047053415 | 0.360340657 | -5.601234144 |
| **P63272** | SUPT4H1 | 0.086007726 | 3.191761946 | 1.416482466 | 0.202924466 | 0.608695713 | -5.60126033 |
| **Q96GX2** | ATXN7L3B | 0.129305099 | 3.184460076 | 1.3035804 | 0.236831319 | 0.643510544 | -5.601451483 |
| **Q96NB2** | SFXN2 | 0.073651653 | 3.176706482 | 1.718211519 | 0.13294605 | 0.519961998 | -5.601463347 |
| **Q17RN3-1** | FAM98C | -0.053700205 | 3.171688556 | -0.664964591 | 0.529047628 | 0.839849063 | -5.601688275 |
| **Q9UMW8** | USP18 | -0.077818915 | 3.199161139 | -0.943405253 | 0.37934926 | 0.750854478 | -5.602096159 |
| **Q96D46** | NMD3 | -0.070038151 | 3.197846195 | -2.114532779 | 0.075522019 | 0.41013154 | -5.602221593 |
| **Q9NQT4** | EXOSC5 | -0.099957859 | 3.172488584 | -1.852705411 | 0.109790284 | 0.480862075 | -5.602553365 |
| **Q5T2T1** | MPP7 | 0.061486506 | 3.213031586 | 1.584310801 | 0.160618769 | 0.560635782 | -5.602815632 |
| **Q96N21-2** | ENTHD2; TEPSIN | -0.057962983 | 3.110451306 | -0.769284388 | 0.468865354 | 0.805725242 | -5.603439294 |
| **Q9NR55** | BATF3 | -0.008574489 | 3.160242048 | -0.068601799 | 0.947377714 | 0.989472196 | -5.603785389 |
| **Q8IW41** | MAPKAPK5 | -0.088505761 | 3.216034524 | -1.032060756 | 0.339063759 | 0.721920542 | -5.60471949 |
| **Q8N5K1** | CISD2 | -0.044915589 | 3.201206772 | -1.901063612 | 0.102467495 | 0.467580037 | -5.605110657 |
| **P49641** | MAN2A2 | -0.030210274 | 3.176753229 | -0.299375989 | 0.774037583 | 0.943158294 | -5.605504147 |
| **Q16643-3** | DBN1 | -0.033891234 | 3.185442568 | -2.178539201 | 0.068925691 | 0.399844221 | -5.605980822 |
| **Q643R3** | LPCAT4 | -0.077405923 | 3.20161802 | -1.09271754 | 0.313518597 | 0.701109468 | -5.607460834 |
| **Q86YV0** | RASAL3 | -0.062936576 | 3.202924372 | -1.903952199 | 0.102045631 | 0.46702335 | -5.607876798 |
| **P08174-7** | CD55 | -0.05922011 | 3.147689123 | -1.359823164 | 0.219356457 | 0.625602181 | -5.608526148 |
| **P17655-1** | CAPN2 | -0.020912693 | 3.214584225 | -0.219878906 | 0.832739055 | 0.960584638 | -5.608754829 |
| **P17483** | HOXB4 | -0.058734827 | 3.193884598 | -0.977191043 | 0.363579256 | 0.738781889 | -5.608798899 |
| **P61803** | DAD1 | -0.198606283 | 3.23369263 | -1.390973819 | 0.210180174 | 0.615470161 | -5.609147999 |
| **Q03405** | PLAUR | -0.053197179 | 3.167419942 | -0.932129787 | 0.384727498 | 0.754667212 | -5.609604257 |
| **Q15035** | TRAM2 | 0.081691723 | 3.219746726 | 0.723228556 | 0.494841175 | 0.822007449 | -5.610831628 |
| **P98169** | ZXDB | 0.019613281 | 3.241604241 | 0.203626651 | 0.844899045 | 0.962551891 | -5.610969366 |
| **Q13794** | PMAIP1 | -0.064087278 | 3.228221549 | -0.617605248 | 0.557931585 | 0.851117349 | -5.611004998 |
| **Q8NCM8-2** | DYNC2H1 | 0.023424913 | 3.103071098 | 0.20214753 | 0.846008076 | 0.963216816 | -5.611044271 |
| **Q14353-1** | GAMT | 0.107446695 | 3.203308487 | 1.808346762 | 0.116956901 | 0.492866972 | -5.611404288 |
| **P51809** | VAMP7 | -0.076650452 | 3.192738021 | -1.312617045 | 0.23394324 | 0.641144892 | -5.612211119 |
| **Q5T3F8** | TMEM63B | -0.019853994 | 3.20252411 | -0.220845458 | 0.83201741 | 0.960474408 | -5.612507761 |
| **Q86XP1** | DGKH | -0.048354707 | 3.205017627 | -1.552425808 | 0.167966861 | 0.566424749 | -5.612903806 |
| **Q00978** | IRF9 | 0.045633384 | 3.121573932 | 0.487401927 | 0.642047073 | 0.890981968 | -5.61317712 |
| **Q96GW9** | MARS2 | 0.1182027 | 3.211643751 | 1.875966679 | 0.10620584 | 0.475403904 | -5.613713671 |
| **Q9NRD1** | FBXO6 | -0.062390286 | 3.183739215 | -0.942381978 | 0.379834964 | 0.750948043 | -5.613924651 |
| **Q08AN1** | ZNF616 | -0.077044256 | 3.169144259 | -0.762059859 | 0.47287718 | 0.808597546 | -5.614114779 |
| **Q8NHU6** | TDRD7 | -0.048870415 | 3.201579529 | -1.78608583 | 0.120723221 | 0.499295152 | -5.614471689 |
| **Q9BRU9** | UTP23 | -0.058168546 | 3.162185744 | -1.472096167 | 0.187886986 | 0.591197824 | -5.615411396 |

| | | | | | | | |
|---|---|---|---|---|---|---|---|
| **P07919** | UQCRH | -0.294241106 | 3.192405732 | -1.044229617 | 0.333808899 | 0.719204247 | -5.616288143 |
| **Q9Y5E7** | PCDHB2 | -0.020794785 | 3.140028234 | -0.244381879 | 0.814500875 | 0.95620584 | -5.616524644 |
| **Q5T3J3** | LRIF1 | -0.059313714 | 3.16880257 | -0.746469743 | 0.481614365 | 0.81235548 | -5.616624373 |
| **O95147** | DUSP14 | -0.029940662 | 3.198541076 | -0.308862414 | 0.767130332 | 0.940879515 | -5.616642592 |
| **Q96Q11-1** | TRNT1 | 0.042961263 | 3.188611295 | 1.938889026 | 0.097077139 | 0.455254009 | -5.617007947 |
| **Q8N9Z2-1** | CCDC71L | -0.049407653 | 3.198632219 | -0.435587005 | 0.677293091 | 0.903662615 | -5.617275716 |
| **Q96NT3-3** | GUCD1 | 0.012765726 | 3.160006272 | 0.06443706 | 0.950567091 | 0.989472196 | -5.617324512 |
| **Q6ZXV5** | TMTC3 | -0.064244796 | 3.15916267 | -1.532905482 | 0.172618976 | 0.572872916 | -5.617420582 |
| **Q9UBU6** | FAM8A1 | -0.029333464 | 3.211478911 | -0.373622884 | 0.720621903 | 0.925644897 | -5.617700072 |
| **Q07890** | SOS 2 | 0.054909692 | 3.158364566 | 1.325882836 | 0.229759759 | 0.637461728 | -5.617740299 |
| **Q86V85** | GPR180 | 0.002544453 | 3.227349018 | 0.022840541 | 0.982465634 | 0.994932161 | -5.618372519 |
| **Q9NXR1-1** | NDE1 | -0.041536797 | 3.177490669 | -1.902470512 | 0.10226181 | 0.467205335 | -5.618925092 |
| **Q9NXR5** | ANKRD10 | -0.046392721 | 3.141194761 | -1.263374961 | 0.250062882 | 0.654603944 | -5.619746642 |
| **Q5T1R4-1** | HIVEP3 | -0.046219695 | 3.197194524 | -0.506048622 | 0.629598567 | 0.886324146 | -5.620151194 |
| **Q9H867-4** | METTL21D; VCPKM | 0.026736299 | 3.170404355 | 0.302079051 | 0.772067125 | 0.942543532 | -5.620749779 |
| **Q5HYA8** | TMEM67 | 0.057497036 | 3.199168197 | 0.737139478 | 0.486895391 | 0.815942602 | -5.621254214 |
| **Q9HCK5** | AGO4; EIF2C4 | 0.05536264 | 3.15736295 | 1.172369035 | 0.282393062 | 0.679496093 | -5.621495349 |
| **Q63HQ0** | AP1AR | -0.06123659 | 3.163413049 | -1.243018315 | 0.257004114 | 0.659782078 | -5.622048901 |
| **O75177** | SS18L1 | 0.061619237 | 3.190726089 | 1.225873136 | 0.262978706 | 0.66662532 | -5.622260545 |
| **Q9Y2V7-1** | COG6 | -0.046311327 | 3.184960219 | -1.581188445 | 0.161324818 | 0.561414897 | -5.622584341 |
| **O43752** | STX6 | 0.072059546 | 3.149820958 | 1.284253741 | 0.243113295 | 0.649360189 | -5.622689774 |
| **Q86YS7-3** | C2CD5; KIAA0528 | 0.034375169 | 3.153477732 | 2.1273794 | 0.074148638 | 0.406910627 | -5.622707148 |
| **Q9UQL6-3** | HDAC5 | -0.018344475 | 3.15833561 | -0.165622422 | 0.873508368 | 0.97237605 | -5.622752737 |
| **Q9Y666** | SLC12A7 | -0.057210295 | 3.238852647 | -1.504223388 | 0.179671818 | 0.580990067 | -5.62420889 |
| **Q9H967** | WDR76 | -0.044266336 | 3.183052453 | -2.028016807 | 0.085463539 | 0.431389971 | -5.625171337 |
| **O95625** | ZBTB11 | -0.042135418 | 3.205857704 | -2.04189549 | 0.083784385 | 0.426983283 | -5.625241655 |
| **Q9P000-1** | COMMD9 | -0.043490647 | 3.182650664 | -1.675722573 | 0.141194046 | 0.536483353 | -5.625369337 |
| **Q9H992** | 7-Mar | -0.018359537 | 3.188101283 | -0.202020647 | 0.84610323 | 0.963216816 | -5.625795389 |
| **Q01094** | E2F1 | 0.103178302 | 3.205222368 | 0.935909706 | 0.382918089 | 0.754258979 | -5.627318117 |
| **P52741-1** | ZNF134 | -0.052188969 | 3.256200436 | -0.561664428 | 0.593251781 | 0.869935985 | -5.627593915 |
| **Q7L622** | G2E3 | -0.032205866 | 3.244008937 | -0.352240102 | 0.735848157 | 0.929681266 | -5.628731951 |
| **O43299** | AP5Z1 | 0.092799181 | 3.209673308 | 1.503704166 | 0.179801914 | 0.581174211 | -5.629180939 |
| **Q6PCB7** | SLC27A1 | 0.053632499 | 3.143220904 | 0.65722425 | 0.533703056 | 0.840132297 | -5.629467363 |
| **Q9NRX5** | SERINC1 | 0.049602599 | 3.167421794 | 0.360910926 | 0.729657745 | 0.927974736 | -5.62953344 |
| **L0R6Q1** | SLC35A4 | 0.136710155 | 3.216154454 | 1.5185266 | 0.176122016 | 0.577741296 | -5.630269147 |
| **P41223** | BUD31 | 0.075293157 | 3.160382989 | 1.529674736 | 0.173400387 | 0.574167672 | -5.630539812 |
| **Q9BS40** | LXN | 0.039376988 | 3.145041567 | 0.282564798 | 0.786332705 | 0.946749713 | -5.630583884 |
| **Q5JTH9-1** | RRP12 | -0.081678977 | 3.191859222 | -2.344026196 | 0.054461755 | 0.376021618 | -5.63150668 |
| **Q14159** | KIAA0146; SPIDR | -0.053081933 | 3.180639611 | -1.089389348 | 0.314878436 | 0.701837878 | -5.632701523 |

| | | | | | | | |
|---|---|---|---|---|---|---|---|
| **Q9H0I2-1** | ENKD1 | -0.056727907 | 3.109747195 | -0.894414043 | 0.40313812 | 0.766336431 | -5.633081503 |
| **Q9NX47** | 5-Mar | 0.075844692 | 3.186161283 | 1.639720227 | 0.14856277 | 0.546231571 | -5.633824796 |
| **Q9NZL9-2** | MAT2B | 0.085336479 | 3.111871517 | 1.154786377 | 0.289032124 | 0.683685579 | -5.634149218 |
| **Q7L2E3-2** | DHX30 | 0.087831378 | 3.219313587 | 1.063345051 | 0.325686584 | 0.710703247 | -5.634907566 |
| **P32942** | ICAM3 | -0.047914582 | 3.152195515 | -1.555459731 | 0.167254383 | 0.566424749 | -5.635322273 |
| **Q7RTN6-1** | STRADA | -0.048107578 | 3.201098224 | -1.562683356 | 0.165569344 | 0.564812553 | -5.635820604 |
| **Q15118** | PDK1 | 0.138300393 | 3.213789618 | 1.610353048 | 0.15484196 | 0.553156191 | -5.636111323 |
| **Q13616** | CUL1 | -0.047650322 | 3.176293196 | -2.265777945 | 0.060867288 | 0.383378328 | -5.63638797 |
| **O95819-6** | MAP4K4 | 0.040636059 | 3.154867284 | 2.00968194 | 0.087733599 | 0.437017835 | -5.636563449 |
| **O94886** | TMEM63A | 0.007735144 | 3.164729175 | 0.041148397 | 0.968418066 | 0.992306703 | -5.6369866 |
| **Q6P444** | FAM54A; MTFR2 | -0.046946378 | 3.178323718 | -1.424986015 | 0.200556495 | 0.606094087 | -5.638387344 |
| **O43663-1** | PRC1 | -0.054346184 | 3.182200434 | -0.934062376 | 0.383801575 | 0.754456977 | -5.638743353 |
| **P53567** | CEBPG | -0.032016842 | 3.273784939 | -0.375979477 | 0.718952078 | 0.925182096 | -5.639121112 |
| **Q12899** | TRIM26 | -0.044021387 | 3.18107898 | -1.93854646 | 0.097124675 | 0.455254009 | -5.639232799 |
| **Q5VWQ0** | RSBN1 | -0.054779384 | 3.180279469 | -1.912048671 | 0.10087229 | 0.463471034 | -5.639333326 |
| **Q9NRG0** | CHRAC1 | -0.046302823 | 3.174553545 | -1.229759848 | 0.261613933 | 0.665340172 | -5.639726964 |
| **P52907** | CAPZA1 | 0.037566045 | 3.198378388 | 2.228201024 | 0.064213317 | 0.390938548 | -5.640144348 |
| **P43243** | MATR3 | 0.033554379 | 3.181325938 | 2.347653941 | 0.054182189 | 0.376021618 | -5.640821908 |
| **Q9BXM9-1** | FSD1L | -0.054950486 | 3.186074878 | -1.64880319 | 0.146670006 | 0.545761036 | -5.641537273 |
| **Q8TEQ8-1** | PIGO | -0.071356348 | 3.208305863 | -1.397242018 | 0.208375892 | 0.614231901 | -5.641692304 |
| **Q71RG4-1** | TMUB2 | 0.051038132 | 3.163399106 | 0.88315472 | 0.408760053 | 0.768662396 | -5.642094978 |
| **Q99963-3** | SH3GL3 | 0.058660453 | 3.180298275 | 0.511606533 | 0.625913079 | 0.885387512 | -5.642131098 |
| **Q9H0B6** | KLC2 | 0.030288137 | 3.200457648 | 1.921448706 | 0.099526744 | 0.459999988 | -5.642455503 |
| **Q86WV6** | TMEM173 | 0.066311992 | 3.187527795 | 0.886397577 | 0.407134915 | 0.767181929 | -5.643681374 |
| **Q9P287-2** | BCCIP | 0.063997194 | 3.168186626 | 1.028940923 | 0.340421584 | 0.722975002 | -5.644364417 |
| **Q99470** | SDF2 | 0.056457366 | 3.19931302 | 1.37138791 | 0.215908706 | 0.621519808 | -5.644409083 |
| **O95400** | CD2BP2 | -0.072990486 | 3.163774485 | -1.785474149 | 0.120828356 | 0.499469974 | -5.64499592 |
| **P0C1Z6** | TFPT | -0.040717297 | 3.181038042 | -1.565377193 | 0.164945026 | 0.563893388 | -5.645956985 |
| **Q9Y608** | LRRFIP2 | -0.054524365 | 3.149120254 | -1.538563516 | 0.171258357 | 0.571100234 | -5.646003228 |
| **Q6PJG6-1** | BRAT1 | -0.041929693 | 3.160062209 | -1.87932079 | 0.10569856 | 0.475403904 | -5.646246113 |
| **Q9NYZ1** | FAM18B1; TVP23B | 0.080841198 | 3.175775349 | 1.154285342 | 0.289223214 | 0.683892392 | -5.646408649 |
| **A6NM62** | LRRC53; LOC1053 | -0.058728551 | 3.251948553 | -0.256790593 | 0.805311531 | 0.954249678 | -5.64641979 |
| **P81274** | GPSM2 | -0.060114996 | 3.14127395 | -1.034314644 | 0.338085506 | 0.721894046 | -5.648312181 |
| **Q9BSF8-2** | BTBD10 | -0.070173302 | 3.221642383 | -1.261677198 | 0.250635511 | 0.65503261 | -5.649596543 |
| **P51648-2** | ALDH3A2 | -0.05908043 | 3.150104205 | -0.656412007 | 0.534193072 | 0.840262117 | -5.650045601 |
| **Q2TAA5** | ALG11 | -0.072550644 | 3.196475809 | -1.255547741 | 0.25271236 | 0.656572825 | -5.650331947 |
| **P35251-1** | RFC1 | -0.066844933 | 3.174617501 | -2.145683539 | 0.072235351 | 0.404094866 | -5.650426895 |
| **Q9GZP9** | DERL2 | 0.049345417 | 3.18292374 | 0.737885829 | 0.486471518 | 0.815404264 | -5.650887307 |
| **Q5JY77** | GPRASP1 | -0.049903847 | 3.198762952 | -0.659066888 | 0.532592466 | 0.839951983 | -5.651276052 |

| | | | | | | | |
|---|---|---|---|---|---|---|---|
| **Q02833-1** | RASSF7 | -0.053059908 | 3.204702251 | -0.908352733 | 0.396258508 | 0.762021989 | -5.652324049 |
| **P46940** | IQGAP1 | -0.034494498 | 3.162944189 | -2.531460177 | 0.04179836 | 0.355540374 | -5.652793249 |
| **Q9H9D4** | ZNF408 | -0.056147989 | 3.20510591 | -0.904573233 | 0.398115167 | 0.762880385 | -5.65391121 |
| **Q8NHQ1-1** | CEP70 | 0.0479642 | 3.183256808 | 1.300902371 | 0.237693206 | 0.644090219 | -5.654264289 |
| **Q15758-1** | SLC1A5 | -0.059253014 | 3.231682569 | -1.178422374 | 0.2801373 | 0.677109587 | -5.654869157 |
| **Q14764** | MVP | 0.047772353 | 3.131770576 | 0.365110694 | 0.72666722 | 0.92710315 | -5.6551551 |
| **O43502** | RAD51C | -0.07786283 | 3.213257034 | -1.248655332 | 0.255065511 | 0.659782078 | -5.655277819 |
| **Q9Y4X0** | AMMECR1 | -0.027253864 | 3.217579898 | -0.335878687 | 0.747587245 | 0.934974996 | -5.655356188 |
| **Q9UJ55** | MAGEL2 | 0.008573481 | 3.139202211 | 0.1048432 | 0.919675101 | 0.983742622 | -5.655631118 |
| **Q96AA3** | RFT1 | 0.027097178 | 3.120853475 | 0.335016945 | 0.748207594 | 0.935183323 | -5.655793683 |
| **Q9UBF6** | RNF7 | -0.061278568 | 3.179919558 | -1.077314906 | 0.319852336 | 0.708257023 | -5.656184364 |
| **Q9NS37** | CREBZF | -0.067071502 | 3.188653981 | -1.328245822 | 0.229021544 | 0.636883503 | -5.656570537 |
| **Q8WUM0** | NUP133 | -0.03401095 | 3.188018291 | -2.329215684 | 0.055618628 | 0.376021618 | -5.657427198 |
| **Q9BXS9-1** | SLC26A6 | 0.05357105 | 3.163716729 | 1.006319602 | 0.350396751 | 0.731407657 | -5.657975269 |
| **Q53GI3** | ZNF394 | -0.017414908 | 3.167671576 | -0.158016023 | 0.879260829 | 0.97338977 | -5.659689796 |
| **Q5T6J7-2** | IDNK | 0.052754334 | 3.180567811 | 1.172341022 | 0.282403537 | 0.679496093 | -5.659813131 |
| **P08047** | SP1 | 0.050457062 | 3.199295217 | 1.808904655 | 0.116863995 | 0.492826136 | -5.65998058 |
| **Q96MG8-1** | PCMTD1 | 0.047870262 | 3.115911497 | 1.517807534 | 0.176298916 | 0.577844424 | -5.660381705 |
| **Q9NR12-1** | PDLIM7 | -0.047374828 | 3.244604554 | -1.248404185 | 0.255151612 | 0.659782078 | -5.660595535 |
| **Q6P2D0-1** | ZFP1 | 0.077179109 | 3.190868788 | 0.976193417 | 0.364037512 | 0.739334875 | -5.660627105 |
| **P07737** | PFN1 | 0.030233087 | 3.169142345 | 2.620993907 | 0.036873958 | 0.348943902 | -5.66077037 |
| **Q9H3R0** | KDM4C | -0.047993942 | 3.191442308 | -1.434457827 | 0.197948413 | 0.603708078 | -5.661456546 |
| **Q8TC44-1** | POC1B | -0.012424702 | 3.155314535 | -0.122216769 | 0.906435282 | 0.98075504 | -5.661494868 |
| **Q8IUF1** | CBWD2 | -0.058340802 | 3.19398141 | -1.210209591 | 0.268540819 | 0.67097152 | -5.661856255 |
| **Q8NHG8** | ZNRF2 | -0.066454088 | 3.217749159 | -1.297179773 | 0.238895847 | 0.644491512 | -5.662598958 |
| **Q14249** | ENDOG | 0.107929532 | 3.178215203 | 1.394823692 | 0.209070342 | 0.614302382 | -5.662666647 |
| **P0C7X2-1** | ZNF688 | -0.049201774 | 3.158584007 | -0.720934308 | 0.496159887 | 0.822932915 | -5.663921427 |
| **Q9H2Y7** | ZFP106; ZNF106 | -0.036882986 | 3.214848528 | -1.767357775 | 0.12398277 | 0.506448898 | -5.664890425 |
| **Q5T2R2-1** | PDSS1 | 0.137692233 | 3.230402322 | 1.79434273 | 0.119312684 | 0.498071065 | -5.665009657 |
| **Q9NQG7-4** | HPS4 | 0.050041094 | 3.202087616 | 1.3609782 | 0.219009915 | 0.625602181 | -5.665238906 |
| **Q9H079** | KATNBL1 | -0.118971064 | 3.168761352 | -1.232605369 | 0.260618625 | 0.664084341 | -5.666103622 |
| **Q9NPH0-1** | ACP6 | 0.085284176 | 3.153348965 | 1.562692569 | 0.165567205 | 0.564812553 | -5.666528946 |
| **Q9P035** | PTPLAD1; HACD3 | -0.079654666 | 3.170437752 | -1.655634664 | 0.145261526 | 0.543107211 | -5.667340424 |
| **Q16394** | EXT1 | 0.048534827 | 3.192908225 | 0.987768956 | 0.358747939 | 0.735928834 | -5.667457458 |
| **Q14789-2** | GOLGB1 | -0.059783137 | 3.151428724 | -0.970706106 | 0.366566127 | 0.741119482 | -5.667952595 |
| **Q92993-3** | KAT5 | 0.05790796 | 3.199085272 | 1.664765167 | 0.143399107 | 0.539313633 | -5.668070068 |
| **O75934** | BCAS2 | -0.027046918 | 3.191388028 | -1.96296989 | 0.093792759 | 0.448493945 | -5.668131099 |
| **Q9BXP2-1** | SLC12A9 | 0.051274907 | 3.286258549 | 0.489846085 | 0.640408077 | 0.890256071 | -5.668571995 |
| **Q9NVX2** | NLE1 | -0.039252885 | 3.19457779 | -1.824187412 | 0.114346736 | 0.488695437 | -5.669034185 |

| | | | | | | | |
|---|---|---|---|---|---|---|---|
| **Q99571-2** | P2RX4 | 0.046641479 | 3.233084148 | 0.7247514 | 0.493967147 | 0.821555157 | -5.669279217 |
| **O14967** | CLGN | -0.030224026 | 3.187715031 | -2.109536805 | 0.076063038 | 0.411304347 | -5.669891318 |
| **Q7Z7N9** | TMEM179B | -0.086448482 | 3.130106157 | -0.667724922 | 0.527393667 | 0.839849063 | -5.669898239 |
| **P49356** | FNTB; CHURC1-FN | 0.100645406 | 3.208283967 | 0.915264651 | 0.392879919 | 0.760129045 | -5.669932877 |
| **Q9HCK1** | ZDBF2 | 0.078439765 | 3.258664971 | 0.448951159 | 0.668112837 | 0.90001396 | -5.670899865 |
| **Q6NSZ9** | ZNF498; ZSCAN25 | 0.085833016 | 3.227544799 | 1.250641272 | 0.254385551 | 0.659262133 | -5.67132163 |
| **Q9UFN0** | NIPSNAP3A | 0.167368372 | 3.2106489 | 1.775140077 | 0.122618062 | 0.50211742 | -5.672122557 |
| **Q8IUQ4-2** | SIAH1 | -0.047078045 | 3.195836698 | -0.841333103 | 0.430149036 | 0.782603406 | -5.672356874 |
| **Q9BWL3** | C1orf43 | -0.072368743 | 3.195421433 | -0.835556333 | 0.433166224 | 0.783962934 | -5.672366711 |
| **P10412** | HIST1H1E | 0.007626396 | 3.247838368 | 0.04872809 | 0.962605453 | 0.991283985 | -5.672438714 |
| **Q9H400** | LIME1 | -0.075899644 | 3.222882467 | -1.136786721 | 0.295963621 | 0.688880164 | -5.672819058 |
| **P22061-2** | PCMT1 | 0.104323278 | 3.187540096 | 2.05726671 | 0.081963236 | 0.423947857 | -5.67312446 |
| **O94916-5** | NFAT5 | -0.054634603 | 3.19694942 | -1.185021423 | 0.277695545 | 0.676093886 | -5.67322037 |
| **P36954** | POLR2I | -0.375811871 | 3.21443097 | -1.49787314 | 0.181268918 | 0.58362594 | -5.674642962 |
| **Q7Z6K1** | THAP5 | -0.058918165 | 3.188991099 | -0.790950178 | 0.456975387 | 0.798548386 | -5.674889112 |
| **Q5T7V8** | GORAB | -0.056906685 | 3.208324221 | -0.96291544 | 0.370179535 | 0.744424726 | -5.674962036 |
| **Q9NZJ5** | EIF2AK3 | 0.055658389 | 3.176091043 | 1.025829878 | 0.341779898 | 0.724080487 | -5.675230839 |
| **O14672** | ADAM10 | 0.057557853 | 3.161632806 | 2.175760392 | 0.069199547 | 0.399872475 | -5.675444955 |
| **Q6ZN55-1** | ZNF574 | -0.070264026 | 3.187719518 | -1.833402348 | 0.11285457 | 0.486505458 | -5.675512208 |
| **Q9BUU2-1** | METTL22 | 0.019330156 | 3.152373313 | 0.185658148 | 0.858396829 | 0.968855822 | -5.675699794 |
| **Q9UBV7** | B4GALT7 | -0.026732963 | 3.178218008 | -0.29356526 | 0.778279578 | 0.944053983 | -5.67581859 |
| **Q9H8P0** | SRD5A3 | 0.080348568 | 3.161284113 | 1.142226981 | 0.293854151 | 0.686466107 | -5.676152639 |
| **Q99550** | MPHOSPH9 | -0.082583041 | 3.196444881 | -1.406625838 | 0.205700861 | 0.611865721 | -5.676301392 |
| **Q504T8** | MIDN | -0.034390062 | 3.204490497 | -0.370056343 | 0.723152226 | 0.926648409 | -5.676308082 |
| **Q5JPF3** | ANKRD36C | -0.11331744 | 3.128851136 | -1.093197762 | 0.313322785 | 0.701109468 | -5.676587642 |
| **Q14511-1** | NEDD9 | -0.046498051 | 3.19865498 | -1.169517726 | 0.283460892 | 0.679787821 | -5.676904875 |
| **O43505** | B3GNT1; B4GAT1 | 0.046791434 | 3.193954616 | 0.648653079 | 0.53888817 | 0.843540088 | -5.677978072 |
| **Q92878-2** | RAD50 | 0.022825034 | 3.166608446 | 2.316471027 | 0.056634395 | 0.377614679 | -5.678233157 |
| **Q969W3-2** | FAM104A | 0.025363935 | 3.246455771 | 0.270904572 | 0.794900027 | 0.949994286 | -5.678234332 |
| **Q9Y5L0-1** | TNPO3 | -0.038565316 | 3.181020125 | -2.218971753 | 0.065063531 | 0.393401641 | -5.678450342 |
| **Q6ZV73-1** | FGD6 | -0.034284173 | 3.195934175 | -0.463390156 | 0.658263356 | 0.896852471 | -5.678550331 |
| **Q9NPA3** | MID1IP1 | -0.050852486 | 3.236996192 | -1.433487247 | 0.198214241 | 0.603708078 | -5.678781707 |
| **Q9H116** | GZF1 | -0.053263954 | 3.234190277 | -1.221482269 | 0.264527853 | 0.66823064 | -5.678788161 |
| **Q86WX3** | RPS19BP1 | -0.070519903 | 3.157745312 | -1.264829734 | 0.249573113 | 0.65453932 | -5.678862623 |
| **Q6P6B7-1** | ANKRD16 | 0.065686212 | 3.179114565 | 1.018120018 | 0.345164691 | 0.726061285 | -5.678867492 |
| **Q13315** | ATM | 0.031786643 | 3.184017128 | 2.143829795 | 0.07242682 | 0.404094866 | -5.679566448 |
| **O75419-3** | CDC45 | -0.040305673 | 3.170623821 | -1.933045121 | 0.097891231 | 0.456510811 | -5.679691325 |
| **Q9HBM6** | TAF9B | 0.053730286 | 3.180546391 | 1.513986507 | 0.177241699 | 0.578452275 | -5.679880289 |
| **Q8WU67** | ABHD3 | -0.032989111 | 3.171933013 | -0.354581862 | 0.734174158 | 0.928971761 | -5.679918799 |

| | | | | | | | |
|---|---|---|---|---|---|---|---|
| **P33897** | ABCD1 | 0.052574196 | 3.169115395 | 0.969358464 | 0.367189216 | 0.741942605 | -5.679937644 |
| **P38435-1** | GGCX | -0.05680601 | 3.148813335 | -0.671778065 | 0.52497103 | 0.837910607 | -5.680823777 |
| **Q8NDX6** | ZNF740 | -0.056517624 | 3.152452512 | -1.382293838 | 0.212701871 | 0.618337492 | -5.681673686 |
| **P18621-3** | RPL17-C18ORF32; | -0.109827462 | 3.179833678 | -2.297476904 | 0.058183813 | 0.379089796 | -5.683047961 |
| **Q8IWJ2** | GCC2 | -0.032153597 | 3.192123618 | -2.330320458 | 0.055531464 | 0.376021618 | -5.68326072 |
| **Q5U3C3** | TMEM164 | 0.072118655 | 3.212261964 | 0.74244851 | 0.483885654 | 0.814028593 | -5.683333233 |
| **Q8N5D0-1** | WDTC1 | 0.055221991 | 3.137015947 | 0.839763229 | 0.43096747 | 0.782692325 | -5.683795745 |
| **O76064** | RNF8 | -0.062757965 | 3.214001201 | -1.344182342 | 0.224097476 | 0.629881158 | -5.684015199 |
| **Q9NRF2-2** | SH2B1 | -0.054643452 | 3.160102789 | -0.630633297 | 0.54989121 | 0.846638465 | -5.685787314 |
| **P78345** | RPP38 | -0.036784678 | 3.162179807 | -1.60248916 | 0.15656543 | 0.554834464 | -5.68595573 |
| **Q9BQ39** | DDX50 | -0.037524778 | 3.176809282 | -2.216225791 | 0.065318705 | 0.393496543 | -5.686081177 |
| **P15907** | ST6GAL1 | 0.050771292 | 3.112241991 | 0.835577822 | 0.433154972 | 0.783962934 | -5.686097124 |
| **A7E2V4-2** | ZSWIM8 | 0.074254945 | 3.177633617 | 1.081829835 | 0.317985029 | 0.705893003 | -5.687009921 |
| **Q8TET4** | GANC | 0.057326783 | 3.17549353 | 0.88183972 | 0.409420426 | 0.769470301 | -5.687377338 |
| **O15315-2** | RAD51B | 0.015047126 | 3.173059667 | 0.154042077 | 0.882269377 | 0.97451158 | -5.687692394 |
| **Q96DT6** | ATG4C | -0.047878053 | 3.175507344 | -0.922411113 | 0.389409581 | 0.75855365 | -5.688059387 |
| **Q9BT30** | ALKBH7 | 0.087238424 | 3.191164488 | 0.557827515 | 0.595720674 | 0.871378088 | -5.688685448 |
| **Q9H2D1** | SLC25A32 | 0.110084121 | 3.232567681 | 1.549917648 | 0.168558004 | 0.566974321 | -5.688973099 |
| **Q9UM00-1** | TMCO1 | -0.051990223 | 3.198304694 | -1.427596908 | 0.199834485 | 0.605177234 | -5.689263332 |
| **Q96ED9-2** | HOOK2 | 0.041386848 | 3.206472769 | 1.34406529 | 0.224133298 | 0.629881158 | -5.691124353 |
| **Q99418** | CYTH2 | -0.062119328 | 3.183013303 | -1.131544972 | 0.298008028 | 0.689670498 | -5.691791713 |
| **P35611-3** | ADD1 | -0.023808439 | 3.211031375 | -2.088131619 | 0.078425673 | 0.415949247 | -5.692283322 |
| **Q08209-1** | PPP3CA | 0.054553785 | 3.121422518 | 0.415770736 | 0.691015793 | 0.909522034 | -5.693832123 |
| **Q5FWF5-1** | ESCO1 | -0.052850675 | 3.169049904 | -1.076510023 | 0.320186159 | 0.708257023 | -5.694286087 |
| **Q9BT43** | POLR3GL; LOC101 | 0.059492618 | 3.197841766 | 1.10384182 | 0.309008297 | 0.696978291 | -5.694628971 |
| **O95271** | TNKS | -0.049428837 | 3.148406216 | -1.244920022 | 0.256348689 | 0.659782078 | -5.695491801 |
| **Q14669-2** | TRIP12 | -0.044386157 | 3.187289607 | -0.527761891 | 0.615266561 | 0.880583268 | -5.695609953 |
| **O75123** | ZNF623 | 0.006243279 | 3.209591947 | 0.052817227 | 0.959470633 | 0.990513797 | -5.695794301 |
| **Q9NQW6** | ANLN | -0.048980388 | 3.178562761 | -1.945513736 | 0.096162388 | 0.454227211 | -5.695871061 |
| **Q9NQS1** | AVEN | 0.049383257 | 3.204108428 | 1.783375853 | 0.121189688 | 0.499487166 | -5.696151501 |
| **Q14676** | MDC 1.00 | -0.040525711 | 3.201891241 | -2.206564358 | 0.066224657 | 0.394427892 | -5.69660543 |
| **O94941-1** | UBOX5 | -0.01669647 | 3.175602196 | -0.214142556 | 0.837025513 | 0.960912428 | -5.69702521 |
| **Q13472-1** | TOP3A | -0.034314333 | 3.150984213 | -1.938639196 | 0.097111805 | 0.455254009 | -5.697069731 |
| **Q96JM2-3** | ZNF462 | 0.057837783 | 3.205691465 | 1.40513268 | 0.20612443 | 0.612058044 | -5.697081104 |
| **Q13428-7** | TCOF1 | 0.050377911 | 3.190647165 | 0.974193983 | 0.364957299 | 0.740073081 | -5.697200688 |
| **Q9H0V1** | TMEM168 | 0.039353854 | 3.234265814 | 0.400112983 | 0.701949293 | 0.914845949 | -5.697415792 |
| **Q5BKX5-1** | C19orf54 | 0.043133053 | 3.169606638 | 0.779098743 | 0.463453063 | 0.802800142 | -5.697724632 |
| **O15530-1** | PDPK1 | 0.055817466 | 3.228805609 | 0.922764919 | 0.389238376 | 0.75855365 | -5.698325963 |
| **Q9BQ48** | MRPL34 | 0.071701343 | 3.182671458 | 0.921712044 | 0.389748023 | 0.75855365 | -5.698622504 |

| | | | | | | | |
|---|---|---|---|---|---|---|---|
| **Q15853** | USF2 | 0.068281404 | 3.1972702 | 1.223056995 | 0.263971375 | 0.667489679 | -5.698636377 |
| **Q14678-1** | KANK1 | -0.053203955 | 3.257638587 | -1.262983388 | 0.250194853 | 0.654603944 | -5.698730264 |
| **Q9H4K7** | GTPBP5; MTG2 | 0.126992452 | 3.197994086 | 1.595496687 | 0.158113071 | 0.556829942 | -5.698776158 |
| **Q9NVR5** | DNAAF2 | -0.058850798 | 3.228776856 | -1.492559737 | 0.182615294 | 0.585267652 | -5.698864812 |
| **Q9H3S4** | TPK1 | 0.056215883 | 3.168142106 | 1.037169431 | 0.336849685 | 0.721128781 | -5.699093581 |
| **Q6PL18-1** | ATAD2 | -0.042069059 | 3.183887877 | -2.285856386 | 0.05915316 | 0.379928744 | -5.699150693 |
| **Q9BYX4-1** | IFIH1 | -0.046991057 | 3.188656587 | -1.87470113 | 0.106397864 | 0.475403904 | -5.699187128 |
| **Q86UF1** | TSPAN33 | 0.029024891 | 3.161804947 | 0.243137519 | 0.815424179 | 0.95624343 | -5.699203862 |
| **Q6PCD5** | RFWD3 | 0.04703659 | 3.191964062 | 1.573458126 | 0.163085388 | 0.561580888 | -5.699478508 |
| **Q9H668** | OBFC1; STN1 | 0.062786454 | 3.214199683 | 1.316786351 | 0.232621231 | 0.639692206 | -5.700003482 |
| **Q9H792** | PEAK1 | -0.047480429 | 3.18959042 | -0.581330326 | 0.580689911 | 0.863643082 | -5.701537033 |
| **O95479** | H6PD | 0.048699979 | 3.192716702 | 1.351210877 | 0.221955831 | 0.627993355 | -5.702393059 |
| **Q92794** | KAT6A | -0.042646437 | 3.200269768 | -2.046770991 | 0.083202391 | 0.425585663 | -5.702826025 |
| **Q8WWP7** | GIMAP1 | 0.040375366 | 3.169518908 | 1.869137343 | 0.107246123 | 0.475805886 | -5.702934685 |
| **Q96P50** | ACAP3 | 0.072182602 | 3.184263497 | 1.024290308 | 0.342453684 | 0.724485042 | -5.703968426 |
| **Q15628** | TRADD | -0.038534593 | 3.195243813 | -1.578884801 | 0.161847598 | 0.561414897 | -5.705717514 |
| **Q6AZZ1** | TRIM68 | 0.048927191 | 3.159781549 | 1.297878565 | 0.238669685 | 0.644491512 | -5.706514631 |
| **P50053-2** | KHK | -0.031774446 | 3.199352327 | -0.436835011 | 0.67643323 | 0.903662615 | -5.7065783 |
| **Q16842** | ST3GAL2 | 0.039843511 | 3.168384702 | 0.527107836 | 0.615695659 | 0.880583268 | -5.707363233 |
| **O75356** | ENTPD5 | 0.033448452 | 3.169175563 | 0.436634331 | 0.67657146 | 0.903662615 | -5.707637407 |
| **Q9UBZ9** | REV1 | -0.071690768 | 3.186708827 | -0.751459021 | 0.478806359 | 0.811537935 | -5.707696643 |
| **P06756** | ITGAV | 0.051326437 | 3.188293332 | 0.70628543 | 0.504634834 | 0.828715122 | -5.708013859 |
| **Q9H8T0-2** | AKTIP | -0.044958879 | 3.17029747 | -0.617487554 | 0.558004545 | 0.851117349 | -5.70879927 |
| **P02545** | LMNA | -0.153972525 | 0.640269801 | -0.586318291 | 0.577528561 | 0.861313941 | -5.709474795 |
| **Q13487** | SNAPC2 | -0.027757156 | 3.164859354 | -0.255509521 | 0.806258714 | 0.954293355 | -5.710806511 |
| **Q7Z698** | SPRED2 | -0.034323181 | 3.20445464 | -0.375981207 | 0.718950853 | 0.925182096 | -5.710828071 |
| **Q96T68-1** | SETDB2 | -0.05250547 | 3.181417361 | -0.591214528 | 0.57443519 | 0.85911412 | -5.710924505 |
| **Q9BTP7** | C19orf40; FAAP24 | -0.046992197 | 3.183788188 | -0.793572107 | 0.455550898 | 0.798084321 | -5.711092459 |
| **Q9GZX9** | TWSG1 | -0.0551791 | 3.213199551 | -0.659859354 | 0.532115281 | 0.839951983 | -5.711903833 |
| **Q8NCS4** | ZMYM6NB; TMEM | -0.019840885 | 3.13561484 | -0.182584195 | 0.860711278 | 0.968855822 | -5.712023718 |
| **Q15714-1** | TSC22D1 | 0.051812865 | 3.19729433 | 0.942111049 | 0.379963642 | 0.750948043 | -5.713648609 |
| **P18850** | ATF6 | -0.064671748 | 3.214277283 | -0.807998256 | 0.447768964 | 0.792674781 | -5.713742765 |
| **Q96FN4** | CPNE2 | 0.042365552 | 3.218949409 | 1.377390876 | 0.214138244 | 0.619843797 | -5.715084616 |
| **P98172** | EFNB1 | 0.008490697 | 3.181880104 | 0.102636152 | 0.921359104 | 0.984164603 | -5.715935327 |
| **Q8WZ73-1** | RFFL | 0.010152952 | 3.213841693 | 0.132806487 | 0.898380788 | 0.978757548 | -5.716223764 |
| **P61923-4** | COPZ1 | -0.051924103 | 3.193149455 | -1.52373791 | 0.174844872 | 0.574976204 | -5.716312387 |
| **Q96ME1** | FBXL18 | 0.057192467 | 3.164820322 | 1.432743866 | 0.198418061 | 0.603708078 | -5.716580518 |
| **Q9BPX5** | ARPC5L | 0.045830569 | 3.155147265 | 1.579621151 | 0.161680321 | 0.561414897 | -5.717115061 |
| **Q9NX05-1** | FAM120C | 0.042895174 | 3.18595171 | 1.792715144 | 0.119589454 | 0.49841973 | -5.717144373 |

| | | | | | | | |
|---|---|---|---|---|---|---|---|
| **P35270** | SPR | -0.038884841 | 3.196731725 | -1.405019483 | 0.206156573 | 0.612058044 | -5.718511754 |
| **O75326-1** | SEMA7A | -0.066843857 | 3.288157789 | -0.9952416 | 0.355365285 | 0.734489383 | -5.718645035 |
| **Q5TEU4** | NDUFAF5 | 0.104524728 | 3.155329599 | 1.168811617 | 0.283725858 | 0.679787821 | -5.718911332 |
| **A2RUS2** | DENND3 | -0.021909089 | 3.187982381 | -0.217144238 | 0.834781759 | 0.960912428 | -5.719441316 |
| **Q9NY93** | DDX56 | -0.094252285 | 3.196306591 | -2.053373095 | 0.082420751 | 0.424664634 | -5.719631347 |
| **Q96C12** | ARMC5 | 0.048383694 | 3.217109752 | 1.03773025 | 0.336607335 | 0.721128781 | -5.720491039 |
| **Q9NR56-1** | MBNL1 | 0.126035915 | 3.188793521 | 1.421507849 | 0.201522013 | 0.607855882 | -5.720736928 |
| **O00409** | FOXN3 | 0.053904658 | 3.133807202 | 1.013731565 | 0.347103166 | 0.728788229 | -5.720875138 |
| **Q96FA3** | PELI1 | -0.009549473 | 3.170546471 | -0.065640972 | 0.949645025 | 0.989472196 | -5.721381025 |
| **P49789** | FHIT | 0.056800014 | 3.132529724 | 0.602066379 | 0.567614113 | 0.854108736 | -5.721481391 |
| **Q9NV64-1** | TMEM39A | 0.032879217 | 3.251579478 | 0.483368552 | 0.644756518 | 0.891546795 | -5.721567249 |
| **Q04724** | TLE1 | -0.056529652 | 3.18610835 | -0.971649502 | 0.366130431 | 0.740745168 | -5.721743568 |
| **O95873** | C6orf47 | -0.049043514 | 3.181961621 | -0.553393301 | 0.598581163 | 0.871871293 | -5.7218963 |
| **O60307** | MAST3 | -0.005879057 | 3.196268425 | -0.072104817 | 0.944695898 | 0.98904859 | -5.722262786 |
| **Q58FF7** | HSP90AB3P | 0.060894016 | 3.186789053 | 0.667751349 | 0.527377848 | 0.839849063 | -5.722360518 |
| **O43295** | SRGAP3 | 0.050475872 | 3.159633262 | 0.550061238 | 0.600735761 | 0.87241889 | -5.722429774 |
| **O60268-1** | KIAA0513 | 0.040905403 | 3.242515242 | 0.640480325 | 0.543861457 | 0.844271059 | -5.722522263 |
| **O60331-3** | PIP5K1C | -0.085062866 | 3.264087228 | -0.822710564 | 0.439930046 | 0.788662011 | -5.722661229 |
| **O14782** | KIF3C | 0.048654804 | 3.079767853 | 0.700277983 | 0.508137739 | 0.82913633 | -5.722996763 |
| **P28906** | CD34 | 0.041459214 | 3.149990918 | 0.429981105 | 0.681161939 | 0.903980599 | -5.723199178 |
| **P07195** | LDHB | 0.019722526 | 3.180175544 | 2.642040657 | 0.035807352 | 0.347992906 | -5.725345274 |
| **P0C860** | MSL3P1 | -0.003122811 | 3.197447004 | -0.029656665 | 0.977234559 | 0.99349054 | -5.72589576 |
| **Q9HCM9** | TRIM39 | 0.043977408 | 3.207011865 | 0.655846157 | 0.534534609 | 0.84036998 | -5.725988249 |
| **Q15527** | SURF2 | -0.037804666 | 3.193744056 | -1.704484762 | 0.135558896 | 0.524095099 | -5.726510175 |
| **P52657** | GTF2A2 | -0.115495916 | 3.146415687 | -0.857973837 | 0.42154274 | 0.777315429 | -5.727313239 |
| **Q8N165** | PDIK1L | 0.050425249 | 3.204990788 | 1.444989582 | 0.195084692 | 0.600645673 | -5.727793344 |
| **Q9H2M9** | RAB3GAP2 | -0.032464556 | 3.178265337 | -2.200946025 | 0.066757369 | 0.394427892 | -5.728181106 |
| **Q9NPE2-2** | NGRN | 0.114466195 | 3.179042675 | 1.00717813 | 0.350013996 | 0.731407657 | -5.729098466 |
| **O14494** | PPAP2A; PLPP1 | -0.077909877 | 3.181808501 | -0.720571354 | 0.496368723 | 0.822965256 | -5.72928582 |
| **Q13322-1** | GRB10 | -0.073691298 | 3.25103691 | -0.779795448 | 0.463070506 | 0.802800142 | -5.729378176 |
| **P07305** | H1F0 | -0.056513578 | 3.166714549 | -0.993525269 | 0.356139997 | 0.734600528 | -5.72965889 |
| **Q2VPB7** | AP5B1 | 0.00910942 | 3.178723728 | 0.101726424 | 0.922053367 | 0.984164603 | -5.73031193 |
| **Q06787-5** | FMR1 | 0.068004025 | 3.175883817 | 1.85582025 | 0.109303509 | 0.479873319 | -5.731041702 |
| **Q8N9B5-1** | JMY | 0.038949775 | 3.191978285 | 1.552564764 | 0.167934167 | 0.566424749 | -5.731269506 |
| **Q16635-1** | TAZ | 0.095659754 | 3.205748712 | 0.531696329 | 0.612688804 | 0.878766193 | -5.731404764 |
| **Q86YV5** | SGK223; PRAG1 | 0.048042446 | 3.11342815 | 1.332568209 | 0.227676627 | 0.634253435 | -5.731456341 |
| **Q9Y2V0-1** | C15orf41 | 0.033274204 | 3.181891375 | 0.436178786 | 0.676885295 | 0.903662615 | -5.73256597 |
| **Q9H0X9** | OSBPL5 | -0.059956548 | 3.182408455 | -1.369373198 | 0.216505838 | 0.622562027 | -5.73329735 |
| **Q9Y6X4-1** | FAM169A | 0.037209321 | 3.188610015 | 1.928808241 | 0.098485677 | 0.457316603 | -5.733390443 |

| | | | | | | | |
|---|---|---|---|---|---|---|---|
| **Q9BYG5** | PARD6B | -0.049977445 | 3.191776434 | -0.891320914 | 0.404676795 | 0.766336431 | -5.734291697 |
| **Q86WT1** | TTC30A | -0.061277078 | 3.139141778 | -1.000112366 | 0.35317394 | 0.732738778 | -5.734631047 |
| **Q9UJX6** | ANAPC2 | -0.048744813 | 3.17676734 | -1.993089102 | 0.089839869 | 0.441148184 | -5.734659363 |
| **Q86V97** | KBTBD6 | 0.054984347 | 3.207182936 | 0.536853686 | 0.609318847 | 0.876646363 | -5.734952659 |
| **O95816** | BAG2 | 0.039314586 | 3.19008464 | 1.828314223 | 0.113676118 | 0.487929098 | -5.735161305 |
| **Q9UBM7** | DHCR7 | 0.091808483 | 3.15235516 | 1.44223247 | 0.195830721 | 0.601527622 | -5.736173936 |
| **Q96G01** | BICD1 | -0.050659107 | 3.230429352 | -1.652475703 | 0.145911221 | 0.544000305 | -5.737355067 |
| **Q9NWX6** | THG1L | -0.050487241 | 3.195281076 | -1.219376154 | 0.265273686 | 0.66823064 | -5.737517852 |
| **Q9H446** | RWDD1 | 0.047977369 | 3.153746824 | 1.581480202 | 0.16125872 | 0.561414897 | -5.737739909 |
| **Q9GZR5** | ELOVL4 | -0.051756079 | 3.206804029 | -0.831587917 | 0.435247731 | 0.784298014 | -5.737774733 |
| **Q9NWD8-1** | TMEM248 | 0.011653854 | 3.124757242 | 0.112153329 | 0.914100642 | 0.982219313 | -5.737868418 |
| **Q96G04-1** | FAM86A; EEF2KMT | -0.051240429 | 3.188512976 | -0.557248117 | 0.596094 | 0.871378088 | -5.737876161 |
| **Q53HC9** | TSSC1; EIPR1 | -0.046538066 | 3.158678587 | -1.688086469 | 0.138744779 | 0.53252525 | -5.738211056 |
| **Q92503-2** | SEC14L1 | -0.052477406 | 3.222834723 | -1.837223217 | 0.112241455 | 0.484388026 | -5.738328885 |
| **Q9BVK8-1** | TMEM147 | 0.071704008 | 3.18742703 | 0.715017264 | 0.49957171 | 0.825004623 | -5.739187372 |
| **P49768-1** | PSEN1 | -0.049833533 | 3.116107895 | -0.911970589 | 0.39448736 | 0.760544061 | -5.739215593 |
| **Q8WTT2** | NOC3L | -0.073063144 | 3.197803195 | -1.9611789 | 0.094033188 | 0.448519793 | -5.739405121 |
| **Q5TBK1-1** | N4BP2L1 | -0.008219422 | 3.167034217 | -0.082211225 | 0.936963216 | 0.987232823 | -5.739509082 |
| **Q6ZN04-1** | MEX3B | -0.049499284 | 3.217586194 | -0.855148437 | 0.422995078 | 0.777884596 | -5.739589371 |
| **Q96PV6-2** | LENG8 | 0.041765682 | 3.212664653 | 1.811716718 | 0.116396794 | 0.492686484 | -5.739781972 |
| **O95260-2** | ATE1 | 0.04495797 | 3.189806551 | 1.321058528 | 0.231273442 | 0.638769945 | -5.739962823 |
| **P30453** | HLA-A | 0.053478576 | 3.172511094 | 1.756511259 | 0.125909361 | 0.509142907 | -5.740021589 |
| **Q6P3W2** | DNAJC24 | -0.040560974 | 3.226861033 | -0.551838219 | 0.599586176 | 0.872097298 | -5.740068196 |
| **Q8TB61** | SLC35B2 | -0.070944997 | 3.18580127 | -1.434701177 | 0.197881814 | 0.603708078 | -5.740092342 |
| **Q4AE62** | GTDC1 | 0.041432367 | 3.18527591 | 0.387097814 | 0.71109657 | 0.921051693 | -5.741675496 |
| **P49674** | CSNK1E; LOC4009 | -0.048227728 | 3.206950527 | -0.77432629 | 0.466079485 | 0.803951695 | -5.742147548 |
| **Q6PCB5-1** | RSBN1L | -0.056056102 | 3.200712194 | -1.94895492 | 0.095690601 | 0.452536799 | -5.742334313 |
| **Q96LX8** | ZNF597 | 0.052687755 | 3.189476438 | 0.991873335 | 0.356886894 | 0.734600528 | -5.742380874 |
| **Q02952-1** | AKAP12 | -0.042503757 | 3.169453332 | -0.646497354 | 0.540197212 | 0.843540088 | -5.74241762 |
| **P07947** | YES1 | -0.004140431 | 3.162620299 | -0.042500152 | 0.967381286 | 0.992306703 | -5.743288686 |
| **O75380** | NDUFS6 | 0.09059383 | 3.217649791 | 1.416023215 | 0.203053071 | 0.608695713 | -5.743438281 |
| **P28328** | PEX2 | 0.054367722 | 3.180619725 | 1.325959535 | 0.229735764 | 0.637461728 | -5.744038803 |
| **Q9H0A0** | NAT10 | -0.068543644 | 3.19838734 | -2.385379404 | 0.051361283 | 0.371644256 | -5.744494666 |
| **Q6UXB8-1** | PI16 | 0.057183592 | 3.110897542 | 0.677964237 | 0.521287153 | 0.834809173 | -5.744523326 |
| **Q9H3M7** | TXNIP; LOC101060 | -0.075464735 | 3.157966722 | -1.074407876 | 0.321059356 | 0.708954025 | -5.744776951 |
| **Q6EMB2** | TTLL5 | 0.049370875 | 3.17388768 | 0.88933967 | 0.405664659 | 0.766336431 | -5.744831457 |
| **Q9H7X7** | RABL5; IFT22 | -0.048349904 | 3.199621173 | -1.41384925 | 0.203662849 | 0.608995611 | -5.745258288 |
| **Q8IV53-1** | DENND1C | 0.054365423 | 3.17536371 | 1.667137223 | 0.142918992 | 0.539313633 | -5.745674536 |
| **Q9NRY6** | PLSCR3 | -0.029854191 | 3.14981842 | -0.474879502 | 0.6504783 | 0.893885477 | -5.746526542 |

| | | | | | | | |
|---|---|---|---|---|---|---|---|
| **Q5JS54-2** | PSMG4 | 0.040399066 | 3.18205833 | 1.414752378 | 0.203409329 | 0.608695713 | -5.747085955 |
| **Q9Y388** | RBMX2 | -0.046967416 | 3.187137262 | -1.738667013 | 0.129141989 | 0.514283314 | -5.747305468 |
| **Q9BQ69** | MACROD1 | 0.122296436 | 3.191204822 | 1.617474898 | 0.153296562 | 0.552031202 | -5.747467708 |
| **Q99720** | SIGMAR1 | -0.056526939 | 3.157254747 | -0.819221331 | 0.441780249 | 0.789146094 | -5.748793652 |
| **Q8WXG6-2** | MADD | 0.043938399 | 3.178239235 | 1.577595126 | 0.162140965 | 0.561489222 | -5.751228478 |
| **O15033-1** | AREL1; KIAA0317 | -0.003970489 | 3.175103996 | -0.04423341 | 0.966051998 | 0.992001409 | -5.751542969 |
| **Q9NUI1** | DECR2 | -0.040475489 | 3.165185892 | -0.663035413 | 0.530205521 | 0.839849063 | -5.752260412 |
| **Q6SJ93-1** | FAM111B | -0.056747474 | 3.17260027 | -1.628140414 | 0.151009419 | 0.548684491 | -5.752261442 |
| **Q96H55-1** | MYO19 | -0.039610635 | 3.192177864 | -1.486356791 | 0.184198719 | 0.586791828 | -5.752501663 |
| **P37108** | SRP14 | -0.079391274 | 3.151207804 | -1.985547914 | 0.090813737 | 0.443191118 | -5.752556321 |
| **O43609** | SPRY1 | 0.052193294 | 3.154942995 | 0.776546547 | 0.464856333 | 0.803074528 | -5.752584492 |
| **O43293-1** | DAPK3 | -0.04012063 | 3.201509455 | -1.503121196 | 0.179948087 | 0.581254067 | -5.752897221 |
| **O60238** | BNIP3L | -0.108774902 | 3.0663697 | -0.575033036 | 0.584695487 | 0.8653224 | -5.753088556 |
| **P61024** | CKS1B | 0.038235702 | 3.185770099 | 1.456294187 | 0.192052865 | 0.596505086 | -5.7541627 |
| **Q4VC31** | CCDC58 | 0.108593737 | 3.185596297 | 1.628966641 | 0.150833594 | 0.548546897 | -5.754539567 |
| **P68431** | HIST1H3F; HIST1H | -0.031124935 | 3.185399553 | -0.319169951 | 0.759651292 | 0.938422437 | -5.754889761 |
| **Q9BZI7-1** | UPF3B | -0.03586511 | 3.163568431 | -1.718299518 | 0.132929456 | 0.519961998 | -5.755057357 |
| **Q16670-1** | ZNF187; ZSCAN26 | -0.018811171 | 3.130861951 | -0.200060904 | 0.847573257 | 0.964069922 | -5.755068978 |
| **O00566** | MPHOSPH10 | -0.046773195 | 3.179091083 | -1.960750682 | 0.094090764 | 0.448519793 | -5.75569382 |
| **Q12894-2** | IFRD2 | 0.048203966 | 3.246328437 | 1.105843161 | 0.308202546 | 0.696616637 | -5.75572523 |
| **O75460** | ERN 1.00 | 0.014680511 | 3.064527101 | 0.151892548 | 0.883897604 | 0.975295424 | -5.756206976 |
| **P19532** | TFE3 | 0.061078463 | 3.156772487 | 0.984178706 | 0.360382082 | 0.73661838 | -5.756852662 |
| **O43439-4** | CBFA2T2 | -0.070705695 | 3.209975476 | -1.251943562 | 0.253940515 | 0.659039362 | -5.756977196 |
| **Q9HB07** | C12orf10 | 0.069285676 | 3.153836721 | 1.844308552 | 0.111113115 | 0.48266468 | -5.758038116 |
| **Q9NU22** | MDN1 | -0.029719688 | 3.185422951 | -2.522782352 | 0.042310966 | 0.355540374 | -5.758214075 |
| **Q9NRR6-1** | INPP5E | 0.0064963 | 3.200193993 | 0.069698078 | 0.946538351 | 0.989472196 | -5.75825168 |
| **Q6TCH7-1** | PAQR3 | 0.047078042 | 3.220912157 | 0.866907692 | 0.416974461 | 0.773310479 | -5.758609077 |
| **P30049** | ATP5D | 0.093882315 | 3.169003797 | 1.127417953 | 0.299625909 | 0.690882899 | -5.759013166 |
| **Q9H2W6** | MRPL46 | 0.191750884 | 3.136470568 | 1.509197789 | 0.178429829 | 0.57989621 | -5.759504559 |
| **O15534-1** | PER1 | 0.042418905 | 3.200443872 | 0.552119532 | 0.5994043 | 0.872097298 | -5.759562781 |
| **Q15388** | TOMM20 | 0.038369823 | 3.195033521 | 0.677770198 | 0.521402452 | 0.834809173 | -5.759688734 |
| **Q9Y6A4** | C16orf80; CFAP20 | -0.04488425 | 3.169372957 | -1.350593466 | 0.222143228 | 0.627993355 | -5.759818478 |
| **Q96G97-4** | BSCL2 | -0.012810888 | 3.172617231 | -0.136554216 | 0.895533352 | 0.978757548 | -5.760014706 |
| **Q8NBK3** | SUMF1 | 0.037689844 | 3.129900361 | 0.498102646 | 0.634887601 | 0.887591411 | -5.760902503 |
| **Q4VC05-2** | BCL7A | 0.038093395 | 3.253008188 | 0.297678243 | 0.775276124 | 0.94357883 | -5.76109646 |
| **Q9NWM8** | FKBP14 | 0.054287234 | 3.210674331 | 1.064886627 | 0.32503857 | 0.710703247 | -5.761613158 |
| **Q9Y5A9-1** | YTHDF2 | -0.040791652 | 3.197765513 | -2.18355857 | 0.068433814 | 0.398319894 | -5.761981958 |
| **Q9P2F8-1** | SIPA1L2 | -0.040156423 | 3.213451585 | -0.548901922 | 0.601486428 | 0.872497827 | -5.76254138 |
| **P00558** | PGK 1.00 | 0.038922433 | 3.172320989 | 2.867182635 | 0.026237843 | 0.328952544 | -5.762723548 |

| | | | | | | | |
|---|---|---|---|---|---|---|---|
| **P40925-3** | MDH1 | 0.025963664 | 3.165393388 | 2.314021425 | 0.05683181 | 0.377614679 | -5.763219893 |
| **Q15274** | QPRT | 0.035394698 | 3.188428564 | 1.497915203 | 0.181258296 | 0.58362594 | -5.763460344 |
| **Q9H4B7** | TUBB1 | -0.047254164 | 3.218513712 | -0.423795067 | 0.685443324 | 0.906272103 | -5.764337567 |
| **Q9BST9** | RTKN | 0.051315305 | 3.192811451 | 1.00155996 | 0.352524725 | 0.732738778 | -5.764432565 |
| **Q9Y6R1-5** | SLC4A4 | 0.041072979 | 3.162337653 | 0.62025037 | 0.556293371 | 0.850606395 | -5.764663486 |
| **Q9Y324** | FCF1 | -0.052194069 | 3.148437404 | -1.504283989 | 0.179656639 | 0.580990067 | -5.764928449 |
| **P61244** | MAX | -0.075583802 | 3.179119187 | -1.521197596 | 0.175466356 | 0.576466876 | -5.765651991 |
| **Q9NX24** | NHP2 | 0.078914371 | 3.170715094 | 1.598197079 | 0.157513697 | 0.555457754 | -5.766268447 |
| **Q08170** | SRSF4 | 0.066841392 | 3.139103413 | 1.855449748 | 0.109361298 | 0.479873319 | -5.766436468 |
| **Q8WXD5** | GEMIN6 | -0.06120821 | 3.212265299 | -1.628926598 | 0.15084211 | 0.548546897 | -5.76697615 |
| **Q14118** | DAG1 | -0.061636436 | 3.208053729 | -1.499996066 | 0.180733547 | 0.583236405 | -5.767068592 |
| **O43805** | SSNA1 | -0.11503495 | 3.159192868 | -1.182622498 | 0.278581095 | 0.676427621 | -5.767079621 |
| **Q9BYC9** | MRPL20 | 0.115566321 | 3.204698848 | 1.667397875 | 0.142866328 | 0.539313633 | -5.767633469 |
| **P26232** | CTNNA2 | -0.110787064 | 3.161980964 | -0.730928482 | 0.490432413 | 0.818649279 | -5.768092391 |
| **Q86SE9** | PCGF5 | 0.065027892 | 3.181246493 | 1.322556879 | 0.230802376 | 0.638705983 | -5.768151508 |
| **Q96EU7** | C1GALT1C1 | 0.038812966 | 3.205961282 | 0.741821095 | 0.484240686 | 0.814322316 | -5.769517485 |
| **Q9BVC6** | TMEM109 | -0.05193182 | 3.1624093 | -1.37575554 | 0.214619263 | 0.619913059 | -5.769552424 |
| **Q9Y4E1-4** | FAM21C; WASHC2 | 0.059497424 | 3.14761083 | 1.262822834 | 0.250248981 | 0.654603944 | -5.76984895 |
| **P53701** | HCCS | 0.121794234 | 3.181480656 | 1.58739138 | 0.159925008 | 0.55963744 | -5.770077604 |
| **Q6NUS6** | TCTN3 | -0.045477906 | 3.223804523 | -0.6461451 | 0.540411303 | 0.843540088 | -5.770094417 |
| **Q8NAA4** | ATG16L2 | -0.031813339 | 3.163306444 | -0.518069814 | 0.621641842 | 0.883385606 | -5.770381959 |
| **Q8IYD8** | FANCM | 0.018173819 | 3.219003412 | 0.205052505 | 0.843830316 | 0.962543524 | -5.770469356 |
| **Q9NUG6** | PDRG1 | -0.074945804 | 3.206670558 | -1.446405346 | 0.194702617 | 0.600645673 | -5.771209382 |
| **Q92784-2** | DPF3 | -0.081165737 | 3.176632675 | -1.179549873 | 0.279718826 | 0.676725357 | -5.772225358 |
| **O75586-2** | MED6 | 0.146882243 | 3.206996748 | 1.49524436 | 0.181933884 | 0.584742029 | -5.772251983 |
| **O94854** | KIAA0754 | -0.050195593 | 3.149271192 | -1.052625458 | 0.330221574 | 0.7158555 | -5.772255446 |
| **P14621** | ACYP2 | -0.054915351 | 3.182730627 | -1.655182502 | 0.145354352 | 0.543198649 | -5.772977683 |
| **P81877-1** | SSBP2 | 0.04189467 | 3.150474987 | 0.547856804 | 0.602163602 | 0.872497827 | -5.773619488 |
| **Q9H173** | SIL1 | -0.051191433 | 3.162881454 | -1.375256404 | 0.214766271 | 0.619913059 | -5.773768332 |
| **Q96GE6** | CALML4 | -0.089355737 | 3.151064245 | -0.349500747 | 0.737808361 | 0.930015458 | -5.77423478 |
| **O43768-4** | ENSA | -0.273428437 | 3.153537698 | -1.604297562 | 0.156167504 | 0.554834464 | -5.776193316 |
| **Q9HCG8** | CWC22 | -0.042106019 | 3.170836004 | -2.121287448 | 0.074796727 | 0.407645581 | -5.776951462 |
| **P13807** | GYS1 | -0.047669028 | 3.192295808 | -1.801888106 | 0.118037711 | 0.495933166 | -5.777065974 |
| **Q92793-1** | CREBBP | 0.031245806 | 3.171633826 | 1.766879678 | 0.124067087 | 0.506532891 | -5.777259316 |
| **Q96PB1** | CASD1 | -0.091313062 | 3.20459093 | -1.286291599 | 0.242444097 | 0.649360189 | -5.777733567 |
| **Q05BQ5** | MBTD1 | 0.04380749 | 3.144764346 | 1.471805128 | 0.187962958 | 0.591197824 | -5.778840962 |
| **Q96LZ7-1** | FAM82A1; RMDN2 | -0.031508318 | 3.148793856 | -0.188394003 | 0.856338217 | 0.968855822 | -5.779409974 |
| **Q15637-6** | SF1 | -0.043273342 | 3.212906671 | -0.82205914 | 0.44027505 | 0.788722722 | -5.779633594 |
| **P50548** | ERF | -0.033016635 | 3.196634376 | -0.3982982 | 0.703221573 | 0.915549813 | -5.780330266 |

| | | | | | | | |
|---|---|---|---|---|---|---|---|
| **Q96CJ1-1** | EAF2 | 0.036584944 | 3.211840015 | 0.417755757 | 0.68963534 | 0.908760279 | -5.780490283 |
| **Q641Q2** | FAM21A; WASHC2/ | 0.046819579 | 3.169137973 | 1.329678687 | 0.228574929 | 0.635864079 | -5.781966326 |
| **Q9BYC5** | FUT8 | -0.040703219 | 3.177323615 | -1.71422962 | 0.133698974 | 0.521508161 | -5.782268695 |
| **Q9H0E7** | USP44 | 0.043401844 | 3.167560944 | 0.430468484 | 0.680825162 | 0.903980599 | -5.78287303 |
| **Q8IUC6** | TICAM1 | 0.016872781 | 3.179646287 | 0.177348236 | 0.864656972 | 0.969885592 | -5.783250535 |
| **Q6ZNC8-1** | MBOAT1 | -0.05788011 | 3.204646446 | -0.761937012 | 0.4729456 | 0.808597546 | -5.783291031 |
| **P27144** | AK4; LOC10050785 | 0.137647532 | 3.192519984 | 1.786251992 | 0.120694676 | 0.499295152 | -5.783328317 |
| **P19256** | CD58 | -0.057575287 | 3.171611533 | -1.073467039 | 0.32145079 | 0.709227029 | -5.784226014 |
| **P51797-5** | CLCN6 | 0.000269387 | 3.103497961 | 0.002318361 | 0.99822005 | 0.998597116 | -5.784455086 |
| **Q9Y282-3** | ERGIC3 | -0.038684781 | 3.18727628 | -1.774236405 | 0.122775781 | 0.502293294 | -5.785031822 |
| **Q9UL45** | BLOC1S6 | 0.059525921 | 3.187072921 | 0.903595726 | 0.398596423 | 0.762880385 | -5.785348441 |
| **Q6VN20** | RANBP10 | -0.039763816 | 3.184869122 | -1.537323183 | 0.171555771 | 0.571732635 | -5.785407255 |
| **Q92800-2** | EZH1 | 0.047752862 | 3.211979286 | 0.699887199 | 0.508366153 | 0.82913633 | -5.785414255 |
| **O75915** | ARL6IP5 | -0.102414525 | 3.207457462 | -1.278501308 | 0.245011007 | 0.65017116 | -5.786406533 |
| **Q8IXI2-3** | RHOT1 | 0.076727267 | 3.178222908 | 1.70794857 | 0.134894963 | 0.523800962 | -5.786590416 |
| **Q96PU5** | NEDD4L | 0.065647148 | 3.204682061 | 0.801453955 | 0.45128747 | 0.79513757 | -5.786735981 |
| **Q96DX5** | ASB9 | 0.057459642 | 3.19043965 | 1.250566013 | 0.25441129 | 0.659262133 | -5.786767255 |
| **Q6PJ69** | TRIM65 | -0.055462397 | 3.166873691 | -1.43594487 | 0.197541762 | 0.603708078 | -5.786801807 |
| **Q9UJN7** | ZNF391 | 0.044960314 | 3.214443053 | 0.485228469 | 0.643506373 | 0.891474831 | -5.787555601 |
| **Q70Z53** | FRA10AC1 | -0.038727693 | 3.183583394 | -1.534938218 | 0.172129001 | 0.571776747 | -5.788482416 |
| **Q9BXC9** | BBS2 | 0.054783646 | 3.175843934 | 1.343589594 | 0.224278928 | 0.629881158 | -5.788810967 |
| **Q9Y6K9-2** | IKBKG | -0.049446646 | 3.186150056 | -1.763999071 | 0.124576285 | 0.507048456 | -5.789177433 |
| **P15260** | IFNGR1 | -0.060311343 | 3.224771239 | -0.79644835 | 0.453991824 | 0.796767184 | -5.789544329 |
| **Q15072** | ZNF146 | -0.017356547 | 3.170494342 | -0.183308657 | 0.860165678 | 0.968855822 | -5.789617678 |
| **P04424** | ASL | 0.035460996 | 3.184329057 | 1.692653376 | 0.137850389 | 0.529905456 | -5.789722721 |
| **Q9P0J7** | KCMF1 | 0.064485999 | 3.187063933 | 1.199702874 | 0.272327808 | 0.672884265 | -5.790560553 |
| **Q5UE93** | PIK3R6 | -0.058190549 | 3.198415971 | -1.31295469 | 0.233835934 | 0.641071944 | -5.791345035 |
| **Q12979** | ABR | 0.034571242 | 3.218642105 | 0.449324401 | 0.667857322 | 0.90001396 | -5.792469862 |
| **Q9UJX5-3** | ANAPC4 | -0.037715555 | 3.192540024 | -1.533130942 | 0.172564566 | 0.572872916 | -5.792689634 |
| **Q8WYA0-1** | IFT81 | 0.032468747 | 3.16192606 | 1.846612752 | 0.110748568 | 0.482135549 | -5.793163582 |
| **Q9Y2M5** | KLHL20 | 0.01107811 | 3.258006582 | 0.109973968 | 0.915762014 | 0.98295196 | -5.793475463 |
| **Q9BUV8-5** | C20orf24 | 0.10987229 | 3.200629882 | 1.078894721 | 0.319197935 | 0.707596986 | -5.793503332 |
| **P20591** | MX1 | -0.082424809 | 3.210485645 | -2.00723842 | 0.088040652 | 0.437724019 | -5.793763549 |
| **P02649** | APOE | 0.085052613 | 3.200343025 | 1.450887397 | 0.19349753 | 0.598629202 | -5.793977728 |
| **Q14586** | ZNF267 | -0.045973085 | 3.184553256 | -0.814702745 | 0.444184512 | 0.790556887 | -5.794688118 |
| **Q9NZQ3-1** | NCKIPSD | -0.03626824 | 3.196325595 | -1.526534855 | 0.174162965 | 0.574454961 | -5.795459409 |
| **Q9NSK7-4** | C19orf12 | -0.020007833 | 3.107901764 | -0.243828777 | 0.814911232 | 0.95620584 | -5.795545367 |
| **Q6ZWT7** | MBOAT2 | -0.028765118 | 3.229950485 | -0.468202503 | 0.654996862 | 0.896497734 | -5.795584919 |
| **Q9UGK3-2** | STAP2 | 0.046491648 | 3.186430645 | 1.04516509 | 0.333407652 | 0.719098069 | -5.796011521 |

| | | | | | | | |
|---|---|---|---|---|---|---|---|
| **Q6SPF0** | SAMD1 | -0.041019886 | 3.197834691 | -1.511093335 | 0.177958647 | 0.57969719 | -5.796158905 |
| **Q5BKT4** | ALG10 | 0.046611984 | 3.189641145 | 0.776584487 | 0.464835451 | 0.803074528 | -5.798541243 |
| **Q8IZU2-1** | WDR17 | -0.051758962 | 3.179747444 | -0.571142729 | 0.587178014 | 0.86631928 | -5.798782831 |
| **Q86WW8** | COA5 | 0.069905216 | 3.194349678 | 1.102815385 | 0.309422218 | 0.697209166 | -5.79907147 |
| **Q9BWF2** | TRAIP | -0.02111359 | 3.214914445 | -0.295519795 | 0.776851785 | 0.94357883 | -5.799265567 |
| **Q9H1X3** | DNAJC25; DNAJC2 | 0.041537466 | 3.211564244 | 0.499220349 | 0.634142215 | 0.887509451 | -5.800069485 |
| **Q5U623** | ATF7IP2 | -0.000361541 | 3.210826787 | -0.004748601 | 0.996354215 | 0.998480498 | -5.800483336 |
| **Q8IXQ6-2** | PARP9 | -0.034385216 | 3.172506733 | -1.918044175 | 0.100012017 | 0.461169746 | -5.80048724 |
| **Q9H0M0-1** | WWP1 | -0.050264997 | 3.202488945 | -0.647146112 | 0.539803052 | 0.843540088 | -5.800493938 |
| **Q9Y3E0** | GOLT1B | -0.087674389 | 3.24086288 | -0.53888602 | 0.607993689 | 0.875726951 | -5.800898631 |
| **O75764** | TCEA3 | 0.043110696 | 3.188147263 | 0.728660308 | 0.491728368 | 0.820113219 | -5.801411862 |
| **Q4W5G0** | TIGD2 | -0.011575766 | 3.174996969 | -0.131777644 | 0.899162767 | 0.978757548 | -5.801510538 |
| **Q00839** | HNRNPU | 0.047168202 | 3.194565804 | 0.609818864 | 0.562770889 | 0.852576894 | -5.801619505 |
| **Q9UJ78-4** | ZMYM5 | 0.047907244 | 3.158100472 | 0.6708345 | 0.525534381 | 0.838233243 | -5.801841103 |
| **Q9UM82** | SPATA2 | -0.04472831 | 3.214317486 | -1.120859288 | 0.302212033 | 0.691344519 | -5.801853521 |
| **Q13439-5** | GOLGA4 | 0.032900182 | 3.183462066 | 0.300426256 | 0.773271751 | 0.943127531 | -5.802632828 |
| **O75436-1** | VPS26A | -0.051641537 | 3.166644073 | -2.012244421 | 0.087412747 | 0.435966276 | -5.802692567 |
| **Q14157-1** | UBAP2L | -0.126505209 | 3.217129492 | -0.467485614 | 0.655482948 | 0.896497734 | -5.804040941 |
| **Q9HDC5** | JPH1 | -0.045285388 | 3.209074175 | -1.765498485 | 0.124310985 | 0.50674745 | -5.804052084 |
| **Q9NPA8-1** | ENY2 | -0.058487377 | 3.167360746 | -1.7232322 | 0.132002508 | 0.519814431 | -5.804881986 |
| **O60336** | MAPKBP1 | -0.040692307 | 3.121707271 | -0.531974696 | 0.61250665 | 0.878766193 | -5.80509503 |
| **P49590** | HARS2 | 0.147082722 | 3.126848689 | 1.417068713 | 0.202760406 | 0.608695713 | -5.805794317 |
| **Q6Y1H2** | PTPLB; HACD2 | -0.013688081 | 3.199376922 | -0.130950265 | 0.89979171 | 0.978757548 | -5.806440965 |
| **P29353-6** | SHC1 | 0.066917034 | 3.192582458 | 1.629303885 | 0.150761882 | 0.548546897 | -5.8066175 |
| **A8MW92** | PHF20L1 | -0.042592607 | 3.174357815 | -1.435134235 | 0.197763347 | 0.603708078 | -5.806743049 |
| **Q9NUP7-1** | TRMT13 | -0.029797421 | 3.208191209 | -1.676759626 | 0.140987032 | 0.536483353 | -5.808009334 |
| **Q9Y4P8-1** | WIPI2 | 0.04398093 | 3.178649017 | 1.367621107 | 0.217026332 | 0.623156562 | -5.808519661 |
| **P29590-13** | PML | -0.01390776 | 3.194017419 | -0.141703882 | 0.891623501 | 0.977905675 | -5.809553829 |
| **Q5SRI9** | MANEA | 0.039113096 | 3.175523954 | 0.588146222 | 0.576372558 | 0.860605145 | -5.809770802 |
| **O75438** | NDUFB1 | 0.142639339 | 3.184900101 | 1.391030322 | 0.210163847 | 0.615470161 | -5.809996566 |
| **O14681** | EI24 | 0.051792296 | 3.220242396 | 1.16909409 | 0.283619836 | 0.679787821 | -5.809999521 |
| **O75496** | GMNN | -0.082164646 | 3.153476875 | -1.602474005 | 0.156568769 | 0.554834464 | -5.810365974 |
| **P50897** | PPT1 | 0.044090004 | 3.165759436 | 1.628020338 | 0.151034988 | 0.548684491 | -5.810618719 |
| **O76024** | WFS1 | -0.002909943 | 3.227561223 | -0.033263414 | 0.974467012 | 0.99316605 | -5.810929914 |
| **Q9UKN5** | PRDM4 | 0.017245343 | 3.187655506 | 0.090826881 | 0.930376962 | 0.986707862 | -5.811812148 |
| **A4D2B0** | MBLAC1 | -0.000818007 | 3.134310482 | -0.005873426 | 0.99549063 | 0.998254835 | -5.812417762 |
| **P06737-1** | PYGL | 0.047143062 | 3.203787512 | 1.220564321 | 0.264852703 | 0.66823064 | -5.812506758 |
| **P61020** | RAB5B | -0.041140371 | 3.168784997 | -1.23989227 | 0.258084651 | 0.660717666 | -5.812763001 |
| **Q9Y5A6** | ZSCAN21 | -0.02641692 | 3.175019941 | -0.30376783 | 0.77083698 | 0.94219997 | -5.814639117 |

| | | | | | | | |
|---|---|---|---|---|---|---|---|
| **Q9H9L7** | AKIRIN1 | -0.042895817 | 3.254939832 | -0.484910914 | 0.643719729 | 0.891474831 | -5.815021259 |
| **P62079** | TSPAN5 | 0.073151816 | 3.13770366 | 0.820818493 | 0.44093265 | 0.788971713 | -5.815508406 |
| **Q9NYB9-2** | ABI2 | 0.055754811 | 3.164198907 | 1.456847471 | 0.191905587 | 0.596505086 | -5.816112835 |
| **Q9BWM7** | SFXN3 | 0.13692588 | 3.149763152 | 0.779993941 | 0.462961555 | 0.802800142 | -5.816330006 |
| **Q6ZT21-1** | TMPPE | -0.009144587 | 3.209162913 | -0.131495236 | 0.899377434 | 0.978757548 | -5.816399715 |
| **O15527-4** | OGG1 | 0.029510966 | 3.174624092 | 0.420871712 | 0.687470993 | 0.907431059 | -5.816433179 |
| **Q9H2A2-1** | ALDH8A1 | 0.027054979 | 3.215795349 | 0.378633734 | 0.717073337 | 0.924679646 | -5.816449707 |
| **Q9BTV7** | CABLES2 | -0.052053347 | 3.19565076 | -1.115215367 | 0.304452233 | 0.693086302 | -5.816581659 |
| **Q6ICL3-4** | C22orf25; TANGO2 | 0.035367409 | 3.18265592 | 0.339017396 | 0.74532948 | 0.934897808 | -5.816704609 |
| **P15813** | CD1D | 0.015437869 | 3.168532356 | 0.201595061 | 0.846422411 | 0.963442128 | -5.81684424 |
| **Q8WVB3-2** | HEXDC | 0.039948749 | 3.195291293 | 0.911262234 | 0.394833671 | 0.761027053 | -5.81705688 |
| **P55196-5** | MLLT4; AFDN | -0.013642997 | 3.239231488 | -0.177829553 | 0.864294086 | 0.969738239 | -5.817281945 |
| **Q9H165-1** | BCL11A | 0.042997394 | 3.174374007 | 0.700290163 | 0.508130621 | 0.82913633 | -5.817283633 |
| **Q5JSZ5-1** | PRRC2B | -0.032956448 | 3.182843846 | -1.743643254 | 0.128232541 | 0.512600477 | -5.817682719 |
| **Q9NPB6-1** | PARD6A | -0.022917203 | 3.210825543 | -0.31875079 | 0.759954891 | 0.938422437 | -5.817852237 |
| **Q96BR1-1** | SGK3; C8orf44-SG | 0.030843274 | 3.194323363 | 0.444332643 | 0.671278556 | 0.901175787 | -5.81794767 |
| **P35711** | SOX5 | 0.068775312 | 3.174492747 | 0.958136519 | 0.372409654 | 0.745276848 | -5.818297803 |
| **Q9NW75-1** | GPATCH2 | -0.075619484 | 3.186302822 | -1.130811367 | 0.298295086 | 0.689670498 | -5.818695511 |
| **Q8TBZ3-2** | WDR20 | 0.051382162 | 3.225599418 | 0.741736441 | 0.484288602 | 0.814322316 | -5.818817085 |
| **Q9H4P4** | RNF41 | 0.044647478 | 3.184410142 | 1.242855147 | 0.257060417 | 0.659782078 | -5.819150297 |
| **O14628-1** | ZNF195 | -0.03956871 | 3.186339397 | -1.158662567 | 0.287557354 | 0.682522792 | -5.81948535 |
| **P55145** | MANF | 0.032517318 | 3.200967948 | 1.877909461 | 0.105911721 | 0.475403904 | -5.819946705 |
| **O60337** | 6-Mar | -0.046955002 | 3.209892843 | -0.921416787 | 0.389891034 | 0.75855365 | -5.820122151 |
| **Q969Q6-1** | PPP2R3C | 0.043420883 | 3.180565994 | 0.688329546 | 0.515151868 | 0.83239406 | -5.820157196 |
| **P17676** | CEBPB | -0.001132068 | 3.226382697 | -0.015754005 | 0.987905233 | 0.996054198 | -5.820196946 |
| **Q8NEY8-3** | PPHLN1 | -0.018319794 | 3.18092728 | -0.206448365 | 0.842784414 | 0.962543524 | -5.82030108 |
| **O75582-1** | RPS6KA5 | 0.054119602 | 3.187604652 | 1.67813949 | 0.140712036 | 0.53619047 | -5.820356755 |
| **O60506** | SYNCRIP | 0.046026679 | 3.154263417 | 1.321035112 | 0.23128081 | 0.638769945 | -5.82125858 |
| **Q6UWZ7** | FAM175A; ABRAXA | 0.032509714 | 3.169468117 | 1.560464693 | 0.166085193 | 0.565363245 | -5.82127668 |
| **Q8IX21-2** | FAM178A; SLF2 | -0.048591253 | 3.206366884 | -0.769089449 | 0.468973297 | 0.805725242 | -5.821848124 |
| **P67812-3** | SEC11A | -0.057848807 | 3.191417551 | -1.661634815 | 0.144035056 | 0.539645535 | -5.822095761 |
| **Q9Y3B9** | RRP15 | -0.044190224 | 3.2212714 | -1.735552319 | 0.129714384 | 0.515290389 | -5.823816931 |
| **Q9P281** | BAHCC1 | -0.045112323 | 3.182210328 | -1.554872503 | 0.167392066 | 0.566424749 | -5.824542459 |
| **Q9NRY4** | ARHGAP35 | 0.029024029 | 3.18877401 | 2.019371538 | 0.086526505 | 0.434379005 | -5.8250706 |
| **P30039** | PBLD | -0.020274491 | 3.173804259 | -0.26194195 | 0.801506392 | 0.953003335 | -5.825124821 |
| **Q6IEG0-1** | SNRNP48 | -0.040849983 | 3.198033941 | -1.348345326 | 0.222826777 | 0.628740079 | -5.825126335 |
| **Q9Y3C0** | CCDC53; WASHC3 | -0.052029304 | 3.179852367 | -1.494005469 | 0.182248047 | 0.584895208 | -5.825178893 |
| **Q9UJW8** | ZNF180 | -0.007336635 | 3.194567422 | -0.083964954 | 0.935622121 | 0.986793807 | -5.825221071 |
| **Q8WYH8-1** | ING5 | 0.01162822 | 3.210343572 | 0.151415279 | 0.88425921 | 0.97534908 | -5.825461347 |

| | | | | | | | |
|---|---|---|---|---|---|---|---|
| **P54886** | ALDH18A1 | 0.165381782 | 3.19736791 | 2.202759746 | 0.066584922 | 0.394427892 | -5.825844627 |
| **P05107** | ITGB2 | -0.05041091 | 3.210372493 | -2.2246512 | 0.064538981 | 0.392319975 | -5.825955425 |
| **Q9HA47-4** | UCK1 | -0.046196862 | 3.201966508 | -1.398972374 | 0.207880277 | 0.614231901 | -5.826797702 |
| **Q9Y6J9** | TAF6L | -0.046803579 | 3.145282432 | -1.387673036 | 0.211135931 | 0.61590617 | -5.827324526 |
| **Q15750-1** | TAB1 | 0.031283921 | 3.198876343 | 1.757728313 | 0.125691751 | 0.509142907 | -5.82740543 |
| **Q96S38** | RPS6KC1 | -0.04557031 | 3.16905623 | -0.618365325 | 0.557460544 | 0.851117349 | -5.827542122 |
| **P09497-2** | CLTB | -0.43546048 | 3.162162629 | -1.724034691 | 0.131852293 | 0.519814431 | -5.827654936 |
| **Q9NYG5** | ANAPC11 | -0.102156616 | 3.205192386 | -0.962308505 | 0.370462193 | 0.744424726 | -5.82786101 |
| **Q9H3E2** | SNX25 | -0.003620791 | 3.217798212 | -0.047515507 | 0.963535185 | 0.991283985 | -5.827878478 |
| **Q7Z3E5** | ARMC9 | -0.039059354 | 3.177206125 | -1.20144932 | 0.271695193 | 0.671838876 | -5.828413178 |
| **P28074-1** | PSMB5 | -0.066492855 | 3.216787839 | -1.293291033 | 0.240157857 | 0.645958461 | -5.82893621 |
| **Q2YD98-1** | UVSSA | -0.04434965 | 3.192770014 | -1.427982744 | 0.199727987 | 0.605177234 | -5.829373845 |
| **Q92797-1** | SYMPK | 0.020475346 | 3.179890733 | 2.18394791 | 0.06839581 | 0.398319894 | -5.829924608 |
| **P20393** | NR1D1 | -0.048672718 | 3.235101112 | -0.86088035 | 0.42005251 | 0.776300813 | -5.830017059 |
| **Q13233** | MAP3K1 | -0.048051784 | 3.197033748 | -0.831674647 | 0.435202162 | 0.784298014 | -5.831630683 |
| **Q14527** | HLTF | -0.041201627 | 3.194212536 | -2.142376971 | 0.072577239 | 0.404366176 | -5.831758095 |
| **Q8NCH0** | CHST14 | -0.038530516 | 3.151326784 | -1.149081139 | 0.291214313 | 0.685264687 | -5.832185458 |
| **Q86XI8** | C19orf68; ZSWIM9 | -0.008576456 | 3.148673609 | -0.086863461 | 0.933406102 | 0.986793807 | -5.832533386 |
| **Q9P2E9-1** | RRBP1 | -0.061847877 | 3.196605993 | -1.120288199 | 0.30243809 | 0.691344519 | -5.833524092 |
| **Q16540** | MRPL23 | 0.080002135 | 3.194867559 | 0.973940201 | 0.365074173 | 0.740115924 | -5.833808649 |
| **A8CG34-1** | POM121C | 0.040002351 | 3.226229465 | 0.856713154 | 0.422190318 | 0.777719006 | -5.834856474 |
| **O60725** | ICMT | -0.049132534 | 3.207086653 | -0.58591557 | 0.577783429 | 0.861313941 | -5.835709831 |
| **O15015-2** | ZNF646 | -0.061150269 | 3.220624351 | -0.889616937 | 0.405526303 | 0.766336431 | -5.83573293 |
| **P57775** | FBXW4 | -0.101438218 | 3.153003152 | -0.964291494 | 0.369539308 | 0.74415747 | -5.835886893 |
| **Q13867** | BLMH | 0.03556188 | 3.176338977 | 2.074328139 | 0.079988355 | 0.419060309 | -5.836939977 |
| **Q8N983-4** | MRPL43 | 0.086406047 | 3.199453898 | 1.73413424 | 0.129975796 | 0.515715456 | -5.837186438 |
| **O60831** | PRAF2 | -0.043844334 | 3.200407398 | -0.81637376 | 0.443294311 | 0.789680112 | -5.837616536 |
| **Q9UDV7** | ZNF282 | -0.049857958 | 3.178909768 | -1.255716322 | 0.25265504 | 0.656572825 | -5.838089843 |
| **Q8TEB9** | RHBDD1 | 0.016183756 | 3.15937898 | 0.208323974 | 0.841379588 | 0.962112957 | -5.838477193 |
| **Q9BWF3-3** | RBM4 | -0.059165522 | 3.213728413 | -1.298453828 | 0.238483645 | 0.644491512 | -5.838733824 |
| **Q8NB37** | PDDC1; GATD1 | 0.04470639 | 3.219158153 | 0.921411771 | 0.389893464 | 0.75855365 | -5.839209053 |
| **P41236** | PPP1R2 | -0.133761867 | 3.17013237 | -1.52741685 | 0.173948442 | 0.574454961 | -5.839456785 |
| **Q9BWC9** | CCDC106 | 0.058640128 | 3.140760657 | 0.832055573 | 0.435002063 | 0.784298014 | -5.84015181 |
| **Q6NS38-1** | ALKBH2 | -0.040553649 | 3.169505691 | -0.948948984 | 0.376726163 | 0.749049177 | -5.840572253 |
| **P26641** | EEF1G | -0.071459687 | 3.192116115 | -2.386408508 | 0.051286495 | 0.37144139 | -5.840726111 |
| **Q96DZ5** | CLIP3 | 0.039443173 | 3.088792378 | 0.721201195 | 0.496006363 | 0.822932915 | -5.841837505 |
| **P06753-4** | TPM3 | -0.059926385 | 3.16964906 | -0.978016507 | 0.363200421 | 0.738441741 | -5.841923305 |
| **Q9NX01** | TXNL4B | 0.042283345 | 3.229666572 | 0.800935652 | 0.451566963 | 0.79513757 | -5.842409729 |
| **P42773** | CDKN2C | 0.055603045 | 3.149025313 | 1.096528736 | 0.311967321 | 0.699768595 | -5.842961973 |

| | | | | | | | |
|---|---|---|---|---|---|---|---|
| **Q9Y4X4-1** | KLF12 | -0.050503157 | 3.176310408 | -0.831873176 | 0.435097868 | 0.784298014 | -5.843013599 |
| **Q9H9L3** | ISG20L2 | -0.100588605 | 3.199821397 | -1.350407919 | 0.222199573 | 0.627993355 | -5.843069642 |
| **Q9HAU8** | RNPEPL1 | 0.005540968 | 3.139115447 | 0.077422883 | 0.940626036 | 0.988136171 | -5.843106644 |
| **O60287** | URB1 | -0.041093865 | 3.202769027 | -2.144230312 | 0.072385408 | 0.404094866 | -5.843111309 |
| **Q9NQR4** | NIT2 | 0.042937285 | 3.174507383 | 2.032780757 | 0.084883391 | 0.430367743 | -5.843778301 |
| **A3KN83-1** | SBNO1 | -0.034452493 | 3.160411897 | -1.854055509 | 0.109579035 | 0.480201565 | -5.843997467 |
| **O60762** | DPM1 | -0.054420217 | 3.210077578 | -1.730816256 | 0.130589425 | 0.516957143 | -5.844220252 |
| **Q8TBZ6** | TRMT10A | 0.03650984 | 3.197674362 | 1.337631943 | 0.226109939 | 0.63121696 | -5.844770016 |
| **P17036-1** | ZNF3 | 0.037554186 | 3.213451003 | 0.691167927 | 0.513479985 | 0.83148099 | -5.84525868 |
| **Q9GZL7** | WDR12 | -0.038802853 | 3.196502537 | -1.972828206 | 0.092480244 | 0.446659583 | -5.845462833 |
| **Q8WW59** | SPRYD4 | 0.136556624 | 3.182804683 | 1.527587104 | 0.173907061 | 0.574454961 | -5.846358309 |
| **A6NED2** | RCCD1 | -0.038209167 | 3.192493036 | -1.160705054 | 0.286782795 | 0.682301253 | -5.847547289 |
| **Q9HBD1-1** | RC3H2 | -0.039518683 | 3.22606949 | -0.690945309 | 0.513610986 | 0.83148099 | -5.848013041 |
| **Q7Z7L9-1** | ZSCAN2 | 0.0316711 | 3.218006873 | 0.400688312 | 0.701546167 | 0.914634772 | -5.848063363 |
| **Q9BUE0** | MED18 | -0.042975441 | 3.183238041 | -1.592635096 | 0.158750564 | 0.557345662 | -5.848372235 |
| **Q9P2Y4** | ZNF219 | 0.000238686 | 3.157952778 | 0.003124955 | 0.997600779 | 0.998480498 | -5.84865475 |
| **Q9HBE1-1** | PATZ1 | 0.038640268 | 3.230966557 | 0.464595978 | 0.657444108 | 0.896852471 | -5.849148348 |
| **Q969W8-2** | ZNF566 | -0.017025505 | 3.174167655 | -0.20431963 | 0.844379588 | 0.962543524 | -5.849183696 |
| **Q9BZR6** | RTN4R | 0.046639863 | 3.098085218 | 0.711357347 | 0.501689804 | 0.826806835 | -5.849923725 |
| **Q6P1L5-1** | FAM117B | 0.049908347 | 3.169336237 | 1.617488324 | 0.153293663 | 0.552031202 | -5.849935889 |
| **Q9H4B6** | SAV1 | -0.050288169 | 3.192285825 | -1.135981801 | 0.296276798 | 0.688880164 | -5.850411811 |
| **A6NKF1** | SAC3D1 | -0.035161581 | 3.183805688 | -1.335682923 | 0.226711819 | 0.632674886 | -5.850491651 |
| **Q6AHZ1-1** | ZNF518A | 0.042674343 | 3.186104008 | 1.309679991 | 0.234878494 | 0.64164013 | -5.850845571 |
| **Q96GZ6-9** | SLC41A3 | -0.012905483 | 3.172161085 | -0.167556212 | 0.872047221 | 0.971931713 | -5.851030736 |
| **Q9P1Z2-1** | CALCOCO1 | -0.043091217 | 3.182515065 | -1.404081452 | 0.206423108 | 0.612058044 | -5.851176329 |
| **P63241-1** | EIF5A | -0.094488221 | 3.162566539 | -1.586891955 | 0.160037289 | 0.55963744 | -5.851854757 |
| **Q5M7Z0** | RNFT1 | -0.043765748 | 3.208036318 | -0.726070737 | 0.493210754 | 0.821156631 | -5.853452502 |
| **P54252** | ATXN3 | 0.042476768 | 3.17051447 | 1.415791046 | 0.203118113 | 0.608695713 | -5.853601083 |
| **P46379-3** | BAG6 | 0.070903305 | 3.201924048 | 2.02368702 | 0.085994257 | 0.432694348 | -5.854035602 |
| **P19793** | RXRA | 0.040087314 | 3.230475269 | 0.594724918 | 0.572223402 | 0.857633451 | -5.85437908 |
| **O43766-1** | LIAS | 0.136006398 | 3.177011513 | 1.259224272 | 0.251464855 | 0.655935708 | -5.854453717 |
| **Q9NRH1** | YAE1D1 | -0.048784788 | 3.20218936 | -1.105644492 | 0.308282454 | 0.696616637 | -5.854778734 |
| **Q9P209** | CEP72 | 0.03575784 | 3.209687715 | 1.269943168 | 0.247858217 | 0.652280071 | -5.855579684 |
| **O95716** | RAB3D | -0.049433282 | 3.2533387 | -0.75125629 | 0.47892024 | 0.811537935 | -5.855658986 |
| **P19438-1** | TNFRSF1A | 0.032497248 | 3.15035583 | 0.442651276 | 0.672432846 | 0.90142081 | -5.855685565 |
| **Q96B86-4** | RGMA | 0.014775618 | 3.202091552 | 0.19761261 | 0.849410679 | 0.964964904 | -5.856663927 |
| **Q8N1A6** | C4orf33 | -0.035765786 | 3.135371564 | -0.509857073 | 0.62707191 | 0.885387512 | -5.856913793 |
| **O95298-1** | NDUFC2; NDUFC2- | 0.147055909 | 3.173921032 | 1.713218628 | 0.13389079 | 0.521767816 | -5.857060062 |
| **P51397** | DAP | 0.06762432 | 3.181615415 | 1.244767842 | 0.256401084 | 0.659782078 | -5.857106167 |

| | | | | | | | |
|---|---|---|---|---|---|---|---|
| **Q16828-1** | DUSP6 | -0.029422018 | 3.156456979 | -0.214615157 | 0.836672134 | 0.960912428 | -5.857369953 |
| **Q6ZWJ1-1** | STXBP4 | -0.114197384 | 3.221357505 | -0.790689027 | 0.45711744 | 0.798548386 | -5.857675944 |
| **P30291-1** | WEE1 | -0.033721845 | 3.233550106 | -1.720221379 | 0.13256756 | 0.519961998 | -5.858385355 |
| **Q8NDT2** | RBM15B | 0.08930669 | 3.171430282 | 1.706677284 | 0.135138277 | 0.523998833 | -5.858735719 |
| **P30305-1** | CDC25B | -0.042201271 | 3.149377358 | -1.24659894 | 0.255771247 | 0.659782078 | -5.858855313 |
| **P63173** | RPL38 | -0.056624809 | 3.200493473 | -1.741618418 | 0.128601848 | 0.512665171 | -5.859497392 |
| **Q9NWS9** | ZNF446 | -0.039774268 | 3.149021942 | -0.645395608 | 0.540867 | 0.843540088 | -5.859687359 |
| **Q9H0F6-1** | SHARPIN | 0.047896194 | 3.194616893 | 1.245909471 | 0.256008245 | 0.659782078 | -5.860795099 |
| **Q8IY37** | DHX37 | -0.060926071 | 3.198732866 | -2.082925838 | 0.079011372 | 0.416709711 | -5.861438942 |
| **P01024** | C3 | 0.048384113 | 3.170960866 | 0.547282975 | 0.602535591 | 0.872497827 | -5.861467531 |
| **Q9HC44** | GPBP1L1 | -0.045977285 | 3.185348971 | -1.27173481 | 0.247259782 | 0.652280071 | -5.861775946 |
| **Q4U2R6** | MRPL51 | 0.129292322 | 3.188645251 | 1.435400216 | 0.197690617 | 0.603708078 | -5.861955042 |
| **P37840-1** | SNCA | 0.048033954 | 3.177956902 | 0.662255733 | 0.530673939 | 0.839849063 | -5.862401774 |
| **O96013-1** | PAK4 | -0.035317912 | 3.161078419 | -0.627547283 | 0.551789331 | 0.847634615 | -5.863183632 |
| **Q9NYL2-2** | ZAK; MAP3K20 | -0.023303897 | 3.165380894 | -0.336960067 | 0.74680907 | 0.934974996 | -5.863683832 |
| **Q8N3J3** | C17orf53 | 0.058283789 | 3.195802255 | 1.036586798 | 0.33710161 | 0.721128781 | -5.86391193 |
| **Q9BY11** | PACSIN1 | -0.066011231 | 3.168008224 | -0.530748812 | 0.613309053 | 0.879238619 | -5.864433618 |
| **Q15048** | LRRC14 | 0.072087918 | 3.191474149 | 1.429237949 | 0.199381884 | 0.604614147 | -5.864614697 |
| **Q86WH2-1** | RASSF3 | -0.035665511 | 3.210393088 | -0.749170626 | 0.480092902 | 0.81235548 | -5.866070537 |
| **Q99835** | SMO | 0.019478736 | 3.126043221 | 0.254682369 | 0.806870472 | 0.954666553 | -5.866938686 |
| **Q9H8Y5** | ANKZF1 | -0.036936931 | 3.206346076 | -1.811320608 | 0.116462494 | 0.492702086 | -5.867883708 |
| **Q969E2-1** | SCAMP4 | -0.047298256 | 3.17812883 | -0.71969388 | 0.496873845 | 0.823442885 | -5.868806012 |
| **Q9NX57** | RAB20 | -0.010698857 | 3.157344078 | -0.133582793 | 0.897790834 | 0.978757548 | -5.868872964 |
| **Q9BYX7** | POTEKP | -0.185076163 | 3.081552864 | -0.461792563 | 0.659349564 | 0.896852471 | -5.870867201 |
| **P31321** | PRKAR1B | -0.059892568 | 3.209019083 | -1.437822718 | 0.197029326 | 0.603701501 | -5.871782008 |
| **Q93015-2** | NAT6 | 0.046368857 | 3.19107357 | 0.634142214 | 0.54773785 | 0.846115385 | -5.872652804 |
| **A6NFY4** | TMEM194B; NEMP | -0.018566849 | 3.233013107 | -0.278267014 | 0.789486815 | 0.947865984 | -5.873238363 |
| **Q8N6N7** | ACBD7 | -0.121083736 | 3.198423352 | -1.206333265 | 0.269932722 | 0.671731232 | -5.87330062 |
| **P98173-1** | FAM3A | 0.043135407 | 3.180541725 | 1.317358145 | 0.23244044 | 0.63967416 | -5.873395596 |
| **Q53T59** | HS1BP3 | 0.048947244 | 3.201355166 | 1.116850795 | 0.303801687 | 0.692399426 | -5.874089127 |
| **Q86YI8** | PHF13 | -0.062548922 | 3.176618607 | -0.904149255 | 0.39832385 | 0.762880385 | -5.875009346 |
| **Q13257-1** | MAD2L1 | -0.062549439 | 3.210970467 | -1.817597934 | 0.115425555 | 0.491192304 | -5.876129074 |
| **Q8IVS2** | MCAT | 0.069627351 | 3.166503286 | 1.356902469 | 0.220234929 | 0.625644783 | -5.876268545 |
| **Q8WVK7-1** | SKA2 | -0.067805405 | 3.178776275 | -1.343193835 | 0.22440015 | 0.629985581 | -5.876305473 |
| **Q8N3J2** | METTL4 | 0.007370889 | 3.130784032 | 0.086074073 | 0.934009556 | 0.986793807 | -5.876492498 |
| **P26373-1** | RPL13 | -0.157193499 | 3.15743996 | -1.926171941 | 0.09885736 | 0.458056247 | -5.876552494 |
| **Q99613** | EIF3C | -0.044254021 | 3.205199566 | -2.352159436 | 0.05383704 | 0.376021618 | -5.876788219 |
| **A6NDU8** | C5orf51 | -0.037971506 | 3.159542603 | -1.557882427 | 0.166687471 | 0.566132944 | -5.876791637 |
| **Q14241** | TCEB3; ELOA | -0.063512454 | 3.191188077 | -1.606634683 | 0.155654647 | 0.554834464 | -5.87704136 |

| | | | | | | | |
|---|---|---|---|---|---|---|---|
| **Q6PGQ7** | BORA | -0.054596076 | 3.230563751 | -0.861647542 | 0.419659798 | 0.775968093 | -5.877088467 |
| **Q5TGY3** | AHDC1 | -0.055454685 | 3.189330543 | -1.538414226 | 0.171294129 | 0.571100234 | -5.87741748 |
| **O94806** | PRKD3 | -0.010888133 | 3.223227913 | -0.151728562 | 0.884021846 | 0.975295424 | -5.877457156 |
| **Q9Y4F9** | FAM65B; RIPOR2 | -0.032162573 | 3.161044124 | -0.449128072 | 0.667991719 | 0.90001396 | -5.877684293 |
| **P15884-3** | TCF4 | -0.014815886 | 3.186081567 | -0.190683185 | 0.854616642 | 0.967777825 | -5.877818384 |
| **P51955** | NEK2 | -0.012693014 | 3.149285197 | -0.131559482 | 0.899328598 | 0.978757548 | -5.877921439 |
| **P56199** | ITGA1 | 0.007929242 | 3.194772707 | 0.114770919 | 0.912105819 | 0.981900848 | -5.878014257 |
| **O75716** | STK16 | 0.004943288 | 3.147739504 | 0.05198741 | 0.960106728 | 0.990784251 | -5.878120556 |
| **Q9NUJ3** | TCP11L1 | 0.054035007 | 3.180701812 | 1.396110364 | 0.2087006 | 0.614302382 | -5.878441581 |
| **P48551-1** | IFNAR2 | -0.044857639 | 3.180586152 | -0.710151952 | 0.502388694 | 0.826906604 | -5.879336065 |
| **Q9Y606** | PUS1 | 0.07443535 | 3.216227325 | 1.642408765 | 0.148000116 | 0.546231571 | -5.879653753 |
| **Q15121-2** | PEA15 | -0.203811285 | 3.231551931 | -1.239781673 | 0.258122951 | 0.660717666 | -5.879672429 |
| **Q9BQI0-2** | AIF1L | -0.033447193 | 3.218515685 | -0.5505048 | 0.60044869 | 0.872406697 | -5.88034103 |
| **Q08050-2** | FOXM1 | -0.00573326 | 3.0888641 | -0.081477751 | 0.937524178 | 0.987401606 | -5.880975439 |
| **O95294-3** | RASAL1 | 0.022065552 | 3.182302953 | 0.306195563 | 0.769069822 | 0.941649022 | -5.881211191 |
| **O14975** | SLC27A2 | -0.036382521 | 3.193062959 | -0.775838182 | 0.465246333 | 0.803299278 | -5.88122861 |
| **Q9Y483** | MTF2 | 0.065616983 | 3.206919078 | 1.07581108 | 0.320476274 | 0.708257023 | -5.881999114 |
| **P13196** | ALAS1 | 0.032205147 | 3.240405889 | 0.367959664 | 0.724641505 | 0.926648409 | -5.88293164 |
| **Q15427** | SF3B4 | 0.043280343 | 3.173171234 | 1.311281545 | 0.2343681 | 0.64164013 | -5.882975483 |
| **Q7Z7E8-1** | UBE2Q1 | -0.044850057 | 3.185742238 | -1.475133989 | 0.187095671 | 0.590381098 | -5.883381476 |
| **Q70EL1-1** | USP54 | -0.042350568 | 3.211286339 | -0.738897146 | 0.485897561 | 0.81473183 | -5.883791191 |
| **Q9BUK0-4** | CHCHD7 | -0.015627991 | 3.160773873 | -0.186590261 | 0.857695319 | 0.968855822 | -5.884124131 |
| **Q15116** | PDCD1 | 0.011253693 | 3.26119926 | 0.145057415 | 0.88907911 | 0.976409291 | -5.884749475 |
| **Q14155-5** | ARHGEF7 | -0.047779499 | 3.175471958 | -1.720448555 | 0.132524844 | 0.519961998 | -5.885430615 |
| **O15116** | LSM 1.00 | 0.04274584 | 3.167833532 | 1.613158962 | 0.154231347 | 0.552714504 | -5.885876873 |
| **Q9NS00-1** | C1GALT1 | -0.033069838 | 3.197359753 | -0.61717207 | 0.558200146 | 0.851117349 | -5.885961478 |
| **P04632** | CAPNS1 | -0.039832659 | 3.154402262 | -1.672468981 | 0.141845401 | 0.538186108 | -5.886420801 |
| **Q9Y6I7** | WSB1 | -0.040797583 | 3.205591315 | -1.097608529 | 0.311528957 | 0.699575343 | -5.886460849 |
| **P22670** | RFX1 | -0.164001435 | 3.221666736 | -1.575032839 | 0.162725296 | 0.561580888 | -5.886922641 |
| **O43150-1** | ASAP2 | 0.036971148 | 3.149589783 | 0.682751473 | 0.518447747 | 0.833791899 | -5.887242291 |
| **Q86Y13** | DZIP3 | -0.024706197 | 3.182587824 | -0.420695166 | 0.687593537 | 0.907431059 | -5.887366681 |
| **P52824** | DGKQ | -0.025694264 | 3.127316001 | -0.466331792 | 0.656265679 | 0.896497734 | -5.887468606 |
| **P11308-3** | ERG | 0.033720248 | 3.141064047 | 0.633870179 | 0.547904607 | 0.846115385 | -5.888044569 |
| **E7ETH6** | ZNF587B | -0.029984995 | 3.229337828 | -0.538456736 | 0.608273465 | 0.875812373 | -5.888391005 |
| **Q9NWQ4** | GPATCH2L | 0.072409163 | 3.224129983 | 0.44953568 | 0.667712705 | 0.90001396 | -5.888700898 |
| **Q499Z4** | ZNF672 | -0.010159644 | 3.177490077 | -0.120897317 | 0.907439714 | 0.98075504 | -5.88905953 |
| **Q9BY42** | RTFDC1 | -0.034686671 | 3.180863039 | -1.951250924 | 0.095377096 | 0.452130684 | -5.889301388 |
| **Q8TB96** | ITFG1 | 0.045340403 | 3.174586182 | 1.141474145 | 0.294145315 | 0.686742442 | -5.889593057 |
| **Q9P2J3** | KLHL9 | -0.036395926 | 3.198580766 | -0.688649651 | 0.514963141 | 0.832258372 | -5.89026602 |

| | | | | | | | |
|---|---|---|---|---|---|---|---|
| **Q8N3Y1** | FBXW8 | -0.06904981 | 3.215824097 | -0.906103623 | 0.397362586 | 0.762571436 | -5.890609718 |
| **Q07108** | CD69 | 0.010936208 | 3.176071383 | 0.110104335 | 0.91566262 | 0.98295196 | -5.890801968 |
| **Q96CW6** | SLC7A6OS | -0.065879714 | 3.224519007 | -1.038865223 | 0.336117299 | 0.720858943 | -5.890868882 |
| **Q92796-2** | DLG3 | -0.041014416 | 3.167968279 | -1.605701614 | 0.155859208 | 0.554834464 | -5.891101728 |
| **O14678** | ABCD4 | 0.008944705 | 3.204767226 | 0.146258833 | 0.888167915 | 0.97627201 | -5.891328948 |
| **A1L0T0** | ILVBL | 0.033654398 | 3.17492611 | 1.368841242 | 0.216663748 | 0.622765864 | -5.891565373 |
| **Q9Y2A7-2** | NCKAP1 | 0.055542518 | 3.123140835 | 0.50946964 | 0.627328698 | 0.885387512 | -5.891602049 |
| **Q8WV44-1** | TRIM41 | -0.046454278 | 3.176315602 | -1.297367599 | 0.238835039 | 0.644491512 | -5.893277134 |
| **Q6ZYL4** | GTF2H5 | 0.143930222 | 3.185553935 | 0.708646958 | 0.503262192 | 0.827688992 | -5.893281085 |
| **Q8NFQ8** | TOR1AIP2 | -0.034475672 | 3.206976333 | -1.756267338 | 0.125953018 | 0.509142907 | -5.893679661 |
| **Q8NC42** | RNF149 | -0.045320703 | 3.163483394 | -1.291295252 | 0.240807812 | 0.646357456 | -5.894117114 |
| **Q96DC7-2** | TMCO6 | 0.043671913 | 3.166106312 | 0.856102869 | 0.422504065 | 0.777861044 | -5.894596671 |
| **Q96NI8** | ZNF570 | 0.018832114 | 3.20725792 | 0.226818544 | 0.827561737 | 0.959077751 | -5.89504311 |
| **O14653-2** | GOSR2 | -0.054017862 | 3.182023607 | -1.544412159 | 0.169862387 | 0.569040056 | -5.895819851 |
| **Q8WY07** | SLC7A3 | 0.048331201 | 3.282844152 | 0.626098591 | 0.552681762 | 0.847927742 | -5.895909267 |
| **Q7Z2K6** | ERMP1 | -0.045283313 | 3.21556503 | -0.468573117 | 0.654745639 | 0.896497734 | -5.896617297 |
| **Q9P0K7-2** | RAI14 | -0.028069804 | 3.200182305 | -0.369657333 | 0.723435544 | 0.926648409 | -5.896884196 |
| **Q9NX09** | DDIT4 | -0.019251591 | 3.112734802 | -0.196048262 | 0.850585243 | 0.965552187 | -5.897878755 |
| **Q4AC94-5** | C2CD3 | 0.041793716 | 3.208260665 | 0.721775853 | 0.495675903 | 0.822797039 | -5.898695298 |
| **Q8N3Z6** | ZCCHC7 | 0.035898261 | 3.183119467 | 1.368342543 | 0.216811879 | 0.622765864 | -5.900071864 |
| **Q05209-1** | PTPN12 | -0.036617736 | 3.208667274 | -0.593993945 | 0.572683549 | 0.857999396 | -5.900311387 |
| **Q96AZ6-1** | ISG20 | -0.044495068 | 3.23542217 | -1.106032868 | 0.308126259 | 0.696616637 | -5.900480592 |
| **P06400** | RB1 | -0.029612284 | 3.193411602 | -2.308912926 | 0.057245789 | 0.377614679 | -5.901647908 |
| **O95202-1** | LETM1 | 0.203179538 | 3.182049586 | 1.993924105 | 0.089732679 | 0.440894334 | -5.901954609 |
| **O75525-1** | KHDRBS3 | 0.054120859 | 3.216032805 | 0.624403816 | 0.553726906 | 0.848478354 | -5.902051879 |
| **Q96B02-3** | UBE2W | -0.033064008 | 3.229307882 | -0.520772675 | 0.619860351 | 0.882737137 | -5.902848354 |
| **Q92664** | GTF3A | -0.046399982 | 3.182163421 | -1.557656656 | 0.166740225 | 0.566132944 | -5.903388118 |
| **Q8IUX4-1** | APOBEC3F | -0.059255034 | 3.191065693 | -0.433990878 | 0.678393567 | 0.903662615 | -5.903930781 |
| **Q14257-2** | RCN2 | -0.050123445 | 3.172852371 | -1.614388196 | 0.153964562 | 0.552139704 | -5.904325958 |
| **Q8NCN2** | ZBTB34 | 0.037335031 | 3.15607823 | 0.573370119 | 0.585755903 | 0.8653224 | -5.904540945 |
| **P78536-1** | ADAM17 | -0.089932152 | 3.20578337 | -1.097933284 | 0.311397215 | 0.699477204 | -5.90502641 |
| **Q9H7U1-3** | CCSER2; FAM190E | -0.002381235 | 3.216826759 | -0.025274406 | 0.980597627 | 0.994229224 | -5.905078201 |
| **P49720** | PSMB3 | -0.03356028 | 3.169915806 | -1.744947591 | 0.127995186 | 0.512600477 | -5.905327686 |
| **Q9HCP0-1** | CSNK1G1 | 0.041644439 | 3.186937084 | 1.121693279 | 0.301882163 | 0.691344519 | -5.905661484 |
| **Q96JA3** | PLEKHA8 | -0.035327426 | 3.174827399 | -0.614837697 | 0.559648751 | 0.851665881 | -5.906604634 |
| **P20226-1** | TBP | -0.020064647 | 3.171281542 | -0.255563154 | 0.806219052 | 0.954293355 | -5.906672935 |
| **Q96GY3** | LIN37 | 0.03680773 | 3.18338129 | 0.701803214 | 0.507246884 | 0.82913633 | -5.906771269 |
| **Q8IXK2** | GALNT12 | -0.039187124 | 3.200850231 | -1.2659701 | 0.249189777 | 0.654483563 | -5.907068234 |
| **P13995** | MTHFD2 | 0.099912617 | 3.222910942 | 1.676020512 | 0.141134542 | 0.536483353 | -5.90707396 |

| | | | | | | | |
|---|---|---|---|---|---|---|---|
| **P42704** | LRPPRC | 0.152263686 | 3.195527766 | 2.521194694 | 0.04240546 | 0.355540374 | -5.907279976 |
| **Q9UBC2-2** | EPS15L1 | -0.038100808 | 3.188175014 | -1.554638656 | 0.167446924 | 0.566424749 | -5.908575034 |
| **Q5RL73-2** | RBM48 | 0.050582112 | 3.155444054 | 0.678058146 | 0.521231358 | 0.834809173 | -5.908716022 |
| **Q9Y6R0** | NUMBL | 0.018701713 | 3.198839956 | 0.276648747 | 0.790675567 | 0.948368382 | -5.908965566 |
| **Q93096** | PTP4A1 | 0.016657216 | 3.212683764 | 0.218995767 | 0.833398577 | 0.96068017 | -5.909177256 |
| **Q9UNY4-1** | TTF2 | -0.0454108 | 3.168380582 | -2.061802981 | 0.081433423 | 0.423686898 | -5.909568933 |
| **Q9H0H3** | KLHL25 | 0.065402849 | 3.18159208 | 0.997726777 | 0.354245879 | 0.734057823 | -5.909581672 |
| **Q9H425** | C1orf198 | -0.049097427 | 3.257455907 | -1.110422979 | 0.30636519 | 0.694853393 | -5.909589919 |
| **P31942** | HNRNPH3 | -0.004712227 | 3.19703667 | -0.062535426 | 0.952023703 | 0.989472196 | -5.910345937 |
| **Q9BVC3** | DSCC1 | 0.029546542 | 3.187785507 | 1.655897918 | 0.145207508 | 0.543107211 | -5.911559348 |
| **Q96IF1** | AJUBA | -0.036328667 | 3.163298582 | -0.453413418 | 0.665061195 | 0.898624353 | -5.91173898 |
| **O00221** | NFKBIE | 0.066761229 | 3.181044622 | 1.085003902 | 0.316677609 | 0.703974147 | -5.91178439 |
| **P33316-3** | DUT | 0.114246584 | 3.162048082 | 0.647027106 | 0.539875342 | 0.843540088 | -5.911786268 |
| **P54198** | HIRA | 0.026753834 | 3.170556455 | 1.729183758 | 0.130892363 | 0.517898318 | -5.912045526 |
| **Q9BT22** | ALG1 | -0.053882931 | 3.182018071 | -0.905404613 | 0.397706197 | 0.762880385 | -5.912051768 |
| **Q8N556-1** | AFAP1 | -0.050113716 | 3.240087134 | -0.414792995 | 0.691696219 | 0.909855374 | -5.912648269 |
| **P07205** | PGK 2.00 | 0.048547448 | 3.176218029 | 1.137219589 | 0.295795315 | 0.688880164 | -5.912916266 |
| **Q92569** | PIK3R3 | -0.059837345 | 3.156009854 | -0.837881739 | 0.431949842 | 0.783333621 | -5.914359226 |
| **Q8IWP9** | CCDC28A | -0.033230775 | 3.055269273 | -0.432488336 | 0.6794303 | 0.903893384 | -5.91521394 |
| **P51825-2** | AFF1 | 0.030693618 | 3.126926118 | 0.577877014 | 0.582884509 | 0.864802507 | -5.915270406 |
| **O00763** | ACACB | 0.037685832 | 3.170622635 | 0.556654105 | 0.59647688 | 0.871378088 | -5.915505459 |
| **Q9NPB0** | SAYSD1 | -0.001172438 | 3.210811055 | -0.018477679 | 0.985814451 | 0.995531556 | -5.91620769 |
| **Q9NYF3** | FAM53C | -0.019802744 | 3.233497738 | -0.251327439 | 0.809353272 | 0.955498404 | -5.917631907 |
| **Q99836-6** | MYD88 | -0.029981759 | 3.197593753 | -1.636083464 | 0.149327094 | 0.546464774 | -5.917683678 |
| **P09132-1** | SRP19 | -0.042856802 | 3.167179534 | -1.31038559 | 0.234653508 | 0.64164013 | -5.918077532 |
| **Q7L5Y6-2** | DET1 | -0.028542149 | 3.22853109 | -0.432212203 | 0.679620911 | 0.903893384 | -5.918209205 |
| **Q7Z4W1** | DCXR | 0.091999839 | 3.194529799 | 1.793505198 | 0.119455028 | 0.49819958 | -5.918461031 |
| **O60292** | SIPA1L3 | -0.040032326 | 3.204328645 | -1.107717413 | 0.30744953 | 0.696120409 | -5.918516277 |
| **P11021** | HSPA5 | 0.055420664 | 3.212944205 | 2.460647616 | 0.046178485 | 0.358190865 | -5.918615291 |
| **Q96B97-2** | SH3KBP1 | -0.048830975 | 3.180426697 | -0.984533316 | 0.360220419 | 0.73661838 | -5.919034393 |
| **Q96BF3** | TMIGD2 | -0.027349732 | 3.186504972 | -0.16758693 | 0.872024015 | 0.971931713 | -5.919063356 |
| **P60953** | CDC42 | 0.08864042 | 3.132612717 | 1.124691132 | 0.300698875 | 0.691292041 | -5.919162779 |
| **Q04727-3** | TLE4 | 0.043603277 | 3.140055589 | 0.565614189 | 0.590716387 | 0.868764508 | -5.919880765 |
| **P32119** | PRDX2 | 0.115615453 | 3.168176961 | 1.810866003 | 0.116537941 | 0.492758883 | -5.920251252 |
| **P49006** | MARCKSL1 | -0.38952713 | 3.115902303 | -1.750548855 | 0.126980713 | 0.510041336 | -5.920600071 |
| **Q96DU3-1** | SLAMF6 | -0.04242241 | 3.165817392 | -1.076201395 | 0.320314237 | 0.708257023 | -5.920924993 |
| **O15439-1** | ABCC4 | 0.056062582 | 3.184700844 | 0.969810052 | 0.366980331 | 0.741709165 | -5.921018222 |
| **Q92833** | JARID2 | -0.024364296 | 3.221312131 | -0.347575364 | 0.739187384 | 0.931081875 | -5.921236412 |
| **Q14938-3** | NFIX | -0.04515563 | 3.220846293 | -1.205011655 | 0.270408687 | 0.671731232 | -5.921821219 |

| | | | | | | | |
|---|---|---|---|---|---|---|---|
| **Q69YL0** | LOC152217; NCBP | 0.095935825 | 3.146589006 | 0.87362707 | 0.41356256 | 0.771122868 | -5.922294798 |
| **Q5H9F3-3** | BCORL1 | -0.044181933 | 3.209643635 | -1.213211016 | 0.267467289 | 0.67010832 | -5.922619097 |
| **Q02153** | GUCY1B3 | 0.048441212 | 3.14266734 | 0.827999789 | 0.437135948 | 0.786646683 | -5.922659124 |
| **Q8IZ13** | C5orf54; ZBED8 | 0.01695044 | 3.207515739 | 0.241810307 | 0.816409311 | 0.956409905 | -5.922803084 |
| **A6NHX0** | GATSL2; CASTOR2 | -0.007222842 | 3.178625952 | -0.078273115 | 0.939975538 | 0.988002681 | -5.922952031 |
| **P30042-1** | C21orf33; LOC1027 | 0.175912644 | 3.162927877 | 1.829999991 | 0.113403283 | 0.487284523 | -5.923199094 |
| **O75398** | DEAF1 | 0.047869188 | 3.201062145 | 0.817954892 | 0.442453163 | 0.789550692 | -5.923467477 |
| **Q92536** | SLC7A6 | 0.019029795 | 3.235123675 | 0.198352319 | 0.848855426 | 0.964964904 | -5.924351116 |
| **Q9BVN2-2** | RUSC1 | 0.039000879 | 3.148346409 | 0.369834533 | 0.723309717 | 0.926648409 | -5.924584208 |
| **Q8NC67** | NETO2 | -0.023373012 | 3.160419352 | -0.227442747 | 0.827096503 | 0.959077751 | -5.924805776 |
| **Q9NZC9** | SMARCAL1 | 0.031072167 | 3.190150726 | 1.714180647 | 0.13370826 | 0.521508161 | -5.924947193 |
| **Q9C040-2** | TRIM2 | -0.036857943 | 3.231201732 | -0.910134994 | 0.395385243 | 0.761244496 | -5.925414169 |
| **Q9H930-4** | SP140L | 0.035365119 | 3.197430982 | 1.438419527 | 0.196866719 | 0.603701501 | -5.92667986 |
| **P10124** | SRGN | -0.018201044 | 3.162415017 | -0.236860356 | 0.820086621 | 0.958501895 | -5.928033449 |
| **P49327** | FASN | -0.041155871 | 3.174760608 | -2.562979345 | 0.039990497 | 0.355476279 | -5.928127177 |
| **Q96BU1-1** | S100PBP | -0.034343519 | 3.102464726 | -1.243912085 | 0.256695895 | 0.659782078 | -5.92933713 |
| **Q96S90-1** | LYSMD1 | -0.052618723 | 3.20343412 | -1.219734886 | 0.265146523 | 0.66823064 | -5.930375264 |
| **Q9UK76** | HN1; JPT1 | -0.419772473 | 3.137195843 | -1.589555165 | 0.159439396 | 0.558776357 | -5.93059133 |
| **O95297-1** | MPZL1 | -0.044626663 | 3.164488347 | -0.855991882 | 0.422561141 | 0.777861044 | -5.93400232 |
| **Q5JSJ4** | DDX26B; INTS6L | -0.040595481 | 3.183989985 | -1.141015108 | 0.294322969 | 0.686955342 | -5.934142988 |
| **Q9UKF6** | CPSF3 | -0.048708076 | 3.192787203 | -2.028035328 | 0.085461275 | 0.431389971 | -5.934598533 |
| **Q8IWA5** | SLC44A2 | -0.019795948 | 3.121982798 | -0.202651896 | 0.845629865 | 0.963091926 | -5.934621412 |
| **Q8IYX8** | CEP57L1 | 0.004169638 | 3.178209019 | 0.057899662 | 0.955575428 | 0.989472196 | -5.934876268 |
| **Q6W2J9-1** | BCOR | -0.030762289 | 3.210624811 | -1.867157748 | 0.107549533 | 0.476564996 | -5.93506006 |
| **O43463-2** | SUV39H1 | -0.03534187 | 3.201468846 | -1.539831895 | 0.170954714 | 0.570709295 | -5.935124604 |
| **Q9GZU7-2** | CTDSP1 | 0.055264816 | 3.17977512 | 0.477926131 | 0.64842183 | 0.893481222 | -5.93545541 |
| **Q9Y296** | TRAPPC4 | -0.080477491 | 3.205414013 | -1.551980202 | 0.168071743 | 0.566515005 | -5.935609299 |
| **Q9BV23** | ABHD6 | -0.054395676 | 3.198375159 | -0.916980156 | 0.392044743 | 0.759806823 | -5.935869576 |
| **Q6ZVD8-1** | PHLPP2 | -0.026485359 | 3.211500481 | -0.484088574 | 0.644272406 | 0.891474831 | -5.93595489 |
| **P62847-4** | RPS24 | -0.111631577 | 3.171967702 | -1.984346543 | 0.09096985 | 0.443577837 | -5.936172686 |
| **Q8N1I0-3** | DOCK4 | -0.063044912 | 3.18039325 | -0.747034005 | 0.481296235 | 0.81235548 | -5.936290427 |
| **Q5JXC2** | MIIP | -0.03783973 | 3.21318983 | -0.399799866 | 0.702168734 | 0.914845949 | -5.937075428 |
| **Q92478** | CLEC2B | 0.038484336 | 3.146648095 | 0.853855163 | 0.423661076 | 0.778083043 | -5.93761448 |
| **P62854** | RPS26; LOC10192 | -0.154172732 | 3.149332651 | -1.559788505 | 0.166242707 | 0.565410236 | -5.938019914 |
| **Q9H0C5** | BTBD1 | -0.05319741 | 3.202012558 | -1.130668924 | 0.29835085 | 0.689670498 | -5.938022814 |
| **Q14168-4** | MPP2 | -0.012510029 | 3.227835084 | -0.167475238 | 0.872108393 | 0.971931713 | -5.938091664 |
| **Q9UJK0** | TSR3 | -0.108692252 | 3.217243951 | -1.212821996 | 0.267606223 | 0.67010832 | -5.938263104 |
| **Q13191-1** | CBLB | 0.03730222 | 3.149925182 | 1.361686196 | 0.218797738 | 0.625479172 | -5.938371378 |
| **Q8TCN5** | ZNF507 | -0.036579594 | 3.235028205 | -1.242734432 | 0.257102079 | 0.659782078 | -5.938442748 |

| | | | | | | | |
|---|---|---|---|---|---|---|---|
| **Q9BY66-3** | KDM5D | -0.017896981 | 3.123235117 | -0.175504685 | 0.866047237 | 0.970432241 | -5.939261729 |
| **P33240** | CSTF2 | -0.061178076 | 3.188785925 | -1.470956252 | 0.18818471 | 0.591661069 | -5.939272871 |
| **Q6P4I2** | WDR73 | 0.039568276 | 3.199900199 | 1.313507072 | 0.233660475 | 0.640812038 | -5.939829262 |
| **Q9BR11** | ZSWIM1 | -0.041228443 | 3.199366251 | -0.660875302 | 0.531503921 | 0.839849063 | -5.939968528 |
| **Q9NZE8** | MRPL35 | 0.105418564 | 3.199382174 | 1.20310502 | 0.271096605 | 0.671838876 | -5.940394753 |
| **O95674-1** | CDS2 | -0.03669022 | 3.194040075 | -1.45787629 | 0.191632002 | 0.596505086 | -5.940452511 |
| **Q86YB8** | ERO1LB; ERO1B | -0.038848513 | 3.139604404 | -0.369142154 | 0.723801418 | 0.926648409 | -5.94080329 |
| **Q9H2L5-1** | RASSF4 | -0.030124701 | 3.229638212 | -0.330373702 | 0.751553653 | 0.935635612 | -5.940863324 |
| **P07738** | BPGM | 0.042392059 | 3.155029506 | 1.117377859 | 0.303592274 | 0.692399426 | -5.941008415 |
| **Q16831** | UPP1 | 0.030524307 | 3.207285237 | 0.503267339 | 0.631447188 | 0.887152592 | -5.94113308 |
| **Q9UJJ7** | RPUSD1 | 0.00989866 | 3.267958025 | 0.112724959 | 0.913664954 | 0.982219313 | -5.94158487 |
| **Q86XR7-2** | TMED7-TICAM2; TI | -0.037036619 | 3.181211584 | -1.539992659 | 0.170916264 | 0.570709295 | -5.941872403 |
| **Q9BYN8** | MRPS26 | 0.10008161 | 3.162974826 | 0.831928065 | 0.435069035 | 0.784298014 | -5.942462139 |
| **Q8N7R7-1** | CCNYL1 | -0.026736987 | 3.154733766 | -0.356825884 | 0.732571503 | 0.928164811 | -5.943237936 |
| **P21796** | VDAC1 | 0.121545603 | 3.170133406 | 2.028990038 | 0.085344698 | 0.431338184 | -5.943332215 |
| **Q9UKE5-4** | TNIK | 0.037395305 | 3.210115343 | 0.679495014 | 0.520378135 | 0.834650661 | -5.944527987 |
| **Q9NQG6** | SMCR7L; MIEF1 | -0.049530241 | 3.198803959 | -1.018170862 | 0.345142283 | 0.726061285 | -5.944577486 |
| **Q9BUJ0** | ABHD14A | -0.016610907 | 3.273132 | -0.295580802 | 0.776807234 | 0.94357883 | -5.944762238 |
| **O60675** | MAFK | -0.033365057 | 3.206274127 | -0.37411703 | 0.720271625 | 0.925644897 | -5.945406505 |
| **Q14689** | DIP2A | -0.031783866 | 3.18826288 | -1.681054112 | 0.140132857 | 0.535444609 | -5.946584142 |
| **Q9UHN1** | POLG2 | 0.096354119 | 3.107925308 | 1.10697854 | 0.307746205 | 0.696395215 | -5.94663003 |
| **Q9P243-1** | ZFAT | -0.010106704 | 3.196175209 | -0.155459443 | 0.881196089 | 0.974405418 | -5.946783137 |
| **P29375-1** | KDM5A | 0.046983372 | 3.20177985 | 1.911086971 | 0.10101096 | 0.463623383 | -5.947071769 |
| **Q8TD31-2** | CCHCR1 | -0.034711476 | 3.212735846 | -0.33255561 | 0.749980574 | 0.935635612 | -5.947156839 |
| **Q9BQE9-1** | BCL7B | -0.184131325 | 3.190811732 | -1.445047776 | 0.195068974 | 0.600645673 | -5.947368159 |
| **Q99942** | RNF5 | 0.025792972 | 3.202301322 | 0.45768657 | 0.662145347 | 0.897125645 | -5.947428634 |
| **A6QL63-1** | BTBD11 | 0.024477802 | 3.191622529 | 0.357472537 | 0.732109939 | 0.928164811 | -5.947450886 |
| **Q8IVT5-1** | KSR1 | 0.0140096 | 3.217197369 | 0.165139164 | 0.873873596 | 0.97239856 | -5.949781351 |
| **Q9P2M4-1** | TBC1D14 | -0.041245625 | 3.182226009 | -1.17202971 | 0.282519963 | 0.679557562 | -5.950860156 |
| **Q8N2G8-1** | GHDC | 0.004560783 | 3.236695231 | 0.060057058 | 0.953922381 | 0.989472196 | -5.950954602 |
| **Q969Q4** | ARL 11.00 | 0.050791617 | 3.239779361 | 0.7476537 | 0.480947016 | 0.81235548 | -5.951331287 |
| **Q8IY81** | FTSJ3 | -0.082401266 | 3.174480378 | -1.980514202 | 0.091469637 | 0.444209209 | -5.951565073 |
| **Q96JC1-1** | VPS39 | -0.030906323 | 3.15633551 | -1.664653904 | 0.143421665 | 0.539313633 | -5.951884533 |
| **Q6ZSJ8** | C1orf122 | -0.012104556 | 3.200878789 | -0.174886322 | 0.866513676 | 0.970722967 | -5.953247009 |
| **Q9H8G1** | ZNF430 | 0.021566655 | 3.121432492 | 0.385797286 | 0.712013491 | 0.921627108 | -5.953802629 |
| **Q93050-3** | ATP6V0A1 | -0.035151571 | 3.214899241 | -1.201671184 | 0.271614917 | 0.671838876 | -5.953880726 |
| **Q9NWB7** | IFT57 | -0.03425653 | 3.183998197 | -1.072517939 | 0.321846055 | 0.709476351 | -5.953924873 |
| **Q6MZP7** | LIN54 | 0.033044503 | 3.185436218 | 1.756071741 | 0.125988037 | 0.509142907 | -5.954201997 |
| **Q96M11** | HYLS1 | -0.036073138 | 3.267112857 | -0.277530858 | 0.790027507 | 0.947865984 | -5.955412242 |

| | | | | | | | |
|---|---|---|---|---|---|---|---|
| **Q16877** | PFKFB4 | -0.066337127 | 3.205403978 | -0.745615204 | 0.482096424 | 0.81235548 | -5.95597166 |
| **Q9HB96** | FANCE | 0.0454766 | 3.195769128 | 0.726292586 | 0.493083641 | 0.821156631 | -5.956454543 |
| **Q07812-5** | BAX | 0.049468159 | 3.158474015 | 1.32561851 | 0.229842467 | 0.637461728 | -5.957144302 |
| **Q00597** | FANCC | -0.018484422 | 3.188662016 | -0.206461296 | 0.842774727 | 0.962543524 | -5.957296044 |
| **O95081** | AGFG2 | 0.034872296 | 3.229924663 | 0.393305563 | 0.706727072 | 0.917924896 | -5.957370104 |
| **Q8IZD2-1** | KMT2E; MLL5 | -0.045800129 | 3.175117572 | -1.593746443 | 0.158502696 | 0.557345662 | -5.95763905 |
| **P18077** | RPL35A | -0.074652505 | 3.186803813 | -1.850443944 | 0.110145027 | 0.481294133 | -5.958275507 |
| **Q9H6N6** | MYH16 | -0.035842459 | 2.987136246 | -0.72933195 | 0.491344377 | 0.81976713 | -5.961717713 |
| **Q15669** | RHOH | -0.013899307 | 3.115366915 | -0.238941454 | 0.81853997 | 0.957415004 | -5.961925711 |
| **Q9NZT1** | CALML5 | -0.362963702 | -0.156590319 | -0.879593573 | 0.410550233 | 0.769906516 | -5.962442239 |
| **Q7L099-3** | RUFY3 | -0.035571722 | 3.149900749 | -0.71388394 | 0.500226963 | 0.825209342 | -5.962452099 |
| **Q96D09** | GPRASP2; ARMCX | -0.036508301 | 3.181600349 | -0.930012509 | 0.385743858 | 0.755232861 | -5.963668562 |
| **Q9NX31** | C20orf111; OSER1 | -0.013096141 | 3.194748142 | -0.081423754 | 0.937565477 | 0.987401606 | -5.964076599 |
| **Q9NPF0** | CD320 | -0.062435883 | 3.239654335 | -1.411809407 | 0.204236513 | 0.610251635 | -5.964773772 |
| **Q02078-3** | MEF2A | -0.043827295 | 3.16088192 | -0.408893154 | 0.695808586 | 0.911956208 | -5.964950994 |
| **P51649-2** | ALDH5A1 | 0.145872977 | 3.17860538 | 1.719557042 | 0.132692551 | 0.519961998 | -5.965420327 |
| **Q9BUZ4** | TRAF4 | 0.051008089 | 3.247678084 | 0.816625252 | 0.443160443 | 0.789680112 | -5.965597808 |
| **O60784-2** | TOM1 | -0.049635678 | 3.207253405 | -0.662429742 | 0.530569375 | 0.839849063 | -5.965936455 |
| **Q8WV37-1** | ZNF480 | -0.014908318 | 3.180638799 | -0.206909221 | 0.842439177 | 0.962493422 | -5.966188391 |
| **Q8IX15-3** | HOMEZ | -0.007038531 | 3.176567417 | -0.107144747 | 0.917919471 | 0.983529359 | -5.966243441 |
| **P62341** | SELT; SELENOT | 0.058119929 | 3.149488063 | 0.758494504 | 0.474865695 | 0.809268114 | -5.966414803 |
| **Q96SB3** | PPP1R9B | 0.0296932 | 3.185367727 | 0.61158258 | 0.561672515 | 0.852576894 | -5.966497495 |
| **O14965** | AURKA | -0.033604622 | 3.15592496 | -1.368526773 | 0.216757146 | 0.622765864 | -5.96672407 |
| **O43482** | OIP5 | -0.046996678 | 3.201809544 | -1.334167585 | 0.227180753 | 0.633538465 | -5.966968923 |
| **P54136-1** | RARS | -0.073249714 | 3.153425471 | -1.320693656 | 0.231388282 | 0.638769945 | -5.96703163 |
| **Q9BT92** | TCHP | -0.051825519 | 3.207887468 | -1.090844278 | 0.314283386 | 0.701837878 | -5.968011434 |
| **Q7Z5G4-1** | GOLGA7 | -0.074933482 | 3.160934005 | -1.117209282 | 0.30365924 | 0.692399426 | -5.968505637 |
| **Q96RF0** | SNX18 | -0.043415279 | 3.185863685 | -0.788079291 | 0.458538705 | 0.799639769 | -5.968706985 |
| **Q3MHD2-2** | LSM 12.00 | -0.060428859 | 3.177308813 | -1.063293946 | 0.325708084 | 0.710703247 | -5.969173507 |
| **Q16543** | CDC37 | -0.038162734 | 3.168297723 | -1.914215659 | 0.100560517 | 0.46262496 | -5.969365346 |
| **Q8IW19** | APLF | 0.01709322 | 3.159279975 | 0.243497136 | 0.815157314 | 0.956212145 | -5.970826045 |
| **A6NKD9** | CCDC85C | 0.037026153 | 3.192994078 | 0.849775402 | 0.425767032 | 0.779559304 | -5.970941814 |
| **Q2KHR2-1** | RFX7 | -0.039615934 | 3.183675668 | -1.764378433 | 0.124509111 | 0.507034795 | -5.971522458 |
| **Q2TAZ0-1** | ATG2A | -0.036057624 | 3.132771374 | -0.607125061 | 0.564450974 | 0.852578515 | -5.971540902 |
| **Q8IYN0** | ZNF100 | -0.03451716 | 3.195939511 | -1.14885946 | 0.29129938 | 0.685264687 | -5.971962668 |
| **Q16610-4** | ECM1 | -0.231099797 | 3.019747272 | -0.391495654 | 0.707999793 | 0.91822696 | -5.972092975 |
| **Q99439** | CNN2 | 0.00905313 | 3.255338646 | 0.095879132 | 0.926517486 | 0.985528933 | -5.972872657 |
| **P07093-3** | SERPINE2 | -0.048522609 | 3.211981071 | -0.702989143 | 0.506554914 | 0.82913633 | -5.973668186 |
| **Q9BUL5-1** | PHF23 | -0.040334311 | 3.198760545 | -1.20414377 | 0.270721637 | 0.671731232 | -5.974909306 |

| | | | | | | | |
|---|---|---|---|---|---|---|---|
| **Q9HC16** | APOBEC3G | -0.044723017 | 3.167086399 | -1.593778583 | 0.158495534 | 0.557345662 | -5.974909384 |
| **Q13439-1** | GOLGA4 | -0.008179493 | 3.179100616 | -0.133961186 | 0.897503299 | 0.978757548 | -5.975263027 |
| **Q6ZTN6-3** | ANKRD13D | 0.03667118 | 3.180137589 | 0.698976012 | 0.508899002 | 0.82913633 | -5.975732045 |
| **Q8N9I9-2** | DTX3 | 0.038621591 | 3.184843523 | 0.705307465 | 0.505203997 | 0.828977366 | -5.977614318 |
| **P36957** | DLST | 0.152081816 | 3.177690517 | 1.310155947 | 0.234726711 | 0.64164013 | -5.978422525 |
| **P61968** | LMO4 | 0.022509576 | 3.160966813 | 0.340783475 | 0.744060285 | 0.934118682 | -5.978450076 |
| **Q9H0R1** | AP5M1 | -0.032748408 | 3.219644704 | -1.544795876 | 0.169771171 | 0.569040056 | -5.978632276 |
| **Q9BRR9-2** | ARHGAP9 | 0.036566408 | 3.174551469 | 1.246227549 | 0.255898885 | 0.659782078 | -5.979092303 |
| **Q5SRE5** | NUP188 | -0.046374877 | 3.202452396 | -2.140829178 | 0.072737839 | 0.404796855 | -5.979096576 |
| **Q8N5I2** | ARRDC1 | -0.105129261 | 3.234998332 | -1.135096281 | 0.296621654 | 0.688880164 | -5.979979822 |
| **Q96HA8** | WDYHV1 | 0.031612078 | 3.220922335 | 0.680866665 | 0.519564475 | 0.834217303 | -5.981210945 |
| **Q5U5X0** | LYRM7 | 0.152995034 | 3.155724804 | 1.006210663 | 0.350445343 | 0.731407657 | -5.98122693 |
| **Q96S44** | TP53RK | -0.049340566 | 3.174177119 | -1.065776539 | 0.324664964 | 0.710703247 | -5.981449599 |
| **Q8WVE0** | N6AMT2; EEF1AKM | 0.035775733 | 3.198693085 | 0.691175428 | 0.513475571 | 0.83148099 | -5.98224472 |
| **Q6NTE8** | C5orf45; MRNIP | -0.017052928 | 3.221721137 | -0.225994069 | 0.828176354 | 0.959077751 | -5.982525616 |
| **Q9NP66** | HMG20A | 0.046585867 | 3.173526208 | 1.432615351 | 0.198453317 | 0.603708078 | -5.982734257 |
| **Q96GC9** | VMP1 | 0.013335948 | 3.220746311 | 0.161957019 | 0.876279371 | 0.972759481 | -5.982873329 |
| **Q6QNY1** | BLOC1S2 | -0.072671349 | 3.21723796 | -1.202394163 | 0.271353464 | 0.671838876 | -5.984419327 |
| **P46926** | GNPDA1 | 0.039348071 | 3.178997991 | 1.76021243 | 0.125248717 | 0.508724344 | -5.984432211 |
| **Q9ULR3** | PPM1H | -0.013005036 | 3.112131295 | -0.064167292 | 0.950773715 | 0.989472196 | -5.984691718 |
| **P82932** | MRPS6 | 0.060587409 | 3.207292593 | 1.265552359 | 0.249330142 | 0.65453932 | -5.984709599 |
| **Q96Q83-1** | ALKBH3 | 0.037984918 | 3.203385121 | 1.292854087 | 0.240300024 | 0.64608585 | -5.984778336 |
| **Q9BRX9** | WDR83 | -0.032893665 | 3.169616026 | -0.92166272 | 0.389771911 | 0.75855365 | -5.984851529 |
| **Q96JP0** | FEM1C | -0.042968317 | 3.216030788 | -0.844565487 | 0.428467414 | 0.78113208 | -5.984910398 |
| **Q6ZMG9-2** | CERS6 | -0.034738171 | 3.198447141 | -0.59584774 | 0.571517011 | 0.857059768 | -5.985316768 |
| **P01732-3** | CD8A | 0.022166587 | 3.174816615 | 0.442673737 | 0.672417419 | 0.90142081 | -5.985940637 |
| **P04075** | ALDOA | 0.168328554 | 3.156063176 | 1.453286135 | 0.19285538 | 0.597362961 | -5.985956847 |
| **P48507** | GCLM | 0.0377257 | 3.181769721 | 1.671165507 | 0.142107154 | 0.538432214 | -5.986161543 |
| **Q7Z3E2** | C10orf118; CCDC1 | -0.046752063 | 3.191066387 | -1.651701458 | 0.146070878 | 0.544179725 | -5.986651417 |
| **O75907** | DGAT1 | 0.033620297 | 3.207698752 | 0.7560224 | 0.47624782 | 0.809713018 | -5.986880885 |
| **Q14789-4** | GOLGB1 | 0.031660651 | 3.163343938 | 0.678690704 | 0.52085563 | 0.834650661 | -5.987120038 |
| **O95696-2** | BRD1; LOC90834 | -0.028068291 | 3.200953571 | -0.536499694 | 0.609549826 | 0.876646363 | -5.98801799 |
| **Q00722** | PLCB2 | 0.028540818 | 3.164823716 | 0.343628999 | 0.742017167 | 0.932804809 | -5.988187166 |
| **Q7Z422** | SZRD1 | 0.043933502 | 3.177507265 | 1.188752835 | 0.276322854 | 0.675087662 | -5.988779301 |
| **Q9Y546** | LRRC42 | -0.040866235 | 3.204651735 | -1.45552655 | 0.192257372 | 0.596793208 | -5.988787901 |
| **P46020-2** | PHKA1 | -0.046324395 | 3.21235029 | -1.578897409 | 0.161844733 | 0.561414897 | -5.989090572 |
| **Q6PH81-1** | C16orf87 | -0.098624184 | 3.213177506 | -1.048679181 | 0.331903834 | 0.717739783 | -5.990968351 |
| **P83369** | LSM 11.00 | -0.042496447 | 3.174624069 | -0.906968495 | 0.396937751 | 0.762492852 | -5.991539196 |
| **O95834-3** | EML2 | 0.058270854 | 3.188678518 | 1.536706117 | 0.171703913 | 0.571776747 | -5.992022067 |

| | | | | | | | |
|---|---|---|---|---|---|---|---|
| **O95365** | ZBTB7A | -0.043786402 | 3.223203321 | -1.111245968 | 0.306035978 | 0.694598791 | -5.992745727 |
| **Q9NYL4** | FKBP11 | -0.043434673 | 3.214891975 | -1.342341041 | 0.224661562 | 0.630229362 | -5.99283584 |
| **P16070** | CD44 | -0.069783623 | 3.170798374 | -1.872337084 | 0.10675748 | 0.475441804 | -5.993423019 |
| **Q06330-1** | RBPJ | -0.040471561 | 3.186829435 | -1.805224781 | 0.117478134 | 0.494104699 | -5.994892745 |
| **P15311** | EZR | 0.022077273 | 3.178829681 | 2.06371729 | 0.081210875 | 0.422818087 | -5.995335038 |
| **A2RUB6-1** | CCDC66 | -0.024356189 | 3.244747357 | -0.415486091 | 0.69121385 | 0.909522034 | -5.996002299 |
| **Q9H2C0** | GAN | 0.037033518 | 3.21795945 | 1.284392869 | 0.243067556 | 0.649360189 | -5.996553272 |
| **Q969Q5** | RAB24 | 0.04487723 | 3.176549938 | 0.799920058 | 0.452114973 | 0.795197094 | -5.996605222 |
| **O96033** | MOCS2 | -0.070493326 | 3.150058116 | -0.458455152 | 0.661621569 | 0.897027878 | -5.99670693 |
| **Q9UH99** | SUN2 | 0.083913278 | 3.182417576 | 1.560153219 | 0.166157732 | 0.565363245 | -5.99721347 |
| **P04234** | CD3D | -0.034749477 | 3.220200656 | -1.456457042 | 0.192009504 | 0.596505086 | -5.997452096 |
| **Q8WXA3** | RUFY2 | -0.038956903 | 3.161224269 | -0.904929258 | 0.397939993 | 0.762880385 | -5.997690212 |
| **Q9BZJ0** | CRNKL1 | -0.029788862 | 3.178404289 | -1.942476865 | 0.096580659 | 0.454581358 | -5.997857947 |
| **O95628-10** | CNOT4 | -0.044571119 | 3.197062233 | -1.507487819 | 0.17885587 | 0.580540072 | -5.997862679 |
| **P57060** | RWDD2B | -0.0333726 | 3.182315359 | -0.501784144 | 0.632434192 | 0.887490806 | -5.997887269 |
| **Q9NQ89** | C12orf4 | -0.033228917 | 3.196395417 | -0.746958416 | 0.481338843 | 0.81235548 | -5.998240534 |
| **Q96PV7-1** | FAM193B | -0.036972075 | 3.199544675 | -0.9430435 | 0.379520914 | 0.750948043 | -5.9986481 |
| **P40692-1** | MLH1 | 0.033971319 | 3.198266051 | 1.755065509 | 0.126168335 | 0.509218658 | -5.998926783 |
| **Q58A45** | PAN3 | -0.017541244 | 3.173849194 | -0.285601701 | 0.784106575 | 0.946296623 | -5.998935146 |
| **Q13530** | SERINC3 | 0.089834189 | 3.112505295 | 0.866918029 | 0.416969196 | 0.773310479 | -5.999761409 |
| **Q8NCF5-1** | NFATC2IP | 0.034808735 | 3.18890681 | 1.36612544 | 0.217471535 | 0.623815474 | -5.999794568 |
| **Q96G61** | NUDT11 | -0.009511907 | 3.168861909 | -0.159327121 | 0.878268712 | 0.973200128 | -5.999908783 |
| **P58004** | SESN2 | 0.017098458 | 3.187712221 | 0.233143619 | 0.82285103 | 0.959068023 | -6.000152294 |
| **Q9BU89** | DOHH | -0.055973509 | 3.211942857 | -1.508792732 | 0.178530664 | 0.57989621 | -6.000702457 |
| **Q9Y244** | POMP | -0.055550249 | 3.186140332 | -1.552436534 | 0.167964337 | 0.566424749 | -6.001256982 |
| **O60518** | RANBP6 | -0.034259787 | 3.179831823 | -1.767920086 | 0.123883672 | 0.50630441 | -6.001611252 |
| **O43237** | DYNC1LI2 | -0.036306456 | 3.190864227 | -1.580639719 | 0.1614492 | 0.561414897 | -6.001789404 |
| **P28329** | CHAT | 0.038239319 | 3.1739213 | 0.851521963 | 0.424864534 | 0.778986255 | -6.001900528 |
| **Q6ZNA4-3** | RNF111 | 0.01193745 | 3.209566968 | 0.179483183 | 0.863047612 | 0.969524427 | -6.001924671 |
| **Q9BSB4** | C12orf44; ATG101 | 0.03449675 | 3.228523025 | 0.573307549 | 0.585795824 | 0.8653224 | -6.002005002 |
| **Q969Y2-2** | GTPBP3 | 0.064840257 | 3.154308849 | 1.129284746 | 0.298893185 | 0.690723198 | -6.0022615 |
| **Q68D85** | NCR3LG1 | -0.017036735 | 3.172057894 | -0.199911336 | 0.847685476 | 0.964069922 | -6.003078189 |
| **Q8IV50** | LYSMD2 | 0.029057251 | 3.206628892 | 0.468055546 | 0.655096491 | 0.896497734 | -6.003795228 |
| **Q68DQ2** | CRYBG3 | -0.039713599 | 3.223386571 | -1.068284731 | 0.323613837 | 0.710703247 | -6.003902517 |
| **Q8NDA8-1** | HEATR7A; MROH1 | -0.03251438 | 3.200850345 | -0.813882844 | 0.444621762 | 0.790803649 | -6.004174441 |
| **O75146** | HIP1R | -0.039750055 | 3.196720141 | -1.659209434 | 0.144529629 | 0.540879841 | -6.004260026 |
| **Q52LR7** | EPC2 | -0.034097948 | 3.175399519 | -1.193718992 | 0.274504861 | 0.674170362 | -6.004540488 |
| **Q9NZJ7** | MTCH1 | 0.115372563 | 3.170658755 | 1.484061141 | 0.184787923 | 0.587961572 | -6.004702567 |
| **Q70UQ0-4** | IKBIP | -0.038248083 | 3.228517865 | -1.067447869 | 0.323964241 | 0.710703247 | -6.00596111 |

| | | | | | | | |
|---|---|---|---|---|---|---|---|
| **Q2TAA2** | IAH1 | 0.037431376 | 3.177344235 | 1.050741812 | 0.331023696 | 0.71677637 | -6.006148368 |
| **Q9BVG4** | CXorf26; PBDC1 | -0.053774428 | 3.198931775 | -1.718007029 | 0.132984617 | 0.519961998 | -6.006926252 |
| **Q96BD0** | SLCO4A1 | -0.024969605 | 3.119878489 | -0.121585328 | 0.906915943 | 0.98075504 | -6.006933025 |
| **P06241-2** | FYN | 0.039396469 | 3.183712807 | 0.699721791 | 0.508462854 | 0.82913633 | -6.007338493 |
| **Q9NPD8** | UBE2T | -0.063020819 | 3.206773208 | -1.647082968 | 0.147026712 | 0.545853843 | -6.007677957 |
| **Q15022** | SUZ12 | -0.026343539 | 3.179696801 | -1.95779962 | 0.094488502 | 0.450024955 | -6.008057319 |
| **P28072** | PSMB6 | -0.063677955 | 3.132177528 | -1.30895163 | 0.235110937 | 0.64164013 | -6.008272958 |
| **Q6UB98-1** | ANKRD12 | -0.017973955 | 3.215681711 | -0.30884767 | 0.76714105 | 0.940879515 | -6.008728795 |
| **Q96AX9-14** | MIB2 | -0.000704423 | 3.23817491 | -0.009271268 | 0.99288198 | 0.99756865 | -6.008907006 |
| **Q7Z7L7** | ZER1 | -0.042134977 | 3.084215751 | -1.279338415 | 0.244734047 | 0.650150213 | -6.008916215 |
| **Q96TC7** | FAM82A2; RMDN3 | 0.053590546 | 3.177106666 | 1.630844521 | 0.15043469 | 0.548007159 | -6.00910376 |
| **Q9UFG5** | C19orf25 | -0.073023431 | 3.173736182 | -0.754528641 | 0.477084294 | 0.810614781 | -6.009256384 |
| **P06733-1** | ENO1 | 0.044741343 | 3.171379772 | 2.79691153 | 0.028893887 | 0.332216984 | -6.009540895 |
| **O14775-2** | GNB5 | 0.110049209 | 3.186589837 | 0.788871334 | 0.458107031 | 0.799573893 | -6.009695853 |
| **Q9UQC2-1** | GAB2 | -0.034592925 | 3.148377707 | -0.273560207 | 0.792946042 | 0.949070183 | -6.010762017 |
| **Q96CG8** | CTHRC1 | -0.045238232 | 3.151290206 | -1.000465604 | 0.353015434 | 0.732738778 | -6.010959158 |
| **Q6P4E1-4** | CASC4 | -0.041194301 | 3.183818573 | -1.554012635 | 0.167593863 | 0.566424749 | -6.011479191 |
| **P46783** | RPS10 | -0.049869336 | 3.208354968 | -2.129913493 | 0.073880725 | 0.406305157 | -6.011877548 |
| **Q16584** | MAP3K11 | -0.017800662 | 3.190650044 | -0.262244713 | 0.801282932 | 0.952917733 | -6.012296489 |
| **Q9BSJ6** | FAM64A; PIMREG | 0.028538828 | 3.173640547 | 0.371658668 | 0.722014969 | 0.926572271 | -6.012372448 |
| **Q16514** | TAF12 | 0.038525176 | 3.187212793 | 0.500484207 | 0.633299917 | 0.887509451 | -6.012883602 |
| **P25054-1** | APC | -0.035807826 | 3.203329958 | -1.361774761 | 0.218771209 | 0.625479172 | -6.012939526 |
| **P51114-2** | FXR1 | 0.023190113 | 3.163489631 | 0.465883508 | 0.656569913 | 0.896759147 | -6.013372873 |
| **Q92802-2** | N4BP2L2 | 0.068482349 | 3.202282279 | 1.432535577 | 0.198475205 | 0.603708078 | -6.013867803 |
| **Q5VST6-1** | ABHD17B; FAM108 | -0.041341419 | 3.169148384 | -1.519997418 | 0.175760688 | 0.576794163 | -6.014526971 |
| **Q7L2J0-1** | MEPCE | -0.04431856 | 3.169405365 | -1.754084334 | 0.126344384 | 0.509218658 | -6.014605961 |
| **P22001** | KCNA3 | 0.006730439 | 3.105061587 | 0.084268202 | 0.935390246 | 0.986793807 | -6.014834048 |
| **Q9NS87** | KIF15 | -0.043446816 | 3.171872077 | -2.153275074 | 0.071456574 | 0.403638325 | -6.015708908 |
| **Q9NRA0-2** | SPHK2 | -0.024933919 | 3.164049455 | -0.406885128 | 0.697210797 | 0.912576571 | -6.016549049 |
| **Q9UPW0-1** | FOXJ3 | -0.035348112 | 3.191991305 | -1.019334211 | 0.344629876 | 0.725897233 | -6.016695423 |
| **Q5VVQ6-1** | YOD1 | 0.034337015 | 3.173105254 | 0.919024262 | 0.391051342 | 0.759806823 | -6.016793962 |
| **Q5T6S3** | PHF19 | 0.047942602 | 3.218557373 | 0.878474691 | 0.411113888 | 0.769906516 | -6.01753595 |
| **Q49AM3-1** | TTC31 | -0.010910282 | 3.1837296 | -0.128417479 | 0.901717525 | 0.97952714 | -6.017923906 |
| **Q4ZIN3-1** | C19orf6; TMEM259 | -0.028719254 | 3.149035006 | -0.338179069 | 0.745932247 | 0.934974996 | -6.018383658 |
| **Q8WTS1** | ABHD5 | 0.00146486 | 3.170185572 | 0.019301288 | 0.985182245 | 0.995531556 | -6.018401441 |
| **Q5TCZ1** | SH3PXD2A | 0.039386058 | 3.197399462 | 0.822685409 | 0.439943365 | 0.788662011 | -6.019673668 |
| **Q96EP0-1** | RNF31 | 0.046181649 | 3.176720828 | 1.604691718 | 0.156080899 | 0.554834464 | -6.020297755 |
| **Q9P2D1-1** | CHD7 | -0.038208458 | 3.2083797 | -1.48414792 | 0.184765619 | 0.587961572 | -6.020428584 |
| **Q99638** | RAD9A | 0.049820451 | 3.188007454 | 1.049901409 | 0.331382075 | 0.716856834 | -6.020908365 |

| | | | | | | | |
|---|---|---|---|---|---|---|---|
| **Q9BRB3-1** | PIGQ | -0.009624709 | 3.159134736 | -0.121412383 | 0.907047598 | 0.98075504 | -6.021055928 |
| **Q9UPS8-3** | ANKRD26 | -0.006070275 | 3.172992282 | -0.06688141 | 0.948695073 | 0.989472196 | -6.021257132 |
| **Q6NVV7** | CDPF1 | -0.040128301 | 3.178838622 | -0.714034736 | 0.500139745 | 0.825209342 | -6.021394203 |
| **Q86UW6** | N4BP2 | 0.010198441 | 3.162587108 | 0.150607243 | 0.884871491 | 0.975708125 | -6.021717768 |
| **P0C869-6** | JMJD7-PLA2G4B; I | -0.008106628 | 3.197460697 | -0.071179746 | 0.945404033 | 0.98904859 | -6.022421792 |
| **Q6DKI1** | RPL7L1 | -0.126808886 | 3.265508982 | -0.983769361 | 0.360568769 | 0.736630258 | -6.023318417 |
| **P43897** | TSFM | 0.104314261 | 3.194324579 | 1.692593269 | 0.137862124 | 0.529905456 | -6.023356006 |
| **Q92902-1** | HPS1 | -0.027567948 | 3.185101763 | -0.480239051 | 0.64686284 | 0.89271426 | -6.0235694 |
| **Q9P270** | SLAIN2 | -0.038875295 | 3.201861321 | -1.637072376 | 0.14911889 | 0.546464774 | -6.025215644 |
| **P62072** | TIMM10 | -0.106136732 | 3.19877221 | -1.085788221 | 0.316355219 | 0.703848282 | -6.025537191 |
| **Q96MX3** | ZNF48 | -0.006190839 | 3.168580973 | -0.078430852 | 0.939854862 | 0.988002681 | -6.025849613 |
| **O94880-2** | PHF14 | -0.01855427 | 3.169224354 | -0.33791941 | 0.746118984 | 0.934974996 | -6.026058752 |
| **Q8TEP8-3** | CEP192 | -0.02951565 | 3.21162114 | -1.471986572 | 0.187915591 | 0.591197824 | -6.026527973 |
| **Q9BXW6-1** | OSBPL1A | 0.041304148 | 3.192952464 | 1.565651023 | 0.164881687 | 0.563893388 | -6.027113105 |
| **Q49A26-5** | GLYR1 | 0.000674421 | 3.151726806 | 0.009445424 | 0.992748276 | 0.99756865 | -6.027568945 |
| **Q9P258** | RCC2 | 0.022618939 | 3.170327989 | 2.327386685 | 0.05576324 | 0.376021618 | -6.027569172 |
| **P45984-1** | MAPK9 | -0.030334016 | 3.187164183 | -1.48317201 | 0.185016593 | 0.588198522 | -6.027752843 |
| **Q7L804-2** | RAB11FIP2 | 0.007510303 | 3.224451603 | 0.116236158 | 0.910989484 | 0.981900848 | -6.028258169 |
| **Q9NV88-1** | INTS9 | -0.028066488 | 3.164107557 | -1.798988073 | 0.118526167 | 0.496152504 | -6.028872697 |
| **Q9UMS4** | PRPF19 | 0.051056003 | 3.177567043 | 2.094438948 | 0.077721899 | 0.414504124 | -6.02971311 |
| **Q16206** | ENOX2 | 0.022316645 | 3.180948679 | 0.359602144 | 0.730590739 | 0.927984559 | -6.030358977 |
| **Q9UNK0** | STX8 | -0.036618217 | 3.19976228 | -1.33768381 | 0.226093941 | 0.63121696 | -6.030720627 |
| **P06213-1** | INSR | 0.036770929 | 3.167029053 | 1.313945723 | 0.233521224 | 0.640812038 | -6.03102499 |
| **Q2LD37-1** | KIAA1109 | 0.007704213 | 3.193488102 | 0.128555861 | 0.901612287 | 0.97952714 | -6.031048164 |
| **A6NHQ4** | C17orf96; EPOP | 0.021833745 | 3.206027686 | 0.438564096 | 0.67524278 | 0.90314827 | -6.032522843 |
| **P24864-1** | CCNE1 | -0.01008041 | 3.264748139 | -0.085515785 | 0.934436372 | 0.986793807 | -6.032974577 |
| **Q9Y4U1** | MMACHC | -0.021624738 | 3.230873026 | -0.371145627 | 0.722379019 | 0.926635956 | -6.033124892 |
| **Q53TQ3-2** | INO80D | 0.031532533 | 3.20590019 | 0.664867174 | 0.529106059 | 0.839849063 | -6.033219538 |
| **Q6PID6** | TTC33 | 0.046751204 | 3.155304562 | 1.623161065 | 0.152073163 | 0.551195838 | -6.033844218 |
| **Q765P7** | MTSS1L | -0.018789114 | 3.182560345 | -0.344009066 | 0.741744446 | 0.932740726 | -6.033959635 |
| **M5A8F1** | ERVH48-1 | -0.072414454 | 3.145258696 | -0.650247969 | 0.537920965 | 0.843454128 | -6.034497451 |
| **Q96D96-1** | HVCN1 | 0.050116734 | 3.177423644 | 1.190604541 | 0.275643798 | 0.674825663 | -6.034707284 |
| **Q6GMV3** | PTRHD1 | -0.062676482 | 3.181917634 | -1.145277218 | 0.292676913 | 0.68530312 | -6.034832081 |
| **Q13405** | MRPL49 | 0.176532922 | 3.203862514 | 1.394542842 | 0.209151126 | 0.614302382 | -6.035309355 |
| **Q53G59-2** | KLHL12 | 0.032028773 | 3.136987529 | 0.549562815 | 0.601058429 | 0.872497827 | -6.035592494 |
| **P10075** | GLI4 | 0.032284473 | 3.135591321 | 0.711071411 | 0.501855532 | 0.826806835 | -6.036271777 |
| **P07996** | THBS1 | -0.111198197 | 3.242950452 | -1.070912488 | 0.322515565 | 0.710195722 | -6.036642216 |
| **O43513** | MED7 | 0.081126394 | 3.191943022 | 0.760375244 | 0.473816033 | 0.808788313 | -6.036659615 |
| **P47897** | QARS | -0.040544421 | 3.175697262 | -2.198152996 | 0.067023818 | 0.394739983 | -6.037285382 |

| | | | | | | | |
|---|---|---|---|---|---|---|---|
| **Q86UW9** | DTX2 | -0.018046457 | 3.206480702 | -0.305188882 | 0.769802409 | 0.941649022 | -6.037430427 |
| **Q15058** | KIF14 | -0.04891938 | 3.169631501 | -1.774844702 | 0.122669593 | 0.50211742 | -6.037650502 |
| **P49842-1** | STK19 | -0.015222115 | 3.146800783 | -0.296657542 | 0.77602109 | 0.94357883 | -6.038174797 |
| **Q8IZE3** | SCYL3 | -0.032525873 | 3.187462384 | -1.159291253 | 0.287318755 | 0.682522792 | -6.038473649 |
| **Q15776** | ZKSCAN8; ZNF192 | -0.039553068 | 3.156206789 | -0.861613647 | 0.419677143 | 0.775968093 | -6.038512731 |
| **Q92997** | DVL3 | -0.030093281 | 3.191377625 | -0.615104791 | 0.559482889 | 0.851665881 | -6.038681423 |
| **Q12767-2** | KIAA0195; TMEM9 | 0.014608044 | 3.189223252 | 0.191044662 | 0.854344873 | 0.967745939 | -6.039463228 |
| **O14966-1** | RAB7L1; RAB29 | -0.030263984 | 3.201197024 | -0.539621053 | 0.607514814 | 0.875672206 | -6.039777193 |
| **Q7Z309-2** | FAM122B | 0.059119909 | 3.15826748 | 0.593502481 | 0.572993049 | 0.858139449 | -6.039860249 |
| **Q9Y3V2-1** | RWDD3 | 0.000431669 | 3.200954593 | 0.007469658 | 0.994265133 | 0.997608555 | -6.040589251 |
| **Q8N9N2-2** | ASCC1 | 0.027817057 | 3.164738882 | 1.467945984 | 0.188973015 | 0.593033086 | -6.040690915 |
| **Q96AE4** | FUBP1 | 0.049408272 | 3.204118779 | 1.120555778 | 0.302332155 | 0.691344519 | -6.040964944 |
| **Q9Y297** | BTRC | -0.01120158 | 3.162279145 | -0.207506014 | 0.841992161 | 0.962493422 | -6.04266158 |
| **Q9P2Q2** | FRMD4A | 0.022853924 | 3.112184466 | 0.462774896 | 0.658681567 | 0.896852471 | -6.042761546 |
| **Q08431** | MFGE8 | -0.050360722 | 3.211469494 | -1.572845551 | 0.163225667 | 0.561580888 | -6.042973011 |
| **Q07866-6** | KLC1 | 0.050860543 | 3.209211377 | 0.557475411 | 0.595947531 | 0.871378088 | -6.043415379 |
| **Q8N0U8** | VKORC1L1 | -0.030636781 | 3.212040368 | -0.705892367 | 0.504863541 | 0.828775136 | -6.043521589 |
| **Q9H1E5** | TMX4 | -0.004769634 | 3.132097312 | -0.075852432 | 0.941827692 | 0.988377491 | -6.043953399 |
| **P10644** | PRKAR1A | -0.054715114 | 3.15744962 | -1.815781476 | 0.115724683 | 0.492169772 | -6.044706966 |
| **O75925-2** | PIAS1 | -0.030938262 | 3.184803126 | -1.394547716 | 0.209149723 | 0.614302382 | -6.044804327 |
| **Q8IUH3-3** | RBM45 | 0.043588344 | 3.183336122 | 1.617998057 | 0.153183616 | 0.552031202 | -6.045123542 |
| **Q8NEU8-3** | APPL2 | 0.003846974 | 3.139314034 | 0.060777843 | 0.953370153 | 0.989472196 | -6.045226636 |
| **P11474-1** | ESRRA | -0.028918821 | 3.253466965 | -0.547183311 | 0.602600212 | 0.872497827 | -6.045275479 |
| **Q8NFH4** | NUP37 | -0.064559245 | 3.179936424 | -1.555269665 | 0.167298935 | 0.566424749 | -6.045419973 |
| **Q96K21** | ZFYVE19 | -0.033471604 | 3.169301465 | -0.549482082 | 0.601110704 | 0.872497827 | -6.045862648 |
| **Q9BRJ6** | C7orf50 | -0.044077759 | 3.21454872 | -1.065033401 | 0.324976927 | 0.710703247 | -6.045986906 |
| **A5PL33** | KRBA1 | 0.034390162 | 3.20781322 | 0.54025183 | 0.60710403 | 0.87523889 | -6.046090007 |
| **Q9NQC8-2** | IFT46 | 0.039066783 | 3.201644834 | 0.954953193 | 0.373900917 | 0.746240828 | -6.046320981 |
| **Q8TDW0** | LRRC8C | 0.03976582 | 3.186633917 | 1.069976178 | 0.322906549 | 0.710272801 | -6.047008277 |
| **P01040** | CSTA | 0.107275715 | 1.42530131 | 0.189006403 | 0.855877579 | 0.968791476 | -6.048592549 |
| **A4D1U4** | KIAA1147 | -0.001568374 | 3.158295009 | -0.024329512 | 0.981322825 | 0.994579747 | -6.049559241 |
| **Q96P53** | WDFY2 | 0.02281183 | 3.19996304 | 0.47326369 | 0.651570319 | 0.89454401 | -6.04974921 |
| **Q9Y6Y0** | IVNS1ABP | -0.036324242 | 3.172275983 | -1.698472366 | 0.136718776 | 0.527552539 | -6.051287505 |
| **Q8TAQ2-3** | SMARCC2 | -0.021320583 | 3.151421909 | -0.271703576 | 0.79431196 | 0.949806048 | -6.051357923 |
| **Q8TCT9-1** | HM13 | -0.05034472 | 3.197166377 | -0.890648511 | 0.405011859 | 0.766336431 | -6.051429901 |
| **P62166** | NCS1 | -0.007213505 | 3.167526778 | -0.088615443 | 0.932066956 | 0.986711787 | -6.051647519 |
| **Q9H160** | ING2 | -0.012460052 | 3.173752764 | -0.219088452 | 0.833329354 | 0.96068017 | -6.051727245 |
| **Q9Y678** | COPG1 | -0.030174995 | 3.199353933 | -1.957026065 | 0.094593037 | 0.450024955 | -6.051957916 |
| **Q8N6G5-1** | CSGALNACT2 | 0.017214659 | 3.183204577 | 0.313327553 | 0.763887097 | 0.94035993 | -6.052024805 |

| | | | | | | | |
|---|---|---|---|---|---|---|---|
| **Q6DHV7** | ADAL | 0.04185959 | 3.175602486 | 1.108258436 | 0.307232446 | 0.695827189 | -6.052247268 |
| **Q92604** | LPGAT1 | 0.035153595 | 3.17389023 | 0.506169637 | 0.629518198 | 0.886324146 | -6.053749525 |
| **Q96N66-1** | MBOAT7 | 0.015961018 | 3.208736089 | 0.241066962 | 0.816961222 | 0.956633295 | -6.053912178 |
| **Q96T49** | PPP1R16B | 0.001843386 | 3.21181993 | 0.029436751 | 0.977403315 | 0.993521701 | -6.054462868 |
| **Q9H553-1** | ALG2 | -0.029294422 | 3.192003011 | -0.434152484 | 0.678282106 | 0.903662615 | -6.054537622 |
| **Q9H936** | SLC25A22 | 0.074321088 | 3.206422739 | 1.393744728 | 0.20938085 | 0.614302382 | -6.054958713 |
| **Q15645** | TRIP13 | -0.03181211 | 3.180919233 | -1.695333983 | 0.137327985 | 0.528619594 | -6.054980127 |
| **P56277-1** | CMC4; MTCP1 | 0.021028849 | 3.187157347 | 0.291038092 | 0.780127065 | 0.94472083 | -6.054981137 |
| **Q9Y5W7** | SNX14 | -0.019867268 | 3.243589987 | -0.284197366 | 0.785135718 | 0.946571059 | -6.055119772 |
| **Q969M7-2** | UBE2F | 0.037457665 | 3.191367977 | 1.318336369 | 0.232131432 | 0.639266629 | -6.055179842 |
| **Q86V20-2** | FAM35A | 0.004459176 | 3.230835931 | 0.061327502 | 0.952949052 | 0.989472196 | -6.056421391 |
| **O95070** | YIF1A | -0.01311751 | 3.203880046 | -0.222649698 | 0.830670804 | 0.960067847 | -6.056588996 |
| **P54284** | CACNB3 | -0.00890918 | 3.1993702 | -0.125988224 | 0.9035653 | 0.980210603 | -6.05674208 |
| **P53990-5** | IST1 | -0.045751969 | 3.161159438 | -1.670025748 | 0.142336408 | 0.538432214 | -6.056848259 |
| **Q8NBX0** | SCCPDH | 0.039610933 | 3.171781368 | 1.380686681 | 0.213171749 | 0.61901473 | -6.056974055 |
| **Q8N680** | ZBTB2 | 0.038269465 | 3.181932831 | 1.120629338 | 0.302303038 | 0.691344519 | -6.057391652 |
| **Q8IXZ2-1** | ZC3H3 | -0.034436806 | 3.175602441 | -0.855388411 | 0.422871584 | 0.777884596 | -6.057392541 |
| **Q6VEQ5** | | -0.072991048 | 3.186857029 | -1.409262813 | 0.20495474 | 0.611018916 | -6.057519231 |
| **Q96Q89-3** | KIF20B | -0.038688788 | 3.197389456 | -2.041317053 | 0.083853704 | 0.427062614 | -6.057957224 |
| **O75896** | TUSC2 | -0.014733126 | 3.105478736 | -0.224185739 | 0.829524858 | 0.959466443 | -6.05886953 |
| **O75618-2** | DEDD | -0.008218881 | 3.120961297 | -0.128587168 | 0.901588479 | 0.97952714 | -6.058929055 |
| **Q8IYJ3** | SYTL1 | -0.036407066 | 3.195959327 | -1.270174 | 0.247781045 | 0.652280071 | -6.059295538 |
| **Q96T21** | SECISBP2 | -0.041957962 | 3.225719547 | -1.346621911 | 0.223352049 | 0.629489901 | -6.059337179 |
| **Q9P0R6** | GSKIP | -0.060958276 | 3.211433903 | -0.87181922 | 0.414478505 | 0.771744018 | -6.059537087 |
| **O15060** | ZBTB39 | -0.064226119 | 3.180741194 | -0.79502294 | 0.454764002 | 0.797593818 | -6.05959395 |
| **P38159-1** | RBMX | 0.05188632 | 3.165795777 | 1.713137589 | 0.133906177 | 0.521767816 | -6.06040273 |
| **Q9NXF7** | DCAF16 | -0.040343271 | 3.190817049 | -1.298289955 | 0.238536629 | 0.644491512 | -6.060541511 |
| **P53794** | SLC5A3 | -0.037152579 | 3.241666007 | -0.536384398 | 0.609625067 | 0.876646363 | -6.060797034 |
| **Q7L9B9** | EEPD1 | 0.039349446 | 3.194561596 | 0.901137161 | 0.399808774 | 0.763210165 | -6.061205228 |
| **Q8TAU3** | ZNF417 | -0.009106446 | 3.182086177 | -0.159374149 | 0.878233131 | 0.973200128 | -6.061247801 |
| **Q8N567** | ZCCHC9 | -0.040494953 | 3.20430927 | -0.694877731 | 0.511300122 | 0.83054728 | -6.061515437 |
| **Q14161-1** | GIT2 | -0.025304226 | 3.171734842 | -1.578084891 | 0.162029498 | 0.561414897 | -6.06177941 |
| **Q8N8R5** | C2orf69 | -0.040871139 | 3.190335003 | -0.889680578 | 0.405494551 | 0.766336431 | -6.06188539 |
| **Q6IC98-1** | GRAMD4 | -0.028414032 | 3.163903831 | -1.516145251 | 0.176708488 | 0.577967785 | -6.062021305 |
| **Q14344** | GNA13 | -0.038495091 | 3.17568585 | -1.685449428 | 0.139263744 | 0.533333364 | -6.062236115 |
| **Q8N137-2** | CNTROB | 0.044639811 | 3.169890209 | 0.458118539 | 0.661850941 | 0.897032706 | -6.063244456 |
| **Q6PJ61** | FBXO46 | -0.001859127 | 3.194762211 | -0.028289432 | 0.978283761 | 0.993585863 | -6.063446704 |
| **Q9Y2D4** | EXOC6B | -0.036840877 | 3.188260051 | -1.641152376 | 0.1482628 | 0.546231571 | -6.063711555 |
| **Q92994** | BRF1 | -0.061063704 | 3.187635427 | -1.370105077 | 0.216288748 | 0.622162964 | -6.06411314 |

| | | | | | | | |
|---|---|---|---|---|---|---|---|
| **O60906** | SMPD2 | -0.020454472 | 3.206324291 | -0.309161268 | 0.766913101 | 0.940879515 | -6.065720639 |
| **O14734** | ACOT8 | 0.068618026 | 3.184321819 | 1.231820994 | 0.260892658 | 0.664371245 | -6.065724216 |
| **Q9UBZ4** | APEX2 | -0.023104589 | 3.235610204 | -0.487342245 | 0.642087122 | 0.890981968 | -6.065995993 |
| **P55055** | NR1H2 | 0.027508173 | 3.244340761 | 0.680741022 | 0.519638972 | 0.834217303 | -6.06696567 |
| **Q96IR2** | ZNF845 | -0.001460163 | 3.171031011 | -0.016559428 | 0.987286952 | 0.99582925 | -6.067768167 |
| **Q9NQZ6-1** | ZC4H2 | -0.059828382 | 3.17281205 | -1.045676145 | 0.333188612 | 0.719098069 | -6.068298929 |
| **Q7L4P6** | BEND5 | -0.065966874 | 3.198905381 | -1.341819561 | 0.224821547 | 0.630229362 | -6.068715347 |
| **Q9Y478** | PRKAB1 | 0.049805635 | 3.202089515 | 1.480270192 | 0.185764705 | 0.588813889 | -6.069737527 |
| **Q8TCU4-1** | ALMS1 | -0.026621636 | 3.21998041 | -1.547597588 | 0.169106541 | 0.567857763 | -6.069774872 |
| **O00189** | AP4M1 | -0.047478228 | 3.192969915 | -1.387989251 | 0.211044201 | 0.61590617 | -6.069775314 |
| **Q71H61** | ILDR2 | -0.031266006 | 3.180724579 | -0.547383997 | 0.602470093 | 0.872497827 | -6.069802795 |
| **Q6ZMT4** | JHDM1D; KDM7A | -0.003468736 | 3.005959364 | -0.066009435 | 0.949362839 | 0.989472196 | -6.070400165 |
| **Q9UNL4-1** | ING4 | 0.028894096 | 3.15966063 | 0.551449166 | 0.599837761 | 0.872097298 | -6.07112358 |
| **Q02083** | NAAA | -0.039852442 | 3.189596476 | -0.8149422 | 0.444056869 | 0.790506794 | -6.071601948 |
| **Q96AK3** | APOBEC3D | 0.042248924 | 3.159989865 | 0.650531142 | 0.537749351 | 0.84335148 | -6.07206272 |
| **Q8TCJ0-1** | FBXO25 | -0.064168171 | 3.235880315 | -1.022080217 | 0.343422768 | 0.724892107 | -6.072137706 |
| **Q92979** | EMG1 | 0.053054627 | 3.208409576 | 1.639183252 | 0.14867539 | 0.546231571 | -6.072502444 |
| **Q4ADV7** | KIAA1432; RIC1 | 0.020501372 | 3.194883601 | 0.397471947 | 0.703801174 | 0.915921429 | -6.072712422 |
| **Q96LD8** | SENP8 | -0.026227191 | 3.128773434 | -0.345604222 | 0.740600266 | 0.931611328 | -6.073176074 |
| **P61129** | ZC3H6 | 0.054895243 | 3.201935653 | 0.784391417 | 0.460552379 | 0.801377278 | -6.073245058 |
| **Q8TAP8** | PPP1R35 | -0.04359039 | 3.172919146 | -0.807542013 | 0.44801363 | 0.792931229 | -6.073590083 |
| **Q3SXY8** | ARL13B | 0.040887962 | 3.149467091 | 1.092590799 | 0.313570293 | 0.701109468 | -6.074013729 |
| **Q9C0C2** | TNKS1BP1 | -0.036723721 | 3.190635917 | -1.779924895 | 0.121786231 | 0.500047341 | -6.074166481 |
| **Q5TKA1-2** | LIN9 | -0.031849761 | 3.187238139 | -1.433195153 | 0.198294305 | 0.603708078 | -6.074267187 |
| **Q8WYL5-1** | SSH1 | 0.078590445 | 3.237056644 | 0.554103484 | 0.598122506 | 0.87186032 | -6.075332259 |
| **Q9Y291** | MRPS33 | 0.02929503 | 3.170346511 | 0.394923023 | 0.705590544 | 0.917048401 | -6.076107525 |
| **O43504** | LAMTOR5 | -0.294758111 | 3.194155842 | -0.907663268 | 0.396596718 | 0.762021989 | -6.076278665 |
| **O00257-1** | CBX4 | -0.037693089 | 3.202444443 | -1.233027363 | 0.260471297 | 0.663969372 | -6.076646968 |
| **P33908** | MAN1A1 | 0.009477106 | 3.167537985 | 0.140394528 | 0.892617313 | 0.978276697 | -6.077428767 |
| **Q9BU23-1** | LMF2 | -0.042374298 | 3.168887098 | -0.917437229 | 0.391822448 | 0.759806823 | -6.077542082 |
| **Q9NXX6** | NSMCE4A | 0.046223316 | 3.191370096 | 1.321949137 | 0.230993341 | 0.638769945 | -6.077926389 |
| **Q9UBB5-1** | MBD2 | -0.03397101 | 3.15638401 | -1.41681825 | 0.202830483 | 0.608695713 | -6.078414008 |
| **Q8IWY8-1** | ZSCAN29 | 0.004809174 | 3.201346643 | 0.064969862 | 0.950159012 | 0.989472196 | -6.078673668 |
| **Q9UNA1-1** | ARHGAP26 | 0.038030625 | 3.184249006 | 1.23959596 | 0.258187276 | 0.660717666 | -6.078976146 |
| **Q9NZD2** | GLTP | -0.039245887 | 3.168487652 | -1.416345536 | 0.202962803 | 0.608695713 | -6.080011032 |
| **Q8TDB6-1** | DTX3L | -0.054014289 | 3.224887428 | -1.871849434 | 0.106831809 | 0.475471224 | -6.080329283 |
| **Q96EX2-1** | RNFT2 | -0.000895856 | 3.148464811 | -0.016765894 | 0.987128461 | 0.99582925 | -6.08059201 |
| **Q9Y2M0** | FAN1 | -0.031585688 | 3.214507145 | -1.321798911 | 0.231040567 | 0.638769945 | -6.081202659 |
| **P50443** | SLC26A2 | 0.012298067 | 3.206793234 | 0.174482693 | 0.866818169 | 0.970722967 | -6.081363982 |

| | | | | | | | |
|---|---|---|---|---|---|---|---|
| **P68104** | EEF1A1 | -0.047451516 | 3.215337069 | -2.566685376 | 0.039783373 | 0.355476279 | -6.0813806 |
| **Q9UJD0** | RIMS3 | 0.012699586 | 3.169822197 | 0.204470513 | 0.844266497 | 0.962543524 | -6.081390612 |
| **Q9BZM6** | ULBP1 | -0.033004777 | 3.181249103 | -0.738773346 | 0.485967796 | 0.81473183 | -6.081845747 |
| **Q9Y5Z9-1** | UBIAD1 | -0.025844252 | 3.233857 | -0.14767638 | 0.887093039 | 0.975708125 | -6.081910702 |
| **Q9H0K1** | SIK2 | -0.02968447 | 3.207197122 | -0.950707225 | 0.37589714 | 0.748684576 | -6.08204018 |
| **Q8IY31-2** | IFT20 | 0.014151518 | 3.199379587 | 0.233369454 | 0.822682981 | 0.959068023 | -6.082128951 |
| **O14610** | GNGT2 | -0.027237892 | 3.227293157 | -0.498860494 | 0.634382149 | 0.887509451 | -6.082906276 |
| **P28340** | POLD1 | -0.017269155 | 3.18477212 | -2.084559226 | 0.078827128 | 0.416709711 | -6.083972481 |
| **P23142** | FBLN1 | -0.020013382 | 3.200104185 | -0.073572145 | 0.943572781 | 0.988878215 | -6.085141662 |
| **O60684** | KPNA6 | 0.038035939 | 3.175956236 | 0.92249867 | 0.389367207 | 0.75855365 | -6.085676383 |
| **Q9Y276** | BCS1L | 0.059023266 | 3.200177505 | 1.09907089 | 0.310936087 | 0.699034299 | -6.085796947 |
| **P51826-2** | AFF3 | 0.00050754 | 3.133046037 | 0.007818674 | 0.993997181 | 0.997608555 | -6.085798266 |
| **P60660-2** | MYL6 | -0.156479085 | 3.207499723 | -1.752030941 | 0.126713586 | 0.509635359 | -6.085977071 |
| **P78415** | IRX3 | 0.024294083 | 3.139933544 | 0.376962832 | 0.71825579 | 0.924851578 | -6.086194614 |
| **Q8TDR0-1** | TRAF3IP1 | -0.036079897 | 3.172422152 | -0.767100557 | 0.470075571 | 0.806293267 | -6.087191481 |
| **Q96IW7** | SEC22A | 0.00036686 | 3.190277303 | 0.005662362 | 0.995652674 | 0.998291329 | -6.087540312 |
| **Q9H825-1** | METTL8 | -0.036516177 | 3.130989725 | -1.065141166 | 0.324931673 | 0.710703247 | -6.087700633 |
| **O43586-1** | PSTPIP1 | -0.033031071 | 3.203442175 | -0.809720367 | 0.446846314 | 0.792204208 | -6.087751711 |
| **Q99943** | AGPAT1 | 0.041226433 | 3.210989673 | 0.664687804 | 0.529213657 | 0.839849063 | -6.088021473 |
| **Q92889** | ERCC4 | -0.042485288 | 3.201978921 | -1.664124825 | 0.143528979 | 0.539320158 | -6.088215181 |
| **Q9BUH6** | C9orf142; PAXX | -0.047578631 | 3.168165895 | -1.227972617 | 0.262240741 | 0.665868547 | -6.088973921 |
| **Q9Y6M5** | SLC30A1 | -0.024830325 | 3.213069862 | -0.523310446 | 0.6181902 | 0.881939511 | -6.089226739 |
| **Q8IWV1** | LAX1 | -0.022529029 | 3.174607169 | -0.330237703 | 0.751651746 | 0.935635612 | -6.089945656 |
| **P43307** | SSR1 | -0.039714793 | 3.178920129 | -1.144294668 | 0.293055698 | 0.68530312 | -6.090576617 |
| **Q9Y2H1** | STK38L | 0.040157163 | 3.179684595 | 1.618924199 | 0.15298386 | 0.552031202 | -6.090781934 |
| **Q9H7D7** | WDR26 | -0.047838888 | 3.186171137 | -1.67084008 | 0.142172575 | 0.538432214 | -6.091304166 |
| **Q8WXB1-1** | METTL21A | 0.018802723 | 3.212936599 | 0.178423112 | 0.863846624 | 0.969738239 | -6.092074187 |
| **Q93052** | LPP | -0.038363534 | 3.184704779 | -1.485876864 | 0.184321754 | 0.586948432 | -6.092916354 |
| **Q96BD5-3** | PHF21A | 0.030861285 | 3.167745233 | 1.270150545 | 0.247788886 | 0.652280071 | -6.093398953 |
| **Q9BX59** | TAPBPL | 0.021201543 | 3.16552051 | 0.315064577 | 0.762626806 | 0.939644172 | -6.094293362 |
| **Q9Y242** | TCF19 | -0.000232655 | 3.198153903 | -0.003819026 | 0.997067901 | 0.998480498 | -6.094348508 |
| **Q86YS6** | RAB43 | 0.04347505 | 3.194255826 | 0.713725061 | 0.500318868 | 0.825209342 | -6.09484225 |
| **P17181-1** | IFNAR1 | -0.031767867 | 3.160301744 | -0.78015498 | 0.462873174 | 0.802800142 | -6.094994097 |
| **P27348** | YWHAQ | 0.038773787 | 3.156196627 | 2.005183378 | 0.08829972 | 0.437773138 | -6.095519119 |
| **Q9NVE5-3** | USP40 | 0.036012101 | 3.159559213 | 0.645875653 | 0.540575101 | 0.843540088 | -6.095694943 |
| **Q8WUY8** | NAT14 | 0.012551598 | 3.177008986 | 0.182505003 | 0.860770923 | 0.968855822 | -6.096848666 |
| **Q9Y3D0** | FAM96B | 0.005825637 | 3.193635983 | 0.093213788 | 0.928553313 | 0.98601391 | -6.096948708 |
| **Q9BUB5-1** | MKNK1 | -0.036894923 | 3.167728564 | -1.060509649 | 0.326881195 | 0.712111625 | -6.097026166 |
| **O00391** | QSOX1 | 0.066351347 | 3.178523013 | 0.569427174 | 0.588274694 | 0.867233795 | -6.097402033 |

| | | | | | | | |
|---|---|---|---|---|---|---|---|
| **Q9NWT6** | HIF1AN | -0.047710625 | 3.182705157 | -1.337870162 | 0.226036472 | 0.63121696 | -6.097458563 |
| **Q96DN6** | MBD6 | -0.02403078 | 3.200799494 | -0.409448356 | 0.695421115 | 0.911821519 | -6.097734669 |
| **A2RUC4** | TYW5 | -0.024768543 | 3.213371119 | -0.294164758 | 0.777841543 | 0.944053983 | -6.097775842 |
| **O14640-1** | DVL1 | -0.035074265 | 3.155504858 | -0.692221097 | 0.512860535 | 0.831227449 | -6.097966452 |
| **O15228** | GNPAT | 0.028228379 | 3.176391064 | 1.696513845 | 0.137098652 | 0.528248684 | -6.098129636 |
| **Q9BZQ8** | FAM129A | -0.040482455 | 3.167533177 | -1.001700274 | 0.352461847 | 0.732738778 | -6.098283507 |
| **Q9NVV9-1** | THAP1 | 0.024014647 | 3.207162071 | 0.380555665 | 0.715714282 | 0.924254721 | -6.098756296 |
| **Q99795** | GPA33 | 0.039626417 | 3.165025156 | 1.006159677 | 0.350468087 | 0.731407657 | -6.098966866 |
| **Q9UKB1-3** | FBXW11 | 0.032103344 | 3.178481646 | 1.064004637 | 0.325409194 | 0.710703247 | -6.099095096 |
| **Q86UV5** | USP48 | 0.031460778 | 3.182511602 | 0.91334781 | 0.3938147 | 0.760194378 | -6.099121612 |
| **Q11201** | ST3GAL1 | -0.121829456 | 3.112904634 | -0.746059711 | 0.48184563 | 0.81235548 | -6.100307426 |
| **P18859-2** | ATP5J | 0.013521445 | 3.247795912 | 0.120099577 | 0.908047083 | 0.98075504 | -6.100395537 |
| **P29323** | EPHB2 | -0.017261787 | 3.122169216 | -0.341463221 | 0.743572015 | 0.933895938 | -6.101588534 |
| **Q7Z2W9** | MRPL21 | 0.086224348 | 3.150464604 | 0.772718698 | 0.466966505 | 0.804608302 | -6.102205107 |
| **O95685** | PPP1R3D | -0.006813618 | 3.185314216 | -0.109452937 | 0.916159276 | 0.982975929 | -6.10234919 |
| **P59190-1** | RAB15 | 0.02871547 | 3.189816625 | 0.405320577 | 0.698304223 | 0.913150177 | -6.102878319 |
| **Q8TDR2** | STK35 | 0.036637038 | 3.191843082 | 1.044885463 | 0.33352755 | 0.719098069 | -6.103302696 |
| **Q6AI12** | ANKRD40 | -0.031128948 | 3.202358846 | -1.152511083 | 0.289900751 | 0.68467939 | -6.103516036 |
| **Q8N9R8-2** | SCAI | 0.050314895 | 3.183621375 | 1.678987009 | 0.140543387 | 0.536062029 | -6.103810889 |
| **Q8TD30-1** | GPT2 | 0.107669308 | 3.247158093 | 1.255046257 | 0.252882936 | 0.656801219 | -6.104158213 |
| **O00232-1** | PSMD12 | -0.036283067 | 3.17635408 | -1.787620703 | 0.120459797 | 0.499295152 | -6.104775196 |
| **Q9Y5K8** | ATP6V1D | -0.029781281 | 3.179978126 | -1.453962809 | 0.192674585 | 0.597035716 | -6.10497725 |
| **Q6YP21** | CCBL2; KYAT3 | 0.027806413 | 3.174214077 | 1.869056845 | 0.107258444 | 0.475805886 | -6.105338234 |
| **Q9Y467** | SALL2 | -0.020930526 | 3.139499044 | -0.250156078 | 0.810220696 | 0.955498404 | -6.105399108 |
| **Q9Y2Q0-2** | ATP8A1 | -0.032208745 | 3.175609366 | -0.358299085 | 0.731520148 | 0.92815165 | -6.106126976 |
| **P38432** | COIL | -0.032016189 | 3.165785141 | -1.540809873 | 0.170720933 | 0.57062592 | -6.106318483 |
| **A8MT69-1** | STRA13; CENPX | -0.001515556 | 3.172933308 | -0.020275309 | 0.984434596 | 0.995531556 | -6.106710016 |
| **Q9P289** | MST4; STK26 | -0.053088754 | 3.155124141 | -1.526503554 | 0.174170582 | 0.574454961 | -6.107502176 |
| **Q9H5I1** | SUV39H2 | -0.043196596 | 3.172175868 | -1.372248119 | 0.215654202 | 0.621237359 | -6.107511516 |
| **Q9UK23** | NAGPA | 0.026286233 | 3.160589477 | 0.641115572 | 0.543473879 | 0.844164217 | -6.108849007 |
| **Q8N5N7** | MRPL50 | 0.238887252 | 3.244599313 | 1.276820972 | 0.245567779 | 0.650779188 | -6.109050438 |
| **Q9BW66-3** | CINP | -0.070802576 | 3.189843435 | -1.119991263 | 0.302555682 | 0.691344519 | -6.109600779 |
| **Q9NXU5** | ARL 15.00 | -0.026331358 | 3.209806332 | -0.681575803 | 0.519144137 | 0.834217303 | -6.109954936 |
| **Q92628** | KIAA0232 | 0.031140256 | 3.197910359 | 0.569575797 | 0.588179639 | 0.867233795 | -6.1102371 |
| **P55795** | HNRNPH2 | 0.030505532 | 3.159182717 | 1.478118072 | 0.186321331 | 0.588831734 | -6.110307815 |
| **P62857** | RPS28 | -0.09912497 | 3.173388467 | -1.588336097 | 0.159712817 | 0.559241222 | -6.110374668 |
| **P49795** | RGS19 | 0.015690261 | 3.11772059 | 0.215934007 | 0.835686205 | 0.960912428 | -6.111536594 |
| **Q07820** | MCL1 | -0.023347886 | 3.206324784 | -0.343863642 | 0.741848791 | 0.932740726 | -6.111874468 |
| **Q8TD17** | ZNF398 | -0.067158412 | 3.24172148 | -0.840883632 | 0.430383247 | 0.782649325 | -6.111917749 |

| | | | | | | | |
|---|---|---|---|---|---|---|---|
| **Q9UP95-7** | SLC12A4 | 0.032156164 | 3.177688695 | 0.46214296 | 0.659111251 | 0.896852471 | -6.111949408 |
| **Q9BR77-1** | CCDC77 | 0.051074731 | 3.10570752 | 0.984417479 | 0.360273222 | 0.73661838 | -6.112024237 |
| **P62318** | SNRPD3 | 0.044803322 | 3.159202635 | 1.426916254 | 0.200022484 | 0.605177234 | -6.112611788 |
| **Q9H223** | EHD4 | -0.053649239 | 3.187113692 | -1.58435951 | 0.160607777 | 0.560635782 | -6.112630564 |
| **Q9BRR8** | GPATCH1 | -0.028026032 | 3.188712199 | -1.432620681 | 0.198451855 | 0.603708078 | -6.112910025 |
| **Q9NUV9** | GIMAP4 | -0.00819394 | 3.177594767 | -0.142006033 | 0.891394195 | 0.977905675 | -6.113412529 |
| **P98155-2** | VLDLR | 0.034300848 | 3.185128453 | 1.520995674 | 0.175515843 | 0.576466876 | -6.114342064 |
| **P29590** | PML | -0.04112968 | 3.207402494 | -0.655772685 | 0.534578965 | 0.84036998 | -6.114438836 |
| **Q13009** | TIAM1 | -0.029296443 | 3.199089809 | -1.027835719 | 0.340903634 | 0.722995142 | -6.114759525 |
| **O94923** | GLCE | -0.023067388 | 3.190258745 | -0.501418001 | 0.63267797 | 0.887490806 | -6.115183344 |
| **P15336-1** | ATF2 | -0.027948153 | 3.195061126 | -1.288466704 | 0.241731608 | 0.647961411 | -6.115308296 |
| **Q9UG22** | GIMAP2 | 0.054395214 | 3.212723476 | 0.965881315 | 0.368800688 | 0.743611616 | -6.115492107 |
| **Q9H920** | RNF121 | 0.006535059 | 3.228013482 | 0.068528133 | 0.947434119 | 0.989472196 | -6.115729666 |
| **Q8NAF0** | ZNF579 | -0.030491603 | 3.162760986 | -0.554761761 | 0.597697549 | 0.871651708 | -6.116749806 |
| **P50750-2** | CDK9 | -0.039432925 | 3.152242581 | -1.089457697 | 0.314850461 | 0.701837878 | -6.116969659 |
| **Q14999-2** | CUL7 | -0.035653335 | 3.207414118 | -0.895429669 | 0.402633851 | 0.766026328 | -6.117443348 |
| **Q14644** | RASA3 | -0.035328181 | 3.183424661 | -1.480842216 | 0.185617013 | 0.588713441 | -6.11897105 |
| **Q96AC1-3** | FERMT2 | -0.039980744 | 3.184789617 | -1.044137651 | 0.333848366 | 0.719204247 | -6.119615941 |
| **Q8WUA7** | TBC1D22A | 0.035982442 | 3.168627718 | 1.283472028 | 0.243370424 | 0.649360189 | -6.120312362 |
| **Q9BVG9** | PTDSS2 | -0.0265699 | 3.162951342 | -0.515608177 | 0.623266747 | 0.884417629 | -6.120709211 |
| **Q8WXB4** | ZNF606 | -0.037136532 | 3.237361566 | -0.404028666 | 0.699207696 | 0.913245694 | -6.121103817 |
| **O43422-1** | PRKRIR; THAP12 | 0.03211423 | 3.186874575 | 1.384665374 | 0.212010212 | 0.616778152 | -6.12115083 |
| **Q9BVL4** | SELO; SELENOO | 0.002752549 | 3.19994324 | 0.04751425 | 0.963536149 | 0.991283985 | -6.1214654 |
| **Q8WU17** | RNF139 | 0.032524385 | 3.149159995 | 0.519551511 | 0.620664892 | 0.883249609 | -6.121831156 |
| **O76003** | GLRX3 | -0.033787713 | 3.167926537 | -1.952554507 | 0.095199554 | 0.451948632 | -6.12213682 |
| **P62380** | TBPL1 | -0.042051832 | 3.170435231 | -1.029101738 | 0.340351488 | 0.722975002 | -6.123195237 |
| **P26045** | PTPN3 | 0.039529414 | 3.157641722 | 1.179725661 | 0.279653629 | 0.676725357 | -6.123248803 |
| **Q9H8M2-5** | BRD9 | -0.045083821 | 3.218057009 | -1.640895354 | 0.148316592 | 0.546231571 | -6.123326359 |
| **Q9BTX7** | TTPAL | -0.005696104 | 3.209947248 | -0.112833432 | 0.913582281 | 0.982219313 | -6.123650474 |
| **O60306** | AQR | -0.024379139 | 3.189636066 | -2.049127873 | 0.082922503 | 0.425319102 | -6.123777114 |
| **Q96EC8-1** | YIPF6 | -0.025125723 | 3.219060878 | -0.524812979 | 0.617202513 | 0.881353777 | -6.126063556 |
| **Q92754** | TFAP2C | 0.032583985 | 3.174724549 | 0.479395624 | 0.647431117 | 0.892717845 | -6.126166275 |
| **P15882** | CHN1 | -0.02935178 | 3.182767243 | -0.839493489 | 0.431108208 | 0.782692325 | -6.126167674 |
| **O43896** | KIF1C | -0.03328268 | 3.152366872 | -0.510462229 | 0.626670928 | 0.885387512 | -6.126884114 |
| **Q13469** | NFATC2 | -0.048357374 | 3.186465765 | -1.432644282 | 0.19844538 | 0.603708078 | -6.126913557 |
| **Q8WYN0** | ATG4A | 0.023821958 | 3.197210393 | 0.501653433 | 0.632521214 | 0.887490806 | -6.127049661 |
| **Q6PCT2-1** | FBXL19 | -0.046310978 | 3.174478913 | -1.062327866 | 0.326114737 | 0.710831711 | -6.12783286 |
| **Q9NZB8** | MOCS1 | 0.038224 | 3.193075343 | 0.61991661 | 0.556499918 | 0.850691425 | -6.128294223 |
| **Q7L5Y1-1** | ENOSF1 | 0.034365603 | 3.179187431 | 1.467992472 | 0.188960818 | 0.593033086 | -6.128574265 |

| | | | | | | | |
|---|---|---|---|---|---|---|---|
| **O15431** | SLC31A1 | 0.035407191 | 3.182936588 | 0.403531792 | 0.699555317 | 0.913245694 | -6.12868894 |
| **Q86SX3-1** | C14orf80; TEDC1 | -0.030281327 | 3.189516777 | -0.752191706 | 0.478394937 | 0.811328097 | -6.128884319 |
| **Q96AE4-2** | FUBP1 | 0.038048566 | 3.184283153 | 0.626061124 | 0.552704854 | 0.847927742 | -6.129246229 |
| **Q9H8H0** | NOL11 | -0.028005698 | 3.187127656 | -1.808101616 | 0.116997748 | 0.492866972 | -6.129492097 |
| **Q3T906** | GNPTAB | -0.039044474 | 3.169241426 | -1.197252103 | 0.273217665 | 0.672884265 | -6.12962432 |
| **Q9NVE7** | PANK4 | 0.02463368 | 3.191808377 | 1.764517528 | 0.12448449 | 0.507034795 | -6.130270628 |
| **Q01629** | IFITM2 | 0.043305946 | 3.179543143 | 0.377209779 | 0.718080979 | 0.924851578 | -6.130527669 |
| **Q6NXT4-2** | SLC30A6 | 0.000530526 | 3.189572017 | 0.008691457 | 0.993327117 | 0.99756865 | -6.130653862 |
| **Q9UHA3** | RSL24D1 | -0.049640838 | 3.220037674 | -0.993453419 | 0.356172458 | 0.734600528 | -6.130793045 |
| **Q8N960** | CEP120 | -0.043383222 | 3.194222441 | -1.248430648 | 0.255142538 | 0.659782078 | -6.130808958 |
| **Q5FBB7-1** | SGOL1; SGO1 | -0.006091908 | 3.120244308 | -0.090796014 | 0.930400548 | 0.986707862 | -6.130992999 |
| **Q96DU7** | ITPKC | 0.038250863 | 3.184346086 | 0.963042188 | 0.370120528 | 0.744424726 | -6.131283081 |
| **O75197** | LRP5 | 0.011441692 | 3.218160791 | 0.18813724 | 0.856531368 | 0.968855822 | -6.131763801 |
| **Q9Y5U2-1** | TSSC4 | 0.030381913 | 3.197694539 | 0.560131495 | 0.594237459 | 0.870591299 | -6.132042533 |
| **Q14690** | PDCD11 | -0.036079524 | 3.193344813 | -2.182588178 | 0.06852863 | 0.398579767 | -6.132266358 |
| **Q92625** | ANKS1A | -0.04394802 | 3.179028675 | -1.712330233 | 0.134059564 | 0.521853617 | -6.132569156 |
| **Q9BRR0-1** | ZKSCAN3 | 0.027488157 | 3.174255733 | 0.507933115 | 0.628347651 | 0.885908173 | -6.132780628 |
| **Q9UHK0** | NUFIP1 | -0.05932188 | 3.235038939 | -1.525998301 | 0.174293587 | 0.57459027 | -6.133024456 |
| **Q9NZL4-1** | HSPBP1 | 0.02229354 | 3.182173301 | 1.66625534 | 0.14309731 | 0.539313633 | -6.13375159 |
| **Q8TE99-1** | ACPL2; PXYLP1 | 0.027877705 | 3.154960503 | 0.719337026 | 0.497079367 | 0.823456124 | -6.133925226 |
| **Q96DT7** | ZBTB10 | -0.04184365 | 3.172930649 | -1.066706942 | 0.324274731 | 0.710703247 | -6.134228775 |
| **Q9GZR1** | SENP6 | -0.06074761 | 3.208646783 | -1.478949817 | 0.186106027 | 0.588831734 | -6.134270995 |
| **O43313** | ATMIN | 0.014417862 | 3.220091309 | 0.227372966 | 0.827148509 | 0.959077751 | -6.134517507 |
| **C9JLW8** | FAM195B; MCRIP1 | 0.042297343 | 3.172770026 | 1.179612787 | 0.279695491 | 0.676725357 | -6.134782558 |
| **Q9Y330** | ZBTB12 | -0.03300389 | 3.183388317 | -0.809714595 | 0.446849404 | 0.792204208 | -6.134882274 |
| **O15050** | TRANK1 | -0.021606154 | 3.046532767 | -0.34051416 | 0.744253772 | 0.934138424 | -6.135171235 |
| **Q02809-2** | PLOD1 | -0.028178944 | 3.231322511 | -1.390421053 | 0.210339958 | 0.615470161 | -6.135290253 |
| **Q6IBS0** | TWF2 | 0.053058777 | 3.175747067 | 1.685344716 | 0.139284389 | 0.533333364 | -6.135506435 |
| **Q9NVV0** | TMEM38B | 0.025766544 | 3.196836007 | 0.52401799 | 0.61772499 | 0.881750816 | -6.135932121 |
| **P98171-2** | ARHGAP4 | 0.027155754 | 3.178177307 | 1.806993854 | 0.1171825 | 0.493122332 | -6.136191485 |
| **O75161** | NPHP4 | 0.001464052 | 3.184040403 | 0.018985374 | 0.985424741 | 0.995531556 | -6.136311068 |
| **O75953-5** | DNAJB5 | 0.034477878 | 3.179409781 | 1.133287351 | 0.297327159 | 0.689506809 | -6.136539031 |
| **Q9BQA5** | HINFP | -0.026862445 | 3.159802877 | -0.598117291 | 0.570090769 | 0.855566898 | -6.136715765 |
| **P42167** | TMPO | 0.058338993 | 3.214945234 | 1.776237436 | 0.122426803 | 0.501640511 | -6.136814717 |
| **O95140** | MFN2 | 0.040390865 | 3.167446114 | 1.707632643 | 0.134955389 | 0.523800962 | -6.136878967 |
| **Q9Y2D8** | SSX2IP | -0.040443121 | 3.208008944 | -1.176599535 | 0.28081497 | 0.677979347 | -6.137406079 |
| **Q9Y6G5** | COMMD10 | -0.03540848 | 3.205857342 | -1.036588946 | 0.337100681 | 0.721128781 | -6.138042601 |
| **Q86TU7** | SETD3 | -0.032220521 | 3.20492115 | -1.570138347 | 0.163846971 | 0.562848368 | -6.138045517 |
| **Q9Y573-1** | IPP | -0.037904612 | 3.179604745 | -0.600498697 | 0.568596511 | 0.854816036 | -6.138324182 |

| | | | | | | | |
|---|---|---|---|---|---|---|---|
| **Q8TBF5-2** | PIGX | 0.028113087 | 3.20813764 | 0.573121025 | 0.585914842 | 0.8653224 | -6.138668057 |
| **Q8IVG5** | SAMD9L | 0.031279812 | 3.168379962 | 1.601283067 | 0.156831353 | 0.555269652 | -6.139034048 |
| **Q9H0X4** | ITFG3; FAM234A | 0.032389413 | 3.156514641 | 0.633286726 | 0.548262369 | 0.846224652 | -6.139121569 |
| **A2RTX5** | TARSL2 | -0.033543012 | 3.224405751 | -0.657905134 | 0.533292506 | 0.840132297 | -6.139288914 |
| **Q6DN90-1** | IQSEC1 | -0.03508198 | 3.198479435 | -0.965972387 | 0.368758412 | 0.743611616 | -6.139657141 |
| **Q5W111** | SPRYD7 | -0.013213851 | 3.190285093 | -0.19481821 | 0.851509097 | 0.966218452 | -6.140263962 |
| **P54296** | MYOM2 | 0.017842801 | 3.195021925 | 0.359987185 | 0.730316202 | 0.927974736 | -6.140724344 |
| **O43829** | ZBTB14; ZFP161 | 0.040353916 | 3.186830427 | 1.260823617 | 0.25092384 | 0.655139636 | -6.140758569 |
| **Q96MP5** | ZSWIM3 | -0.002011055 | 3.177771265 | -0.028527781 | 0.97810085 | 0.993585863 | -6.141641737 |
| **Q07912-1** | TNK2 | 0.014050923 | 3.171740007 | 0.297313598 | 0.775542233 | 0.94357883 | -6.141831177 |
| **O15182** | CETN3 | -0.118298794 | 3.188595077 | -1.617725518 | 0.153242445 | 0.552031202 | -6.142190162 |
| **P62244** | RPS15A | -0.040808705 | 3.189421707 | -1.878699799 | 0.1057923 | 0.475403904 | -6.142429007 |
| **Q7Z6J0-1** | SH3RF1 | 0.007649035 | 3.183196503 | 0.134446342 | 0.897134662 | 0.978757548 | -6.142789868 |
| **O15173-2** | PGRMC2 | -0.031221781 | 3.199669435 | -1.217434447 | 0.265962893 | 0.668906358 | -6.143660683 |
| **O95239-1** | KIF4A | -0.031506155 | 3.18659337 | -1.855535485 | 0.109347923 | 0.479873319 | -6.143719738 |
| **Q9NW38-2** | FANCL | 0.007094506 | 3.20156046 | 0.131400042 | 0.899449796 | 0.978757548 | -6.143859221 |
| **Q96QT6-1** | PHF12 | 0.035803016 | 3.184062266 | 1.450021552 | 0.193729798 | 0.598629202 | -6.144037796 |
| **Q9BWJ5** | SF3B5 | -0.129892964 | 3.20164087 | -1.516339379 | 0.176660611 | 0.577967785 | -6.14452779 |
| **P48723** | HSPA13 | 0.051379936 | 3.183122982 | 1.308501815 | 0.235254588 | 0.64164013 | -6.144641778 |
| **Q5T5U3** | ARHGAP21 | -0.006038157 | 3.099987029 | -0.080260938 | 0.938454884 | 0.987739899 | -6.146894393 |
| **Q9UHB4-2** | NDOR1 | -0.008478538 | 3.150436462 | -0.1014234 | 0.922284636 | 0.984164603 | -6.146979876 |
| **O75528** | TADA3 | -0.034850209 | 3.165834488 | -1.247015963 | 0.255627992 | 0.659782078 | -6.147036357 |
| **Q9H1B5** | XYLT2 | -0.034034972 | 3.179093471 | -0.622151461 | 0.555117766 | 0.849626402 | -6.147183133 |
| **O15550** | KDM6A | 0.052040294 | 3.166979159 | 1.127029412 | 0.2997786 | 0.690882899 | -6.147535738 |
| **Q9Y3B7-1** | MRPL11 | 0.173949482 | 3.155051336 | 1.602729532 | 0.156512483 | 0.554834464 | -6.148376357 |
| **P30519** | HMOX2 | -0.022086071 | 3.206803781 | -1.573490887 | 0.163077888 | 0.561580888 | -6.1484278 |
| **Q8NEJ9** | NGDN | -0.095618857 | 3.143431162 | -1.515883631 | 0.176773029 | 0.577967785 | -6.148575694 |
| **P50238** | CRIP1 | -0.026564451 | 3.232653911 | -0.564673018 | 0.591319971 | 0.868764508 | -6.149050488 |
| **Q8IWE4** | DCUN1D3 | -0.013822173 | 3.214489367 | -0.225607572 | 0.828464519 | 0.959077751 | -6.149208772 |
| **Q9NPE3** | NOP10 | 0.028970977 | 3.206500205 | 1.386782484 | 0.211394463 | 0.61590617 | -6.149299456 |
| **Q96ER3** | SAAL1 | -0.032381936 | 3.171022822 | -1.579139549 | 0.161789708 | 0.561414897 | -6.150047259 |
| **Q66K89** | E4F1 | -0.012579625 | 3.190078955 | -0.253644361 | 0.807638387 | 0.954797691 | -6.150119335 |
| **O75564** | JRK | -0.042212334 | 3.173446651 | -0.88502683 | 0.407821271 | 0.767987674 | -6.150624982 |
| **Q9BSY9** | DESI2 | -0.030524862 | 3.168528578 | -0.425241027 | 0.684441432 | 0.905978473 | -6.150717545 |
| **Q96IK5** | GMCL1 | -0.05211825 | 3.136477646 | -0.529094982 | 0.614392484 | 0.879839272 | -6.151014238 |
| **Q5H8A4-1** | PIGG | -0.054438801 | 3.20167412 | -0.765404766 | 0.471016812 | 0.807027916 | -6.151334466 |
| **Q9UBK9** | UXT | -0.018676649 | 3.230601701 | -0.336268125 | 0.747306963 | 0.934974996 | -6.152114631 |
| **Q8TCT8** | SPPL2A | -0.023720798 | 3.17728031 | -0.378210657 | 0.717372658 | 0.924679646 | -6.152155091 |
| **Q12841** | FSTL1 | -0.068485125 | 3.151640566 | -1.011306697 | 0.348177976 | 0.730105045 | -6.152165865 |

| | | | | | | | |
|---|---|---|---|---|---|---|---|
| **Q9UHW5-2** | GPN3 | -0.02986981 | 3.170506838 | -1.127142504 | 0.29973415 | 0.690882899 | -6.152642824 |
| **Q8N720-3** | ZNF655 | -0.087848284 | 3.241544309 | -1.044670257 | 0.333619849 | 0.719101926 | -6.15271945 |
| **Q9NX70** | MED29 | -0.072412856 | 3.20152033 | -1.454818832 | 0.192446092 | 0.596793208 | -6.15282242 |
| **P67809** | YBX1 | -0.035157566 | 3.175470788 | -1.929636071 | 0.09836925 | 0.457310529 | -6.15289387 |
| **Q92968** | PEX13 | -0.000609868 | 3.192275649 | -0.007203535 | 0.994469446 | 0.997608555 | -6.153733581 |
| **Q3B820-3** | FAM161A | -0.091583962 | 3.108465542 | -0.556303557 | 0.596702896 | 0.871378088 | -6.153750508 |
| **Q14192** | FHL2 | -0.044195228 | 3.236021738 | -0.973010219 | 0.365502709 | 0.740417905 | -6.154296425 |
| **P78537** | BLOC1S1 | 0.058105214 | 3.206322339 | 0.890209144 | 0.405230912 | 0.766336431 | -6.154824085 |
| **Q9P0K8** | FOXJ2 | 0.009569083 | 3.198047589 | 0.165327867 | 0.873730978 | 0.97237605 | -6.15599687 |
| **Q8N5C8** | TAB3 | -0.031568572 | 3.198044246 | -0.822379287 | 0.440105471 | 0.788699726 | -6.156262606 |
| **Q8NDX5-7** | PHC3 | 0.027578877 | 3.177918412 | 1.136158655 | 0.296207964 | 0.688880164 | -6.156789899 |
| **Q8IYB5-2** | SMAP1 | -0.076974332 | 3.219315746 | -0.979940331 | 0.362318704 | 0.738054037 | -6.156819773 |
| **Q9UNA4** | POLI | 0.01897174 | 3.178145556 | 0.253919724 | 0.807434652 | 0.954765339 | -6.157786803 |
| **O14523** | C2CD2L | -0.002870518 | 3.181687874 | -0.039375017 | 0.969778329 | 0.992476747 | -6.157825858 |
| **P04637** | TP53 | -0.027223307 | 3.137689818 | -0.632766821 | 0.548581285 | 0.846404495 | -6.158225129 |
| **P55789** | GFER | -0.037283542 | 3.201733434 | -0.557068798 | 0.596209568 | 0.871378088 | -6.158560238 |
| **Q6P2C8-1** | MED27 | -0.032064646 | 3.210439161 | -1.340531392 | 0.225217178 | 0.630229362 | -6.159063312 |
| **P49589-3** | CARS | 0.026047895 | 3.189471753 | 1.970376782 | 0.092804905 | 0.44685719 | -6.159365899 |
| **Q9HAJ7** | SAP30L | 0.017364986 | 3.161911369 | 0.401087403 | 0.701266591 | 0.914420329 | -6.159883408 |
| **O15151** | MDM4 | 0.015040159 | 3.195675443 | 0.254114791 | 0.807290336 | 0.954765339 | -6.160068053 |
| **O15400** | STX7 | -0.033144557 | 3.184856964 | -1.040233448 | 0.335527314 | 0.720533987 | -6.160232401 |
| **Q5T4F4-2** | ZFYVE27 | -0.002094844 | 3.192555962 | -0.03806315 | 0.970784661 | 0.992521442 | -6.160383977 |
| **Q96PP9** | GBP4 | 0.045014976 | 3.095030032 | 1.638228356 | 0.148875861 | 0.546231571 | -6.160437029 |
| **Q5D1E8** | ZC3H12A | 0.024151678 | 3.217174271 | 0.430319476 | 0.680928118 | 0.903980599 | -6.160767491 |
| **Q9Y4A0** | JRKL | -0.029897625 | 3.187246239 | -0.634924211 | 0.547258661 | 0.846115385 | -6.160944673 |
| **Q9BQD3** | KXD1 | -0.035274523 | 3.23942365 | -0.51709643 | 0.62228409 | 0.883385606 | -6.161204021 |
| **Q6Y7W6-1** | GIGYF2 | -0.016491317 | 3.200559218 | -0.29896301 | 0.774338794 | 0.943158294 | -6.161215462 |
| **Q6P3S6** | FBXO42 | 0.035766368 | 3.222515106 | 0.915156408 | 0.39293266 | 0.760129045 | -6.161540149 |
| **Q86VQ6** | TXNRD3 | -0.040979386 | 3.183756401 | -1.018197708 | 0.345130452 | 0.726061285 | -6.161648946 |
| **Q13163-1** | MAP2K5 | 0.031500234 | 3.166489801 | 0.877987011 | 0.411359743 | 0.769906516 | -6.1621266 |
| **Q969F1** | GTF3C6 | -0.033178381 | 3.193031101 | -1.151323351 | 0.290355053 | 0.684803131 | -6.16219201 |
| **Q9NWA0** | MED9 | -0.165740582 | 3.158720647 | -0.924316781 | 0.388488113 | 0.75788842 | -6.162809278 |
| **Q9NPB8** | GPCPD1 | 0.029880298 | 3.230550462 | 1.145834994 | 0.292462066 | 0.68530312 | -6.163081568 |
| **Q9Y3B1** | SLMO2; PRELID3B | -0.03463542 | 3.260769932 | -0.607588035 | 0.56416201 | 0.852578515 | -6.163296479 |
| **O15260-1** | SURF4 | 0.024318416 | 3.148197473 | 0.330032146 | 0.75180002 | 0.935635612 | -6.163566844 |
| **P07992-3** | ERCC1 | -0.033805749 | 3.205469865 | -1.030219289 | 0.339864685 | 0.722564871 | -6.165162215 |
| **Q96RY7** | IFT140 | -0.031683224 | 3.173086931 | -1.153193424 | 0.289640028 | 0.68447056 | -6.165465951 |
| **P53805** | RCAN1 | 0.02437413 | 3.215632622 | 0.268448631 | 0.79670851 | 0.950757673 | -6.165672813 |
| **Q9BXL7** | CARD11 | 0.030003919 | 3.193729093 | 1.279425772 | 0.24470516 | 0.650150213 | -6.166592379 |

| | | | | | | | |
|---|---|---|---|---|---|---|---|
| **Q13112** | CHAF1B | -0.051109333 | 3.184290552 | -1.782381669 | 0.121361254 | 0.499487166 | -6.16695777 |
| **Q92783** | STAM | 0.060018149 | 3.169502241 | 1.495235622 | 0.181936098 | 0.584742029 | -6.168473172 |
| **Q8NI36** | WDR36 | -0.039447889 | 3.199284113 | -1.888622999 | 0.10430412 | 0.472077308 | -6.169421916 |
| **Q12766-1** | HMGXB3 | -0.027415081 | 3.193630674 | -0.820136342 | 0.441294519 | 0.788971713 | -6.169852826 |
| **Q53GT1** | KLHL22 | -0.052471527 | 3.19567764 | -0.670155647 | 0.525939926 | 0.838233243 | -6.171062224 |
| **O94972** | TRIM37 | -0.018521439 | 3.163483714 | -0.336553233 | 0.747101795 | 0.934974996 | -6.171155293 |
| **O43396** | TXNL1 | -0.056361008 | 3.164781553 | -1.788776834 | 0.120261744 | 0.499295152 | -6.171267238 |
| **Q5VWZ2** | LYPLAL1 | 0.046331284 | 3.198439608 | 1.426908611 | 0.200024596 | 0.605177234 | -6.171699242 |
| **Q15126** | PMVK | -0.053295345 | 3.179496771 | -1.449895002 | 0.193763767 | 0.598629202 | -6.171987197 |
| **P82979** | SARNP | -0.036533017 | 3.157336403 | -1.639436389 | 0.14862229 | 0.546231571 | -6.173238298 |
| **Q9UBB6-3** | NCDN | -0.092622161 | 3.187208186 | -1.621156597 | 0.152503375 | 0.552031202 | -6.17333157 |
| **O94759-1** | TRPM2 | 0.006600887 | 3.225791252 | 0.129867404 | 0.900614979 | 0.97938489 | -6.173554407 |
| **Q9UHR6** | ZNHIT2 | -0.036258993 | 3.214910484 | -0.936581282 | 0.382597292 | 0.754089677 | -6.173829651 |
| **Q8IWB7** | WDFY1 | -0.035241458 | 3.214456533 | -1.388626997 | 0.210859307 | 0.61590617 | -6.173845642 |
| **Q96EY9** | ADAT3 | -0.001235354 | 3.183507981 | -0.018065943 | 0.986130506 | 0.995531556 | -6.173892279 |
| **Q8NFI3-1** | ENGASE | -0.004475342 | 3.198165359 | -0.068953838 | 0.947108169 | 0.989472196 | -6.174061777 |
| **Q5MNZ6** | WDR45B; WDR45L | -0.036673222 | 3.210560008 | -1.163295236 | 0.285803057 | 0.681580409 | -6.174991663 |
| **Q9Y2K5-1** | R3HDM2 | -0.021322321 | 3.197154012 | -0.492016877 | 0.638954224 | 0.88909352 | -6.175081228 |
| **Q9H9Y4** | GPN2 | 0.031367643 | 3.185569099 | 0.689648654 | 0.51437444 | 0.831675214 | -6.175760579 |
| **P13798** | APEH | -0.061849097 | 3.176678729 | -1.696087549 | 0.13718147 | 0.528311574 | -6.175839421 |
| **Q96FJ2** | DYNLL2 | 0.035616364 | 3.173820041 | 1.420832462 | 0.201709984 | 0.608191963 | -6.176843228 |
| **Q9HAT8** | PELI2 | -0.162540629 | 3.205088056 | -1.096787382 | 0.311862272 | 0.699730515 | -6.176961824 |
| **P00488** | F13A1 | -0.019099219 | 3.201148732 | -0.150444294 | 0.884994974 | 0.975708125 | -6.177202235 |
| **Q9UER7-1** | DAXX | 0.03437757 | 3.180896776 | 1.176547081 | 0.280834491 | 0.677979347 | -6.177920974 |
| **Q9P2B4** | CTTNBP2NL | -0.038397973 | 3.181677752 | -1.251725174 | 0.254015098 | 0.659039362 | -6.178754211 |
| **Q9NX07-1** | TRNAU1AP | -0.071274127 | 3.207255905 | -1.107020849 | 0.30772921 | 0.696395215 | -6.178989416 |
| **O14732-1** | IMPA2 | 0.025696033 | 3.177596956 | 0.385177656 | 0.712450537 | 0.921892429 | -6.179263711 |
| **Q96SY0** | VWA9; INTS14 | -0.02897438 | 3.154899895 | -1.294545173 | 0.239750215 | 0.645958461 | -6.179714997 |
| **Q9NWQ8** | PAG1 | -0.097783679 | 3.156552208 | -1.377180214 | 0.214200154 | 0.619843797 | -6.180240167 |
| **Q6ULP2** | AFTPH | -0.030539802 | 3.212049807 | -1.491099828 | 0.182986833 | 0.585513649 | -6.180363395 |
| **O00571** | DDX3X | -0.052764765 | 3.196994928 | -1.641460825 | 0.148198269 | 0.546231571 | -6.180677318 |
| **A0A5B9** | TRBC2 | 0.068054007 | 3.239331298 | 1.437133463 | 0.197217272 | 0.603708078 | -6.180857234 |
| **Q9UQ49-2** | NEU3 | -0.052019131 | 3.141601876 | -0.632489637 | 0.54875136 | 0.846404495 | -6.18094772 |
| **Q9Y3L5** | RAP2C | 0.021507306 | 3.157071651 | 0.31858756 | 0.760073132 | 0.938422437 | -6.181376315 |
| **Q9NVC6** | MED17 | -0.024788873 | 3.187557628 | -1.403081155 | 0.206707677 | 0.612110509 | -6.181521844 |
| **Q96QT4** | TRPM7 | 0.030392904 | 3.170762349 | 1.055388851 | 0.329047659 | 0.714675683 | -6.18236232 |
| **Q9BVV6-3** | KIAA0586 | -0.026118194 | 3.167105678 | -0.561794823 | 0.593167981 | 0.869935985 | -6.183108779 |
| **Q15637-5** | SF1 | -0.0286619 | 3.162727128 | -0.829665821 | 0.436258485 | 0.785483221 | -6.183473364 |
| **Q96AX1** | VPS33A | 0.036843117 | 3.205962504 | 1.598713399 | 0.157399339 | 0.555457754 | -6.183933306 |

| | | | | | | | |
|---|---|---|---|---|---|---|---|
| **Q9Y6I3** | EPN1 | -0.029924752 | 3.17643116 | -1.160346536 | 0.286918626 | 0.682301253 | -6.184048166 |
| **Q6XZF7** | DNMBP | -0.040577752 | 3.204949284 | -0.910052656 | 0.395425555 | 0.761244496 | -6.184104214 |
| **Q9NWQ9** | C14orf119 | -0.009605467 | 3.227940105 | -0.154459504 | 0.881953256 | 0.97451158 | -6.184160795 |
| **Q9Y222-1** | DMTF1 | -0.025294627 | 3.176211672 | -0.556208694 | 0.596764067 | 0.871378088 | -6.184341059 |
| **Q9Y6M4-4** | CSNK1G3 | -0.006823805 | 3.185784054 | -0.13600139 | 0.895953271 | 0.978757548 | -6.185762812 |
| **Q9BWU0** | SLC4A1AP | 0.03949038 | 3.181245008 | 1.647271847 | 0.146987505 | 0.545853843 | -6.185815493 |
| **Q14331** | FRG1 | 0.022976973 | 3.178834938 | 1.75146517 | 0.126815495 | 0.509635359 | -6.186031342 |
| **P63167** | DYNLL1 | 0.044131094 | 3.188376477 | 1.473433106 | 0.187538352 | 0.590797862 | -6.186098138 |
| **Q67FW5** | B3GNTL1 | -0.024137302 | 3.188326473 | -0.573340333 | 0.585774907 | 0.8653224 | -6.187126821 |
| **Q9UJU2-6** | LEF1 | 0.02335546 | 3.263694088 | 0.285632658 | 0.784083894 | 0.946296623 | -6.187200283 |
| **Q9HCM7** | FBRSL1 | -0.031496943 | 3.220611789 | -1.061751739 | 0.326357442 | 0.711165628 | -6.187422715 |
| **O75061-2** | DNAJC6 | -0.026031597 | 3.226950736 | -0.71522576 | 0.499451226 | 0.825004623 | -6.187510249 |
| **O60941-1** | DTNB | -0.006001119 | 3.185168451 | -0.114650428 | 0.912197629 | 0.981900848 | -6.187817352 |
| **O14519** | CDK2AP1 | 0.034071749 | 3.210527224 | 0.687842625 | 0.515439031 | 0.832688716 | -6.187986572 |
| **P49116-1** | NR2C2 | 0.049925765 | 3.182145775 | 1.572770395 | 0.163242886 | 0.561580888 | -6.18801588 |
| **P24278** | ZBTB25 | -0.007713818 | 3.19204471 | -0.129311327 | 0.901037801 | 0.97952714 | -6.188477128 |
| **P48378** | RFX2 | -0.002388067 | 3.205589521 | -0.033510881 | 0.974277139 | 0.99316605 | -6.188608416 |
| **Q01082-3** | SPTBN1 | 0.047547118 | 3.173642347 | 0.960184385 | 0.371452735 | 0.744717822 | -6.188679877 |
| **Q99996-2** | AKAP9 | -0.031001451 | 3.18782312 | -2.181003589 | 0.068683744 | 0.398897914 | -6.188728205 |
| **O43189** | PHF1 | -0.020446721 | 3.193055787 | -0.376835386 | 0.718346015 | 0.924851578 | -6.188913992 |
| **Q9NW13-1** | RBM28 | -0.039931884 | 3.152997863 | -1.733564162 | 0.130081029 | 0.515715456 | -6.188956556 |
| **O76094** | SRP72 | -0.03715361 | 3.188526134 | -1.937841467 | 0.097222576 | 0.455444201 | -6.189175574 |
| **Q15738** | NSDHL | -0.028537407 | 3.17490446 | -1.518126919 | 0.176220323 | 0.577825201 | -6.189199702 |
| **Q8NHP6-1** | MOSPD2 | 0.028106071 | 3.208091053 | 1.093479375 | 0.313208002 | 0.701109468 | -6.189503563 |
| **O95199-1** | RCBTB2 | -0.048699968 | 3.209481261 | -0.85843991 | 0.421303515 | 0.777315429 | -6.189586806 |
| **Q96I76** | GPATCH3 | 0.000981348 | 3.216369318 | 0.020700089 | 0.984108545 | 0.995510872 | -6.189673183 |
| **O15379-2** | HDAC3 | 0.036500177 | 3.186230339 | 1.464601637 | 0.189852351 | 0.593849183 | -6.189859994 |
| **Q8N3J9** | ZNF664 | 0.007991667 | 3.179288136 | 0.095851877 | 0.926538301 | 0.985528933 | -6.190629747 |
| **Q13615** | MTMR3 | -0.052828225 | 3.185967241 | -1.372672032 | 0.21552888 | 0.621101543 | -6.190831257 |
| **P29692-2** | EEF1D | 0.034116347 | 3.194579406 | 1.041386084 | 0.335030937 | 0.720384517 | -6.191275946 |
| **P01111** | NRAS | -0.105458593 | 3.140761676 | -0.975073988 | 0.364552253 | 0.739812938 | -6.191354362 |
| **Q7L2K0-1** | C16orf59; TEDC2 | -0.024152796 | 3.194265731 | -0.493006826 | 0.638291794 | 0.889065271 | -6.191536857 |
| **P49247** | LOC101060545; RF | -0.030962936 | 3.166349917 | -1.463170677 | 0.190229738 | 0.594291452 | -6.191582841 |
| **Q6NW29** | RWDD4 | -0.046763097 | 3.168420798 | -0.862043064 | 0.419457442 | 0.775923022 | -6.192269657 |
| **P09543-1** | CNP | -0.025471787 | 3.176657474 | -1.837767919 | 0.112154315 | 0.484388026 | -6.192561142 |
| **Q12980** | NPRL3 | 0.038557226 | 3.21374881 | 0.80157379 | 0.451222867 | 0.79513757 | -6.193351268 |
| **Q00535** | CDK5 | -0.029160805 | 3.175010024 | -1.49903414 | 0.18097595 | 0.58354461 | -6.193465283 |
| **O15504** | NUPL2 | -0.033146944 | 3.193430479 | -1.196386002 | 0.273532731 | 0.673239636 | -6.194643378 |
| **Q9BXI6-2** | TBC1D10A | -0.027707618 | 3.183038822 | -1.22445888 | 0.263476819 | 0.666875862 | -6.195036236 |

| | | | | | | | |
|---|---|---|---|---|---|---|---|
| **O43760-2** | SYNGR2 | -0.005538963 | 3.133828175 | -0.055460094 | 0.957444968 | 0.989500907 | -6.195491334 |
| **Q8WY22** | BRI3BP | -0.052783272 | 3.154229373 | -1.010429751 | 0.348567326 | 0.730176717 | -6.195542749 |
| **Q9NQB0-11** | TCF7L2 | 0.041418987 | 3.144627962 | 0.646267215 | 0.540337078 | 0.843540088 | -6.195702762 |
| **Q5JPI9** | METTL10; EEF1AKI | 0.024928151 | 3.167990903 | 0.425187119 | 0.684478772 | 0.905978473 | -6.195783576 |
| **Q7L2E3-1** | DHX30 | 0.052587486 | 3.209636485 | 0.320614479 | 0.75860537 | 0.9384213 | -6.196009212 |
| **Q13432** | UNC119 | -0.025770358 | 3.18570821 | -0.335710854 | 0.747708048 | 0.934974996 | -6.196042735 |
| **O14776-1** | TCERG1 | 0.027739717 | 3.173896486 | 1.949469048 | 0.095620311 | 0.452536799 | -6.196554377 |
| **Q9NY74** | ETAA1 | 0.038130247 | 3.192600195 | 0.662298445 | 0.530648272 | 0.839849063 | -6.197100991 |
| **Q9H777** | ELAC1 | -0.029689563 | 3.18432146 | -0.792799532 | 0.455970313 | 0.798084321 | -6.197267289 |
| **Q8NBW4** | SLC38A9 | -0.049326693 | 3.207791116 | -0.691835195 | 0.513087458 | 0.831256087 | -6.197777024 |
| **Q4VCS5** | AMOT | 0.033619821 | 3.146259957 | 0.792790771 | 0.45597507 | 0.798084321 | -6.197890082 |
| **Q96DR4** | STARD4 | 0.045093051 | 3.127005859 | 0.791508254 | 0.456671924 | 0.798472367 | -6.198036079 |
| **Q9BUR4** | WRAP53 | 0.036887024 | 3.195342858 | 1.189109531 | 0.276191937 | 0.674975373 | -6.198088362 |
| **O15126-1** | SCAMP1 | -0.049066002 | 3.176309487 | -0.979038542 | 0.362731799 | 0.738054037 | -6.198466395 |
| **Q9UGH3** | SLC23A2 | 0.029342624 | 3.233884625 | 0.447843615 | 0.668871325 | 0.90001396 | -6.200098636 |
| **Q5VWJ9** | SNX30 | 0.042297069 | 3.178189054 | 0.837252772 | 0.432278601 | 0.783333621 | -6.200211235 |
| **Q8N300** | CCDC23; SVBP | -0.043360753 | 3.094777008 | -0.627973214 | 0.551527114 | 0.847559559 | -6.200650526 |
| **Q03113** | GNA12 | 0.018013104 | 3.203986709 | 0.372923664 | 0.721117673 | 0.925869411 | -6.201330225 |
| **Q96NB1** | FOPNL | 0.026873525 | 3.179069274 | 0.710435882 | 0.502224014 | 0.826806835 | -6.201389773 |
| **O75427** | LRCH4 | -0.030194901 | 3.196890451 | -1.184350372 | 0.277943019 | 0.676093886 | -6.202069981 |
| **Q9H7B2** | RPF2 | -0.110914346 | 3.172107796 | -1.220872029 | 0.264743771 | 0.66823064 | -6.202445536 |
| **Q86X52** | CHSY1 | 0.027526558 | 3.194443076 | 0.61191334 | 0.561466675 | 0.852576894 | -6.202824711 |
| **Q9UMY4-1** | SNX12 | -0.067575071 | 3.177892731 | -1.294142241 | 0.239881117 | 0.645958461 | -6.204067935 |
| **Q14146** | URB2 | -0.027771597 | 3.199660988 | -1.690927765 | 0.138187685 | 0.53089998 | -6.204936421 |
| **Q96L73-1** | NSD1 | -0.020519082 | 3.193381263 | -1.602957853 | 0.156462206 | 0.554834464 | -6.205192014 |
| **P19022** | CDH2 | -0.021323442 | 3.224060253 | -0.448732512 | 0.668262541 | 0.90001396 | -6.205844561 |
| **Q9H6U8-3** | ALG9 | 0.038944109 | 3.212893351 | 0.876518263 | 0.412100844 | 0.770575992 | -6.206174342 |
| **P78381-4** | SLC35A2 | 0.017426208 | 3.18086196 | 0.312092745 | 0.764783482 | 0.940369694 | -6.206436522 |
| **Q9Y2H2-1** | INPP5F | -0.030002246 | 3.206407959 | -0.800388178 | 0.451862319 | 0.79513757 | -6.206507559 |
| **Q13643** | FHL3 | 0.031448613 | 3.18497974 | 0.98786322 | 0.358705112 | 0.735928834 | -6.206638942 |
| **O95445-2** | APOM | -0.014094416 | 3.148220039 | -0.138323683 | 0.894189532 | 0.97851842 | -6.206796169 |
| **Q86TN4-4** | TRPT1 | -0.027026994 | 3.216208832 | -0.693007391 | 0.512398372 | 0.831227449 | -6.206897581 |
| **Q92575** | UBXN4 | 0.032896049 | 3.162638731 | 1.276341258 | 0.245726933 | 0.650983821 | -6.20720126 |
| **P35625** | TIMP3 | -0.005320164 | 3.221311938 | -0.113009315 | 0.913448235 | 0.982219313 | -6.207580081 |
| **Q7Z2E3-1** | APTX | -0.035428699 | 3.19001647 | -1.504316142 | 0.179648587 | 0.580990067 | -6.207728571 |
| **Q8IXM6-1** | NRM | 0.041607825 | 3.199757646 | 0.66088573 | 0.531497648 | 0.839849063 | -6.208345306 |
| **Q9H1Y0-1** | ATG5 | -0.027961587 | 3.18664202 | -1.377332841 | 0.214155298 | 0.619843797 | -6.208372583 |
| **P51686** | CCR9 | -0.009309361 | 3.1931005 | -0.071114959 | 0.945453629 | 0.98904859 | -6.208795685 |
| **O60427** | FADS1 | -0.048260968 | 3.148734841 | -1.038837274 | 0.33612936 | 0.720858943 | -6.208872321 |

| | | | | | | | |
|---|---|---|---|---|---|---|---|
| **Q12849** | GRSF1 | 0.073959652 | 3.203099445 | 1.79824315 | 0.118651952 | 0.496152504 | -6.20972114 |
| **P30260-2** | CDC27 | -0.050313221 | 3.17826845 | -1.735702082 | 0.129686806 | 0.515290389 | -6.209830051 |
| **P54725-1** | RAD23A | -0.031130813 | 3.17574178 | -1.199528082 | 0.272391192 | 0.672884265 | -6.210189847 |
| **Q8WZ60** | KLHL6 | -0.022089882 | 3.116388416 | -0.44907598 | 0.668027381 | 0.90001396 | -6.211468679 |
| **Q96A73-1** | KIAA1191 | 0.030746373 | 3.205510126 | 0.603520738 | 0.566703635 | 0.853395233 | -6.211600541 |
| **O60704** | TPST2 | -0.013939819 | 3.146512853 | -0.195857337 | 0.850728624 | 0.965576989 | -6.211630983 |
| **Q8NEZ5-1** | FBXO22 | 0.033149929 | 3.18332915 | 1.147976569 | 0.291638387 | 0.685264687 | -6.21178181 |
| **Q7Z2Z1** | TICRR | -0.03279467 | 3.175854431 | -0.456959866 | 0.662640776 | 0.897184895 | -6.212425189 |
| **P98198-3** | ATP8B2 | -0.001838913 | 3.168279023 | -0.038445063 | 0.970491689 | 0.992521442 | -6.212793003 |
| **Q9Y4P1-2** | ATG4B | 0.065889551 | 3.195510443 | 1.18944221 | 0.276069882 | 0.67488468 | -6.212928762 |
| **Q92828** | CORO2A | -0.029951261 | 3.190943381 | -0.812986668 | 0.44510004 | 0.791127304 | -6.213624055 |
| **Q03426** | MVK | 0.054876162 | 3.154594954 | 1.168756884 | 0.283746406 | 0.679787821 | -6.213830596 |
| **O43716** | GATC | -0.024600056 | 3.17537271 | -0.21764159 | 0.83441015 | 0.960780962 | -6.214335446 |
| **Q9BY27** | DGCR6L | 0.019088316 | 3.188898666 | 0.305054277 | 0.769900384 | 0.941649022 | -6.214626229 |
| **Q9NPR9** | GPR108 | 0.008374803 | 3.193118428 | 0.183536021 | 0.859994466 | 0.968855822 | -6.215266046 |
| **Q8WVL7** | ANKRD49 | 0.025909976 | 3.182428395 | 0.485738382 | 0.643163854 | 0.891474831 | -6.216829596 |
| **Q96B54** | ZNF428 | -0.06098652 | 3.149384256 | -0.834960681 | 0.433478196 | 0.783962934 | -6.216830978 |
| **Q56A73** | SPIN4 | -0.036481196 | 3.119329887 | -0.839743476 | 0.430977775 | 0.782692325 | -6.217914285 |
| **O14713-1** | ITGB1BP1 | -0.028302697 | 3.222312275 | -0.499461537 | 0.63398143 | 0.887509451 | -6.218405334 |
| **O15021** | MAST4 | -0.012671332 | 3.144581431 | -0.262423517 | 0.801150972 | 0.952917733 | -6.218884703 |
| **Q9H672-1** | ASB7 | -0.010035332 | 3.177837841 | -0.134824637 | 0.89684724 | 0.978757548 | -6.21916231 |
| **O95785-3** | WIZ | 0.028773376 | 3.16087205 | 0.689991755 | 0.514172355 | 0.831657036 | -6.21961796 |
| **Q9NRW3** | APOBEC3C | -0.031980025 | 3.217598701 | -0.705293921 | 0.505211883 | 0.828977366 | -6.21995486 |
| **P49427** | CDC34 | -0.058621549 | 3.239351387 | -1.072727625 | 0.321758695 | 0.709476351 | -6.220019333 |
| **Q9H2K0** | MTIF3 | 0.143970347 | 3.165803534 | 0.982775358 | 0.36102241 | 0.736789892 | -6.221406799 |
| **Q9BXJ9** | NAA15 | -0.043953654 | 3.184543163 | -2.030754401 | 0.085129674 | 0.4310741 | -6.221588558 |
| **Q8NG08** | HELB | -0.035180922 | 3.205773746 | -0.696638698 | 0.510267496 | 0.829456831 | -6.221772766 |
| **Q15796-1** | SMAD2 | -0.047531361 | 3.19852737 | -0.745768667 | 0.482009829 | 0.81235548 | -6.22191963 |
| **Q32M88** | ATHL1; PGGHG | 0.034759989 | 3.176850794 | 0.916961665 | 0.392053738 | 0.759806823 | -6.222112529 |
| **Q8NF99-1** | ZNF397 | 0.012258623 | 3.160393717 | 0.216695943 | 0.835116753 | 0.960912428 | -6.222322659 |
| **Q86T03-2** | TMEM55B; PIP4P1 | -0.035564723 | 3.173614635 | -0.636469049 | 0.546312781 | 0.845412981 | -6.222406066 |
| **Q32MZ4** | LRRFIP1 | -0.076902822 | 3.169865764 | -1.848722503 | 0.110415813 | 0.481625912 | -6.222741027 |
| **P28161** | GSTM2 | 0.032114301 | 3.167642379 | 1.421820677 | 0.201435002 | 0.607824189 | -6.223353295 |
| **P10275** | AR | 0.012740189 | 3.173299264 | 0.289317083 | 0.781386096 | 0.944918194 | -6.223939951 |
| **Q9NWW6-1** | NMRK1 | 0.009092253 | 3.151971788 | 0.156506263 | 0.880403568 | 0.974002142 | -6.224237677 |
| **Q6NSJ2-1** | PHLDB3 | -0.025694323 | 3.174100733 | -0.586467366 | 0.577434233 | 0.861313941 | -6.224827077 |
| **Q96AB6** | NTAN1 | 0.026180665 | 3.173181357 | 0.750829968 | 0.47915978 | 0.811537935 | -6.224945583 |
| **O75446** | SAP30 | -0.030712736 | 3.180031545 | -1.106136476 | 0.308084601 | 0.696616637 | -6.225806707 |
| **O00400** | SLC33A1 | -0.010727934 | 3.186140102 | -0.235355811 | 0.821205328 | 0.95877683 | -6.228189699 |

| | | | | | | | |
|---|---|---|---|---|---|---|---|
| **Q8N3A8-1** | PARP8 | -0.011535944 | 3.170696578 | -0.214998568 | 0.836385475 | 0.960912428 | -6.228380084 |
| **Q8N129** | CNPY4 | -0.035632769 | 3.191677078 | -0.459925767 | 0.66061994 | 0.896934512 | -6.228867679 |
| **Q6IQ19-1** | CCSAP | -0.073218497 | 3.12590052 | -0.391557016 | 0.707956627 | 0.91822696 | -6.228904216 |
| **P49770** | EIF2B2 | -0.037244834 | 3.195936618 | -1.559205505 | 0.166378626 | 0.565630375 | -6.228983143 |
| **P78560** | CRADD | -0.028760944 | 3.205136521 | -0.677413129 | 0.521614665 | 0.834856672 | -6.229067572 |
| **Q9H944** | MED20 | -0.157681508 | 3.221976446 | -1.127766498 | 0.299488991 | 0.690882899 | -6.231055463 |
| **P06729** | CD2 | -0.065082168 | 3.160414384 | -1.279853778 | 0.244563672 | 0.650150213 | -6.231225653 |
| **P23025** | XPA | 0.043577818 | 3.207268314 | 1.174783124 | 0.281491627 | 0.678740812 | -6.231242744 |
| **Q9HCS7** | XAB2 | -0.034789663 | 3.174675528 | -1.725896127 | 0.131504491 | 0.519544096 | -6.231327206 |
| **P28908-1** | TNFRSF8 | -0.043207556 | 3.222413404 | -0.461564243 | 0.659504873 | 0.896852471 | -6.231447597 |
| **Q96SU4** | OSBPL9 | 0.041206692 | 3.181951802 | 1.568789259 | 0.164157413 | 0.563603735 | -6.231486867 |
| **Q8NBZ0** | INO80E | 0.055129238 | 3.201354564 | 0.784964737 | 0.460238928 | 0.801007291 | -6.231806551 |
| **P57768** | SNX16 | 0.021933981 | 3.180679128 | 0.350194485 | 0.73731174 | 0.929681266 | -6.232013627 |
| **Q9UMX5** | NENF | -0.253860546 | 3.188460693 | -1.401492072 | 0.207160474 | 0.612994402 | -6.232053543 |
| **O43567** | RNF13 | -0.046870158 | 3.16567622 | -0.958495814 | 0.372241626 | 0.745139764 | -6.232173387 |
| **Q8NBI5-2** | SLC43A3 | -0.021868372 | 3.211143066 | -0.327082358 | 0.75392903 | 0.936379818 | -6.232529064 |
| **Q15528-1** | MED22 | -0.043555189 | 3.225938944 | -0.868450743 | 0.416189121 | 0.772686182 | -6.232979237 |
| **P48730-2** | CSNK1D | -0.029484883 | 3.227173166 | -0.819179205 | 0.441802621 | 0.789146094 | -6.233125628 |
| **Q96CX2** | KCTD12 | -0.028890553 | 3.182800823 | -1.071816918 | 0.322138258 | 0.7096696 | -6.233379223 |
| **Q96PY6-3** | NEK1 | -0.04602102 | 3.18248725 | -1.492660887 | 0.182589577 | 0.585267652 | -6.233560418 |
| **Q6FI13** | HIST2H2AA4; HIST | 0.028307278 | 3.159256257 | 0.851500532 | 0.4248756 | 0.778986255 | -6.233649391 |
| **Q9Y4B6** | VPRBP; DCAF1 | -0.033454692 | 3.180531401 | -1.839907321 | 0.111812699 | 0.483851792 | -6.233903341 |
| **Q9ULJ1-4** | ODF2L | -0.030142595 | 3.155522619 | -0.764952636 | 0.471267983 | 0.807118802 | -6.234027038 |
| **Q9P086** | MED11 | 0.011033299 | 3.195575859 | 0.218244322 | 0.833959866 | 0.96068017 | -6.234538451 |
| **Q96N64** | PWWP2A | -0.025573618 | 3.229827366 | -0.765868346 | 0.470759375 | 0.806943524 | -6.234727864 |
| **P60891** | PRPS1 | 0.032413775 | 3.168530655 | 1.586986151 | 0.160016106 | 0.55963744 | -6.23493107 |
| **Q15543** | TAF13 | -0.026681917 | 3.178373787 | -0.629565795 | 0.550547346 | 0.84701277 | -6.235199699 |
| **Q96L34-1** | MARK4 | -0.025913313 | 3.163412276 | -0.45171133 | 0.666224405 | 0.899889986 | -6.236292632 |
| **Q8WW01** | TSEN15 | -0.029398541 | 3.212472623 | -0.73478776 | 0.488232619 | 0.817149391 | -6.237327841 |
| **Q9Y365** | STARD10 | -0.025002087 | 3.190307493 | -0.335716622 | 0.747703896 | 0.934974996 | -6.23784147 |
| **Q96AJ1-1** | CLUAP1 | 0.026353583 | 3.149539316 | 0.733571979 | 0.4889249 | 0.817619097 | -6.237949489 |
| **Q49A26-2** | GLYR1 | -0.041824073 | 3.155988871 | -0.938497919 | 0.381682886 | 0.752847698 | -6.238288877 |
| **Q99592-2** | ZBTB18; ZNF238 | -0.0028149 | 3.192457882 | -0.04977996 | 0.961798995 | 0.991283985 | -6.23856156 |
| **Q9BVK6** | TMED9 | 0.045462781 | 3.166498199 | 1.258702601 | 0.25164154 | 0.655935708 | -6.238638915 |
| **Q96HS1-1** | PGAM5 | 0.091661514 | 3.190458633 | 1.401655368 | 0.207113903 | 0.612994402 | -6.238871048 |
| **Q14703** | MBTPS1 | -0.045673602 | 3.170800804 | -0.676234362 | 0.522315626 | 0.835810201 | -6.238964306 |
| **Q9BY41** | HDAC8 | 0.025013943 | 3.194825611 | 0.776350068 | 0.464964484 | 0.803074528 | -6.239091762 |
| **Q01664** | TFAP4 | 0.026647759 | 3.259465835 | 1.589684266 | 0.159410466 | 0.558776357 | -6.239490672 |
| **P49760-1** | CLK2 | 0.033589331 | 3.16439875 | 0.873867187 | 0.413441018 | 0.771122868 | -6.239768085 |

| | | | | | | | |
|---|---|---|---|---|---|---|---|
| **Q8IZC7** | ZNF101 | -0.018259136 | 3.10301764 | -0.125116931 | 0.904228194 | 0.980210603 | -6.239912673 |
| **Q9HD34** | LYRM4 | 0.055225374 | 3.210898717 | 0.901307097 | 0.399724888 | 0.763210165 | -6.239973881 |
| **P28062-1** | PSMB8 | 0.0209674 | 3.185047628 | 1.787150681 | 0.120540405 | 0.499295152 | -6.240589584 |
| **O75828** | CBR3 | 0.032096693 | 3.165018878 | 1.287632937 | 0.242004504 | 0.648474127 | -6.241006261 |
| **Q6P0Q8** | MAST2 | 0.043679782 | 3.17154741 | 1.208207469 | 0.26925897 | 0.671667979 | -6.241111554 |
| **Q8IUD2-1** | ERC1 | -0.031639897 | 3.187032367 | -1.620336548 | 0.152679711 | 0.552031202 | -6.241280779 |
| **Q6Y2X3** | DNAJC14 | 0.019834782 | 3.178177188 | 0.417469598 | 0.689834266 | 0.908760279 | -6.242355139 |
| **O00273-1** | DFFA | 0.000557784 | 3.183206056 | 0.012357924 | 0.99051232 | 0.996659117 | -6.242543963 |
| **P53671-3** | LIMK2 | -0.028090325 | 3.173663958 | -1.361482828 | 0.218858666 | 0.625479172 | -6.242951454 |
| **P19338** | NCL | 0.060024585 | 3.171808316 | 2.373718216 | 0.052216658 | 0.374493198 | -6.242966854 |
| **Q7Z4K8-1** | TRIM46 | -0.003433812 | 3.187701858 | -0.060740284 | 0.953398928 | 0.989472196 | -6.243132331 |
| **Q9HCJ1** | ANKH | 0.028341147 | 3.190547429 | 0.343184212 | 0.74233638 | 0.933058463 | -6.244086572 |
| **Q03468** | ERCC6 | 0.000409875 | 3.173946581 | 0.008575572 | 0.993416085 | 0.99756865 | -6.244178524 |
| **Q8TF68-1** | ZNF384 | 0.051534652 | 3.182725444 | 0.810228115 | 0.446574536 | 0.792204208 | -6.246352525 |
| **O43493-1** | TGOLN2 | -0.094331117 | 3.220587084 | -0.639696775 | 0.544339755 | 0.844518523 | -6.247053896 |
| **Q96HE9** | PRR11 | -0.019518173 | 3.075855802 | -0.381880154 | 0.71477835 | 0.9235376 | -6.247110864 |
| **Q8WXE0-1** | CASKIN2 | -0.030131538 | 3.172378503 | -1.193120875 | 0.274723279 | 0.674207874 | -6.248190405 |
| **Q86YW9-1** | MED12L | 0.005194252 | 3.20584771 | 0.092052708 | 0.929440346 | 0.986692083 | -6.248436891 |
| **Q9NR19-2** | ACSS2 | 0.029052123 | 3.170318649 | 1.0885761 | 0.315211447 | 0.702089976 | -6.248751103 |
| **Q9UFF9** | CNOT8 | 0.038298839 | 3.153897543 | 0.907951057 | 0.39645552 | 0.762021989 | -6.248826723 |
| **Q9Y366** | IFT52 | 0.027399063 | 3.188446581 | 0.648104159 | 0.539221308 | 0.843540088 | -6.249027108 |
| **P82675** | MRPS5 | 0.095774041 | 3.179047651 | 1.354288116 | 0.221023919 | 0.626484139 | -6.249249259 |
| **Q8IXQ5** | KLHL7 | -0.02557426 | 3.156004003 | -1.316281513 | 0.232780955 | 0.6397249 | -6.249700747 |
| **Q99549** | MPHOSPH8 | -0.027138209 | 3.179688165 | -1.550195189 | 0.168492496 | 0.566974321 | -6.249910353 |
| **P20265** | POU3F2 | -0.038167367 | 3.210571202 | -1.156792691 | 0.288267994 | 0.68315575 | -6.249915734 |
| **Q9BTV5** | FSD1 | 0.024048421 | 3.204517932 | 0.578109959 | 0.58273632 | 0.864744128 | -6.25005863 |
| **P10398-1** | ARAF | 0.067133596 | 3.192250348 | 1.170531893 | 0.283080692 | 0.679787821 | -6.250388608 |
| **Q9H063** | MAF 1.00 | -0.022061762 | 3.175381851 | -0.656473466 | 0.534155984 | 0.840262117 | -6.250728956 |
| **Q8NDV7-6** | TNRC6A | -0.03022418 | 3.20459635 | -1.300050507 | 0.237967943 | 0.64439513 | -6.251968791 |
| **O15265-2** | ATXN7 | -0.025065429 | 3.087049418 | -0.716205223 | 0.498885478 | 0.824970816 | -6.252132308 |
| **Q9BPX7-2** | C7orf25 | -0.032002888 | 3.191758671 | -1.124674291 | 0.300705512 | 0.691292041 | -6.252619807 |
| **Q6ZNE5** | ATG14 | 0.015245258 | 3.102093431 | 0.331990572 | 0.750387823 | 0.935635612 | -6.252643511 |
| **Q9H019-1** | FAM54B; MTFR1L | -0.008202154 | 3.191936165 | -0.057862737 | 0.955603722 | 0.989472196 | -6.252668315 |
| **Q5T6C5** | ATXN7L2 | 0.027204786 | 3.213249107 | 0.386918408 | 0.711223027 | 0.921051693 | -6.252672421 |
| **Q8N4Q1-2** | CHCHD4 | -0.092368394 | 3.208099642 | -0.953852411 | 0.37441766 | 0.746861238 | -6.25282747 |
| **Q13425-1** | SNTB2 | 0.04183186 | 3.185370598 | 1.258457613 | 0.251724553 | 0.655936888 | -6.253268339 |
| **Q14202-2** | ZMYM3 | -0.043762836 | 3.167395549 | -1.799813889 | 0.118386874 | 0.496152504 | -6.253735043 |
| **Q9NXF1** | TEX10 | -0.037668886 | 3.185892078 | -1.685907933 | 0.139173378 | 0.533333364 | -6.254033314 |
| **Q03169** | TNFAIP2 | -0.032241304 | 3.181637736 | -0.673572968 | 0.523900454 | 0.837176492 | -6.254083063 |

| | | | | | | | |
|---|---|---|---|---|---|---|---|
| **P10586-1** | PTPRF | -0.011269492 | 3.199039619 | -0.24824854 | 0.811633902 | 0.955606306 | -6.25408628 |
| **Q3SY77-1** | UGT3A2 | 0.020104134 | 3.204573124 | 0.316071294 | 0.761896746 | 0.939498834 | -6.254655697 |
| **P42680** | TEC | 0.013901294 | 3.200667827 | 0.235753962 | 0.820909238 | 0.958712906 | -6.255094669 |
| **Q53GS7-1** | GLE1 | -0.029909261 | 3.186698557 | -1.524918788 | 0.174556668 | 0.5748429 | -6.256009577 |
| **Q16539** | MAPK14 | -0.010917292 | 3.183261362 | -0.192898449 | 0.852951478 | 0.966856826 | -6.25634827 |
| **Q92844** | TANK | 0.041458313 | 3.181362975 | 0.816987475 | 0.442967685 | 0.789629405 | -6.256385856 |
| **Q8WU10** | PYROXD1 | -0.032067279 | 3.223891473 | -1.074517849 | 0.321013627 | 0.708954025 | -6.257094647 |
| **P55198** | MLLT6 | -0.012906836 | 3.218330845 | -0.187663792 | 0.856887551 | 0.968855822 | -6.257732192 |
| **A0AVK6** | E2F8 | -0.019766445 | 3.158228801 | -0.408877251 | 0.695819686 | 0.911956208 | -6.258028578 |
| **Q9NYF8-1** | BCLAF1 | -0.130166381 | 3.109853435 | -0.680918685 | 0.519533633 | 0.834217303 | -6.258160043 |
| **Q8IYU2-1** | HACE1 | 0.025929971 | 3.190085134 | 0.637256487 | 0.545831033 | 0.845346503 | -6.258362773 |
| **P07902** | GALT | 0.024641219 | 3.188160891 | 1.304868417 | 0.236417767 | 0.642606624 | -6.258603279 |
| **Q96BK5** | PINX1 | -0.072336168 | 3.183933681 | -0.886266969 | 0.407200276 | 0.767181929 | -6.258887352 |
| **Q13045-1** | FLII | 0.028005956 | 3.168255453 | 0.835545718 | 0.433171782 | 0.783962934 | -6.258923201 |
| **Q8N9T8** | KRI1 | -0.021533606 | 3.187718039 | -1.579782129 | 0.161643773 | 0.561414897 | -6.259413545 |
| **P18440** | NAT1 | -0.0234525 | 3.145265971 | -0.604280572 | 0.566228301 | 0.853395233 | -6.259657572 |
| **Q16352** | INA | -0.052065672 | 3.132026244 | -0.769566585 | 0.468709125 | 0.805725242 | -6.260083396 |
| **Q5JTY5** | CBWD3; CBWD5 | -0.042838608 | 3.154501174 | -0.868892925 | 0.415964272 | 0.772518968 | -6.260165107 |
| **P62995** | TRA2B | -0.056800847 | 3.188784945 | -1.246803931 | 0.255700821 | 0.659782078 | -6.260451771 |
| **Q7Z6I6-1** | ARHGAP30 | -0.028043566 | 3.176796983 | -1.357214624 | 0.220140891 | 0.625644783 | -6.26102967 |
| **Q01826-2** | SATB1 | -0.036541582 | 3.177300284 | -1.734781396 | 0.129856435 | 0.515596889 | -6.262011347 |
| **Q86WQ0-2** | NR2C2AP | -0.009630959 | 3.210622493 | -0.14412325 | 0.889787737 | 0.976971196 | -6.262121207 |
| **Q96IT1-2** | ZNF496 | 0.015782458 | 3.126636865 | 0.325439487 | 0.755115786 | 0.937112609 | -6.262681292 |
| **O00308** | WWP2 | 0.027448383 | 3.180403067 | 1.218771312 | 0.26548821 | 0.668347222 | -6.262754657 |
| **Q9BY77** | POLDIP3 | -0.022588801 | 3.179015677 | -0.586017945 | 0.577718633 | 0.861313941 | -6.263160779 |
| **Q9NSU2-1** | TREX1 | 0.022095805 | 3.164030818 | 0.410363042 | 0.69478298 | 0.911200194 | -6.263510806 |
| **Q05469-1** | LIPE | -0.034479515 | 3.177248187 | -1.102530092 | 0.309537346 | 0.697270829 | -6.264155657 |
| **P61225** | RAP2B | 0.026060405 | 3.199234294 | 0.660768479 | 0.531568182 | 0.839849063 | -6.264382389 |
| **Q8NCY6** | MSANTD4 | 0.005027682 | 3.214507044 | 0.089615495 | 0.931302662 | 0.986707862 | -6.264426487 |
| **P62273-2** | RPS29 | -0.046338754 | 3.192449747 | -1.077286885 | 0.319863953 | 0.708257023 | -6.264648462 |
| **Q9BWH2** | FUNDC2 | -0.031592037 | 3.141448845 | -0.203315995 | 0.845131941 | 0.962662834 | -6.264840634 |
| **Q8N9M1-1** | C19orf47 | -0.041072166 | 3.189174518 | -1.249427119 | 0.254801074 | 0.659782078 | -6.26581765 |
| **P62304** | SNRPE | -0.031222069 | 3.18172369 | -1.404563424 | 0.20628612 | 0.612058044 | -6.265938034 |
| **P37235** | HPCAL1 | 0.076177621 | 3.21105033 | 1.195517916 | 0.273848828 | 0.673600291 | -6.26618837 |
| **P07099** | EPHX1 | -0.02908426 | 3.191085713 | -1.298069353 | 0.23860797 | 0.644491512 | -6.266195517 |
| **Q8WV92** | MITD1 | -0.046175696 | 3.16612786 | -1.069544446 | 0.323086961 | 0.710272801 | -6.266987477 |
| **Q9BYK8-1** | HELZ2 | -0.055068277 | 3.19552449 | -0.819924234 | 0.441407082 | 0.788971713 | -6.267400367 |
| **Q6UUV9-2** | CRTC1 | 0.009424161 | 3.197275029 | 0.200743822 | 0.847060916 | 0.963754687 | -6.268630797 |
| **Q9H5X1-1** | FAM96A | -0.04259087 | 3.175178644 | -0.833262518 | 0.434368496 | 0.784298014 | -6.26885371 |

| | | | | | | | |
|---|---|---|---|---|---|---|---|
| **Q15642-2** | TRIP10 | 0.032505437 | 3.176476448 | 1.029047633 | 0.34037507 | 0.722975002 | -6.2689995 |
| **P37287-1** | PIGA | -0.01353901 | 3.174264259 | -0.210229089 | 0.839953306 | 0.961392936 | -6.26922335 |
| **Q6ZNW5** | GDPGP1 | 0.015348186 | 3.185255361 | 0.251001924 | 0.809594295 | 0.955498404 | -6.269229237 |
| **Q9HAZ1** | CLK4 | 0.040368978 | 3.222139163 | 0.996906489 | 0.354615057 | 0.734091082 | -6.269784475 |
| **Q9NYM9** | BET1L | -0.0283327 | 3.185939189 | -0.466672069 | 0.656034793 | 0.896497734 | -6.269937059 |
| **C9JLR9** | C11orf95 | 0.015372784 | 3.192805409 | 0.075858865 | 0.941822769 | 0.988377491 | -6.270333263 |
| **Q96CD2-1** | PPCDC | -0.040374565 | 3.177197147 | -0.769534081 | 0.468727119 | 0.805725242 | -6.270341231 |
| **Q9H4Z2** | ZNF335 | -0.02968844 | 3.207447232 | -0.75747305 | 0.475436446 | 0.809713018 | -6.270566418 |
| **Q969H6** | POP5 | -0.028088474 | 3.11085191 | -0.351434144 | 0.736424656 | 0.929681266 | -6.270672466 |
| **Q9NWS6** | FAM118A | -0.038983453 | 3.174123898 | -1.216980164 | 0.266124362 | 0.66910065 | -6.271915106 |
| **Q9P0N9** | TBC1D7; TBC1D7-I | -0.03728869 | 3.212395297 | -1.04057654 | 0.335379502 | 0.720533987 | -6.272941671 |
| **Q9H6R6-1** | ZDHHC6 | -0.016009539 | 3.198700433 | -0.334462047 | 0.748607161 | 0.935412899 | -6.273274715 |
| **O95803-1** | NDST3 | -0.030642468 | 3.192931712 | -0.72319601 | 0.494859865 | 0.822007449 | -6.273382221 |
| **Q96IX5** | USMG5 | 0.092459045 | 3.224696001 | 0.912537633 | 0.394210301 | 0.760194378 | -6.273440197 |
| **P61457** | PCBD1 | -0.000931119 | 3.198279602 | -0.01620792 | 0.987556785 | 0.99582925 | -6.274043908 |
| **Q9H0L4** | CSTF2T | -0.049680035 | 3.196199639 | -1.548604222 | 0.168868336 | 0.56753762 | -6.274076138 |
| **Q9H3F6-1** | KCTD10 | -0.069733974 | 3.151671173 | -0.817062749 | 0.442927635 | 0.789629405 | -6.274416315 |
| **Q5T0D9** | TPRG1L | -0.089265816 | 3.222167277 | -0.665407933 | 0.528781761 | 0.839849063 | -6.27466331 |
| **P55345-1** | PRMT2 | 0.021732784 | 3.192054601 | 0.463236737 | 0.658367626 | 0.896852471 | -6.274680618 |
| **Q9P0T7** | TMEM9 | 0.055674668 | 3.179988698 | 0.689617229 | 0.514392951 | 0.831675214 | -6.275020832 |
| **Q8WTS6** | SETD7 | 0.055489388 | 3.17267835 | 0.984825657 | 0.360087186 | 0.73661838 | -6.275845381 |
| **P28715-1** | ERCC5 | 0.024944601 | 3.164137576 | 1.598274188 | 0.157496613 | 0.555457754 | -6.277329692 |
| **Q8WXE9-3** | STON2 | -0.027995372 | 3.209118249 | -0.752566694 | 0.478184464 | 0.811328097 | -6.277406782 |
| **Q9Y2K1** | ZBTB1 | -0.035800522 | 3.180409978 | -1.04566733 | 0.333192389 | 0.719098069 | -6.277482359 |
| **O15213** | WDR46 | -0.045555804 | 3.1756652 | -1.166758099 | 0.284497623 | 0.680618374 | -6.277493897 |
| **P51151** | RAB9A | 0.024535647 | 3.171644062 | 0.460874104 | 0.659974433 | 0.896934512 | -6.277705635 |
| **Q3ZCW2** | LGALSL | 0.010968227 | 3.21298289 | 0.267926246 | 0.797093355 | 0.950886893 | -6.277771389 |
| **O43298** | ZBTB43 | -0.029112027 | 3.200896363 | -0.854972174 | 0.423085804 | 0.777884596 | -6.278695285 |
| **O75864-1** | PPP1R37 | 0.022565385 | 3.206642135 | 0.281469717 | 0.787135964 | 0.947114226 | -6.278923807 |
| **Q6XUX3-1** | DSTYK | -0.012961548 | 3.197163917 | -0.290374718 | 0.780612282 | 0.944845305 | -6.279328726 |
| **P53992** | SEC24C | 0.030260373 | 3.186720443 | 1.822043555 | 0.114696641 | 0.489548681 | -6.280361064 |
| **Q9P2S5** | WRAP73 | 0.030378746 | 3.205511999 | 0.792284083 | 0.456250289 | 0.798116417 | -6.280490485 |
| **Q14839-2** | CHD4 | 0.020109469 | 3.173611083 | 2.251721574 | 0.062097533 | 0.385441328 | -6.280589858 |
| **Q9NQE9** | HINT3 | 0.030948862 | 3.170980259 | 0.902397752 | 0.399186815 | 0.763210165 | -6.280667187 |
| **Q6QNY0** | BLOC1S3 | 0.022173982 | 3.175638907 | 0.510129608 | 0.626891309 | 0.885387512 | -6.280995216 |
| **Q6ZS17-4** | FAM65A; RIPOR1 | -0.043565354 | 3.189529097 | -1.021575773 | 0.343644262 | 0.72497442 | -6.281039624 |
| **Q8IWZ8-1** | SUGP1 | -0.030376023 | 3.18652762 | -1.759878489 | 0.125308186 | 0.508724344 | -6.281551306 |
| **Q99676** | ZNF184 | 0.029023803 | 3.157383939 | 0.930568171 | 0.385476925 | 0.754896271 | -6.281774443 |
| **Q8WZA9** | IRGQ | -0.045425984 | 3.164231096 | -1.513434819 | 0.177378204 | 0.578452275 | -6.282134004 |

| | | | | | | | |
|---|---|---|---|---|---|---|---|
| **Q03393** | PTS | -0.005867787 | 3.234483025 | -0.11256872 | 0.913784034 | 0.982219313 | -6.282181452 |
| **Q9BZQ6** | EDEM3 | 0.034463342 | 3.174579995 | 1.423810432 | 0.20088236 | 0.606617388 | -6.282475615 |
| **Q9GZU0** | C6orf62 | -0.00346361 | 3.23402469 | -0.05702858 | 0.956242939 | 0.989472196 | -6.282664581 |
| **P33121** | ACSL1 | 0.025374738 | 3.205005372 | 0.646018091 | 0.540488508 | 0.843540088 | -6.28272556 |
| **Q99856** | ARID3A | -0.008330746 | 3.183630427 | -0.149795036 | 0.885487019 | 0.975708125 | -6.283210751 |
| **Q8N565-2** | MREG | -0.012624388 | 3.194955622 | -0.221972159 | 0.831176417 | 0.960187608 | -6.285493313 |
| **Q96S66** | CLCC1 | -0.03289317 | 3.178070718 | -0.697574923 | 0.50971905 | 0.82918056 | -6.285833944 |
| **Q96TA1** | FAM129B | -0.029009134 | 3.179550906 | -0.81499911 | 0.444026536 | 0.790506794 | -6.285886785 |
| **Q6UXH1-5** | CRELD2 | -0.005292813 | 3.194559785 | -0.11619412 | 0.911021509 | 0.981900848 | -6.285995289 |
| **Q9C073** | FAM117A | -0.032847473 | 3.181358312 | -0.992893272 | 0.3564256 | 0.734600528 | -6.286394857 |
| **Q9BQC6** | MRP63; MRPL57 | 0.06711722 | 3.160561408 | 0.863580585 | 0.4186715 | 0.775191113 | -6.286807589 |
| **Q14872** | MTF1 | -0.012134986 | 3.196588388 | -0.297352352 | 0.77551395 | 0.94357883 | -6.286982166 |
| **O15427** | SLC16A3 | -0.04939244 | 3.248880003 | -0.769762954 | 0.468600434 | 0.805725242 | -6.287396363 |
| **Q9NZC3** | GDE1 | -0.03021909 | 3.236701853 | -0.509495746 | 0.627311393 | 0.885387512 | -6.28745142 |
| **Q969U7** | PSMG2 | -0.058717566 | 3.176249906 | -1.459560241 | 0.191184972 | 0.595771959 | -6.287544359 |
| **O43164** | PJA2 | 0.026824645 | 3.185538039 | 1.204535341 | 0.270580402 | 0.671731232 | -6.287617183 |
| **O95433** | AHSA1 | -0.039093952 | 3.198489776 | -1.891248013 | 0.103913905 | 0.470961767 | -6.288946306 |
| **Q9Y680-1** | FKBP7 | 0.047201188 | 3.17741711 | 1.190138546 | 0.275814554 | 0.67488347 | -6.289289672 |
| **O94992** | HEXIM1 | -0.194485326 | 3.196429005 | -1.42954783 | 0.199296522 | 0.604614147 | -6.289436291 |
| **O75390** | CS | 0.110263985 | 3.18207352 | 1.615779351 | 0.15366316 | 0.552139704 | -6.289496275 |
| **Q9NTM9** | CUTC | -0.033056153 | 3.156395114 | -1.089068148 | 0.315009927 | 0.701837878 | -6.289625971 |
| **Q9ULH1-2** | ASAP1 | 0.024875279 | 3.178313112 | 1.560338496 | 0.166114579 | 0.565363245 | -6.290100312 |
| **Q96MY1** | C20orf112; NOL4L | -0.042291748 | 3.168070598 | -0.850621923 | 0.425329437 | 0.778986255 | -6.290114108 |
| **Q6ZVM7** | TOM1L2 | 0.008895453 | 3.155735832 | 0.1657619 | 0.873402962 | 0.97237605 | -6.290395587 |
| **Q6ZSS7** | MFSD6 | -0.019593881 | 3.15415825 | -0.236808263 | 0.820125347 | 0.958501895 | -6.290438002 |
| **Q8IWI9-4** | MGA | 0.008846591 | 3.204206815 | 0.210228379 | 0.839953838 | 0.961392936 | -6.290898108 |
| **P13639** | EEF2 | -0.016238591 | 3.194370342 | -2.478512434 | 0.045030142 | 0.357262465 | -6.291237956 |
| **Q8N6S5** | ARL6IP6 | -0.006987385 | 3.195139662 | -0.131218673 | 0.899587667 | 0.978757548 | -6.291397124 |
| **P62906** | RPL10A | -0.046626674 | 3.166706443 | -2.006248789 | 0.088165315 | 0.437773138 | -6.291695579 |
| **Q04721** | NOTCH2 | -0.038919572 | 3.226208135 | -1.126451604 | 0.30000579 | 0.690882899 | -6.292347235 |
| **Q9UI95** | MAD2L2 | -0.035929423 | 3.169380424 | -1.135936985 | 0.296294243 | 0.688880164 | -6.293527511 |
| **O14578-4** | CIT | -0.032230906 | 3.16828336 | -1.380221313 | 0.213307981 | 0.61901473 | -6.295639679 |
| **Q9BW85** | CCDC94 | -0.095465152 | 3.207576979 | -1.305751131 | 0.236134715 | 0.642276724 | -6.295741903 |
| **Q96IK1** | BOD1 | -0.037449786 | 3.21159632 | -1.028028137 | 0.340819669 | 0.722995142 | -6.296558694 |
| **O15397-1** | IPO8 | -0.033468782 | 3.180396908 | -1.610643605 | 0.154778624 | 0.553156191 | -6.296997664 |
| **P60520** | GABARAPL2 | -0.073522071 | 3.182653986 | -1.001325339 | 0.352629884 | 0.732738778 | -6.297083964 |
| **Q96CW1** | AP2M1 | -0.048895424 | 3.186164736 | -1.641422938 | 0.148206194 | 0.546231571 | -6.29812636 |
| **O00625** | PIR | -0.028624693 | 3.213435786 | -0.637632177 | 0.545601282 | 0.845155427 | -6.298133178 |
| **O14974** | PPP1R12A | -0.03473195 | 3.200228906 | -1.93892835 | 0.097071684 | 0.455254009 | -6.298210255 |

| | | | | | | | |
|---|---|---|---|---|---|---|---|
| **P26885** | FKBP2 | 0.048000112 | 3.220978162 | 1.263666516 | 0.249964659 | 0.654571265 | -6.298397195 |
| **Q9H0R8-2** | GABARAPL1 | -0.006060441 | 3.179975973 | -0.098562859 | 0.924468224 | 0.985099254 | -6.298581029 |
| **Q7Z5W3** | BCDIN3D | 0.003159636 | 3.194842341 | 0.076746555 | 0.941143519 | 0.988377491 | -6.299217045 |
| **Q99543-1** | DNAJC2 | 0.044002742 | 3.190321962 | 1.710114156 | 0.134481454 | 0.522983434 | -6.299404515 |
| **P22694-2** | PRKACB | 0.018121241 | 3.19702753 | 0.345442802 | 0.740716018 | 0.931611328 | -6.299564883 |
| **P09496-2** | CLTA | -0.248966682 | 3.133806981 | -1.543832864 | 0.170000181 | 0.569040056 | -6.299956947 |
| **Q9UKT8-1** | FBXW2 | -0.006431051 | 3.188998404 | -0.113332677 | 0.913201796 | 0.982219313 | -6.300320742 |
| **P15291** | B4GALT1 | -0.007807144 | 3.187623149 | -0.184259677 | 0.859449583 | 0.968855822 | -6.300357016 |
| **Q9Y462-3** | ZNF711 | -0.002013723 | 3.179611265 | -0.033821182 | 0.974039057 | 0.99316605 | -6.300405225 |
| **Q8NEW0** | SLC30A7 | 0.033986591 | 3.168434802 | 0.5474505 | 0.602426978 | 0.872497827 | -6.301208871 |
| **Q8NCP5** | ZBTB44 | 0.036893671 | 3.178148472 | 1.090103978 | 0.314586045 | 0.701837878 | -6.302033321 |
| **Q13595-1** | TRA2A | 0.035667025 | 3.182516695 | 1.06684243 | 0.324217936 | 0.710703247 | -6.303197618 |
| **Q9BYV8** | CEP41 | 0.037304507 | 3.228523076 | 1.535345353 | 0.172031019 | 0.571776747 | -6.303374218 |
| **Q9UBH6** | XPR1 | 0.027326831 | 3.217054821 | 0.612268261 | 0.561245849 | 0.852576894 | -6.303495611 |
| **Q9H7D0-1** | DOCK5 | 0.044633123 | 3.184681604 | 0.846193735 | 0.427622136 | 0.78052568 | -6.303618687 |
| **Q15111-1** | PLCL1 | 0.034973223 | 3.179163236 | 1.258799801 | 0.251608611 | 0.655935708 | -6.303939595 |
| **Q9UJT0** | TUBE1 | -0.035880697 | 3.195612475 | -0.942764193 | 0.379653488 | 0.750948043 | -6.304131893 |
| **P49757** | NUMB | -0.025456939 | 3.198091645 | -1.246886785 | 0.25567236 | 0.659782078 | -6.304286707 |
| **Q0PNE2** | ELP6 | -0.049952902 | 3.208603618 | -1.279339725 | 0.244733613 | 0.650150213 | -6.304327722 |
| **Q96BZ8** | LENG1 | -0.02626004 | 3.214629472 | -0.825870422 | 0.43825928 | 0.786897076 | -6.304424498 |
| **Q6NUK4** | REEP3 | 0.027762616 | 3.19693438 | 0.602614532 | 0.567270848 | 0.854077485 | -6.304646822 |
| **Q2T9J0** | TYSND1 | -0.016725152 | 3.175848079 | -0.346555467 | 0.739918293 | 0.931345189 | -6.306070831 |
| **O43592** | XPOT | -0.068398073 | 3.175146592 | -1.59895633 | 0.15734556 | 0.555457754 | -6.306319845 |
| **Q15831-1** | STK11 | -0.010054103 | 3.185323411 | -0.164546587 | 0.874321489 | 0.972624507 | -6.306697127 |
| **Q8IWV8** | UBR2 | -0.035408216 | 3.158750586 | -1.529500093 | 0.17344272 | 0.574167672 | -6.307321193 |
| **Q14814-1** | MEF2D | -0.021114494 | 3.210196631 | -0.495736406 | 0.636467146 | 0.88885491 | -6.307492813 |
| **P11172** | UMPS | -0.024709583 | 3.180066735 | -1.664601403 | 0.14343231 | 0.539313633 | -6.307820923 |
| **Q96J42-1** | TXNDC15 | 0.00134502 | 3.210275486 | 0.021649732 | 0.983379632 | 0.995280405 | -6.308072164 |
| **Q9UGY1** | NOL12 | -0.048313867 | 3.215222039 | -0.780782767 | 0.462528749 | 0.802800142 | -6.308101116 |
| **Q9UBD5-2** | ORC3 | 0.027421711 | 3.189842827 | 1.493564391 | 0.182360018 | 0.584921415 | -6.308346322 |
| **Q96S99** | PLEKHF1 | -0.005369665 | 3.224768101 | -0.084317604 | 0.935352472 | 0.986793807 | -6.308971461 |
| **Q8WXK3-1** | ASB13 | 0.002179315 | 3.186811769 | 0.032938752 | 0.974716119 | 0.99316605 | -6.309557849 |
| **Q53H80** | AKIRIN2 | 0.001758315 | 3.203605052 | 0.024182676 | 0.981435521 | 0.994579747 | -6.310060464 |
| **Q8NB15-1** | ZNF511 | -0.029072938 | 3.180621585 | -0.493393838 | 0.638032921 | 0.889065271 | -6.310066738 |
| **O00329** | PIK3CD | 0.039452199 | 3.146713383 | 0.988916747 | 0.358226733 | 0.735429301 | -6.310753171 |
| **Q8IXQ4** | GPALPP1; KIAA170 | -0.030952135 | 3.184190843 | -1.178110253 | 0.280253238 | 0.677109587 | -6.310849318 |
| **P30825** | SLC7A1 | 0.058736089 | 3.205842725 | 0.963087109 | 0.370099617 | 0.744424726 | -6.310868207 |
| **O15155** | BET1 | -0.041503173 | 3.179882 | -0.880934375 | 0.409875535 | 0.769831818 | -6.310904078 |
| **P42677** | RPS27 | -0.082177664 | 3.212156004 | -1.317035471 | 0.232542449 | 0.639692206 | -6.310910259 |

| | | | | | | | |
|---|---|---|---|---|---|---|---|
| **Q96C11-3** | FGGY | 0.036851084 | 3.193757373 | 0.943855231 | 0.379135826 | 0.750619023 | -6.310953773 |
| **Q14114** | LRP8 | 0.020909745 | 3.173270656 | 0.499111692 | 0.634214657 | 0.887509451 | -6.310989487 |
| **O60880-1** | SH2D1A | -0.102805451 | 3.182095922 | -1.243712817 | 0.256764585 | 0.659782078 | -6.311032233 |
| **Q9UBE0-1** | SAE1 | -0.033892861 | 3.158477053 | -1.724825932 | 0.131704345 | 0.519814431 | -6.311047267 |
| **O14524-1** | TMEM194A; NEMP | 0.060469534 | 3.181002865 | 0.72271695 | 0.495135039 | 0.822292618 | -6.311192667 |
| **Q6ZMK1** | CYHR1 | 0.065140107 | 3.202498466 | 0.656899314 | 0.533899052 | 0.840132297 | -6.311325464 |
| **Q9BRF8-1** | CPPED1 | -0.026876748 | 3.204323071 | -0.578182262 | 0.582690328 | 0.864744128 | -6.311766159 |
| **P42336** | PIK3CA | 0.021920054 | 3.170896098 | 1.404733118 | 0.206237909 | 0.612058044 | -6.311773119 |
| **Q9UM22** | EPDR1 | 0.022914698 | 3.204343746 | 0.377952316 | 0.717555457 | 0.924679646 | -6.312030469 |
| **Q5VTB9-3** | RNF220 | -0.029876427 | 3.169910653 | -1.292525065 | 0.240407124 | 0.646142333 | -6.312040476 |
| **Q9HAH7-1** | FBRS | -0.018475851 | 3.180570865 | -0.471295991 | 0.652901415 | 0.895616288 | -6.312138714 |
| **A4D161-1** | FAM221A | -0.025994972 | 3.200795225 | -1.042667638 | 0.334479732 | 0.719394009 | -6.312277972 |
| **Q9Y2V2** | CARHSP1 | 0.025625452 | 3.201653612 | 1.39116048 | 0.210126242 | 0.615470161 | -6.312362261 |
| **Q96M27-1** | PRRC1 | 0.037517457 | 3.179616149 | 1.544387992 | 0.169868133 | 0.569040056 | -6.312677281 |
| **Q5VSY0** | GKAP1 | -0.026964396 | 3.184904971 | -0.926210228 | 0.387574194 | 0.757123654 | -6.313061011 |
| **Q8IYA6** | CKAP2L | -0.033641527 | 3.149717617 | -1.025388254 | 0.341973064 | 0.724140724 | -6.313769412 |
| **Q8NFH3** | NUP43 | -0.026808135 | 3.18244848 | -1.320799578 | 0.231354939 | 0.638769945 | -6.314717912 |
| **Q14781-1** | CBX2 | 0.02602951 | 3.175921085 | 1.387292691 | 0.211246312 | 0.61590617 | -6.315118553 |
| **Q9BTE0-2** | NAT9 | 0.036245559 | 3.193142181 | 0.479743868 | 0.647196449 | 0.89271426 | -6.315320872 |
| **Q6RFH5-1** | WDR74 | -0.036451596 | 3.17106546 | -1.120076147 | 0.302522062 | 0.691344519 | -6.315475117 |
| **Q587I9** | SFT2D3 | -0.014911107 | 3.172028746 | -0.210134795 | 0.840023884 | 0.961392936 | -6.315645022 |
| **Q8IWA0** | WDR75 | -0.031657538 | 3.18094727 | -1.618606186 | 0.153052423 | 0.552031202 | -6.315937619 |
| **Q9NRZ5** | AGPAT4 | -0.001059798 | 3.12846104 | -0.009026581 | 0.993069832 | 0.99756865 | -6.316010645 |
| **Q14966-1** | ZNF638 | 0.01615823 | 3.188591621 | 1.913189817 | 0.10070799 | 0.46303529 | -6.316283953 |
| **O95936-4** | EOMES | -0.02338389 | 3.215176054 | -0.442377636 | 0.672620795 | 0.901479794 | -6.316285971 |
| **Q9P246-2** | STIM2 | -0.0355925 | 3.184505578 | -0.778994963 | 0.463510066 | 0.802800142 | -6.316472539 |
| **O43708** | GSTZ1 | 0.03670836 | 3.164923326 | 1.028455063 | 0.340633431 | 0.722995142 | -6.316502103 |
| **O00522** | KRIT1 | 0.067068315 | 3.208763599 | 1.343856116 | 0.224197324 | 0.629881158 | -6.317387169 |
| **Q9BW61** | DDA1 | -0.046837642 | 3.190571514 | -1.127459764 | 0.299609482 | 0.690882899 | -6.317949508 |
| **Q7Z7K0** | CMC1 | 0.055349286 | 3.201535514 | 0.275160406 | 0.791769415 | 0.948542293 | -6.318403212 |
| **Q7Z7A3** | CTU1 | -0.056498893 | 3.184379706 | -0.948131099 | 0.37711228 | 0.749413973 | -6.318536479 |
| **Q8N2Z9** | APITD1; CENPS; AI | -0.02122212 | 3.173221337 | -0.350459054 | 0.737122382 | 0.929681266 | -6.318793445 |
| **Q86XR8-1** | CEP57 | -0.059176612 | 3.214727664 | -1.03208726 | 0.339052242 | 0.721920542 | -6.319028587 |
| **P11532-1** | DMD | 0.009907922 | 3.194867997 | 0.181282041 | 0.86169215 | 0.968885385 | -6.31965315 |
| **Q9HCL2** | GPAM | -0.030811514 | 3.218813062 | -0.851645377 | 0.424800815 | 0.778986255 | -6.319890076 |
| **P0C7T5** | ATXN1L | 0.025059823 | 3.192226197 | 0.786953236 | 0.459152907 | 0.800168863 | -6.320090216 |
| **Q8IZP0** | ABI1 | -0.036117461 | 3.182599245 | -1.108917426 | 0.306968198 | 0.695615278 | -6.320388179 |
| **Q9Y2G2-5** | CARD8 | -0.01313513 | 3.178210333 | -0.212885481 | 0.837965668 | 0.96111408 | -6.320803879 |
| **Q8WVJ2** | NUDCD2 | -0.035278785 | 3.170900668 | -0.554648764 | 0.597770483 | 0.871651708 | -6.321106604 |

| | | | | | | | |
|---|---|---|---|---|---|---|---|
| **Q9UIB8** | CD84 | -0.056375399 | 3.164763253 | -1.271689072 | 0.247275044 | 0.652280071 | -6.321634366 |
| **P62068-1** | USP46 | -0.010064124 | 3.175450296 | -0.244914498 | 0.814105774 | 0.95620584 | -6.321693581 |
| **O75947-1** | ATP5H | 0.100960315 | 3.170180819 | 1.190885586 | 0.275540857 | 0.674825663 | -6.322199419 |
| **P30876** | POLR2B | 0.023064838 | 3.183383801 | 1.922062241 | 0.09943954 | 0.459999988 | -6.322239601 |
| **O95197** | RTN3 | -0.030608431 | 3.19388189 | -0.764495013 | 0.471522299 | 0.807380316 | -6.322426932 |
| **Q8IWY9** | CDAN1 | 0.023743281 | 3.197335431 | 0.611294066 | 0.561852102 | 0.852576894 | -6.322577407 |
| **P51116** | FXR2 | 0.038398532 | 3.167091426 | 1.34050745 | 0.225224537 | 0.630229362 | -6.322602764 |
| **P10619** | CTSA | 0.039998458 | 3.169025158 | 1.359675448 | 0.219400811 | 0.625602181 | -6.322708272 |
| **Q7Z2W4-3** | ZC3HAV1 | -0.005426062 | 3.255804234 | -0.034133117 | 0.973799724 | 0.99316605 | -6.322911596 |
| **Q9HB90** | RRAGC | 0.037790792 | 3.207888545 | 0.794259422 | 0.455177997 | 0.797877083 | -6.323119238 |
| **Q99583** | MNT | 0.02763336 | 3.207012454 | 0.562742128 | 0.592559378 | 0.869575962 | -6.323294003 |
| **O95503** | CBX6 | -0.000474855 | 3.127067158 | -0.007514788 | 0.994230485 | 0.997608555 | -6.323580235 |
| **Q1RMZ1** | C7orf60; BMT2 | -0.025124437 | 3.145016538 | -0.517426281 | 0.622066411 | 0.883385606 | -6.3236878 |
| **Q6XQN6-2** | NAPRT1; NAPRT | -0.013080127 | 3.149428167 | -0.273240254 | 0.793181374 | 0.949070183 | -6.324274257 |
| **Q8NAT1** | GTDC2; POMGNT2 | 0.01488741 | 3.211662148 | 0.366716693 | 0.725525006 | 0.926648409 | -6.32434165 |
| **Q9Y3E1** | HDGFRP3; HDGFL[illegible] | -0.040491024 | 3.146483952 | -0.768795798 | 0.46913593 | 0.805725242 | -6.325324204 |
| **Q9H7F0-1** | ATP13A3 | 0.03927309 | 3.186359377 | 0.849637087 | 0.425838563 | 0.779559304 | -6.325328155 |
| **P06126** | CD1A | 0.031345682 | 3.173443369 | 0.840303029 | 0.430685925 | 0.782692325 | -6.32571102 |
| **Q8N766-1** | EMC1 | -0.03425858 | 3.183727252 | -1.360256517 | 0.219226382 | 0.625602181 | -6.326158931 |
| **Q9BSJ8** | ESYT1 | -0.047088246 | 3.186316645 | -1.697046557 | 0.136995227 | 0.528106296 | -6.326248807 |
| **Q66LE6** | PPP2R2D | -0.034261039 | 3.207227129 | -0.242592874 | 0.815828401 | 0.956293397 | -6.327281348 |
| **P09493-2** | TPM1 | -0.001689831 | 3.145289677 | -0.036093162 | 0.972295944 | 0.992928787 | -6.328716876 |
| **Q9HAD4** | WDR41 | 0.028210532 | 3.201747696 | 1.309042506 | 0.235081924 | 0.64164013 | -6.328762734 |
| **P37275-2** | ZEB1 | -0.030761921 | 3.196155816 | -1.212673134 | 0.267659404 | 0.67010832 | -6.328762942 |
| **P51957** | NEK4 | 0.041149617 | 3.179952648 | 1.161297247 | 0.286558549 | 0.682057421 | -6.328998459 |
| **Q9H469** | FBXL15 | -0.006359748 | 3.205925188 | -0.127771663 | 0.902208691 | 0.979567411 | -6.32946721 |
| **Q9NXD2** | MTMR10 | -0.027117421 | 3.166644814 | -0.887153398 | 0.40675683 | 0.766710087 | -6.329531049 |
| **O95248-4** | SBF1 | 0.027899865 | 3.173908076 | 1.226312273 | 0.262824202 | 0.666498017 | -6.32958093 |
| **P29372** | MPG | -0.026712697 | 3.197564224 | -1.108710069 | 0.307051325 | 0.695615278 | -6.329702604 |
| **Q9Y4G2** | PLEKHM1 | 0.00102161 | 3.167554423 | 0.018394642 | 0.985878191 | 0.995531556 | -6.329944507 |
| **Q8TCG1** | KIAA1524; CIP2A | -0.062109792 | 3.190273197 | -1.65353773 | 0.145692489 | 0.543674934 | -6.330356233 |
| **P05388** | RPLP0 | -0.036400936 | 3.188136056 | -1.928182245 | 0.098573809 | 0.457458476 | -6.33053287 |
| **Q9NWM0-5** | SMOX | -0.044029954 | 3.169474537 | -1.077489732 | 0.319779865 | 0.708257023 | -6.331348073 |
| **Q6ICG6-1** | KIAA0930 | -0.028067192 | 3.184101956 | -0.604513801 | 0.566082447 | 0.853395233 | -6.331410872 |
| **Q8NFH5** | NUP35 | 0.030298303 | 3.192696112 | 1.245449737 | 0.256166379 | 0.659782078 | -6.331882366 |
| **Q9Y6I8-1** | PXMP4 | 0.064252356 | 3.188364841 | 0.847335795 | 0.427029977 | 0.779764001 | -6.331986995 |
| **Q9Y6I9** | TEX264 | -0.03986963 | 3.203577386 | -1.338995591 | 0.225689672 | 0.631141413 | -6.333324419 |
| **Q86YP4-3** | GATAD2A | 0.039885067 | 3.200642233 | 1.751806623 | 0.126753981 | 0.509635359 | -6.33335998 |
| **P20340-1** | RAB6A | -0.033008545 | 3.19093386 | -1.308542513 | 0.235241587 | 0.64164013 | -6.334040483 |

| | | | | | | | |
|---|---|---|---|---|---|---|---|
| **Q9BUB7** | TMEM70 | 0.114665894 | 3.217317456 | 1.437109258 | 0.197223876 | 0.603708078 | -6.334276082 |
| **Q15007** | WTAP | 0.029873808 | 3.172021629 | 1.577369708 | 0.162192293 | 0.561489222 | -6.334825261 |
| **O95405-1** | ZFYVE9 | -0.018323936 | 3.19037001 | -0.453420842 | 0.665056124 | 0.898624353 | -6.334835074 |
| **Q6ZUT1-2** | C11orf57; NKAPD1 | 0.022739875 | 3.159466194 | 0.600236902 | 0.568760665 | 0.854862558 | -6.334843805 |
| **Q9BTV6** | DPH7; WDR85 | 0.005212092 | 3.189880646 | 0.115752318 | 0.911358088 | 0.981900848 | -6.335266295 |
| **P46527** | CDKN1B | 0.038761933 | 3.089958725 | 0.203607515 | 0.844913391 | 0.962551891 | -6.335510115 |
| **Q8TCT7** | SPPL2B | -0.02189274 | 3.161686721 | -0.504806923 | 0.630423524 | 0.886812128 | -6.3356954 |
| **P51817** | PRKX | 0.006304375 | 3.185247755 | 0.12347542 | 0.905477309 | 0.98075504 | -6.335808177 |
| **Q9H0Z9** | RBM38 | -0.005216304 | 3.215929549 | -0.134874934 | 0.896809026 | 0.978757548 | -6.335949116 |
| **P61088** | UBE2N | -0.055889911 | 3.176102584 | -1.491852764 | 0.182795128 | 0.585371742 | -6.336184397 |
| **Q8WXW3** | PIBF1 | 0.034409398 | 3.167767009 | 1.251300087 | 0.254160327 | 0.659041711 | -6.336641878 |
| **Q96RD7-1** | PANX1 | 0.008636792 | 3.183531642 | 0.123622178 | 0.905365621 | 0.98075504 | -6.3369807 |
| **P33316-2** | DUT | 0.146655211 | 3.165738447 | 1.118606093 | 0.303104737 | 0.692399426 | -6.337062326 |
| **Q13188-2** | STK3 | 0.037074049 | 3.19436717 | 1.181915456 | 0.278842551 | 0.676427621 | -6.337431334 |
| **Q6NT76-5** | HMBOX1 | 0.01377597 | 3.158049163 | 0.373454301 | 0.72074142 | 0.925644897 | -6.337859744 |
| **P06241-1** | FYN | -0.014654512 | 3.166618595 | -0.352367077 | 0.735757349 | 0.929681266 | -6.338000621 |
| **Q6UUV7-1** | CRTC3 | 0.006532613 | 3.203664723 | 0.16355011 | 0.875074778 | 0.972759481 | -6.3382805 |
| **P27487** | DPP4 | 0.020648704 | 3.207673101 | 0.496697167 | 0.635825557 | 0.888277483 | -6.338364448 |
| **Q7RTS9-1** | DYM | 0.035879475 | 3.180364649 | 1.064720908 | 0.325108181 | 0.710703247 | -6.33883751 |
| **P41440** | SLC19A1 | -0.026769217 | 3.23914626 | -0.415546968 | 0.691171489 | 0.909522034 | -6.33888502 |
| **Q9H2J4** | PDCL3 | -0.07895919 | 3.203764955 | -1.396330909 | 0.208637283 | 0.614302382 | -6.338989604 |
| **Q9H3H1** | TRIT1 | 0.046215266 | 3.203890007 | 0.965713688 | 0.368878513 | 0.743611616 | -6.339334349 |
| **Q96FS4** | SIPA1 | -0.002025024 | 3.178973797 | -0.036822408 | 0.971736486 | 0.992928787 | -6.33973408 |
| **P24539** | ATP5F1 | 0.199021585 | 3.188788476 | 1.596338679 | 0.157925954 | 0.556664466 | -6.340497034 |
| **Q9H089** | LSG1 | -0.041089234 | 3.18733306 | -1.631950616 | 0.150200198 | 0.547403934 | -6.341084068 |
| **Q9Y3A5** | SBDS | -0.043879512 | 3.175912933 | -1.460138663 | 0.191031641 | 0.595661847 | -6.341346917 |
| **P51610-4** | HCFC1 | 0.023310822 | 3.195661614 | 0.475503854 | 0.650056592 | 0.893733406 | -6.342029676 |
| **Q9Y5Z7** | HCFC2 | 0.042455933 | 3.19966746 | 0.8876359 | 0.406515605 | 0.766437228 | -6.342113057 |
| **Q96GG9** | DCUN1D1 | -0.054021852 | 3.189653642 | -1.26948185 | 0.248012508 | 0.652351802 | -6.342407945 |
| **Q6IAN0** | DHRS7B | -0.027491059 | 3.184677901 | -0.626024835 | 0.552727221 | 0.847927742 | -6.342518208 |
| **Q9UF56-1** | FBXL17 | 0.044020297 | 3.191530341 | 0.857877611 | 0.421592143 | 0.777315429 | -6.342851417 |
| **Q99757** | TXN2 | 0.008558105 | 3.207892665 | 0.179112306 | 0.863327135 | 0.969625966 | -6.343063962 |
| **Q9H6S1-1** | AZI2 | 0.004723099 | 3.218475123 | 0.043826438 | 0.966364107 | 0.992001409 | -6.343553881 |
| **Q5VZK9** | LRRC16A; CARMIL | 0.015885021 | 3.181299629 | 0.333417763 | 0.749359349 | 0.935521689 | -6.344423157 |
| **Q9ULD4** | BRPF3 | -0.004429698 | 3.205299347 | -0.07842228 | 0.93986142 | 0.988002681 | -6.345736672 |
| **Q8WWK9-1** | CKAP2 | -0.053953188 | 3.158844753 | -1.305345125 | 0.236264868 | 0.642410807 | -6.345857145 |
| **Q9H813-2** | TMEM206 | -0.00502443 | 3.168039405 | -0.085528029 | 0.934427011 | 0.986793807 | -6.346051503 |
| **Q8NG11-1** | TSPAN14 | -0.012943586 | 3.120377137 | -0.161137034 | 0.876899531 | 0.973040052 | -6.347012901 |
| **Q96EQ0** | SGTB | -0.031376522 | 3.196487233 | -0.938653512 | 0.381608727 | 0.752847698 | -6.347528196 |

| | | | | | | | |
|---|---|---|---|---|---|---|---|
| **P20338** | RAB4A | -0.025161025 | 3.19907548 | -1.306968577 | 0.235744816 | 0.641994289 | -6.34753168 |
| **Q96HQ2-1** | CDKN2AIPNL | 0.052087545 | 3.177739934 | 0.692766511 | 0.512539926 | 0.831227449 | -6.348168769 |
| **Q9BSM1** | PCGF1 | -0.033049697 | 3.190603296 | -0.279905181 | 0.788284066 | 0.947805127 | -6.348633101 |
| **Q8IYS2-2** | KIAA2013 | -0.040116235 | 3.191333169 | -0.968434031 | 0.367617108 | 0.742429568 | -6.348894302 |
| **Q13127-1** | REST | -0.025730691 | 3.197537312 | -1.214384997 | 0.267048385 | 0.67010832 | -6.349105092 |
| **P10747-7** | CD28 | 0.026346817 | 3.141059708 | 0.403714678 | 0.699427358 | 0.913245694 | -6.349439091 |
| **Q92564-3** | DCUN1D4 | -0.021088829 | 3.179312508 | -0.374729687 | 0.719837442 | 0.925644897 | -6.349592743 |
| **Q13574** | DGKZ | 0.033219002 | 3.173076332 | 1.344386442 | 0.224035028 | 0.629881158 | -6.349711793 |
| **O95295** | SNAPIN | -0.006222524 | 3.164878687 | -0.109960563 | 0.915772235 | 0.98295196 | -6.349720235 |
| **Q9UJA3-4** | MCM8 | -0.033866693 | 3.197469739 | -1.000497583 | 0.353001087 | 0.732738778 | -6.349907418 |
| **Q0VG06** | C17orf70; FAAP100 | -0.023134922 | 3.200238754 | -0.62173003 | 0.555378242 | 0.849861351 | -6.349998614 |
| **Q9H0W9** | C11orf54 | 0.02885944 | 3.18087001 | 1.298308453 | 0.238530647 | 0.644491512 | -6.350014293 |
| **O15417-1** | TNRC18 | -0.001434829 | 3.170587887 | -0.027700413 | 0.978735783 | 0.99361817 | -6.350251711 |
| **Q12912-1** | LRMP | 0.058338187 | 3.172193259 | 1.172787305 | 0.282236704 | 0.679496093 | -6.350335717 |
| **Q9UHB7-1** | AFF4 | -0.033433205 | 3.173919063 | -1.333433115 | 0.227408351 | 0.633950648 | -6.350541519 |
| **P51991-1** | HNRNPA3 | 0.134279311 | 3.196758462 | 1.87439828 | 0.106443867 | 0.475403904 | -6.352222166 |
| **Q5SWX8-1** | C1orf27; ODR4 | 0.023456832 | 3.184613049 | 1.032923794 | 0.338688909 | 0.721920542 | -6.352371944 |
| **Q12962** | TAF10 | 0.030296508 | 3.219057976 | 0.869104611 | 0.415856662 | 0.772499691 | -6.353520013 |
| **O95427** | PIGN | -0.009802318 | 3.217238 | -0.165564202 | 0.873552367 | 0.97237605 | -6.353697819 |
| **Q96NT0** | CCDC115 | 0.017786308 | 3.191417268 | 0.337996209 | 0.746063751 | 0.934974996 | -6.354195903 |
| **Q9NPF5** | DMAP1 | -0.03278255 | 3.186668716 | -1.340439271 | 0.225245495 | 0.630229362 | -6.354282557 |
| **Q8TAF3-1** | WDR48 | 0.023433417 | 3.160857978 | 1.509694192 | 0.178306325 | 0.57989621 | -6.354336332 |
| **O75808-1** | CAPN15; SOLH | 0.038143243 | 3.173003354 | 1.127187718 | 0.29971638 | 0.690882899 | -6.354538332 |
| **O60921** | HUS1 | -0.033870042 | 3.208717407 | -1.395138633 | 0.208979785 | 0.614302382 | -6.354559266 |
| **Q8NEL9-1** | DDHD1 | -0.021786332 | 3.187224834 | -1.163953391 | 0.285554558 | 0.681580409 | -6.354804195 |
| **P61026** | RAB10 | -0.054455633 | 3.178393731 | -1.417652454 | 0.202597166 | 0.608695713 | -6.355499898 |
| **Q9H3S5** | PIGM | -0.008498942 | 3.209350027 | -0.192159499 | 0.85350684 | 0.967210362 | -6.355863182 |
| **Q9Y2R4** | DDX52 | -0.034997459 | 3.161099791 | -0.889361284 | 0.405653872 | 0.766336431 | -6.355867712 |
| **P34059** | GALNS | -0.015804753 | 3.198230459 | -0.380864532 | 0.715495978 | 0.924177458 | -6.356107795 |
| **Q9NTJ4-4** | MAN2C1 | -0.010972863 | 3.159024697 | -0.267185175 | 0.797639413 | 0.951252648 | -6.356193078 |
| **Q9H1K0** | ZFYVE20; RBSN | 0.00511183 | 3.175151063 | 0.070376097 | 0.946019265 | 0.989472196 | -6.356693546 |
| **Q8NFJ9-2** | DPP3; BBS1 | 0.059463 | 3.150058318 | 0.267518675 | 0.797393658 | 0.951102329 | -6.3574593 |
| **P78563-5** | ADARB1 | 0.027026539 | 3.199892158 | 0.775329773 | 0.465526385 | 0.803432467 | -6.357476017 |
| **O75665** | OFD1 | -0.025777879 | 3.165001239 | -0.945792229 | 0.378218117 | 0.750267026 | -6.358318289 |
| **Q14318-2** | FKBP8 | 0.039139793 | 3.172236188 | 1.416510102 | 0.20291673 | 0.608695713 | -6.358438604 |
| **P08648** | ITGA5 | -0.032624237 | 3.147458146 | -1.059142154 | 0.327458613 | 0.712392847 | -6.358520452 |
| **Q99728** | BARD1 | -0.029468974 | 3.175183323 | -1.41011843 | 0.204713173 | 0.610527836 | -6.35899497 |
| **Q16760-1** | DGKD | -0.015565033 | 3.176573011 | -0.279549392 | 0.788545234 | 0.947805127 | -6.359817019 |
| **Q16658** | FSCN1 | -0.042353197 | 3.161760817 | -0.690341921 | 0.513966161 | 0.831492802 | -6.359952283 |

| | | | | | | | |
|---|---|---|---|---|---|---|---|
| **O43670-4** | ZNF207 | -0.038794167 | 3.204644557 | -1.36186142 | 0.218745254 | 0.625479172 | -6.360011488 |
| **Q9H5U6-1** | ZCCHC4 | -0.018365761 | 3.215216293 | -0.542910534 | 0.605374288 | 0.874292777 | -6.360154447 |
| **Q9NX76** | CMTM6 | -0.044483708 | 3.168927427 | -0.802047279 | 0.450967671 | 0.79513757 | -6.36049883 |
| **P32019-1** | INPP5B | 0.022550238 | 3.172383625 | 1.388032529 | 0.211031649 | 0.61590617 | -6.360861266 |
| **P38398-7** | BRCA1 | -0.029929844 | 3.170807341 | -1.512380253 | 0.177639409 | 0.578894629 | -6.362123447 |
| **Q8WUY1** | THEM6 | -0.058030763 | 3.163456725 | -0.834780778 | 0.433572452 | 0.783962934 | -6.36295551 |
| **Q8TCB7-1** | METTL6 | 0.012003311 | 3.195117761 | 0.269510034 | 0.795926756 | 0.950729912 | -6.363248682 |
| **A1L020** | MEX3A | -0.000735541 | 3.179563969 | -0.018061601 | 0.986133839 | 0.995531556 | -6.363517726 |
| **Q9BZM5** | ULBP2 | 0.01910753 | 3.209933129 | 0.545576409 | 0.603642647 | 0.873076436 | -6.364417636 |
| **Q5JRA6-1** | MIA3 | -0.032215197 | 3.20444167 | -1.849434553 | 0.110303727 | 0.481518193 | -6.36498315 |
| **Q9BQI3-1** | EIF2AK1 | 0.012747603 | 3.17429343 | 0.328067626 | 0.753217652 | 0.935926532 | -6.365514118 |
| **P61073-2** | CXCR4 | 0.015082358 | 3.14825298 | 0.107951221 | 0.917304407 | 0.983235402 | -6.36575174 |
| **P08582-1** | MFI2; MELTF | -0.014545413 | 3.137337429 | -0.350279543 | 0.73725086 | 0.929681266 | -6.365787657 |
| **O43861-1** | ATP9B | -0.018907602 | 3.171797367 | -0.455217149 | 0.663829619 | 0.898182275 | -6.366018237 |
| **Q9Y3C7** | MED31 | 0.010314635 | 3.194739181 | 0.230888062 | 0.824529993 | 0.959077751 | -6.36614537 |
| **Q17RS7** | GEN1 | -0.040799912 | 3.200915794 | -0.647945321 | 0.53931773 | 0.843540088 | -6.366177169 |
| **Q9UMZ2-1** | SYNRG | 0.00370658 | 3.213977312 | 0.09453823 | 0.927541612 | 0.985861954 | -6.366856935 |
| **Q96H22-3** | CENPN | -0.025342269 | 3.192594747 | -0.959922621 | 0.371574946 | 0.744717822 | -6.367224367 |
| **Q96MD7-1** | C9orf85 | 0.035561677 | 3.191986564 | 0.667198801 | 0.52770866 | 0.839849063 | -6.367525577 |
| **Q8NBZ7-2** | UXS1 | -0.009806575 | 3.165990306 | -0.205337543 | 0.843616712 | 0.962543524 | -6.367928781 |
| **Q6ZTU2-6** | | 0.026397781 | 3.167911144 | 0.388347289 | 0.710216133 | 0.920650543 | -6.368159743 |
| **P49137** | MAPKAPK2 | 0.02705244 | 3.16569868 | 1.405651629 | 0.205977129 | 0.612058044 | -6.3684893 |
| **Q9Y2S0-2** | POLR1D | -0.045387946 | 3.210209671 | -1.062901696 | 0.325873145 | 0.710703247 | -6.368607618 |
| **Q9Y2W2** | WBP11 | -0.027625811 | 3.176801276 | -1.635212841 | 0.149510621 | 0.546464774 | -6.368788959 |
| **Q9Y5N5** | N6AMT1 | 0.023922989 | 3.193629083 | 0.418753929 | 0.688941663 | 0.908052188 | -6.36965829 |
| **Q86SQ0-1** | PHLDB2 | -0.019072282 | 3.199156874 | -0.443395248 | 0.671921976 | 0.901355316 | -6.370438979 |
| **Q64LD2-1** | WDR25 | -0.021924134 | 3.188414627 | -0.587922175 | 0.576514175 | 0.860654851 | -6.37049151 |
| **O00507-1** | USP9Y | -0.009966082 | 3.151305165 | -0.18323412 | 0.860221809 | 0.968855822 | -6.371382301 |
| **Q9NRH2** | SNRK | 0.033244537 | 3.219880817 | 0.766061156 | 0.470652332 | 0.806934135 | -6.371395883 |
| **O00478** | BTN3A3 | 0.023624438 | 3.183606225 | 0.607726861 | 0.56407538 | 0.852578515 | -6.371486358 |
| **P19484-1** | TFEB | 0.00186037 | 3.231047451 | 0.045954868 | 0.964731877 | 0.991821268 | -6.371950639 |
| **Q6ZU35** | KIAA1211 | 0.023715565 | 3.187600215 | 0.61680386 | 0.558428489 | 0.851249874 | -6.372058846 |
| **Q6UW68** | TMEM205 | -0.017219163 | 3.174440482 | -0.322225157 | 0.757439795 | 0.9382381 | -6.372100723 |
| **Q9BV20-1** | MRI1 | -0.022984123 | 3.178870119 | -1.131307936 | 0.298100754 | 0.689670498 | -6.372120654 |
| **Q969T4** | UBE2E3 | 0.011507354 | 3.186696232 | 0.153017632 | 0.883045298 | 0.974883858 | -6.372399147 |
| **Q12893** | TMEM115 | -0.019020935 | 3.172810162 | -0.350476661 | 0.73710978 | 0.929681266 | -6.373119658 |
| **Q53GQ0** | HSD17B12 | 0.060889821 | 3.147816077 | 1.280594605 | 0.244318943 | 0.650150213 | -6.37414658 |
| **Q2TAK8-1** | MUM1 | -0.025429836 | 3.217952238 | -0.928851013 | 0.386302279 | 0.755953597 | -6.374474601 |
| **Q96SZ6-3** | CDK5RAP1 | 0.087971396 | 3.140989559 | 1.091010198 | 0.314215585 | 0.701837878 | -6.374886098 |

| | | | | | | | |
|---|---|---|---|---|---|---|---|
| **Q5T7M9-1** | FAM69A | -0.010794466 | 3.17925291 | -0.145388122 | 0.888828273 | 0.976409291 | -6.375332379 |
| **Q9P2E5-1** | CHPF2 | -0.034146849 | 3.172038126 | -0.982026896 | 0.361364286 | 0.736919727 | -6.375399588 |
| **Q7Z5Y7** | KCTD20 | -0.02917303 | 3.154545767 | -0.759320215 | 0.47440466 | 0.809053352 | -6.376009234 |
| **Q9Y3T9** | NOC2L | -0.034066339 | 3.184092383 | -1.568147285 | 0.164305332 | 0.563646744 | -6.376011836 |
| **Q9BSW2-2** | EFCAB4B; CRACR2 | -0.037164079 | 3.168792752 | -1.370648116 | 0.216127799 | 0.621925158 | -6.376861139 |
| **P20936-1** | RASA1 | 0.02347393 | 3.194892289 | 1.357683962 | 0.219999569 | 0.625644783 | -6.3771939 |
| **Q99717** | SMAD5 | 0.000486556 | 3.184026678 | 0.007928145 | 0.993913136 | 0.997608555 | -6.377385666 |
| **Q01831** | XPC | 0.026495148 | 3.208991842 | 0.934247929 | 0.383712764 | 0.754456977 | -6.377539542 |
| **O14893** | GEMIN2 | -0.025305016 | 3.170183652 | -1.266607442 | 0.248975756 | 0.654137693 | -6.378169688 |
| **Q9Y6R4-2** | MAP3K4 | -0.020434148 | 3.188601096 | -1.500770194 | 0.180538686 | 0.582844315 | -6.378189942 |
| **Q9NPH2** | ISYNA1 | -0.013999462 | 3.195447202 | -0.199889299 | 0.847702011 | 0.964069922 | -6.378737066 |
| **Q13042** | CDC16 | 0.02284048 | 3.186828614 | 1.639241999 | 0.148663065 | 0.546231571 | -6.378757508 |
| **Q9H082** | RAB33B | 0.013996539 | 3.182625714 | 0.329719592 | 0.752025495 | 0.93570301 | -6.379185657 |
| **P58546** | LUZP6; MTPN | -0.120587173 | 3.219954599 | -1.48288735 | 0.185089858 | 0.588198522 | -6.379477888 |
| **Q92558** | WASF1 | -0.026549556 | 3.135086317 | -0.942155305 | 0.37994262 | 0.750948043 | -6.380048148 |
| **Q96MD2** | C12orf66 | -0.023685551 | 3.163181129 | -0.46940115 | 0.654184527 | 0.896370708 | -6.380599065 |
| **Q8N3R9-1** | MPP5 | 0.026347007 | 3.219058518 | 0.742407236 | 0.483909005 | 0.814028593 | -6.380666425 |
| **Q14119** | VEZF1 | 0.030339998 | 3.176095059 | 0.83696082 | 0.432431264 | 0.783333621 | -6.380826649 |
| **P04843** | RPN1 | -0.040194768 | 3.198418442 | -1.850338763 | 0.110161554 | 0.481294133 | -6.381153024 |
| **O94964-2** | SOGA1 | -0.024037546 | 3.180759552 | -1.427832047 | 0.199769576 | 0.605177234 | -6.381425595 |
| **Q49AN0-1** | CRY2 | 0.012525073 | 3.232152966 | 0.278269273 | 0.789485156 | 0.947865984 | -6.381453269 |
| **Q68DK7** | MSL1 | -0.032222923 | 3.176683223 | -1.151125891 | 0.290430638 | 0.684803131 | -6.381506679 |
| **Q9UDW1** | UQCR10 | 0.026095961 | 3.193291661 | 0.608427038 | 0.563638575 | 0.852578515 | -6.381728 |
| **Q13885** | TUBB2A | 0.021744742 | 3.168648807 | 0.675084786 | 0.522999807 | 0.836627689 | -6.382249751 |
| **Q9NUP1** | BLOC1S4 | -0.021882782 | 3.175215056 | -0.727215952 | 0.492554814 | 0.820752517 | -6.382545439 |
| **Q92562** | FIG4 | 0.005488734 | 3.180225865 | 0.096375925 | 0.926138095 | 0.985528933 | -6.382751728 |
| **O96006** | ZBED1 | 0.02811387 | 3.206290591 | 0.858724926 | 0.421157271 | 0.777257728 | -6.383094832 |
| **Q9H4G4** | GLIPR2 | -0.020459322 | 3.195117865 | -0.801134286 | 0.451459836 | 0.79513757 | -6.383290321 |
| **Q96I24-1** | FUBP3 | 0.033860982 | 3.173231667 | 1.635121824 | 0.14952982 | 0.546464774 | -6.38378142 |
| **Q76N32** | CEP68 | 0.001672905 | 3.127849868 | 0.03972539 | 0.969509568 | 0.992476747 | -6.384138646 |
| **O60234** | GMFG | -0.092788665 | 3.16722936 | -0.945781109 | 0.378223381 | 0.750267026 | -6.384612132 |
| **P05114** | HMGN1 | 0.028622362 | 3.172158204 | 0.363504028 | 0.727810668 | 0.92710315 | -6.384989391 |
| **P38571** | LIPA | -0.032000192 | 3.168622224 | -1.205952411 | 0.270069809 | 0.671731232 | -6.386347299 |
| **O95455** | TGDS | 0.014698365 | 3.214902139 | 0.31995074 | 0.759085887 | 0.9384213 | -6.386691437 |
| **Q8WUU5** | GATAD1 | 0.009402416 | 3.207380283 | 0.264550456 | 0.799581803 | 0.95228263 | -6.387078447 |
| **Q96GE4** | CEP95 | -0.038125843 | 3.2062123 | -0.931629933 | 0.38496726 | 0.754667212 | -6.387419924 |
| **Q8WUX2** | CHAC2 | -0.043121358 | 3.19794453 | -0.777525514 | 0.464317721 | 0.803074528 | -6.388109432 |
| **Q8WYB5** | KAT6B | -0.034738666 | 3.204359113 | -0.899531601 | 0.400601986 | 0.763806762 | -6.388288909 |
| **Q9BU02-1** | THTPA | 0.026079538 | 3.170850973 | 0.985891166 | 0.359601912 | 0.73661838 | -6.388982919 |

| | | | | | | | |
|---|---|---|---|---|---|---|---|
| **Q9NV35** | NUDT15 | 0.023436777 | 3.196913837 | 1.491146248 | 0.182975009 | 0.585513649 | -6.389271333 |
| **O75843** | AP1G2 | -0.026021948 | 3.181768844 | -1.386637688 | 0.211436524 | 0.61590617 | -6.389490314 |
| **Q96K12** | FAR2 | 0.039570331 | 3.214599337 | 0.630507225 | 0.549968675 | 0.846638465 | -6.389648941 |
| **Q9NTZ6** | RBM12 | -0.034535721 | 3.198220257 | -1.926042975 | 0.098875578 | 0.458056247 | -6.390885982 |
| **Q8WVX9** | FAR1 | 0.043602574 | 3.233239115 | 1.324837847 | 0.230086891 | 0.637575603 | -6.391115901 |
| **Q86UL3** | AGPAT6; GPAT4 | -0.019802908 | 3.191136944 | -0.392809284 | 0.707075952 | 0.91801515 | -6.391502879 |
| **Q52LJ0-2** | FAM98B | 0.025081759 | 3.17928541 | 1.341674303 | 0.224866129 | 0.630229362 | -6.391613764 |
| **Q86YC3** | LRRC33; NRROS | 0.019007693 | 3.233315417 | 0.302574443 | 0.771706196 | 0.942488971 | -6.392306399 |
| **P23511** | NFYA | -0.007953509 | 3.217383891 | -0.188470594 | 0.856280603 | 0.968855822 | -6.39325399 |
| **Q14592** | ZNF460 | 0.015793424 | 3.222233936 | 0.132356682 | 0.89872265 | 0.978757548 | -6.3934155 |
| **Q96HR3** | MED30 | 0.022213362 | 3.194284892 | 0.696661887 | 0.510253907 | 0.829456831 | -6.393679775 |
| **Q96S82** | UBL7 | -0.002952052 | 3.186727811 | -0.061982301 | 0.952447423 | 0.989472196 | -6.393839901 |
| **O75140-4** | DEPDC5 | 0.019038108 | 3.157965563 | 0.603889018 | 0.566473218 | 0.853395233 | -6.394523932 |
| **O15541** | RNF113A | -0.056317413 | 3.168120746 | -1.122309776 | 0.30163851 | 0.691344519 | -6.394969812 |
| **Q10589** | BST2 | -0.07579076 | 3.118953652 | -0.311702979 | 0.765066506 | 0.940369694 | -6.395296246 |
| **O95059** | RPP14 | 0.038619156 | 3.196733552 | 0.638514209 | 0.545062115 | 0.84497922 | -6.395707652 |
| **Q9H6P5** | TASP1 | -0.024225667 | 3.175892353 | -0.628034045 | 0.551489671 | 0.847559559 | -6.396138042 |
| **P49189** | ALDH9A1 | 0.056960889 | 3.164812855 | 1.703312814 | 0.13578424 | 0.524710986 | -6.39638297 |
| **Q9BV94-1** | EDEM2; MMP24-AS | 0.018190668 | 3.206932052 | 0.413789551 | 0.692394856 | 0.909991705 | -6.396411334 |
| **Q96Q07-1** | BTBD9 | 0.01711744 | 3.205322594 | 0.439106259 | 0.674869718 | 0.90314827 | -6.396612745 |
| **Q09328** | MGAT5 | 0.023341387 | 3.199089135 | 0.827110693 | 0.437604733 | 0.786862931 | -6.396832227 |
| **Q32P28-3** | LEPRE1; P3H1 | 0.029503543 | 3.18973253 | 1.204556029 | 0.270572942 | 0.671731232 | -6.397119269 |
| **Q8ND71** | GIMAP8 | -0.030573523 | 3.180733099 | -0.708824637 | 0.503159015 | 0.827688992 | -6.398584127 |
| **Q15043** | SLC39A14 | 0.046077383 | 3.239267889 | 0.879366098 | 0.410664781 | 0.769906516 | -6.398744627 |
| **Q13162** | PRDX4 | 0.031154746 | 3.180658659 | 1.49776984 | 0.181295006 | 0.58362594 | -6.398783451 |
| **Q86VR2** | FAM134C; RETREG | -0.015065559 | 3.172505208 | -0.230960551 | 0.824476019 | 0.959077751 | -6.399454329 |
| **Q96HD9** | ACY3 | 0.027450325 | 3.189448684 | 0.76523099 | 0.471113338 | 0.807027916 | -6.400168111 |
| **P35658-1** | NUP214 | 0.013829086 | 3.178883738 | 1.907940525 | 0.101465969 | 0.464906068 | -6.400737983 |
| **Q9Y5S1** | TRPV2 | -0.027664523 | 3.182993381 | -0.644737829 | 0.541267133 | 0.843540088 | -6.401125285 |
| **Q9C099-1** | LRRCC1 | 0.02302853 | 3.207071688 | 1.118131571 | 0.303293018 | 0.692399426 | -6.401659253 |
| **Q9P2J9** | PDP2 | 0.008602302 | 3.216917284 | 0.102974516 | 0.9211009 | 0.984164603 | -6.401660716 |
| **Q86X76-2** | NIT1 | 0.039959637 | 3.193785792 | 1.205124677 | 0.270367956 | 0.671731232 | -6.401959533 |
| **Q27J81** | INF2 | -0.023889442 | 3.19791168 | -1.280388139 | 0.244387127 | 0.650150213 | -6.402270671 |
| **Q8TE04-1** | PANK1 | -0.02066035 | 3.187369022 | -0.726080019 | 0.493205435 | 0.821156631 | -6.402529274 |
| **Q9ULT0-4** | TTC7A | 0.023277047 | 3.15159078 | 1.342360891 | 0.224655474 | 0.630229362 | -6.402590699 |
| **Q14680** | MELK | -0.023018251 | 3.161861671 | -1.309001809 | 0.235094916 | 0.64164013 | -6.402672523 |
| **Q9Y624** | F11R | 0.023947026 | 3.172068222 | 0.571545229 | 0.586920885 | 0.866100748 | -6.402938587 |
| **O75928-1** | PIAS2 | -0.016603309 | 3.190876701 | -0.308128303 | 0.76766404 | 0.941217716 | -6.403084239 |
| **Q5W0Z9-1** | ZDHHC20 | 0.010421482 | 3.176810507 | 0.163081687 | 0.875428932 | 0.972759481 | -6.403098265 |

| | | | | | | | |
|---|---|---|---|---|---|---|---|
| **P62249** | RPS16 | -0.043636402 | 3.201687071 | -1.645011201 | 0.147457406 | 0.546231571 | -6.403227617 |
| **Q9H0D6-1** | XRN2 | 0.028378451 | 3.167459435 | 1.855308529 | 0.109383333 | 0.479873319 | -6.403567374 |
| **P09668** | CTSH | 0.03870007 | 3.184226935 | 1.310085285 | 0.23474924 | 0.64164013 | -6.403712371 |
| **Q5T5C0-1** | STXBP5 | 0.047368074 | 3.195008811 | 0.753259203 | 0.47779594 | 0.811322085 | -6.403796073 |
| **P83916** | CBX1 | 0.041678874 | 3.154076696 | 1.55444567 | 0.167492208 | 0.566424749 | -6.403846416 |
| **Q8N653** | LZTR1 | -0.021530737 | 3.181594765 | -0.719530849 | 0.496967732 | 0.823442885 | -6.404213791 |
| **O95149** | SNUPN | -0.044743675 | 3.212782139 | -1.144656485 | 0.292916166 | 0.68530312 | -6.405054852 |
| **Q13948** | CUX1 | -0.019577024 | 3.179292181 | -0.511830522 | 0.625764794 | 0.885387512 | -6.405254253 |
| **P10914** | IRF1 | -0.057257124 | 3.196387592 | -0.940515665 | 0.380722046 | 0.751698969 | -6.405426766 |
| **Q96AJ9-2** | VTI1A | 0.017226702 | 3.172570444 | 0.324175042 | 0.756029671 | 0.937424764 | -6.405942708 |
| **Q9BQ52-1** | ELAC2 | 0.029288907 | 3.185592765 | 1.683960446 | 0.139557591 | 0.533923294 | -6.407128025 |
| **P15812-1** | CD1E | -0.032549013 | 3.179709329 | -0.692515434 | 0.5126875 | 0.831227449 | -6.407341875 |
| **Q8TB52** | FBXO30 | 0.027125838 | 3.200612899 | 1.116082797 | 0.30410704 | 0.692697946 | -6.4075421 |
| **O75817** | POP7 | -0.007847217 | 3.253184964 | -0.173213117 | 0.867776083 | 0.970868803 | -6.408401468 |
| **Q9H6H4-1** | REEP4 | -0.022416571 | 3.202029611 | -0.562124637 | 0.592956049 | 0.869935985 | -6.40874271 |
| **Q7L9L4-2** | MOB1B | -0.029546022 | 3.19144554 | -1.228756673 | 0.261965602 | 0.665595366 | -6.40883413 |
| **O00299** | CLIC1 | -0.028471177 | 3.171666483 | -1.671031158 | 0.142134159 | 0.538432214 | -6.409225115 |
| **Q8IXU6-1** | SLC35F2 | 0.00260557 | 3.145590026 | 0.031344533 | 0.975939371 | 0.99316605 | -6.410422518 |
| **Q9H299** | SH3BGRL3 | -0.012934225 | 3.191733676 | -0.363925141 | 0.727510893 | 0.92710315 | -6.41044439 |
| **Q15418-1** | RPS6KA1 | 0.034575828 | 3.188350538 | 1.425290242 | 0.200472244 | 0.606070006 | -6.41073745 |
| **P19397** | CD53 | 0.041861792 | 3.169400213 | 0.329396105 | 0.752258884 | 0.935759643 | -6.411076201 |
| **O43572** | AKAP10 | -0.028750473 | 3.185404028 | -1.103063061 | 0.309322298 | 0.697181747 | -6.411121775 |
| **P55211-1** | CASP9 | 0.013340209 | 3.198583684 | 0.277774101 | 0.789848836 | 0.947865984 | -6.411166236 |
| **O15344-1** | MID1 | 0.0349811 | 3.203976412 | 0.96147771 | 0.370849375 | 0.744717822 | -6.411612808 |
| **Q13064** | MKRN3 | 0.009546349 | 3.211922425 | 0.215584663 | 0.835947332 | 0.960912428 | -6.411620205 |
| **Q15061** | WDR43 | 0.025998066 | 3.199760852 | 1.663589232 | 0.143637691 | 0.539320158 | -6.411953965 |
| **O75071** | EFCAB14; KIAA049 | 0.019978621 | 3.166987809 | 0.510753864 | 0.626477738 | 0.885387512 | -6.412999792 |
| **Q92766-2** | RREB1 | 0.02008341 | 3.197561179 | 1.415759708 | 0.203126894 | 0.608695713 | -6.413012676 |
| **Q8NFU3-1** | TSTD1 | -0.017492923 | 3.215600021 | -0.224407069 | 0.829359775 | 0.959415173 | -6.413072706 |
| **O95758-4** | PTBP3 | 0.026358346 | 3.168151916 | 0.932166293 | 0.384709992 | 0.754667212 | -6.413665627 |
| **O60613-1** | SEP15; SELENOF | 0.036632539 | 3.153023632 | 0.305000961 | 0.769939192 | 0.941649022 | -6.413979799 |
| **Q9GZU3** | TMEM39B | -0.016233425 | 3.201018493 | -0.298945784 | 0.77435136 | 0.943158294 | -6.415038641 |
| **Q9NUX5-1** | POT1 | -0.027976025 | 3.173948527 | -1.293969046 | 0.239937402 | 0.645958461 | -6.41548092 |
| **Q9UKG9-1** | CROT | -0.004581212 | 3.189010225 | -0.088824232 | 0.931907381 | 0.986711787 | -6.415745305 |
| **P07741-1** | APRT | -0.055290749 | 3.179326409 | -1.221266411 | 0.264604212 | 0.66823064 | -6.415862208 |
| **Q8N1K5-4** | THEMIS | -0.037350505 | 3.198436634 | -1.759247177 | 0.125420687 | 0.508921022 | -6.416744963 |
| **Q9P2K2** | TXNDC16 | 0.007973954 | 3.159387558 | 0.178149924 | 0.864052563 | 0.969738239 | -6.416844359 |
| **Q5TZA2** | CROCC | -0.0216252 | 3.184827902 | -1.30312411 | 0.236977976 | 0.643518711 | -6.416999427 |
| **P31153** | MAT2A | 0.057091622 | 3.19697472 | 1.297770611 | 0.238704612 | 0.644491512 | -6.417026858 |

| | | | | | | | |
|---|---|---|---|---|---|---|---|
| **Q7Z406-2** | MYH14 | 0.004567268 | 3.187076199 | 0.083613627 | 0.935890766 | 0.986946129 | -6.417412694 |
| **Q13620** | CUL4B | 0.026419026 | 3.168571073 | 1.716923492 | 0.133189152 | 0.520505564 | -6.417437799 |
| **Q13686** | ALKBH1 | 0.013022923 | 3.19727439 | 0.175406279 | 0.866121463 | 0.970432241 | -6.417867836 |
| **Q99653** | CHP1 | 0.021691723 | 3.191393779 | 0.5183689 | 0.621444574 | 0.883385606 | -6.417959205 |
| **O75865-2** | TRAPPC6A | 0.022696848 | 3.175732116 | 0.49557569 | 0.636574505 | 0.88885491 | -6.418001889 |
| **Q9BVJ7** | DUSP23 | -0.017912319 | 3.197296732 | -0.413771397 | 0.692407498 | 0.909991705 | -6.418642972 |
| **Q13541** | EIF4EBP1 | 0.028064235 | 3.18566867 | 0.111381994 | 0.914688594 | 0.982718171 | -6.418714987 |
| **Q02108** | GUCY1A3 | 0.025048028 | 3.202256619 | 0.788905481 | 0.458088427 | 0.799573893 | -6.418725352 |
| **Q9NPA0** | EMC7 | 0.029038656 | 3.195529596 | 0.900888899 | 0.399931349 | 0.763260765 | -6.418939607 |
| **P32856** | STX2 | -0.012437366 | 3.141809451 | -0.244731525 | 0.814241499 | 0.95620584 | -6.419013355 |
| **P50552** | VASP | -0.031514282 | 3.187401261 | -1.215791729 | 0.26654717 | 0.669528065 | -6.419033946 |
| **Q5SW79-2** | CEP170 | 0.007252579 | 3.186605863 | 0.101361273 | 0.922332054 | 0.984164603 | -6.419109204 |
| **Q9H0R4-1** | HDHD2 | 0.025081465 | 3.183373801 | 1.190717689 | 0.27560235 | 0.674825663 | -6.419482935 |
| **Q8TF50** | ZNF526 | 0.012155598 | 3.191623711 | 0.280222394 | 0.788051239 | 0.947802649 | -6.41975179 |
| **P17480-1** | UBTF | -0.025004601 | 3.14598323 | -0.561191387 | 0.593555848 | 0.869935985 | -6.419936643 |
| **Q08J23** | NSUN2 | 0.012098583 | 3.205141106 | 0.276276424 | 0.790949156 | 0.948368382 | -6.420239328 |
| **Q9BTA9-1** | WAC | 0.032352599 | 3.178799199 | 1.364694315 | 0.21789829 | 0.624306497 | -6.42058986 |
| **O15145** | ARPC3 | 0.200119279 | 3.161360481 | 1.504564113 | 0.179586493 | 0.580990067 | -6.420866832 |
| **Q8NCJ5** | SPRYD3 | 0.01348612 | 3.180001321 | 0.323574462 | 0.756463893 | 0.937613982 | -6.420951446 |
| **P20339** | RAB5A | 0.025341477 | 3.160837604 | 0.656997614 | 0.533839754 | 0.840132297 | -6.421690271 |
| **O43909** | EXTL3 | -0.009341029 | 3.169853277 | -0.193092088 | 0.852805963 | 0.96682982 | -6.422180924 |
| **Q7L945** | ZNF627 | 0.013847304 | 3.130037675 | 0.251608923 | 0.809144868 | 0.955498404 | -6.423079662 |
| **Q9NYJ8-1** | TAB2 | 0.013081048 | 3.177219402 | 0.373927078 | 0.720406265 | 0.925644897 | -6.423169103 |
| **Q6GPH4-1** | XAF1 | -0.024999744 | 3.210828632 | -0.712271527 | 0.501160189 | 0.826425425 | -6.423655194 |
| **P20290-1** | BTF3 | -0.026707591 | 3.236860643 | -1.417423442 | 0.202661194 | 0.608695713 | -6.424103333 |
| **O00562** | PITPNM1 | 0.019402929 | 3.175348703 | 0.670232905 | 0.525893763 | 0.838233243 | -6.424554329 |
| **Q9Y217-1** | MTMR6 | 0.033305648 | 3.190337302 | 1.224909827 | 0.263317904 | 0.666686026 | -6.425078639 |
| **Q8N5S9-2** | CAMKK1 | 0.02936105 | 3.235217154 | 0.703851283 | 0.506052257 | 0.82913633 | -6.425181163 |
| **Q15366-2** | PCBP2 | -0.037954514 | 3.182880028 | -1.315599784 | 0.232996797 | 0.640096665 | -6.426749276 |
| **O94766** | B3GAT3 | 0.021591455 | 3.168010984 | 0.665369449 | 0.528804836 | 0.839849063 | -6.427153954 |
| **Q8NFZ0-2** | FBXO18 | 0.020320286 | 3.199899727 | 0.386950803 | 0.711200192 | 0.921051693 | -6.42776632 |
| **Q5T8D3-3** | ACBD5 | 0.023103579 | 3.179925152 | 1.04513495 | 0.333420573 | 0.719098069 | -6.428083534 |
| **Q8TDX7-1** | NEK7 | 0.029712522 | 3.177583216 | 1.444019576 | 0.195346864 | 0.601096374 | -6.428581836 |
| **Q9P2H3-3** | TRIM59; IFT80 | 0.028923995 | 3.163533203 | 0.965036573 | 0.369193007 | 0.743721714 | -6.429012486 |
| **Q9NVP1** | DDX18 | -0.057579462 | 3.179548552 | -1.830438481 | 0.11333242 | 0.487243547 | -6.429627945 |
| **Q96CN4-2** | EVI5L | -0.013897942 | 3.16633904 | -0.347047589 | 0.739565578 | 0.931196278 | -6.429971663 |
| **Q9Y4L5** | RNF115; LOC1010( | -0.003341143 | 3.176338148 | -0.07314372 | 0.943900691 | 0.98895887 | -6.429993034 |
| **P57071** | PRDM15 | -0.000819488 | 3.240981521 | -0.018585739 | 0.985731503 | 0.995531556 | -6.430012522 |
| **O14562** | UBFD1 | -0.024580916 | 3.195937924 | -1.265215962 | 0.249443225 | 0.65453932 | -6.430395624 |

| | | | | | | | |
|---|---|---|---|---|---|---|---|
| **Q6PFW1-1** | PPIP5K1 | 0.010953905 | 3.189453152 | 0.27598619 | 0.791162447 | 0.948368382 | -6.43135525 |
| **Q92930** | RAB8B | 0.02302833 | 3.149343575 | 0.439602389 | 0.674528417 | 0.90314827 | -6.431637371 |
| **Q8ND76** | CCNY | 0.024816331 | 3.168014023 | 0.526730384 | 0.615943364 | 0.880583268 | -6.432820325 |
| **Q99683** | MAP3K5 | 0.013077305 | 3.231069333 | 0.244028535 | 0.81476302 | 0.95620584 | -6.433040102 |
| **O00401** | WASL; ASB15 | 0.021642251 | 3.19260468 | 0.556331386 | 0.596684951 | 0.871378088 | -6.433603302 |
| **P62913** | RPL11 | -0.035043885 | 3.186563368 | -1.60026674 | 0.157055765 | 0.555457754 | -6.433937654 |
| **Q8IYL3** | C1orf174 | 0.008678985 | 3.201893687 | 0.167757581 | 0.871895101 | 0.971931713 | -6.433993835 |
| **P55771** | PAX9 | -0.003488976 | 3.236875686 | -0.064058162 | 0.950857302 | 0.989472196 | -6.434010416 |
| **P42330** | AKR1C3 | -0.011760966 | 3.1989103 | -0.125531971 | 0.903912414 | 0.980210603 | -6.434447042 |
| **P55265-1** | ADAR | 0.026052906 | 3.176659523 | 0.916773055 | 0.392145496 | 0.759806823 | -6.434662108 |
| **Q96BN2** | TADA1 | -0.028669453 | 3.196922155 | -1.006474666 | 0.350327595 | 0.731407657 | -6.435288344 |
| **P40616** | ARL 1.00 | -0.025414707 | 3.178830606 | -0.957747769 | 0.372591522 | 0.745276848 | -6.435787581 |
| **Q9NWX5** | ASB6 | -0.040990531 | 3.192864904 | -1.113142464 | 0.305278451 | 0.693599527 | -6.435906034 |
| **Q96Q15-1** | SMG1 | 0.016441839 | 3.188575694 | 1.756742929 | 0.125867911 | 0.509142907 | -6.43602075 |
| **Q06136** | KDSR | 0.014031106 | 3.176014131 | 0.241935267 | 0.816316543 | 0.956409905 | -6.436091546 |
| **Q86VK4-5** | ZNF410 | -0.024036186 | 3.198572953 | -0.685388639 | 0.516887851 | 0.833280927 | -6.436342014 |
| **P42568** | MLLT3 | -0.021699851 | 3.168018364 | -0.360152336 | 0.730198461 | 0.927974736 | -6.436376962 |
| **O75882-1** | ATRN | 0.022841882 | 3.153289293 | 1.076191345 | 0.320318409 | 0.708257023 | -6.436462241 |
| **Q8TBC3** | SHKBP1 | 0.020149077 | 3.219042283 | 0.426029936 | 0.683895097 | 0.905978473 | -6.436506085 |
| **Q9BV19** | C1orf50 | -0.027451819 | 3.222775952 | -1.168364441 | 0.283893768 | 0.679787821 | -6.437356072 |
| **Q5T653** | MRPL2 | 0.094068352 | 3.160996307 | 1.009901274 | 0.348802127 | 0.730176717 | -6.437382696 |
| **P60604** | UBE2G2 | -0.020955945 | 3.242027342 | -0.936371372 | 0.38269754 | 0.754100187 | -6.437542116 |
| **Q9Y3Y2-3** | CHTOP | -0.058385935 | 3.120794786 | -0.773003056 | 0.466809519 | 0.804563762 | -6.437549193 |
| **Q9BXS5-2** | AP1M1 | -0.035061155 | 3.184837479 | -1.614465271 | 0.153947849 | 0.552139704 | -6.437808846 |
| **O15037** | KHNYN | -0.030327766 | 3.19063582 | -0.647947001 | 0.53931671 | 0.843540088 | -6.438222943 |
| **Q96L58** | B3GALT6 | -0.029467104 | 3.190358353 | -0.579444392 | 0.581887834 | 0.864278602 | -6.438295739 |
| **Q9NVA2-2** | 11-Sep | -0.030426459 | 3.188023212 | -0.90350381 | 0.398641698 | 0.762880385 | -6.438741409 |
| **Q2KHT3** | CLEC16A | 0.02111566 | 3.181636383 | 0.763934358 | 0.471834002 | 0.807565951 | -6.438838577 |
| **Q16526** | CRY1 | 0.014589752 | 3.204948231 | 0.318430186 | 0.760187137 | 0.938422437 | -6.439397302 |
| **Q9UGL1-2** | KDM5B | -0.048431772 | 3.054008627 | -1.249115472 | 0.254907825 | 0.659782078 | -6.439624309 |
| **P07766** | CD3E | -0.029605819 | 3.202904049 | -1.338750611 | 0.225765122 | 0.631141413 | -6.439648799 |
| **Q9NVP2** | ASF1B | 0.025561814 | 3.169791503 | 0.684770821 | 0.517253022 | 0.833280927 | -6.439710114 |
| **Q8TBF2-8** | FAM213B | 0.035980621 | 3.175777269 | 1.059498533 | 0.327308054 | 0.712260338 | -6.439806607 |
| **Q14164** | IKBKE | 0.020720448 | 3.167356319 | 0.683026347 | 0.518285016 | 0.833726353 | -6.440037368 |
| **Q96BY6-1** | DOCK10 | 0.022231269 | 3.181531535 | 1.378572586 | 0.213791257 | 0.619843797 | -6.440299493 |
| **Q12846** | STX4 | 0.028652612 | 3.129507388 | 0.940589551 | 0.380686896 | 0.751698969 | -6.440832469 |
| **Q8NBJ5** | COLGALT1; GLT25I | -0.055625227 | 3.193271677 | -1.466199932 | 0.189431641 | 0.59346782 | -6.44130396 |
| **Q9Y285** | FARSA | 0.024385801 | 3.197646042 | 1.615442392 | 0.153736113 | 0.552139704 | -6.441358596 |
| **Q7Z6I8-1** | C5orf24 | 0.032440196 | 3.222651461 | 0.424555017 | 0.684916676 | 0.905978473 | -6.442780899 |

| | | | | | | | |
|---|---|---|---|---|---|---|---|
| **Q9H1A3** | METTL9 | 0.00862653 | 3.183561204 | 0.158036671 | 0.879245203 | 0.97338977 | -6.44296727 |
| **Q9ULM3** | YEATS2 | 0.024635638 | 3.153479268 | 1.539134308 | 0.171121651 | 0.571004416 | -6.443559227 |
| **O15318** | POLR3G | -0.013184881 | 3.210507013 | -0.353913549 | 0.734651739 | 0.92925216 | -6.443609689 |
| **P16298-4** | PPP3CB | -0.014754222 | 3.174694949 | -0.380039974 | 0.716078833 | 0.924439954 | -6.443877769 |
| **Q13216-1** | ERCC8 | -0.009745545 | 3.194229354 | -0.242205709 | 0.816115783 | 0.956348067 | -6.443922485 |
| **Q9H2K8** | TAOK3 | -0.050422014 | 3.180253119 | -1.397839239 | 0.208204714 | 0.614231901 | -6.444336329 |
| **Q9Y2U8** | LEMD3 | 0.045080008 | 3.193378006 | 1.377064239 | 0.214234244 | 0.619843797 | -6.444889201 |
| **Q13813-3** | SPTAN1 | 0.034659959 | 3.138995127 | 0.547489842 | 0.602401473 | 0.872497827 | -6.445399627 |
| **Q9Y3C4-3** | TPRKB | -0.053894167 | 3.15251706 | -0.982284733 | 0.361246486 | 0.736919727 | -6.446022763 |
| **Q9Y2Y9** | KLF13 | 0.005816728 | 3.189108556 | 0.139235741 | 0.893497019 | 0.978276697 | -6.446147732 |
| **O14777** | NDC80 | -0.02605619 | 3.159233387 | -1.645184591 | 0.147421314 | 0.546231571 | -6.447278093 |
| **P29597** | TYK2 | -0.014392123 | 3.157199007 | -0.422946492 | 0.686031615 | 0.906701749 | -6.447435786 |
| **Q9Y6K0** | CEPT1 | 0.004941438 | 3.191466466 | 0.132290936 | 0.89877262 | 0.978757548 | -6.447906336 |
| **Q9H7C9-1** | AAMDC | 0.01995355 | 3.198783795 | 1.154834937 | 0.289013609 | 0.683685579 | -6.448343415 |
| **Q96AY2-2** | EME1 | 0.013838233 | 3.136083887 | 0.258595996 | 0.803977278 | 0.95391847 | -6.448839939 |
| **Q8TEB1** | DCAF11 | 0.033721599 | 3.178135285 | 1.08511592 | 0.316631548 | 0.703974147 | -6.448866884 |
| **O94822-3** | LTN1 | -0.028781418 | 3.195548809 | -1.611107123 | 0.154677637 | 0.553156191 | -6.448961003 |
| **O15525** | MAFG | 0.016559196 | 3.176526564 | 0.457057308 | 0.662574334 | 0.897184895 | -6.449191738 |
| **Q96C23** | GALM | 0.045438392 | 3.194594284 | 1.213944642 | 0.267205448 | 0.67010832 | -6.449416157 |
| **P62875** | POLR2L | -0.037336127 | 3.217904085 | -0.972064181 | 0.365939044 | 0.740729864 | -6.449759627 |
| **O14715** | RGPD8 | -0.010652629 | 3.208418796 | -0.147259255 | 0.887409303 | 0.975708125 | -6.449831025 |
| **O75791** | GRAP2 | -0.025708775 | 3.179063732 | -1.465794917 | 0.18953817 | 0.593538505 | -6.450258236 |
| **P52655-1** | GTF2A1 | -0.026506206 | 3.186624619 | -1.034291883 | 0.338095374 | 0.721894046 | -6.450928 |
| **Q9NQ34** | TMEM9B | -0.02112394 | 3.196358757 | -0.526854636 | 0.615861817 | 0.880583268 | -6.451015283 |
| **Q96BZ9-3** | TBC1D20 | -0.01972458 | 3.213452143 | -0.461136574 | 0.659795833 | 0.896852471 | -6.451407912 |
| **Q0VD83-4** | APOBR | 0.029234004 | 3.208490963 | 0.812982135 | 0.44510246 | 0.791127304 | -6.453710047 |
| **Q5VYV7** | SLX4IP | -0.024949928 | 3.153491024 | -0.551947259 | 0.599515675 | 0.872097298 | -6.453773326 |
| **P18074-1** | ERCC2 | -0.030251604 | 3.174835956 | -1.364764282 | 0.217877409 | 0.624306497 | -6.454148494 |
| **Q92508** | PIEZO1 | 0.011671408 | 3.180739363 | 0.182312774 | 0.860915708 | 0.968855822 | -6.454356347 |
| **Q96B23-1** | C18orf25 | 0.014882283 | 3.212309634 | 0.313066224 | 0.764076771 | 0.940369694 | -6.454605251 |
| **Q9H490** | PIGU | 0.002859842 | 3.18599979 | 0.054846186 | 0.957915477 | 0.989679905 | -6.454957752 |
| **Q9H5V9-1** | CXorf56 | -0.033253563 | 3.168001671 | -1.218843202 | 0.265462705 | 0.668347222 | -6.455403356 |
| **P36873-2** | PPP1CC | -0.032812876 | 3.222358948 | -0.609707552 | 0.562840253 | 0.852576894 | -6.456191904 |
| **P29992** | GNA11 | 0.000307031 | 3.209040905 | 0.004485368 | 0.996556313 | 0.998480498 | -6.457159578 |
| **Q8IY17-4** | PNPLA6 | -0.025224569 | 3.182184942 | -1.005213333 | 0.350890444 | 0.73209679 | -6.457505509 |
| **O95486-1** | SEC24A | 0.030687572 | 3.200196715 | 1.158454538 | 0.287636342 | 0.682522792 | -6.457786911 |
| **Q9UBL0-3** | ARPP21 | -0.006118885 | 3.185849293 | -0.128703629 | 0.901499914 | 0.97952714 | -6.457826955 |
| **P17050** | NAGA | 0.021236794 | 3.170704903 | 0.70032698 | 0.508109105 | 0.82913633 | -6.457976624 |
| **Q8N9N5-1** | BANP | -0.054489139 | 3.173736965 | -1.073728751 | 0.321341867 | 0.709183647 | -6.458846394 |

| | | | | | | | |
|---|---|---|---|---|---|---|---|
| **Q9Y259-1** | CHKB | 0.004878164 | 3.184730951 | 0.108629987 | 0.916786788 | 0.983235402 | -6.459071163 |
| **P20248** | CCNA2 | 0.041609965 | 3.134649763 | 1.351716488 | 0.221802472 | 0.627993355 | -6.459325435 |
| **P08133-1** | ANXA6 | 0.043258329 | 3.183441545 | 2.214094275 | 0.065517484 | 0.394047248 | -6.459330715 |
| **Q9HA65** | TBC1D17 | 0.013959604 | 3.177141153 | 0.314432061 | 0.763085635 | 0.939644172 | -6.459471021 |
| **Q8TEV9** | SMCR8 | 0.006204732 | 3.186979111 | 0.127701122 | 0.902262343 | 0.979567411 | -6.460246727 |
| **Q9UNX4** | WDR3 | -0.045945861 | 3.188703464 | -1.645639069 | 0.147326754 | 0.546231571 | -6.46071988 |
| **Q9NWF9-1** | RNF216 | 0.008809946 | 3.176964145 | 0.23253853 | 0.823301339 | 0.959068023 | -6.461301762 |
| **Q7Z434** | MAVS | 0.002389252 | 3.18180666 | 0.035789823 | 0.972528664 | 0.992928787 | -6.461645085 |
| **P34910-2** | EVI2B | 0.007495838 | 3.194744841 | 0.182165835 | 0.861026387 | 0.968855822 | -6.462610197 |
| **O60344-4** | EEF1AKMT4; ECE2 | -0.015093594 | 3.226049714 | -0.386959632 | 0.711193969 | 0.921051693 | -6.462946666 |
| **Q99719-1** | 5-Sep | -0.024768547 | 3.143337508 | -0.484552958 | 0.643960273 | 0.891474831 | -6.463023565 |
| **P23469** | PTPRE | 0.009550182 | 3.195921834 | 0.260740981 | 0.802392988 | 0.953157309 | -6.463049192 |
| **O14936** | CASK | 0.00467063 | 3.221426102 | 0.105644914 | 0.919063494 | 0.983742622 | -6.463413671 |
| **Q9HCU9** | BRMS1 | -0.031624497 | 3.194873919 | -1.086179575 | 0.316194455 | 0.703753313 | -6.463579806 |
| **P23468-2** | PTPRD | 0.029915153 | 3.164442045 | 0.963619874 | 0.369851681 | 0.744424726 | -6.46388939 |
| **Q8N697** | SLC15A4 | -0.033918236 | 3.212625459 | -0.509235941 | 0.62748362 | 0.885387512 | -6.464170479 |
| **Q9BSY4** | CHCHD5 | -0.014375319 | 3.203321077 | -0.177971303 | 0.864187221 | 0.969738239 | -6.464265073 |
| **Q76L83** | ASXL2 | 0.041728832 | 3.186044957 | 0.99789424 | 0.354170548 | 0.734057823 | -6.464902611 |
| **Q8TBB5** | KLHDC4 | -0.026227639 | 3.197673859 | -0.939269101 | 0.381315434 | 0.752683509 | -6.464938271 |
| **P52630-3** | STAT2 | -0.029753596 | 3.160733337 | -1.197115102 | 0.273267482 | 0.672884265 | -6.46497471 |
| **Q14457** | BECN1 | -0.029848281 | 3.162709407 | -0.757294651 | 0.475536177 | 0.809713018 | -6.465089716 |
| **O95985** | TOP3B | -0.022899983 | 3.182900354 | -1.527202564 | 0.174000539 | 0.574454961 | -6.465107904 |
| **Q9BQ04** | RBM4B | -0.037994057 | 3.16306585 | -0.356964047 | 0.732472875 | 0.928164811 | -6.465372617 |
| **Q96H79** | ZC3HAV1L | -0.038390085 | 3.216636682 | -1.113117152 | 0.305288552 | 0.693599527 | -6.466537751 |
| **Q99708-2** | RBBP8 | 0.030592454 | 3.196980225 | 0.97906271 | 0.362720723 | 0.738054037 | -6.467559482 |
| **O94966-6** | USP19 | 0.035233711 | 3.159348923 | 0.700550792 | 0.507978322 | 0.82913633 | -6.468526108 |
| **Q9H9V9** | JMJD4 | 0.010835467 | 3.230448819 | 0.115659546 | 0.911428766 | 0.981900848 | -6.468835702 |
| **Q13085-4** | ACACA | -0.039236564 | 3.195721022 | -1.895993746 | 0.103212087 | 0.469656376 | -6.469536912 |
| **O95297-3** | MPZL1 | 0.016896018 | 3.188863206 | 0.143693585 | 0.890113704 | 0.977017047 | -6.469690532 |
| **Q92674** | CENPI | -0.026157112 | 3.170003756 | -0.769084145 | 0.468976234 | 0.805725242 | -6.470276071 |
| **P10074** | ZBTB48 | -0.027870824 | 3.190597271 | -0.616290847 | 0.558746725 | 0.851363273 | -6.470840253 |
| **Q8WUA2** | PPIL4 | -0.033358855 | 3.185236758 | -1.398324706 | 0.20806566 | 0.614231901 | -6.47091748 |
| **P36871-1** | PGM1 | -0.031347739 | 3.163196354 | -1.831607491 | 0.11314371 | 0.487223187 | -6.472401548 |
| **Q8N8N7-1** | PTGR2 | -0.046985574 | 3.179745167 | -0.658843122 | 0.532727255 | 0.839951983 | -6.472455179 |
| **Q2NKJ3-1** | CTC1 | -0.041967821 | 3.150914025 | -0.772040147 | 0.467341258 | 0.805079422 | -6.472490602 |
| **Q13158** | FADD | -0.009586263 | 3.210387441 | -0.215391862 | 0.836091455 | 0.960912428 | -6.473021285 |
| **P11233** | RALA | 0.038621411 | 3.144573637 | 0.815204714 | 0.443916964 | 0.790506794 | -6.473290742 |
| **Q53GS9** | USP39 | -0.019609575 | 3.179171496 | -1.633837529 | 0.149800972 | 0.546701296 | -6.47369997 |
| **Q92499** | DDX1 | 0.017841102 | 3.166981368 | 1.949894955 | 0.095562121 | 0.452536799 | -6.473809476 |

| | | | | | | | |
|---|---|---|---|---|---|---|---|
| **Q13642-4** | FHL1 | 0.022838338 | 3.202990606 | 0.749164759 | 0.480096204 | 0.81235548 | -6.473909645 |
| **Q9UIF7-1** | MUTYH | -0.02860557 | 3.19928714 | -0.998929321 | 0.353705209 | 0.733449655 | -6.473934148 |
| **P23610** | F8A2; F8A1; F8A3 | -0.035355163 | 3.181252758 | -0.697794358 | 0.509590559 | 0.829141305 | -6.474131702 |
| **Q7Z5U6** | WDR53 | -0.018285316 | 3.200822719 | -0.54234603 | 0.605741323 | 0.87454385 | -6.474622518 |
| **Q96BH1** | RNF25 | -0.059196965 | 3.208890869 | -1.125167933 | 0.300511031 | 0.691292041 | -6.475001283 |
| **Q6P1K2** | PMF1; PMF1-BGLA | -0.027388205 | 3.216394247 | -1.057807467 | 0.328022972 | 0.713117775 | -6.475252958 |
| **Q96RY5** | CRAMP1L; CRAMF | 0.014045469 | 3.187565963 | 0.336264054 | 0.747309892 | 0.934974996 | -6.475253164 |
| **Q8IV08** | PLD3 | 0.006358574 | 3.149388337 | 0.182547649 | 0.860738803 | 0.968855822 | -6.475394838 |
| **Q02878** | RPL6 | -0.103076687 | 3.154240604 | -1.948994402 | 0.095685201 | 0.452536799 | -6.47562089 |
| **Q9NRG1** | PRTFDC1 | -0.004914966 | 3.174377343 | -0.114449397 | 0.912350809 | 0.981932698 | -6.47675846 |
| **Q96H35** | RBM18 | -0.021816379 | 3.212325884 | -0.527103298 | 0.615698637 | 0.880583268 | -6.477295801 |
| **Q9UJV9** | DDX41 | -0.032113721 | 3.180982739 | -1.608296225 | 0.155291004 | 0.554148197 | -6.477791968 |
| **Q8WZA1-2** | POMGNT1 | 0.029657433 | 3.205457802 | 0.642226252 | 0.542796639 | 0.843669625 | -6.478085806 |
| **Q9BSJ5-1** | C17orf80 | -0.052906732 | 3.224593589 | -0.489220333 | 0.640827484 | 0.890430235 | -6.478456391 |
| **Q9NUM4** | TMEM106B | 0.011275899 | 3.1793351 | 0.292406164 | 0.779126742 | 0.944053983 | -6.478479259 |
| **Q9C0B5** | ZDHHC5 | -0.04881461 | 3.178458213 | -0.853338978 | 0.423927108 | 0.778391698 | -6.478644507 |
| **P35670** | ATP7B | -0.050243581 | 2.971635441 | -0.420742648 | 0.687560578 | 0.907431059 | -6.478883968 |
| **O14832** | PHYH | 0.005407963 | 3.157154216 | 0.166187876 | 0.873081063 | 0.97237605 | -6.479036878 |
| **P17900** | GM2A | -0.040904562 | 3.179429475 | -0.992114536 | 0.356777762 | 0.734600528 | -6.479632614 |
| **P68371** | TUBB4B | 0.037865466 | 3.205928125 | 0.814414268 | 0.444338321 | 0.790653518 | -6.479898954 |
| **O60443** | DFNA5; GSDME | 0.030822934 | 3.185640227 | 0.985435512 | 0.359809372 | 0.73661838 | -6.481103584 |
| **Q9UK58** | CCNL1 | 0.030563069 | 3.199740935 | 1.002946092 | 0.351903955 | 0.732738778 | -6.481115084 |
| **P10253** | GAA | -0.022307301 | 3.193564986 | -0.972440089 | 0.365765617 | 0.740729864 | -6.481594969 |
| **Q9HAF1-3** | MEAF6 | 0.01649083 | 3.162751136 | 0.476062322 | 0.649679503 | 0.893646754 | -6.481980553 |
| **Q15363** | TMED2 | -0.02134362 | 3.207037302 | -1.273542843 | 0.246657148 | 0.652280071 | -6.4822775 |
| **P53667-1** | LIMK1 | -0.023658777 | 3.183696275 | -0.413385977 | 0.692675931 | 0.909991705 | -6.482496099 |
| **Q2TAY7** | SMU1 | -0.033922325 | 3.184761992 | -1.283950425 | 0.243213036 | 0.649360189 | -6.482725763 |
| **Q684P5** | RAP1GAP2 | -0.027509258 | 3.181049165 | -0.878199865 | 0.411252423 | 0.769906516 | -6.482929545 |
| **O75771** | RAD51D | 0.000580718 | 3.185030685 | 0.014052747 | 0.989211229 | 0.996219143 | -6.482990722 |
| **Q96EI5-2** | TCEAL4 | -0.098749399 | 3.159436857 | -1.293774158 | 0.240000752 | 0.645958461 | -6.48303883 |
| **Q9Y5Y0** | FLVCR1 | -0.033228545 | 3.186812248 | -0.350556082 | 0.737052941 | 0.929681266 | -6.483380631 |
| **Q14596-1** | NBR1 | -0.024898153 | 3.121409564 | -0.889429671 | 0.405619744 | 0.766336431 | -6.483831629 |
| **Q6ZVK8-1** | NUDT18 | 0.011025377 | 3.166887694 | 0.296372994 | 0.776228814 | 0.94357883 | -6.484090918 |
| **Q8NB91** | FANCB | 0.012610321 | 3.189603774 | 0.374353848 | 0.720103782 | 0.925644897 | -6.484452753 |
| **P62258-1** | YWHAE | 0.028435898 | 3.179318604 | 2.00795347 | 0.087950688 | 0.43768911 | -6.484728737 |
| **Q7RTR2-2** | NLRC3 | 0.000153806 | 3.193193865 | 0.003343567 | 0.997432938 | 0.998480498 | -6.484863348 |
| **Q49MG5** | MAP9 | -0.022376318 | 3.227770591 | -0.87139952 | 0.414691359 | 0.771959431 | -6.485173861 |
| **Q9BW83-1** | IFT27 | -0.037678791 | 3.181321174 | -0.994406597 | 0.355742021 | 0.734600528 | -6.486067132 |
| **O15254** | ACOX3 | 0.026618589 | 3.167935618 | 0.831259501 | 0.435420312 | 0.784298014 | -6.48698074 |

| | | | | | | | |
|---|---|---|---|---|---|---|---|
| **Q8IUH5-1** | ZDHHC17 | -0.025531798 | 3.17417511 | -0.759267328 | 0.474434181 | 0.809053352 | -6.487341035 |
| **Q9Y3D6** | FIS1 | -0.043286508 | 3.132838415 | -0.608063769 | 0.563865175 | 0.852578515 | -6.488022111 |
| **Q5SQN1-1** | SNAP47 | -0.04638204 | 3.216734757 | -0.64493304 | 0.541148365 | 0.843540088 | -6.488182332 |
| **Q96JB5-4** | CDK5RAP3 | 0.033624202 | 3.188284899 | 1.456482948 | 0.192002607 | 0.596505086 | -6.48930691 |
| **Q96EE3-1** | SEH1L | 0.0354164 | 3.195190602 | 1.503022122 | 0.17997294 | 0.581254067 | -6.489360713 |
| **Q6GTX8** | LAIR1 | -0.008654464 | 3.174974471 | -0.18013342 | 0.862557592 | 0.969309769 | -6.489744867 |
| **Q15397** | KIAA0020; PUM3 | -0.020714281 | 3.196474295 | -1.66555502 | 0.143239067 | 0.539313633 | -6.490549251 |
| **Q02156** | PRKCE | -0.012046688 | 3.181762035 | -0.273360189 | 0.793093157 | 0.949070183 | -6.490701383 |
| **Q7Z7C8-2** | TAF8 | -0.00725688 | 3.156944496 | -0.175563022 | 0.866003236 | 0.970432241 | -6.491096331 |
| **P13929-1** | ENO3 | -0.01651922 | 3.176000188 | -0.366648789 | 0.725573285 | 0.926648409 | -6.492211782 |
| **Q9UHR5-1** | SAP30BP | -0.033307166 | 3.190030154 | -1.19858826 | 0.27273221 | 0.672884265 | -6.492453238 |
| **P49585** | PCYT1A | -0.018784422 | 3.181843099 | -0.801014026 | 0.451524693 | 0.79513757 | -6.494098098 |
| **P35236-3** | PTPN7 | 0.026412807 | 3.211722724 | 0.751169496 | 0.478969001 | 0.811537935 | -6.49551905 |
| **Q8N5M4** | TTC9C | -0.017525318 | 3.159972392 | -0.557989522 | 0.595616312 | 0.871378088 | -6.495655563 |
| **Q9GZQ3** | COMMD5 | -0.023983297 | 3.162075756 | -0.517046439 | 0.622317085 | 0.883385606 | -6.49570914 |
| **P51608-2** | MECP2 | -0.034434379 | 3.172075177 | -1.081494128 | 0.318123566 | 0.706003277 | -6.495962318 |
| **Q00537** | CDK17 | 0.022201742 | 3.135260075 | 0.986435271 | 0.359354303 | 0.73661838 | -6.496121959 |
| **Q96C10** | DHX58 | 0.004285331 | 3.130936273 | 0.112283096 | 0.914001733 | 0.982219313 | -6.497445734 |
| **Q6N063-1** | OGFOD2 | -0.009697496 | 3.181386629 | -0.259497976 | 0.803310951 | 0.953721684 | -6.497633601 |
| **Q9UHC7** | MKRN1 | -0.014020773 | 3.181372814 | -0.312528157 | 0.764467358 | 0.940369694 | -6.497850617 |
| **P53582** | METAP1 | -0.028208353 | 3.191082583 | -1.431375769 | 0.198793665 | 0.603983046 | -6.49796383 |
| **O43543** | XRCC2 | -0.004924643 | 3.198008039 | -0.109444466 | 0.916165734 | 0.982975929 | -6.498164348 |
| **P30203-1** | CD6 | -0.012271707 | 3.195034108 | -0.363002203 | 0.728167966 | 0.927130528 | -6.498329996 |
| **Q96K80** | ZC3H10 | -0.017094398 | 3.21263994 | -0.454507928 | 0.664313735 | 0.89837832 | -6.498548541 |
| **Q96KC8** | DNAJC1 | 0.000936853 | 3.203010776 | 0.018328657 | 0.985928842 | 0.995531556 | -6.498658713 |
| **Q8N1G0** | ZNF687 | -0.02487022 | 3.179410072 | -0.709307322 | 0.502878794 | 0.827541843 | -6.498895921 |
| **O15446-2** | CD3EAP | -0.03796658 | 3.187106339 | -1.204874956 | 0.270457959 | 0.671731232 | -6.499328904 |
| **Q96CN5** | LRRC45 | 0.013765839 | 3.178096344 | 0.318655582 | 0.760023857 | 0.938422437 | -6.49958764 |
| **Q7RTV0** | PHF5A | -0.029878081 | 3.162694152 | -1.143039018 | 0.293540361 | 0.685934755 | -6.499598252 |
| **P15924-1** | DSP | 0.034568839 | 1.755352436 | 0.165505478 | 0.873596747 | 0.97237605 | -6.499664948 |
| **Q96EK4** | THAP11 | 0.028889324 | 3.158886034 | 0.539418134 | 0.607646995 | 0.875703859 | -6.499733989 |
| **O75787** | ATP6AP2 | 0.033654225 | 3.165108956 | 0.98543906 | 0.359807756 | 0.73661838 | -6.5002039 |
| **Q9P2N6-3** | KANSL3 | -0.024477593 | 3.161728013 | -1.163873794 | 0.285584602 | 0.681580409 | -6.500417065 |
| **Q96SL4** | GPX7 | 0.022038786 | 3.195838757 | 0.681445394 | 0.51922142 | 0.834217303 | -6.501711582 |
| **Q00536-2** | CDK16 | -0.001819146 | 3.174761449 | -0.047693204 | 0.963398935 | 0.991283985 | -6.501791615 |
| **O75063** | FAM20B | 0.024814695 | 3.181636458 | 0.663357605 | 0.53001203 | 0.839849063 | -6.501900584 |
| **Q969T7** | NT5C3B; NT5C3L | -0.013395926 | 3.185000026 | -0.31437404 | 0.763127729 | 0.939644172 | -6.5020807 |
| **P21127-1** | CDK11B | -0.080661031 | 3.123214308 | -0.647140191 | 0.539806648 | 0.843540088 | -6.502777866 |
| **Q9GZN1** | ACTR6 | 0.019652048 | 3.183571663 | 0.731543962 | 0.490081144 | 0.818503298 | -6.50285304 |

| | | | | | | | |
|---|---|---|---|---|---|---|---|
| **Q8WUN7** | UBTD2 | 0.022353044 | 3.161030258 | 0.645729153 | 0.540664172 | 0.843540088 | -6.503054473 |
| **Q9H9H4** | VPS37B | 0.017009246 | 3.155063973 | 0.513665793 | 0.624550514 | 0.885387512 | -6.503231821 |
| **P15822-1** | HIVEP1 | -0.000501852 | 3.226174892 | -0.014162792 | 0.98912675 | 0.996219143 | -6.503629124 |
| **P23229-1** | ITGA6 | 0.016813136 | 3.170028083 | 0.585332221 | 0.578152727 | 0.861366689 | -6.50413323 |
| **Q9Y6X9-1** | MORC2 | -0.035735293 | 3.194653589 | -1.346254397 | 0.223464204 | 0.629582661 | -6.504150723 |
| **Q9Y692-2** | GMEB1 | 0.025867876 | 3.181168053 | 1.010976466 | 0.348324553 | 0.730167181 | -6.504858992 |
| **Q96JM3** | CHAMP1 | 0.030771384 | 3.189794353 | 1.718451452 | 0.132900812 | 0.519961998 | -6.504984708 |
| **Q96LW4** | CCDC111; PRIMPC | -0.018184956 | 3.200533082 | -0.437946459 | 0.675667899 | 0.903340705 | -6.505056679 |
| **O75821** | EIF3G | -0.042380315 | 3.193181329 | -1.529837304 | 0.173360989 | 0.574167672 | -6.505105756 |
| **Q13601** | KRR1 | -0.06161519 | 3.190608458 | -1.110649485 | 0.306274554 | 0.694846183 | -6.50516429 |
| **Q96EK9** | KTI12 | -0.025446444 | 3.213999896 | -1.212221656 | 0.267820751 | 0.670184524 | -6.505685439 |
| **Q8TCB0** | IFI44 | -0.003007771 | 3.166584703 | -0.064706677 | 0.950360586 | 0.989472196 | -6.506004792 |
| **Q9Y487** | ATP6V0A2 | -0.034491348 | 3.18340892 | -1.092459032 | 0.313624046 | 0.701109468 | -6.506787382 |
| **Q9H7E9-2** | C8orf33 | -0.054838455 | 3.136497554 | -0.750855054 | 0.479145683 | 0.811537935 | -6.507019521 |
| **P62158** | CALM3; CALM2; C/ | -0.284912242 | 3.214069688 | -0.924683627 | 0.388310919 | 0.75788842 | -6.507059667 |
| **Q9HBH5** | RDH14 | 0.032452396 | 3.152181289 | 0.918782453 | 0.391168758 | 0.759806823 | -6.508598858 |
| **Q9H1I8-1** | ASCC2 | 0.024644248 | 3.178786899 | 1.308763031 | 0.235171157 | 0.64164013 | -6.508734039 |
| **Q7Z3U7-1** | MON2 | -0.047454412 | 3.190296253 | -1.433025448 | 0.198340835 | 0.603708078 | -6.50884616 |
| **Q99504** | EYA3 | -0.028374667 | 3.194108442 | -0.555022561 | 0.597529234 | 0.871651708 | -6.509259636 |
| **Q8IW35-2** | CEP97 | 0.022092176 | 3.181327809 | 1.405888679 | 0.205909875 | 0.612058044 | -6.50951645 |
| **Q13200** | PSMD2 | 0.02967713 | 3.176161643 | 1.873572121 | 0.106569459 | 0.475403904 | -6.509694819 |
| **Q9NWL6** | ASNSD1; ASDURF | 0.012418462 | 3.154446495 | 0.179394169 | 0.863114698 | 0.969524427 | -6.509766345 |
| **Q14207** | NPAT | 0.026812022 | 3.168555047 | 1.173427924 | 0.281997368 | 0.679261937 | -6.509950603 |
| **Q8TA86** | RP9 | 0.029552377 | 3.196850458 | 1.135450027 | 0.296483851 | 0.688880164 | -6.510194124 |
| **Q8WWN9-2** | IPCEF1 | -0.00792878 | 3.186765533 | -0.162692193 | 0.875723435 | 0.972759481 | -6.510196139 |
| **Q9Y3D8** | AK6; TAF9 | -0.034559913 | 3.208848924 | -0.854027903 | 0.423572075 | 0.778083043 | -6.510636171 |
| **Q96EY5-1** | FAM125A; MVB12A | -0.000663682 | 3.153313521 | -0.009729932 | 0.992529853 | 0.99756865 | -6.511324318 |
| **O00182-1** | LGALS9 | -0.034213353 | 3.167884338 | -0.982184329 | 0.361292355 | 0.736919727 | -6.511814108 |
| **Q96HW7** | INTS4 | 0.029568005 | 3.19767334 | 1.532216047 | 0.172785452 | 0.572946752 | -6.512022783 |
| **O00233-1** | PSMD9 | -0.031694736 | 3.198430964 | -1.103555412 | 0.309123749 | 0.696978291 | -6.513089145 |
| **Q9Y3A6-1** | TMED5 | 0.071740646 | 3.221462785 | 0.740029552 | 0.485255422 | 0.81473183 | -6.513413252 |
| **Q9Y4E6-1** | WDR7 | 0.02538655 | 3.191503962 | 1.349146009 | 0.222583114 | 0.6284374 | -6.513582174 |
| **Q11206-4** | ST3GAL4 | 0.003251122 | 3.191613898 | 0.057175481 | 0.956130365 | 0.989472196 | -6.513777016 |
| **Q9H8M7-1** | FAM188A; MINDY3 | 0.002542859 | 3.190483462 | 0.062812181 | 0.951811702 | 0.989472196 | -6.514093071 |
| **Q8IY22-1** | CMIP | -0.016176245 | 3.198975278 | -0.482823613 | 0.645123036 | 0.891546795 | -6.514147728 |
| **Q9NS56** | TOPORS | -0.005010376 | 3.205734386 | -0.128074098 | 0.901978673 | 0.97952714 | -6.514297225 |
| **P08865** | RPSA | -0.026674069 | 3.193750192 | -1.876201258 | 0.106170284 | 0.475403904 | -6.514799948 |
| **Q9UHD2** | TBK1 | -0.033513307 | 3.178351203 | -0.949307526 | 0.376556994 | 0.749049177 | -6.514801582 |
| **Q8TF01** | PNISR | -0.034342561 | 3.193049863 | -1.066905901 | 0.324191332 | 0.710703247 | -6.516119722 |

| | | | | | | | |
|---|---|---|---|---|---|---|---|
| **Q9BSR8** | YIPF4 | -0.035036737 | 3.191332947 | -0.429063345 | 0.681796322 | 0.904168215 | -6.516474511 |
| **P56377** | AP1S2 | -0.011630773 | 3.21812654 | -0.218902471 | 0.833468258 | 0.96068017 | -6.516878217 |
| **Q9Y3I1** | FBXO7 | -0.023651454 | 3.177545481 | -0.599704639 | 0.569094498 | 0.855040807 | -6.517023867 |
| **Q5VT25-6** | CDC42BPA | 0.021705011 | 3.18661222 | 0.913150616 | 0.39391096 | 0.760194378 | -6.517930988 |
| **P10242-4** | MYB | 0.046542379 | 3.187273283 | 1.433908148 | 0.198098923 | 0.603708078 | -6.518183475 |
| **P0C2W1** | FBXO45 | -0.014634224 | 3.182297928 | -0.46800194 | 0.655132836 | 0.896497734 | -6.519935254 |
| **P14854** | COX6B1 | -0.208136999 | 3.178026062 | -1.215358311 | 0.26670151 | 0.669704013 | -6.520133538 |
| **Q96CP2** | FLYWCH2 | 0.021835642 | 3.185313952 | 0.935605868 | 0.383063293 | 0.754258979 | -6.520935328 |
| **Q9Y2X3** | NOP58 | 0.024597283 | 3.171302324 | 1.787117551 | 0.120546089 | 0.499295152 | -6.521356384 |
| **P48426** | PIP4K2A | 0.042614782 | 3.187767349 | 1.253229594 | 0.253501697 | 0.658193131 | -6.52142008 |
| **Q9Y2G9** | SBNO2 | 0.001028843 | 3.195245198 | 0.027248143 | 0.979082868 | 0.99371658 | -6.521619398 |
| **O14545** | TRAFD1 | -0.037835015 | 3.234880458 | -1.263760074 | 0.249933147 | 0.654571265 | -6.522058752 |
| **Q8IZM8** | ZNF654 | -0.021793289 | 3.195883621 | -1.023767354 | 0.342682793 | 0.724485042 | -6.522259472 |
| **P16989-1** | CSDA; YBX3 | -0.011572537 | 3.217288554 | -0.258114957 | 0.804332712 | 0.953936916 | -6.522919524 |
| **Q96RU2-2** | USP28 | -0.019725404 | 3.181381177 | -1.169411626 | 0.283500693 | 0.679787821 | -6.523138199 |
| **Q13503** | MED21 | 0.063399815 | 3.156300762 | 0.477431914 | 0.648755199 | 0.893481222 | -6.523171416 |
| **Q9BX10** | GTPBP2 | 0.022293168 | 3.127911207 | 0.718005803 | 0.497846553 | 0.82455511 | -6.523178856 |
| **Q6PK18-1** | OGFOD3 | 0.004772641 | 3.19839429 | 0.120583656 | 0.907678516 | 0.98075504 | -6.523288456 |
| **Q9H477** | RBKS | 0.045833375 | 3.206759948 | 0.412682797 | 0.693165799 | 0.909991705 | -6.523684477 |
| **O75581** | LRP6 | -0.009687787 | 3.155703162 | -0.231310807 | 0.824215239 | 0.959077751 | -6.524016276 |
| **Q8N0S6-2** | CENPL | -0.023447317 | 3.208378798 | -0.504427921 | 0.630675439 | 0.887009447 | -6.524295758 |
| **Q9Y592** | CCDC41; CEP83 | -0.01882478 | 3.18998119 | -0.457703758 | 0.662133632 | 0.897125645 | -6.524578273 |
| **P04792** | HSPB1 | -0.044978042 | 2.665981469 | -0.295214973 | 0.777074396 | 0.943581855 | -6.525409703 |
| **O75396** | SEC22B | -0.025258637 | 3.180657825 | -1.228103473 | 0.262194804 | 0.665868547 | -6.525422932 |
| **Q9Y450-2** | HBS1L | -0.042959563 | 3.174914408 | -0.793984349 | 0.455327211 | 0.797877083 | -6.525623358 |
| **Q99519** | NEU1 | 0.008992366 | 3.175474713 | 0.22900422 | 0.825933032 | 0.959077751 | -6.525795683 |
| **Q9BZG8** | DPH1 | 0.000450338 | 3.167123246 | 0.01182356 | 0.990922551 | 0.996844685 | -6.526479176 |
| **Q32NB8** | PGS1 | 0.013608522 | 3.186871321 | 0.285483128 | 0.784193451 | 0.946296623 | -6.527583297 |
| **Q5T280** | C9orf114; SPOUT1 | -0.042776348 | 3.146448822 | -0.644069421 | 0.54167392 | 0.843570684 | -6.528491797 |
| **O15357** | INPPL1 | -0.021746104 | 3.161551219 | -1.113461199 | 0.30515129 | 0.693599527 | -6.528528501 |
| **Q9NX63** | CHCHD3 | 0.053246505 | 3.176757863 | 1.261306132 | 0.250760817 | 0.655139636 | -6.529291546 |
| **Q9Y484-3** | WDR45 | -0.022488927 | 3.162726015 | -0.752141402 | 0.478423176 | 0.811328097 | -6.52991013 |
| **Q5BKY9-1** | FAM133B | -0.002028293 | 3.192462272 | -0.055409673 | 0.95748361 | 0.989500907 | -6.530430508 |
| **Q9UKM7** | MAN1B1 | -0.030294552 | 3.177957379 | -0.901973286 | 0.39939616 | 0.763210165 | -6.53063884 |
| **Q6P1M9** | ARMCX5 | -0.024583522 | 3.171168026 | -0.818712441 | 0.442050557 | 0.789324124 | -6.53155398 |
| **P02794** | FTH1 | 0.023641376 | 3.215347892 | 0.689419904 | 0.514509201 | 0.831693917 | -6.531734034 |
| **Q6FIF0** | ZFAND6 | -0.070345205 | 3.175888334 | -0.989478365 | 0.357971921 | 0.735096127 | -6.533021923 |
| **P61224-1** | RAP1B | 0.046829049 | 3.160751248 | 1.053580235 | 0.329815595 | 0.715561141 | -6.533195564 |
| **Q96EK5** | KIAA1279; KIF1BP | -0.024168612 | 3.197469027 | -1.363304013 | 0.218313589 | 0.625083672 | -6.533251666 |

| | | | | | | | |
|---|---|---|---|---|---|---|---|
| **Q16560-2** | SNRNP35 | 0.02777739 | 3.21461083 | 0.741190995 | 0.484597413 | 0.814349276 | -6.534230941 |
| **Q93034** | CUL5 | 0.020052914 | 3.195478906 | 1.572498937 | 0.163305092 | 0.561580888 | -6.53429493 |
| **Q9BZX2-1** | UCK2 | -0.036718613 | 3.209599436 | -0.87442489 | 0.413158821 | 0.771093923 | -6.534491868 |
| **O43524** | FOXO3 | -0.016387186 | 3.21324976 | -0.663240993 | 0.530082055 | 0.839849063 | -6.534841281 |
| **Q9UG63-2** | ABCF2 | -0.048391236 | 3.199681679 | -1.845174514 | 0.110975974 | 0.482457068 | -6.535175454 |
| **Q9BQA9-1** | C17orf62 | -0.036386962 | 3.179081889 | -0.890643333 | 0.40501444 | 0.766336431 | -6.535460222 |
| **Q96BM9** | ARL8A | 0.009175794 | 3.208330223 | 0.093358922 | 0.928442443 | 0.98601391 | -6.536094735 |
| **Q99733-2** | NAP1L4 | -0.033938002 | 3.184389294 | -1.38548611 | 0.211771315 | 0.616308826 | -6.536179168 |
| **Q15286** | RAB35 | -0.082914419 | 3.156607801 | -1.043004892 | 0.334334798 | 0.719277002 | -6.536675747 |
| **P47224** | RABIF | -0.003396976 | 3.189987041 | -0.082228466 | 0.936950031 | 0.987232823 | -6.537224862 |
| **O43379-1** | WDR62 | -0.018869604 | 3.190153159 | -1.156682853 | 0.288309784 | 0.68315575 | -6.538301517 |
| **Q9Y672** | ALG6 | 0.01242904 | 3.171649816 | 0.169344508 | 0.870696492 | 0.971931713 | -6.538918057 |
| **O75461-1** | E2F6 | 0.004030375 | 3.181808825 | 0.060166622 | 0.953838436 | 0.989472196 | -6.539193093 |
| **Q15041** | ARL6IP1 | -0.027072446 | 3.207835929 | -0.966078676 | 0.368709076 | 0.743611616 | -6.539426286 |
| **Q9BT25** | HAUS8 | 0.022709826 | 3.196541957 | 0.687138362 | 0.515854555 | 0.832851949 | -6.539642204 |
| **O60671-1** | RAD1 | 0.022795269 | 3.212755483 | 0.956984554 | 0.372948774 | 0.745722908 | -6.539872882 |
| **Q5VUG0** | SFMBT2 | -0.01594531 | 3.184873982 | -0.461131986 | 0.659798955 | 0.896852471 | -6.540484491 |
| **Q8TF74** | WIPF2 | -0.021059223 | 3.1939994 | -0.221925127 | 0.831211518 | 0.960187608 | -6.540605387 |
| **Q96P70** | IPO9 | -0.024679232 | 3.180125599 | -1.611777937 | 0.154531596 | 0.553156191 | -6.541174169 |
| **Q15773** | MLF 2.00 | 0.004787294 | 3.213010797 | 0.125665921 | 0.903810503 | 0.980210603 | -6.541226263 |
| **O14593** | RFXANK | -0.025163773 | 3.205073314 | -0.684576669 | 0.517367813 | 0.833280927 | -6.541236207 |
| **P23246-1** | SFPQ | 0.023570233 | 3.181836865 | 2.056826499 | 0.082014834 | 0.423947857 | -6.541552463 |
| **Q8TAC1-2** | RFESD | 0.012921566 | 3.202968385 | 0.338288552 | 0.745853517 | 0.934974996 | -6.541608713 |
| **Q8N668** | COMMD1 | -0.026489422 | 3.17743054 | -0.562906943 | 0.592453528 | 0.869575962 | -6.541731673 |
| **Q9Y2F5** | KIAA0947; ICE1 | -0.019187005 | 3.194514188 | -1.395410996 | 0.208901499 | 0.614302382 | -6.542029805 |
| **Q86VV8** | RTTN | -0.031767432 | 3.181884202 | -1.277635814 | 0.245297647 | 0.650552011 | -6.542493885 |
| **P61421** | ATP6V0D1 | -0.037897622 | 3.173915045 | -0.782188728 | 0.461758039 | 0.802800142 | -6.542666457 |
| **Q9HAU5** | UPF2 | 0.021791046 | 3.188134342 | 1.604463088 | 0.156131128 | 0.554834464 | -6.542695715 |
| **Q99941-1** | ATF6B | -0.002471839 | 3.195188435 | -0.03180446 | 0.975586459 | 0.99316605 | -6.543131107 |
| **P49459-1** | UBE2A | -0.012819669 | 3.204113468 | -0.348789012 | 0.738318007 | 0.930510242 | -6.543151357 |
| **Q06481** | APLP2 | 0.020884339 | 3.221053358 | 0.591304905 | 0.574378183 | 0.85911412 | -6.545000923 |
| **Q9Y2X9** | ZNF281 | 0.022656714 | 3.199467669 | 0.955610691 | 0.373592529 | 0.746151997 | -6.545731139 |
| **Q9Y3A2** | UTP11L; UTP11 | -0.062053124 | 3.159727564 | -0.901484641 | 0.399637259 | 0.763210165 | -6.546291817 |
| **Q9NQ86** | TRIM36 | -0.029097371 | 3.21579358 | -0.469957374 | 0.653807742 | 0.896119351 | -6.546637186 |
| **Q13352-5** | ITGB3BP | -0.007367533 | 3.1628647 | -0.207076919 | 0.842313559 | 0.962493422 | -6.546893566 |
| **Q9BY50** | SEC11C | -0.026850703 | 3.146959197 | -0.35186248 | 0.736118246 | 0.929681266 | -6.547205241 |
| **Q9UQ84-1** | EXO1 | -0.021750163 | 3.15358564 | -0.731379853 | 0.490174788 | 0.818503298 | -6.548554321 |
| **Q01432-4** | AMPD3 | -0.023601013 | 3.204824281 | -0.747992479 | 0.480756175 | 0.81235548 | -6.54880339 |
| **Q9NVF7-1** | FBXO28 | 0.008336099 | 3.193710277 | 0.234190014 | 0.82207247 | 0.959068023 | -6.549066509 |

| | | | | | | | |
|---|---|---|---|---|---|---|---|
| **Q8NFU5** | IPMK | -0.013232232 | 3.203242562 | -0.26940453 | 0.796004452 | 0.950729912 | -6.549542643 |
| **Q92896-2** | GLG1 | -0.021014924 | 3.162991949 | -1.753806272 | 0.126394319 | 0.509218658 | -6.549842643 |
| **Q9P0L0** | VAPA | -0.033177253 | 3.193709045 | -0.797542911 | 0.4533995 | 0.796326278 | -6.550030623 |
| **Q9Y5X3** | SNX5 | -0.039440667 | 3.189423767 | -1.431973105 | 0.19862959 | 0.603946458 | -6.550758974 |
| **O96007** | MOCS2 | 0.030056931 | 3.194371688 | 0.778375871 | 0.46385022 | 0.802800142 | -6.551343778 |
| **P32121-4** | ARRB2 | 0.066760264 | 3.173761515 | 1.464729022 | 0.189818788 | 0.593849183 | -6.551793628 |
| **Q9NRV9** | HEBP1 | -0.008929677 | 3.203755221 | -0.136074277 | 0.895897905 | 0.978757548 | -6.552044081 |
| **P37837** | TALDO1 | -0.029770992 | 3.174433027 | -1.810412597 | 0.116613236 | 0.49281498 | -6.55253376 |
| **Q99704** | DOK1 | -0.02156354 | 3.183977402 | -0.554020193 | 0.598176287 | 0.87186032 | -6.5534036 |
| **Q15029** | EFTUD2 | -0.040256746 | 3.178371218 | -1.915986568 | 0.100306435 | 0.461723423 | -6.55343665 |
| **Q96HD1-2** | CRELD1 | -0.029058492 | 3.201605153 | -0.448140141 | 0.668668212 | 0.90001396 | -6.553532992 |
| **P62306** | SNRPF | -0.021577912 | 3.186311787 | -0.626915833 | 0.552178211 | 0.847904115 | -6.554082658 |
| **Q13905-3** | RAPGEF1 | -0.024441748 | 3.169609876 | -0.990065406 | 0.357705727 | 0.734856937 | -6.554994816 |
| **P46459** | NSF | 0.01983005 | 3.21087711 | 0.398507963 | 0.703074462 | 0.915549813 | -6.555419916 |
| **O95104** | SCAF4 | 0.042181553 | 3.165734137 | 1.136313842 | 0.296147574 | 0.688880164 | -6.555639548 |
| **Q9Y6A9** | SPCS1 | 0.045270111 | 3.217499719 | 0.531421141 | 0.612868908 | 0.878766193 | -6.555810658 |
| **P11166** | SLC2A1 | -0.040158255 | 3.145521261 | -0.622606197 | 0.554836789 | 0.849359979 | -6.556999006 |
| **Q6PI47** | KCTD18 | -0.019015204 | 3.163227075 | -0.577186668 | 0.583323805 | 0.864969696 | -6.558270578 |
| **P78312-2** | FAM193A | -0.022940697 | 3.212704313 | -1.022129926 | 0.343400947 | 0.724892107 | -6.558429797 |
| **Q8IVW6** | ARID3B | -0.024140065 | 3.209452697 | -0.540666826 | 0.606833855 | 0.875166995 | -6.55861666 |
| **Q8NEZ4-3** | KMT2C; MLL3 | -0.020303095 | 3.185967299 | -1.456624283 | 0.191964985 | 0.596505086 | -6.558838927 |
| **Q8NFA0** | USP32 | -0.022979848 | 3.198188403 | -1.158634847 | 0.287567878 | 0.682522792 | -6.559177753 |
| **Q99640** | PKMYT1 | -0.010682114 | 3.175047207 | -0.227219172 | 0.82726313 | 0.959077751 | -6.559512446 |
| **Q96IG2-1** | FBXL20 | -0.001999558 | 3.134354667 | -0.041833675 | 0.967892457 | 0.992306703 | -6.559544484 |
| **Q9NRR3** | CDC42SE2 | -0.006611217 | 3.22296468 | -0.173047828 | 0.867900814 | 0.970868803 | -6.559837571 |
| **P25685** | DNAJB1 | 0.034881468 | 3.209835957 | 0.723303443 | 0.49479817 | 0.822007449 | -6.560733506 |
| **Q92585** | MAML1 | 0.020068845 | 3.194567922 | 0.847957505 | 0.42670787 | 0.779764001 | -6.560955528 |
| **Q9H6S0** | YTHDC2 | -0.014928016 | 3.182228033 | -1.611572369 | 0.154576336 | 0.553156191 | -6.560990553 |
| **Q9UBS0** | RPS6KB2 | 0.005351019 | 3.183814174 | 0.155040636 | 0.881513199 | 0.974438769 | -6.561153645 |
| **P16591-1** | FER | 0.029036955 | 3.191806914 | 0.876013449 | 0.41235579 | 0.770644294 | -6.561274482 |
| **Q03111** | MLLT1 | -0.038063504 | 3.179307874 | -0.698925625 | 0.508928478 | 0.82913633 | -6.561744057 |
| **Q15172-1** | PPP2R5A | 0.024349839 | 3.189094471 | 1.186943391 | 0.276987782 | 0.675421056 | -6.561992934 |
| **Q9UBI1** | COMMD3 | -0.064186203 | 3.159817867 | -1.187548399 | 0.276765305 | 0.675408632 | -6.562374635 |
| **Q6PI98-1** | INO80C | 0.032857701 | 3.21073161 | 0.517182983 | 0.622226967 | 0.883385606 | -6.563285444 |
| **Q92859** | NEO1 | -0.029710073 | 3.148145356 | -1.089634181 | 0.314778238 | 0.701837878 | -6.563950805 |
| **O00462** | MANBA | 0.016353463 | 3.190608331 | 0.415906764 | 0.690921153 | 0.909522034 | -6.564052449 |
| **Q13190** | STX5 | 0.025537357 | 3.148635197 | 0.875609912 | 0.412559673 | 0.770644294 | -6.564351052 |
| **Q9Y6P5-2** | SESN1 | -0.019073867 | 3.164201392 | -0.304681345 | 0.770171854 | 0.941676728 | -6.564380512 |
| **Q01970-1** | PLCB3 | -0.021900751 | 3.169730279 | -0.832762171 | 0.434631065 | 0.784298014 | -6.564639509 |

| | | | | | | | |
|---|---|---|---|---|---|---|---|
| **Q15560** | TCEA2 | 0.000865273 | 3.153115157 | 0.020265358 | 0.984442235 | 0.995531556 | -6.564811427 |
| **Q969F9-1** | HPS3 | -0.020911304 | 3.177365705 | -1.115545804 | 0.304320698 | 0.692985939 | -6.5648673 |
| **P55347** | PKNOX1 | -0.007331936 | 3.172428059 | -0.166967608 | 0.872491906 | 0.97208641 | -6.564931558 |
| **P05204** | HMGN2 | -0.009557156 | 3.120611653 | -0.039997895 | 0.96930054 | 0.992469575 | -6.565168355 |
| **Q8N5P1** | ZC3H8 | -0.027902514 | 3.173947038 | -0.900393368 | 0.400176092 | 0.763361117 | -6.56560256 |
| **Q99805** | TM9SF2 | -0.040667656 | 3.193623025 | -1.121215392 | 0.302071146 | 0.691344519 | -6.565653858 |
| **P61009** | SPCS3 | -0.049543284 | 3.169019858 | -1.068426002 | 0.323554716 | 0.710703247 | -6.565788455 |
| **P42566-1** | EPS15 | -0.025350525 | 3.186972282 | -1.561117367 | 0.165933287 | 0.565325887 | -6.565871111 |
| **P61018-2** | RAB4B | 0.025386821 | 3.199671548 | 0.646008968 | 0.540494054 | 0.843540088 | -6.566544577 |
| **O15258** | RER1 | 0.063252861 | 3.192617458 | 1.113821947 | 0.305007419 | 0.693599527 | -6.566634631 |
| **Q16594** | TAF9 | -0.01474593 | 3.191960532 | -0.430063192 | 0.681105211 | 0.903980599 | -6.566666832 |
| **Q8WZ64** | ARAP2 | 0.008891842 | 3.207614692 | 0.218257162 | 0.833950274 | 0.96068017 | -6.566783175 |
| **Q70UQ0** | IKBIP | 0.011341555 | 3.201822307 | 0.262394893 | 0.801172096 | 0.952917733 | -6.567193709 |
| **P15735-1** | PHKG2 | 0.024557181 | 3.200284881 | 1.262651644 | 0.250306706 | 0.654603944 | -6.567318832 |
| **P01023** | A2M | 0.068340716 | 3.195836211 | 0.421790004 | 0.686833753 | 0.907305098 | -6.567478568 |
| **Q6ZMZ3** | SYNE3 | -0.02816042 | 3.197374569 | -0.961117045 | 0.371017555 | 0.744717822 | -6.567940149 |
| **O14879** | IFIT3 | -0.018047347 | 3.17287147 | -1.201801949 | 0.271567612 | 0.671838876 | -6.568438213 |
| **P48454-3** | PPP3CC | 0.013696654 | 3.180738282 | 0.3920602 | 0.707602697 | 0.918150554 | -6.568644029 |
| **Q9BRQ0** | PYGO2 | -0.028172137 | 3.199465798 | -0.663877782 | 0.529699734 | 0.839849063 | -6.568805487 |
| **Q02750** | MAP2K1 | -0.038630227 | 3.177972427 | -1.326460797 | 0.229579005 | 0.637461728 | -6.569277686 |
| **P22415** | USF1 | -0.038429692 | 3.17139595 | -0.822270775 | 0.440162943 | 0.788699726 | -6.569324051 |
| **Q04656-3** | ATP7A | 0.026990656 | 3.191311683 | 0.596374914 | 0.571185533 | 0.856724384 | -6.569326471 |
| **P51970** | NDUFA8 | 0.071987508 | 3.147948579 | 0.57928762 | 0.581987479 | 0.864278602 | -6.569387151 |
| **Q75QN2** | INTS8 | -0.023897508 | 3.188617897 | -1.010064436 | 0.348729621 | 0.730176717 | -6.569439556 |
| **Q9NR46** | SH3GLB2 | 0.022357564 | 3.19148784 | 1.2599136 | 0.25123155 | 0.655727551 | -6.569628827 |
| **O43929** | ORC4 | 0.027401286 | 3.175759909 | 0.978984474 | 0.362756578 | 0.738054037 | -6.570055831 |
| **Q8N6M0** | OTUD6B | -0.030684569 | 3.180527988 | -0.975704258 | 0.36426237 | 0.739413523 | -6.570353118 |
| **Q96PM5-1** | RCHY1 | -0.018157907 | 3.217606122 | -0.945941389 | 0.378147519 | 0.750267026 | -6.570785809 |
| **Q01968-1** | OCRL | -0.003380876 | 3.190198309 | -0.098962206 | 0.92416334 | 0.985099254 | -6.571191337 |
| **Q8N5L8** | RPP25L | -0.020529854 | 3.209655826 | -0.378181227 | 0.717393482 | 0.924679646 | -6.571313637 |
| **Q6P9B9** | INTS5 | 0.017758942 | 3.171828224 | 0.741280987 | 0.484546454 | 0.814349276 | -6.571668228 |
| **O15162** | PLSCR1 | 0.001795217 | 3.1646646 | 0.030363264 | 0.97669234 | 0.99316605 | -6.571783975 |
| **Q8N1G1** | REXO1 | -0.010931099 | 3.20889467 | -0.334101092 | 0.74886712 | 0.935495168 | -6.57189315 |
| **P57737-3** | CORO7-PAM16; C( | 0.117177081 | 3.189567459 | 1.171815364 | 0.282600149 | 0.679557562 | -6.572174309 |
| **Q14693-7** | LPIN1 | -0.025928759 | 3.198823955 | -0.818365214 | 0.442235061 | 0.789386107 | -6.572179505 |
| **Q9NX74** | DUS2; DUS2L | 0.034871716 | 3.18405293 | 0.996680988 | 0.354716599 | 0.734091082 | -6.572341349 |
| **Q9NRB3** | CHST12 | -0.012877224 | 3.1956279 | -0.262641907 | 0.800989807 | 0.952917733 | -6.572356022 |
| **Q8WUI4-7** | HDAC7 | -0.022915951 | 3.181745027 | -1.075834717 | 0.32046646 | 0.708257023 | -6.5726264 |
| **O43151** | TET3 | -0.000371659 | 3.1848867 | -0.009674651 | 0.992572293 | 0.99756865 | -6.572695792 |

| | | | | | | | |
|---|---|---|---|---|---|---|---|
| **Q9NS18-2** | GLRX2 | 0.005743502 | 3.238126706 | 0.140204381 | 0.892761654 | 0.978276697 | -6.572827051 |
| **P16930** | FAH | 0.01328356 | 3.195572472 | 0.335144523 | 0.748115741 | 0.935183323 | -6.573129338 |
| **Q9NP77** | SSU72 | -0.063207651 | 3.202697067 | -1.1466833 | 0.292135564 | 0.685264687 | -6.573171932 |
| **Q14103-3** | HNRNPD | 0.08187607 | 3.206172635 | 0.703090838 | 0.506495605 | 0.82913633 | -6.573284384 |
| **P14151-2** | SELL | -0.023841593 | 3.140254634 | -0.255433645 | 0.806314826 | 0.954293355 | -6.573484559 |
| **Q8NF64** | ZMIZ2 | -0.005898139 | 3.196723473 | -0.15937728 | 0.878230761 | 0.973200128 | -6.573663465 |
| **Q5JUR7** | TEX30 | -0.026445753 | 3.184884377 | -1.120481969 | 0.302361373 | 0.691344519 | -6.573726849 |
| **Q96DZ1** | ERLEC1 | 0.06205025 | 3.173225942 | 1.289339857 | 0.241446108 | 0.647414555 | -6.574764766 |
| **Q96DB5** | FAM82B; RMDN1 | 0.038222827 | 3.176374857 | 0.913025171 | 0.393972205 | 0.760194378 | -6.575072028 |
| **Q9Y5X2** | SNX8 | 0.009493464 | 3.185224295 | 0.314772139 | 0.762838929 | 0.939644172 | -6.575227983 |
| **Q86UA6-8** | RPAIN | -0.008042658 | 3.156933876 | -0.096799958 | 0.925814285 | 0.985476218 | -6.575334886 |
| **O75379-1** | VAMP4 | -0.02368775 | 3.179242593 | -0.44159371 | 0.673159377 | 0.901897345 | -6.575442902 |
| **Q8NDX1** | PSD4 | -0.034037598 | 3.19773633 | -1.444973081 | 0.19508915 | 0.600645673 | -6.575692803 |
| **O95544** | NADK | -0.019868513 | 3.19793023 | -0.794023156 | 0.455306158 | 0.797877083 | -6.575949458 |
| **Q2M296** | MTHFSD | 0.019961278 | 3.193769053 | 0.567927815 | 0.589234134 | 0.868063266 | -6.577459344 |
| **Q8IX01-1** | SUGP2 | -0.035527911 | 3.195421916 | -1.40848947 | 0.205173301 | 0.611215566 | -6.577859673 |
| **Q8N6T3-1** | ARFGAP1 | 0.04965606 | 3.195073606 | 1.313828476 | 0.233558437 | 0.640812038 | -6.577928332 |
| **O75143-2** | ATG13 | 0.030837443 | 3.162221804 | 0.64004961 | 0.544124344 | 0.844397822 | -6.578382746 |
| **Q96BT3-1** | CENPT | -0.016218972 | 3.213081537 | -0.51963164 | 0.620612084 | 0.883249609 | -6.578730412 |
| **Q9UBG0** | MRC2 | 0.017685478 | 3.201269485 | 0.394682913 | 0.705759208 | 0.917117584 | -6.578894399 |
| **Q13608** | PEX6 | -0.001173364 | 3.196838433 | -0.028429043 | 0.978176623 | 0.993585863 | -6.580179055 |
| **P18615** | NELFE; RDBP | -0.03600737 | 3.204740342 | -1.354731762 | 0.220889853 | 0.626484139 | -6.581090035 |
| **Q969J2** | ZKSCAN4 | 0.026891574 | 3.195929735 | 0.845921444 | 0.427763408 | 0.78056506 | -6.581859861 |
| **Q9H008-1** | LHPP | 0.035845232 | 3.179105751 | 0.636636226 | 0.546210481 | 0.845412981 | -6.582061242 |
| **Q8NC51-1** | SERBP1 | -0.020823573 | 3.185127279 | -0.788476016 | 0.458322449 | 0.799639769 | -6.582073284 |
| **Q9NXW2-1** | DNAJB12 | -0.025527439 | 3.199479681 | -0.996474053 | 0.354809801 | 0.734091082 | -6.582594422 |
| **P60468** | SEC61B | 0.018139774 | 3.199647455 | 0.37023292 | 0.723026863 | 0.926648409 | -6.582772465 |
| **Q8IY63-1** | AMOTL1 | 0.020676831 | 3.191050887 | 0.79683727 | 0.453781297 | 0.796605553 | -6.582915735 |
| **Q9NXA8-1** | SIRT5 | 0.028700077 | 3.175324831 | 0.957808603 | 0.372563057 | 0.745276848 | -6.583284734 |
| **Q2NL82** | TSR1 | -0.022633084 | 3.193206955 | -1.64319771 | 0.14783539 | 0.546231571 | -6.583369076 |
| **Q9NRX4** | PHPT1 | 0.036415523 | 3.17801085 | 1.038558621 | 0.336249621 | 0.720858943 | -6.58365689 |
| **Q8TEX9-2** | IPO4 | -0.026958267 | 3.205470795 | -1.306832205 | 0.235788462 | 0.641994289 | -6.583851893 |
| **Q15572-6** | TAF1C | 0.027280832 | 3.181605716 | 0.649472975 | 0.538390817 | 0.843524953 | -6.584217269 |
| **Q14CW9-2** | ATXN7L3 | -0.03265362 | 3.165417419 | -0.353264089 | 0.73511597 | 0.929571285 | -6.584461601 |
| **Q9NUP9** | LIN7C | -0.043568354 | 3.227108957 | -0.778329089 | 0.463875931 | 0.802800142 | -6.584887076 |
| **Q9C0J8** | WDR33 | 0.018008198 | 3.170422404 | 1.539804049 | 0.170961375 | 0.570709295 | -6.585185523 |
| **Q8N371-3** | KDM8 | 0.02014474 | 3.190118652 | 0.71711478 | 0.49836049 | 0.824890436 | -6.585487173 |
| **O94915-1** | FRYL | -0.029719619 | 3.182169796 | -1.579167384 | 0.161783384 | 0.561414897 | -6.585628723 |
| **Q13907** | IDI1 | 0.031761948 | 3.127970341 | 1.151012098 | 0.290474205 | 0.684803131 | -6.585631381 |

| | | | | | | | |
|---|---|---|---|---|---|---|---|
| **O75319-1** | DUSP11 | 0.036589753 | 3.179269209 | 0.910368354 | 0.39527101 | 0.761244496 | -6.585656297 |
| **Q15293** | RCN1 | -0.104851118 | 3.190383882 | -1.297338193 | 0.238844558 | 0.644491512 | -6.586361993 |
| **Q9UGJ1** | TUBGCP4 | -0.022770299 | 3.208678811 | -1.193482919 | 0.274591051 | 0.674173641 | -6.586447552 |
| **P09693** | CD3G | -0.025537698 | 3.189326373 | -1.181720583 | 0.278914649 | 0.676427621 | -6.586726147 |
| **Q5UCC4-1** | EMC10 | -0.014557754 | 3.212629172 | -0.292496385 | 0.779060789 | 0.944053983 | -6.586814527 |
| **Q9NPJ6-1** | MED4 | -0.025448058 | 3.179446166 | -0.834834017 | 0.433544557 | 0.783962934 | -6.587311705 |
| **Q9NW15-1** | ANO10 | -0.025211088 | 3.174319077 | -0.550382776 | 0.600527656 | 0.872406697 | -6.58761225 |
| **Q9NUQ3-1** | TXLNG | -0.02996517 | 3.190977506 | -1.133069576 | 0.297412188 | 0.689506809 | -6.58808147 |
| **P49917** | LIG4 | 0.037548603 | 3.181215936 | 1.3414664 | 0.224929951 | 0.630229362 | -6.58862507 |
| **Q68CP4-1** | HGSNAT | 0.016164096 | 3.188172164 | 0.131836397 | 0.899118108 | 0.978757548 | -6.588696337 |
| **Q5FWF4-1** | ZRANB3 | -0.030197762 | 3.193027044 | -0.741165007 | 0.48461213 | 0.814349276 | -6.588729361 |
| **Q6IA17-1** | SIGIRR | -0.002144035 | 3.186538167 | -0.066834298 | 0.94873115 | 0.989472196 | -6.588856321 |
| **Q9NQP4** | PFDN4 | -0.014296632 | 3.169703179 | -0.329103568 | 0.752469968 | 0.935782019 | -6.589418718 |
| **Q15696** | ZRSR2 | 0.008172766 | 3.165472116 | 0.117793134 | 0.909803497 | 0.981717884 | -6.589434105 |
| **Q9NPI6** | DCP1A | 0.019134975 | 3.188604548 | 1.390581915 | 0.210293448 | 0.615470161 | -6.589486036 |
| **P04114** | APOB | 0.051554442 | 2.996812319 | 1.204263303 | 0.270678516 | 0.671731232 | -6.589521784 |
| **Q9NXR8-1** | ING3 | -0.021544557 | 3.205397426 | -0.635750319 | 0.546752722 | 0.845784731 | -6.589797809 |
| **P78549** | NTHL1 | 0.025628567 | 3.201562807 | 0.620984592 | 0.55583916 | 0.850075481 | -6.589962822 |
| **Q86Y37** | CACUL1 | -0.026009921 | 3.201323433 | -0.979342613 | 0.362592468 | 0.738054037 | -6.58998853 |
| **Q6Q0C0-1** | TRAF7 | 0.01234367 | 3.213879692 | 0.392713638 | 0.707143199 | 0.91801515 | -6.590090692 |
| **Q01658** | DR1 | -0.013491995 | 3.185227708 | -0.209025817 | 0.840854071 | 0.961788885 | -6.590491767 |
| **Q9H9G7** | AGO3; EIF2C3 | 0.01351325 | 3.182361815 | 0.316766177 | 0.761392978 | 0.939181371 | -6.591114308 |
| **Q9NT62** | ATG3 | -0.031100066 | 3.206275469 | -1.136206478 | 0.296189353 | 0.688880164 | -6.591251441 |
| **Q96JI7-1** | SPG11 | -0.025992001 | 3.155018664 | -1.069184787 | 0.323237318 | 0.710406774 | -6.59125681 |
| **P27361-1** | MAPK3 | 0.049781643 | 3.181097882 | 0.998819749 | 0.353754447 | 0.733449655 | -6.591263059 |
| **Q7Z4G1-1** | COMMD6 | -0.013987909 | 3.192345176 | -0.45899174 | 0.661256014 | 0.896934512 | -6.591820849 |
| **P61966** | AP1S1 | 0.023938623 | 3.150319266 | 0.810736213 | 0.446302689 | 0.792020296 | -6.592748235 |
| **Q14088** | RAB33A | 0.023071427 | 3.147157522 | 0.580816964 | 0.58101585 | 0.86380444 | -6.592861344 |
| **Q96DX4** | RSPRY1 | 0.015071282 | 3.210072479 | 0.260628996 | 0.802475675 | 0.953157309 | -6.59290478 |
| **Q9P013** | CWC15 | -0.04753416 | 3.142691327 | -1.051303212 | 0.330784468 | 0.716489955 | -6.593058362 |
| **Q15390** | MTFR1 | 0.012604141 | 3.215740539 | 0.419346306 | 0.688530145 | 0.908052188 | -6.593307977 |
| **Q96IZ5** | RBM41 | 0.014270467 | 3.193077176 | 0.483952927 | 0.644363594 | 0.891474831 | -6.593413563 |
| **P49755** | TMED10 | -0.034743645 | 3.180055925 | -1.366580474 | 0.217336002 | 0.623815474 | -6.594082627 |
| **P07199** | CENPB | 0.029404318 | 3.188285999 | 0.984633445 | 0.360174781 | 0.73661838 | -6.594338681 |
| **Q9HBM0-1** | VEZT | -0.011008238 | 3.202202325 | -0.293672387 | 0.778201297 | 0.944053983 | -6.594538282 |
| **O95793** | STAU1 | 0.026824186 | 3.19317227 | 1.277573514 | 0.245318291 | 0.650552011 | -6.594757493 |
| **Q9UHY1** | NRBP1 | -0.021741444 | 3.16580574 | -0.700535463 | 0.507987279 | 0.82913633 | -6.594907883 |
| **Q9C0D3** | ZYG11B | 0.009085726 | 3.162231009 | 0.22804102 | 0.826650667 | 0.959077751 | -6.595110245 |
| **Q9UI30** | TRMT112 | -0.02857858 | 3.235846927 | -0.402538006 | 0.70025082 | 0.913245694 | -6.595685994 |

| | | | | | | | |
|---|---|---|---|---|---|---|---|
| **Q9H307** | PNN | -0.040669615 | 3.175996723 | -1.743913137 | 0.128183394 | 0.512600477 | -6.595900545 |
| **P55199** | ELL | -0.03197768 | 3.198148427 | -1.100799866 | 0.310236322 | 0.698251438 | -6.596010054 |
| **P22681** | CBL | -0.027323699 | 3.198767984 | -1.096223299 | 0.312091411 | 0.699835744 | -6.596095674 |
| **Q7L0J3** | SV2A | -0.010272766 | 3.159567378 | -0.369582942 | 0.723488371 | 0.926648409 | -6.596203785 |
| **Q12815-1** | TROAP | -0.01513751 | 3.139477907 | -0.340699699 | 0.744120471 | 0.934118682 | -6.596218772 |
| **P05023** | ATP1A1 | -0.052908781 | 3.185558896 | -1.610544064 | 0.154800319 | 0.553156191 | -6.596773796 |
| **Q86UP2-1** | KTN1 | 0.020121745 | 3.181172491 | 1.837299247 | 0.112229288 | 0.484388026 | -6.597015509 |
| **O60291-2** | MGRN1 | 0.011256851 | 3.177652064 | 0.222516965 | 0.830769849 | 0.960067847 | -6.597404547 |
| **O75607** | NPM3 | -0.018542948 | 3.160789088 | -0.255827362 | 0.80602368 | 0.954293355 | -6.598024834 |
| **P61916** | NPC2 | -0.027882987 | 3.153300663 | -0.664081207 | 0.529577636 | 0.839849063 | -6.598208982 |
| **Q9NUY8** | TBC1D23 | 0.019711047 | 3.176964601 | 0.756875122 | 0.475770763 | 0.809713018 | -6.598309735 |
| **Q6PJW8** | CNST | -0.01425562 | 3.177703768 | -0.358357295 | 0.731478619 | 0.92815165 | -6.598674786 |
| **Q8IZ73** | RPUSD2 | -0.025577088 | 3.196641407 | -1.054957787 | 0.329230557 | 0.714877501 | -6.598737962 |
| **Q14697-2** | GANAB | -0.005761451 | 3.16380934 | -0.11309076 | 0.913386163 | 0.982219313 | -6.599263595 |
| **P53803** | POLR2K | -0.01621837 | 3.211788881 | -0.438745925 | 0.675117652 | 0.90314827 | -6.599613938 |
| **Q9Y6M7-13** | SLC4A7 | -0.037366881 | 3.212372542 | -1.064963133 | 0.325006437 | 0.710703247 | -6.600097285 |
| **Q8IXH7** | NELFCD; TH1L | -0.028594732 | 3.180621195 | -1.275748319 | 0.245923775 | 0.651206185 | -6.600853262 |
| **Q9NXW9-1** | ALKBH4 | -0.037999106 | 3.149435467 | -0.792656146 | 0.456048184 | 0.798084321 | -6.601245776 |
| **Q86W34-4** | AMZ2 | -0.00351811 | 3.170466171 | -0.058275425 | 0.955287492 | 0.989472196 | -6.601774134 |
| **P54819** | AK2 | 0.05274907 | 3.175481404 | 1.507306419 | 0.178901122 | 0.580540072 | -6.601799563 |
| **O00311-1** | CDC7 | 0.014809016 | 3.172094443 | 0.606512141 | 0.564833664 | 0.852624219 | -6.602005241 |
| **Q12772** | SREBF2 | -0.021403908 | 3.184431507 | -0.552958633 | 0.598861983 | 0.871871293 | -6.602644917 |
| **Q8TEQ0-1** | SNX29 | -0.002987627 | 3.208622516 | -0.076540406 | 0.941301257 | 0.988377491 | -6.602668369 |
| **Q13506-1** | NAB1 | 0.022786368 | 3.180253219 | 0.670554805 | 0.525701447 | 0.838233243 | -6.603088426 |
| **Q29963** | HLA-C | -0.080822405 | 3.146819028 | -0.564656587 | 0.591330511 | 0.868764508 | -6.603327893 |
| **Q07960** | ARHGAP1 | -0.021575302 | 3.172970598 | -1.226449784 | 0.262775837 | 0.666498017 | -6.603496016 |
| **Q9NX04** | C1orf109 | -0.05212662 | 3.158276237 | -0.562949648 | 0.592426103 | 0.869575962 | -6.603629132 |
| **O60308** | CEP104 | 0.002315569 | 3.214420699 | 0.080008786 | 0.938647761 | 0.987755823 | -6.604082824 |
| **Q9UDW3** | ZMAT5 | -0.001710363 | 3.183076793 | -0.034560973 | 0.973471456 | 0.99316605 | -6.604203732 |
| **O95817** | BAG3 | -0.035795671 | 3.169089262 | -0.5706368 | 0.58750131 | 0.866635334 | -6.604409787 |
| **Q8IUR0** | TRAPPC5 | -0.033145838 | 3.208112437 | -0.790793494 | 0.457060611 | 0.798548386 | -6.604478652 |
| **P48200** | IREB2 | -0.041272683 | 3.068315812 | -1.258732147 | 0.25163153 | 0.655935708 | -6.604579436 |
| **Q6PIU2-2** | NCEH1 | 0.016609215 | 3.210351647 | 0.367428459 | 0.725019028 | 0.926648409 | -6.604786578 |
| **O60551** | NMT2 | 0.027134191 | 3.182986594 | 0.606115348 | 0.565081492 | 0.852720314 | -6.605279951 |
| **P08579** | SNRPB2 | 0.034296904 | 3.168169571 | 0.61808918 | 0.557631651 | 0.851117349 | -6.605536405 |
| **Q9H0F7-2** | ARL 6.00 | 0.00512049 | 3.191836127 | 0.136372508 | 0.895671371 | 0.978757548 | -6.605559299 |
| **P16150** | SPN; LOC10192988 | -0.06454012 | 3.193227021 | -0.866979075 | 0.416938106 | 0.773310479 | -6.605655954 |
| **Q92530** | PSMF1 | -0.032929032 | 3.187829184 | -1.290882985 | 0.240942265 | 0.64649993 | -6.605920021 |
| **Q9NSI6-2** | BRWD1 | 0.013458224 | 3.176652639 | 0.310673342 | 0.765814355 | 0.940484723 | -6.606184371 |

| | | | | | | | |
|---|---|---|---|---|---|---|---|
| **P27987** | ITPKB | -0.003922828 | 3.194084432 | -0.098579207 | 0.924455743 | 0.985099254 | -6.606254031 |
| **Q9NP61-1** | ARFGAP3 | -0.074395307 | 3.211633853 | -1.296558237 | 0.239097162 | 0.644815667 | -6.606526861 |
| **Q8N2K1-3** | UBE2J2 | -0.014075454 | 3.222044966 | -0.455503395 | 0.663634275 | 0.898182275 | -6.60691472 |
| **Q53FA7-1** | TP53I3 | 0.027202413 | 3.185151606 | 0.643658949 | 0.541923823 | 0.843570684 | -6.607207813 |
| **Q7L273** | KCTD9 | 0.031867663 | 3.170061469 | 0.510589615 | 0.626586539 | 0.885387512 | -6.607528794 |
| **P15923-2** | TCF3 | -0.00871576 | 3.213993988 | -0.305250337 | 0.76975768 | 0.941649022 | -6.608051314 |
| **Q53H47** | SETMAR | 0.025867272 | 3.188152577 | 1.197385073 | 0.273169321 | 0.672884265 | -6.608261409 |
| **Q15025** | TNIP1 | -0.019765356 | 3.181613496 | -0.785599614 | 0.459891995 | 0.800754306 | -6.608331667 |
| **P19388** | POLR2E | 0.023483138 | 3.19921869 | 0.660981945 | 0.531439772 | 0.839849063 | -6.608841759 |
| **Q96K37-1** | SLC35E1 | 0.00160193 | 3.201095661 | 0.044343936 | 0.965967237 | 0.992001409 | -6.609475893 |
| **Q9Y289** | SLC5A6 | -0.027731013 | 3.190951738 | -0.439289394 | 0.674743724 | 0.90314827 | -6.609683227 |
| **Q9H8W3** | FAM204A | -0.028228041 | 3.203491642 | -0.777031754 | 0.464589326 | 0.803074528 | -6.609721029 |
| **Q5C9Z4** | NOM1 | -0.02973299 | 3.182297767 | -1.082266815 | 0.317804773 | 0.705887314 | -6.609961092 |
| **O15160** | POLR1C | -0.03813457 | 3.189508977 | -1.040565825 | 0.335384117 | 0.720533987 | -6.610030707 |
| **Q96QR8** | PURB | 0.034744209 | 3.191837465 | 0.873236405 | 0.413760364 | 0.771310674 | -6.610545518 |
| **Q8N3X1-2** | FNBP4 | -0.030452778 | 3.181705827 | -1.18784931 | 0.276654709 | 0.675408632 | -6.610726685 |
| **Q86XK2-5** | FBXO11 | -0.004741354 | 3.176002068 | -0.099394281 | 0.923833486 | 0.985083485 | -6.610910353 |
| **Q9NY35-1** | CLDND1 | -0.012815162 | 3.159302025 | -0.26063176 | 0.802473635 | 0.953157309 | -6.610973961 |
| **Q2M1P5** | KIF7 | -0.023672952 | 3.206984511 | -0.848817527 | 0.426262584 | 0.779649091 | -6.612659895 |
| **Q8NCG7** | DAGLB | -0.003580387 | 3.213515356 | -0.071581406 | 0.945096559 | 0.98904859 | -6.613747763 |
| **P15814** | IGLL1 | 0.034728704 | 3.135707587 | 1.057621897 | 0.3281015 | 0.713117775 | -6.614068717 |
| **Q9BZ72** | PITPNM2 | 0.020834726 | 3.185680947 | 0.775746503 | 0.465296825 | 0.803299278 | -6.614304644 |
| **Q9UK97** | FBXO9 | -0.017776346 | 3.194392208 | -0.522231439 | 0.618900014 | 0.882318429 | -6.614676022 |
| **Q99487** | PAFAH2 | 0.004290267 | 3.141701548 | 0.11965357 | 0.908386685 | 0.980855153 | -6.614730514 |
| **Q96MH6-1** | TMEM68 | -0.019677593 | 3.213392348 | -0.355407954 | 0.733584004 | 0.928755685 | -6.614858325 |
| **Q96A72** | MAGOHB | 0.018828255 | 3.175282291 | 1.524948211 | 0.174549492 | 0.5748429 | -6.614971436 |
| **Q9Y3E7** | CHMP3; RNF103-C | 0.01007893 | 3.210737165 | 0.273472533 | 0.793010526 | 0.949070183 | -6.615271316 |
| **Q9NXR7-2** | BRE; BABAM2 | -0.025408994 | 3.174787761 | -0.731897899 | 0.48987922 | 0.818496205 | -6.615276429 |
| **Q9BZL1** | UBL5 | 0.070082211 | 3.199121362 | 0.9079467 | 0.396457657 | 0.762021989 | -6.615472253 |
| **A5PLN9-5** | C5orf44; TRAPPC1 | -0.019049749 | 3.217273528 | -0.493500711 | 0.637961443 | 0.889065271 | -6.615701796 |
| **O00505** | KPNA3 | 0.024104787 | 3.197399985 | 0.206289856 | 0.842903166 | 0.962543524 | -6.615727721 |
| **Q2TAL8** | QRICH1 | -0.026949607 | 3.19818202 | -1.212564966 | 0.267698053 | 0.67010832 | -6.615798688 |
| **P53634-1** | CTSC | -0.037580085 | 3.170601548 | -1.279199736 | 0.24477991 | 0.650150213 | -6.615802538 |
| **Q96E22** | NUS1 | 0.014349838 | 3.247676491 | 0.225424095 | 0.828601325 | 0.959096377 | -6.616234942 |
| **Q9UPY3** | DICER1 | -0.029086311 | 3.191470589 | -1.51644219 | 0.17663526 | 0.577967785 | -6.616249149 |
| **P33241** | LSP1 | -0.058918915 | 3.162342381 | -0.381730095 | 0.71488436 | 0.9235376 | -6.61635332 |
| **Q9H9Q2** | COPS7B | -0.049026751 | 3.167356276 | -1.14418618 | 0.293097546 | 0.68530312 | -6.61669542 |
| **Q9BQS8-4** | FYCO1 | 0.020826434 | 3.203903112 | 1.301302824 | 0.237564151 | 0.644090219 | -6.616777055 |
| **O00291** | HIP1 | 0.009804632 | 3.189392599 | 0.303309301 | 0.771170912 | 0.942318194 | -6.617131199 |

| | | | | | | | |
|---|---|---|---|---|---|---|---|
| **O00267-2** | SUPT5H | -0.02445189 | 3.189136651 | -1.6515781 | 0.146096332 | 0.544179725 | -6.61762556 |
| **Q9HCM1** | KIAA1551 | 0.030066029 | 3.191245393 | 1.008666356 | 0.349351285 | 0.730997093 | -6.620451001 |
| **Q9UHD9** | UBQLN2 | -0.028796225 | 3.169795834 | -0.821691951 | 0.440469604 | 0.788893373 | -6.620466728 |
| **Q9NVW2** | RLIM | -0.028458173 | 3.180048966 | -0.609884521 | 0.562729977 | 0.852576894 | -6.621153363 |
| **Q70IA6-3** | MOB2 | 0.026051843 | 3.150411869 | 0.501494156 | 0.632627262 | 0.887490806 | -6.621278872 |
| **Q96A47** | ISL2 | 0.017879738 | 3.160683938 | 0.603733676 | 0.566570403 | 0.853395233 | -6.621519409 |
| **Q14684** | RRP1B | -0.061900911 | 3.184607741 | -1.442191001 | 0.195841961 | 0.601527622 | -6.621577812 |
| **Q96GX5-1** | MASTL | 0.02080278 | 3.179269993 | 1.000369146 | 0.353058711 | 0.732738778 | -6.621874075 |
| **Q13426-1** | XRCC4 | -0.021290706 | 3.193808091 | -1.190434785 | 0.275705991 | 0.674825663 | -6.622365044 |
| **Q96BY9** | TMEM66; SARAF | 0.019581493 | 3.211034448 | 0.481153217 | 0.646247191 | 0.89263455 | -6.622404597 |
| **Q6PIJ6-2** | FBXO38 | -0.018624186 | 3.170160969 | -0.955727381 | 0.373537818 | 0.746151997 | -6.622881683 |
| **P14678-3** | SNRPB | -0.030821365 | 3.176531348 | -1.623374221 | 0.152027481 | 0.551195838 | -6.623033663 |
| **Q9Y6B6** | SAR1B | 0.012008442 | 3.193758087 | 0.284529536 | 0.78489225 | 0.94654395 | -6.623058029 |
| **Q7Z3B3** | KANSL1 | 0.029877052 | 3.180325643 | 1.340481245 | 0.225232593 | 0.630229362 | -6.623347052 |
| **P98196** | ATP11A | 0.014997795 | 3.171591622 | 0.611051151 | 0.562003332 | 0.852576894 | -6.623633382 |
| **Q5T2E6-1** | C10orf76 | -0.028336103 | 3.210512431 | -0.841091509 | 0.430274915 | 0.782631456 | -6.623931371 |
| **Q7Z6L1-2** | TECPR1 | 0.024245997 | 3.134989299 | 0.841599229 | 0.430010406 | 0.782603406 | -6.624157095 |
| **Q15154-1** | PCM1 | 0.013186186 | 3.190914724 | 0.268964187 | 0.796328757 | 0.950757673 | -6.62425428 |
| **Q9UET6** | FTSJ1 | -0.016119053 | 3.216448968 | -0.450937247 | 0.666753746 | 0.90001396 | -6.624402926 |
| **Q96C57** | C12orf43 | -0.020665686 | 3.179284213 | -0.964168174 | 0.369596649 | 0.74415747 | -6.624479159 |
| **Q13445** | TMED1 | 0.032096255 | 3.162080015 | 0.333113027 | 0.749578904 | 0.935635612 | -6.624526798 |
| **P26572** | MGAT1 | -0.008762758 | 3.205165414 | -0.25193082 | 0.808906564 | 0.955498404 | -6.624629538 |
| **P52758** | HRSP12; RIDA | -0.022960391 | 3.18872778 | -0.639998284 | 0.544155676 | 0.844397822 | -6.625022359 |
| **Q14142** | TRIM14 | -0.018595513 | 3.192174246 | -0.438533261 | 0.675264001 | 0.90314827 | -6.62533688 |
| **O60294** | LCMT2 | -0.022519274 | 3.175698456 | -0.615787013 | 0.559059374 | 0.851394811 | -6.625908357 |
| **Q9NVI1-3** | FANCI | 0.012446551 | 3.186677808 | 0.369378965 | 0.723633228 | 0.926648409 | -6.626097274 |
| **Q9UHY7** | ENOPH1 | 0.031918794 | 3.189745181 | 0.541271564 | 0.606440271 | 0.874996148 | -6.626859011 |
| **P09488** | GSTM1 | -0.022733435 | 3.170794179 | -1.053003779 | 0.33006066 | 0.715729292 | -6.627471662 |
| **Q9H1Z4** | WDR13 | 0.00623318 | 3.188326679 | 0.147350489 | 0.887340128 | 0.975708125 | -6.62773815 |
| **P0CG12** | CHTF8 | 0.000989908 | 3.135052826 | 0.02844802 | 0.978162059 | 0.993585863 | -6.627745977 |
| **Q13148-1** | TARDBP | 0.055455207 | 3.188738392 | 1.343574094 | 0.224283675 | 0.629881158 | -6.628205752 |
| **P61927** | RPL37 | -0.025935252 | 3.17570008 | -0.404961073 | 0.698555582 | 0.913150177 | -6.628261606 |
| **Q7Z628** | NET1 | 0.02073457 | 3.153038473 | 0.641379714 | 0.54331277 | 0.844078991 | -6.628291788 |
| **P30307-1** | CDC25C | -0.019846173 | 3.119859391 | -0.548602042 | 0.601680689 | 0.872497827 | -6.630196493 |
| **Q9BV68-2** | RNF126 | -0.017820646 | 3.190014419 | -0.5224973 | 0.618725078 | 0.882227341 | -6.630219402 |
| **Q15042-1** | RAB3GAP1 | 0.034698625 | 3.1783351 | 1.504920867 | 0.179497194 | 0.580990067 | -6.63118076 |
| **O15127** | SCAMP2 | -0.014791954 | 3.177028433 | -0.345605326 | 0.740599474 | 0.931611328 | -6.631705732 |
| **Q16773** | CCBL1; KYAT1 | -0.004327678 | 3.195848996 | -0.139167174 | 0.893549077 | 0.978276697 | -6.63188508 |
| **O95163** | IKBKAP; ELP1 | 0.021011692 | 3.171417877 | 1.548215474 | 0.168960291 | 0.567606558 | -6.631887472 |

| | | | | | | | |
|---|---|---|---|---|---|---|---|
| **O95881** | TXNDC12 | -0.022451413 | 3.182598284 | -1.247247626 | 0.255548441 | 0.659782078 | -6.632622631 |
| **P28482** | MAPK1 | -0.054925244 | 3.197328557 | -1.438037127 | 0.196970894 | 0.603701501 | -6.632992049 |
| **Q9Y508-1** | RNF114 | -0.053576615 | 3.180233662 | -1.13176694 | 0.297921217 | 0.689670498 | -6.633189571 |
| **Q96GD4-5** | AURKB | 0.023301816 | 3.184615149 | 0.949830193 | 0.376310491 | 0.748944602 | -6.633194168 |
| **Q96E39** | RBMXL1 | 0.016548309 | 3.179096016 | 0.332516446 | 0.750008799 | 0.935635612 | -6.633558196 |
| **Q8TF76-1** | GSG2; HASPIN | -0.007825038 | 3.226809915 | -0.225832878 | 0.828296531 | 0.959077751 | -6.633697569 |
| **Q9Y4E5** | ZNF451 | 0.027440532 | 3.18394977 | 1.2384887 | 0.258571077 | 0.661376267 | -6.634106399 |
| **Q9BRA0-2** | LSMD1; NAA38 | 0.00235471 | 3.152632835 | 0.066222671 | 0.949199537 | 0.989472196 | -6.634345118 |
| **Q9HCJ3** | RAVER2 | -0.026363055 | 3.211647812 | -0.27609617 | 0.791081621 | 0.948368382 | -6.635133425 |
| **O75340** | PDCD6 | -0.030789628 | 3.195050508 | -1.032488575 | 0.3388779 | 0.721920542 | -6.635381251 |
| **Q7L8J4** | SH3BP5L | 0.021270046 | 3.164439639 | 0.388521888 | 0.710093142 | 0.92064132 | -6.635619209 |
| **Q86V88** | MDP1 | -0.020520867 | 3.181666151 | -0.533821815 | 0.61129871 | 0.878099485 | -6.636001987 |
| **P78318** | IGBP1 | -0.029025467 | 3.205714025 | -0.715632891 | 0.499216012 | 0.824970816 | -6.636042308 |
| **P27707** | DCK | 0.092581421 | 3.179567733 | 1.51322211 | 0.177430861 | 0.578452275 | -6.636073643 |
| **P40337** | VHL | 0.005857996 | 3.209781883 | 0.109758889 | 0.915925998 | 0.982975929 | -6.636335835 |
| **Q13637** | RAB32 | -0.021162029 | 3.194217724 | -1.063873693 | 0.325464247 | 0.710703247 | -6.636608359 |
| **Q6N043** | ZNF280D | 0.027354415 | 3.190910311 | 1.007087687 | 0.350054302 | 0.731407657 | -6.636853301 |
| **P17252** | PRKCA | 0.018698151 | 3.175332798 | 1.182911141 | 0.278474417 | 0.676427621 | -6.637201625 |
| **Q8TBF4** | ZCRB1 | 0.01244365 | 3.179545521 | 0.314291802 | 0.763187393 | 0.939644172 | -6.63732084 |
| **Q9NXH9-1** | TRMT1 | -0.023794051 | 3.189417053 | -1.195555931 | 0.273834979 | 0.673600291 | -6.63759331 |
| **P62834** | RAP1A | 0.034254248 | 3.1569246 | 1.038717372 | 0.336181103 | 0.720858943 | -6.637803716 |
| **Q5SY16** | NOL9 | -0.084637509 | 3.154705788 | -0.850661751 | 0.425308856 | 0.778986255 | -6.637935738 |
| **Q9NVM6** | DNAJC17 | 0.019360662 | 3.175492741 | 0.395960873 | 0.704861717 | 0.916550956 | -6.637952634 |
| **Q96MW5** | COG8 | -0.020369009 | 3.184932811 | -0.795290725 | 0.454618867 | 0.797515322 | -6.638512774 |
| **Q8WUF5** | PPP1R13L | 0.004549756 | 3.213433924 | 0.143944787 | 0.889923126 | 0.976984833 | -6.639000574 |
| **Q969R8** | ITFG2 | -0.027177824 | 3.187049662 | -0.661537278 | 0.531105807 | 0.839849063 | -6.639089463 |
| **O75718** | CRTAP | 0.028178887 | 3.198548519 | 1.021758136 | 0.343564176 | 0.72497442 | -6.63924609 |
| **Q9NYL2** | ZAK; MAP3K20 | 0.004692709 | 3.196529576 | 0.134831046 | 0.89684237 | 0.978757548 | -6.639842109 |
| **Q96C01** | FAM136A | 0.085089273 | 3.171861636 | 1.121682364 | 0.301886478 | 0.691344519 | -6.639971506 |
| **Q5QJ74** | TBCEL | 0.012264744 | 3.089445895 | 0.320006015 | 0.759045866 | 0.9384213 | -6.640197851 |
| **O15360** | FANCA | 0.000909858 | 3.201810577 | 0.022507501 | 0.982721254 | 0.994932161 | -6.640443473 |
| **Q9H8U3** | ZFAND3 | -0.02406795 | 3.17464543 | -0.658894358 | 0.532696391 | 0.839951983 | -6.640515758 |
| **P26374** | CHML | 0.020238773 | 3.191879824 | 1.070000589 | 0.322896351 | 0.710272801 | -6.640637509 |
| **O14795-2** | UNC13B | -0.011756521 | 3.203098365 | -0.412988835 | 0.692952579 | 0.909991705 | -6.640895139 |
| **Q06547-3** | GABPB1 | 0.009611961 | 3.226676199 | 0.180484654 | 0.862292929 | 0.969286548 | -6.641810435 |
| **P78346-2** | RPP30 | 0.021204285 | 3.169680762 | 0.962807544 | 0.370229772 | 0.744424726 | -6.641921697 |
| **P10244** | MYBL2 | 0.020702087 | 3.162282321 | 1.251545509 | 0.254076472 | 0.659039362 | -6.642016614 |
| **P48736** | PIK3CG | 0.026704627 | 3.149225691 | 1.280379755 | 0.244389896 | 0.650150213 | -6.642028343 |
| **Q8TBC5-3** | ZSCAN18 | 0.007887845 | 3.189766263 | 0.214143818 | 0.837024569 | 0.960912428 | -6.642410768 |

| | | | | | | | |
|---|---|---|---|---|---|---|---|
| **Q7RTP6** | MICAL3 | -0.018465264 | 3.222310018 | -0.872389096 | 0.414189617 | 0.771567763 | -6.642519638 |
| **Q8NEM0-3** | MCPH1 | 0.015745907 | 3.225269273 | 0.310772876 | 0.765742049 | 0.940484723 | -6.642577453 |
| **P29762** | CRABP1 | 0.024769785 | 3.159514193 | 1.178289659 | 0.280186592 | 0.677109587 | -6.642688771 |
| **P55209** | NAP1L1 | 0.026417332 | 3.186237861 | 0.312597782 | 0.764416812 | 0.940369694 | -6.643282167 |
| **Q9H8H2** | DDX31 | -0.053768641 | 3.174114353 | -1.231326042 | 0.261065705 | 0.664371245 | -6.643419057 |
| **Q9BTC8** | MTA3 | 0.029612648 | 3.194630508 | 0.825582917 | 0.43841111 | 0.786897076 | -6.643445956 |
| **Q9ULH0-1** | KIDINS220 | -0.021785217 | 3.220614246 | -0.985075359 | 0.359973416 | 0.73661838 | -6.643667517 |
| **O15440** | ABCC5 | 0.013475971 | 3.18404999 | 0.378758818 | 0.716984852 | 0.924679646 | -6.64453421 |
| **P35610** | SOAT1 | 0.015620056 | 3.196979668 | 0.583336215 | 0.579417374 | 0.862073227 | -6.644716852 |
| **O15270** | SPTLC2 | -0.023334771 | 3.190642472 | -1.1468663 | 0.29206517 | 0.685264687 | -6.644771859 |
| **P14859-6** | POU2F1 | -0.029694409 | 3.166270397 | -0.696394041 | 0.510410881 | 0.829456831 | -6.645269527 |
| **Q9BRA2** | TXNDC17 | 0.018920625 | 3.18072276 | 0.590566414 | 0.574844097 | 0.859195843 | -6.645505492 |
| **Q96MN5** | TCEANC2 | -0.034696693 | 3.147473411 | -0.498360661 | 0.634715492 | 0.887591411 | -6.645589522 |
| **Q9UKA9-3** | PTBP2 | 0.001228765 | 3.162714984 | 0.039270118 | 0.969858795 | 0.992476747 | -6.645835386 |
| **P57088** | TMEM33 | -0.021037444 | 3.192494141 | -0.898608244 | 0.401058695 | 0.764494082 | -6.645996513 |
| **P04818** | TYMS | -0.035443164 | 3.205462612 | -1.23951223 | 0.258216281 | 0.660717666 | -6.646152027 |
| **Q8NB78-1** | KDM1B | 0.021469021 | 3.215331787 | 1.063973336 | 0.325422353 | 0.710703247 | -6.646203278 |
| **Q9UPS6-1** | SETD1B | 0.002193518 | 3.205982603 | 0.073908399 | 0.943315427 | 0.988738927 | -6.646342635 |
| **Q8IXS8** | FAM126B | 0.024460449 | 3.171809465 | 0.58648304 | 0.577424316 | 0.861313941 | -6.64686103 |
| **Q9Y664** | KPTN | 0.00863186 | 3.170005678 | 0.289329384 | 0.781377095 | 0.944918194 | -6.647706238 |
| **Q96QU8** | XPO6 | 0.020965214 | 3.19892226 | 0.869787485 | 0.415509662 | 0.77203561 | -6.648197338 |
| **O94952-2** | FBXO21 | -0.03192683 | 3.150522437 | -0.310672149 | 0.765815222 | 0.940484723 | -6.648569372 |
| **Q16576-2** | RBBP7 | -0.032850195 | 3.200949221 | -1.39901989 | 0.207866682 | 0.614231901 | -6.648999994 |
| **Q8N6R0-5** | METTL13 | 0.03642802 | 3.194288591 | 1.147580911 | 0.291790417 | 0.685264687 | -6.649068338 |
| **P52747** | ZNF143 | 0.018234044 | 3.195583501 | 0.654926808 | 0.535089805 | 0.840674016 | -6.64959376 |
| **P11831** | SRF | 0.024390595 | 3.209704959 | 0.268398146 | 0.7967457 | 0.950757673 | -6.649755194 |
| **Q9NRF9** | POLE3 | -0.013121383 | 3.191330439 | -0.398848873 | 0.702835406 | 0.915414312 | -6.649793954 |
| **Q96B26** | EXOSC8 | -0.028531804 | 3.17719643 | -0.9498881 | 0.376283188 | 0.748944602 | -6.650287391 |
| **Q9UII2-1** | ATPIF1 | 0.053635151 | 3.175551516 | 0.649690868 | 0.53825869 | 0.843524953 | -6.65130128 |
| **Q15904** | ATP6AP1 | 0.008158417 | 3.201342923 | 0.184580724 | 0.859207875 | 0.968855822 | -6.651332013 |
| **Q9GZN7** | ROGDI | 0.01421826 | 3.185052 | 0.565044179 | 0.591081899 | 0.868764508 | -6.651414116 |
| **Q86U70** | LDB1 | -0.020705398 | 3.188833239 | -0.558461925 | 0.595312055 | 0.871378088 | -6.6515673 |
| **Q9H1U4** | MEGF9 | 0.013875954 | 3.155586135 | 0.228692927 | 0.826164942 | 0.959077751 | -6.651794202 |
| **Q9Y620-1** | RAD54B | 0.020252673 | 3.18633603 | 1.090004675 | 0.314626662 | 0.701837878 | -6.652855892 |
| **Q01415-1** | GALK2 | 0.02974548 | 3.171346868 | 0.975901444 | 0.364171714 | 0.739413523 | -6.653210468 |
| **Q86TP1-1** | PRUNE; PRUNE1 | 0.021221629 | 3.162275101 | 0.704493916 | 0.505677793 | 0.82913633 | -6.653404753 |
| **Q9NU19** | TBC1D22B | 0.020346605 | 3.165459358 | 0.700058975 | 0.508265741 | 0.82913633 | -6.653987552 |
| **Q9UL33-1** | TRAPPC2L | -0.010884802 | 3.24435323 | -0.230534682 | 0.824793129 | 0.959077751 | -6.654016159 |
| **Q68DH5** | LMBRD2 | -0.001628283 | 3.181895356 | -0.047191562 | 0.963783577 | 0.991359466 | -6.654086468 |

| | | | | | | | |
|---|---|---|---|---|---|---|---|
| **Q9Y2R2-1** | PTPN22 | -0.024601831 | 3.209704474 | -0.857592651 | 0.421738468 | 0.777315429 | -6.65420003 |
| **P27701** | CD82 | -0.042663004 | 3.191687096 | -0.45432977 | 0.664435374 | 0.898389899 | -6.654374727 |
| **Q8WUD4** | CCDC12 | 0.100182737 | 3.176843534 | 1.206590109 | 0.269840305 | 0.671731232 | -6.655241828 |
| **Q13418** | ILK | -0.028188944 | 3.178880768 | -1.393889372 | 0.209339199 | 0.614302382 | -6.656190693 |
| **Q9BZK7** | TBL1XR1 | -0.028141969 | 3.174349021 | -1.220786174 | 0.264774161 | 0.66823064 | -6.656342786 |
| **Q99622** | C12orf57 | 0.023780713 | 3.310207015 | 0.256162952 | 0.805775544 | 0.954293355 | -6.657323015 |
| **Q6NXT6** | TAPT1 | 0.015522861 | 3.184213665 | 0.660537494 | 0.531707152 | 0.839849063 | -6.657811044 |
| **Q96JQ2** | CLMN | 0.009760407 | 3.222553175 | 0.252567499 | 0.808435291 | 0.955329057 | -6.657830837 |
| **Q8N4C6-7** | NIN | 0.020228665 | 3.194768233 | 1.071699269 | 0.322187319 | 0.7096696 | -6.658079577 |
| **Q6NUQ1** | RINT1 | -0.016751151 | 3.189701732 | -1.026726884 | 0.341387814 | 0.723479909 | -6.65821952 |
| **Q15361** | TTF1 | -0.054758935 | 3.184212232 | -1.036841992 | 0.336991248 | 0.721128781 | -6.658792676 |
| **O00628** | PEX7 | -0.011665805 | 3.210440543 | -0.257365555 | 0.804886537 | 0.954249678 | -6.658881535 |
| **O75792** | RNASEH2A | -0.051520131 | 3.184761783 | -1.242731041 | 0.257103249 | 0.659782078 | -6.659524061 |
| **Q9C0K1-1** | SLC39A8 | 0.029659155 | 3.166560244 | 0.296802874 | 0.775915002 | 0.94357883 | -6.65977335 |
| **P07108-5** | DBI | -0.497564587 | 3.163987612 | -1.325274945 | 0.229950008 | 0.637461728 | -6.659833046 |
| **Q8WUW1-2** | BRK1 | -0.013253484 | 3.157691402 | -0.234359731 | 0.821946215 | 0.959068023 | -6.660176743 |
| **Q8N5F7** | NKAP | 0.012405928 | 3.170375833 | 0.418736342 | 0.688953882 | 0.908052188 | -6.661355973 |
| **Q99523** | SORT1 | -0.029970588 | 3.204233915 | -0.962410606 | 0.370414631 | 0.744424726 | -6.661363082 |
| **Q9UEG4** | ZNF629 | 0.009464587 | 3.147822194 | 0.283070699 | 0.785961714 | 0.946704945 | -6.661966974 |
| **P42898-2** | MTHFR | -0.029932745 | 3.1711655 | -1.220479879 | 0.264882603 | 0.66823064 | -6.662076242 |
| **Q5TFE4** | NT5DC1 | -0.023638346 | 3.182871348 | -1.257943283 | 0.251898907 | 0.656176006 | -6.662159931 |
| **Q9C0K0-1** | BCL11B | -0.038726735 | 3.201855274 | -1.360415151 | 0.219178784 | 0.625602181 | -6.66263118 |
| **Q6PII3** | CCDC174 | 0.015412686 | 3.198640159 | 0.747580139 | 0.480988461 | 0.81235548 | -6.66366716 |
| **O14907** | TAX1BP3 | 0.007568118 | 3.141365143 | 0.247728469 | 0.812019332 | 0.955776828 | -6.663748036 |
| **Q9NQS7** | INCENP | -0.030845094 | 3.163482449 | -1.165801191 | 0.284857857 | 0.68101215 | -6.664019295 |
| **Q9Y6X5** | ENPP4 | 0.032602529 | 3.169999242 | 0.868333375 | 0.416248818 | 0.772686182 | -6.664446722 |
| **Q96JN8-1** | NEURL4 | -0.010162718 | 3.175950861 | -0.278553734 | 0.789276259 | 0.947865984 | -6.664482664 |
| **Q8TB24-1** | RIN3 | 0.004579231 | 3.194311262 | 0.136420474 | 0.895634937 | 0.978757548 | -6.664539153 |
| **Q96GQ7** | DDX27 | -0.0424675 | 3.189731306 | -1.526463875 | 0.174180239 | 0.574454961 | -6.66507244 |
| **Q86Y82** | STX12 | 0.014860722 | 3.174699001 | 0.477656166 | 0.648603921 | 0.893481222 | -6.665103819 |
| **P49840** | GSK3A | 0.036336348 | 3.180955271 | 0.813939626 | 0.444591471 | 0.790803649 | -6.665493826 |
| **Q5SNT2-1** | TMEM201 | 0.025128752 | 3.24095546 | 0.888213005 | 0.406227222 | 0.766336431 | -6.66585411 |
| **Q96GQ5-1** | C16orf58 | -0.021889116 | 3.192022053 | -0.507409307 | 0.628695218 | 0.886106708 | -6.666013596 |
| **Q8WXF1** | PSPC1 | 0.047001163 | 3.150990571 | 1.35401892 | 0.221105304 | 0.626491311 | -6.666514199 |
| **Q9P1U1-1** | ACTR3B | 0.009077877 | 3.172599777 | 0.180909998 | 0.861972448 | 0.969063407 | -6.666614359 |
| **Q9Y2P8** | RCL1 | 0.050760765 | 3.171796551 | 1.079597144 | 0.318907324 | 0.707150066 | -6.666817029 |
| **Q4KMP7** | TBC1D10B | -0.024552436 | 3.183012631 | -0.710584452 | 0.502137856 | 0.826806835 | -6.66721986 |
| **Q9UKK6** | NXT1 | -0.027013118 | 3.203551145 | -0.463254514 | 0.658355544 | 0.896852471 | -6.667637509 |
| **Q68DI1** | ZNF776 | -0.014431613 | 3.207760413 | -0.487932198 | 0.641691301 | 0.890911725 | -6.668946269 |

| | | | | | | | |
|---|---|---|---|---|---|---|---|
| **O00750** | PIK3C2B | -0.014207993 | 3.166738871 | -0.458721929 | 0.661439812 | 0.896934512 | -6.669092531 |
| **Q969G3-2** | SMARCE1 | 0.040468119 | 3.167761848 | 1.282785708 | 0.243596371 | 0.649672095 | -6.669176169 |
| **Q8N806** | UBR7 | 0.022936188 | 3.174473913 | 1.403549245 | 0.206574468 | 0.612058044 | -6.669339332 |
| **Q8IUH4** | ZDHHC13 | -0.031396599 | 3.182047874 | -0.760122616 | 0.473956935 | 0.808788313 | -6.669446701 |
| **Q8NBA8** | DTWD2 | 0.006837151 | 3.19359303 | 0.132652295 | 0.898497975 | 0.978757548 | -6.669569068 |
| **Q9HA64** | FN3KRP | 0.092216554 | 3.212637312 | 0.589573993 | 0.575470567 | 0.859904769 | -6.670094381 |
| **Q9NPF2-1** | CHST11 | -0.002908797 | 3.199503004 | -0.101210252 | 0.922447318 | 0.984164603 | -6.670890177 |
| **Q14674-1** | ESPL1 | -0.016449745 | 3.144789648 | -0.919231901 | 0.390950539 | 0.759806823 | -6.671239133 |
| **P59768** | GNG2 | 0.047720603 | 3.200065518 | 0.42461616 | 0.684874312 | 0.905978473 | -6.673862178 |
| **Q96BJ3-1** | AIDA | 0.016226523 | 3.150047473 | 0.685783775 | 0.516654388 | 0.833280927 | -6.674854452 |
| **Q8WUH1** | CHURC1 | -0.024114432 | 3.198541313 | -0.484353818 | 0.644094114 | 0.891474831 | -6.674927363 |
| **Q9Y5M8** | SRPRB | -0.024716181 | 3.207602941 | -1.147595837 | 0.29178468 | 0.685264687 | -6.675112886 |
| **O43399-3** | TPD52L2 | 0.062474917 | 3.212990913 | 0.778605474 | 0.463724047 | 0.802800142 | -6.67615761 |
| **Q13542** | EIF4EBP2 | -0.149494898 | 3.191190763 | -0.526912779 | 0.615823659 | 0.880583268 | -6.67624374 |
| **Q10471** | GALNT2 | -0.059257312 | 3.149981383 | -1.589872248 | 0.159368351 | 0.558776357 | -6.676369778 |
| **Q9UK59** | DBR1 | 0.035835286 | 3.167198803 | 1.429412455 | 0.199333809 | 0.604614147 | -6.676489991 |
| **Q53ET0** | CRTC2 | 0.006999471 | 3.180233598 | 0.209170483 | 0.840745761 | 0.961788885 | -6.677301431 |
| **Q6WKZ4-4** | RAB11FIP1 | -0.025407825 | 3.188692318 | -1.187379583 | 0.276827368 | 0.675408632 | -6.677615546 |
| **Q8TAS1** | UHMK1 | 0.012389518 | 3.208965263 | 0.361617077 | 0.729154556 | 0.927643386 | -6.677756933 |
| **Q5T011** | SZT2 | 0.00094553 | 3.182908104 | 0.030485137 | 0.97659882 | 0.99316605 | -6.678485734 |
| **P55210-3** | CASP7 | 0.024741744 | 3.18802189 | 0.948601643 | 0.376890103 | 0.749159836 | -6.678671743 |
| **Q9UQ80** | PA2G4 | 0.031536319 | 3.168753773 | 1.159008732 | 0.287425957 | 0.682522792 | -6.678927595 |
| **Q8IZQ5** | C11orf31; SELENO | -0.00241156 | 3.168729318 | -0.081958238 | 0.937156697 | 0.987232823 | -6.679806068 |
| **Q53H96** | PYCRL; PYCR3 | -0.01484739 | 3.178273338 | -0.585853138 | 0.577822946 | 0.861313941 | -6.679817696 |
| **Q66GS9-1** | CEP135 | 0.021311468 | 3.173183872 | 1.397831747 | 0.20820686 | 0.614231901 | -6.68074752 |
| **Q9NRX1** | PNO1 | -0.028149415 | 3.179049893 | -0.959299538 | 0.37186597 | 0.744950865 | -6.680929282 |
| **Q9NVM4** | PRMT7 | -0.029892478 | 3.16251381 | -0.841719446 | 0.429947793 | 0.782603406 | -6.680970958 |
| **Q9P253** | VPS18 | -0.028205792 | 3.193716694 | -1.031191769 | 0.339441528 | 0.722416219 | -6.681331493 |
| **Q9P021** | CRIPT | -0.000155271 | 3.173893869 | -0.004203747 | 0.996772529 | 0.998480498 | -6.681461356 |
| **Q6NXT1-1** | ANKRD54 | -0.009409251 | 3.181009924 | -0.234082725 | 0.822152288 | 0.959068023 | -6.681843107 |
| **Q9HCE0-1** | EPG5 | -0.003668988 | 3.189511063 | -0.096801484 | 0.92581312 | 0.985476218 | -6.682843067 |
| **Q5T0N5-4** | FNBP1L | -0.024500827 | 3.178700238 | -1.00170026 | 0.352461853 | 0.732738778 | -6.684350886 |
| **O75478** | TADA2A | 0.016483699 | 3.18773273 | 0.523625607 | 0.617982958 | 0.881802191 | -6.684440785 |
| **Q96JG6** | CCDC132; VPS50 | 0.033801973 | 3.195061726 | 1.41502575 | 0.203332646 | 0.608695713 | -6.68498592 |
| **A0PJZ3** | GXYLT2 | -0.01480476 | 3.19641968 | -0.321153238 | 0.758215418 | 0.9384213 | -6.685202723 |
| **Q6GMV2** | SMYD5 | 0.032688577 | 3.192151232 | 0.891133651 | 0.404770089 | 0.766336431 | -6.685215712 |
| **Q9UJY5-6** | GGA1 | -0.040829267 | 3.167449258 | -0.744457883 | 0.482749808 | 0.812781066 | -6.685422006 |
| **Q5XUX1-3** | FBXW9 | -0.004249075 | 3.153818547 | -0.128770968 | 0.901448705 | 0.97952714 | -6.686460846 |
| **P27816-4** | MAP4 | -0.026372747 | 3.201940774 | -0.974779283 | 0.36468786 | 0.739899144 | -6.686909193 |

| | | | | | | | |
|---|---|---|---|---|---|---|---|
| **O95219** | SNX4 | 0.015143351 | 3.19863911 | 0.250361659 | 0.810068437 | 0.955498404 | -6.687469296 |
| **Q9NUL5** | C19orf66 | 0.004691128 | 3.176136242 | 0.089162778 | 0.931648643 | 0.986711787 | -6.68823559 |
| **Q8NBR6-1** | FAM63B; MINDY2 | 0.001160071 | 3.175419245 | 0.027080444 | 0.979211567 | 0.993720258 | -6.688323327 |
| **O95159** | ZFPL1 | 0.017622814 | 3.178845899 | 0.524863836 | 0.617169098 | 0.881353777 | -6.688714891 |
| **Q8TAA9** | VANGL1 | -0.020438741 | 3.20068535 | -0.623537674 | 0.554261509 | 0.848970058 | -6.68922608 |
| **Q6P158-1** | DHX57 | -0.033109011 | 3.185397857 | -1.331546981 | 0.227993751 | 0.634914248 | -6.689534933 |
| **Q9H9Q4-1** | NHEJ1 | 0.01476241 | 3.201121003 | 0.379918635 | 0.716164621 | 0.924439954 | -6.689582547 |
| **Q8IV04-1** | TBC1D10C | -0.026741137 | 3.178471966 | -0.644751155 | 0.541259024 | 0.843540088 | -6.689902446 |
| **Q16342-1** | PDCD2 | -0.028708927 | 3.2033947 | -0.743808843 | 0.483116496 | 0.813039729 | -6.689955392 |
| **Q9H6K1** | C6orf106 | -0.022035926 | 3.206596733 | -0.474805771 | 0.650528109 | 0.893885477 | -6.690381746 |
| **Q9NZR1** | TMOD2 | 0.002354985 | 3.180567143 | 0.080577343 | 0.938212865 | 0.987739899 | -6.690420003 |
| **Q5JTD0-1** | TJAP1 | -0.016398707 | 3.178084392 | -0.215097734 | 0.836311338 | 0.960912428 | -6.690530352 |
| **Q9P2M7-1** | CGN | -0.026495161 | 3.200835585 | -0.8519364 | 0.424650587 | 0.778986255 | -6.690553345 |
| **O15078-1** | CEP290 | 0.019733442 | 3.161489113 | 1.145183577 | 0.292712995 | 0.68530312 | -6.690925747 |
| **Q13496** | MTM1 | 0.024155462 | 3.182982492 | 0.687436967 | 0.515678348 | 0.832736682 | -6.69100237 |
| **O00399** | DCTN6 | -0.008032114 | 3.172367477 | -0.171665849 | 0.868943847 | 0.971129394 | -6.69109958 |
| **Q03188** | CENPC; CENPC1 | -0.022299955 | 3.175404608 | -1.135445398 | 0.296485654 | 0.688880164 | -6.691664843 |
| **Q2KHR3-1** | QSER1 | -0.022920217 | 3.210352019 | -1.109348714 | 0.306795358 | 0.695431988 | -6.69179531 |
| **Q92804-1** | TAF15 | -0.039461763 | 3.178783935 | -0.992633901 | 0.356542863 | 0.734600528 | -6.691972162 |
| **Q96QG7** | MTMR9 | -0.021688638 | 3.178862982 | -0.779709971 | 0.463117429 | 0.802800142 | -6.692162431 |
| **A6H8Y1** | BDP1 | 0.018179818 | 3.214469697 | 0.684818917 | 0.517224588 | 0.833280927 | -6.692555945 |
| **Q86VI3** | IQGAP3 | 0.041235975 | 3.196818474 | 0.75991068 | 0.474075163 | 0.808788313 | -6.69289786 |
| **Q9H211** | CDT1 | -0.016902516 | 3.229453551 | -0.422105082 | 0.686615172 | 0.907227265 | -6.692900829 |
| **Q9BRK5-1** | SDF4 | -0.008880357 | 3.231370709 | -0.341437627 | 0.743590397 | 0.933895938 | -6.69345408 |
| **O95402-1** | MED26 | -0.006982156 | 3.214201434 | -0.152254692 | 0.883623244 | 0.975295424 | -6.693826825 |
| **Q71SY5-4** | MED25 | 0.017620602 | 3.161048329 | 0.663588822 | 0.529873201 | 0.839849063 | -6.693894963 |
| **Q00765** | REEP5 | 0.005055688 | 3.203121921 | 0.067156935 | 0.948484082 | 0.989472196 | -6.693948041 |
| **Q9NWZ8** | GEMIN8 | 0.023249843 | 3.212243869 | 0.211198362 | 0.839227902 | 0.961364456 | -6.694580861 |
| **P21964-1** | COMT | 0.028061968 | 3.195818881 | 1.012659684 | 0.347577948 | 0.729399576 | -6.694667112 |
| **Q8N9F7** | GDPD1 | -0.001450479 | 3.202207222 | -0.046417742 | 0.964376936 | 0.991584672 | -6.694789373 |
| **Q9H269** | VPS16 | -0.028814573 | 3.182065324 | -1.017505857 | 0.345435462 | 0.726438525 | -6.694803512 |
| **Q8N2W9** | PIAS4 | 0.014542652 | 3.203458748 | 0.408414936 | 0.69614241 | 0.912062672 | -6.694984474 |
| **Q9NR77** | PXMP2 | -0.038409996 | 3.196241999 | -0.692669277 | 0.512597073 | 0.831227449 | -6.695259494 |
| **Q9UNE7-1** | STUB1 | -0.035097126 | 3.202681108 | -0.70290789 | 0.506602303 | 0.82913633 | -6.695683981 |
| **P13489** | RNH1 | -0.033952892 | 3.175067726 | -1.307617252 | 0.235537303 | 0.641994289 | -6.695879308 |
| **O60828** | PQBP1 | -0.025435743 | 3.193105243 | -1.121537765 | 0.301943651 | 0.691344519 | -6.696075571 |
| **P30154-2** | PPP2R1B | -0.025988689 | 3.168300784 | -1.144539356 | 0.292961329 | 0.68530312 | -6.696620046 |
| **Q9H497-1** | TOR3A | 0.012401381 | 3.188545381 | 0.431928547 | 0.67981674 | 0.903893384 | -6.696929321 |
| **P34913-1** | EPHX2 | 0.002082611 | 3.166255114 | 0.059770979 | 0.954141567 | 0.989472196 | -6.697056148 |

| | | | | | | | |
|---|---|---|---|---|---|---|---|
| **Q13459-1** | MYO9B | -0.025906355 | 3.169587547 | -1.574423362 | 0.162864577 | 0.561580888 | -6.697427317 |
| **Q99698-1** | LYST | 0.01179182 | 3.164151225 | 0.270792364 | 0.794982624 | 0.949994286 | -6.697507311 |
| **Q96I36** | COX14 | -0.016677931 | 3.208073515 | -0.325081174 | 0.755374716 | 0.937287539 | -6.697598454 |
| **P26196** | DDX6 | 0.018586513 | 3.173967655 | 1.565830162 | 0.164840264 | 0.563893388 | -6.697710922 |
| **Q8N1F8** | STK11IP | 0.014703428 | 3.205699543 | 0.414165263 | 0.692133232 | 0.909991705 | -6.697790229 |
| **Q9HC07** | TMEM165 | -0.009486435 | 3.160311676 | -0.315853618 | 0.762054579 | 0.939498834 | -6.698271245 |
| **Q9P2Y5** | UVRAG | -0.024464154 | 3.177224678 | -0.518034484 | 0.621665146 | 0.883385606 | -6.698457154 |
| **Q6NZY4** | ZCCHC8 | -0.039613426 | 3.187373893 | -1.48725478 | 0.18396871 | 0.586529454 | -6.699142504 |
| **Q13557** | CAMK2D | -0.013532658 | 3.170115844 | -0.369870762 | 0.723283993 | 0.926648409 | -6.699515576 |
| **P62879** | GNB2 | -0.018638895 | 3.160182584 | -0.752295035 | 0.478336934 | 0.811328097 | -6.699711335 |
| **O94953** | KDM4B | -0.068621195 | 3.163092961 | -0.829586253 | 0.436300363 | 0.785483221 | -6.70036226 |
| **O43913** | ORC5 | -0.025475121 | 3.195832413 | -1.069568144 | 0.323077056 | 0.710272801 | -6.700418729 |
| **Q9Y639-2** | NPTN | 0.02198236 | 3.176018272 | 0.952061948 | 0.375259335 | 0.74816447 | -6.700501721 |
| **Q15404-1** | RSU1 | -0.015567452 | 3.171788582 | -0.463937548 | 0.657891388 | 0.896852471 | -6.700530864 |
| **O94901-9** | SUN1 | 0.003033789 | 3.241294074 | 0.056675346 | 0.956513634 | 0.989472196 | -6.700830933 |
| **Q9BYT8** | NLN | 0.03621375 | 3.184384916 | 1.351000089 | 0.222019793 | 0.627993355 | -6.701092425 |
| **P51665** | PSMD7 | 0.019744994 | 3.172844773 | 1.197265567 | 0.273212769 | 0.672884265 | -6.701361376 |
| **Q92544** | TM9SF4 | -0.035599026 | 3.189548803 | -1.051690194 | 0.330619646 | 0.716489955 | -6.701385865 |
| **Q9H9B1** | EHMT1 | -0.028409325 | 3.193199474 | -1.410223247 | 0.204683597 | 0.610527836 | -6.702399834 |
| **Q8WTW4** | NPRL2 | 0.007911798 | 3.204913955 | 0.244515386 | 0.814401833 | 0.95620584 | -6.702924802 |
| **Q13889** | GTF2H3 | -0.020383302 | 3.189609644 | -0.894558771 | 0.403066232 | 0.766336431 | -6.702955229 |
| **Q9HD15** | SRA1 | 0.018559496 | 3.186922442 | 0.486567929 | 0.642606832 | 0.891449441 | -6.703198571 |
| **P46109** | CRKL | -0.021510197 | 3.170772947 | -1.350397171 | 0.222202837 | 0.627993355 | -6.703277315 |
| **Q7Z624** | CAMKMT | 0.007239723 | 3.19085884 | 0.25924971 | 0.803494337 | 0.953796878 | -6.703323327 |
| **P13987** | CD59 | 0.009201066 | 3.17698767 | 0.114844496 | 0.912049757 | 0.981900848 | -6.703496557 |
| **P11171-1** | EPB41 | -0.022722842 | 3.208081369 | -0.574279255 | 0.585176023 | 0.8653224 | -6.703781489 |
| **Q8WVC6-1** | DCAKD | 0.013653401 | 3.194537362 | 0.499639711 | 0.633862666 | 0.887509451 | -6.703930297 |
| **P49903-1** | SEPHS1 | -0.030306291 | 3.16101616 | -0.727336178 | 0.492485987 | 0.820752517 | -6.703966786 |
| **Q9NY97-1** | B3GNT2 | -0.01996248 | 3.202580032 | -0.71421197 | 0.500037248 | 0.825209342 | -6.705229176 |
| **Q12974** | PTP4A2 | -0.012929357 | 3.134949751 | -0.147680446 | 0.887089956 | 0.975708125 | -6.705814548 |
| **O15372** | EIF3H | -0.035795621 | 3.20296121 | -1.474686503 | 0.187212043 | 0.590446574 | -6.706114506 |
| **O94864-1** | SUPT7L | 0.016840645 | 3.177359828 | 0.740247312 | 0.485132005 | 0.81473183 | -6.706135033 |
| **Q9NQZ5** | STARD7 | 0.018407719 | 3.21391871 | 0.608394031 | 0.563659162 | 0.852578515 | -6.706589373 |
| **P31946** | YWHAB | 0.016234036 | 3.169179916 | 1.40440257 | 0.206331829 | 0.612058044 | -6.707286327 |
| **P62070-4** | RRAS2 | 0.013908128 | 3.152973191 | 0.646638377 | 0.540111516 | 0.843540088 | -6.707452115 |
| **Q5QP82** | DCAF10 | 0.009998412 | 3.177325558 | 0.35194861 | 0.736056639 | 0.929681266 | -6.70784191 |
| **Q9ULV3** | CIZ1 | 0.027736239 | 3.208344497 | 0.94445196 | 0.378852927 | 0.750267026 | -6.708575363 |
| **Q9NZ63** | C9orf78 | -0.062351662 | 3.228880367 | -1.380860508 | 0.213120883 | 0.61901473 | -6.70930004 |
| **Q96JC9** | EAF1 | 0.006433165 | 3.191502899 | 0.132059337 | 0.898948652 | 0.978757548 | -6.70973958 |

| | | | | | | | |
|---|---|---|---|---|---|---|---|
| **Q9BT23** | LIMD2 | -0.035491772 | 3.189852327 | -0.842469028 | 0.429557537 | 0.782580746 | -6.710041707 |
| **O95249-1** | GOSR1 | -0.037043727 | 3.191769105 | -0.767752755 | 0.469713917 | 0.806020965 | -6.710936346 |
| **Q9P0W2** | HMG20B | -0.018080177 | 3.198810554 | -0.664034671 | 0.529605566 | 0.839849063 | -6.711300532 |
| **Q7L4I2-1** | RSRC2 | 0.014109464 | 3.189006961 | 0.393798075 | 0.706380916 | 0.917775369 | -6.712494114 |
| **P34947** | GRK5 | 0.031342765 | 3.179830134 | 0.674748586 | 0.523200009 | 0.836627689 | -6.712602581 |
| **Q9BZF2** | OSBPL7 | 0.00785906 | 3.162508906 | 0.148975875 | 0.886107904 | 0.975708125 | -6.712732877 |
| **Q9P287** | BCCIP | -0.019505051 | 3.156219729 | -0.529741545 | 0.613968792 | 0.879837049 | -6.713190621 |
| **Q9UM21** | MGAT4A | 0.013437742 | 3.180292785 | 0.35583829 | 0.733276652 | 0.92857555 | -6.713426914 |
| **P15374** | UCHL3 | 0.026019241 | 3.183062573 | 1.245398908 | 0.256183867 | 0.659782078 | -6.713585756 |
| **Q9H0P0-4** | NT5C3; NT5C3A | 0.024384966 | 3.173508092 | 1.20435488 | 0.270645485 | 0.671731232 | -6.713781485 |
| **O75529-1** | TAF5L | -0.027094849 | 3.168890017 | -0.934936595 | 0.383383286 | 0.754258979 | -6.713858476 |
| **Q9NRG4** | SMYD2 | -0.017783122 | 3.217470458 | -1.124031118 | 0.30095906 | 0.691344519 | -6.71415659 |
| **Q1MSJ5-3** | CSPP1 | -0.024165069 | 3.198904518 | -0.970642994 | 0.366595288 | 0.741119482 | -6.714175722 |
| **Q8TDJ6-3** | DMXL2 | -0.017716892 | 3.181201128 | -1.10987083 | 0.306586225 | 0.695156266 | -6.714280962 |
| **O94900** | TOX | 0.058989078 | 3.206770933 | 0.685905262 | 0.516582622 | 0.833280927 | -6.71436428 |
| **Q7Z7K6-1** | CENPV | 0.016969751 | 3.216409094 | 0.876101095 | 0.412311518 | 0.770644294 | -6.714416614 |
| **P51398-1** | DAP3 | 0.132380161 | 3.221582435 | 1.043735202 | 0.334021121 | 0.719277002 | -6.714599428 |
| **Q96AT9** | RPE | -0.004594537 | 3.185551035 | -0.149892246 | 0.885413344 | 0.975708125 | -6.714830594 |
| **O60888-2** | CUTA | 0.026237804 | 3.178828779 | 0.684560055 | 0.517377636 | 0.833280927 | -6.715950682 |
| **O43818** | RRP9 | -0.029586627 | 3.194626437 | -0.724882351 | 0.493892037 | 0.821555157 | -6.716313206 |
| **Q8IWT3** | CUL9 | 0.000535332 | 3.171148175 | 0.013654586 | 0.989516888 | 0.996219143 | -6.716679243 |
| **P31629** | HIVEP2 | 0.011161487 | 3.157476035 | 0.293249772 | 0.778510131 | 0.944053983 | -6.71687972 |
| **Q9GZT9-1** | EGLN1 | -0.040219325 | 3.183412083 | -1.154740284 | 0.289049699 | 0.683685579 | -6.71703393 |
| **Q7Z392** | TRAPPC11 | -0.016510042 | 3.193549023 | -1.104573742 | 0.30871342 | 0.696978291 | -6.717454152 |
| **Q8WY54** | PPM1E | 0.015542528 | 3.196911207 | 0.770972035 | 0.46793158 | 0.80548741 | -6.717476744 |
| **P15586** | GNS | -0.021567451 | 3.172409874 | -0.590496856 | 0.574887993 | 0.859195843 | -6.717927454 |
| **O94855-2** | SEC24D | 0.021332232 | 3.176598875 | 1.246313683 | 0.255869278 | 0.659782078 | -6.71806126 |
| **P13674-1** | P4HA1 | -0.084025792 | 3.191362208 | -1.298377241 | 0.238508406 | 0.644491512 | -6.718362126 |
| **Q5T8I9** | HENMT1 | -0.002056122 | 3.180494535 | -0.058210379 | 0.955337334 | 0.989472196 | -6.718639114 |
| **O00116** | AGPS | -0.030523899 | 3.177639799 | -1.375271745 | 0.214761752 | 0.619913059 | -6.718726947 |
| **Q8WXX5** | DNAJC9 | -0.027524718 | 3.17752337 | -1.184214388 | 0.277993191 | 0.676093886 | -6.718856283 |
| **P46108-1** | CRK | -0.010013841 | 3.199674808 | -0.395539327 | 0.705157706 | 0.916737455 | -6.719110525 |
| **Q9P0I2** | EMC3 | -0.019665466 | 3.201318255 | -0.701263796 | 0.507561829 | 0.82913633 | -6.719947756 |
| **Q8N5Y2-1** | MSL3 | 0.013920115 | 3.173121019 | 0.459754222 | 0.660736739 | 0.896934512 | -6.720160286 |
| **Q8WUD1** | RAB2B | -0.004763714 | 3.195550452 | -0.074561155 | 0.942815856 | 0.988606569 | -6.720230517 |
| **Q86UT8** | CCDC84 | 0.008733383 | 3.184633361 | 0.232452498 | 0.823365371 | 0.959068023 | -6.720248109 |
| **Q8IYW5** | RNF168 | -0.042298106 | 3.182012885 | -0.924296867 | 0.388497734 | 0.75788842 | -6.720487847 |
| **P17812** | CTPS1 | -0.031983859 | 3.184419872 | -1.610633145 | 0.154780904 | 0.553156191 | -6.720620785 |
| **P17152** | TMEM11 | 0.02232572 | 3.211404935 | 0.289581326 | 0.781192739 | 0.944918194 | -6.720957281 |

| | | | | | | | |
|---|---|---|---|---|---|---|---|
| **Q8NI77** | KIF18A | 0.020566854 | 3.114189884 | 0.515785768 | 0.623149444 | 0.884409135 | -6.721647622 |
| **P09326** | CD48 | -0.033474684 | 3.143423563 | -0.578127577 | 0.582725113 | 0.864744128 | -6.721997426 |
| **P80303** | NUCB2 | -0.128122687 | 3.180936154 | -1.492065126 | 0.182741092 | 0.585371742 | -6.722377082 |
| **O14828** | SCAMP3 | 0.025528968 | 3.179084875 | 0.521367784 | 0.619468479 | 0.88249544 | -6.722524624 |
| **P08240-1** | SRPR; SRPRA | 0.019377258 | 3.176135625 | 1.311742383 | 0.234221418 | 0.64164013 | -6.722604489 |
| **O14579-1** | COPE | 0.017136792 | 3.179696237 | 1.198733283 | 0.272679565 | 0.672884265 | -6.722828206 |
| **Q9GZM5** | YIPF3 | -0.036718491 | 3.162556685 | -0.463378268 | 0.658271435 | 0.896852471 | -6.722847469 |
| **Q9BQA1** | WDR77 | -0.019740441 | 3.202081378 | -0.833187573 | 0.434407818 | 0.784298014 | -6.722856297 |
| **Q96BT7-4** | ALKBH8 | 0.021310305 | 3.183295371 | 0.933720671 | 0.383965166 | 0.754456977 | -6.723012803 |
| **Q99490-2** | AGAP2 | -0.01378307 | 3.16723656 | -0.658879054 | 0.53270561 | 0.839951983 | -6.723028008 |
| **Q15392** | DHCR24 | -0.004117772 | 3.152340692 | -0.087728027 | 0.932745232 | 0.986793807 | -6.723231787 |
| **O75354-1** | ENTPD6 | -0.007119514 | 3.175339133 | -0.258832666 | 0.803802423 | 0.95391847 | -6.723484995 |
| **O60232** | SSSCA1 | -0.022945277 | 3.167702864 | -0.602214855 | 0.567521122 | 0.854108736 | -6.723672085 |
| **P29466-1** | CASP1 | -0.015639349 | 3.177762635 | -0.521719037 | 0.619237247 | 0.882418852 | -6.724042217 |
| **Q9BU76-1** | C1orf35 | -0.020759487 | 3.188942649 | -0.729589101 | 0.491197412 | 0.819694064 | -6.724443685 |
| **Q92845** | KIFAP3 | 0.015256329 | 3.165818557 | 0.350756816 | 0.736909289 | 0.929681266 | -6.724768253 |
| **Q9UNP9-1** | PPIE | -0.051803488 | 3.143691558 | -0.94514458 | 0.37852477 | 0.750267026 | -6.724942701 |
| **P55769** | NHP2L1; SNU13 | 0.031909426 | 3.18696748 | 1.150927907 | 0.290506441 | 0.684803131 | -6.724957584 |
| **Q8TD55** | PLEKHO2 | -0.012306176 | 3.196251726 | -0.352055084 | 0.735980483 | 0.929681266 | -6.725148453 |
| **O14717-1** | TRDMT1 | -0.013537923 | 3.183520052 | -0.629417949 | 0.550638258 | 0.84701277 | -6.726029629 |
| **Q9BYI3-1** | FAM126A | -0.010154886 | 3.183612398 | -0.3582712 | 0.731540042 | 0.92815165 | -6.726123393 |
| **Q9NWB6** | ARGLU1 | -0.039786304 | 3.192697493 | -0.49098291 | 0.639646492 | 0.88954864 | -6.726399585 |
| **O00264** | PGRMC1 | 0.003970008 | 3.189356429 | 0.115065109 | 0.911881664 | 0.981900848 | -6.727166843 |
| **Q15814** | TBCC | 0.016163358 | 3.192653267 | 0.486267492 | 0.64280854 | 0.891449441 | -6.72731883 |
| **Q9H6V9-2** | C2orf43; LDAH | 0.018137572 | 3.185261012 | 0.956669966 | 0.373096106 | 0.745722908 | -6.727479417 |
| **Q5T7W0** | ZNF618 | -0.018227607 | 3.184772651 | -0.626223801 | 0.552604595 | 0.847927742 | -6.727902901 |
| **P12829** | MYL4 | -0.059330502 | 3.240610672 | -0.802936518 | 0.450488674 | 0.79513757 | -6.728012359 |
| **Q5VTE6-1** | ANGEL2 | 0.015699538 | 3.171270569 | 0.480279456 | 0.646835623 | 0.89271426 | -6.728117981 |
| **Q92551** | IP6K1 | -0.006522651 | 3.182162605 | -0.186465676 | 0.857789074 | 0.968855822 | -6.728638205 |
| **Q96Q42-1** | ALS2 | -0.007165875 | 3.185083278 | -0.232493998 | 0.823334484 | 0.959068023 | -6.728695299 |
| **Q8TDP1** | RNASEH2C | -0.01709292 | 3.212098172 | -0.256379465 | 0.805615466 | 0.954293355 | -6.728765498 |
| **P51532-4** | SMARCA4 | -0.023205863 | 3.175461496 | -1.476118932 | 0.186839762 | 0.590235352 | -6.729058005 |
| **Q9Y316-3** | MEMO1 | -0.020349065 | 3.198116597 | -0.860291199 | 0.420354268 | 0.77652027 | -6.729475084 |
| **Q9BQE3** | TUBA1C | 0.021532487 | 3.183617356 | 0.988555609 | 0.35839066 | 0.735575767 | -6.729523605 |
| **Q8IYT2** | CMTR2; FTSJD1 | 0.010222049 | 3.205050648 | 0.396625785 | 0.704394962 | 0.916268778 | -6.730497941 |
| **Q8NBM8** | PCYOX1L | -0.039854331 | 3.210374608 | -0.615815356 | 0.559041784 | 0.851394811 | -6.730525778 |
| **Q14554** | PDIA5 | 0.0285334 | 3.180273049 | 1.049846958 | 0.331405305 | 0.716856834 | -6.730653326 |
| **O95267-1** | RASGRP1 | 0.006620194 | 3.230631578 | 0.197009029 | 0.849863819 | 0.96500901 | -6.730954689 |
| **P51692** | STAT5B | 0.01678541 | 3.184513571 | 0.994236295 | 0.355818896 | 0.734600528 | -6.731023686 |

| | | | | | | | |
|---|---|---|---|---|---|---|---|
| **Q9C0B0** | UNK | 0.023548003 | 3.195985791 | 1.212542855 | 0.267705954 | 0.67010832 | -6.731612849 |
| **Q9Y5Y5-2** | PEX16 | -0.043067795 | 3.162696009 | -0.534598867 | 0.610790943 | 0.87752876 | -6.732048417 |
| **Q01130** | SRSF2 | 0.028276514 | 3.167470004 | 0.947225133 | 0.377540334 | 0.749625692 | -6.732114941 |
| **P09234** | SNRPC | -0.025480732 | 3.164170567 | -0.870065881 | 0.415368258 | 0.77203561 | -6.732302553 |
| **Q9HDC9** | APMAP | -0.035315245 | 3.197250228 | -1.040466905 | 0.33542673 | 0.720533987 | -6.732407883 |
| **Q8N4J0** | C9orf41; CARNMT1 | 0.00014959 | 3.199554149 | 0.00295658 | 0.997730051 | 0.998482186 | -6.732570147 |
| **Q9UEU0** | VTI1B | 0.015714113 | 3.208472639 | 0.358379912 | 0.731462484 | 0.92815165 | -6.732932901 |
| **O00559-2** | EBAG9 | -0.005496148 | 3.169638975 | -0.145817322 | 0.88850275 | 0.976290023 | -6.733430554 |
| **P42768** | WAS | -0.019938111 | 3.188342485 | -1.241505638 | 0.257526492 | 0.660441569 | -6.733522929 |
| **O43598-1** | C6orf108; DNPH1 | 0.001455106 | 3.165076521 | 0.041002421 | 0.968530032 | 0.992306703 | -6.733829343 |
| **P60033** | CD81 | 0.012681963 | 3.144052744 | 0.229330062 | 0.825690303 | 0.959077751 | -6.733858522 |
| **Q96PZ2** | FAM111A | -0.028231566 | 3.190201521 | -1.184609009 | 0.277847616 | 0.676093886 | -6.734397096 |
| **P19634-1** | SLC9A1 | 0.000292518 | 3.18756863 | 0.007259135 | 0.99442676 | 0.997608555 | -6.734862159 |
| **O43432-3** | EIF4G3 | -0.019136642 | 3.198376162 | -1.292289941 | 0.240483685 | 0.646142333 | -6.736035616 |
| **O00151** | PDLIM1 | -0.003568337 | 3.201417113 | -0.099429942 | 0.923806262 | 0.985083485 | -6.736442674 |
| **Q8NAV1-1** | PRPF38A | -0.035283281 | 3.194456135 | -0.852041187 | 0.424596505 | 0.778986255 | -6.736904519 |
| **P00734** | F2 | 0.036302285 | 3.222933731 | 0.839725673 | 0.430987063 | 0.782692325 | -6.738090323 |
| **P42356** | PI4KA | -0.019701741 | 3.183594476 | -1.536238427 | 0.171816273 | 0.571776747 | -6.739407049 |
| **Q15013-3** | MAD2L1BP | -0.001438525 | 3.16290945 | -0.026756568 | 0.979460123 | 0.993783815 | -6.739806084 |
| **Q9HCN4-5** | GPN1 | 0.009733377 | 3.19938386 | 0.367787013 | 0.724764198 | 0.926648409 | -6.739920723 |
| **O95218-2** | ZRANB2 | -0.027436823 | 3.178882066 | -1.135166789 | 0.296594183 | 0.688880164 | -6.740176644 |
| **Q8NBM4-1** | UBAC2 | -0.010880506 | 3.188993646 | -0.331611361 | 0.750661186 | 0.935635612 | -6.740478502 |
| **Q9Y587-2** | AP4S1 | -0.00287089 | 3.209110921 | -0.05439079 | 0.958264512 | 0.989911786 | -6.740538323 |
| **Q8WW12** | PCNP | -0.03856569 | 3.182296246 | -1.177974254 | 0.280303768 | 0.677109587 | -6.741216748 |
| **Q93008-3** | USP9X | -0.023105512 | 3.183425675 | -1.488022498 | 0.183772278 | 0.586138398 | -6.742167385 |
| **Q9H7Z3** | NRDE2 | 0.019582493 | 3.171936172 | 1.147370976 | 0.29187111 | 0.685264687 | -6.742464224 |
| **Q9UL18** | AGO1; EIF2C1 | 0.02335358 | 3.168165897 | 1.007601268 | 0.349825471 | 0.731407657 | -6.742665287 |
| **Q9NR45** | NANS | -0.021239657 | 3.183528955 | -1.226348833 | 0.262811343 | 0.666498017 | -6.742778087 |
| **P62253** | UBE2G1 | 0.032139669 | 3.181850705 | 1.059788615 | 0.327185545 | 0.71218881 | -6.743184489 |
| **P49841-2** | GSK3B | -0.020122752 | 3.187499875 | -0.983759371 | 0.360573326 | 0.736630258 | -6.743205748 |
| **Q9Y295** | DRG1 | -0.0342408 | 3.196259122 | -1.097016239 | 0.311769346 | 0.699730515 | -6.743429223 |
| **Q9HCD6-2** | TANC2 | -0.006605285 | 3.192290221 | -0.19766543 | 0.849371027 | 0.964964904 | -6.744041393 |
| **P25445-1** | FAS | -0.00378788 | 3.162348384 | -0.115056661 | 0.911888101 | 0.981900848 | -6.745062616 |
| **Q58EX7** | PLEKHG4 | -0.020152852 | 3.194866015 | -0.520218301 | 0.62022552 | 0.882940648 | -6.745234555 |
| **Q5TA45-1** | CPSF3L; INTS11 | 0.02852845 | 3.195448883 | 1.10399172 | 0.308947887 | 0.696978291 | -6.745363108 |
| **Q8WXE1** | ATRIP | 0.015262802 | 3.18640361 | 0.355073874 | 0.733822644 | 0.928822831 | -6.746357887 |
| **O95789** | ZMYM6 | -0.011770508 | 3.190603475 | -0.504116264 | 0.630882631 | 0.887123867 | -6.746583526 |
| **Q12905** | ILF2 | -0.019275403 | 3.171604265 | -1.514392432 | 0.177141321 | 0.578452275 | -6.746606684 |
| **Q9BTF0** | THUMPD2 | -0.00892689 | 3.203222855 | -0.204411255 | 0.844310912 | 0.962543524 | -6.747236014 |

| | | | | | | | |
|---|---|---|---|---|---|---|---|
| **Q7Z5J4** | RAI1 | -0.015972599 | 3.198361983 | -0.642371047 | 0.542708389 | 0.843669625 | -6.74816274 |
| **P56270-2** | MAZ | -0.015058249 | 3.196563981 | -0.43841779 | 0.675343471 | 0.90314827 | -6.748492936 |
| **Q68D91-1** | MBLAC2 | 0.014334987 | 3.175614747 | 0.738964556 | 0.485859319 | 0.81473183 | -6.748552765 |
| **Q6Y288** | B3GALTL; B3GLCT | 0.018090846 | 3.208399045 | 0.878667733 | 0.411016599 | 0.769906516 | -6.748554742 |
| **Q96DN5-1** | TBC1D31; WDR67 | -0.015708404 | 3.178614526 | -0.584309493 | 0.578800514 | 0.861567376 | -6.748689273 |
| **Q9BVI0** | PHF20 | -0.020328319 | 3.182402187 | -1.067175109 | 0.324078514 | 0.710703247 | -6.748794785 |
| **Q9NV66** | TYW1 | -0.018477952 | 3.198246657 | -0.574392067 | 0.585104091 | 0.8653224 | -6.748841589 |
| **Q86YT6** | MIB1 | -0.016117139 | 3.162811433 | -0.857329951 | 0.421873396 | 0.777315429 | -6.749076596 |
| **Q9UL25** | RAB21 | -0.018433158 | 3.198131505 | -1.233097708 | 0.260446745 | 0.663969372 | -6.749354845 |
| **Q9HB71** | CACYBP | 0.079775053 | 3.181388454 | 1.486855671 | 0.184070904 | 0.586619868 | -6.749416228 |
| **Q9H1D9** | POLR3F | -0.025829931 | 3.19199942 | -0.708256288 | 0.5034891 | 0.827688992 | -6.749703598 |
| **Q92698** | RAD54L | -0.017499701 | 3.151368995 | -0.726381212 | 0.493032867 | 0.821156631 | -6.749817614 |
| **Q96BW1** | UPRT | -0.021561392 | 3.183194934 | -0.480470947 | 0.64670664 | 0.89271426 | -6.750021186 |
| **Q6P3X3** | TTC27 | -0.026586894 | 3.180057658 | -1.240532905 | 0.257862893 | 0.660717666 | -6.750340776 |
| **Q96RT1-8** | ERBB2IP; ERBIN | -0.037292612 | 3.188986067 | -1.543160411 | 0.170160264 | 0.569105131 | -6.750514948 |
| **Q9P2N5** | RBM27 | 0.029215101 | 3.178086531 | 1.203063689 | 0.271111533 | 0.671838876 | -6.750584777 |
| **Q9NUQ8** | ABCF3 | -0.034669385 | 3.173798453 | -1.216353331 | 0.266347298 | 0.669449314 | -6.75147433 |
| **Q9H098-2** | FAM107B | -0.047087954 | 3.15709468 | -0.650572148 | 0.537724503 | 0.84335148 | -6.751584629 |
| **Q13370** | PDE3B | -0.013092035 | 3.188035488 | -0.403313398 | 0.699708133 | 0.913245694 | -6.751605671 |
| **Q14653** | IRF3 | 0.006061488 | 3.210972812 | 0.167930933 | 0.871764149 | 0.971931713 | -6.752360406 |
| **Q99848** | EBNA1BP2 | -0.055836453 | 3.18520544 | -1.219542581 | 0.265214685 | 0.66823064 | -6.752668393 |
| **Q9UEY8-1** | ADD3 | -0.035776932 | 3.210901243 | -1.096842227 | 0.311840001 | 0.699730515 | -6.752756987 |
| **Q7L775** | EPM2AIP1 | 0.055577137 | 3.208677396 | 0.690532088 | 0.513854205 | 0.83148099 | -6.75313891 |
| **Q9Y6X3-1** | MAU2 | -0.008835401 | 3.196492946 | -0.24946722 | 0.810730951 | 0.955498404 | -6.753634236 |
| **P09914** | IFIT1 | 0.028922692 | 3.196914024 | 0.992285807 | 0.356700286 | 0.734600528 | -6.754446656 |
| **P49593** | PPM1F | -0.01953275 | 3.17160048 | -0.472935535 | 0.651792212 | 0.894547168 | -6.754804201 |
| **Q86VQ1** | GLCCI1 | 0.002058968 | 3.181812445 | 0.049328434 | 0.962145169 | 0.991283985 | -6.754856982 |
| **Q6ZW49** | PAXIP1 | -0.01764248 | 3.18791419 | -1.11696964 | 0.303754457 | 0.692399426 | -6.754905493 |
| **Q9H115** | NAPB | 0.000881132 | 3.205301341 | 0.033602982 | 0.974206473 | 0.99316605 | -6.755647425 |
| **Q9NP87** | POLM | 0.010572147 | 3.168985816 | 0.326002767 | 0.75470881 | 0.937046647 | -6.755757659 |
| **P57076** | C21orf59 | -0.026231314 | 3.199615211 | -1.158187743 | 0.28773767 | 0.682522792 | -6.757226839 |
| **P61599** | NAA20 | 0.015024989 | 3.186757153 | 0.323948788 | 0.756193242 | 0.937424764 | -6.757305077 |
| **Q9NSK0-3** | KLC4 | 0.017115895 | 3.17604118 | 0.661102942 | 0.531366996 | 0.839849063 | -6.757491365 |
| **Q8TAD4-1** | SLC30A5 | -0.028500392 | 3.197711891 | -0.660471688 | 0.531746747 | 0.839849063 | -6.757583861 |
| **Q9C0F1-1** | CEP44 | 0.017182019 | 3.181209167 | 0.613392835 | 0.560546503 | 0.852515691 | -6.757600632 |
| **O15287** | FANCG | -0.001323938 | 3.186350895 | -0.037198458 | 0.971447997 | 0.992928787 | -6.757606137 |
| **Q9Y6R9** | CCDC61 | 0.016902621 | 3.205476267 | 0.469297549 | 0.654254718 | 0.896370708 | -6.758028165 |
| **Q04323-2** | UBXN1 | 0.036431238 | 3.18835392 | 0.780151716 | 0.462874966 | 0.802800142 | -6.758404172 |
| **Q86UY6** | NAA40 | 0.003725346 | 3.19012816 | 0.085859155 | 0.93417386 | 0.986793807 | -6.759044809 |

| | | | | | | | |
|---|---|---|---|---|---|---|---|
| **Q68DK2** | ZFYVE26 | -0.010849983 | 3.159670553 | -0.3285551 | 0.752865785 | 0.935782019 | -6.75926288 |
| **P62308** | SNRPG | 0.012428074 | 3.171140205 | 0.415611098 | 0.691126866 | 0.909522034 | -6.759380892 |
| **Q9Y5W9** | SNX11 | -0.009360473 | 3.200656252 | -0.213060434 | 0.837834805 | 0.96111408 | -6.759784314 |
| **Q6P5R6** | RPL22L1 | -0.035922601 | 3.210345482 | -0.705864072 | 0.504880008 | 0.828775136 | -6.759911143 |
| **Q96FK6** | WDR89 | -0.015656447 | 3.186242293 | -0.848594043 | 0.426378263 | 0.779649091 | -6.760015078 |
| **P42575-1** | CASP2 | -0.040265753 | 3.206785719 | -1.313817353 | 0.233561968 | 0.640812038 | -6.76046397 |
| **P63218** | GNG5 | -0.041620374 | 3.160807935 | -0.526009051 | 0.616416895 | 0.880991586 | -6.761189907 |
| **Q9H7B4-1** | SMYD3 | -0.025045232 | 3.168126846 | -1.05434653 | 0.329490051 | 0.715245479 | -6.762797377 |
| **P21266** | GSTM3 | 0.01690022 | 3.167002519 | 1.210772002 | 0.26833938 | 0.67097152 | -6.762797436 |
| **Q9P0K1** | ADAM22 | 0.004980796 | 3.189232857 | 0.180224863 | 0.862488686 | 0.969309769 | -6.76317777 |
| **O60879** | DIAPH2 | 0.023742847 | 3.167322332 | 1.182531169 | 0.278614855 | 0.676427621 | -6.76424587 |
| **Q96HR8** | NAF1 | 0.024643472 | 3.212130938 | 0.51511437 | 0.623592981 | 0.884722542 | -6.764420943 |
| **Q9Y2L9-2** | LRCH1 | -0.018847675 | 3.170891911 | -0.885500491 | 0.407584004 | 0.767722833 | -6.764504289 |
| **Q9NRS6** | SNX15 | 0.008857639 | 3.199518247 | 0.26132822 | 0.801959428 | 0.953157309 | -6.765679099 |
| **P78364** | PHC1 | -1.79E-05 | 3.17977518 | -0.000503175 | 0.999613681 | 0.999613681 | -6.765832911 |
| **Q9H9J4-1** | USP42 | 0.012422567 | 3.184008799 | 0.444280271 | 0.671314496 | 0.901175787 | -6.767059096 |
| **Q96T88-2** | UHRF1 | -0.021761981 | 3.178870257 | -1.701102373 | 0.136210244 | 0.526101308 | -6.767317583 |
| **Q02040-1** | AKAP17A | -0.018826698 | 3.201251219 | -0.812539171 | 0.445339 | 0.79137069 | -6.767331204 |
| **Q7L1W4** | LRRC8D | -0.009160756 | 3.204937209 | -0.248396725 | 0.811524092 | 0.955606306 | -6.767718951 |
| **Q96GX9-1** | APIP | -0.025580598 | 3.199989628 | -0.432477131 | 0.679438035 | 0.903893384 | -6.767931964 |
| **Q7Z6J9** | TSEN54 | -0.021361292 | 3.182443946 | -0.610980457 | 0.562047349 | 0.852576894 | -6.768595138 |
| **Q9UIQ6-1** | LNPEP | -0.040844968 | 3.171214746 | -1.209004691 | 0.268972814 | 0.671425933 | -6.768823019 |
| **Q8ND56** | LSM14A | -0.017473691 | 3.191789104 | -0.660344955 | 0.531823008 | 0.839849063 | -6.768926092 |
| **O43920** | NDUFS5 | 0.019998954 | 3.196661968 | 0.578921952 | 0.582219937 | 0.864462231 | -6.76926728 |
| **P24386-1** | CHM | -0.006345128 | 3.164739345 | -0.219812293 | 0.832788796 | 0.960584638 | -6.769345555 |
| **Q86U28** | ISCA2 | -0.008859478 | 3.208209724 | -0.147572829 | 0.887171549 | 0.975708125 | -6.769512001 |
| **P16104** | H2AFX | -0.004088928 | 3.178670336 | -0.118536686 | 0.909237203 | 0.981506736 | -6.7696548 |
| **Q9NX38** | FAM206A | 0.006109402 | 3.167689598 | 0.230674754 | 0.824688824 | 0.959077751 | -6.771421826 |
| **Q96FV2** | SCRN2 | -0.008774892 | 3.161345172 | -0.305687884 | 0.76943924 | 0.941649022 | -6.772070439 |
| **P28290** | SSFA2 | -0.013762142 | 3.188428517 | -0.353780766 | 0.734746642 | 0.92925216 | -6.772300586 |
| **Q9Y4G8** | RAPGEF2 | -0.005058976 | 3.175550014 | -0.104504203 | 0.919933731 | 0.983742622 | -6.77243211 |
| **O14641** | DVL2 | -0.006310708 | 3.173954252 | -0.246853174 | 0.81266815 | 0.956167049 | -6.772661077 |
| **P06493** | CDK1 | -0.036288811 | 3.16534002 | -1.419457317 | 0.2020932 | 0.608654462 | -6.773800988 |
| **Q9Y6J0** | CABIN1 | -0.01874981 | 3.179366901 | -1.24398858 | 0.256669531 | 0.659782078 | -6.774498397 |
| **Q9UPZ3** | HPS5 | -0.003775473 | 3.208158834 | -0.137885532 | 0.894522251 | 0.97851842 | -6.775293958 |
| **O60861-3** | GAS7 | -0.059428603 | 3.171838702 | -0.874655734 | 0.413042056 | 0.771057127 | -6.775484195 |
| **P56937-1** | HSD17B7 | 0.026020419 | 3.156460957 | 0.770286909 | 0.468310505 | 0.805700945 | -6.775769058 |
| **Q96N16** | JAKMIP1 | -0.004292512 | 3.197914657 | -0.163691534 | 0.87496786 | 0.972759481 | -6.775835425 |
| **O96011-1** | LOC101060567; PE | -0.004581184 | 3.181197637 | -0.124474863 | 0.904716743 | 0.980490319 | -6.776778792 |

| | | | | | | | |
|---|---|---|---|---|---|---|---|
| **Q9Y547** | HSPB11 | 0.00739963 | 3.16544876 | 0.225059542 | 0.828873169 | 0.959254681 | -6.77696439 |
| **O94919** | ENDOD1 | -0.011811723 | 3.160079932 | -0.363915362 | 0.727517854 | 0.92710315 | -6.777465378 |
| **Q2PPJ7-1** | RALGAPA2 | 0.016734737 | 3.208337137 | 0.539005098 | 0.607916095 | 0.875726951 | -6.777784676 |
| **Q8IWZ6-1** | BBS7 | -0.00609588 | 3.172113557 | -0.201052735 | 0.84682919 | 0.963629033 | -6.778159247 |
| **Q92520** | FAM3C; WNT16 | -0.027490602 | 3.174118011 | -0.990348388 | 0.357577464 | 0.73485591 | -6.778626389 |
| **Q9BXS6** | NUSAP1 | 0.021703281 | 3.123290788 | 0.640688266 | 0.543734569 | 0.844239037 | -6.778641122 |
| **P21675-12** | TAF1 | 0.025337657 | 3.190796226 | 1.300395016 | 0.237856801 | 0.64431377 | -6.778798014 |
| **Q00403** | GTF2B | -0.032491838 | 3.191133428 | -1.036980877 | 0.336931198 | 0.721128781 | -6.779719622 |
| **P61221** | ABCE1 | 0.036617252 | 3.171295719 | 1.427414612 | 0.19988482 | 0.605177234 | -6.779843112 |
| **Q9P2K3-3** | RCOR3 | -0.011969688 | 3.188254963 | -0.483407207 | 0.644730523 | 0.891546795 | -6.779923123 |
| **Q13685** | AAMP | -0.032351176 | 3.187151243 | -1.161720772 | 0.286398263 | 0.681992638 | -6.780221891 |
| **Q9H3Z4** | DNAJC5 | 0.051959317 | 3.1890605 | 0.634223712 | 0.547687898 | 0.846115385 | -6.780420412 |
| **P49759-3** | CLK1 | 0.043840341 | 3.200614394 | 0.71098735 | 0.501904261 | 0.826806835 | -6.781040355 |
| **P51571** | SSR4 | -0.011706962 | 3.200975595 | -0.421548009 | 0.687001657 | 0.907305098 | -6.781262633 |
| **Q3L8U1-1** | CHD9 | -0.025378078 | 3.218229243 | -0.693759611 | 0.51195649 | 0.831118576 | -6.781660064 |
| **Q5T6F0** | DCAF12 | -0.01595121 | 3.187070544 | -0.649949185 | 0.538102077 | 0.843524953 | -6.78176157 |
| **P47813** | EIF1AX; LOC10106 | 0.013452525 | 3.190920804 | 0.120555155 | 0.907700215 | 0.98075504 | -6.781887612 |
| **O60493-1** | SNX3 | -0.037282102 | 3.206657597 | -0.824375773 | 0.439049005 | 0.787769725 | -6.78192881 |
| **Q96CV9-1** | OPTN | -0.020133159 | 3.167363654 | -0.610041529 | 0.562632151 | 0.852576894 | -6.781945632 |
| **Q96LT9-2** | RNPC3 | 0.016152288 | 3.181780177 | 0.9032508 | 0.398766344 | 0.762880385 | -6.782507166 |
| **Q9Y2G8** | DNAJC16 | 0.008105453 | 3.207052732 | 0.21159645 | 0.838930022 | 0.961364456 | -6.782952584 |
| **Q9NZI7** | UBP1 | 0.018594496 | 3.206347408 | 0.732464536 | 0.489556065 | 0.81832087 | -6.783152786 |
| **Q6N069** | NAA16 | -0.023466721 | 3.166501251 | -1.224048299 | 0.263621581 | 0.666941931 | -6.783312008 |
| **P50613** | CDK7 | 0.00340332 | 3.18298486 | 0.105941043 | 0.9188376 | 0.983742622 | -6.783327711 |
| **Q96MU7-1** | YTHDC1 | 0.019848783 | 3.180374874 | 0.931405393 | 0.385075 | 0.754667212 | -6.78345402 |
| **P49458** | SRP9; SRP9P1 | 0.028409978 | 3.185988724 | 1.169908744 | 0.283314253 | 0.679787821 | -6.783692984 |
| **Q86T82** | USP37 | -0.011777002 | 3.174644126 | -0.318454921 | 0.760169217 | 0.938422437 | -6.784093011 |
| **Q10472** | GALNT1 | 0.011628117 | 3.171568756 | 0.392524655 | 0.707276079 | 0.918037648 | -6.785315628 |
| **Q9UKZ1** | C2orf29; CNOT11 | -0.028241935 | 3.185564305 | -0.691916943 | 0.513039382 | 0.831256087 | -6.785689744 |
| **Q16864-2** | ATP6V1F | -0.016932431 | 3.19746253 | -0.471275944 | 0.652914983 | 0.895616288 | -6.786449752 |
| **P41212** | ETV6 | 0.007454797 | 3.132305386 | 0.177684359 | 0.864403551 | 0.969738239 | -6.786753253 |
| **Q8TAV0-3** | FAM76A | -0.016321068 | 3.186746086 | -0.459156179 | 0.661144008 | 0.896934512 | -6.787412979 |
| **Q12873-3** | CHD3 | 0.000804666 | 3.193942325 | 0.03250128 | 0.975051786 | 0.99316605 | -6.787530415 |
| **P35232** | PHB | 0.070261829 | 3.16399586 | 1.2334137 | 0.260336481 | 0.663969372 | -6.787833583 |
| **Q9Y5P6-2** | GMPPB | -0.024588691 | 3.195716833 | -0.901554134 | 0.399602965 | 0.763210165 | -6.788093342 |
| **Q15596** | NCOA2 | 0.027911133 | 3.190426139 | 0.927514576 | 0.386945567 | 0.756467159 | -6.788310425 |
| **Q96G23** | CERS2 | -0.002649697 | 3.212538923 | -0.066505304 | 0.948983092 | 0.989472196 | -6.788443716 |
| **Q96RR4-1** | CAMKK2 | 0.0165208 | 3.184883169 | 0.540714315 | 0.606802942 | 0.875166995 | -6.788646343 |
| **Q86X83** | COMMD2 | -0.005945803 | 3.182387545 | -0.111193188 | 0.91483252 | 0.98273991 | -6.788885796 |

| | | | | | | | |
|---|---|---|---|---|---|---|---|
| **Q8N128-2** | FAM177A1 | -0.01053607 | 3.204424307 | -0.359940614 | 0.730349405 | 0.927974736 | -6.788985618 |
| **Q99598** | TSNAX | 0.021701589 | 3.155375723 | 1.010799221 | 0.348403245 | 0.730167181 | -6.789059633 |
| **O00483** | NDUFA4 | 0.078757499 | 3.176484553 | 1.009754602 | 0.348867315 | 0.730176717 | -6.789519682 |
| **P61081** | UBE2M | -0.048255469 | 3.186980347 | -1.111140671 | 0.306078083 | 0.694598791 | -6.789883945 |
| **O00512** | BCL9; LOC101060 | -0.018840817 | 3.191042842 | -0.48220172 | 0.645541444 | 0.891969874 | -6.789890725 |
| **Q13464** | ROCK1 | -0.021332526 | 3.165708315 | -1.454943756 | 0.192412767 | 0.596793208 | -6.789941294 |
| **Q92574-1** | TSC1 | 0.018146426 | 3.16699282 | 0.668629661 | 0.526852273 | 0.839483378 | -6.790171502 |
| **Q9Y3E2** | BOLA1 | 0.003152897 | 3.173137274 | 0.121693404 | 0.906833671 | 0.98075504 | -6.790544633 |
| **Q14694-2** | USP10 | -0.03420691 | 3.194696321 | -1.30909642 | 0.235064714 | 0.64164013 | -6.790649954 |
| **P15056** | BRAF | -0.019181088 | 3.171883279 | -0.825483524 | 0.438463607 | 0.786897076 | -6.791685988 |
| **Q05048** | CSTF1 | -0.025227025 | 3.174847231 | -1.237809784 | 0.258806647 | 0.661376267 | -6.791852641 |
| **Q8IZD4** | DCP1B | 0.01807065 | 3.156158994 | 0.702305381 | 0.506953802 | 0.82913633 | -6.792063623 |
| **Q92616** | GCN1L1; GCN1 | -0.027302927 | 3.183943639 | -1.889842118 | 0.104122716 | 0.471639097 | -6.793619453 |
| **Q8IZA0** | KIAA0319L | 0.002288691 | 3.201459179 | 0.062563856 | 0.952001925 | 0.989472196 | -6.793936869 |
| **Q8IZ07** | ANKRD13A | -0.024067026 | 3.172167752 | -0.736084268 | 0.487495097 | 0.816430975 | -6.793995475 |
| **Q9UBB4-1** | ATXN10 | -0.029420378 | 3.183277553 | -1.478564681 | 0.186205694 | 0.588831734 | -6.794204769 |
| **P51452** | DUSP3 | -0.016462718 | 3.202535484 | -0.509872032 | 0.627061996 | 0.885387512 | -6.794224295 |
| **Q15435** | PPP1R7 | 0.003331255 | 3.171500431 | 0.069837258 | 0.946431794 | 0.989472196 | -6.795676827 |
| **P27797** | CALR | 0.026485575 | 3.206302419 | 1.932911431 | 0.097909934 | 0.456510811 | -6.795717413 |
| **O43852-3** | CALU | -0.010203673 | 3.2022531 | -0.28494185 | 0.784590077 | 0.946344035 | -6.796115004 |
| **Q16851** | UGP2 | -0.022996385 | 3.169619459 | -1.575195668 | 0.162688104 | 0.561580888 | -6.796567592 |
| **Q86TM6** | SYVN1 | 0.002138298 | 3.218829938 | 0.053096783 | 0.959256348 | 0.990513797 | -6.797234765 |
| **Q13490-1** | BIRC2 | -0.015635484 | 3.177084421 | -0.614637088 | 0.559773345 | 0.851665881 | -6.797265058 |
| **Q96EZ8-2** | MCRS1 | 0.021835527 | 3.178162742 | 0.884310031 | 0.408180526 | 0.768299994 | -6.797552722 |
| **Q96B45** | C10orf32; BORCS7 | 0.002621106 | 3.183023864 | 0.07218603 | 0.944633733 | 0.98904859 | -6.798806183 |
| **Q8TF05** | PPP4R1 | 0.020139266 | 3.186044629 | 0.993778349 | 0.35602568 | 0.734600528 | -6.798888411 |
| **Q9H2H9** | SLC38A1 | 0.03904337 | 3.178794359 | 0.437033671 | 0.676296404 | 0.903662615 | -6.798959291 |
| **Q5H9U9-1** | DDX60L | 0.008681057 | 3.194421047 | 0.218563335 | 0.833721567 | 0.96068017 | -6.7993379 |
| **P57740** | NUP107 | -0.03325294 | 3.191672578 | -1.462931264 | 0.190292946 | 0.594291452 | -6.799623637 |
| **Q9Y5R8** | TRAPPC1 | -0.000920398 | 3.181542991 | -0.02776273 | 0.97868796 | 0.99361817 | -6.799934592 |
| **Q7L3B6** | CDC37L1 | -0.028718126 | 3.147035741 | -0.347366616 | 0.73933696 | 0.931081875 | -6.800130575 |
| **P35626** | ADRBK2; GRK3 | -0.017234471 | 3.22478491 | -0.846152389 | 0.427643585 | 0.78052568 | -6.800415439 |
| **P00519-2** | ABL1 | 0.023338856 | 3.205340494 | 1.079732089 | 0.318851518 | 0.707150066 | -6.800778003 |
| **P08754** | GNAI3 | -0.053233337 | 3.131217699 | -0.682425791 | 0.5186406 | 0.833791899 | -6.802524984 |
| **Q96C19** | EFHD2 | -0.051228141 | 3.190117716 | -0.715918877 | 0.499050831 | 0.824970816 | -6.802843326 |
| **Q6UWE0** | LRSAM1 | -0.016813393 | 3.177312424 | -0.710931085 | 0.501936878 | 0.826806835 | -6.803130125 |
| **Q53S33** | BOLA3 | 0.028723741 | 3.173593673 | 0.295381004 | 0.776953141 | 0.94357883 | -6.803162374 |
| **Q96N96-6** | SPATA13 | -0.011674101 | 3.19701241 | -0.363196851 | 0.728029368 | 0.92710315 | -6.803285325 |
| **Q9H1P3-1** | OSBPL2 | 0.002996636 | 3.181132533 | 0.116422143 | 0.910847802 | 0.981900848 | -6.803303127 |

| | | | | | | | |
|---|---|---|---|---|---|---|---|
| **Q9HCD5** | NCOA5 | -0.02537467 | 3.184453558 | -1.310311436 | 0.234677144 | 0.64164013 | -6.803401531 |
| **Q9BTE1** | DCTN5 | -0.001604826 | 3.144941444 | -0.032904335 | 0.974742526 | 0.99316605 | -6.803554726 |
| **Q8N8S7** | ENAH | 0.021251382 | 3.182800498 | 0.802714584 | 0.450608188 | 0.79513757 | -6.804118067 |
| **Q92973-1** | TNPO1 | -0.02645361 | 3.179842803 | -1.210439988 | 0.268458282 | 0.67097152 | -6.805118964 |
| **P56159-2** | GFRA1 | 0.050350346 | 3.175772729 | 0.78072442 | 0.462560753 | 0.802800142 | -6.805209997 |
| **Q86WP2-2** | GPBP1 | -0.005227347 | 3.220310665 | -0.191546004 | 0.853967984 | 0.967456956 | -6.805327162 |
| **Q9HCH0** | NCKAP5L | -0.008407741 | 3.181097578 | -0.336321816 | 0.747268325 | 0.934974996 | -6.805580157 |
| **O15197-1** | EPHB6 | 0.009624789 | 3.187868869 | 0.320388646 | 0.75876885 | 0.9384213 | -6.806748421 |
| **Q9Y294** | ASF1A | -0.056088534 | 3.149914486 | -0.933108254 | 0.384258492 | 0.754667212 | -6.807306352 |
| **Q8NG68** | TTL | 0.012836919 | 3.179419554 | 0.464179959 | 0.657726697 | 0.896852471 | -6.807778258 |
| **Q9BQC3** | DPH2 | -0.017507655 | 3.191069749 | -0.683852592 | 0.517796062 | 0.83340353 | -6.808347507 |
| **O76054** | SEC14L2 | 0.007293901 | 3.214829969 | 0.147632723 | 0.887126138 | 0.975708125 | -6.808517214 |
| **Q9Y2U5** | MAP3K2 | -0.00862217 | 3.199512362 | -0.249279475 | 0.810870035 | 0.955498404 | -6.808893402 |
| **Q13404-1** | TMEM189-UBE2V1 | -0.025203467 | 3.18596138 | -1.117707061 | 0.303461537 | 0.692399426 | -6.809630501 |
| **Q9NQC7** | CYLD | 0.027171208 | 3.171276678 | 0.845318336 | 0.428076435 | 0.780598411 | -6.809817925 |
| **P49023** | PXN | -0.015336999 | 3.218728705 | -0.702246156 | 0.506988361 | 0.82913633 | -6.81001841 |
| **O75155-1** | CAND2 | 0.013157405 | 3.184378548 | 0.532335479 | 0.612270609 | 0.878766193 | -6.810247666 |
| **Q9UM13** | ANAPC10 | 0.004706154 | 3.188963554 | 0.082049776 | 0.937086689 | 0.987232823 | -6.810423645 |
| **Q92989-1** | CLP 1 | -0.020161966 | 3.182744189 | -0.917955503 | 0.391570503 | 0.759806823 | -6.810492671 |
| **Q8N4S0** | CCDC82 | 0.006985667 | 3.147373173 | 0.18759031 | 0.856942836 | 0.968855822 | -6.8107045 |
| **Q9UIC8** | LCMT1 | -0.032811802 | 3.165542981 | -0.851769473 | 0.424736751 | 0.778986255 | -6.811732371 |
| **Q5VW38** | GPR107 | 0.003691717 | 3.172803034 | 0.077730388 | 0.940390764 | 0.988019653 | -6.812179964 |
| **Q9UIM3** | FKBPL | 0.014844715 | 3.157380254 | 0.22491687 | 0.828979565 | 0.959254681 | -6.81253815 |
| **Q8IU81** | IRF2BP1 | -0.021859025 | 3.200786179 | -1.168514485 | 0.283837419 | 0.679787821 | -6.812569968 |
| **Q9Y3Q3** | TMED3 | -0.005007431 | 3.227031644 | -0.042578898 | 0.967320891 | 0.992306703 | -6.812570649 |
| **P14649** | MYL6B | -0.02374101 | 3.188852342 | -0.960044514 | 0.371518033 | 0.744717822 | -6.81297158 |
| **O95749** | GGPS1 | -0.018885645 | 3.189897261 | -0.859900281 | 0.420554581 | 0.77668669 | -6.813118016 |
| **Q14289-2** | PTK2B | -0.031713672 | 3.219347224 | -1.178966516 | 0.279935274 | 0.677042847 | -6.814035883 |
| **Q5VZE5** | NAA35 | -0.028370703 | 3.173041982 | -1.208948655 | 0.26899292 | 0.671425933 | -6.814070364 |
| **O00560-1** | SDCBP | -0.02462885 | 3.157299809 | -0.913961535 | 0.393515224 | 0.760194378 | -6.814304308 |
| **Q92879-4** | CELF1 | -0.022856127 | 3.194872562 | -1.065385291 | 0.324829176 | 0.710703247 | -6.814629731 |
| **Q15788** | NCOA1 | -0.014245814 | 3.191941314 | -0.616165737 | 0.558824351 | 0.851363273 | -6.815606952 |
| **Q9NZN4** | EHD2 | -0.019769206 | 3.201923728 | -0.63336834 | 0.548212316 | 0.846224652 | -6.815653742 |
| **Q14247-1** | CTTN | -0.010090085 | 3.19775385 | -0.330464723 | 0.751488005 | 0.935635612 | -6.816811207 |
| **Q9P2R6** | RERE | 0.003071517 | 3.174955256 | 0.116362222 | 0.910893449 | 0.981900848 | -6.816851676 |
| **Q92734-1** | TFG | 0.027492204 | 3.195375001 | 0.912610806 | 0.394174558 | 0.760194378 | -6.817003126 |
| **P16284** | PECAM1 | -0.021829685 | 3.17952921 | -1.21374568 | 0.267276438 | 0.67010832 | -6.817259545 |
| **Q96ES7** | CCDC101; SGF29 | 0.01870852 | 3.249387878 | 0.124976203 | 0.90433527 | 0.980210603 | -6.81765925 |
| **Q92995** | USP13 | -0.029575609 | 3.211226878 | -0.811080539 | 0.446118531 | 0.792020296 | -6.817968318 |

| | | | | | | | |
|---|---|---|---|---|---|---|---|
| **Q8WWI5** | SLC44A1 | -0.025693077 | 3.184189884 | -0.576408942 | 0.583818936 | 0.8653224 | -6.818238352 |
| **O75874** | IDH1 | 0.020041118 | 3.145252701 | 1.337835537 | 0.226047149 | 0.63121696 | -6.818295537 |
| **Q7LDG7-2** | RASGRP2 | -0.012306958 | 3.18740619 | -0.296538125 | 0.776108264 | 0.94357883 | -6.818412315 |
| **Q8NA72** | POC5 | -0.014240311 | 3.212279231 | -0.476111329 | 0.649646418 | 0.893646754 | -6.819264563 |
| **Q9BYB4-1** | GNB1L | 0.070087865 | 3.227521663 | 0.836952429 | 0.432435653 | 0.783333621 | -6.819900776 |
| **Q8IWD4-1** | CCDC117 | 0.01767216 | 3.186426253 | 0.287691545 | 0.782575941 | 0.94563233 | -6.820196131 |
| **Q8IV63** | VRK3 | 0.015447823 | 3.153308992 | 0.613908119 | 0.560226236 | 0.852191738 | -6.820301383 |
| **Q9Y2S2-1** | CRYL1 | -0.016676349 | 3.203170793 | -0.517392572 | 0.622088655 | 0.883385606 | -6.820332849 |
| **Q96DM3** | C18orf8; WDR98 | 0.012639149 | 3.183356771 | 0.610733903 | 0.562200877 | 0.852576894 | -6.820595219 |
| **Q9Y3T6** | R3HCC1 | -0.013836332 | 3.191130468 | -0.442963791 | 0.672218225 | 0.90142081 | -6.821358625 |
| **Q8TC12** | RDH11 | -0.020483027 | 3.181935647 | -1.036790838 | 0.337013368 | 0.721128781 | -6.821376734 |
| **Q9GZM8-2** | NDEL1 | 0.016957494 | 3.203827573 | 0.28236043 | 0.786482591 | 0.946749713 | -6.821778694 |
| **Q9BWD1** | ACAT2 | 0.023396727 | 3.161854995 | 0.865210057 | 0.417839732 | 0.774192321 | -6.822073686 |
| **Q9UBU7** | DBF4 | -0.013212344 | 3.168904178 | -0.461543195 | 0.659519191 | 0.896852471 | -6.82253052 |
| **Q92610** | ZNF592 | -0.016762016 | 3.196677968 | -0.795337432 | 0.454593555 | 0.797515322 | -6.822547242 |
| **O14727-4** | APAF1 | 0.037951147 | 3.186778978 | 0.781238824 | 0.462278653 | 0.802800142 | -6.822586083 |
| **O75368** | SH3BGRL | 0.014082444 | 3.165781174 | 0.692350377 | 0.512784529 | 0.831227449 | -6.823454672 |
| **Q5T6V5** | C9orf64 | 0.017496992 | 3.182481326 | 0.617691392 | 0.557878187 | 0.851117349 | -6.823481835 |
| **P20674** | COX5A | -0.033080036 | 3.23451246 | -0.218490697 | 0.833775825 | 0.96068017 | -6.824743111 |
| **Q13409-1** | DYNC1I2 | -0.02142678 | 3.180589253 | -1.413470041 | 0.203769384 | 0.609084934 | -6.82496112 |
| **O95870-1** | ABHD16A | 0.019860848 | 3.182233537 | 0.607210757 | 0.564397481 | 0.852578515 | -6.825097226 |
| **Q9Y4D1** | DAAM1 | -0.010320415 | 3.173047871 | -0.330528182 | 0.751442236 | 0.935635612 | -6.825161148 |
| **Q9NSI8-3** | SAMSN1 | 0.048936393 | 3.212492001 | 1.094912619 | 0.312624361 | 0.700451368 | -6.825188307 |
| **P50583** | NUDT2 | 0.008572827 | 3.162250085 | 0.283227128 | 0.785847013 | 0.946704945 | -6.825762021 |
| **Q8N2K0-2** | ABHD12 | 0.010041817 | 3.202463455 | 0.264974049 | 0.799269414 | 0.952055634 | -6.825905655 |
| **O94913** | PCF11 | -0.021568658 | 3.191106665 | -1.459439742 | 0.191216928 | 0.595771959 | -6.825975859 |
| **Q9BWT6** | MND1 | -0.01949963 | 3.191313373 | -0.797468206 | 0.453439909 | 0.796326278 | -6.82636015 |
| **Q9ULJ6-1** | ZMIZ1 | -0.023387726 | 3.178221521 | -0.679115823 | 0.520603214 | 0.834650661 | -6.826455177 |
| **Q86U90** | YRDC | -0.028675768 | 3.246291679 | -0.54701847 | 0.602707102 | 0.872497827 | -6.826622202 |
| **Q9NYZ3** | GTSE1 | -0.027685789 | 3.186578351 | -0.694266878 | 0.511658642 | 0.830804805 | -6.826715003 |
| **P36969** | GPX4 | -0.033214904 | 3.187834268 | -1.02931631 | 0.340257977 | 0.722975002 | -6.826958143 |
| **Q96BY7** | ATG2B | -0.015402697 | 3.17481839 | -1.086107749 | 0.316223956 | 0.703753313 | -6.826987271 |
| **Q9HB21-1** | PLEKHA1 | -0.012146049 | 3.17057587 | -0.466684037 | 0.656026673 | 0.896497734 | -6.827282932 |
| **P41214** | EIF2D | -0.035110045 | 3.160429614 | -1.173408018 | 0.282004802 | 0.679261937 | -6.828320808 |
| **Q96RT7** | TUBGCP6 | 0.005997882 | 3.189744202 | 0.242784851 | 0.815685914 | 0.956267462 | -6.828539075 |
| **O14867** | BACH1 | 0.022061753 | 3.189382739 | 0.575860198 | 0.584168434 | 0.8653224 | -6.828786502 |
| **P51159-1** | RAB27A | 0.003691171 | 3.191652517 | 0.104639544 | 0.919830474 | 0.983742622 | -6.828842397 |
| **Q6P2P2** | PRMT10; PRMT9 | -0.022996027 | 3.175486894 | -0.991468227 | 0.357070244 | 0.734600528 | -6.82939516 |
| **Q5HYI8** | RABL3 | 0.034356074 | 3.188991423 | 0.538652135 | 0.608146109 | 0.875787718 | -6.830115568 |

| | | | | | | | |
|---|---|---|---|---|---|---|---|
| **P49354** | FNTA | -0.019134864 | 3.185473735 | -0.564911943 | 0.591166712 | 0.868764508 | -6.830566766 |
| **Q7LGA3-1** | HS2ST1 | -0.017115024 | 3.186933699 | -0.990826729 | 0.357360735 | 0.734600528 | -6.830645691 |
| **Q86UB9** | TMEM135 | 0.005345192 | 3.191323156 | 0.108337798 | 0.917009603 | 0.983235402 | -6.830825742 |
| **Q9UKL6** | PCTP | 0.000784151 | 3.183701395 | 0.028145332 | 0.978394345 | 0.993585863 | -6.830873578 |
| **Q9UGU5** | HMGXB4 | 0.004647514 | 3.166159814 | 0.115239815 | 0.911748553 | 0.981900848 | -6.830993995 |
| **Q13057-2** | COASY | 0.003699223 | 3.182793655 | 0.139164967 | 0.893550752 | 0.978276697 | -6.831384042 |
| **Q7Z333** | SETX | -0.032268594 | 3.193682468 | -1.396567075 | 0.2085695 | 0.614302382 | -6.831671738 |
| **Q9UK45** | LSM 7.00 | -0.026231032 | 3.154922923 | -0.564759141 | 0.591264724 | 0.868764508 | -6.83187298 |
| **Q96DY7** | MTBP | -0.004481098 | 3.219116554 | -0.125608478 | 0.903854207 | 0.980210603 | -6.831976206 |
| **Q14BN4** | SLMAP | 0.018848279 | 3.197956313 | 0.73890793 | 0.485891442 | 0.81473183 | -6.832111109 |
| **P13667** | PDIA4 | 0.028039373 | 3.202274332 | 1.829118792 | 0.113545822 | 0.487633273 | -6.832361569 |
| **P04844-1** | RPN2 | -0.058086447 | 3.178609818 | -1.155035208 | 0.288937261 | 0.683685579 | -6.832610264 |
| **P51148-2** | RAB5C | 0.011942554 | 3.17290555 | 0.472927265 | 0.651797804 | 0.894547168 | -6.832647301 |
| **Q8IYM9-1** | TRIM22 | -0.012301641 | 3.195480087 | -0.30813691 | 0.767657782 | 0.941217716 | -6.833052648 |
| **Q9UKT7** | FBXL3 | 0.002771419 | 3.172248859 | 0.06626888 | 0.949164149 | 0.989472196 | -6.833611299 |
| **P09110-1** | ACAA1 | -0.016762977 | 3.173496635 | -0.916888059 | 0.392089545 | 0.759806823 | -6.834309889 |
| **Q9BUT1-1** | BDH2 | 0.033985491 | 3.157970215 | 0.892900959 | 0.403890257 | 0.766336431 | -6.834884754 |
| **Q8IU60** | DCP2 | -0.017865145 | 3.180130275 | -0.935189302 | 0.383262438 | 0.754258979 | -6.834904155 |
| **Q06587** | RING1 | 0.015125423 | 3.181424222 | 0.498673521 | 0.634506832 | 0.887527602 | -6.835308294 |
| **Q9NVZ3** | NECAP2 | 0.017091258 | 3.170531544 | 0.599729205 | 0.569079088 | 0.855040807 | -6.835616375 |
| **Q8IU85-1** | CAMK1D | 0.028149235 | 3.200620925 | 0.848876594 | 0.426232014 | 0.779649091 | -6.835934014 |
| **Q9NQC3** | RTN4 | 0.054723728 | 3.225330977 | 1.165830003 | 0.284847005 | 0.68101215 | -6.835997624 |
| **Q8TCA0-1** | LRRC20 | -0.00140709 | 3.162906049 | -0.033308844 | 0.974432155 | 0.99316605 | -6.836614509 |
| **Q13427** | PPIG | -0.001276711 | 3.179184013 | -0.031199693 | 0.976050511 | 0.99316605 | -6.837112801 |
| **Q69YN2-1** | CWF19L1 | -0.015031559 | 3.168721205 | -1.247621166 | 0.255420217 | 0.659782078 | -6.837169164 |
| **P50990** | CCT8 | 0.000838609 | 3.192057364 | 0.030845818 | 0.976322052 | 0.99316605 | -6.837426825 |
| **Q9BW91-1** | NUDT9 | 0.016355132 | 3.199802892 | 0.960978016 | 0.371082402 | 0.744717822 | -6.838220435 |
| **Q9H6Q4** | NARFL | 0.016282125 | 3.209606315 | 0.249732865 | 0.810534169 | 0.955498404 | -6.838238944 |
| **P51157-1** | RAB28 | -0.013469764 | 3.299972674 | -0.57171382 | 0.586813203 | 0.866100748 | -6.838559726 |
| **Q13952-1** | NFYC | -0.040662435 | 3.193392871 | -0.798837638 | 0.452699557 | 0.795729641 | -6.83858348 |
| **Q8N3D4** | EHBP1L1 | -0.000694388 | 3.180812986 | -0.019626605 | 0.984932533 | 0.995531556 | -6.839520239 |
| **P17028-1** | ZNF24 | 0.006144465 | 3.206903363 | 0.169739127 | 0.870398493 | 0.971798205 | -6.839612207 |
| **Q9H0N5** | PCBD2 | -0.006452633 | 3.156379085 | -0.114936203 | 0.911979881 | 0.981900848 | -6.839864874 |
| **P53602** | MVD | 0.01799385 | 3.16699715 | 0.789470812 | 0.457780497 | 0.799355176 | -6.84100519 |
| **P63302** | SEPW1; SELENOW | 0.01083182 | 3.203210639 | 0.293064814 | 0.778645306 | 0.944053983 | -6.841308549 |
| **Q9HAU0-6** | PLEKHA5 | 0.025370279 | 3.175978929 | 0.762098354 | 0.472855741 | 0.808597546 | -6.841920867 |
| **P17544-6** | ATF7 | -0.011695466 | 3.184379572 | -0.360711297 | 0.729800022 | 0.927974736 | -6.842129578 |
| **P55263-1** | ADK | 0.018121678 | 3.157346209 | 1.264530368 | 0.249673831 | 0.65453932 | -6.842646313 |
| **P28065** | PSMB9 | 0.014029295 | 3.194279585 | 0.615267745 | 0.559381712 | 0.851665881 | -6.842680004 |

| | | | | | | | |
|---|---|---|---|---|---|---|---|
| **Q09161** | NCBP1 | -0.021990487 | 3.175444095 | -1.378926989 | 0.213687291 | 0.619841376 | -6.842890253 |
| **Q86W92-1** | PPFIBP1 | 0.016324373 | 3.199277822 | 0.494325975 | 0.637409638 | 0.889065271 | -6.844030449 |
| **P41567** | EIF1 | -0.038595204 | 3.22056729 | -1.126860535 | 0.299844987 | 0.690882899 | -6.844628508 |
| **P12931-2** | SRC | -0.023439764 | 3.151644437 | -0.358828932 | 0.731142174 | 0.92815165 | -6.845294494 |
| **Q6ZUM4-1** | ARHGAP27 | -0.01498767 | 3.171942682 | -0.958678256 | 0.372156328 | 0.745139764 | -6.846107293 |
| **Q6IE81-1** | PHF17; JADE1 | 0.011966052 | 3.199930988 | 0.536002234 | 0.6098745 | 0.876846345 | -6.846276118 |
| **Q5PRF9** | SAMD4B | -0.00207214 | 3.166925051 | -0.065337289 | 0.949877605 | 0.989472196 | -6.846379255 |
| **Q8IUI8** | CRLF3 | -0.014356531 | 3.166964465 | -0.435253024 | 0.677523288 | 0.903662615 | -6.846671218 |
| **P14618-2** | PKM | -0.001121938 | 3.181213579 | -0.038388318 | 0.970535219 | 0.992521442 | -6.846925968 |
| **Q9Y6B7** | AP4B1 | -0.021938278 | 3.18114206 | -0.65315763 | 0.536159231 | 0.842021168 | -6.847233184 |
| **Q9HAN9** | NMNAT1 | 0.014570073 | 3.18174608 | 0.274814295 | 0.79202386 | 0.948542293 | -6.847236632 |
| **Q9BQF6** | SENP7 | 0.001912734 | 3.167309663 | 0.078375765 | 0.939897006 | 0.988002681 | -6.847978561 |
| **Q07617** | SPAG1 | 0.001994334 | 3.220398234 | 0.062315189 | 0.952192413 | 0.989472196 | -6.848387411 |
| **P11940-1** | PABPC1 | -0.037974986 | 3.201934398 | -1.551403644 | 0.168207539 | 0.566515005 | -6.848793823 |
| **Q01780** | EXOSC10 | -0.022543973 | 3.183485343 | -1.444910784 | 0.195105978 | 0.600645673 | -6.848873915 |
| **Q8NFY9** | KBTBD8 | -0.009880837 | 3.225195123 | -0.361844108 | 0.728992809 | 0.927586142 | -6.849061601 |
| **Q96S15-2** | WDR24 | 0.004262839 | 3.169964054 | 0.099550194 | 0.923714463 | 0.985083485 | -6.850003316 |
| **Q99961** | SH3GL1 | -0.020028391 | 3.192688584 | -0.83123451 | 0.435433447 | 0.784298014 | -6.850487183 |
| **O75093-1** | SLIT1 | 0.022788276 | 3.191011637 | 0.721647927 | 0.495749455 | 0.822797039 | -6.85193436 |
| **P50148** | GNAQ | 0.015394378 | 3.18361259 | 0.162754629 | 0.875676224 | 0.972759481 | -6.852242533 |
| **P04080** | CSTB | -0.040870482 | 3.185183658 | -0.848635756 | 0.42635667 | 0.779649091 | -6.852756811 |
| **Q8N302-1** | AGGF1 | -0.017603176 | 3.187472173 | -0.698156106 | 0.509378782 | 0.82913633 | -6.852775207 |
| **Q13613-1** | MTMR1 | 0.042753195 | 3.207767081 | 0.930608805 | 0.385457411 | 0.754896271 | -6.85354897 |
| **Q5MIZ7-1** | SMEK2; PPP4R3B | -0.018056338 | 3.204954081 | -0.907676987 | 0.396589986 | 0.762021989 | -6.853748841 |
| **Q9Y679-1** | AUP1 | -0.023673938 | 3.202400274 | -0.736366302 | 0.48733476 | 0.816334529 | -6.853888085 |
| **Q8NCE0-1** | TSEN2 | 0.01332456 | 3.19916758 | 0.451154362 | 0.666605255 | 0.90001396 | -6.854250362 |
| **O14737** | PDCD5 | -0.059703287 | 3.17745999 | -0.845583088 | 0.427939002 | 0.780598411 | -6.854434507 |
| **Q6PIY7-1** | PAPD4 | -0.014727897 | 3.162740805 | -0.589196208 | 0.575709151 | 0.860065716 | -6.854532743 |
| **Q5RKV6** | EXOSC6 | -0.015677921 | 3.196219892 | -0.630185312 | 0.550166504 | 0.846778937 | -6.855075495 |
| **Q8IYB4-3** | PEX5L | 0.039540742 | 3.201870186 | 0.604674629 | 0.565981882 | 0.853395233 | -6.855987338 |
| **Q8TDH9** | BLOC1S5 | 0.004085198 | 3.191965844 | 0.056501533 | 0.956646835 | 0.989472196 | -6.856711529 |
| **P29320** | EPHA3 | 0.015765491 | 3.208696428 | 0.632079223 | 0.549003241 | 0.84662864 | -6.857568287 |
| **O60333-2** | KIF1B | 0.020845891 | 3.147044774 | 0.778823826 | 0.463604079 | 0.802800142 | -6.857603297 |
| **Q9NUA8-1** | ZBTB40 | 0.037237743 | 3.181987971 | 0.773501919 | 0.466534201 | 0.804518266 | -6.857716789 |
| **Q9BYE7-1** | PCGF6 | 0.000805273 | 3.199701494 | 0.013762774 | 0.989433835 | 0.996219143 | -6.858110122 |
| **O14657** | TOR1B | 0.022378932 | 3.189815257 | 0.588404513 | 0.576209322 | 0.860523132 | -6.858261033 |
| **O95379-4** | TNFAIP8 | -0.022166447 | 3.157225854 | -0.540386011 | 0.607016667 | 0.87523889 | -6.858357758 |
| **Q8TE02-4** | ELP5 | 0.003258436 | 3.203149554 | 0.094983981 | 0.927201149 | 0.985631941 | -6.858456054 |
| **Q96A33-1** | CCDC47 | -0.021690003 | 3.186535139 | -1.217492217 | 0.265942366 | 0.668906358 | -6.858582458 |

| | | | | | | | |
|---|---|---|---|---|---|---|---|
| **P46782** | RPS5 | -0.037438377 | 3.198315938 | -1.136046635 | 0.296251562 | 0.688880164 | -6.858792791 |
| **Q9Y4C4** | MFHAS1 | 0.019423479 | 3.18542706 | 0.821486835 | 0.440578311 | 0.788895128 | -6.858809409 |
| **Q7Z3K3-1** | POGZ | 0.021904235 | 3.174134717 | 1.323576876 | 0.230482185 | 0.638264538 | -6.858893961 |
| **Q8N3C7-1** | CLIP4 | 0.002203892 | 3.19593723 | 0.076065544 | 0.941664615 | 0.988377491 | -6.859232082 |
| **Q147X3** | NAA30 | -0.009622426 | 3.18921376 | -0.214491553 | 0.836764553 | 0.960912428 | -6.859621656 |
| **Q9BVC4** | MLST8 | 0.005075839 | 3.181852238 | 0.170217046 | 0.87003762 | 0.971622708 | -6.859636622 |
| **Q9NRR5-1** | UBQLN4 | 0.085520116 | 3.211717814 | 0.744261932 | 0.482860495 | 0.812781066 | -6.860102158 |
| **O15304** | SIVA1 | -0.014022763 | 3.17423657 | -0.379089422 | 0.716751005 | 0.924679646 | -6.86025199 |
| **Q9NS73** | MBIP | -0.006307536 | 3.162299112 | -0.088287529 | 0.932317582 | 0.986793807 | -6.860272647 |
| **Q92615** | LARP4B | 0.01903296 | 3.179755154 | 1.151906152 | 0.29013206 | 0.684803131 | -6.860336339 |
| **Q9UK61** | FAM208A | 0.017615384 | 3.186701745 | 1.494779539 | 0.182051696 | 0.584756817 | -6.860511557 |
| **O00170** | AIP | 0.093262701 | 3.183527121 | 0.984375138 | 0.360292523 | 0.73661838 | -6.860517141 |
| **O00754** | MAN2B1 | -0.021993545 | 3.191527133 | -0.635430784 | 0.546948383 | 0.845922698 | -6.861114814 |
| **Q15005** | SPCS2 | -0.048109533 | 3.157443001 | -0.411950647 | 0.693676018 | 0.910349515 | -6.861218222 |
| **Q9BSJ2-4** | TUBGCP2 | 0.023253761 | 3.185151432 | 1.386228481 | 0.211555436 | 0.61590617 | -6.861411028 |
| **Q9P2A4** | ABI3 | -0.009840335 | 3.180226958 | -0.356074718 | 0.733107815 | 0.928509739 | -6.861417458 |
| **P50416** | CPT1A | 0.012507308 | 3.199322797 | 0.326243226 | 0.754535101 | 0.936977396 | -6.861629836 |
| **Q9BVS4** | RIOK2 | 0.023518764 | 3.188447675 | 0.831842889 | 0.435113777 | 0.784298014 | -6.86175996 |
| **Q14671-1** | PUM1 | 0.018418449 | 3.182214902 | 1.040171747 | 0.335553902 | 0.720533987 | -6.861966925 |
| **Q9P2T1-1** | GMPR2 | 0.029511749 | 3.176765364 | 0.96528973 | 0.369075401 | 0.743673361 | -6.862453901 |
| **Q9H583** | HEATR1 | -0.02804133 | 3.181404749 | -1.637710215 | 0.148984747 | 0.546231571 | -6.863307601 |
| **Q6NW34** | C3orf17; NEPRO | 0.004494509 | 3.176312966 | 0.145751137 | 0.888552946 | 0.976290023 | -6.864395775 |
| **Q9BSV6** | TSEN34 | -0.014962911 | 3.171923502 | -0.338734535 | 0.745532839 | 0.934974996 | -6.865315108 |
| **Q9Y6Y8** | SEC23IP | -0.013609498 | 3.176659345 | -1.392745023 | 0.209668919 | 0.614920472 | -6.86564948 |
| **Q9HAU4** | SMURF2 | -0.002763221 | 3.192506834 | -0.078162937 | 0.940059831 | 0.988002681 | -6.865961916 |
| **Q99615-1** | DNAJC7 | -0.019778188 | 3.189769279 | -1.505465617 | 0.179360918 | 0.580990067 | -6.866198688 |
| **Q8IYS1** | PM20D2 | 0.023674117 | 3.1782404 | 0.807152405 | 0.448222636 | 0.793124464 | -6.866861436 |
| **Q9UI12-1** | ATP6V1H | -0.018716291 | 3.190658586 | -1.194280249 | 0.274300038 | 0.674084072 | -6.867484454 |
| **Q8TBM8** | DNAJB14 | 0.01259963 | 3.203212397 | 0.47740708 | 0.648771954 | 0.893481222 | -6.867883673 |
| **Q7Z4H3-1** | HDDC2 | 0.010493714 | 3.150587716 | 0.462831816 | 0.658642871 | 0.896852471 | -6.868508989 |
| **Q9NP79** | VTA1 | 0.019058794 | 3.182530287 | 1.124741222 | 0.300679137 | 0.691292041 | -6.868581174 |
| **O75037** | KIF21B | -0.009334016 | 3.210014451 | -0.27964619 | 0.788474176 | 0.947805127 | -6.869524026 |
| **Q96DF8** | DGCR14; ESS2 | -0.017938668 | 3.16266861 | -0.406933373 | 0.697177093 | 0.912576571 | -6.869547023 |
| **Q9NZ08-2** | ERAP1 | -0.020471584 | 3.179804135 | -1.303031505 | 0.237007749 | 0.643518711 | -6.869548851 |
| **P61313-1** | RPL15 | -0.051188882 | 3.145729703 | -1.013392358 | 0.34725336 | 0.72891095 | -6.869900211 |
| **P49916-1** | LIG3 | 0.043015366 | 3.177541602 | 1.564101548 | 0.165240392 | 0.564660179 | -6.870315345 |
| **Q8WVY7** | UBLCP1 | 0.029335891 | 3.161381961 | 0.692794049 | 0.512523742 | 0.831227449 | -6.871007831 |
| **P07311** | ACYP1 | -0.001468429 | 3.171002405 | -0.038389341 | 0.970534434 | 0.992521442 | -6.871521557 |
| **Q9UJ70-2** | NAGK | -0.020683626 | 3.174498421 | -0.989939185 | 0.357762949 | 0.734856937 | -6.874047285 |

| | | | | | | | |
|---|---|---|---|---|---|---|---|
| **Q13099-1** | IFT88 | -0.010226008 | 3.190060062 | -0.536593445 | 0.609488649 | 0.876646363 | -6.874386365 |
| **Q9Y2X0-1** | MED16 | 0.055977899 | 3.197172415 | 0.981121767 | 0.361778061 | 0.737384991 | -6.874621527 |
| **Q8NCA5** | FAM98A | -0.02172536 | 3.195609193 | -0.722228359 | 0.495415792 | 0.822586931 | -6.875099135 |
| **P32929** | CTH | -0.003886848 | 3.162402543 | -0.142113281 | 0.891312807 | 0.977905675 | -6.875506707 |
| **P16035** | TIMP2 | 0.008117884 | 3.161211294 | 0.176174481 | 0.865542068 | 0.970330426 | -6.875940713 |
| **Q15067** | ACOX1 | -0.031254876 | 3.156756365 | -0.88347348 | 0.408600096 | 0.768662396 | -6.876302373 |
| **Q504Q3-2** | PAN2 | -0.040989545 | 3.213287595 | -0.642486633 | 0.542637947 | 0.843669625 | -6.876478686 |
| **Q9Y320** | TMX2 | -0.024186045 | 3.163171158 | -0.429884339 | 0.681228813 | 0.903980599 | -6.876642938 |
| **Q8TAK5** | GABPB2 | 0.028467161 | 3.212202276 | 0.373377552 | 0.720795835 | 0.925644897 | -6.877884054 |
| **Q9P260-1** | KIAA1468 | 0.003266231 | 3.159018746 | 0.071082905 | 0.945478167 | 0.98904859 | -6.87815818 |
| **Q9UDY2-7** | TJP2 | -0.000163276 | 3.16588531 | -0.006335249 | 0.995136067 | 0.99815125 | -6.878704906 |
| **P14635** | CCNB1 | -0.019588381 | 3.170514021 | -0.488276591 | 0.641460294 | 0.890911725 | -6.878726065 |
| **Q96CN9** | GCC1 | -0.03263076 | 3.204564695 | -1.199285152 | 0.272479306 | 0.672884265 | -6.878855832 |
| **P19838-2** | NFKB1 | 0.026995665 | 3.163697763 | 0.860266872 | 0.420366732 | 0.77652027 | -6.879103668 |
| **Q8TAP6** | CEP76 | 0.014388139 | 3.184971838 | 0.462268972 | 0.659025558 | 0.896852471 | -6.879183278 |
| **Q9NZ32** | ACTR10 | 0.015395206 | 3.157478992 | 0.524778796 | 0.617224974 | 0.881353777 | -6.879217396 |
| **Q8N3P4-1** | VPS8 | 0.032013392 | 3.183944528 | 0.944576799 | 0.378793763 | 0.750267026 | -6.879459175 |
| **Q96F15** | GIMAP5 | -0.000917201 | 3.153793514 | -0.030603435 | 0.976508044 | 0.99316605 | -6.8794917 |
| **Q92609-2** | TBC1D5 | -0.016571323 | 3.190252816 | -1.232973967 | 0.260489935 | 0.663969372 | -6.879949402 |
| **Q03164-3** | KMT2A; MLL | -0.0278594 | 3.188398581 | -1.051302458 | 0.33078479 | 0.716489955 | -6.880944317 |
| **Q9Y2S0-1** | POLR1D | -0.002312337 | 3.195903954 | -0.078761776 | 0.939601694 | 0.988002681 | -6.881131294 |
| **Q8TF46** | DIS3L | -0.018703706 | 3.198575378 | -0.944429439 | 0.378863601 | 0.750267026 | -6.882214613 |
| **Q9UH62** | ARMCX3 | 0.015059095 | 3.18912137 | 0.420062768 | 0.688032581 | 0.907741425 | -6.882224676 |
| **Q96F44-1** | TRIM11 | 0.015819133 | 3.234651959 | 0.250073907 | 0.810281557 | 0.955498404 | -6.882236859 |
| **Q7L2Z9** | CENPQ | 0.018091507 | 3.166004843 | 0.746141594 | 0.48179944 | 0.81235548 | -6.883058524 |
| **Q8NHV1** | GIMAP7 | 0.025268253 | 3.174511269 | 0.46344891 | 0.658223426 | 0.896852471 | -6.885009123 |
| **O76071** | CIAO1 | -0.011041913 | 3.173849633 | -0.50337882 | 0.631373035 | 0.887152592 | -6.885343252 |
| **Q14651** | PLS1 | -0.015923656 | 3.181475529 | -1.099143005 | 0.310906874 | 0.699034299 | -6.885563217 |
| **Q9ULE6** | PALD1 | 0.01857548 | 3.205302598 | 0.507176606 | 0.628849658 | 0.886167175 | -6.885902733 |
| **O75394** | MRPL33 | 0.003728039 | 3.19502733 | 0.088686473 | 0.932012668 | 0.986711787 | -6.886020896 |
| **P60866** | RPS20 | -0.050336022 | 3.19448892 | -1.030628724 | 0.339686475 | 0.722416219 | -6.886098651 |
| **Q92572** | AP3S1 | -0.017012052 | 3.192705313 | -0.854467041 | 0.423345882 | 0.777884596 | -6.886616953 |
| **O60583** | CCNT2 | 0.014611718 | 3.193399074 | 0.202235674 | 0.845941977 | 0.963216816 | -6.887309684 |
| **Q16881-1** | TXNRD1 | 0.0235619 | 3.174674547 | 1.321135665 | 0.23124917 | 0.638769945 | -6.888164255 |
| **O15066** | KIF3B | -0.022452764 | 3.179275796 | -0.876900484 | 0.411907888 | 0.770569383 | -6.888261887 |
| **Q92545** | TMEM131 | 0.005826369 | 3.197839919 | 0.12161181 | 0.906895784 | 0.98075504 | -6.888850315 |
| **Q04864** | REL | -0.021476377 | 3.183777695 | -0.953877463 | 0.374405893 | 0.746861238 | -6.889321149 |
| **Q9NQT8** | KIF13B | -0.015814238 | 3.17932472 | -1.376054377 | 0.21453129 | 0.619913059 | -6.889325045 |
| **Q14376** | GALE | 0.023714329 | 3.206212344 | 0.876509244 | 0.412105398 | 0.770575992 | -6.889446681 |

| | | | | | | | |
|---|---|---|---|---|---|---|---|
| **P13994** | CCDC130 | 0.015932121 | 3.193222643 | 0.500903229 | 0.633020791 | 0.887490806 | -6.889551833 |
| **Q01650** | SLC7A5 | -0.013232857 | 3.224111414 | -0.265032957 | 0.799225973 | 0.952055634 | -6.889624971 |
| **Q9UL26** | RAB22A | 0.014701233 | 3.183972864 | 0.419819548 | 0.688201472 | 0.907813498 | -6.889639042 |
| **Q9UIW2** | PLXNA1 | 0.012714054 | 3.193654901 | 0.523654262 | 0.617964117 | 0.881802191 | -6.890075818 |
| **Q969P0** | IGSF8 | -0.029317623 | 3.177374763 | -0.757817676 | 0.475243829 | 0.809713018 | -6.890224627 |
| **Q99436** | PSMB7 | -0.058953858 | 3.145089044 | -1.050101918 | 0.331296542 | 0.716856834 | -6.890600619 |
| **Q5VW32** | BROX | 0.027748243 | 3.197390202 | 0.474112598 | 0.650996483 | 0.894219754 | -6.890703993 |
| **Q96F63** | CCDC97 | -0.01264724 | 3.207964737 | -0.29653157 | 0.776113049 | 0.94357883 | -6.893403732 |
| **Q9Y5L4** | TIMM13 | -0.0595564 | 3.172659067 | -0.735238342 | 0.487976219 | 0.816892343 | -6.893470283 |
| **P09104** | ENO2 | -0.044215734 | 3.190827513 | -0.732293584 | 0.489653545 | 0.81832087 | -6.894279681 |
| **A0AVF1** | TTC26 | 0.02575665 | 3.177604554 | 0.511642778 | 0.625889083 | 0.885387512 | -6.894750628 |
| **Q9UDY8** | MALT1 | 0.017903989 | 3.200108024 | 0.509658068 | 0.627203802 | 0.885387512 | -6.895414931 |
| **P07900-2** | HSP90AA1 | 0.03337897 | 3.193287131 | 1.877429519 | 0.105984305 | 0.475403904 | -6.895485142 |
| **Q7Z6K3** | PTAR1 | 0.013267145 | 3.186746689 | 0.55120068 | 0.599998477 | 0.872116337 | -6.896459648 |
| **Q96KM6** | ZNF512B | -0.014702949 | 3.171526588 | -0.2226347 | 0.830681995 | 0.960067847 | -6.897390953 |
| **Q9BUK6** | MSTO1 | 0.005335107 | 3.169809675 | 0.215084179 | 0.836321472 | 0.960912428 | -6.89739247 |
| **P16278** | GLB1 | 0.011822472 | 3.184817146 | 0.617329285 | 0.558102667 | 0.851117349 | -6.897465368 |
| **Q8WVR3** | C7orf43 | -0.000330623 | 3.192790297 | -0.014769354 | 0.98866111 | 0.996184213 | -6.897990885 |
| **Q9BX67** | JAM3 | -0.043966921 | 3.185477269 | -0.498869203 | 0.634376341 | 0.887509451 | -6.898554047 |
| **Q8WUF8** | FAM172A | 0.005112128 | 3.179734972 | 0.185502535 | 0.858513958 | 0.968855822 | -6.898643229 |
| **Q9BTY7** | FAM203B; FAM203 | -0.038591637 | 3.197337508 | -0.835283711 | 0.433308989 | 0.783962934 | -6.899063975 |
| **Q9Y2Q5-2** | LAMTOR2 | 0.01176889 | 3.163503904 | 0.377887312 | 0.717601456 | 0.924679646 | -6.899201284 |
| **Q8N5G2-1** | TMEM57; MACO1 | -0.05732566 | 3.156714634 | -0.625344507 | 0.553146646 | 0.848243603 | -6.899345194 |
| **Q9NV56** | MRGBP | 0.009617247 | 3.173541242 | 0.248957678 | 0.811108446 | 0.955498404 | -6.900395901 |
| **Q9H9A7** | RMI1 | -0.002703566 | 3.196680221 | -0.07915258 | 0.939302727 | 0.988002681 | -6.900559736 |
| **Q8IVH2-1** | FOXP4 | -0.003789139 | 3.213228737 | -0.171897176 | 0.868769235 | 0.97108181 | -6.900567606 |
| **O00115** | DNASE2 | -0.034460824 | 3.181881906 | -0.728076089 | 0.492062541 | 0.820448454 | -6.901956814 |
| **O14618** | CCS | -0.006958439 | 3.168415186 | -0.225758335 | 0.828352109 | 0.959077751 | -6.902117005 |
| **Q9C0G0** | ZNF407 | -0.010297996 | 3.183901042 | -0.328699435 | 0.752761614 | 0.935782019 | -6.902215417 |
| **P63000-2** | RAC1 | 0.01748243 | 3.159762393 | 0.731748407 | 0.4899645 | 0.818496205 | -6.902255932 |
| **O43633** | CHMP2A | -0.04010483 | 3.159063989 | -0.591290077 | 0.574387536 | 0.85911412 | -6.902473614 |
| **Q9Y613** | FHOD1 | 0.018043132 | 3.184589805 | 0.715967039 | 0.499023017 | 0.824970816 | -6.902529191 |
| **O00214-2** | LGALS8 | -0.041382478 | 3.163460595 | -0.65759309 | 0.533480633 | 0.840132297 | -6.902695737 |
| **Q92598** | HSPH1 | -0.018940992 | 3.172317951 | -0.357002521 | 0.732445412 | 0.928164811 | -6.902900198 |
| **Q5SZL2-4** | CEP85L | 0.002990002 | 3.177452925 | 0.089576156 | 0.931332726 | 0.986707862 | -6.903381953 |
| **Q9H7Z6** | KAT8 | 0.015928295 | 3.144134904 | 1.002283683 | 0.352200503 | 0.732738778 | -6.903910925 |
| **P51570-2** | GALK1 | 0.006518315 | 3.156979715 | 0.157643848 | 0.8795425 | 0.97338977 | -6.903986795 |
| **Q12982-2** | BNIP2 | 0.019832797 | 3.168283774 | 0.419131086 | 0.688679643 | 0.908052188 | -6.904154554 |
| **O60508** | CDC40 | -0.028576028 | 3.186235056 | -1.203735909 | 0.270868814 | 0.671838876 | -6.904205178 |

| | | | | | | | |
|---|---|---|---|---|---|---|---|
| **P51946** | CCNH | 0.013415701 | 3.206349787 | 0.617202274 | 0.558181417 | 0.851117349 | -6.904326603 |
| **Q9Y5J7** | TIMM9 | -0.032173214 | 3.206127941 | -0.222578787 | 0.830723717 | 0.960067847 | -6.905084625 |
| **P05161** | ISG15 | 0.007693741 | 3.199274216 | 0.272855615 | 0.793464316 | 0.949122853 | -6.905657687 |
| **Q9UBP6** | METTL1 | -0.010681872 | 3.177410737 | -0.552819348 | 0.598951985 | 0.871871293 | -6.905692561 |
| **P36543** | ATP6V1E1 | 0.029558561 | 3.147022863 | 0.82377354 | 0.439367493 | 0.787985266 | -6.905930721 |
| **Q9UBW8** | COPS7A | -0.012335018 | 3.17548457 | -0.678996935 | 0.520673796 | 0.834650661 | -6.906011755 |
| **Q92834** | RPGR | -0.002286193 | 3.196969676 | -0.087291489 | 0.933078912 | 0.986793807 | -6.906078161 |
| **Q14746** | COG2 | 0.023642142 | 3.184529651 | 0.800961555 | 0.451552992 | 0.79513757 | -6.906167141 |
| **Q92600-2** | RQCD1; CNOT9 | -0.01996722 | 3.180509391 | -0.82026628 | 0.441225573 | 0.788971713 | -6.906620625 |
| **Q16204** | CCDC6 | -0.023582199 | 3.192271307 | -1.025661625 | 0.341853481 | 0.724080487 | -6.906851282 |
| **Q14686** | NCOA6 | -0.021472647 | 3.193961568 | -0.631755956 | 0.549201689 | 0.846638465 | -6.906945119 |
| **O95475** | SIX6 | -0.015760518 | 3.207308283 | -0.55322595 | 0.598689273 | 0.871871293 | -6.90764173 |
| **Q9UBS4** | DNAJB11 | 0.035304591 | 3.168210213 | 1.101319405 | 0.310026303 | 0.698174312 | -6.907976298 |
| **Q14186** | TFDP1 | -0.007627142 | 3.194480405 | -0.29601167 | 0.776492617 | 0.94357883 | -6.908706049 |
| **Q96K17-1** | BTF3L4 | 0.035550285 | 3.181469911 | 0.794332652 | 0.455138278 | 0.797877083 | -6.909119126 |
| **Q14807** | KIF22 | -0.03476952 | 3.126338305 | -1.240050606 | 0.258029827 | 0.660717666 | -6.909620052 |
| **P49005** | POLD2 | -0.019836462 | 3.182940598 | -0.891704082 | 0.404485952 | 0.766336431 | -6.910012404 |
| **Q0VDG4-1** | SCRN3 | -0.011605228 | 3.169384611 | -0.289788843 | 0.781040901 | 0.944918194 | -6.910076772 |
| **Q96MF7** | NSMCE2 | -0.019473927 | 3.177736625 | -0.561385561 | 0.593431024 | 0.869935985 | -6.910176527 |
| **Q9Y232-1** | CDYL | -0.024132904 | 3.16663938 | -0.324685254 | 0.755660861 | 0.937349811 | -6.91036602 |
| **Q13227** | GPS2 | 0.017937231 | 3.180796921 | 0.262201996 | 0.801314459 | 0.952917733 | -6.91038119 |
| **Q8N954-1** | CCDC75; GPATCH1 | 0.001577375 | 3.190944831 | 0.049986293 | 0.961640807 | 0.991283985 | -6.91071015 |
| **Q6ZU80** | CEP128 | 0.0061928 | 3.19050542 | 0.293250363 | 0.778509699 | 0.944053983 | -6.911074756 |
| **Q8TC07-1** | TBC1D15 | 0.014913661 | 3.178551238 | 1.20429757 | 0.270666156 | 0.671731232 | -6.911244462 |
| **Q9BVI4** | NOC4L | -0.027320261 | 3.179601867 | -1.175567963 | 0.281199087 | 0.678447235 | -6.911873425 |
| **P13056-1** | NR2C1 | 0.006591227 | 3.19919833 | 0.262596412 | 0.80102338 | 0.952917733 | -6.911892188 |
| **Q9Y4G6** | TLN2 | 0.018135628 | 3.212534624 | 0.870807742 | 0.414991621 | 0.77203561 | -6.912248936 |
| **O60293** | ZFC3H1 | -0.016267867 | 3.193245682 | -0.873854568 | 0.413447405 | 0.771122868 | -6.91245297 |
| **Q9BRP8** | WIBG; PYM1 | -0.053720721 | 3.149516323 | -0.680788114 | 0.519611049 | 0.834217303 | -6.912575057 |
| **Q86XN7** | PROSER1 | 0.001074282 | 3.184978386 | 0.035481168 | 0.972765464 | 0.992928787 | -6.912599418 |
| **Q96RQ1** | ERGIC2 | -0.012219911 | 3.187044277 | -0.335660025 | 0.747744635 | 0.934974996 | -6.913055021 |
| **Q8WUH2** | TGFBRAP1 | 0.001579173 | 3.186597118 | 0.064850053 | 0.950250773 | 0.989472196 | -6.913300402 |
| **Q6ZSR9** | | -0.000676999 | 3.179141691 | -0.023266844 | 0.982138435 | 0.994932161 | -6.913448528 |
| **O95394-3** | PGM3 | -0.012548487 | 3.172838795 | -0.450848502 | 0.666814445 | 0.90001396 | -6.913468139 |
| **O95551-2** | TDP2 | -0.019427147 | 3.178868378 | -0.579593198 | 0.581793263 | 0.864278602 | -6.914221024 |
| **Q96EX3** | WDR34 | -0.010430384 | 3.199010524 | -0.425385635 | 0.684341273 | 0.905978473 | -6.914583878 |
| **O15392** | BIRC5 | -0.017711191 | 3.190088184 | -0.570238201 | 0.587756093 | 0.866850224 | -6.914863537 |
| **P0CAP2-1** | POLR2M | 0.004634072 | 3.170861882 | 0.131503664 | 0.899371028 | 0.978757548 | -6.915558377 |
| **Q9GZT4** | SRR | 0.005342382 | 3.20016455 | 0.089675627 | 0.931256709 | 0.986707862 | -6.916359349 |

| | | | | | | | |
|---|---|---|---|---|---|---|---|
| **Q8TC05** | MDM1 | -0.003325539 | 3.1960976 | -0.102815427 | 0.921222299 | 0.984164603 | -6.917534139 |
| **Q9UN37** | VPS4A | -0.034577324 | 3.165956176 | -0.921155074 | 0.39001783 | 0.75855365 | -6.917538185 |
| **Q12972-1** | PPP1R8 | 0.0541344 | 3.156400365 | 1.023372895 | 0.342855688 | 0.724657739 | -6.917835129 |
| **Q9Y2G5** | POFUT2 | -0.019984382 | 3.202546022 | -0.330546316 | 0.751429157 | 0.935635612 | -6.91834672 |
| **Q6P1X5** | TAF2 | -0.023205161 | 3.180440392 | -0.95302401 | 0.374806901 | 0.747450006 | -6.918400907 |
| **Q8NB16** | MLKL | -0.013424747 | 3.203596884 | -0.889143021 | 0.405762807 | 0.766336431 | -6.918680894 |
| **Q8N5A5-2** | ZGPAT | -0.017743243 | 3.20234581 | -0.950805291 | 0.375850942 | 0.748684576 | -6.919440745 |
| **Q9UPN6** | SCAF8 | 0.018350855 | 3.17978747 | 1.132598861 | 0.297596045 | 0.689530645 | -6.920011275 |
| **P35754** | GLRX | -0.019016033 | 3.159761892 | -0.727879861 | 0.492174817 | 0.820463475 | -6.920133391 |
| **Q5H9R7-5** | PPP6R3 | -0.021712374 | 3.179446901 | -1.358707893 | 0.219691533 | 0.625609044 | -6.920567768 |
| **P21926** | CD9 | 0.021773461 | 3.216792476 | 0.10782439 | 0.917401131 | 0.983235402 | -6.921027486 |
| **Q86X95** | CIR1 | 0.04958376 | 3.100910564 | 0.414279279 | 0.692053848 | 0.909991705 | -6.921072194 |
| **Q8NBQ5** | HSD17B11 | -0.014050958 | 3.178365854 | -0.698325062 | 0.509279891 | 0.82913633 | -6.92109621 |
| **Q13243-1** | SRSF5 | -0.021505806 | 3.197947631 | -0.599485982 | 0.569231673 | 0.855085204 | -6.921388074 |
| **Q8IYH5-1** | ZZZ3 | -0.018825228 | 3.185585921 | -0.747734431 | 0.480901534 | 0.81235548 | -6.92152357 |
| **Q9BW62** | KATNAL1 | -0.001454256 | 3.160285523 | -0.041167487 | 0.968403424 | 0.992306703 | -6.921672397 |
| **P30443** | HLA-A | -0.006495958 | 3.192287571 | -0.088846852 | 0.931890093 | 0.986711787 | -6.921931509 |
| **O43242** | PSMD3 | -0.020673936 | 3.186670536 | -1.527800728 | 0.173855151 | 0.574454961 | -6.922337235 |
| **Q86YV9** | HPS6 | 0.013127021 | 3.18077367 | 0.702894658 | 0.506610021 | 0.82913633 | -6.922716476 |
| **Q86X55** | CARM1 | -0.01490969 | 3.189016198 | -1.170887378 | 0.282947527 | 0.679787821 | -6.922716799 |
| **Q15003** | NCAPH | -0.020348085 | 3.181920899 | -1.270679641 | 0.247612072 | 0.652280071 | -6.923046726 |
| **Q8NBI6** | XXYLT1 | -0.012448953 | 3.189381505 | -0.300745459 | 0.773039049 | 0.943005565 | -6.923602032 |
| **Q8N999-1** | C12orf29 | 0.018307306 | 3.20009316 | 0.674650117 | 0.523258655 | 0.836627689 | -6.924517759 |
| **Q9HAB8-1** | PPCS | -0.011472399 | 3.177241187 | -0.498244554 | 0.634792938 | 0.887591411 | -6.924522987 |
| **P33992** | MCM5 | 0.010721443 | 3.174041722 | 1.697760942 | 0.136856647 | 0.527828187 | -6.924589659 |
| **Q969M3-3** | YIPF5 | 0.010151263 | 3.209292659 | 0.399811444 | 0.70216062 | 0.914845949 | -6.924707126 |
| **Q9P219** | CCDC88C | -0.026209597 | 3.173960916 | -1.194593238 | 0.274185874 | 0.674011995 | -6.924976516 |
| **Q8NEC7-1** | GSTCD | -0.013865229 | 3.189136708 | -0.360573478 | 0.729898254 | 0.927974736 | -6.925246106 |
| **Q8TAG9** | EXOC6 | -0.016539725 | 3.197895973 | -0.841310156 | 0.430160991 | 0.782603406 | -6.926206525 |
| **Q86X27-1** | RALGPS2 | -0.011172368 | 3.21235469 | -0.459294152 | 0.661050037 | 0.896934512 | -6.926217197 |
| **P18085** | ARF4 | -0.016193905 | 3.180027492 | -0.813331016 | 0.444916223 | 0.791127304 | -6.926302179 |
| **Q99627-1** | COPS8 | 0.012340133 | 3.192159651 | 0.449422053 | 0.667790479 | 0.90001396 | -6.926336427 |
| **Q9NZ45** | CISD1 | 0.068227184 | 3.193411767 | 0.746662579 | 0.481505628 | 0.81235548 | -6.92741609 |
| **O95757** | HSPA4L | -0.026992567 | 3.208700413 | -1.209135414 | 0.268925917 | 0.671425933 | -6.927579636 |
| **Q14147** | DHX34 | -0.020189785 | 3.196723369 | -1.047000277 | 0.332621622 | 0.718705136 | -6.928028372 |
| **Q96MT8-3** | CEP63 | -0.008979892 | 3.179289089 | -0.26368143 | 0.800222814 | 0.952507937 | -6.928300006 |
| **P49257** | LMAN1 | -0.018270698 | 3.200108734 | -0.487270866 | 0.642135022 | 0.890981968 | -6.928337815 |
| **Q9Y248** | GINS2 | -0.014424894 | 3.190133896 | -0.479566217 | 0.647316155 | 0.89271426 | -6.928439145 |
| **P12814-1** | ACTN1 | 0.020907409 | 3.193538543 | 1.186762305 | 0.277054402 | 0.675421056 | -6.928618384 |

| | | | | | | | |
|---|---|---|---|---|---|---|---|
| **Q86W50-1** | METTL16 | -0.029127891 | 3.165056288 | -0.900486372 | 0.400130149 | 0.763361117 | -6.930175767 |
| **P42785-1** | PRCP | 0.005615188 | 3.18925277 | 0.158027587 | 0.879252078 | 0.97338977 | -6.930513605 |
| **O43414-1** | ERI3 | 0.013910014 | 3.151760112 | 0.730865635 | 0.490468291 | 0.818649279 | -6.930548831 |
| **P54727** | RAD23B | -0.038214661 | 3.151172331 | -0.720785261 | 0.496245638 | 0.822932915 | -6.931060022 |
| **Q5HYJ3-1** | FAM76B | 0.018840903 | 3.178693059 | 0.699212179 | 0.50876086 | 0.82913633 | -6.931268732 |
| **Q9NRW7** | VPS45 | -0.010538549 | 3.185867477 | -0.504921064 | 0.630347668 | 0.886812128 | -6.931466594 |
| **P84095** | RHOG | 0.038684104 | 3.171882041 | 0.806563756 | 0.44853855 | 0.793506743 | -6.932118513 |
| **Q99471-1** | PFDN5 | -0.032683541 | 3.174269014 | -1.016364946 | 0.345938913 | 0.727112344 | -6.932156897 |
| **P56556** | NDUFA6 | 0.07495164 | 3.189708583 | 0.762617653 | 0.472566596 | 0.808597546 | -6.932640553 |
| **O60504-1** | SORBS3 | -0.001461394 | 3.139857879 | -0.040859279 | 0.968639825 | 0.992306703 | -6.932670337 |
| **Q9NVU0** | POLR3E; LOC1010 | -0.01435487 | 3.197954902 | -0.915427584 | 0.392800539 | 0.760129045 | -6.932745645 |
| **Q49B96** | COX19 | -0.013253678 | 3.187249778 | -0.367193352 | 0.725186144 | 0.926648409 | -6.934111661 |
| **Q92820** | GGH | -0.022216236 | 3.187971985 | -0.958779356 | 0.372109067 | 0.745139764 | -6.934345171 |
| **O00499-2** | BIN1 | 0.003274322 | 3.198825211 | 0.118351142 | 0.909378509 | 0.981525914 | -6.934745116 |
| **P06127** | CD5 | -0.017742359 | 3.185537267 | -0.85539891 | 0.422866181 | 0.777884596 | -6.935176966 |
| **Q00341-1** | HDLBP | -0.035005294 | 3.186168261 | -1.58490322 | 0.160485134 | 0.560635782 | -6.936285779 |
| **Q9BTD8-1** | RBM42 | -0.00268909 | 3.177156493 | -0.080191097 | 0.938508307 | 0.987739899 | -6.937112379 |
| **Q9UNQ2** | DIMT1 | -0.027009076 | 3.171293749 | -0.928447311 | 0.386496514 | 0.7559613 | -6.93751054 |
| **P30044-1** | PRDX5 | 0.033782287 | 3.158202809 | 0.990946112 | 0.357306661 | 0.734600528 | -6.937661274 |
| **Q53RE8** | ANKRD39 | 0.002701287 | 3.223370722 | 0.057733547 | 0.955702719 | 0.989472196 | -6.937714806 |
| **P55036-1** | PSMD4 | -0.001715992 | 3.166504393 | -0.044432664 | 0.965899192 | 0.992001409 | -6.937958161 |
| **Q8NHQ9** | DDX55 | -0.022326857 | 3.174646003 | -0.897243728 | 0.401734322 | 0.765047743 | -6.938090233 |
| **Q6IBW4-4** | NCAPH2 | -0.002463468 | 3.17844648 | -0.06647648 | 0.949005166 | 0.989472196 | -6.938508716 |
| **Q6V1X1** | DPP8 | -0.017209264 | 3.177638552 | -0.536423622 | 0.609599469 | 0.876646363 | -6.938512809 |
| **Q719H9** | KCTD1 | 0.002954683 | 3.164520058 | 0.116001011 | 0.911168623 | 0.981900848 | -6.938530531 |
| **Q9NQZ2** | UTP3 | -0.024074654 | 3.186915302 | -0.567037184 | 0.589804474 | 0.868583051 | -6.939304018 |
| **Q96P16-1** | RPRD1A | 0.006702689 | 3.200574576 | 0.185064084 | 0.858843996 | 0.968855822 | -6.939629141 |
| **Q8N1G4** | LRRC47 | -0.021556115 | 3.179930659 | -1.319142244 | 0.231877138 | 0.639101249 | -6.939779042 |
| **Q13572-1** | ITPK1 | -0.017286503 | 3.212187168 | -0.507789624 | 0.628442853 | 0.885908173 | -6.940548218 |
| **Q9BQ15-1** | NABP2 | 0.006473288 | 3.191320132 | 0.272193807 | 0.793951222 | 0.949562315 | -6.940782456 |
| **P25440-2** | BRD2 | -0.021859572 | 3.190048708 | -1.122660082 | 0.301500134 | 0.691344519 | -6.940797702 |
| **Q7KZI7-14** | MARK2 | -0.014007258 | 3.177887059 | -0.87218638 | 0.414292364 | 0.771578253 | -6.940906552 |
| **Q8NEN9** | PDZD8 | -0.013626657 | 3.186452857 | -0.600593346 | 0.568537169 | 0.854816036 | -6.940950649 |
| **Q01804-1** | OTUD4 | 0.015020713 | 3.201220637 | 0.912703397 | 0.394129336 | 0.760194378 | -6.941137131 |
| **Q5JSH3** | WDR44 | -0.027351205 | 3.181025337 | -1.131037007 | 0.298206769 | 0.689670498 | -6.941657932 |
| **Q12789-2** | GTF3C1 | -0.033653965 | 3.180849472 | -1.448537281 | 0.194128554 | 0.599436985 | -6.941947767 |
| **Q5JSL3** | DOCK11 | 0.020458029 | 3.174627821 | 1.266755001 | 0.248926228 | 0.654137693 | -6.941974898 |
| **Q92918** | MAP4K1 | -0.0159824 | 3.202712785 | -0.914423818 | 0.393289759 | 0.760194378 | -6.942046339 |
| **P18206** | VCL | -0.012942027 | 3.178365041 | -1.271130642 | 0.247461443 | 0.652280071 | -6.942123582 |

| | | | | | | | |
|---|---|---|---|---|---|---|---|
| **O00203-1** | AP3B1 | -0.030503347 | 3.190509604 | -1.316573561 | 0.232688543 | 0.639692206 | -6.942208746 |
| **Q92542-1** | NCSTN | 0.000548178 | 3.189359996 | 0.014807529 | 0.988631805 | 0.996184213 | -6.944397516 |
| **Q5TCQ9-4** | MAGI3 | -0.012408011 | 3.212180901 | -0.458201277 | 0.661794559 | 0.897032706 | -6.944399344 |
| **Q12874** | SF3A3 | 0.018565643 | 3.175234225 | 1.256267532 | 0.252467702 | 0.656572825 | -6.945353241 |
| **Q9H8G2-1** | CAAP1 | 0.035763494 | 3.19677189 | 0.879146866 | 0.4107752 | 0.769906516 | -6.945549215 |
| **Q9H8W4** | PLEKHF2 | 0.022415656 | 3.190059511 | 0.576213552 | 0.583943367 | 0.8653224 | -6.945603721 |
| **O75608** | LYPLA1 | 0.007056863 | 3.170348494 | 0.205220652 | 0.843704307 | 0.962543524 | -6.945724053 |
| **Q5VVJ2** | MYSM1 | -0.002785331 | 3.171844029 | -0.107837419 | 0.917391195 | 0.983235402 | -6.945768834 |
| **Q9NRR4-1** | DROSHA | 0.017550659 | 3.163377965 | 0.612509056 | 0.56109606 | 0.852576894 | -6.945911257 |
| **Q5VTR2** | RNF20 | -0.021747453 | 3.180092683 | -1.403414088 | 0.206612923 | 0.612058044 | -6.946612096 |
| **P30043** | BLVRB | 0.011885429 | 3.179366331 | 0.392725414 | 0.70713492 | 0.91801515 | -6.946640741 |
| **Q6UW02** | CYP20A1 | -0.027990432 | 3.203475404 | -0.90436888 | 0.39821574 | 0.762880385 | -6.946833558 |
| **O95568** | METTL18 | 0.007886324 | 3.197016511 | 0.397536165 | 0.703756118 | 0.915921429 | -6.947096413 |
| **Q96PC5** | CTAGE5; MIA2 | 0.040662518 | 3.181604889 | 1.035790211 | 0.33744629 | 0.721283501 | -6.94713852 |
| **Q16775-1** | HAGH | 0.026318729 | 3.170467624 | 0.748405318 | 0.480523684 | 0.81235548 | -6.947692478 |
| **P16402** | HIST1H1D | 0.00680229 | 3.18714656 | 0.163389941 | 0.875195871 | 0.972759481 | -6.94822217 |
| **Q9GZT8-1** | NIF3L1 | 0.020423951 | 3.189216636 | 0.555023756 | 0.597528463 | 0.871651708 | -6.948242509 |
| **P53384** | NUBP1 | -0.000113092 | 3.18921191 | -0.002626252 | 0.997983663 | 0.998486363 | -6.948639396 |
| **Q9NVX0** | HAUS2 | -0.013827186 | 3.190561816 | -0.753979698 | 0.477391943 | 0.810964076 | -6.94866419 |
| **P04899-6** | GNAI2 | -0.034677456 | 3.1778023 | -0.842796325 | 0.429387216 | 0.782449869 | -6.94904006 |
| **Q96GS4** | C17orf59; BORCS6 | 0.002601398 | 3.166320124 | 0.099566605 | 0.923701935 | 0.985083485 | -6.949582365 |
| **P49407-1** | ARRB1 | 0.004836532 | 3.170010113 | 0.236110542 | 0.82064409 | 0.958568719 | -6.949736538 |
| **P57678** | GEMIN4 | -0.023544403 | 3.190241013 | -1.271927322 | 0.247195555 | 0.652280071 | -6.949947133 |
| **Q9Y5A7** | NUB1 | -0.020805132 | 3.180125753 | -1.123758886 | 0.301066432 | 0.691344519 | -6.950737626 |
| **Q9UGK8** | SERGEF | 0.00302654 | 3.187712095 | 0.05658579 | 0.956582264 | 0.989472196 | -6.95129403 |
| **Q8NEZ3-1** | WDR19 | 0.007935573 | 3.16725758 | 0.149423543 | 0.885768582 | 0.975708125 | -6.951909697 |
| **Q03701** | CEBPZ | -0.020350931 | 3.17833932 | -1.425388361 | 0.200445078 | 0.606070006 | -6.952056599 |
| **Q96EV8-1** | DTNBP1 | -0.009703719 | 3.204904641 | -0.361094929 | 0.729526614 | 0.927968131 | -6.952073879 |
| **P09382** | LGALS1 | 0.015072345 | 3.199719925 | 0.294206386 | 0.777811131 | 0.944053983 | -6.952652969 |
| **Q32P44** | EML3 | 0.010570554 | 3.152681736 | 0.213322409 | 0.837638861 | 0.96111408 | -6.952751192 |
| **P40855-1** | PEX19 | -0.033756993 | 3.196469535 | -0.791937089 | 0.456438834 | 0.798240489 | -6.953025034 |
| **Q96NW4** | ANKRD27 | 0.010323262 | 3.188443175 | 0.377846203 | 0.717630547 | 0.924679646 | -6.953257482 |
| **Q9UKY1** | ZHX1 | 0.017571667 | 3.197365705 | 0.764111949 | 0.471735253 | 0.807565951 | -6.953589851 |
| **Q96T37-1** | RBM15 | -0.033741857 | 3.184890622 | -1.133744065 | 0.297148903 | 0.689298697 | -6.955273497 |
| **Q9ULC4-3** | MCTS1 | -0.025429343 | 3.193636695 | -0.895818847 | 0.402440744 | 0.765842327 | -6.95543204 |
| **Q15019-2** | 2-Sep | -0.032098029 | 3.168385508 | -1.264002213 | 0.249851607 | 0.654571265 | -6.955759176 |
| **Q5VWG9** | TAF3 | -0.016006822 | 3.198023912 | -0.455010193 | 0.66397087 | 0.898220426 | -6.955868395 |
| **O15297** | PPM1D | -0.008378321 | 3.193554006 | -0.362793178 | 0.728316814 | 0.927171461 | -6.956038304 |
| **Q8N3U4-1** | STAG2 | -0.022601974 | 3.164540405 | -1.371741136 | 0.215804167 | 0.62144404 | -6.95623217 |

| | | | | | | | |
|---|---|---|---|---|---|---|---|
| **Q8IWE5** | PLEKHM2 | -0.011892159 | 3.183090353 | -0.628165976 | 0.551408468 | 0.847559559 | -6.956335029 |
| **Q9UHA4** | LAMTOR3 | -0.027425577 | 3.164985237 | -0.724180251 | 0.494294834 | 0.821686278 | -6.956540169 |
| **Q9P2D0** | IBTK | 0.013690826 | 3.205584069 | 0.923006698 | 0.389121414 | 0.75855365 | -6.956856877 |
| **Q49AR2-1** | C5orf22 | -0.015969394 | 3.182441118 | -0.467879466 | 0.655215874 | 0.896497734 | -6.957280866 |
| **P50747** | HLCS | 0.008387555 | 3.183536718 | 0.282701806 | 0.786232228 | 0.946749713 | -6.957922332 |
| **P53609** | PGGT1B | 0.011646831 | 3.197840991 | 0.47610803 | 0.649648645 | 0.893646754 | -6.959497345 |
| **A0JNW5-1** | UHRF1BP1L | 0.004516428 | 3.17109057 | 0.115789904 | 0.911329452 | 0.981900848 | -6.959504607 |
| **Q7Z7H5** | TMED4 | 0.013051061 | 3.177904618 | 0.310579592 | 0.765882461 | 0.940484723 | -6.959703587 |
| **O14950** | MYL12B; MYL12A | 0.002150211 | 3.18268196 | 0.074935207 | 0.942529597 | 0.988606569 | -6.960147561 |
| **O60568** | PLOD3 | -0.003549789 | 3.211821316 | -0.094246689 | 0.927764299 | 0.9859101 | -6.960398144 |
| **Q8WWQ0** | PHIP | -0.013105187 | 3.18277517 | -1.32462633 | 0.230153156 | 0.637575603 | -6.960456971 |
| **Q13415** | ORC1 | -0.020972535 | 3.2260108 | -0.790114864 | 0.457429865 | 0.798918505 | -6.960773375 |
| **Q9HBM1** | SPC25 | -0.011018475 | 3.159996088 | -0.407777667 | 0.696587375 | 0.912062672 | -6.961614166 |
| **O43303** | CCP110 | 0.016171442 | 3.192511035 | 0.992345364 | 0.356673348 | 0.734600528 | -6.962094395 |
| **Q8NBJ7-1** | SUMF2 | -0.013320851 | 3.156838191 | -0.522576574 | 0.618672921 | 0.882227341 | -6.962433814 |
| **Q9ULC3** | RAB23 | 0.006906185 | 3.167695784 | 0.216511759 | 0.835254397 | 0.960912428 | -6.962462905 |
| **Q15006** | EMC2 | 0.001102447 | 3.186063348 | 0.040616471 | 0.968826066 | 0.992306703 | -6.962876555 |
| **Q7Z2W4** | ZC3HAV1 | 0.024130293 | 3.181301212 | 1.146370156 | 0.292256054 | 0.685264687 | -6.962933326 |
| **Q12933-2** | TRAF2 | -0.038827172 | 3.178116985 | -0.8208337 | 0.440924585 | 0.788971713 | -6.963023654 |
| **Q9NX08** | COMMD8 | -0.003748723 | 3.168556042 | -0.135099691 | 0.896638269 | 0.978757548 | -6.963073372 |
| **Q15542** | TAF5 | 0.019139925 | 3.184125783 | 0.820123585 | 0.441301289 | 0.788971713 | -6.96344104 |
| **Q99081-3** | TCF12 | -0.019839752 | 3.189230572 | -0.661286105 | 0.531256841 | 0.839849063 | -6.963597048 |
| **Q86X53** | ERICH1 | 0.003474339 | 3.211303002 | 0.083965203 | 0.93562193 | 0.986793807 | -6.963830481 |
| **Q14240-2** | EIF4A2 | -0.036903199 | 3.208867125 | -1.034742281 | 0.337900154 | 0.721865213 | -6.964224941 |
| **Q96S52** | PIGS | 0.014302345 | 3.175869245 | 0.491972592 | 0.638983866 | 0.88909352 | -6.964379248 |
| **O14920-1** | IKBKB | -0.012587711 | 3.17071579 | -0.801835475 | 0.451081814 | 0.79513757 | -6.964451772 |
| **Q96A57-2** | TMEM230 | -0.01596514 | 3.188035707 | -0.24222978 | 0.816097915 | 0.956348067 | -6.964463664 |
| **Q9UKI8-2** | TLK1 | 0.005883013 | 3.18492454 | 0.227288393 | 0.82721154 | 0.959077751 | -6.964790632 |
| **Q9BSC4-1** | NOL10 | 0.012568799 | 3.197939552 | 1.018180758 | 0.345137922 | 0.726061285 | -6.964984914 |
| **Q14919-2** | DRAP1 | -0.004438308 | 3.178743875 | -0.13073627 | 0.899954394 | 0.978800501 | -6.965914304 |
| **Q9NVH2** | INTS7 | 0.024634537 | 3.195461652 | 0.832615334 | 0.434708143 | 0.784298014 | -6.965960498 |
| **P56962** | STX17 | -0.006963542 | 3.181786835 | -0.248483258 | 0.81145997 | 0.955606306 | -6.966260035 |
| **Q15653-1** | NFKBIB | 0.001508268 | 3.198857908 | 0.059218323 | 0.95456501 | 0.989472196 | -6.966549874 |
| **Q96JH7** | VCPIP1 | 0.018809761 | 3.177050403 | 1.438779292 | 0.196768756 | 0.603701501 | -6.967356133 |
| **P25205** | MCM3 | 0.013408717 | 3.178183246 | 1.628847069 | 0.150859027 | 0.548546897 | -6.967540989 |
| **O95630** | STAMBP | -0.014950545 | 3.160506247 | -0.878828921 | 0.410935377 | 0.769906516 | -6.96855494 |
| **P36405** | ARL 3.00 | -0.006482937 | 3.169391219 | -0.332159456 | 0.750266092 | 0.935635612 | -6.968740616 |
| **Q96CS3** | FAF2 | 0.021278832 | 3.192719335 | 0.61016901 | 0.562552729 | 0.852576894 | -6.969057575 |
| **Q9UBP0-1** | SPAST | 0.012826941 | 3.15662962 | 0.493992655 | 0.637632479 | 0.889065271 | -6.969114865 |

| | | | | | | | |
|---|---|---|---|---|---|---|---|
| **O95376** | ARIH2 | 0.020575528 | 3.18494773 | 0.85787293 | 0.421594546 | 0.777315429 | -6.969413719 |
| **Q9HCI7** | MSL2 | 0.018879309 | 3.20219813 | 0.703459715 | 0.506280514 | 0.82913633 | -6.970044201 |
| **Q9UKT5** | FBXO4 | -0.001272699 | 3.182560021 | -0.039026077 | 0.970045995 | 0.992521442 | -6.970117092 |
| **Q6NUQ4** | TMEM214 | 0.014899905 | 3.194431292 | 0.396598577 | 0.704414059 | 0.916268778 | -6.970419121 |
| **Q969X6** | CIRH1A; UTP4 | -0.028053529 | 3.183872004 | -1.089451783 | 0.314852882 | 0.701837878 | -6.971032475 |
| **Q9HD42-1** | CHMP1A | 0.010553489 | 3.183818286 | 0.342577755 | 0.742771709 | 0.933447653 | -6.97112526 |
| **Q96IU2** | ZBED3 | 0.002789085 | 3.174076772 | 0.122927762 | 0.905894116 | 0.98075504 | -6.971547728 |
| **Q9H6R0-1** | DHX33 | -0.02118282 | 3.200307057 | -1.000064782 | 0.353195296 | 0.732738778 | -6.97174658 |
| **Q9Y4K3** | TRAF6 | -0.008226578 | 3.198377687 | -0.378710989 | 0.717018687 | 0.924679646 | -6.971812862 |
| **P49914** | MTHFS | 0.007807169 | 3.173708425 | 0.405237787 | 0.698362105 | 0.913150177 | -6.97184955 |
| **O95777** | NAA38; LSM8 | 0.015600532 | 3.181584461 | 0.548951068 | 0.601454595 | 0.872497827 | -6.972133161 |
| **Q96HN2** | AHCYL2 | 0.011720384 | 3.190616917 | 0.263643857 | 0.800250532 | 0.952507937 | -6.973251286 |
| **Q9H0G5** | NSRP1 | -0.021793299 | 3.191388018 | -0.875476857 | 0.412626913 | 0.770644294 | -6.973452305 |
| **P20020** | ATP2B1 | 0.012570457 | 3.143254743 | 0.314550069 | 0.763000024 | 0.939644172 | -6.973504867 |
| **Q8IZ21-2** | PHACTR4 | 0.028472223 | 3.169996442 | 0.766859679 | 0.470209191 | 0.806348374 | -6.973505253 |
| **O00629** | KPNA4 | 0.008398389 | 3.16470868 | 0.337249356 | 0.746600949 | 0.934974996 | -6.973703334 |
| **Q92599-1** | 8-Sep | 0.009692562 | 3.190893596 | 0.293572484 | 0.778274299 | 0.944053983 | -6.974329928 |
| **Q9BRS2** | RIOK1 | -0.041700012 | 3.200591295 | -0.872559513 | 0.414103256 | 0.771567763 | -6.974514707 |
| **Q99633-1** | PRPF18 | -0.031691227 | 3.183029752 | -0.749544659 | 0.479882459 | 0.81235548 | -6.974729388 |
| **A6NIH7** | UNC119B | -0.01059754 | 3.161567552 | -0.517283936 | 0.622160344 | 0.883385606 | -6.975296069 |
| **Q5SXM2** | SNAPC4 | -0.008026379 | 3.202628672 | -0.277657609 | 0.789934402 | 0.947865984 | -6.975382641 |
| **Q15545** | TAF7 | -0.011185875 | 3.1831278 | -0.616380165 | 0.558691311 | 0.851363273 | -6.97543341 |
| **P09874** | PARP1 | 0.048829959 | 3.174724177 | 1.789655746 | 0.120111391 | 0.499295152 | -6.975501557 |
| **Q9Y3X0** | CCDC9 | 0.001924184 | 3.178627427 | 0.044336948 | 0.965972595 | 0.992001409 | -6.976229155 |
| **Q6JBY9** | RCSD1 | -0.119638391 | 3.165986124 | -0.978560955 | 0.362950723 | 0.738259983 | -6.977625085 |
| **Q9BZE9-2** | ASPSCR1 | 0.019222841 | 3.172680749 | 0.624684227 | 0.553553897 | 0.848478354 | -6.978712338 |
| **Q8WY91** | THAP4 | -0.000499106 | 3.200826442 | -0.022530935 | 0.982703268 | 0.994932161 | -6.978905394 |
| **Q8WVQ1** | CANT1 | -0.019453972 | 3.206776202 | -0.664196112 | 0.529508677 | 0.839849063 | -6.978909107 |
| **Q9UQ13** | SHOC2 | -0.011215763 | 3.181651242 | -0.608841003 | 0.563380419 | 0.852578515 | -6.979683327 |
| **Q8N5C7-1** | DTWD1 | 0.009021912 | 3.193257779 | 0.199682491 | 0.847857185 | 0.964108392 | -6.979901434 |
| **Q5TAQ9-1** | DCAF8 | -0.013813237 | 3.169318583 | -0.546432978 | 0.603086845 | 0.872497827 | -6.979988382 |
| **Q9NVU7** | SDAD1 | -0.027344918 | 3.178739821 | -0.909477264 | 0.395707347 | 0.761602439 | -6.980569608 |
| **Q13795** | ARFRP1 | 0.001011434 | 3.191783628 | 0.047449983 | 0.963585427 | 0.991283985 | -6.980682124 |
| **Q08AF3-1** | SLFN5 | 0.016131362 | 3.165438855 | 1.103715653 | 0.309059151 | 0.696978291 | -6.98084386 |
| **Q14126** | DSG2 | -0.004812511 | 3.176786613 | -0.210731766 | 0.839577081 | 0.961392936 | -6.981315227 |
| **Q9P107-2** | GMIP | -0.011161233 | 3.182118489 | -0.461319345 | 0.659671478 | 0.896852471 | -6.981522454 |
| **Q9BV57** | ADI1 | 0.022423265 | 3.175538697 | 0.707444176 | 0.503961004 | 0.828293376 | -6.982798072 |
| **P39687** | ANP32A | 0.02464238 | 3.172819811 | 1.257164684 | 0.252163046 | 0.656218606 | -6.983424165 |
| **Q9Y3E5** | PTRH2 | 0.000765259 | 3.200261188 | 0.016269495 | 0.987509517 | 0.99582925 | -6.98353166 |

| | | | | | | | |
|---|---|---|---|---|---|---|---|
| **Q9H9F9** | ACTR5 | 0.010842909 | 3.194108839 | 0.491809088 | 0.639093313 | 0.88909352 | -6.983532323 |
| **Q08722** | CD47 | -0.050551044 | 3.163626688 | -0.671730792 | 0.524999245 | 0.837910607 | -6.983542546 |
| **O94761** | RECQL4 | -0.005365803 | 3.167142258 | -0.245738155 | 0.813494896 | 0.95620584 | -6.9835683 |
| **Q7L1T6** | CYB5R4 | 0.015104122 | 3.180063182 | 0.617116788 | 0.558234425 | 0.851117349 | -6.985128652 |
| **Q3B726** | TWISTNB | 0.054677839 | 3.235137963 | 0.425902064 | 0.683983637 | 0.905978473 | -6.985184837 |
| **Q6UW63** | KDELC1 | -0.01019537 | 3.183708374 | -0.593048148 | 0.573279254 | 0.85840627 | -6.98547465 |
| **P11441** | UBL4A | 0.011953567 | 3.202680718 | 0.366076567 | 0.725980184 | 0.926870088 | -6.985486228 |
| **Q9UHI6** | DDX20 | -0.020490613 | 3.193158733 | -1.223908711 | 0.263670813 | 0.666941931 | -6.985938912 |
| **Q7Z739** | YTHDF3 | -0.02076744 | 3.202315001 | -0.674491716 | 0.523353004 | 0.836627689 | -6.986124367 |
| **Q15813-2** | TBCE | -0.017798136 | 3.181853405 | -1.045227833 | 0.333380754 | 0.719098069 | -6.986402979 |
| **P23443-1** | RPS6KB1 | -0.009865464 | 3.170798845 | -0.289531008 | 0.781229557 | 0.944918194 | -6.986533574 |
| **Q86T24** | ZBTB33 | 0.005057722 | 3.194752465 | 0.239182647 | 0.818360774 | 0.957415004 | -6.987248983 |
| **P78356** | PIP4K2B | 0.006564266 | 3.166522757 | 0.24662604 | 0.81283654 | 0.956167049 | -6.987601205 |
| **Q9H501** | ESF1 | -0.028809994 | 3.198138241 | -0.959749249 | 0.371655905 | 0.744717822 | -6.987611704 |
| **Q9BSU1** | C16orf70 | -0.002163362 | 3.206161617 | -0.064002649 | 0.950899823 | 0.989472196 | -6.987795799 |
| **O00178** | GTPBP1 | -0.025237708 | 3.187240771 | -1.038869172 | 0.336115596 | 0.720858943 | -6.988595317 |
| **Q99547** | MPHOSPH6 | -0.021358017 | 3.20366669 | -0.347810658 | 0.739018803 | 0.931081875 | -6.988656587 |
| **Q9BXP5-3** | SRRT | 0.045599969 | 3.191595079 | 1.293715448 | 0.240019839 | 0.645958461 | -6.988724828 |
| **A4D1P6-1** | WDR91 | 0.013712238 | 3.181669766 | 0.612084639 | 0.561360089 | 0.852576894 | -6.988746098 |
| **Q92522** | H1FX | -0.024362502 | 3.145074072 | -0.493287672 | 0.63810393 | 0.889065271 | -6.98880715 |
| **Q5EBM0-1** | CMPK2 | 0.027575854 | 3.191932249 | 0.334356007 | 0.748683527 | 0.935412899 | -6.989117937 |
| **Q92643** | PIGK | -0.007156386 | 3.202132409 | -0.122507131 | 0.906214269 | 0.98075504 | -6.989674714 |
| **O94830** | DDHD2 | -0.012271225 | 3.179436104 | -0.738786909 | 0.485960102 | 0.81473183 | -6.991087995 |
| **Q16512-2** | PKN1 | -0.020068251 | 3.168909148 | -0.92847606 | 0.386482679 | 0.7559613 | -6.991413723 |
| **Q9H6A0-1** | DENND2D | -0.010352199 | 3.185314769 | -0.403336607 | 0.699691893 | 0.913245694 | -6.992129947 |
| **Q3LXA3** | DAK; TKFC | -0.01515873 | 3.187919565 | -0.94891934 | 0.376740152 | 0.749049177 | -6.992631897 |
| **O75943-1** | RAD17 | -0.001943288 | 3.19348418 | -0.079356958 | 0.939146381 | 0.988002681 | -6.993201328 |
| **P29558-1** | RBMS1 | -0.013247551 | 3.182386954 | -0.580582634 | 0.581164664 | 0.863864033 | -6.99368567 |
| **Q8NB49-1** | ATP11C | 0.009866045 | 3.154836223 | 0.609973509 | 0.56267453 | 0.852576894 | -6.993899426 |
| **Q7LBR1** | CHMP1B | 0.008346496 | 3.167894829 | 0.229692784 | 0.825420126 | 0.959077751 | -6.994373017 |
| **O95361-1** | TRIM16 | -0.011138779 | 3.191170037 | -0.319535201 | 0.759386778 | 0.938422437 | -6.994919175 |
| **Q9H8E8-1** | PET117; CSRP2BP | 0.009409038 | 3.181943587 | 0.381900692 | 0.714763841 | 0.9235376 | -6.995124352 |
| **Q9H2H8-1** | PPIL3 | 0.018863704 | 3.172539671 | 0.854534939 | 0.423310916 | 0.777884596 | -6.995424183 |
| **Q9UI10-1** | EIF2B4 | 0.015917826 | 3.181769836 | 0.782985586 | 0.461321619 | 0.802540018 | -6.995554421 |
| **Q6IPU0** | CENPP | 0.001340772 | 3.178749063 | 0.062363175 | 0.952155654 | 0.989472196 | -6.996626182 |
| **P14316-1** | IRF2 | 0.023341164 | 3.165404124 | 0.642118962 | 0.542862037 | 0.843669625 | -6.99768242 |
| **Q06265-2** | EXOSC9 | -0.032187882 | 3.187791089 | -0.627558761 | 0.551782263 | 0.847634615 | -6.997905835 |
| **Q5JSP0-1** | FGD3 | 0.015423462 | 3.157586351 | 1.07241955 | 0.321887054 | 0.709476351 | -6.998051579 |
| **O60927** | PPP1R11 | 0.001656597 | 3.191953873 | 0.062609681 | 0.951966821 | 0.989472196 | -6.998102963 |

| | | | | | | | |
|---|---|---|---|---|---|---|---|
| **Q16513** | PKN2 | -0.0150567 | 3.191794272 | -0.569102162 | 0.588482593 | 0.867233795 | -6.998448311 |
| **Q96P11-2** | NSUN5 | -0.020559831 | 3.189524969 | -0.892289063 | 0.40419472 | 0.766336431 | -6.998683826 |
| **Q16186** | ADRM1 | -0.010961724 | 3.189831751 | -0.250900343 | 0.809669514 | 0.955498404 | -6.99890045 |
| **Q9UJ41-1** | RABGEF1 | -0.031147891 | 3.208906939 | -0.88903299 | 0.405817731 | 0.766336431 | -6.999436477 |
| **P10768** | ESD | -0.019065634 | 3.174526262 | -1.322876534 | 0.23070199 | 0.63865063 | -6.999687086 |
| **Q15811** | ITSN1 | 0.008739728 | 3.165714474 | 0.276417428 | 0.79084554 | 0.948368382 | -6.999737795 |
| **Q86VX2-1** | COMMD7 | 0.001053638 | 3.182915443 | 0.036110544 | 0.97228261 | 0.992928787 | -6.99981524 |
| **P17029** | ZKSCAN1 | -0.028064629 | 3.195399155 | -0.717635915 | 0.498059859 | 0.824674598 | -7.001229962 |
| **Q07065** | CKAP4 | 0.014256461 | 3.1554433 | 0.295731269 | 0.776697358 | 0.94357883 | -7.001792119 |
| **Q16537** | PPP2R5E | 0.010717011 | 3.14953466 | 0.386413112 | 0.711579246 | 0.921215107 | -7.001843793 |
| **Q9GZR2-1** | REXO4 | -0.027748646 | 3.183919009 | -0.857513868 | 0.421778929 | 0.777315429 | -7.002274411 |
| **Q15631-1** | TSN | 0.016940457 | 3.172246143 | 0.27807327 | 0.789629104 | 0.947865984 | -7.002321505 |
| **Q86SX6** | GLRX5 | -0.01977926 | 3.202396047 | -0.10649233 | 0.918417087 | 0.983742622 | -7.003549552 |
| **Q8TAT6-2** | NPLOC4 | 0.040227817 | 3.183550034 | 1.13075351 | 0.298317735 | 0.689670498 | -7.003769045 |
| **O00443** | PIK3C2A | -0.022426196 | 3.204056321 | -0.83492601 | 0.43349636 | 0.783962934 | -7.004061242 |
| **Q96RG2-2** | PASK | -0.021164185 | 3.178802651 | -0.672933142 | 0.52428192 | 0.837438652 | -7.005344061 |
| **P52298-1** | NCBP2 | 0.002825582 | 3.188312657 | 0.032693813 | 0.974904056 | 0.99316605 | -7.005957041 |
| **Q13610** | PWP1 | 0.009195489 | 3.181104351 | 0.466497272 | 0.656153392 | 0.896497734 | -7.005979255 |
| **Q9UHP3** | USP25 | -0.018547957 | 3.177251257 | -0.894347025 | 0.403171412 | 0.766336431 | -7.007167222 |
| **O95721** | SNAP29 | -0.021450654 | 3.215833113 | -0.792931538 | 0.45589863 | 0.798084321 | -7.007681469 |
| **Q6P2Q9** | PRPF8 | -0.034649325 | 3.175722456 | -1.813360323 | 0.116124566 | 0.492573307 | -7.007859359 |
| **O75044** | SRGAP2 | -0.003990777 | 3.176902826 | -0.1520924 | 0.883746195 | 0.975295424 | -7.007923536 |
| **Q9NY61** | AATF | -0.014579168 | 3.192649066 | -0.931128153 | 0.385208059 | 0.754741808 | -7.008102477 |
| **P49902** | NT5C2 | 0.070272846 | 3.191359212 | 0.800542204 | 0.451779211 | 0.79513757 | -7.008133265 |
| **Q9UFC0** | LRWD1 | 0.015082377 | 3.196696169 | 0.883937159 | 0.408367499 | 0.768469868 | -7.008197563 |
| **P60174** | TPI1 | 0.018607599 | 3.188765402 | 1.842819768 | 0.111349279 | 0.483014501 | -7.008443822 |
| **O15020** | SPTBN2 | -0.014262172 | 3.17355855 | -1.242030244 | 0.257345224 | 0.660189799 | -7.008558248 |
| **Q8NB14** | USP38 | 0.008699049 | 3.199641481 | 0.328342674 | 0.75301911 | 0.93582619 | -7.008778075 |
| **Q13535** | ATR | -0.016682868 | 3.186254547 | -1.055728437 | 0.328903632 | 0.714558204 | -7.00898855 |
| **Q9H0V9-2** | LMAN2L | -0.007120915 | 3.190027726 | -0.226566783 | 0.827749402 | 0.959077751 | -7.009049438 |
| **O43488** | AKR7A2 | 0.007018893 | 3.18955273 | 0.082351081 | 0.936856259 | 0.987232823 | -7.009052023 |
| **Q8IYI6** | EXOC8 | -0.021680593 | 3.187562397 | -0.926027008 | 0.387662558 | 0.757123654 | -7.009152295 |
| **P51948-1** | MNAT1 | 0.004346948 | 3.182649596 | 0.095598099 | 0.926732115 | 0.985528933 | -7.009309958 |
| **Q9UK99** | FBXO3 | -0.00749237 | 3.180409639 | -0.423641061 | 0.685550074 | 0.906272103 | -7.009431588 |
| **Q96IY1** | NSL1 | -0.014782412 | 3.170786695 | -0.680738943 | 0.519640205 | 0.834217303 | -7.009510474 |
| **O95453** | PARN | 0.033524109 | 3.176246775 | 1.185448254 | 0.277538233 | 0.675997904 | -7.009573271 |
| **Q9H9Y6-3** | POLR1B | 0.013452267 | 3.196580899 | 0.915727246 | 0.392654577 | 0.760129045 | -7.009716468 |
| **Q15208** | STK38 | 0.011418597 | 3.143419962 | 0.774685749 | 0.465881306 | 0.80378436 | -7.009958356 |
| **Q9P2I0** | CPSF2 | -0.024246007 | 3.194965524 | -1.264436826 | 0.249705309 | 0.65453932 | -7.010433102 |

| | | | | | | | |
|---|---|---|---|---|---|---|---|
| **P10746** | UROS | 0.0122096 | 3.176738037 | 0.364624652 | 0.727013051 | 0.92710315 | -7.0124282 |
| **Q8WUH6** | C12orf23; TMEM26 | 0.029800946 | 3.12087918 | 0.716157453 | 0.498913061 | 0.824970816 | -7.012804359 |
| **P53985** | SLC16A1 | 0.012283952 | 3.212673487 | 0.164921535 | 0.874038082 | 0.972445395 | -7.013019722 |
| **O75151** | PHF2 | -0.0095018 | 3.18753482 | -0.49336768 | 0.638050416 | 0.889065271 | -7.013075916 |
| **O14656** | TOR1A | -0.025595111 | 3.177911318 | -0.378391057 | 0.717245021 | 0.924679646 | -7.01327109 |
| **Q9Y315** | DERA | -0.012849155 | 3.172415153 | -0.48784769 | 0.641747993 | 0.890911725 | -7.013982184 |
| **P33527** | ABCC1 | -0.018737527 | 3.163479505 | -1.000425443 | 0.353033452 | 0.732738778 | -7.014383769 |
| **Q9ULJ7-1** | ANKRD50 | -0.020667037 | 3.182426726 | -0.760841615 | 0.473555991 | 0.808788313 | -7.014406269 |
| **Q9BZV1** | UBXN6 | -0.002122477 | 3.155703328 | -0.045497632 | 0.965082502 | 0.992001409 | -7.014440447 |
| **Q9ULP9** | TBC1D24 | 0.0189563 | 3.188755933 | 0.45524568 | 0.663810147 | 0.898182275 | -7.014738638 |
| **Q5VW36** | FOCAD | -0.021835183 | 3.192017132 | -0.941850407 | 0.380087465 | 0.751005945 | -7.014806504 |
| **Q9UHG3** | PCYOX1 | -0.012513824 | 3.191513725 | -0.64369575 | 0.541901415 | 0.843570684 | -7.014911891 |
| **Q69YQ0-1** | SPECC1L | 0.000759176 | 3.191268149 | 0.022717625 | 0.982559976 | 0.994932161 | -7.015325275 |
| **Q5JPI3** | C3orf38 | -0.017796308 | 3.180414655 | -0.380450727 | 0.715788458 | 0.924254721 | -7.016423052 |
| **P47756-2** | CAPZB | 0.040160061 | 3.170780226 | 1.380165255 | 0.213324397 | 0.61901473 | -7.017914162 |
| **P18858** | LIG1 | 0.017799435 | 3.158827476 | 1.376184424 | 0.214493017 | 0.619913059 | -7.018458765 |
| **Q6NSI4-1** | CXorf57 | -0.002302916 | 3.15065719 | -0.084674619 | 0.935079496 | 0.986793807 | -7.01851024 |
| **Q969R5-1** | L3MBTL2 | -0.024294047 | 3.208558133 | -0.841342758 | 0.430144006 | 0.782603406 | -7.019363333 |
| **Q53FT3** | C11orf73; HIKESHI | 0.005483682 | 3.183073082 | 0.175557235 | 0.866007601 | 0.970432241 | -7.019442836 |
| **Q6IPR3-1** | TYW3 | -0.009205813 | 3.217099882 | -0.220148206 | 0.832537973 | 0.960582935 | -7.019795916 |
| **O75886** | STAM2 | 0.009874926 | 3.168567283 | 0.328668491 | 0.752783947 | 0.935782019 | -7.019952327 |
| **P0DJD1** | RGPD2 | 0.005071681 | 3.139279118 | 0.155034674 | 0.881517713 | 0.974438769 | -7.020283335 |
| **Q8NI27-1** | THOC2 | -0.023586629 | 3.186584869 | -1.464228694 | 0.189950641 | 0.593922803 | -7.020491938 |
| **P13637-3** | ATP1A3 | 0.002862208 | 3.206246757 | 0.10123224 | 0.922430536 | 0.984164603 | -7.020802551 |
| **Q8N5C6** | SRBD1 | -0.02454152 | 3.166225438 | -1.117386322 | 0.303588913 | 0.692399426 | -7.021035257 |
| **Q6ZMI0-1** | PPP1R21 | -0.016045896 | 3.195560631 | -0.879264972 | 0.410715712 | 0.769906516 | -7.021097934 |
| **Q15326** | ZMYND11 | 0.004910438 | 3.200208995 | 0.161199652 | 0.87685217 | 0.973040052 | -7.021435777 |
| **Q7Z569** | BRAP | -0.013236516 | 3.18867667 | -0.773363708 | 0.466610468 | 0.804518266 | -7.021849199 |
| **Q96RT8** | TUBGCP5 | 0.016172278 | 3.189601436 | 0.777401514 | 0.46438592 | 0.803074528 | -7.021856759 |
| **Q9Y4R8** | TELO2 | -0.034772328 | 3.197871995 | -1.167358797 | 0.284271683 | 0.680282687 | -7.022024749 |
| **Q99873-2** | PRMT1 | 0.026035304 | 3.17618632 | 1.272911723 | 0.246867361 | 0.652280071 | -7.022041441 |
| **P19387** | POLR2C | -0.023225451 | 3.196274365 | -0.668490317 | 0.526935633 | 0.839483378 | -7.022064678 |
| **Q13868** | EXOSC2 | -0.013029549 | 3.186250048 | -0.356619761 | 0.732718652 | 0.928164811 | -7.022249634 |
| **O75191** | XYLB | -0.013859161 | 3.182438374 | -0.564514739 | 0.591421512 | 0.868764508 | -7.022389917 |
| **Q9BYJ9-1** | YTHDF1 | -0.01059384 | 3.19893105 | -0.636919487 | 0.546037175 | 0.845412981 | -7.022401738 |
| **O75976** | CPD | 0.023092306 | 3.185819765 | 1.16298997 | 0.285918378 | 0.681580409 | -7.022808335 |
| **O43399-7** | TPD52L2 | -0.030045347 | 3.200645925 | -0.787922571 | 0.458624152 | 0.799639769 | -7.023183921 |
| **Q13206** | DDX10 | -0.031746371 | 3.16682996 | -1.023914746 | 0.342618207 | 0.724485042 | -7.023216007 |
| **P35813-3** | PPM1A | 0.012808773 | 3.165195458 | 1.062876891 | 0.325883585 | 0.710703247 | -7.023701515 |

| | | | | | | | |
|---|---|---|---|---|---|---|---|
| **Q9NXC5-1** | MIOS | 0.010906364 | 3.176138701 | 0.632624143 | 0.548668826 | 0.846404495 | -7.02371568 |
| **O43813** | LANCL1 | 0.050318029 | 3.145133291 | 0.792252526 | 0.456267434 | 0.798116417 | -7.023810175 |
| **Q16254** | E2F4 | -0.029885799 | 3.177299806 | -0.435180065 | 0.677573581 | 0.903662615 | -7.023913153 |
| **Q8NHP8** | PLBD2 | 0.025144228 | 3.201283674 | 0.312146362 | 0.764744551 | 0.940369694 | -7.024097998 |
| **Q86YM7-1** | HOMER1 | 0.015897644 | 3.203805025 | 0.49189845 | 0.639033494 | 0.88909352 | -7.02467126 |
| **Q8NCD3-1** | HJURP | -0.003356687 | 3.14167623 | -0.128231266 | 0.901859143 | 0.97952714 | -7.024720138 |
| **Q9BTZ2** | DHRS4 | -0.003894177 | 3.173346926 | -0.17236506 | 0.86841609 | 0.970949315 | -7.024875366 |
| **Q7L7X3** | TAOK1 | -0.019835496 | 3.190802848 | -0.480039284 | 0.646997414 | 0.89271426 | -7.024878068 |
| **Q5VTL8** | PRPF38B | -0.011484805 | 3.190420543 | -0.666586171 | 0.528075596 | 0.839849063 | -7.025182299 |
| **Q58WW2-3** | DCAF6 | -0.00470988 | 3.198619984 | -0.181584987 | 0.861463926 | 0.968855822 | -7.025565457 |
| **Q6IQ49-1** | SDE2 | -0.020730817 | 3.195415552 | -0.572760997 | 0.586144608 | 0.8653224 | -7.02605048 |
| **P40425** | PBX2 | -0.010290189 | 3.186775042 | -0.512177254 | 0.625535287 | 0.885387512 | -7.026848458 |
| **P54578** | USP14 | 0.044751824 | 3.175435933 | 1.180593053 | 0.279332117 | 0.676725357 | -7.027458319 |
| **Q13569** | TDG | 0.006727932 | 3.200017169 | 0.229097509 | 0.825863536 | 0.959077751 | -7.029042656 |
| **O43292-1** | GPAA1 | 0.002239622 | 3.215884593 | 0.051623224 | 0.960385903 | 0.990943637 | -7.029774336 |
| **P78332** | RBM6 | -0.05002009 | 3.174143295 | -1.183903779 | 0.27810782 | 0.67612198 | -7.029980248 |
| **O75818-1** | RPP40 | 0.008860865 | 3.207502385 | 0.125341593 | 0.904057259 | 0.980210603 | -7.030086777 |
| **P38935** | IGHMBP2 | -0.005500385 | 3.191119106 | -0.101852756 | 0.921956951 | 0.984164603 | -7.030563735 |
| **Q7KZ85** | SUPT6H | -0.017505263 | 3.192105189 | -1.35435444 | 0.221003872 | 0.626484139 | -7.030919259 |
| **Q9C0D5** | TANC1 | 0.014029956 | 3.181404213 | 0.444969858 | 0.670841348 | 0.901175787 | -7.031159206 |
| **Q9HCI6** | KIAA1586 | 0.005259752 | 3.221254262 | 0.125541074 | 0.903905488 | 0.980210603 | -7.031513699 |
| **P63220** | RPS21 | -0.057236116 | 3.197311688 | -1.318053459 | 0.232220762 | 0.639291044 | -7.031797039 |
| **Q8TEA7** | TBCK | -0.000870806 | 3.230488208 | -0.029290806 | 0.97751531 | 0.993521701 | -7.032000909 |
| **Q9NXE4-4** | SMPD4 | 0.036839412 | 3.196363897 | 0.786384221 | 0.459463493 | 0.800359011 | -7.032207575 |
| **Q96KA5-1** | CLPTM1L | -0.013806937 | 3.223133801 | -0.240162988 | 0.817632548 | 0.956855294 | -7.032420645 |
| **Q9H6W3** | C14orf169; RIOX1 | 0.021453946 | 3.18073176 | 0.684931327 | 0.517158137 | 0.833280927 | -7.03272005 |
| **Q6NUM9-1** | RETSAT | -0.024789522 | 3.185699409 | -0.642900413 | 0.542385824 | 0.843669625 | -7.03301023 |
| **Q8WYJ6-1** | 1-Sep | -0.029483513 | 3.179231027 | -1.146578244 | 0.292175982 | 0.685264687 | -7.033311425 |
| **Q8N1B4** | VPS52 | -0.01608864 | 3.18069112 | -0.659661507 | 0.53223439 | 0.839951983 | -7.033446549 |
| **P29083** | GTF2E1 | -0.04531948 | 3.17686679 | -0.698460769 | 0.50920047 | 0.82913633 | -7.034275509 |
| **Q9H0H0** | INTS2 | 0.014230908 | 3.181082782 | 0.991184622 | 0.357198647 | 0.734600528 | -7.034654975 |
| **O95243** | MBD4 | -0.00252274 | 3.207917376 | -0.084740887 | 0.935028829 | 0.986793807 | -7.034880348 |
| **Q9NXG2** | THUMPD1 | -0.022996967 | 3.181660851 | -0.997345707 | 0.354417345 | 0.734057823 | -7.035385339 |
| **Q5T1V6** | DDX59 | 0.00344197 | 3.19000807 | 0.107666732 | 0.917521368 | 0.983235402 | -7.035424192 |
| **Q04917** | YWHAH | 0.076279258 | 3.1931379 | 0.980198574 | 0.362200474 | 0.738054037 | -7.035537648 |
| **Q8WWI1-3** | LMO7 | -0.017755881 | 3.190554426 | -0.503801738 | 0.631091768 | 0.887123867 | -7.035950212 |
| **P55884-2** | EIF3B | -0.020677453 | 3.184033781 | -1.466790269 | 0.189276467 | 0.593449695 | -7.036852617 |
| **Q15287-1** | RNPS1 | -0.014083301 | 3.200797171 | -0.881384116 | 0.409649407 | 0.769606181 | -7.036937461 |
| **O60563** | CCNT1 | -0.007326389 | 3.170486009 | -0.246448998 | 0.812967801 | 0.956167049 | -7.037831112 |

| | | | | | | | |
|---|---|---|---|---|---|---|---|
| **Q8IV38** | ANKMY2 | -0.008163146 | 3.175079191 | -0.386756596 | 0.711337091 | 0.921051693 | -7.037852787 |
| **Q6P0N0** | MIS18BP1 | 0.016498663 | 3.150334397 | 0.590798456 | 0.574697678 | 0.859195843 | -7.038464722 |
| **Q8IWF2** | FOXRED2 | -0.009035524 | 3.20440061 | -0.264971388 | 0.799271376 | 0.952055634 | -7.039061944 |
| **Q96EY4** | TMA16 | -0.018916315 | 3.156974235 | -0.435120601 | 0.677614572 | 0.903662615 | -7.039298812 |
| **Q9H1B7** | IRF2BPL | -0.001700065 | 3.184078649 | -0.063378374 | 0.951377997 | 0.989472196 | -7.039548952 |
| **Q8TAD8** | SNIP1 | 0.042046697 | 3.134240106 | 0.445189067 | 0.670690976 | 0.901175787 | -7.039787539 |
| **Q8WU76-1** | SCFD2 | -0.02011296 | 3.16807293 | -0.647259718 | 0.539734047 | 0.843540088 | -7.039837561 |
| **Q13573** | SNW1 | -0.02089237 | 3.174699332 | -1.007423077 | 0.349904852 | 0.731407657 | -7.040053127 |
| **Q96SK2** | TMEM209 | 0.007145947 | 3.215075577 | 0.186913636 | 0.857451979 | 0.968855822 | -7.040208396 |
| **O00154** | ACOT7 | 0.028629989 | 3.181374392 | 1.168073309 | 0.28400313 | 0.679844794 | -7.040484791 |
| **Q5T6F2** | UBAP2 | -0.030258425 | 3.17706222 | -1.076129821 | 0.320343946 | 0.708257023 | -7.043667993 |
| **O95352** | ATG7 | -0.011929694 | 3.180135083 | -0.679258915 | 0.520518271 | 0.834650661 | -7.043949732 |
| **Q6P1J9** | CDC73 | -0.023588366 | 3.185965768 | -1.170582031 | 0.283061907 | 0.679787821 | -7.044457766 |
| **O15381-1** | NVL | -0.022086815 | 3.181645995 | -1.160358242 | 0.28691419 | 0.682301253 | -7.045464373 |
| **P62140** | PPP1CB | 0.023999277 | 3.161489853 | 0.488240939 | 0.641484206 | 0.890911725 | -7.04560289 |
| **Q9NX58** | LYAR | -0.031294776 | 3.160905755 | -0.561167548 | 0.593571174 | 0.869935985 | -7.046655402 |
| **P61006** | RAB8A | -0.004415906 | 3.172407344 | -0.132973357 | 0.898253969 | 0.978757548 | -7.047017412 |
| **O43314-1** | PPIP5K2 | 0.01930231 | 3.186662623 | 0.973565723 | 0.365246686 | 0.740245205 | -7.04714159 |
| **Q96SI9** | STRBP | -0.031902858 | 3.188243296 | -1.19274785 | 0.274859573 | 0.674207874 | -7.047182346 |
| **Q9Y394** | DHRS7 | 0.003115654 | 3.209953709 | 0.087857403 | 0.932646342 | 0.986793807 | -7.047716027 |
| **Q9H8V3-3** | ECT2 | 0.005443271 | 3.132675291 | 0.260996528 | 0.802204308 | 0.953157309 | -7.048694017 |
| **Q14204** | DYNC1H1 | -0.016342376 | 3.172941577 | -2.038373064 | 0.084207399 | 0.428040811 | -7.048772458 |
| **P62136-1** | PPP1CA | -0.003570164 | 3.166082707 | -0.129277948 | 0.901063183 | 0.97952714 | -7.049014808 |
| **P61160-1** | ACTR2 | -0.021481701 | 3.180200141 | -1.283969792 | 0.243206667 | 0.649360189 | -7.050455762 |
| **Q9BQG0-2** | MYBBP1A | -0.057721238 | 3.16651566 | -1.271349559 | 0.247388356 | 0.652280071 | -7.050462203 |
| **Q9UPP1** | PHF8 | 0.012320133 | 3.192003723 | 0.619114324 | 0.556996605 | 0.851117349 | -7.051282285 |
| **Q99729-3** | HNRNPAB | -0.045138152 | 3.181984113 | -0.591097656 | 0.574508914 | 0.85911412 | -7.051324785 |
| **Q14320** | FAM50A | -0.029396479 | 3.195387013 | -0.800752633 | 0.451665685 | 0.79513757 | -7.052053768 |
| **Q9Y3Q8** | TSC22D4 | -0.006266839 | 3.168455105 | -0.284390581 | 0.784994095 | 0.94654395 | -7.052122909 |
| **O43159** | RRP8 | 0.015686809 | 3.171637473 | 0.75132842 | 0.47887972 | 0.811537935 | -7.0526213 |
| **Q9NQC1-1** | PHF15; JADE2 | -0.003021137 | 3.172007817 | -0.14729069 | 0.887385468 | 0.975708125 | -7.053263834 |
| **Q9H0Y0** | ATG10 | 0.003980963 | 3.196370562 | 0.103173807 | 0.920948826 | 0.984164603 | -7.053967537 |
| **Q96B01** | RAD51AP1 | -0.011206579 | 3.167749479 | -0.351237295 | 0.736565489 | 0.929681266 | -7.054223746 |
| **Q14667-1** | KIAA0100 | -0.00346759 | 3.184013557 | -0.106166178 | 0.918665867 | 0.983742622 | -7.054695409 |
| **P06132** | UROD | -0.000171746 | 3.161065126 | -0.004468835 | 0.996569006 | 0.998480498 | -7.055397656 |
| **P20073-1** | ANXA7 | 0.034502781 | 3.178526501 | 0.878040374 | 0.411332836 | 0.769906516 | -7.055694513 |
| **Q8NBF6-1** | AVL9 | -0.002142859 | 3.19815471 | -0.116130181 | 0.911070218 | 0.981900848 | -7.05603866 |
| **Q7Z6B0-1** | CCDC91 | -0.014886345 | 3.213513361 | -0.283503072 | 0.785644692 | 0.946704945 | -7.056792267 |
| **O94819** | KBTBD11 | 0.001160749 | 3.183150175 | 0.061660727 | 0.952693772 | 0.989472196 | -7.058493373 |

| | | | | | | | |
|---|---|---|---|---|---|---|---|
| **P56524** | HDAC4 | -0.014581798 | 3.195372654 | -0.92126928 | 0.389962495 | 0.75855365 | -7.059276847 |
| **Q13155** | AIMP2 | 0.005112637 | 3.168732373 | 0.138614643 | 0.893968599 | 0.978451649 | -7.059306469 |
| **P61106** | RAB14 | -0.013791075 | 3.174981498 | -1.013846842 | 0.347052136 | 0.728788229 | -7.059644632 |
| **Q12765-2** | SCRN1 | -0.007308185 | 3.207601765 | -0.131003852 | 0.899750972 | 0.978757548 | -7.06083374 |
| **Q9UKW4-4** | VAV3 | 0.01795687 | 3.170262721 | 0.931862116 | 0.384855876 | 0.754667212 | -7.061025829 |
| **Q8IWF6** | DENND6A | -0.005402472 | 3.17689449 | -0.181854758 | 0.861260707 | 0.968855822 | -7.061125866 |
| **Q9H0W8** | SMG9 | -0.015985983 | 3.192124054 | -0.734412326 | 0.488446326 | 0.817162784 | -7.062330556 |
| **P82970** | HMGN5 | 0.038815851 | 3.143938096 | 0.833242941 | 0.434378767 | 0.784298014 | -7.062664575 |
| **O75884-1** | RBBP9 | 0.011748715 | 3.159222073 | 0.424949155 | 0.684643612 | 0.905978473 | -7.063695726 |
| **Q14CX7-1** | NAA25 | -0.02613199 | 3.190471185 | -1.21464955 | 0.266954064 | 0.67010832 | -7.063910718 |
| **Q9UBK8-1** | MTRR | 0.001567285 | 3.238805986 | 0.020987666 | 0.983887809 | 0.995510872 | -7.064202971 |
| **O43583** | DENR | 0.014049943 | 3.173642269 | 0.476908211 | 0.649108554 | 0.893610512 | -7.064323902 |
| **P46734-3** | MAP2K3 | -0.011003165 | 3.174867595 | -0.233986191 | 0.822224106 | 0.959068023 | -7.064684679 |
| **Q9BVC5** | C2orf49 | 0.017062567 | 3.185098329 | 0.436771754 | 0.6764768 | 0.903662615 | -7.065058155 |
| **Q09472** | EP300 | -0.008113409 | 3.175953334 | -0.240553578 | 0.81734246 | 0.956656724 | -7.065420625 |
| **O95671-1** | ASMTL | -0.014320706 | 3.196029958 | -0.500355319 | 0.633385787 | 0.887509451 | -7.06576893 |
| **O15212** | PFDN6 | 0.014669945 | 3.182606752 | 1.122549809 | 0.301543688 | 0.691344519 | -7.066338908 |
| **Q9UMN6** | KMT2B; MLL4 | 0.018743125 | 3.179774295 | 0.526613067 | 0.616020365 | 0.880583268 | -7.067353224 |
| **Q15291-1** | RBBP5 | -0.020752709 | 3.193960163 | -0.944669422 | 0.378749872 | 0.750267026 | -7.067384071 |
| **Q9BWS9-1** | CHID1 | -0.004135064 | 3.194327579 | -0.121494626 | 0.90698499 | 0.98075504 | -7.067699848 |
| **Q9Y376** | CAB39 | -0.020235183 | 3.194246467 | -1.001227762 | 0.352673625 | 0.732738778 | -7.067931766 |
| **Q9ULK4** | MED23 | -0.021863988 | 3.182012976 | -1.083817427 | 0.317165812 | 0.704862202 | -7.0684714 |
| **P25789** | PSMA4 | -0.011625163 | 3.169304939 | -1.151739475 | 0.29019582 | 0.684803131 | -7.068525512 |
| **P52739** | ZNF131 | -0.007970918 | 3.193811555 | -0.292717467 | 0.778899184 | 0.944053983 | -7.068536589 |
| **Q86VS8** | HOOK3 | -0.024610151 | 3.176550392 | -1.016801468 | 0.345746219 | 0.726899633 | -7.068585984 |
| **Q9NQ48** | LZTFL1 | 0.024662178 | 3.162445432 | 1.027765712 | 0.340934186 | 0.722995142 | -7.069245827 |
| **Q9UEW8** | STK39 | -0.000566095 | 3.177872344 | -0.020724175 | 0.984090057 | 0.995510872 | -7.069432279 |
| **Q9NV31** | IMP3 | -0.001218745 | 3.199500198 | -0.035656593 | 0.972630878 | 0.992928787 | -7.07094171 |
| **Q96CN7** | ISOC1 | -0.021086555 | 3.171833998 | -0.501304605 | 0.63275348 | 0.887490806 | -7.070947127 |
| **O94842** | TOX4 | -0.012924321 | 3.163288223 | -0.564539612 | 0.591405554 | 0.868764508 | -7.071147523 |
| **Q16566** | CAMK4 | 0.009459348 | 3.201377916 | 0.43787838 | 0.675714765 | 0.903340705 | -7.072169793 |
| **Q9UJ83** | HACL1 | 0.018190159 | 3.169975377 | 0.847689716 | 0.426846589 | 0.779764001 | -7.072204398 |
| **Q14699** | RFTN1 | 0.013001573 | 3.179509444 | 0.633837012 | 0.547924941 | 0.846115385 | -7.072339266 |
| **Q96EP5** | DAZAP1 | 0.007757598 | 3.163751394 | 0.247816824 | 0.811953848 | 0.955776828 | -7.072550208 |
| **O43681** | ASNA1 | 0.015512557 | 3.161079458 | 0.755265651 | 0.476671459 | 0.810259893 | -7.072585056 |
| **Q969G6** | RFK | -0.014821042 | 3.241654402 | -0.256952214 | 0.805192058 | 0.954249678 | -7.072935708 |
| **P39656** | DDOST | -0.01897479 | 3.180220576 | -0.454708762 | 0.664176628 | 0.898345814 | -7.073353901 |
| **Q9ULX6** | AKAP8L | 0.021266931 | 3.189025934 | 0.971867876 | 0.366029635 | 0.740729864 | -7.073364662 |
| **Q7L5D6** | GET4 | -0.006448908 | 3.194194786 | -0.327164413 | 0.753869775 | 0.936379818 | -7.073365097 |

| | | | | | | | |
|---|---|---|---|---|---|---|---|
| **Q8N6H7** | ARFGAP2 | 0.014925041 | 3.172790157 | 0.804868739 | 0.449449104 | 0.794586812 | -7.073749128 |
| **Q00613** | HSF1 | -0.012772769 | 3.202476936 | -0.25258848 | 0.808419762 | 0.955329057 | -7.074365478 |
| **P08236** | GUSB | 0.041431635 | 3.199053649 | 0.636543283 | 0.546267354 | 0.845412981 | -7.074598857 |
| **Q9UNN5-1** | FAF1 | -0.015289828 | 3.176399874 | -0.756290919 | 0.476097562 | 0.809713018 | -7.075093376 |
| **Q96JK2** | DCAF5 | -0.001677275 | 3.186554429 | -0.057248885 | 0.956074116 | 0.989472196 | -7.07519299 |
| **Q8IWB9-2** | TEX2 | -0.00124949 | 3.21203227 | -0.038222849 | 0.970662152 | 0.992521442 | -7.075240491 |
| **Q15024** | EXOSC7 | 0.017005414 | 3.19168063 | 0.368067205 | 0.724565087 | 0.926648409 | -7.076594774 |
| **Q96CB8** | INTS12 | 0.010854592 | 3.204194555 | 0.476141622 | 0.649625967 | 0.893646754 | -7.076833078 |
| **Q05823** | RNASEL | 0.014355322 | 3.202576427 | 0.337387988 | 0.746501221 | 0.934974996 | -7.077032774 |
| **P06280** | GLA | -0.004300545 | 3.19544196 | -0.148143 | 0.886739273 | 0.975708125 | -7.077353364 |
| **Q8TCE6-1** | FAM45A | -0.001639269 | 3.159418649 | -0.074680233 | 0.942724725 | 0.988606569 | -7.078027149 |
| **Q6ZU65-1** | UBN2 | 0.012635764 | 3.170710665 | 0.55875947 | 0.595120464 | 0.871378088 | -7.078081559 |
| **Q9UBW5-1** | BIN2 | -0.0192018 | 3.204246053 | -0.791088227 | 0.456900307 | 0.798548386 | -7.078349298 |
| **Q9BSD7** | NTPCR | -0.003786138 | 3.184324506 | -0.145241683 | 0.888939343 | 0.976409291 | -7.078822116 |
| **Q9NUQ7** | UFSP2 | -0.004465603 | 3.192353201 | -0.231170654 | 0.824319586 | 0.959077751 | -7.079065437 |
| **Q92854-1** | SEMA4D | 0.015321778 | 3.163071941 | 0.827343881 | 0.437481747 | 0.786862931 | -7.07950307 |
| **Q13630** | TSTA3 | -0.008390335 | 3.167783435 | -0.331498014 | 0.750742903 | 0.935635612 | -7.079695408 |
| **Q9UJX2-1** | CDC23 | -0.028223461 | 3.185543093 | -0.808082214 | 0.447723951 | 0.792674781 | -7.080451677 |
| **Q14974** | KPNB1 | -0.014792947 | 3.173389058 | -1.494636757 | 0.1820879 | 0.584756817 | -7.080704288 |
| **O75131** | CPNE3 | -0.018442431 | 3.174023861 | -1.202328599 | 0.271377166 | 0.671838876 | -7.081063622 |
| **P13716-1** | ALAD | -0.001797893 | 3.191276731 | -0.079412742 | 0.939103707 | 0.988002681 | -7.081350061 |
| **P34896-1** | SHMT1 | 0.017709183 | 3.170576788 | 1.01876921 | 0.34487866 | 0.726061285 | -7.082505321 |
| **Q96GD0** | PDXP | -0.002643186 | 3.17903448 | -0.123903051 | 0.905151872 | 0.98075504 | -7.083244156 |
| **Q96CT7** | CCDC124 | -0.016172665 | 3.182793811 | -0.258670967 | 0.803921887 | 0.95391847 | -7.08378555 |
| **Q12906-7** | ILF3 | 0.021955758 | 3.175923388 | 1.550370251 | 0.168451188 | 0.566974321 | -7.084459254 |
| **Q96L92** | SNX27 | 0.017855046 | 3.194979547 | 0.608141174 | 0.563816887 | 0.852578515 | -7.084698293 |
| **Q8WYA6** | CTNNBL1 | 0.021204686 | 3.167568867 | 1.19163472 | 0.275266624 | 0.674789673 | -7.084775498 |
| **Q53EZ4-1** | CEP55 | -0.036719922 | 3.18440325 | -0.756196238 | 0.47615054 | 0.809713018 | -7.084850244 |
| **P04150-3** | NR3C1 | -0.009458094 | 3.191817837 | -0.549429722 | 0.601144608 | 0.872497827 | -7.085226231 |
| **Q9UGV2** | NDRG3 | 0.008288966 | 3.194190592 | 0.453711593 | 0.664857525 | 0.898624353 | -7.085368914 |
| **Q86VM9** | ZC3H18 | -0.017393085 | 3.190307343 | -0.771299624 | 0.467750474 | 0.805435093 | -7.08544161 |
| **Q9NZ52** | GGA3 | -0.000957063 | 3.198208652 | -0.039260052 | 0.969866516 | 0.992476747 | -7.085506419 |
| **Q00577** | PURA | -0.000212822 | 3.183098948 | -0.008506664 | 0.993468988 | 0.99756865 | -7.086797192 |
| **Q9NQV6-6** | PRDM10 | -0.004181902 | 3.206899015 | -0.197250641 | 0.84968242 | 0.96500901 | -7.087673558 |
| **Q9H773** | DCTPP1 | -0.026453139 | 3.203224562 | -0.32858179 | 0.752846522 | 0.935782019 | -7.087729244 |
| **P51617** | IRAK1 | 0.001773843 | 3.215005076 | 0.075016246 | 0.942467579 | 0.988606569 | -7.088265027 |
| **P42694** | HELZ | 0.010465898 | 3.186329554 | 0.891379798 | 0.404647463 | 0.766336431 | -7.0885092 |
| **Q9H9E3-1** | COG4 | -0.039530938 | 3.152484607 | -0.888378633 | 0.406144484 | 0.766336431 | -7.088650746 |
| **Q8TBX8-1** | PIP4K2C | -0.004581101 | 3.18887705 | -0.17441444 | 0.86686966 | 0.970722967 | -7.088706343 |

| | | | | | | | |
|---|---|---|---|---|---|---|---|
| **P40200-1** | CD96 | -0.000456829 | 3.184595963 | -0.019789072 | 0.984807825 | 0.995531556 | -7.088823812 |
| **Q96FJ0-2** | STAMBPL1 | 0.000340115 | 3.226120387 | 0.005951817 | 0.995430445 | 0.998254835 | -7.088863577 |
| **Q9NQ29** | LUC7L | -0.035010424 | 3.171202612 | -0.52926632 | 0.61428019 | 0.879837049 | -7.088976695 |
| **Q8NEB9** | PIK3C3 | -0.013864995 | 3.182505757 | -0.765508958 | 0.470958943 | 0.807027916 | -7.089902822 |
| **Q86UA1** | PRPF39 | -0.013787343 | 3.181969071 | -0.49736715 | 0.635378348 | 0.888121213 | -7.090079288 |
| **Q9UGN5-1** | PARP2 | 0.039627642 | 3.190447638 | 0.703031424 | 0.506530254 | 0.82913633 | -7.090399776 |
| **Q9H9C1-1** | VIPAS39 | 0.000456536 | 3.1757724 | 0.017845396 | 0.986299802 | 0.995572599 | -7.090576166 |
| **Q14738-1** | PPP2R5D | -0.028657008 | 3.193784288 | -0.83241289 | 0.434814426 | 0.784298014 | -7.090688227 |
| **Q9BRX2** | PELO | -0.008906537 | 3.174040546 | -0.258460988 | 0.804077028 | 0.95391847 | -7.090700248 |
| **Q9Y496** | KIF3A | -0.016766076 | 3.193979207 | -0.611049093 | 0.562004614 | 0.852576894 | -7.092371704 |
| **Q9H3H3-3** | C11orf68 | 0.009988545 | 3.167641045 | 0.459425132 | 0.660960835 | 0.896934512 | -7.093549325 |
| **P31483** | TIA1 | 0.008535575 | 3.18126526 | 0.22158378 | 0.831466284 | 0.960313945 | -7.093992959 |
| **Q32P41** | TRMT5 | 0.022089817 | 3.18203842 | 0.698702012 | 0.509059304 | 0.82913633 | -7.094451179 |
| **Q7Z4H8** | KDELC2 | -0.014559877 | 3.172103292 | -0.704249996 | 0.505819905 | 0.82913633 | -7.095193642 |
| **Q9BY12** | SCAPER | -0.004795523 | 3.179131909 | -0.238124677 | 0.819146886 | 0.957819113 | -7.095259273 |
| **Q13362-4** | PPP2R5C | -0.002605871 | 3.20725684 | -0.089665177 | 0.931264695 | 0.986707862 | -7.095700838 |
| **Q05655** | PRKCD | 0.009634253 | 3.178990154 | 0.277737088 | 0.789876023 | 0.947865984 | -7.095881762 |
| **O60476** | MAN1A2 | -0.007305155 | 3.167585591 | -0.250540144 | 0.809936253 | 0.955498404 | -7.096470354 |
| **O00303** | EIF3F | -0.028232229 | 3.194804679 | -1.106068017 | 0.308112126 | 0.696616637 | -7.09653982 |
| **P14314** | PRKCSH | 0.032048085 | 3.180537526 | 1.185426426 | 0.277546276 | 0.675997904 | -7.097140156 |
| **Q8IUX7** | AEBP1 | 0.010473203 | 3.234231168 | 0.296415606 | 0.776197706 | 0.94357883 | -7.097462178 |
| **O94829** | IPO13 | -0.010309657 | 3.188121034 | -0.462548829 | 0.658835264 | 0.896852471 | -7.097593008 |
| **Q9Y6V7** | DDX49 | -0.031965621 | 3.16763563 | -0.331658493 | 0.750627208 | 0.935635612 | -7.097812263 |
| **Q9UBN7-1** | HDAC6 | 0.018615241 | 3.178050444 | 0.652658573 | 0.536461142 | 0.842162373 | -7.098291877 |
| **Q53H82** | LACTB2 | 0.018903854 | 3.167051986 | 0.618600863 | 0.557314623 | 0.851117349 | -7.098456473 |
| **P51451** | BLK | -0.008480912 | 3.17923854 | -0.302894331 | 0.771473167 | 0.94239765 | -7.098580037 |
| **Q9Y5Z4-1** | HEBP2 | 0.003432466 | 3.181265072 | 0.089432793 | 0.931442287 | 0.986707862 | -7.098667632 |
| **Q96IU4** | ABHD14B | 0.008039476 | 3.163630792 | 0.261582425 | 0.801771772 | 0.953157309 | -7.098992072 |
| **Q9NVM9** | ASUN; INTS13 | -0.010943125 | 3.179567262 | -1.023806738 | 0.342665534 | 0.724485042 | -7.099330076 |
| **O00255-1** | MEN1 | -0.00178108 | 3.179940843 | -0.035426034 | 0.972807763 | 0.992928787 | -7.099641069 |
| **P14550** | AKR1A1 | -0.025925231 | 3.175361472 | -1.150383434 | 0.290714989 | 0.684973485 | -7.099893305 |
| **Q9H4M9** | EHD1 | 0.040629193 | 3.178985118 | 1.163448835 | 0.285745047 | 0.681580409 | -7.100763973 |
| **Q9H3P7** | ACBD3 | 0.010778937 | 3.187801327 | 1.003281073 | 0.351754066 | 0.732738778 | -7.101597775 |
| **Q9NX62** | IMPAD1 | 0.004242248 | 3.18559106 | 0.159762974 | 0.877938952 | 0.973200128 | -7.102167332 |
| **Q99426** | TBCB | -0.023727294 | 3.187936955 | -0.449130367 | 0.667990148 | 0.90001396 | -7.102232303 |
| **P51531** | SMARCA2 | 0.016532564 | 3.199915736 | 0.435746733 | 0.677183011 | 0.903662615 | -7.102276414 |
| **P21580** | TNFAIP3 | 0.011422296 | 3.184633448 | 0.482949187 | 0.645038567 | 0.891546795 | -7.102970318 |
| **Q9H074** | PAIP1 | -0.00983347 | 3.145498899 | -0.287308885 | 0.782856129 | 0.945688299 | -7.10326648 |
| **O95235** | KIF20A | 0.031200591 | 3.127833616 | 0.88982535 | 0.405422329 | 0.766336431 | -7.103367741 |

| | | | | | | | |
|---|---|---|---|---|---|---|---|
| **Q9HD26-1** | GOPC | -0.019771466 | 3.198436903 | -0.832186559 | 0.434933272 | 0.784298014 | -7.104126229 |
| **P06753-2** | TPM3 | -0.02173956 | 3.166630302 | -0.370787427 | 0.722633242 | 0.926648409 | -7.105058817 |
| **Q9UBF2** | COPG2 | 0.021656659 | 3.178266095 | 1.116584901 | 0.303907377 | 0.692441671 | -7.105081185 |
| **Q9NSY1** | BMP2K | 0.003921694 | 3.20341678 | 0.160422869 | 0.877439736 | 0.973095854 | -7.105378645 |
| **P29728** | OAS2 | -0.001581834 | 3.149426105 | -0.055397243 | 0.957493137 | 0.989500907 | -7.105573187 |
| **Q8WVS4** | WDR60 | -0.007343302 | 3.171879916 | -0.322481174 | 0.75725459 | 0.938154954 | -7.105688985 |
| **Q4KMQ2-2** | ANO6 | 0.018098019 | 3.160379772 | 0.501501791 | 0.632622178 | 0.887490806 | -7.105725362 |
| **P25098** | ADRBK1; GRK2 | -0.020239315 | 3.228494711 | -0.433763753 | 0.678550231 | 0.903662615 | -7.10609233 |
| **Q9Y3B4** | SF3B14; SF3B6 | -0.003779627 | 3.15833616 | -0.181717464 | 0.86136413 | 0.968855822 | -7.106392771 |
| **Q86TI2-2** | DPP9 | -0.009338531 | 3.181004574 | -0.603727344 | 0.566574364 | 0.853395233 | -7.106580155 |
| **Q86W42** | THOC6 | 0.019759108 | 3.1836748 | 0.469925518 | 0.653829317 | 0.896119351 | -7.107015169 |
| **Q8NEY8-2** | PPHLN1 | 0.004575087 | 3.198739562 | 0.211622696 | 0.838910384 | 0.961364456 | -7.107438677 |
| **P48556** | PSMD8 | 0.019611105 | 3.173072865 | 0.862071113 | 0.419443095 | 0.775923022 | -7.107478977 |
| **Q6P4F7** | ARHGAP11A | -0.016148331 | 3.14729986 | -0.529390775 | 0.614198631 | 0.879837049 | -7.10767277 |
| **P07858** | CTSB | -0.027759781 | 3.20933807 | -0.66078598 | 0.531557654 | 0.839849063 | -7.108021247 |
| **Q9Y2B0** | CNPY2 | 0.010493839 | 3.185308667 | 0.305675825 | 0.769448017 | 0.941649022 | -7.108141914 |
| **P20645** | M6PR | 0.020855187 | 3.198655399 | 0.232517943 | 0.823316662 | 0.959068023 | -7.110337339 |
| **Q8WWW0-2** | RASSF5 | -0.006716289 | 3.187350602 | -0.312228454 | 0.764684947 | 0.940369694 | -7.111665489 |
| **O75381-1** | PEX14 | -0.013160238 | 3.180726611 | -0.475113444 | 0.650320272 | 0.893885477 | -7.112125247 |
| **Q9H993** | C6orf211; ARMT1 | -0.008032133 | 3.203278955 | -0.212426554 | 0.838308967 | 0.961230299 | -7.112152003 |
| **Q9Y6E0** | STK24 | 0.003194783 | 3.185607968 | 0.105252529 | 0.919362827 | 0.983742622 | -7.112278049 |
| **P61758** | VBP1 | -0.100804239 | 3.150136196 | -0.918134757 | 0.391483392 | 0.759806823 | -7.11229557 |
| **Q13510-2** | ASAH1 | -0.008572615 | 3.173302989 | -0.265076886 | 0.79919358 | 0.952055634 | -7.112412924 |
| **Q9BUA3** | C11orf84 | 0.001099009 | 3.182686506 | 0.05246361 | 0.959741694 | 0.990664903 | -7.112504021 |
| **Q6VMQ6-4** | ATF7IP | -0.019801827 | 3.183633112 | -0.715591826 | 0.499239733 | 0.824970816 | -7.11283584 |
| **O75688** | PPM1B | 0.012582062 | 3.152985081 | 0.869800611 | 0.415502995 | 0.77203561 | -7.11288859 |
| **Q8N0W3-2** | FUK | -0.001846838 | 3.209482791 | -0.06174205 | 0.952631472 | 0.989472196 | -7.112966859 |
| **O95391** | SLU7 | -0.01694673 | 3.179229949 | -0.864051299 | 0.418431098 | 0.775013192 | -7.113388914 |
| **Q5R372** | RABGAP1L | -0.005235775 | 3.193979655 | -0.268086566 | 0.79697524 | 0.950886893 | -7.114313545 |
| **Q9NQT5** | EXOSC3 | -0.009191105 | 3.184153419 | -0.0969432 | 0.925704903 | 0.985476218 | -7.114619584 |
| **Q7L523** | RRAGA | 0.005061336 | 3.185788569 | 0.253557896 | 0.807702364 | 0.954797691 | -7.114709635 |
| **Q14019** | COTL1 | -0.033369685 | 3.176554618 | -0.739475356 | 0.485569611 | 0.81473183 | -7.115116508 |
| **Q9NVT9** | ARMC1 | -0.003655806 | 3.181182156 | -0.182533014 | 0.860749826 | 0.968855822 | -7.115190788 |
| **Q9P215** | POGK | -0.024590314 | 3.193342348 | -0.690606749 | 0.513810254 | 0.83148099 | -7.115595912 |
| **Q8NEM2** | SHCBP1 | -0.009176308 | 3.189790368 | -0.491923262 | 0.639016886 | 0.88909352 | -7.116039154 |
| **P84085** | ARF5 | -0.009136829 | 3.161185294 | -0.298715085 | 0.774519641 | 0.943218662 | -7.116061349 |
| **Q9C0E8-4** | KIAA1715; LNPK | 0.009477037 | 3.186798571 | 0.443875772 | 0.671592111 | 0.901175787 | -7.116108719 |
| **P62314** | SNRPD1 | -0.038733994 | 3.168728769 | -0.877308541 | 0.411701963 | 0.770365544 | -7.116604564 |
| **P54274-1** | TERF1 | -0.013128706 | 3.186658038 | -0.317657233 | 0.760747175 | 0.938739432 | -7.117891537 |

| | | | | | | | |
|---|---|---|---|---|---|---|---|
| **O60832-1** | DKC1 | 0.031769285 | 3.181219883 | 1.144439475 | 0.292999847 | 0.68530312 | -7.118422671 |
| **Q05D32** | CTDSPL2 | -0.019495782 | 3.158641037 | -0.683584213 | 0.517954851 | 0.83340353 | -7.119503406 |
| **Q13153** | PAK1 | 0.025983621 | 3.196321838 | 0.701184503 | 0.507608136 | 0.82913633 | -7.119997242 |
| **Q86WA8** | LONP2 | 0.004308786 | 3.193822504 | 0.181478811 | 0.861543912 | 0.968855822 | -7.12048697 |
| **Q9P2W1** | PSMC3IP | 0.005210579 | 3.173979265 | 0.16093895 | 0.877049357 | 0.973046824 | -7.120924476 |
| **Q5TDH0-3** | DDI2 | -0.003569104 | 3.211865163 | -0.134943548 | 0.896756896 | 0.978757548 | -7.12093365 |
| **Q5VSL9-1** | STRIP1 | -0.009632637 | 3.172327491 | -0.641986011 | 0.542943082 | 0.843669625 | -7.121067541 |
| **P32780** | GTF2H1 | -0.017425843 | 3.149240017 | -0.45726449 | 0.662433077 | 0.897184895 | -7.121104059 |
| **Q14139-2** | UBE4A | 0.012466373 | 3.19596749 | 0.647618976 | 0.539515871 | 0.843540088 | -7.121142495 |
| **P42226** | STAT6 | 0.004332845 | 3.179367159 | 0.173941269 | 0.867226651 | 0.970868803 | -7.121783412 |
| **Q9H6D7** | HAUS4 | -0.006759707 | 3.179000194 | -0.337224853 | 0.746618576 | 0.934974996 | -7.121905136 |
| **Q9NX46** | ADPRHL2 | -0.010244195 | 3.171778763 | -0.528210705 | 0.614972208 | 0.880352106 | -7.122272913 |
| **Q6UX04** | CWC27 | 0.012945233 | 3.179712555 | 0.651435344 | 0.537201599 | 0.84315818 | -7.122323984 |
| **Q15057** | ACAP2 | 0.013233936 | 3.188029225 | 0.892766723 | 0.403957035 | 0.766336431 | -7.122345518 |
| **Q9BTE6-1** | AARSD1; PTGES3L | 0.018425516 | 3.166401372 | 0.906156274 | 0.397336713 | 0.762571436 | -7.123145544 |
| **Q9UHJ3-1** | SFMBT1 | -0.00065045 | 3.1563281 | -0.025201388 | 0.980653667 | 0.994229224 | -7.123468794 |
| **Q13164-1** | MAPK7 | -0.007992831 | 3.19822219 | -0.239775072 | 0.81792068 | 0.957051517 | -7.124254185 |
| **O75431** | MTX2 | 0.001952469 | 3.177139418 | 0.056916644 | 0.956328718 | 0.989472196 | -7.124405141 |
| **Q9BTU6** | PI4K2A | -0.025126434 | 3.157660602 | -0.655150857 | 0.534954468 | 0.840674016 | -7.124550915 |
| **Q8IWA4-1** | MFN1 | 0.002864174 | 3.17211698 | 0.103019869 | 0.921066292 | 0.984164603 | -7.124582879 |
| **O43665-3** | RGS10 | -0.009505083 | 3.187019559 | -0.197074945 | 0.849814329 | 0.96500901 | -7.124755507 |
| **Q96DA2** | RAB39B | 0.004514996 | 3.191488611 | 0.1547717 | 0.881716843 | 0.974438769 | -7.125274727 |
| **Q86W56** | PARG; BPHL | -0.020145536 | 3.183769055 | -0.911059715 | 0.394932723 | 0.761033346 | -7.12528778 |
| **O95747** | OXSR1 | -0.010938725 | 3.159758775 | -0.543202963 | 0.605184203 | 0.87421609 | -7.125561163 |
| **Q66K74** | MAP1S | -0.012830955 | 3.169536979 | -0.410721253 | 0.694533145 | 0.911173355 | -7.126960114 |
| **Q8IWC1** | MAP7D3 | -0.031264951 | 3.205624346 | -1.01974497 | 0.344449097 | 0.725897233 | -7.127154793 |
| **P0C0S5** | H2AFZ | -0.028628737 | 3.197889411 | -1.02045632 | 0.344136205 | 0.725434371 | -7.127372762 |
| **Q9Y570** | PPME1 | 0.029662115 | 3.203777504 | 0.888249832 | 0.406208824 | 0.766336431 | -7.127545409 |
| **P21399** | ACO1 | -0.021208849 | 3.163284599 | -1.032053444 | 0.339066936 | 0.721920542 | -7.127916405 |
| **P45974-1** | USP5 | 0.026914858 | 3.17857244 | 1.121815519 | 0.301833838 | 0.691344519 | -7.128040725 |
| **O43617-1** | TRAPPC3 | -0.009184119 | 3.193639644 | -0.229750124 | 0.825377417 | 0.959077751 | -7.128143679 |
| **Q9BW71-1** | HIRIP3 | -0.005046438 | 3.162796483 | -0.09629963 | 0.926196358 | 0.985528933 | -7.128245547 |
| **P56282** | POLE2 | -0.009884777 | 3.196384441 | -0.36723384 | 0.725157363 | 0.926648409 | -7.128310893 |
| **Q9H5N1** | RABEP2 | -0.009433744 | 3.152788089 | -0.183817423 | 0.859782572 | 0.968855822 | -7.128340755 |
| **Q8ND04-2** | SMG8 | -0.012246376 | 3.17776549 | -0.849021505 | 0.426157022 | 0.779649091 | -7.128799502 |
| **Q8WX93-1** | PALLD | -0.009208541 | 3.184660837 | -0.508349422 | 0.628071488 | 0.8858562 | -7.128993488 |
| **Q13614-1** | MTMR2 | 0.003480585 | 3.209886236 | 0.119752894 | 0.908311055 | 0.980855153 | -7.128996918 |
| **Q13129** | RLF | -0.019803487 | 3.190550182 | -0.717520736 | 0.498126293 | 0.824674598 | -7.130128575 |
| **P40937-1** | RFC5 | -0.038109922 | 3.178569703 | -1.189784069 | 0.275944506 | 0.67488468 | -7.130238871 |

| | | | | | | | |
|---|---|---|---|---|---|---|---|
| **P42892** | ECE1 | 0.000171743 | 3.227233446 | 0.008445072 | 0.993516274 | 0.99756865 | -7.13042607 |
| **O95470** | SGPL1 | 0.006710616 | 3.188700581 | 0.199252953 | 0.848179503 | 0.964336885 | -7.131645154 |
| **Q8WTW3** | COG1 | -0.015406783 | 3.181916176 | -0.637947661 | 0.545408395 | 0.845033545 | -7.132583333 |
| **Q9NUU7** | DDX19A | -0.031082163 | 3.183369134 | -1.059865384 | 0.32715313 | 0.71218881 | -7.133291938 |
| **Q15276** | RABEP1 | -0.026042637 | 3.185043075 | -1.133757481 | 0.297143669 | 0.689298697 | -7.13342663 |
| **P28066-1** | PSMA5 | -0.018891386 | 3.174559465 | -1.220314151 | 0.264941294 | 0.66823064 | -7.134393034 |
| **Q8NBS9-1** | TXNDC5 | 0.02865788 | 3.195896048 | 1.000535832 | 0.352983927 | 0.732738778 | -7.134459628 |
| **Q6UB99** | ANKRD11 | -0.012134841 | 3.195818591 | -0.556224741 | 0.59675372 | 0.871378088 | -7.134545349 |
| **Q15345-2** | LRRC41 | -0.014881261 | 3.207983925 | -0.566902809 | 0.589890553 | 0.868583051 | -7.134574124 |
| **O75391** | SPAG7 | 0.00119341 | 3.174253537 | 0.043169864 | 0.96686765 | 0.992288899 | -7.134617399 |
| **Q86UE8-2** | TLK2 | 0.006030032 | 3.184531043 | 0.350034546 | 0.737426223 | 0.929681266 | -7.134854189 |
| **Q6P1M0** | SLC27A4 | -0.009258222 | 3.194792772 | -0.364556496 | 0.727061551 | 0.92710315 | -7.136330485 |
| **Q9BT40** | INPP5K | -0.00583318 | 3.15587013 | -0.268903777 | 0.79637325 | 0.950757673 | -7.136770098 |
| **P14373** | TRIM27 | -0.025312551 | 3.212194821 | -0.593998698 | 0.572680556 | 0.857999396 | -7.136897457 |
| **Q9ULG1** | INO80 | 0.007006708 | 3.191030194 | 0.283874514 | 0.78537238 | 0.946704945 | -7.137157065 |
| **O95714** | HERC2 | 0.014767624 | 3.173043038 | 1.066350446 | 0.324424209 | 0.710703247 | -7.137639102 |
| **Q9HCE5** | METTL14 | 0.006584291 | 3.200936466 | 0.161318123 | 0.876762564 | 0.973040052 | -7.137851823 |
| **Q96NB3** | ZNF830 | 0.009506666 | 3.180804443 | 0.466890319 | 0.655886726 | 0.896497734 | -7.137961536 |
| **Q9NPG3-1** | UBN1 | 0.027017126 | 3.189003854 | 0.786028605 | 0.459657673 | 0.800521747 | -7.13828009 |
| **Q86UE4** | MTDH | -0.024723793 | 3.191027354 | -1.165637895 | 0.284919369 | 0.68101215 | -7.138371498 |
| **Q9P1Y6-1** | PHRF1 | -0.007592442 | 3.226293237 | -0.317924844 | 0.760553261 | 0.938728548 | -7.138558717 |
| **O14733-2** | MAP2K7 | -0.00725975 | 3.197111321 | -0.386782498 | 0.711318832 | 0.921051693 | -7.138785925 |
| **Q14558-2** | PRPSAP1 | 0.01007533 | 3.165292491 | 0.575550657 | 0.584365636 | 0.8653224 | -7.138972189 |
| **Q92733** | PRCC | -0.00847999 | 3.186700732 | -0.321016012 | 0.758314734 | 0.9384213 | -7.1395254 |
| **P29016** | CD1B | -0.01627608 | 3.144296108 | -0.359036477 | 0.730994142 | 0.92815165 | -7.139636688 |
| **Q9BRJ7-1** | NUDT16L1 | -0.003092338 | 3.183212527 | -0.090338924 | 0.930749828 | 0.986707862 | -7.139686207 |
| **Q15370** | TCEB2; ELOB | 0.035032004 | 3.160771681 | 0.499671175 | 0.633841694 | 0.887509451 | -7.139866622 |
| **P30520** | ADSS | 0.012123495 | 3.197828667 | 0.809238585 | 0.447104301 | 0.792204208 | -7.140017475 |
| **Q9Y2H6** | FNDC3A | -0.008849105 | 3.196470492 | -0.476767878 | 0.649203257 | 0.893610512 | -7.140863704 |
| **Q9H0U3** | MAGT1 | -0.004428985 | 3.212965321 | -0.136161154 | 0.895831913 | 0.978757548 | -7.140915425 |
| **Q6AI08** | HEATR6 | -0.003894818 | 3.183924922 | -0.154818319 | 0.881681541 | 0.974438769 | -7.141477357 |
| **P61019-1** | RAB2A | -0.011028135 | 3.163111743 | -0.643852322 | 0.541806085 | 0.843570684 | -7.142615689 |
| **Q12986** | NFX1 | -0.016502037 | 3.202645116 | -0.719763988 | 0.496833474 | 0.823442885 | -7.142837264 |
| **O43639** | NCK2 | 0.012280446 | 3.19831357 | 0.333598837 | 0.749228902 | 0.935521689 | -7.142847931 |
| **Q8NCN4** | RNF169 | 0.005294481 | 3.180539919 | 0.244022871 | 0.814767223 | 0.95620584 | -7.14370892 |
| **P53611** | RABGGTB | -0.012324283 | 3.236281399 | -0.246144875 | 0.813193298 | 0.956167049 | -7.14612656 |
| **O43402** | EMC8 | -0.009572366 | 3.178509948 | -0.493913881 | 0.63768515 | 0.889065271 | -7.146270081 |
| **P55060-1** | CSE1L | -0.025071492 | 3.182495892 | -1.53650048 | 0.171753308 | 0.571776747 | -7.146476843 |
| **Q9H3U1** | UNC45A | -0.024794217 | 3.174156368 | -1.135655002 | 0.296404028 | 0.688880164 | -7.146696686 |

| | | | | | | | |
|---|---|---|---|---|---|---|---|
| **Q9NVR2** | INTS10 | 0.010948766 | 3.196526144 | 0.412851771 | 0.693048069 | 0.909991705 | -7.147146008 |
| **Q9NZG7** | NINJ2 | -0.010963219 | 3.189753321 | -0.413088172 | 0.692883376 | 0.909991705 | -7.147227442 |
| **Q9BPZ7** | MAPKAP1 | 0.012186525 | 3.197315332 | 0.310716863 | 0.765782739 | 0.940484723 | -7.147537256 |
| **Q8IXJ6** | SIRT2 | -0.012531419 | 3.193201385 | -0.606873854 | 0.564607802 | 0.852624219 | -7.147770217 |
| **Q8IWS0-1** | PHF6 | 0.009164492 | 3.181514899 | 0.183380245 | 0.860111769 | 0.968855822 | -7.148589829 |
| **Q9ULL5-3** | PRR12 | -0.009490111 | 3.165023533 | -0.47881783 | 0.647820565 | 0.893099842 | -7.148933069 |
| **P45983-3** | MAPK8 | -0.013838627 | 3.185654706 | -0.400328331 | 0.701798389 | 0.914813487 | -7.148969915 |
| **O96017-9** | CHEK2 | 0.012336505 | 3.199763185 | 0.611160283 | 0.561935388 | 0.852576894 | -7.149353177 |
| **Q8WV22** | NSMCE1 | 0.019858879 | 3.173181773 | 0.300901513 | 0.772925294 | 0.943005565 | -7.149562521 |
| **Q6IA69** | NADSYN1 | -0.00184904 | 3.197845081 | -0.035777981 | 0.972537749 | 0.992928787 | -7.150776669 |
| **Q8WVT3** | TRAPPC12 | -0.017932411 | 3.183414345 | -0.674852265 | 0.523138265 | 0.836627689 | -7.150794853 |
| **Q15269** | PWP2; LOC102724 | -0.022055928 | 3.200728751 | -0.947836459 | 0.377251452 | 0.749474764 | -7.150940544 |
| **P49643-1** | PRIM2 | -0.009735272 | 3.181863961 | -0.429898156 | 0.681219264 | 0.903980599 | -7.151106977 |
| **P42229** | STAT5A | 0.004031545 | 3.185002048 | 0.167745394 | 0.871904307 | 0.971931713 | -7.151161384 |
| **Q9BPY3** | FAM118B | 0.002987903 | 3.188448647 | 0.085279542 | 0.93461699 | 0.986793807 | -7.151309133 |
| **Q9UKN8** | GTF3C4 | -0.025751275 | 3.195317873 | -1.146557886 | 0.292183815 | 0.685264687 | -7.1514173 |
| **Q9Y312** | AAR2 | -0.000517398 | 3.201788487 | -0.021300772 | 0.98364748 | 0.995424688 | -7.151656928 |
| **Q9NZD8** | SPG21 | -0.001321714 | 3.187201619 | -0.063076753 | 0.951609037 | 0.989472196 | -7.151870711 |
| **O94927** | HAUS5 | -0.007632279 | 3.176154543 | -0.364521083 | 0.727086753 | 0.92710315 | -7.152172931 |
| **Q96EK6** | GNPNAT1 | 0.007462091 | 3.194677807 | 0.250440523 | 0.81001003 | 0.955498404 | -7.15226421 |
| **Q9Y2K7** | KDM2A | -0.015077261 | 3.187784535 | -0.973397319 | 0.365324286 | 0.740245205 | -7.152271314 |
| **Q15102** | PAFAH1B3 | -0.014795585 | 3.178472764 | -0.661965832 | 0.530848174 | 0.839849063 | -7.152710855 |
| **Q9H4I2** | ZHX3 | 0.006610559 | 3.192240881 | 0.254052188 | 0.807336651 | 0.954765339 | -7.152938047 |
| **Q8N5Y8-3** | PARP16 | -0.01284779 | 3.19211311 | -0.250737419 | 0.809790161 | 0.955498404 | -7.153006076 |
| **O75694** | NUP155 | -0.011699788 | 3.181118609 | -1.179802061 | 0.279625298 | 0.676725357 | -7.153431962 |
| **O75449-1** | KATNA1 | 0.00690043 | 3.163261207 | 0.336176462 | 0.74737293 | 0.934974996 | -7.153806743 |
| **Q05932-1** | FPGS | -0.005673749 | 3.211045706 | -0.324742636 | 0.755619386 | 0.937349811 | -7.153823693 |
| **Q9Y237-2** | PIN4 | 0.009454423 | 3.16707035 | 0.475469238 | 0.650079969 | 0.893733406 | -7.153976429 |
| **Q96FW1** | OTUB1 | 0.011094108 | 3.172853915 | 0.735656089 | 0.487738585 | 0.816666609 | -7.154809266 |
| **O14787** | TNPO2 | -0.014656657 | 3.183380962 | -0.351221346 | 0.7365769 | 0.929681266 | -7.155118305 |
| **P41226** | UBA7 | 0.00859595 | 3.192260262 | 0.465247698 | 0.657001536 | 0.896852471 | -7.155241623 |
| **Q8WUX9** | CHMP7 | 0.004168746 | 3.208192009 | 0.071241348 | 0.945356876 | 0.98904859 | -7.155257176 |
| **Q6WCQ1-2** | MPRIP | -0.012598805 | 3.16904578 | -0.850630574 | 0.425324966 | 0.778986255 | -7.155439766 |
| **Q14181** | POLA2 | -0.01975652 | 3.190180405 | -0.52532412 | 0.616866713 | 0.881353777 | -7.156730648 |
| **Q9Y696** | CLIC4 | 0.012812105 | 3.139100612 | 0.70687868 | 0.504289775 | 0.828319674 | -7.158022662 |
| **Q96S94** | CCNL2 | -0.003686286 | 3.210632573 | -0.13287419 | 0.898329335 | 0.978757548 | -7.158271015 |
| **Q01196-8** | RUNX1 | 0.014977946 | 3.178755046 | 0.46990137 | 0.653845674 | 0.896119351 | -7.15971864 |
| **P62873** | GNB1 | -0.006276985 | 3.171532408 | -0.238870294 | 0.818592841 | 0.957415004 | -7.159908932 |
| **Q9Y2W1** | THRAP3 | 0.050230171 | 3.167542912 | 1.306241403 | 0.235977633 | 0.642113083 | -7.159917883 |

| | | | | | | | |
|---|---|---|---|---|---|---|---|
| **Q9H0K6** | PUS7L | -0.025032453 | 3.167800267 | -0.664811822 | 0.529139262 | 0.839849063 | -7.160067187 |
| **P42285** | SKIV2L2 | -0.014420974 | 3.195175587 | -1.190645259 | 0.275628881 | 0.674825663 | -7.160235879 |
| **Q8N9N7** | LRRC57 | 0.004108231 | 3.164887721 | 0.147637223 | 0.887122726 | 0.975708125 | -7.160949525 |
| **Q9GZZ9-1** | UBA5 | 0.012241768 | 3.179934799 | 0.434634366 | 0.6779498 | 0.903662615 | -7.161208354 |
| **Q9BWT3** | PAPOLG | 0.011952545 | 3.190942481 | 0.606389115 | 0.564910496 | 0.852624219 | -7.161224967 |
| **Q76I76** | SSH2 | -0.013910271 | 3.189333642 | -0.954915115 | 0.373918784 | 0.746240828 | -7.161736916 |
| **Q9NVG8** | TBC1D13 | 0.006555492 | 3.191244269 | 0.226902181 | 0.827499396 | 0.959077751 | -7.162522974 |
| **P55735-1** | SEC13 | 0.012480773 | 3.159407711 | 1.043270097 | 0.334220862 | 0.719277002 | -7.16284569 |
| **A6NC98** | CCDC88B | -0.01278857 | 3.189284307 | -0.495236393 | 0.636801186 | 0.888997073 | -7.16320148 |
| **O14686-3** | KMT2D; MLL2 | 0.013856161 | 3.18026972 | 0.388069357 | 0.710411935 | 0.920754132 | -7.163624485 |
| **P62826** | RAN | 0.018102705 | 3.182210229 | 1.473843518 | 0.187431448 | 0.590695302 | -7.163780014 |
| **Q8WVV9** | HNRNPLL; HNRPLI | 0.034018337 | 3.187240293 | 0.905059404 | 0.397875973 | 0.762880385 | -7.164427698 |
| **Q8TF42** | UBASH3B | 0.016833249 | 3.198585373 | 0.396044631 | 0.704802913 | 0.916550956 | -7.16476253 |
| **P30041** | PRDX6 | 0.033033036 | 3.166851093 | 1.186198231 | 0.277262005 | 0.675719826 | -7.166291986 |
| **O00567** | NOP56 | -0.03870089 | 3.189063659 | -1.293978039 | 0.23993448 | 0.645958461 | -7.167379075 |
| **Q15054** | POLD3 | -0.009114682 | 3.180335502 | -0.517471765 | 0.622036398 | 0.883385606 | -7.168369688 |
| **Q8N2M8** | CLASRP | 0.002067078 | 3.197638896 | 0.124990469 | 0.904324415 | 0.980210603 | -7.168718226 |
| **P32321-2** | DCTD | 0.043508807 | 3.208803272 | 0.412886265 | 0.693024037 | 0.909991705 | -7.169258559 |
| **Q7Z3J2-1** | C16orf62 | -0.000680446 | 3.188644332 | -0.039920591 | 0.969359837 | 0.992469575 | -7.169473693 |
| **Q96EB6** | SIRT1 | -0.006687107 | 3.199292708 | -0.433001954 | 0.679075826 | 0.903893384 | -7.169542647 |
| **Q9H4A5** | GOLPH3L | -0.003542297 | 3.153055552 | -0.110064484 | 0.915693003 | 0.98295196 | -7.169582385 |
| **Q8NBP0-1** | TTC13 | -0.001894632 | 3.202288419 | -0.085543005 | 0.934415562 | 0.986793807 | -7.1698233 |
| **Q9H7E2-3** | TDRD3 | -0.013707808 | 3.194157393 | -0.826645132 | 0.437850348 | 0.786862931 | -7.170510555 |
| **Q9NTI5** | PDS5B | 0.030060034 | 3.178406514 | 1.356042354 | 0.220494227 | 0.625865114 | -7.171146758 |
| **Q86VN1** | VPS36 | -0.0117077 | 3.180388871 | -0.561227614 | 0.593532558 | 0.869935985 | -7.171419379 |
| **Q93100-1** | PHKB | -0.01199992 | 3.185964099 | -0.690569777 | 0.513832019 | 0.83148099 | -7.171439163 |
| **Q9BTX1** | NDC1; TMEM48 | 0.00467235 | 3.206300775 | 0.071114637 | 0.945453876 | 0.98904859 | -7.172082836 |
| **Q9UKL0** | RCOR1 | 0.005078082 | 3.184655253 | 0.167914798 | 0.871776338 | 0.971931713 | -7.172152008 |
| **Q7L5N1** | COPS6 | 0.022293206 | 3.192398409 | 0.819513877 | 0.441624911 | 0.789146094 | -7.172289377 |
| **Q9UBK7-2** | RABL2A | -0.000874617 | 3.1929271 | -0.046716226 | 0.964148058 | 0.991477649 | -7.172949563 |
| **Q9ULR0-1** | ISY1-RAB43; RAB4 | -0.038531195 | 3.166683731 | -0.531432447 | 0.612861508 | 0.878766193 | -7.173090996 |
| **Q5VIR6-4** | VPS53 | -0.010613153 | 3.190033952 | -0.628499745 | 0.55120307 | 0.847553395 | -7.17312374 |
| **Q8IXB1-1** | DNAJC10 | 0.001714908 | 3.202196659 | 0.06349459 | 0.951288978 | 0.989472196 | -7.173527898 |
| **Q9HB40** | SCPEP1 | 0.007877593 | 3.182839454 | 0.346136264 | 0.740218799 | 0.931545879 | -7.173562023 |
| **Q9UNI6** | DUSP12 | 0.003546904 | 3.152829684 | 0.122277911 | 0.906388743 | 0.98075504 | -7.173589429 |
| **Q96MG7** | NDNL2; NSMCE3 | -0.013267803 | 3.211800864 | -0.321120451 | 0.758239147 | 0.9384213 | -7.174026144 |
| **P53804-1** | TTC3 | 0.007213352 | 3.210478332 | 0.097253036 | 0.925468312 | 0.985476218 | -7.174464396 |
| **P53365** | ARFIP2 | -0.006818303 | 3.177420601 | -0.278314989 | 0.789451582 | 0.947865984 | -7.175433041 |
| **O15511-1** | ARPC5 | -0.030056351 | 3.171216868 | -0.932427992 | 0.384584514 | 0.754667212 | -7.175845485 |

| | | | | | | | |
|---|---|---|---|---|---|---|---|
| **O75448** | MED24 | 0.018839593 | 3.171309699 | 0.587392137 | 0.576849285 | 0.860993344 | -7.177341975 |
| **P15927-3** | RPA2 | 0.015660859 | 3.180024953 | 0.575636452 | 0.584310974 | 0.8653224 | -7.177382847 |
| **P52594-4** | AGFG1 | -0.002863021 | 3.189232711 | -0.157710216 | 0.879492269 | 0.97338977 | -7.177722278 |
| **O43933** | PEX1 | -0.014445388 | 3.199182062 | -0.657583429 | 0.533486458 | 0.840132297 | -7.178956715 |
| **Q86U42-1** | PABPN1 | -0.002033132 | 3.195513615 | -0.07807658 | 0.940125899 | 0.988002681 | -7.179712761 |
| **P23634-1** | ATP2B4 | -0.026550745 | 3.169114084 | -0.568978133 | 0.588561941 | 0.867233795 | -7.179743463 |
| **Q05519** | SRSF11 | -0.024217169 | 3.171848665 | -0.827113983 | 0.437602998 | 0.786862931 | -7.180157876 |
| **P52888** | THOP1 | -0.025549388 | 3.172801231 | -0.979739096 | 0.362410854 | 0.738054037 | -7.180376428 |
| **Q9H0E3-3** | SAP130 | 0.002723601 | 3.185192115 | 0.091489854 | 0.92987039 | 0.986707862 | -7.180861139 |
| **O00425** | IGF2BP3 | -0.023153091 | 3.15779088 | -1.021067352 | 0.343867618 | 0.725100733 | -7.180943533 |
| **Q12797** | ASPH | 0.013114071 | 3.18590307 | 0.780852256 | 0.462490637 | 0.802800142 | -7.181034082 |
| **Q6P1N9-1** | TATDN1 | 0.021126236 | 3.165590505 | 0.803934849 | 0.449951342 | 0.79513757 | -7.18138921 |
| **O75150** | RNF40 | -0.02783334 | 3.173736393 | -1.094090258 | 0.312959131 | 0.701003749 | -7.182068223 |
| **Q9UQN3-1** | CHMP2B | 0.004998391 | 3.198163746 | 0.163788394 | 0.874894633 | 0.972759481 | -7.182163552 |
| **Q9Y4K4** | MAP4K5 | -0.008973449 | 3.169340065 | -0.1408032 | 0.892307104 | 0.978206823 | -7.182316381 |
| **Q92636** | NSMAF | 0.013525543 | 3.182821391 | 0.550858787 | 0.600219648 | 0.872278234 | -7.183957668 |
| **Q9C0E2** | XPO4 | -0.020724849 | 3.179561459 | -0.811279163 | 0.446012324 | 0.792020296 | -7.184743584 |
| **O00244** | ATOX1 | -0.022027936 | 3.17067478 | -0.384355557 | 0.713030572 | 0.922407393 | -7.187170633 |
| **Q9H5K3** | POMK; SGK196 | 0.003005748 | 3.204611288 | 0.093736072 | 0.928154339 | 0.985985589 | -7.187232339 |
| **Q8N0Z6** | TTC5 | -0.002467899 | 3.174362527 | -0.092768591 | 0.928893418 | 0.986243245 | -7.187325322 |
| **O75153** | CLUH; KIAA0664 | -0.020245984 | 3.193660712 | -1.169506801 | 0.28346499 | 0.679787821 | -7.18751921 |
| **Q9UKL3** | CASP8AP2 | -0.000589586 | 3.205349469 | -0.035033044 | 0.97310927 | 0.993108947 | -7.187528389 |
| **Q96S19** | C16orf13; METTL26 | 0.001576681 | 3.167599407 | 0.061889338 | 0.952518638 | 0.989472196 | -7.189689868 |
| **Q14145** | KEAP1 | -0.021582282 | 3.1797229 | -0.85079494 | 0.425240038 | 0.778986255 | -7.190213933 |
| **Q16850-1** | CYP51A1 | -0.011859429 | 3.085412199 | -0.546835533 | 0.602825738 | 0.872497827 | -7.190851971 |
| **Q8N9V3** | WDSUB1 | -0.012200966 | 3.219567289 | -0.214090846 | 0.837064181 | 0.960912428 | -7.191747482 |
| **Q7Z2T5-1** | TRMT1L | 0.017664304 | 3.171808498 | 0.831229767 | 0.43543594 | 0.784298014 | -7.191853402 |
| **P43007-1** | SLC1A4 | -0.028388641 | 3.172304249 | -0.558459891 | 0.595313365 | 0.871378088 | -7.192353434 |
| **P53041** | PPP5C | -0.01882904 | 3.175421586 | -1.069750629 | 0.323000791 | 0.710272801 | -7.192602048 |
| **Q14191** | WRN | -0.012876691 | 3.167023297 | -0.997538149 | 0.354330746 | 0.734057823 | -7.192655951 |
| **Q8TEQ6** | GEMIN5 | -0.020891905 | 3.195468917 | -1.390218244 | 0.21039861 | 0.615470161 | -7.193040448 |
| **Q9Y5B6** | GCFC1; PAXBP1 | -0.01752026 | 3.192654795 | -0.6490914 | 0.538622246 | 0.843540088 | -7.193753742 |
| **Q07889** | SOS 1 | -0.004167239 | 3.193137361 | -0.183799379 | 0.859796159 | 0.968855822 | -7.194683655 |
| **P62942** | FKBP1A | 0.040466455 | 3.197685792 | 0.870565988 | 0.41511433 | 0.77203561 | -7.195227478 |
| **Q9Y3A3** | MOB4 | -0.005600146 | 3.16778002 | -0.163744955 | 0.874927473 | 0.972759481 | -7.195278084 |
| **Q9UIV1-1** | CNOT7 | -0.00586266 | 3.151510284 | -0.048827963 | 0.962528879 | 0.991283985 | -7.195426406 |
| **Q8NBT2** | SPC24 | -0.003823436 | 3.187606682 | -0.162138225 | 0.876142337 | 0.972759481 | -7.195671171 |
| **P23258** | TUBG1 | 0.013847374 | 3.193775303 | 0.708425129 | 0.503391026 | 0.827688992 | -7.196300536 |
| **Q8NEM7-3** | FAM48A; SUPT20H | -0.002735948 | 3.179863558 | -0.090172494 | 0.930877008 | 0.986707862 | -7.196341539 |

| | | | | | | | |
|---|---|---|---|---|---|---|---|
| **Q96M89-1** | CCDC138 | -0.001031923 | 3.162383542 | -0.044028371 | 0.966209243 | 0.992001409 | -7.196508823 |
| **Q9NWZ3-1** | IRAK4 | -0.01310027 | 3.184633877 | -0.608851906 | 0.563373621 | 0.852578515 | -7.196871297 |
| **Q8WVK2** | SNRNP27 | 0.004214585 | 3.187960161 | 0.144959895 | 0.88915308 | 0.976409291 | -7.197026715 |
| **O75132** | ZBED4 | 0.000741321 | 3.203509104 | 0.033106953 | 0.974587061 | 0.99316605 | -7.197092085 |
| **Q99816** | TSG101 | -0.008387435 | 3.202200251 | -0.429233944 | 0.681678377 | 0.904168215 | -7.197730423 |
| **Q9NR50-1** | EIF2B3 | 0.036921538 | 3.192708703 | 0.903635242 | 0.398576959 | 0.762880385 | -7.197905755 |
| **P68366** | TUBA4A | 7.68E-05 | 3.153611866 | 0.002795379 | 0.997853815 | 0.998482186 | -7.198026468 |
| **Q8WUB8** | PHF10 | -0.00475388 | 3.200535257 | -0.209226263 | 0.840704 | 0.961788885 | -7.198080698 |
| **P17706-1** | PTPN2 | -0.004068503 | 3.185880576 | -0.146827349 | 0.887736799 | 0.975933149 | -7.198816347 |
| **Q8ND82** | ZNF280C | -0.012414993 | 3.184034647 | -0.646464813 | 0.540216987 | 0.843540088 | -7.199091849 |
| **Q7L8W6** | ATPBD4; DPH6 | -0.0059325 | 3.17505596 | -0.178237559 | 0.863986499 | 0.969738239 | -7.19912388 |
| **Q9H1A4** | ANAPC1 | -0.01396336 | 3.183350785 | -1.022538787 | 0.343221514 | 0.724852454 | -7.199539639 |
| **P11717** | IGF2R | 0.012052715 | 3.160936222 | 1.18416567 | 0.278011168 | 0.676093886 | -7.199785644 |
| **Q567U6** | CCDC93 | 0.009207077 | 3.175630962 | 0.434520031 | 0.678028638 | 0.903662615 | -7.20110063 |
| **Q14156-1** | EFR3A | 0.01261316 | 3.159383243 | 0.490654676 | 0.639866334 | 0.88969864 | -7.201289587 |
| **Q9BZL6** | PRKD2 | 0.008868471 | 3.188459164 | 0.694715187 | 0.511395506 | 0.83054728 | -7.201589079 |
| **Q6IQ32** | ADNP2 | 0.000265772 | 3.191847079 | 0.014379093 | 0.988960701 | 0.996219143 | -7.201744522 |
| **P84098** | RPL19 | -0.051702973 | 3.178401258 | -0.920684844 | 0.390245729 | 0.758811139 | -7.203443356 |
| **Q6UWP7-1** | LCLAT1 | 0.003416975 | 3.188099686 | 0.110711854 | 0.915199457 | 0.9828683 | -7.204901463 |
| **Q8TBA6** | GOLGA5 | -0.030676362 | 3.174318293 | -0.6450706 | 0.541064683 | 0.843540088 | -7.205438143 |
| **Q9H7L9** | SUDS3 | 0.002547485 | 3.152486146 | 0.061581745 | 0.952754278 | 0.989472196 | -7.205553325 |
| **P42338** | PIK3CB | -0.007360098 | 3.187243061 | -0.319899229 | 0.759123184 | 0.9384213 | -7.205791912 |
| **Q9Y657** | SPIN1 | 0.01039498 | 3.178166633 | 0.434173923 | 0.678267319 | 0.903662615 | -7.207181766 |
| **Q15382** | RHEB | -0.008586154 | 3.187825942 | -0.355311783 | 0.733652697 | 0.928755685 | -7.207944815 |
| **Q9Y3L3-1** | SH3BP1 | -0.007425303 | 3.17917489 | -0.306048149 | 0.769177083 | 0.941649022 | -7.208025473 |
| **P11802** | CDK4 | 0.00099845 | 3.271697728 | 0.030767099 | 0.976382457 | 0.99316605 | -7.208051912 |
| **P36639** | NUDT1 | -0.019974632 | 3.197234851 | -0.785261371 | 0.460076807 | 0.800900577 | -7.20869846 |
| **O95865** | DDAH2 | -0.01210716 | 3.171832932 | -0.682574044 | 0.518552806 | 0.833791899 | -7.208714545 |
| **P23458** | JAK1 | -0.010567648 | 3.165679337 | -0.662851532 | 0.53031597 | 0.839849063 | -7.209310554 |
| **Q9GZP4** | PITHD1 | -0.004793376 | 3.187916991 | -0.101172346 | 0.922476249 | 0.984164603 | -7.20961614 |
| **Q9H6U6-8** | BCAS3 | 0.017724602 | 3.189836964 | 0.470064801 | 0.653734983 | 0.896119351 | -7.210039652 |
| **O15226-2** | NKRF | -0.010712006 | 3.183674951 | -0.887696947 | 0.406485092 | 0.766437228 | -7.21145482 |
| **Q6F5E8** | RLTPR; CARMIL2 | 0.011439972 | 3.186978881 | 0.715898436 | 0.499062636 | 0.824970816 | -7.211839224 |
| **Q16563-2** | SYPL1 | -0.045671493 | 3.193430586 | -0.373334579 | 0.720826303 | 0.925644897 | -7.212524654 |
| **Q92890-1** | UFD1L; UFD1 | 0.008136341 | 3.178454988 | 0.223355962 | 0.830143848 | 0.959902907 | -7.212982123 |
| **P98179** | RBM3 | 0.017268463 | 3.173452207 | 0.496292016 | 0.636096072 | 0.888499173 | -7.213709024 |
| **Q6KC79** | NIPBL | 0.011482286 | 3.172639834 | 1.285607661 | 0.242668511 | 0.649360189 | -7.213783011 |
| **Q9GZN8-2** | C20orf27 | -0.000617384 | 3.18686747 | -0.030235509 | 0.976790373 | 0.99316605 | -7.214476967 |
| **P14735** | IDE | -0.017488486 | 3.156615108 | -1.162195392 | 0.286218729 | 0.681861411 | -7.214656311 |

| | | | | | | | |
|---|---|---|---|---|---|---|---|
| **O00442** | RTCA | 0.005672656 | 3.165199923 | 0.118166752 | 0.90951894 | 0.981544143 | -7.216388474 |
| **Q4G0F5** | VPS26B | 0.00759162 | 3.189194907 | 0.246338634 | 0.81304963 | 0.956167049 | -7.217130875 |
| **Q13416** | ORC2 | -0.001083625 | 3.186766328 | -0.043081706 | 0.966935262 | 0.992288899 | -7.217169905 |
| **Q9H0C8** | ILKAP | 0.012207821 | 3.173467801 | 0.662528399 | 0.530510096 | 0.839849063 | -7.217223718 |
| **O15117-3** | FYB; FYB1 | 0.013901214 | 3.181970453 | 0.605797727 | 0.565279919 | 0.852857758 | -7.217837997 |
| **Q9H832** | UBE2Z | 0.006095071 | 3.190147827 | 0.385452404 | 0.712256733 | 0.921791782 | -7.218775497 |
| **Q9UBS8-1** | RNF14 | 0.013454471 | 3.192480578 | 0.302433133 | 0.771809144 | 0.942488971 | -7.219056696 |
| **Q6P1N0** | CC2D1A | -0.012370205 | 3.180302102 | -0.497030254 | 0.635603203 | 0.888277483 | -7.21961809 |
| **Q9H081** | MIS12 | -0.000439693 | 3.192263548 | -0.009080499 | 0.993028438 | 0.99756865 | -7.219686302 |
| **O75695** | RP2 | 0.011328709 | 3.148317489 | 0.286675736 | 0.783319804 | 0.946104568 | -7.220086583 |
| **Q9Y5B0** | CTDP1 | 0.006683831 | 3.193896368 | 0.449845409 | 0.667500728 | 0.90001396 | -7.220089634 |
| **O60825-1** | PFKFB2 | 0.000316621 | 3.171737616 | 0.015220107 | 0.988315084 | 0.996184213 | -7.220196936 |
| **O75955** | FLOT1 | -0.006490869 | 3.175745125 | -0.305792279 | 0.769363271 | 0.941649022 | -7.220354358 |
| **P49761** | CLK3 | 0.004920356 | 3.193283154 | 0.170489509 | 0.869831901 | 0.971575207 | -7.220687944 |
| **Q00169** | PITPNA | 0.018940369 | 3.199630019 | 0.580372127 | 0.581298368 | 0.863901148 | -7.220688016 |
| **Q92673** | SORL1 | -0.000377723 | 3.19145719 | -0.015243089 | 0.988297441 | 0.996184213 | -7.221323744 |
| **O60934** | NBN | -0.012072319 | 3.187386708 | -1.011469791 | 0.348105604 | 0.730105045 | -7.221327977 |
| **Q9UPW5** | AGTPBP1 | 0.007145529 | 3.18034574 | 0.412049329 | 0.693607239 | 0.910349515 | -7.221813695 |
| **Q13247** | SRSF6 | -0.010155172 | 3.184373617 | -0.422843078 | 0.686103325 | 0.906701749 | -7.222350516 |
| **Q9NWS0-1** | PIH1D1 | -0.001580987 | 3.169173443 | -0.0486531 | 0.962662948 | 0.991283985 | -7.222519635 |
| **Q9UQR1** | ZNF148 | -0.018760194 | 3.200301519 | -0.522931135 | 0.618439675 | 0.882137023 | -7.222914076 |
| **Q5XPI4-2** | RNF123 | -0.001946471 | 3.179610985 | -0.095223834 | 0.927017957 | 0.985631941 | -7.22321185 |
| **Q9H6E4** | CCDC134 | -0.000741013 | 3.163732253 | -0.031965696 | 0.975462741 | 0.99316605 | -7.223287613 |
| **O95983-1** | MBD3 | 0.008379967 | 3.1846288 | 0.37823896 | 0.717352632 | 0.924679646 | -7.223786593 |
| **Q6NXE6-1** | ARMC6 | -0.000102861 | 3.176926147 | -0.003291258 | 0.997473099 | 0.998480498 | -7.22458064 |
| **O95251-1** | KAT7 | 0.017929244 | 3.171972168 | 0.754754906 | 0.476957526 | 0.810572738 | -7.225676589 |
| **P06746** | POLB | 0.007035072 | 3.164931352 | 0.438654769 | 0.675180381 | 0.90314827 | -7.226100682 |
| **Q9NUW8** | TDP1 | 0.028708642 | 3.194526673 | 0.938996805 | 0.381445146 | 0.752752531 | -7.226515618 |
| **Q9NPD3** | EXOSC4 | 0.006881013 | 3.181639207 | 0.167743047 | 0.87190608 | 0.971931713 | -7.227551238 |
| **Q9BTV4** | TMEM43 | 0.011786688 | 3.186623932 | 0.734483648 | 0.488405723 | 0.817162784 | -7.228095693 |
| **Q8N108-12** | MIER1 | -0.037298214 | 3.170925411 | -0.744373493 | 0.482797476 | 0.812781066 | -7.228248307 |
| **Q9H6R7** | C2orf44; WDCP | -0.00522769 | 3.203140864 | -0.32962244 | 0.752095585 | 0.93570301 | -7.228641548 |
| **P40227-1** | CCT6A | 0.012509405 | 3.192930529 | 1.257407975 | 0.252080484 | 0.656218606 | -7.228643265 |
| **P50454** | SERPINH1 | -0.015010588 | 3.202759594 | -0.542782727 | 0.605457377 | 0.874292777 | -7.230236178 |
| **O75475-1** | PSIP1 | -0.014831426 | 3.1717067 | -1.063522245 | 0.325612046 | 0.710703247 | -7.231285627 |
| **Q9BUI4** | POLR3C; LOC1010 | 0.01632525 | 3.200472274 | 0.572712438 | 0.586175602 | 0.8653224 | -7.231383909 |
| **Q9BYM8** | RBCK1 | -0.003600443 | 3.193334124 | -0.191715248 | 0.853840763 | 0.967450778 | -7.231988736 |
| **Q15311** | RALBP1 | 0.007426342 | 3.173783174 | 0.259816597 | 0.803075616 | 0.953721684 | -7.232347767 |
| **Q8NC56-1** | LEMD2 | 0.015276238 | 3.187218836 | 0.330602005 | 0.751388994 | 0.935635612 | -7.232496884 |

| | | | | | | | |
|---|---|---|---|---|---|---|---|
| **P10606** | COX5B | 0.002060255 | 3.2106015 | 0.013339436 | 0.989758823 | 0.996253288 | -7.233224888 |
| **P14324** | FDPS | -0.017576165 | 3.158236006 | -0.663911432 | 0.529679535 | 0.839849063 | -7.233613661 |
| **O60749** | SNX2 | 0.012614215 | 3.177484345 | 0.832105572 | 0.434975804 | 0.784298014 | -7.233827824 |
| **Q9Y4Z0** | LSM 4.00 | 0.008276603 | 3.157709293 | 0.512935184 | 0.625033759 | 0.885387512 | -7.234433385 |
| **Q9C0I1** | MTMR12 | 0.003651054 | 3.18831315 | 0.170554863 | 0.869782557 | 0.971575207 | -7.235045182 |
| **Q9Y3C6** | PPIL1 | 0.03015142 | 3.176311859 | 0.932276328 | 0.384657229 | 0.754667212 | -7.235350137 |
| **Q96EK7** | FAM120B | 0.005474272 | 3.207568353 | 0.28555617 | 0.784139934 | 0.946296623 | -7.235892434 |
| **O75367-1** | H2AFY | -0.032487936 | 3.16338246 | -1.229245355 | 0.261794241 | 0.665372759 | -7.236058796 |
| **Q9NWV4** | C1orf123 | 0.033081717 | 3.194847604 | 0.603510104 | 0.566710289 | 0.853395233 | -7.236147812 |
| **Q92990** | GLMN | -0.010418465 | 3.175369652 | -0.420847217 | 0.687487995 | 0.907431059 | -7.236234566 |
| **Q969S3** | ZNF622 | -0.007199756 | 3.190284648 | -0.210332351 | 0.839876017 | 0.961392936 | -7.237618463 |
| **Q8IWR0** | ZC3H7A | -0.040104156 | 3.167908559 | -0.895941374 | 0.402379962 | 0.765842327 | -7.237928044 |
| **P28749-1** | RBL1 | 0.00116363 | 3.176832186 | 0.066283975 | 0.949152589 | 0.989472196 | -7.238580163 |
| **O95396** | MOCS3 | 0.010569273 | 3.168757399 | 0.513879741 | 0.624409041 | 0.885387512 | -7.239470375 |
| **Q15283-1** | RASA2 | 0.024129295 | 3.178873864 | 0.960483776 | 0.371312996 | 0.744717822 | -7.240362537 |
| **P49959-3** | MRE11A; MRE11 | 0.01937422 | 3.163752117 | 0.810906769 | 0.446211462 | 0.792020296 | -7.241142698 |
| **Q9H0E9-2** | BRD8 | -0.005765204 | 3.138697567 | -0.33060356 | 0.751387873 | 0.935635612 | -7.241304137 |
| **Q7Z6M1-1** | RABEPK | 0.001853397 | 3.182353859 | 0.076062863 | 0.941666667 | 0.988377491 | -7.24131296 |
| **Q9H5Z1** | DHX35 | -0.004884708 | 3.179284089 | -0.230691471 | 0.824676377 | 0.959077751 | -7.241553964 |
| **Q96LB3-1** | IFT74 | -0.014288747 | 3.165476411 | -0.701516982 | 0.507413988 | 0.82913633 | -7.241610907 |
| **Q68EM7** | ARHGAP17 | -0.00896591 | 3.152482421 | -0.506267035 | 0.629453518 | 0.886324146 | -7.241921493 |
| **P41229** | KDM5C | -0.002026601 | 3.170964871 | -0.061361941 | 0.952922668 | 0.989472196 | -7.244794079 |
| **O95619** | YEATS4 | -0.013216633 | 3.195229234 | -0.354795697 | 0.734021377 | 0.928926384 | -7.245720385 |
| **Q9NWV8** | BABAM1 | 0.012985598 | 3.18815859 | 0.212189644 | 0.838486202 | 0.961294787 | -7.245843542 |
| **Q8N8A2** | ANKRD44 | 0.013859917 | 3.190404269 | 0.772948947 | 0.466839388 | 0.804563762 | -7.24590038 |
| **O00422** | SAP18 | 0.043892188 | 3.184919237 | 0.744301255 | 0.482838281 | 0.812781066 | -7.247167109 |
| **P55039** | DRG2 | -0.028898351 | 3.204814484 | -0.80840407 | 0.447551421 | 0.792642897 | -7.247274746 |
| **Q15785** | TOMM34 | 0.010249321 | 3.206537815 | 0.244120049 | 0.814695124 | 0.95620584 | -7.247754391 |
| **P55011** | SLC12A2 | -0.009669881 | 3.210760286 | -0.416423229 | 0.690561885 | 0.909522034 | -7.248687293 |
| **P14174** | MIF | 0.02317371 | 3.17067274 | 0.450936109 | 0.666754524 | 0.90001396 | -7.24892497 |
| **Q9UMX0-2** | UBQLN1 | 0.007460428 | 3.178574638 | 0.303380719 | 0.771118897 | 0.942318194 | -7.249416596 |
| **Q96IJ6** | GMPPA | 0.007601567 | 3.197179 | 0.346005941 | 0.740312232 | 0.931545879 | -7.250454132 |
| **P16615** | ATP2A2 | -0.021897164 | 3.189869798 | -0.889875806 | 0.40539716 | 0.766336431 | -7.250679683 |
| **Q8NHV4-3** | NEDD1 | -0.01919664 | 3.177843659 | -0.78787882 | 0.458648008 | 0.799639769 | -7.250929729 |
| **P56192** | MARS | -0.012340972 | 3.180197578 | -1.12349522 | 0.301170455 | 0.691344519 | -7.251399937 |
| **Q969N2** | PIGT | -0.008155825 | 3.196559611 | -0.466583689 | 0.656094757 | 0.896497734 | -7.25155402 |
| **Q8NI08** | NCOA7 | 0.003426853 | 3.18842055 | 0.185105125 | 0.858813102 | 0.968855822 | -7.251788254 |
| **Q06323-1** | PSME1 | -0.026075723 | 3.180021551 | -1.395856494 | 0.208773506 | 0.614302382 | -7.251880951 |
| **P61964** | WDR5 | -0.015772942 | 3.19208795 | -0.566458736 | 0.590175069 | 0.868764508 | -7.252110725 |

| | | | | | | | |
|---|---|---|---|---|---|---|---|
| **Q8WWH5** | TRUB1 | -0.002447163 | 3.205750692 | -0.086893742 | 0.933382954 | 0.986793807 | -7.252806843 |
| **O94763-1** | URI1 | -0.003207958 | 3.188041571 | -0.165403897 | 0.873673517 | 0.97237605 | -7.253664977 |
| **Q9NPF4** | OSGEP | 0.005360993 | 3.179271109 | 0.181533537 | 0.861502685 | 0.968855822 | -7.253739481 |
| **O75569-1** | PRKRA | 0.017265974 | 3.169083453 | 0.506887182 | 0.629041772 | 0.886280701 | -7.253878463 |
| **P49754** | VPS41 | 0.000624239 | 3.197871784 | 0.024548098 | 0.98115506 | 0.994549241 | -7.253933094 |
| **Q9NQX3-2** | GPHN | -0.000706912 | 3.151794558 | -0.016406673 | 0.987404214 | 0.99582925 | -7.255830116 |
| **Q96IZ7-1** | RSRC1 | 0.002584375 | 3.183924647 | 0.107927461 | 0.917322526 | 0.983235402 | -7.256043735 |
| **Q92547** | TOPBP1 | -0.014591929 | 3.177674178 | -1.052974129 | 0.330073269 | 0.715729292 | -7.257484218 |
| **Q5VV42** | CDKAL1 | -0.012272219 | 3.193463099 | -0.287605919 | 0.782638634 | 0.94563233 | -7.258476999 |
| **P00966** | ASS1 | 0.015575618 | 3.198867848 | 0.537081676 | 0.609170109 | 0.876646363 | -7.258587006 |
| **Q9Y2G3** | ATP11B | 0.009356516 | 3.194970454 | 0.402758149 | 0.700096726 | 0.913245694 | -7.25885081 |
| **O60341** | KDM1A | -0.011535096 | 3.189575071 | -1.136954243 | 0.295898476 | 0.688880164 | -7.261576549 |
| **Q8WUA4-1** | GTF3C2 | -0.008934716 | 3.177031259 | -0.460064388 | 0.660525565 | 0.896934512 | -7.262239521 |
| **Q9NQ88** | C12orf5; TIGAR | -0.00087995 | 3.187487378 | -0.016974413 | 0.986968393 | 0.99582925 | -7.263280081 |
| **O75223-1** | GGCT | 0.034951681 | 3.194636569 | 0.565417178 | 0.590842703 | 0.868764508 | -7.26328644 |
| **P00387-3** | CYB5R3 | 0.006446663 | 3.178561069 | 0.232804033 | 0.823103742 | 0.959068023 | -7.26377104 |
| **P14618** | PKM | -0.015628736 | 3.183160455 | -0.364145971 | 0.727353713 | 0.92710315 | -7.263817205 |
| **Q9UKX7** | NUP50 | 0.017681064 | 3.180998242 | 1.035052115 | 0.337765913 | 0.721772506 | -7.264249847 |
| **Q9NRL3-3** | STRN4 | -0.011336304 | 3.171493919 | -0.463156939 | 0.658421864 | 0.896852471 | -7.264389105 |
| **Q8IWV7-1** | UBR1 | -0.015315856 | 3.173534598 | -0.80231255 | 0.450824743 | 0.79513757 | -7.265082317 |
| **Q8TCJ2** | STT3B | 0.008468209 | 3.219774573 | 0.178370766 | 0.863886084 | 0.969738239 | -7.265346076 |
| **O75113** | N4BP1 | -0.007697214 | 3.1929628 | -0.404292144 | 0.699023395 | 0.913245694 | -7.265838065 |
| **Q9HBF4-1** | ZFYVE1 | 0.001306251 | 3.168919115 | 0.058901398 | 0.954807844 | 0.989472196 | -7.26597706 |
| **Q6UN15-1** | FIP1L1 | -0.01683876 | 3.180164391 | -0.520245936 | 0.620207314 | 0.882940648 | -7.266188957 |
| **Q14160** | SCRIB | -0.021115623 | 3.187225846 | -0.918505718 | 0.391303165 | 0.759806823 | -7.267107084 |
| **Q8WXA9-2** | SREK1 | 0.001450992 | 3.185799261 | 0.041091836 | 0.968461449 | 0.992306703 | -7.267710353 |
| **P11274-1** | BCR | -0.005419721 | 3.187115156 | -0.201088686 | 0.846802222 | 0.963629033 | -7.268427176 |
| **O60870** | KIN | -0.009292354 | 3.168599789 | -0.624659466 | 0.553569173 | 0.848478354 | -7.270215029 |
| **P41743** | PRKCI | -0.012175007 | 3.199580767 | -0.558613893 | 0.595214197 | 0.871378088 | -7.270232913 |
| **Q96HA7** | TONSL | 0.009893877 | 3.179436784 | 0.443864308 | 0.671599979 | 0.901175787 | -7.270658316 |
| **Q00653-1** | NFKB2 | -0.001097304 | 3.190223565 | -0.055734713 | 0.957234501 | 0.989500907 | -7.271302398 |
| **Q92830-1** | KAT2A | -0.002657273 | 3.203951325 | -0.140046744 | 0.89288132 | 0.978276697 | -7.272096379 |
| **P50749** | RASSF2 | -0.004530352 | 3.174201343 | -0.153895422 | 0.882380446 | 0.97451158 | -7.272499152 |
| **Q9UKK9** | NUDT5 | -0.034514724 | 3.170556131 | -0.840304255 | 0.430685286 | 0.782692325 | -7.273310154 |
| **Q9UNZ2-5** | NSFL1C | 0.027277979 | 3.162966554 | 0.878738214 | 0.410981083 | 0.769906516 | -7.273328153 |
| **Q9UKG1** | APPL1 | 0.021628239 | 3.166435496 | 0.962272504 | 0.370478964 | 0.744424726 | -7.273475487 |
| **O75663** | TIPRL | -0.026939777 | 3.191878711 | -0.463084138 | 0.658471348 | 0.896852471 | -7.273795574 |
| **Q9BZ29-4** | DOCK9 | 0.008194791 | 3.189236359 | 0.585855692 | 0.577821329 | 0.861313941 | -7.274402726 |
| **Q9Y6E2** | BZW2 | -0.035677471 | 3.205785229 | -0.862636462 | 0.419153984 | 0.775722898 | -7.274657047 |

| | | | | | | | |
|---|---|---|---|---|---|---|---|
| **P05423** | POLR3D | -0.001281606 | 3.202190911 | -0.048099417 | 0.963087472 | 0.991283985 | -7.274672029 |
| **O15371** | EIF3D | -0.013991449 | 3.197402834 | -1.098841094 | 0.31102919 | 0.699045803 | -7.274825162 |
| **Q12959-7** | DLG1 | 0.005465115 | 3.199252299 | 0.18229076 | 0.86093229 | 0.968855822 | -7.27523479 |
| **Q96ST3** | SIN3A | 0.020366622 | 3.174687665 | 1.181063658 | 0.279157812 | 0.676725357 | -7.276089528 |
| **Q9NXV6** | CDKN2AIP | 0.013924286 | 3.165070838 | 0.802591254 | 0.450674611 | 0.79513757 | -7.276318103 |
| **Q00013-1** | MPP1 | -0.030406417 | 3.194445365 | -0.799583841 | 0.452296497 | 0.795197094 | -7.276696791 |
| **Q9NZC7** | WWOX | -0.001063102 | 3.151720019 | -0.028067799 | 0.978453844 | 0.993585863 | -7.276932808 |
| **O75330-3** | HMMR | 0.013715128 | 3.063510615 | 0.359972839 | 0.73032643 | 0.927974736 | -7.277562635 |
| **P33991** | MCM4 | 0.026740503 | 3.184943447 | 1.458874664 | 0.191366853 | 0.59600535 | -7.277803923 |
| **Q6ZNJ1-1** | NBEAL2 | 0.011819227 | 3.188692988 | 0.770470718 | 0.468208825 | 0.805700479 | -7.277946817 |
| **Q9BRD0** | BUD13 | -0.00836986 | 3.194456293 | -0.404440772 | 0.69891944 | 0.913245694 | -7.278074502 |
| **Q9BRG1** | VPS25 | 0.017240946 | 3.193795734 | 0.325690175 | 0.754934651 | 0.937112609 | -7.27851865 |
| **Q5VU43-2** | PDE4DIP; LOC6535 | 0.008318466 | 3.187926115 | 0.211193648 | 0.839231429 | 0.961364456 | -7.27887301 |
| **O00231** | PSMD11 | 0.033510729 | 3.18489613 | 1.293249586 | 0.24017134 | 0.645958461 | -7.279099712 |
| **Q9H410** | DSN1 | 0.008305643 | 3.173895784 | 0.409354988 | 0.695486269 | 0.911821519 | -7.27934078 |
| **Q9ULV4-3** | CORO1C | -0.001439163 | 3.185310764 | -0.027495894 | 0.978892736 | 0.993650542 | -7.279651386 |
| **Q13586** | STIM1 | -0.009748205 | 3.194741086 | -0.471392641 | 0.652836001 | 0.895616288 | -7.280035585 |
| **Q9Y6W5** | WASF2 | 0.008308405 | 3.162692134 | 0.445139451 | 0.67072501 | 0.901175787 | -7.28029736 |
| **P55327-3** | TPD52 | 0.003797419 | 3.199622111 | 0.156238619 | 0.88060618 | 0.974002142 | -7.281128547 |
| **Q9NRG9-1** | AAAS | -0.003308332 | 3.171813843 | -0.090176745 | 0.930873759 | 0.986707862 | -7.281767711 |
| **Q15059** | BRD3 | 0.007237584 | 3.176066077 | 0.450695128 | 0.666919356 | 0.90001396 | -7.281848834 |
| **Q9NQG5** | RPRD1B | 0.003756478 | 3.171208266 | 0.157859707 | 0.879379131 | 0.97338977 | -7.283196914 |
| **P54709** | ATP1B3 | -0.02626706 | 3.20383517 | -0.407963305 | 0.696457742 | 0.912062672 | -7.283349974 |
| **Q9UBQ7** | GRHPR | 0.009262251 | 3.178800371 | 0.363196282 | 0.728029773 | 0.92710315 | -7.283457949 |
| **Q8NCL4** | GALNT6 | 0.011329736 | 3.188510557 | 0.477697749 | 0.648575872 | 0.893481222 | -7.283479862 |
| **Q13094** | LCP2 | -0.00730721 | 3.176545989 | -0.190222958 | 0.854962686 | 0.968031714 | -7.283598932 |
| **Q8WXI9** | GATAD2B | 0.019134495 | 3.152047011 | 0.778513396 | 0.463774643 | 0.802800142 | -7.283658705 |
| **Q15257-1** | PPP2R4; PTPA | 0.020118774 | 3.180707599 | 0.684335508 | 0.51751042 | 0.833280927 | -7.284018718 |
| **P20585** | MSH3 | -0.015643328 | 3.182427939 | -0.888272299 | 0.406197601 | 0.766336431 | -7.284579799 |
| **O00268** | TAF4 | 0.00639804 | 3.187494237 | 0.257397359 | 0.804863031 | 0.954249678 | -7.284932736 |
| **Q3MIT2** | PUS10 | -8.25E-05 | 3.182284486 | -0.003401246 | 0.997388655 | 0.998480498 | -7.285165518 |
| **Q9H270** | VPS11 | -0.008129368 | 3.169278245 | -0.30886887 | 0.76712564 | 0.940879515 | -7.285861888 |
| **P02751** | FN1 | -0.492407642 | 2.907278646 | -0.74666788 | 0.481502639 | 0.81235548 | -7.286816417 |
| **Q9UBB9-1** | TFIP11 | -0.013408406 | 3.194625785 | -1.014807868 | 0.346626944 | 0.728365794 | -7.287423131 |
| **P17858-1** | PFKL | 0.02013918 | 3.188294295 | 0.888128916 | 0.406269232 | 0.766336431 | -7.287802028 |
| **Q15262-4** | PTPRK | 0.009706194 | 3.170627723 | 0.431833231 | 0.679882551 | 0.903893384 | -7.287901536 |
| **Q96H20-1** | SNF8 | 0.014032671 | 3.199855392 | 0.375218743 | 0.719490934 | 0.925575691 | -7.28810965 |
| **Q9H8Y8-3** | GORASP2 | -0.005366204 | 3.192042506 | -0.159342227 | 0.878257283 | 0.973200128 | -7.288126721 |
| **Q99747** | NAPG | 0.018741068 | 3.185947353 | 0.531684354 | 0.612696641 | 0.878766193 | -7.288180986 |

| | | | | | | | |
|---|---|---|---|---|---|---|---|
| **Q96JN0-3** | LCOR | 0.012192123 | 3.206829809 | 0.318882105 | 0.759859774 | 0.938422437 | -7.288603933 |
| **Q8TDZ2-4** | MICAL1 | -0.000543581 | 3.199632334 | -0.019288348 | 0.985192178 | 0.995531556 | -7.288750942 |
| **Q93084-5** | ATP2A3 | 0.018398139 | 3.18533651 | 0.995661669 | 0.355175877 | 0.734480048 | -7.290601783 |
| **Q9Y485** | DMXL1 | -0.01283885 | 3.191465685 | -0.628855415 | 0.550984246 | 0.84738092 | -7.291254465 |
| **Q9BR76** | CORO1B | -9.77E-05 | 3.156917907 | -0.004715199 | 0.996379859 | 0.998480498 | -7.291408012 |
| **O75643-1** | SNRNP200 | -0.021925687 | 3.177177091 | -1.642225934 | 0.148038314 | 0.546231571 | -7.291480661 |
| **P48553** | TRAPPC10 | 0.003111798 | 3.168432154 | 0.14770339 | 0.88707256 | 0.975708125 | -7.291484247 |
| **O75691** | UTP20 | -0.038156365 | 3.184545359 | -1.244453176 | 0.256509454 | 0.659782078 | -7.291654177 |
| **O14929** | HAT1 | 0.011316143 | 3.170871776 | 0.502571256 | 0.6319103 | 0.887490806 | -7.291733935 |
| **Q92759** | GTF2H4 | -0.003504582 | 3.191581889 | -0.123300944 | 0.905610094 | 0.98075504 | -7.292100587 |
| **B0I1T2-1** | MYO1G | -0.016648311 | 3.17378002 | -1.10105331 | 0.310133855 | 0.69821861 | -7.292219325 |
| **Q2M2I8-1** | AAK1 | 0.004342738 | 3.183617696 | 0.247262221 | 0.812364921 | 0.956041963 | -7.292331071 |
| **Q8WY36** | BBX | -0.006320351 | 3.170591948 | -0.32125433 | 0.758142257 | 0.9384213 | -7.293557199 |
| **Q9H9A5-6** | CNOT10 | -0.013441135 | 3.177103176 | -0.643835644 | 0.541816239 | 0.843570684 | -7.293645807 |
| **Q08752** | PPID | -0.013988837 | 3.186449487 | -0.903649211 | 0.398570079 | 0.762880385 | -7.293669281 |
| **O60869-1** | EDF1 | -0.021027464 | 3.203925337 | -0.461622968 | 0.659464925 | 0.896852471 | -7.294429795 |
| **P52564-1** | MAP2K6 | 0.007234524 | 3.160555915 | 0.270813906 | 0.794966767 | 0.949994286 | -7.294456319 |
| **Q9BXB4** | OSBPL11 | 0.01368322 | 3.189982579 | 0.552921014 | 0.59888629 | 0.871871293 | -7.294488726 |
| **O14818-1** | PSMA7 | 0.036219242 | 3.194023973 | 0.817630964 | 0.442625397 | 0.789550692 | -7.294515552 |
| **Q9UJX4** | ANAPC5 | -0.014233857 | 3.19184911 | -0.768888776 | 0.469084432 | 0.805725242 | -7.294875143 |
| **O75995** | SASH3 | 0.012720635 | 3.190087831 | 0.71481061 | 0.499691149 | 0.825030377 | -7.295761674 |
| **Q9P0U4-2** | CXXC1 | 0.006145335 | 3.192124599 | 0.234896831 | 0.821546693 | 0.959013938 | -7.296782581 |
| **P30533** | LRPAP1 | -0.006042245 | 3.187977306 | -0.120917325 | 0.907424482 | 0.98075504 | -7.297668741 |
| **Q9NYH9** | UTP6 | -0.021171584 | 3.173614107 | -0.92422782 | 0.388531093 | 0.75788842 | -7.297804009 |
| **Q53QZ3** | ARHGAP15 | 0.012110547 | 3.204019567 | 0.417614558 | 0.689733493 | 0.908760279 | -7.297964175 |
| **P78330** | PSPH | -0.006763176 | 3.192619878 | -0.375238466 | 0.719476961 | 0.925575691 | -7.299736105 |
| **Q9UII4** | HERC5 | -0.011668123 | 3.168696171 | -0.623123329 | 0.554517364 | 0.849198237 | -7.300568284 |
| **Q8TEA1** | NSUN6 | -0.000963922 | 3.174169571 | -0.043772594 | 0.9664054 | 0.992001409 | -7.300908203 |
| **Q6BDS2** | UHRF1BP1 | -0.004527302 | 3.188543713 | -0.139412809 | 0.893362585 | 0.978276697 | -7.301647338 |
| **Q14997** | PSME4 | -0.019888126 | 3.167648347 | -0.686730837 | 0.516095099 | 0.832901801 | -7.302201276 |
| **O94888** | UBXN7 | -0.003664361 | 3.14889466 | -0.172784305 | 0.868099683 | 0.970868803 | -7.302214878 |
| **Q659A1-1** | NARG2; ICE2 | 0.006096343 | 3.190836657 | 0.255751661 | 0.806079656 | 0.954293355 | -7.302637806 |
| **Q9H875** | PRKRIP1 | -0.015225415 | 3.172047578 | -0.444539739 | 0.671136447 | 0.901175787 | -7.30270032 |
| **Q7Z5K2-2** | WAPAL; WAPL | -0.012550472 | 3.182204518 | -1.053777358 | 0.329731827 | 0.715561141 | -7.303335773 |
| **Q14643-1** | ITPR1 | -0.006483128 | 3.190712372 | -0.211138116 | 0.839272985 | 0.961364456 | -7.303662434 |
| **Q03519-1** | TAP2 | 0.004610693 | 3.213174027 | 0.098778546 | 0.924303555 | 0.985099254 | -7.303674007 |
| **Q06210-1** | GFPT1 | -0.015686242 | 3.185423965 | -1.283389179 | 0.243397689 | 0.649360189 | -7.303905887 |
| **Q9BW19** | KIFC1 | -0.020531922 | 3.152097354 | -0.817690129 | 0.442593935 | 0.789550692 | -7.303936386 |
| **O43809** | NUDT21 | -0.006778847 | 3.185187342 | -0.261002495 | 0.802199903 | 0.953157309 | -7.304127554 |

| | | | | | | | |
|---|---|---|---|---|---|---|---|
| **Q13136** | PPFIA1 | 0.01814878 | 3.195650206 | 0.556712228 | 0.596439409 | 0.871378088 | -7.304412922 |
| **Q8IX18** | DHX40 | -0.001368668 | 3.170825322 | -0.074048589 | 0.943208134 | 0.988738927 | -7.304795031 |
| **Q9UNH7-1** | SNX6 | 0.012161087 | 3.164447972 | 0.687664398 | 0.515544167 | 0.832689247 | -7.305964263 |
| **Q4G0J3-3** | LARP7 | -0.040430014 | 3.155842004 | -0.865698714 | 0.417590533 | 0.774100988 | -7.306005576 |
| **Q9UKV8** | AGO2; EIF2C2 | -0.012076622 | 3.179687138 | -0.640901061 | 0.543604738 | 0.844202432 | -7.306783863 |
| **Q9NWZ5-1** | UCKL1 | 0.004447529 | 3.209676489 | 0.171146807 | 0.869335662 | 0.971430637 | -7.307528492 |
| **O60826** | CCDC22 | 0.004014164 | 3.20086944 | 0.182955872 | 0.860431354 | 0.968855822 | -7.307765549 |
| **Q96PU4-1** | UHRF2 | 0.004012779 | 3.189034342 | 0.216286527 | 0.835422727 | 0.960912428 | -7.308153625 |
| **Q13144** | EIF2B5 | 0.01250972 | 3.180961322 | 0.645338386 | 0.540901802 | 0.843540088 | -7.308701509 |
| **Q96G03** | PGM2 | -0.01771349 | 3.179749588 | -1.20518141 | 0.270347512 | 0.671731232 | -7.308826267 |
| **Q8NBF2-1** | NHLRC2 | 0.01016192 | 3.202634914 | 0.170828147 | 0.869576232 | 0.971562813 | -7.308991652 |
| **Q8WVM7** | STAG1 | 0.010273338 | 3.179651338 | 0.74583428 | 0.481972808 | 0.81235548 | -7.309070251 |
| **Q8IZH2-1** | XRN1 | -0.023096783 | 3.1590244 | -0.922128281 | 0.389546482 | 0.75855365 | -7.309334389 |
| **Q14739** | LBR | 0.005846229 | 3.196304704 | 0.160643219 | 0.877273052 | 0.973046824 | -7.309680521 |
| **O95786** | DDX58 | 0.012282163 | 3.173633454 | 0.7465393 | 0.481575141 | 0.81235548 | -7.310002326 |
| **Q96D71** | REPS1 | 0.01162471 | 3.174192397 | 0.821315417 | 0.440669173 | 0.788895128 | -7.310299887 |
| **Q3YEC7-1** | RABL6 | 0.003762847 | 3.176140608 | 0.149646496 | 0.885599598 | 0.975708125 | -7.311518704 |
| **P51580** | TPMT | -0.002740042 | 3.178659418 | -0.16414959 | 0.874621583 | 0.972759481 | -7.311871469 |
| **Q86XI2-1** | NCAPG2 | 0.011280914 | 3.17590112 | 0.621048548 | 0.555799606 | 0.850075481 | -7.312342439 |
| **P52272** | HNRNPM | 0.034566565 | 3.174808623 | 1.543017473 | 0.17019431 | 0.569105131 | -7.312635654 |
| **O95373** | IPO7 | -0.037327033 | 3.200748031 | -0.837552253 | 0.432122041 | 0.783333621 | -7.312670517 |
| **P16383-1** | GCFC2 | 0.004320091 | 3.190212591 | 0.168844031 | 0.871074463 | 0.971931713 | -7.313783039 |
| **P35637-1** | FUS | 0.010603751 | 3.195682417 | 0.522006769 | 0.619047866 | 0.882370882 | -7.313787769 |
| **O43318** | MAP3K7 | 0.013772739 | 3.182029778 | 0.489200483 | 0.640840791 | 0.890430235 | -7.314222649 |
| **Q9BRR6-1** | ADPGK | -0.002555772 | 3.185589787 | -0.06286523 | 0.951771065 | 0.989472196 | -7.315113837 |
| **Q9BUL8** | PDCD10 | -0.001924084 | 3.17986686 | -0.077757446 | 0.940370062 | 0.988019653 | -7.315337736 |
| **P46019** | PHKA2 | -8.1E-05 | 3.193296515 | -0.004008517 | 0.996922418 | 0.998480498 | -7.316077995 |
| **Q96C86** | DCPS | -0.018594313 | 3.168754243 | -0.781787344 | 0.461977976 | 0.802800142 | -7.316436208 |
| **P83436** | COG7 | 0.011124886 | 3.162861091 | 0.405963489 | 0.697854813 | 0.9129683 | -7.31735745 |
| **Q9UPU5** | USP24 | -0.015900033 | 3.18052645 | -1.202789694 | 0.271210518 | 0.671838876 | -7.3175054 |
| **Q96I15-1** | SCLY | 0.002808041 | 3.175502444 | 0.099108552 | 0.924051615 | 0.985099254 | -7.317544977 |
| **Q9UL15-2** | BAG5 | 0.004156991 | 3.179230714 | 0.156391648 | 0.880490333 | 0.974002142 | -7.317578135 |
| **Q9BRX5** | GINS3 | -0.002006961 | 3.184692011 | -0.065865338 | 0.949473194 | 0.989472196 | -7.317966229 |
| **Q96MX6** | WDR92 | -0.003788177 | 3.157669546 | -0.171884181 | 0.868779044 | 0.97108181 | -7.317999305 |
| **P53004** | BLVRA | 0.011877651 | 3.185676197 | 0.332339507 | 0.750136319 | 0.935635612 | -7.318996052 |
| **P09417** | QDPR | 0.007489747 | 3.179687827 | 0.275002606 | 0.791885419 | 0.948542293 | -7.319259095 |
| **P30414** | NKTR | -0.005298153 | 3.196476615 | -0.263663386 | 0.800236125 | 0.952507937 | -7.320232674 |
| **P51659-1** | HSD17B4 | -0.019663865 | 3.184274174 | -0.898301456 | 0.401210524 | 0.764600051 | -7.320955616 |
| **Q9H3G5** | CPVL | 0.008954558 | 3.165437094 | 0.214702725 | 0.836606661 | 0.960912428 | -7.321783589 |

| | | | | | | | |
|---|---|---|---|---|---|---|---|
| **P21291** | CSRP1 | -0.022180059 | 3.165782699 | -0.4157978 | 0.690996963 | 0.909522034 | -7.321983192 |
| **Q9UKD1** | GMEB2 | -0.001911008 | 3.192570779 | -0.065159135 | 0.95001405 | 0.989472196 | -7.323100493 |
| **Q8N335** | GPD1L | 0.006345469 | 3.175219736 | 0.290869094 | 0.780250666 | 0.94472083 | -7.323397837 |
| **Q7Z4Q2** | HEATR3 | -0.003721867 | 3.187403056 | -0.148798431 | 0.88624241 | 0.975708125 | -7.323988144 |
| **O95425-3** | SVIL | 0.009079982 | 3.192460064 | 0.374044816 | 0.72032281 | 0.925644897 | -7.324178501 |
| **P27448-5** | MARK3 | 0.004393854 | 3.195097104 | 0.132359486 | 0.898720519 | 0.978757548 | -7.324274953 |
| **Q8NDI1-2** | EHBP1 | 0.016001598 | 3.198422396 | 0.333537375 | 0.749273179 | 0.935521689 | -7.324775135 |
| **O60244** | MED14 | -0.018658651 | 3.176065233 | -0.740326235 | 0.485087281 | 0.81473183 | -7.325067139 |
| **P08243-1** | ASNS | 0.009622037 | 3.235223983 | 0.683542661 | 0.517979438 | 0.83340353 | -7.325943367 |
| **P98175-1** | RBM10 | 0.027691395 | 3.190048008 | 0.771520494 | 0.467628395 | 0.805399435 | -7.326979486 |
| **Q9Y221-1** | NIP7 | -0.014991803 | 3.204445048 | -0.250660308 | 0.809847264 | 0.955498404 | -7.327101462 |
| **Q99541** | PLIN2 | 0.010949187 | 3.201861201 | 0.377628227 | 0.717784807 | 0.924728441 | -7.327757697 |
| **Q8NBN3-1** | TMEM87A | -0.013776561 | 3.184822382 | -0.408714408 | 0.695933353 | 0.911956208 | -7.328963236 |
| **P24666-1** | ACP1 | 0.003990907 | 3.174041043 | 0.135830373 | 0.896083182 | 0.978757548 | -7.329331617 |
| **P30566** | ADSL | -0.013540541 | 3.171420153 | -0.503498954 | 0.631293132 | 0.887152592 | -7.329627849 |
| **Q6P2E9-1** | EDC4 | -0.015426607 | 3.19322224 | -1.17328772 | 0.282049733 | 0.679261937 | -7.330047277 |
| **Q12824-2** | SMARCB1 | -0.007556926 | 3.190619996 | -0.259511466 | 0.803300987 | 0.953721684 | -7.330139754 |
| **P50402** | EMD | -0.019231741 | 3.211483995 | -0.280202295 | 0.788065991 | 0.947802649 | -7.330172778 |
| **Q13627-2** | DYRK1A | -0.007547556 | 3.180687682 | -0.395427958 | 0.705235913 | 0.916737455 | -7.330627306 |
| **Q9C0B7** | TANGO6; TMCO7 | -0.017361018 | 3.164833929 | -0.448546853 | 0.668389673 | 0.90001396 | -7.33101619 |
| **Q96BN8** | FAM105B; OTULIN | -0.00403492 | 3.190832015 | -0.279218916 | 0.788787849 | 0.947865984 | -7.331153154 |
| **Q9Y4C8** | RBM19 | -0.014525 | 3.200786572 | -0.698258635 | 0.50931877 | 0.82913633 | -7.331478006 |
| **Q96J02** | ITCH | -0.026235192 | 3.198781198 | -0.779227902 | 0.463382126 | 0.802800142 | -7.331990179 |
| **Q92851-4** | CASP10 | 0.006335815 | 3.172631611 | 0.351680148 | 0.736248671 | 0.929681266 | -7.33200186 |
| **Q96KG9** | SCYL1 | 0.012119072 | 3.181434468 | 0.657265735 | 0.533678036 | 0.840132297 | -7.332710652 |
| **P05556-1** | ITGB1 | -0.025521845 | 3.165382212 | -0.572873311 | 0.586072925 | 0.8653224 | -7.332946287 |
| **Q9BSQ5-1** | CCM2 | -0.001141224 | 3.183545931 | -0.038301103 | 0.970602123 | 0.992521442 | -7.333602962 |
| **Q9BT78** | COPS4 | 0.023787268 | 3.193052264 | 0.99535266 | 0.355315201 | 0.734489383 | -7.334220043 |
| **Q6I9Y2** | THOC7 | 0.010313402 | 3.205684858 | 0.341236901 | 0.743734568 | 0.933929373 | -7.335009161 |
| **P27694** | RPA1 | 0.026760145 | 3.174644553 | 1.046611635 | 0.332787957 | 0.718869037 | -7.335408846 |
| **P11908-2** | PRPS2 | 0.017219856 | 3.167932675 | 0.207838631 | 0.841743049 | 0.962390059 | -7.335425461 |
| **P52434** | POLR2H | -0.000682699 | 3.216964489 | -0.011364957 | 0.991274622 | 0.996921123 | -7.33551114 |
| **Q99567** | NUP88 | 0.016777735 | 3.195949794 | 0.548986074 | 0.601431921 | 0.872497827 | -7.336068606 |
| **O14964** | HGS | -0.026498371 | 3.186055748 | -0.678839814 | 0.520767086 | 0.834650661 | -7.336248493 |
| **Q96S55-1** | WRNIP1 | -0.003908196 | 3.198357225 | -0.206946941 | 0.842410921 | 0.962493422 | -7.336397153 |
| **Q9Y4D8-1** | HECTD4 | 0.008615103 | 3.184351632 | 0.488082694 | 0.641590348 | 0.890911725 | -7.33639744 |
| **Q15910** | EZH2 | -0.009663837 | 3.179765448 | -0.434412277 | 0.678102942 | 0.903662615 | -7.336601426 |
| **Q96JM7** | L3MBTL3 | -0.003334178 | 3.187630013 | -0.215838738 | 0.835757414 | 0.960912428 | -7.336751259 |
| **O75175** | CNOT3 | -0.005369374 | 3.169742902 | -0.274864549 | 0.791986914 | 0.948542293 | -7.336885236 |

| | | | | | | | |
|---|---|---|---|---|---|---|---|
| **Q9BVP2** | GNL3 | -0.015450691 | 3.179341349 | -0.916508843 | 0.392274063 | 0.759806823 | -7.337137226 |
| **Q12830-1** | BPTF | -0.015266208 | 3.188471953 | -1.154959646 | 0.288966065 | 0.683685579 | -7.337434407 |
| **Q5TAX3** | ZCCHC11 | 0.005050072 | 3.181327968 | 0.324084972 | 0.756094785 | 0.937424764 | -7.33750209 |
| **P21283** | ATP6V1C1 | 0.019712586 | 3.167968379 | 0.604241075 | 0.566253004 | 0.853395233 | -7.337603735 |
| **P31146** | CORO1A | 0.011635701 | 3.192960227 | 0.966262112 | 0.368623943 | 0.743611616 | -7.337964504 |
| **P12955** | PEPD | 0.009601563 | 3.166072517 | 0.322753045 | 0.757057935 | 0.938057595 | -7.338042389 |
| **Q96ST2** | IWS1 | -0.010142318 | 3.177933732 | -0.431015938 | 0.680446969 | 0.903980599 | -7.338375224 |
| **P35244** | RPA3 | 0.003711672 | 3.187131402 | 0.147720948 | 0.887059248 | 0.975708125 | -7.338621434 |
| **Q8TED0** | UTP15 | 0.018296824 | 3.196154905 | 0.700449692 | 0.508037396 | 0.82913633 | -7.338649184 |
| **Q96RE7** | NACC1 | -0.001398511 | 3.184508689 | -0.037002654 | 0.971598208 | 0.992928787 | -7.33908985 |
| **Q8NFW8** | CMAS | -0.011917134 | 3.175385316 | -0.511676533 | 0.625866736 | 0.885387512 | -7.340666801 |
| **Q7L2H7** | EIF3M | 0.008486636 | 3.201668546 | 0.36199025 | 0.728888699 | 0.927586142 | -7.340870492 |
| **Q9Y4P3** | TBL2 | 0.002881698 | 3.207953444 | 0.087604482 | 0.932839665 | 0.986793807 | -7.341521158 |
| **Q13363-1** | CTBP1 | 0.004700839 | 3.171203287 | 0.232424575 | 0.823386153 | 0.959068023 | -7.342209993 |
| **Q8IXT5** | RBM12B | 0.004621573 | 3.176753335 | 0.213145927 | 0.837770859 | 0.96111408 | -7.342776565 |
| **Q8WVC0-1** | LEO1 | -0.003127286 | 3.197094665 | -0.1165202 | 0.910773104 | 0.981900848 | -7.342804021 |
| **Q9UKD2** | MRTO4 | -0.013754616 | 3.19667002 | -0.760399055 | 0.473802753 | 0.808788313 | -7.344357362 |
| **Q08380** | LGALS3BP | -0.007975888 | 3.206658244 | -0.44183067 | 0.672996556 | 0.901831276 | -7.344993796 |
| **Q96HC4** | PDLIM5 | -0.011107636 | 3.217516226 | -0.583416318 | 0.57936659 | 0.862073227 | -7.345210049 |
| **Q96JJ7-1** | TMX3 | 0.016539848 | 3.194440947 | 0.47580189 | 0.649855338 | 0.893733406 | -7.345240706 |
| **O60216** | RAD21 | 0.015134607 | 3.166574213 | 0.940554112 | 0.380703755 | 0.751698969 | -7.345425473 |
| **Q9C0B1-1** | FTO | 0.017736615 | 3.193589712 | 0.68673956 | 0.51608995 | 0.832901801 | -7.345548654 |
| **Q6PGP7** | TTC37 | 0.015294665 | 3.192960382 | 1.027257829 | 0.341155905 | 0.72318134 | -7.345666155 |
| **Q53F19** | C17orf85; NCBP3 | -0.004405135 | 3.177177554 | -0.13567817 | 0.896198802 | 0.978757548 | -7.345836391 |
| **Q9Y314** | NOSIP | -0.013065083 | 3.210666342 | -0.447770167 | 0.66892164 | 0.90001396 | -7.347131721 |
| **O43427** | FIBP | -0.012517293 | 3.176877421 | -0.288133386 | 0.782252463 | 0.945534128 | -7.347543298 |
| **O00193** | C11orf58 | -0.019493417 | 3.158418218 | -0.248859126 | 0.811181464 | 0.955498404 | -7.347842109 |
| **Q92696** | RABGGTA | -0.010020832 | 3.174359382 | -0.506197552 | 0.62949966 | 0.886324146 | -7.347995015 |
| **P32455** | GBP1 | -0.017795151 | 3.180400902 | -0.447976114 | 0.668780563 | 0.90001396 | -7.34849962 |
| **Q9NZ01-1** | TECR | 0.022979674 | 3.200637545 | 0.633200536 | 0.548315231 | 0.846224652 | -7.348867903 |
| **Q9Y4W2** | LAS1L | -0.006441489 | 3.185763021 | -0.330024016 | 0.751805884 | 0.935635612 | -7.350336705 |
| **Q9BVL2** | NUPL1; NUP58 | 0.000916061 | 3.186560205 | 0.043930618 | 0.96628421 | 0.992001409 | -7.350492133 |
| **Q9Y3C8** | UFC1 | 0.009291769 | 3.195387879 | 0.290558476 | 0.780477863 | 0.944826546 | -7.350624797 |
| **P55212** | CASP6 | 0.028523359 | 3.191236411 | 0.367679036 | 0.724840935 | 0.926648409 | -7.352184051 |
| **O95319-2** | CELF2 | -0.006581486 | 3.179876247 | -0.159585521 | 0.878073207 | 0.973200128 | -7.352326847 |
| **Q9HAS0** | C17orf75 | -0.003970514 | 3.186849633 | -0.24297464 | 0.815545058 | 0.95624343 | -7.352918367 |
| **Q8IXK0-5** | PHC2 | 0.005036895 | 3.191844566 | 0.207334548 | 0.842120588 | 0.962493422 | -7.353669709 |
| **P68036-3** | UBE2L3 | 0.004889017 | 3.162356791 | 0.101570543 | 0.922172335 | 0.984164603 | -7.353747737 |
| **P27708** | CAD | -0.014055321 | 3.178309002 | -1.332864364 | 0.227584734 | 0.634219821 | -7.354509594 |

| | | | | | | | |
|---|---|---|---|---|---|---|---|
| **Q9Y3F4-2** | STRAP | -0.032420354 | 3.195124602 | -1.066530367 | 0.324348761 | 0.710703247 | -7.355254398 |
| **P61962** | DCAF7 | 0.003358903 | 3.188743594 | 0.089983304 | 0.931021582 | 0.986707862 | -7.355835337 |
| **Q15149-3** | PLEC | 0.009906331 | 3.162211294 | 0.738834245 | 0.485933245 | 0.81473183 | -7.356154844 |
| **P57721-1** | PCBP3 | -0.006352892 | 3.158711507 | -0.186307277 | 0.857908279 | 0.968855822 | -7.356547413 |
| **Q96IV0** | NGLY1 | -0.007603339 | 3.199580314 | -0.370531619 | 0.722814818 | 0.926648409 | -7.356562156 |
| **Q96N67** | DOCK7 | 0.013034076 | 3.184023927 | 0.575891726 | 0.58414835 | 0.8653224 | -7.357659583 |
| **Q12792-2** | TWF1 | 0.0031474 | 3.208395686 | 0.204693229 | 0.844099572 | 0.962543524 | -7.358078378 |
| **Q92785** | DPF2 | -0.01964339 | 3.162239045 | -0.55483188 | 0.597652293 | 0.871651708 | -7.358135271 |
| **Q9Y5Q8-3** | GTF3C5 | -0.013789478 | 3.183541321 | -0.752875543 | 0.478011162 | 0.811322085 | -7.358253623 |
| **Q9Y6D9** | MAD1L1 | 0.01015595 | 3.165963687 | 0.301787214 | 0.772279778 | 0.942658294 | -7.358434621 |
| **P09960-1** | LTA4H | -0.010152178 | 3.186351141 | -0.410394541 | 0.694761009 | 0.911200194 | -7.359442119 |
| **Q9BY32-1** | ITPA | -0.006812052 | 3.16750566 | -0.270728235 | 0.795029831 | 0.949994286 | -7.35970349 |
| **P30040-1** | ERP29 | 0.018230156 | 3.205201499 | 0.518434651 | 0.621401211 | 0.883385606 | -7.361186353 |
| **P06865** | HEXA | 0.009409034 | 3.1864885 | 0.179443551 | 0.863077481 | 0.969524427 | -7.361351689 |
| **Q9NZK5** | CECR1; ADA2 | 0.007093091 | 3.166757744 | 0.311772136 | 0.765016286 | 0.940369694 | -7.361426002 |
| **O14802** | POLR3A | -0.01474146 | 3.186678438 | -1.082022891 | 0.317905382 | 0.705893003 | -7.36150659 |
| **P19447** | ERCC3 | 0.004796546 | 3.1812406 | 0.236230965 | 0.82055455 | 0.958568719 | -7.361851692 |
| **O95292** | VAPB | 0.017790457 | 3.196992693 | 0.627238981 | 0.551979178 | 0.847762337 | -7.36190098 |
| **O75116** | ROCK2 | -0.015344193 | 3.176340235 | -0.897594635 | 0.401560492 | 0.765047743 | -7.361943262 |
| **Q13485** | SMAD4 | -0.005398257 | 3.198908788 | -0.336699048 | 0.746996873 | 0.934974996 | -7.362024843 |
| **Q9UHV9** | PFDN2 | 0.013998394 | 3.180711259 | 0.252463053 | 0.808512596 | 0.955329057 | -7.362094033 |
| **Q9HC52** | CBX8 | -0.004353253 | 3.183289201 | -0.204873398 | 0.843964543 | 0.962543524 | -7.362807516 |
| **O60645** | EXOC3 | 0.011783988 | 3.186476287 | 0.428288349 | 0.68233224 | 0.904576948 | -7.362892277 |
| **P55081** | MFAP1 | 0.012237475 | 3.174395978 | 0.424455164 | 0.684985863 | 0.905978473 | -7.363019925 |
| **P82094** | TMF1 | -0.010928113 | 3.19692636 | -0.554957217 | 0.597571403 | 0.871651708 | -7.363408286 |
| **Q86U44** | METTL3 | -0.011716246 | 3.185908235 | -0.602197785 | 0.567531813 | 0.854108736 | -7.364161673 |
| **Q6PD62** | CTR9 | 0.023319194 | 3.19680168 | 1.161595958 | 0.286445492 | 0.681992638 | -7.364525881 |
| **Q9BWH6** | RPAP1 | 0.003641489 | 3.181281462 | 0.228523764 | 0.826290974 | 0.959077751 | -7.364707556 |
| **Q06546** | GABPA | 0.002187082 | 3.173293834 | 0.112448911 | 0.91387535 | 0.982219313 | -7.365646411 |
| **Q9BQ67** | GRWD1 | 0.013558885 | 3.190013761 | 0.340111763 | 0.744542909 | 0.934206162 | -7.366195427 |
| **Q8WVB6** | CHTF18 | 0.007452807 | 3.187130947 | 0.44428095 | 0.671314029 | 0.901175787 | -7.367037371 |
| **Q9BV38** | WDR18 | -0.007662094 | 3.175023749 | -0.278855422 | 0.789054731 | 0.947865984 | -7.369101067 |
| **P36507** | MAP2K2 | -0.034282729 | 3.195825354 | -0.584821286 | 0.578476295 | 0.861450787 | -7.37134521 |
| **O95067** | CCNB2 | 0.004057844 | 3.117972922 | 0.170109248 | 0.870119014 | 0.971622708 | -7.371831074 |
| **Q96DH6-1** | MSI2 | -0.014094839 | 3.196143286 | -0.254800179 | 0.806783332 | 0.954666553 | -7.371910187 |
| **Q5VWN6-1** | FAM208B | -0.007991484 | 3.200700687 | -0.760160686 | 0.4739357 | 0.808788313 | -7.371952387 |
| **Q96KB5-2** | PBK | -0.010074597 | 3.186786537 | -0.574216544 | 0.585216012 | 0.8653224 | -7.372593434 |
| **Q96T76-8** | MMS19 | 0.013144304 | 3.189557623 | 0.626191292 | 0.552624629 | 0.847927742 | -7.373067601 |
| **P62805** | HIST1H4A; HIST1H | -0.022993319 | 3.207927048 | -0.492014878 | 0.638955562 | 0.88909352 | -7.3734432 |

| | | | | | | | |
|---|---|---|---|---|---|---|---|
| **Q5W0V3** | FAM160B1 | 0.019287374 | 3.186218581 | 0.574193205 | 0.585230895 | 0.8653224 | -7.373501637 |
| **O75844** | ZMPSTE24 | -0.01422169 | 3.178235994 | -0.551697697 | 0.599677039 | 0.872097298 | -7.373785585 |
| **O43252** | PAPSS1 | -0.013039943 | 3.185895742 | -0.802178358 | 0.450897041 | 0.79513757 | -7.374104857 |
| **Q96JJ3** | ELMO2 | 0.004069328 | 3.168758738 | 0.132749923 | 0.898423777 | 0.978757548 | -7.374204881 |
| **Q01813** | PFKP | -0.014228617 | 3.181879504 | -1.127457037 | 0.299610553 | 0.690882899 | -7.374307407 |
| **Q6P1R4** | DUS1L | -0.014247884 | 3.176029396 | -0.489721839 | 0.640491341 | 0.890256071 | -7.374679139 |
| **Q15833-1** | STXBP2 | -0.004764223 | 3.182822726 | -0.182742274 | 0.86059222 | 0.968855822 | -7.375429622 |
| **P40306** | PSMB10 | 0.001586319 | 3.181577256 | 0.096810752 | 0.925806043 | 0.985476218 | -7.375554116 |
| **Q7Z5R6** | APBB1IP | -0.004684513 | 3.198231009 | -0.183378238 | 0.860113281 | 0.968855822 | -7.375881284 |
| **O00148** | DDX39A | 0.01523217 | 3.167257287 | 0.399459625 | 0.70240722 | 0.915006618 | -7.376635773 |
| **Q8WU90** | ZC3H15 | -0.024154085 | 3.199943928 | -0.58506798 | 0.578320054 | 0.861408479 | -7.376940964 |
| **P19784** | CSNK2A2 | 0.011363968 | 3.192664725 | 0.282526551 | 0.786360755 | 0.946749713 | -7.377046104 |
| **O95372** | LYPLA2 | -0.007239652 | 3.165067951 | -0.266407891 | 0.798212287 | 0.951650153 | -7.377591741 |
| **P52732** | KIF11 | -0.01671986 | 3.114416584 | -0.974188354 | 0.364959891 | 0.740073081 | -7.378778168 |
| **Q8IUF8-4** | MINA; RIOX2 | 0.002096986 | 3.183967787 | 0.074571461 | 0.942807969 | 0.988606569 | -7.378779 |
| **P61981** | YWHAG | 0.00092425 | 3.168620269 | 0.051246757 | 0.9606745 | 0.990984017 | -7.379255991 |
| **O95684** | FGFR1OP | 0.001381794 | 3.183728191 | 0.059610116 | 0.954264818 | 0.989472196 | -7.379411176 |
| **Q8WX92** | COBRA1; NELFB | 0.00172426 | 3.179633606 | 0.095631918 | 0.926706287 | 0.985528933 | -7.379530412 |
| **Q96QC0** | PPP1R10 | 0.012134136 | 3.189622119 | 0.233002077 | 0.822956359 | 0.959068023 | -7.380576827 |
| **Q9UJA5-1** | TRMT6 | 0.005692019 | 3.179557834 | 0.299509068 | 0.77394053 | 0.943158294 | -7.381044964 |
| **Q14149** | MORC3 | -0.013641336 | 3.182775451 | -0.935574402 | 0.383078333 | 0.754258979 | -7.381103839 |
| **Q9P2K8-1** | EIF2AK4 | -0.008421563 | 3.186635282 | -0.606559515 | 0.564804079 | 0.852624219 | -7.381201148 |
| **Q15554-3** | TERF2 | 0.004447292 | 3.181771713 | 0.241229966 | 0.816840187 | 0.95663256 | -7.382305551 |
| **P46063** | RECQL | 0.025044198 | 3.185201107 | 1.059979118 | 0.327105112 | 0.71218881 | -7.382861866 |
| **Q92925-1** | SMARCD2 | 0.02883372 | 3.185928929 | 0.600643291 | 0.568505857 | 0.854816036 | -7.383542621 |
| **P00374** | DHFR; DHFRP1 | 0.038196878 | 3.173556209 | 0.563206797 | 0.592260979 | 0.869575962 | -7.383931527 |
| **Q7L266** | ASRGL1 | -0.018230584 | 3.166301913 | -0.351496597 | 0.736379977 | 0.929681266 | -7.384435679 |
| **Q9P0U3-1** | SENP1 | 0.004016237 | 3.196199456 | 0.114097441 | 0.912619001 | 0.982088305 | -7.386272586 |
| **Q86UX7** | FERMT3 | -0.021171219 | 3.178573229 | -0.929429243 | 0.386024202 | 0.755595537 | -7.386273753 |
| **Q96PE3-3** | INPP4A | 0.010885496 | 3.179305628 | 0.499365225 | 0.634045632 | 0.887509451 | -7.387247462 |
| **Q2M389** | KIAA1033; WASHC | -0.020409397 | 3.164189231 | -0.485582257 | 0.643268716 | 0.891474831 | -7.387486698 |
| **Q6NXR4** | TTI2 | 0.002224783 | 3.176278728 | 0.093560596 | 0.928288383 | 0.98599615 | -7.387708562 |
| **P47755** | CAPZA2 | 0.014785721 | 3.199761347 | 0.383777822 | 0.713438319 | 0.92256957 | -7.388161008 |
| **Q9BZD4** | NUF2 | -0.004627706 | 3.13608637 | -0.141769112 | 0.891573996 | 0.977905675 | -7.389358163 |
| **Q8NEF9** | SRFBP1 | -0.001451856 | 3.197243847 | -0.050136028 | 0.961526013 | 0.991283985 | -7.389628145 |
| **Q4J6C6** | PREPL | 0.006837266 | 3.233344987 | 0.197546099 | 0.849460609 | 0.964964904 | -7.390457509 |
| **P56537-1** | EIF6 | -0.009384811 | 3.163866565 | -0.268471154 | 0.796691919 | 0.950757673 | -7.390624993 |
| **Q9NV70-1** | EXOC1 | 0.018728442 | 3.165409756 | 0.48442135 | 0.644048724 | 0.891474831 | -7.390757107 |
| **Q9UI26-2** | IPO11 | -0.00282043 | 3.175982325 | -0.205195526 | 0.843723136 | 0.962543524 | -7.391093416 |

| | | | | | | | |
|---|---|---|---|---|---|---|---|
| **Q9H4L5** | OSBPL3 | -0.002176878 | 3.194835738 | -0.109245132 | 0.916317724 | 0.983006254 | -7.391192065 |
| **Q9UBU9-1** | NXF1 | 0.012316245 | 3.178604412 | 0.368017183 | 0.724600631 | 0.926648409 | -7.392015489 |
| **Q6ZRQ5** | MMS22L | -0.008253959 | 3.195871452 | -0.315978785 | 0.761963821 | 0.939498834 | -7.392078489 |
| **P35613** | BSG | -0.016923419 | 3.17083029 | -0.411366971 | 0.694082893 | 0.910733045 | -7.39213872 |
| **O60547-1** | GMDS | -0.008542206 | 3.176816416 | -0.805027736 | 0.449363636 | 0.794586812 | -7.394981857 |
| **Q9Y6Q9-1** | NCOA3 | -0.004557267 | 3.181947299 | -0.277872057 | 0.789776888 | 0.947865984 | -7.396068163 |
| **Q8NHM5-1** | KDM2B | -0.001246456 | 3.183915354 | -0.042224549 | 0.967592665 | 0.992306703 | -7.39609508 |
| **Q13838-2** | DDX39B | -0.004523878 | 3.171822184 | -0.295456057 | 0.776898331 | 0.94357883 | -7.396245703 |
| **O14617-5** | AP3D1 | -0.009088992 | 3.171568398 | -0.389513193 | 0.70939502 | 0.919886312 | -7.396449136 |
| **P30153** | PPP2R1A | -0.035824039 | 3.173674458 | -1.030588826 | 0.339703838 | 0.722416219 | -7.39698726 |
| **P05771-2** | PRKCB | -0.009729841 | 3.181488512 | -0.463730032 | 0.658032389 | 0.896852471 | -7.397400404 |
| **Q75N03-2** | CBLL1 | -0.004408965 | 3.210190224 | -0.104976091 | 0.919573718 | 0.983742622 | -7.39757069 |
| **P15498-1** | VAV1 | 0.022931846 | 3.186030464 | 0.862855261 | 0.419042133 | 0.775696586 | -7.397716333 |
| **Q92540-4** | SMG7 | 0.002471155 | 3.208848368 | 0.185332413 | 0.858642011 | 0.968855822 | -7.397923587 |
| **Q68E01-2** | INTS3 | 0.007204867 | 3.184639321 | 0.753231239 | 0.477811624 | 0.811322085 | -7.398120502 |
| **O15111** | CHUK | -0.005727027 | 3.186537339 | -0.299398059 | 0.774021488 | 0.943158294 | -7.399014863 |
| **Q92541** | RTF1 | -0.012256441 | 3.161477833 | -0.888139474 | 0.406263957 | 0.766336431 | -7.400749845 |
| **Q99700-1** | ATXN2 | -0.024752964 | 3.173065952 | -0.569176404 | 0.588435099 | 0.867233795 | -7.401127025 |
| **Q12882-1** | DPYD | -0.025079541 | 3.160387611 | -0.917556155 | 0.391764624 | 0.759806823 | -7.401259067 |
| **P23919** | DTYMK | -0.023615069 | 3.175060139 | -0.811481564 | 0.445904117 | 0.792020296 | -7.401645203 |
| **O95573** | ACSL3 | -0.009850569 | 3.183900135 | -0.863959724 | 0.418477859 | 0.775013192 | -7.402119679 |
| **P28907** | CD38 | 0.013182223 | 3.157066206 | 0.565510768 | 0.590782695 | 0.868764508 | -7.40282241 |
| **Q01167** | FOXK2 | 0.011941203 | 3.197603695 | 0.217991701 | 0.834148583 | 0.960758262 | -7.40360593 |
| **Q13492-1** | PICALM | 0.004315001 | 3.225463322 | 0.075012203 | 0.942470674 | 0.988606569 | -7.40393951 |
| **Q92922** | SMARCC1 | 0.00969505 | 3.16643118 | 1.011280039 | 0.348189807 | 0.730105045 | -7.404219491 |
| **Q9UL40-1** | ZNF346 | 0.005033023 | 3.183644091 | 0.270077895 | 0.795508615 | 0.950423451 | -7.404559177 |
| **Q99614** | TTC1 | 0.024174618 | 3.191080475 | 0.724951639 | 0.493852298 | 0.821555157 | -7.404721309 |
| **Q14203** | DCTN1 | 0.009399256 | 3.168686589 | 1.201466686 | 0.271688909 | 0.671838876 | -7.40503352 |
| **Q86V21-1** | AACS | -0.009065705 | 3.179333277 | -0.533398065 | 0.611575709 | 0.878338577 | -7.405989918 |
| **Q9UHB9** | SRP68 | -0.037099636 | 3.167033003 | -0.800170931 | 0.451979559 | 0.795167759 | -7.407067167 |
| **P08397-1** | HMBS | 0.018465377 | 3.193112888 | 0.436926811 | 0.676370002 | 0.903662615 | -7.407146009 |
| **Q12904-2** | AIMP1 | 0.022125316 | 3.180479886 | 0.745394531 | 0.482220962 | 0.812393033 | -7.407527593 |
| **O75362** | ZNF217 | -0.005898046 | 3.1677478 | -0.444760686 | 0.670984851 | 0.901175787 | -7.408798497 |
| **Q14141-1** | 6-Sep | 0.026222263 | 3.19565183 | 0.372608128 | 0.721341447 | 0.926007076 | -7.409632394 |
| **Q92917** | GPKOW | -0.018944385 | 3.176465686 | -0.643275602 | 0.542157277 | 0.843603519 | -7.410082419 |
| **O43166-2** | SIPA1L1 | 0.000640901 | 3.191500775 | 0.031838604 | 0.97556026 | 0.99316605 | -7.41095813 |
| **Q9BT09** | CNPY3 | 0.012921159 | 3.207453388 | 0.299939699 | 0.773626504 | 0.943158294 | -7.411463658 |
| **Q8WUQ7-2** | CACTIN | -0.014773871 | 3.191387435 | -0.309215641 | 0.766873581 | 0.940879515 | -7.411580155 |
| **Q9H3N1** | TMX1 | -0.004782274 | 3.21159186 | -0.304458475 | 0.770334105 | 0.941730183 | -7.411807653 |

| | | | | | | | |
|---|---|---|---|---|---|---|---|
| **Q9HBK9** | AS3MT | 0.000720109 | 3.181595699 | 0.047784526 | 0.963328913 | 0.991283985 | -7.412717852 |
| **O60610-1** | DIAPH1 | -0.010469539 | 3.179666012 | -1.399431331 | 0.207748999 | 0.614231901 | -7.413065001 |
| **Q13451-1** | FKBP5 | 0.026667563 | 3.190865122 | 1.043019905 | 0.334328347 | 0.719277002 | -7.413645486 |
| **O60264** | SMARCA5 | -0.013294595 | 3.180210186 | -1.272808205 | 0.246901855 | 0.652280071 | -7.415059565 |
| **Q9UHD1** | CHORDC1 | 0.029939076 | 3.190828101 | 0.908195684 | 0.396335528 | 0.762021989 | -7.415107379 |
| **P25490** | YY1 | 0.001731512 | 3.178001315 | 0.043887704 | 0.966317122 | 0.992001409 | -7.415591409 |
| **P35520-2** | CBS; LOC10272450 | -0.00806268 | 3.202593905 | -0.235401655 | 0.821171234 | 0.95877683 | -7.416780945 |
| **Q8NF50-1** | DOCK8 | -0.016575445 | 3.164725953 | -0.927297614 | 0.387050078 | 0.75648533 | -7.41691597 |
| **Q96T60** | PNKP | -0.003386687 | 3.168173361 | -0.184936104 | 0.858940337 | 0.968855822 | -7.417677248 |
| **Q9BVQ7-1** | SPATA5L1 | -0.009524277 | 3.203095459 | -0.364672339 | 0.726979118 | 0.92710315 | -7.418366588 |
| **Q9UG01** | IFT172 | -0.000332274 | 3.189619912 | -0.0141559 | 0.98913204 | 0.996219143 | -7.418425081 |
| **Q15084-2** | PDIA6 | 0.022243137 | 3.205974962 | 0.52730421 | 0.615566809 | 0.880583268 | -7.418431618 |
| **Q9NZT2-1** | OGFR | 0.008557672 | 3.199972278 | 0.480005726 | 0.647020022 | 0.89271426 | -7.418780843 |
| **Q92466** | DDB2 | 0.01407137 | 3.182892487 | 0.473897433 | 0.651141904 | 0.894264896 | -7.418849274 |
| **Q5JVF3** | PCID2 | -0.012825727 | 3.184331851 | -0.647196873 | 0.539772219 | 0.843540088 | -7.418878418 |
| **O60318** | MCM3AP | 0.009051692 | 3.183430734 | 0.60999724 | 0.562659744 | 0.852576894 | -7.419069271 |
| **Q13395** | TARBP1 | 0.008655024 | 3.20241382 | 0.444182258 | 0.671381759 | 0.901175787 | -7.420000193 |
| **Q00610-1** | CLTC | 0.014991005 | 3.193877219 | 1.482179016 | 0.185272283 | 0.588324655 | -7.42001568 |
| **Q14166** | TTLL12 | 0.011862697 | 3.186997713 | 0.677791824 | 0.521389601 | 0.834809173 | -7.420202832 |
| **P13612** | ITGA4 | -0.015399727 | 3.162440002 | -0.878472755 | 0.411114863 | 0.769906516 | -7.420328561 |
| **Q9NW64-1** | RBM22 | -0.006075568 | 3.197578539 | -0.406159576 | 0.69771777 | 0.912939342 | -7.420343928 |
| **Q6ZSZ5** | ARHGEF18 | -0.00791812 | 3.192342382 | -0.474222716 | 0.650922065 | 0.894219754 | -7.421301334 |
| **Q9BZH6** | WDR11 | 0.009731271 | 3.187340757 | 0.768007952 | 0.469572458 | 0.805952296 | -7.421695804 |
| **Q9UGR2** | ZC3H7B | -0.004383686 | 3.217094403 | -0.17667224 | 0.865166697 | 0.9700838 | -7.421863742 |
| **Q15386-1** | UBE3C | -0.008505871 | 3.166273019 | -0.366744353 | 0.72550534 | 0.926648409 | -7.424883414 |
| **P55201-2** | BRPF1 | -0.000547611 | 3.187128156 | -0.017593032 | 0.986493523 | 0.995641646 | -7.424897579 |
| **Q13617-2** | CUL2 | -0.008347976 | 3.186404164 | -0.757006272 | 0.47569742 | 0.809713018 | -7.424991518 |
| **Q92974** | ARHGEF2 | 0.012646465 | 3.196663832 | 0.90330865 | 0.398737841 | 0.762880385 | -7.425020816 |
| **Q14232** | EIF2B1 | -0.008819086 | 3.188970169 | -0.318894442 | 0.759850839 | 0.938422437 | -7.428276758 |
| **Q969V3** | NCLN | 0.001307963 | 3.182605597 | 0.06511858 | 0.95004511 | 0.989472196 | -7.429204676 |
| **P48059-5** | LIMS1 | -0.000917177 | 3.204230322 | -0.026103785 | 0.979961103 | 0.99409997 | -7.429507661 |
| **P10155-1** | TROVE2 | 0.008306268 | 3.183001952 | 0.491519615 | 0.639287104 | 0.88920449 | -7.429610746 |
| **Q12872-2** | SFSWAP | -0.00311735 | 3.183522084 | -0.065015844 | 0.950123794 | 0.989472196 | -7.431325247 |
| **Q14696** | MESDC2; MESD | 0.001183705 | 3.19575531 | 0.03269849 | 0.974900468 | 0.99316605 | -7.431383954 |
| **P57764** | GSDMD | 0.001021368 | 3.181049584 | 0.049167835 | 0.962268299 | 0.991283985 | -7.432419538 |
| **Q00796** | SORD | 0.013169259 | 3.176752275 | 0.617308957 | 0.55811527 | 0.851117349 | -7.432831473 |
| **O43747-1** | AP1G1 | -0.009883808 | 3.171946959 | -0.648194584 | 0.53916642 | 0.843540088 | -7.433634432 |
| **P13984** | GTF2F2 | -0.010442209 | 3.18230248 | -0.38299756 | 0.713989165 | 0.922981437 | -7.434316237 |
| **Q9UBQ0** | VPS29 | -0.005084011 | 3.165014401 | -0.10444907 | 0.919975794 | 0.983742622 | -7.434538318 |

| | | | | | | | |
|---|---|---|---|---|---|---|---|
| **O75530** | EED | -0.008832314 | 3.177448433 | -0.597113466 | 0.570721337 | 0.856351468 | -7.434954968 |
| **Q13151** | HNRNPA0 | 0.028110636 | 3.197678684 | 0.889790942 | 0.405439493 | 0.766336431 | -7.435139855 |
| **P60981-1** | DSTN | -0.002537668 | 3.183398044 | -0.140693243 | 0.892390567 | 0.978206823 | -7.435186337 |
| **Q13017-1** | ARHGAP5 | -0.014113889 | 3.203303517 | -0.768550905 | 0.46927159 | 0.805784047 | -7.435317728 |
| **O00743** | PPP6C | -0.001390128 | 3.196139818 | -0.042058064 | 0.967720355 | 0.992306703 | -7.435770871 |
| **P28288** | ABCD3 | -0.0257482 | 3.180764017 | -0.7566775 | 0.475881295 | 0.809713018 | -7.436012735 |
| **O75351** | VPS4B | 0.01737731 | 3.196788718 | 0.256783105 | 0.805317066 | 0.954249678 | -7.43656828 |
| **P26038** | MSN | 0.023427389 | 3.162048412 | 1.291965649 | 0.240589316 | 0.646207612 | -7.437280178 |
| **P00390** | GSR | -0.016818348 | 3.169138885 | -0.8200115 | 0.441360769 | 0.788971713 | -7.437743526 |
| **Q6VY07** | PACS1 | 0.004063174 | 3.184601761 | 0.220828495 | 0.832030073 | 0.960474408 | -7.437814385 |
| **Q9UPN7** | PPP6R1 | 0.002668853 | 3.180115867 | 0.168936827 | 0.871004379 | 0.971931713 | -7.439114156 |
| **Q6YHU6** | THADA | 0.001277689 | 3.211486001 | 0.062308001 | 0.952197919 | 0.989472196 | -7.440976352 |
| **Q9UFW8** | CGGBP1 | -0.007226026 | 3.180446875 | -0.317586558 | 0.76079839 | 0.938739432 | -7.44200029 |
| **P48444** | ARCN1 | -0.036713396 | 3.206029771 | -1.099801938 | 0.310640054 | 0.69876422 | -7.444130312 |
| **Q9UPT8** | ZC3H4 | 0.013309992 | 3.179405681 | 0.431172054 | 0.680339139 | 0.903980599 | -7.445411189 |
| **Q9NYB0** | TERF2IP | -0.00562791 | 3.188598643 | -0.432263395 | 0.679585572 | 0.903893384 | -7.445581477 |
| **Q04759-1** | PRKCQ | 0.018788693 | 3.178941886 | 0.564784763 | 0.591248289 | 0.868764508 | -7.446988516 |
| **Q6GYQ0-2** | RALGAPA1 | 0.007738659 | 3.188486082 | 0.448766472 | 0.668239288 | 0.90001396 | -7.447282578 |
| **P06454-1** | PTMA | -0.103824446 | 3.133692948 | -0.673385135 | 0.524012422 | 0.837176492 | -7.447456053 |
| **Q9BVA0** | KATNB1 | 0.006591817 | 3.150495511 | 0.376885524 | 0.71831052 | 0.924851578 | -7.449207305 |
| **Q15650** | TRIP4 | 0.00178361 | 3.183074025 | 0.133825561 | 0.897606356 | 0.978757548 | -7.449746185 |
| **Q14573** | ITPR3 | -0.000962154 | 3.191489201 | -0.031687185 | 0.975676447 | 0.99316605 | -7.450216705 |
| **P04053** | DNTT | 0.013090017 | 3.123396371 | 0.659542083 | 0.532306294 | 0.839951983 | -7.451550918 |
| **Q6FI81-1** | CIAPIN1 | -0.004098613 | 3.190956018 | -0.094149828 | 0.927838285 | 0.9859101 | -7.453508716 |
| **P08237-3** | PFKM | 0.024741681 | 3.171559049 | 0.816457304 | 0.443249838 | 0.789680112 | -7.453906782 |
| **P46934-4** | NEDD4 | -0.005662877 | 3.191847117 | -0.295695451 | 0.776723514 | 0.94357883 | -7.454498201 |
| **P36915-1** | GNL1 | -0.006586741 | 3.175852776 | -0.214930223 | 0.836436572 | 0.960912428 | -7.455338405 |
| **Q15691** | MAPRE1 | 0.033323621 | 3.190863711 | 0.760542287 | 0.473722881 | 0.808788313 | -7.455873848 |
| **Q13111-1** | CHAF1A | -0.011607736 | 3.199405673 | -0.77636916 | 0.464953974 | 0.803074528 | -7.45805045 |
| **P63010-2** | AP2B1 | -0.005952544 | 3.19871571 | -0.427699082 | 0.682739859 | 0.904966329 | -7.458520202 |
| **Q9H6T3** | RPAP3 | -0.015524951 | 3.181428229 | -0.859343173 | 0.420840172 | 0.776852967 | -7.459026543 |
| **Q03252** | LMNB2 | 0.010627204 | 3.176323632 | 0.777243631 | 0.464472764 | 0.803074528 | -7.459348625 |
| **Q15047** | SETDB1 | 0.003657519 | 3.207335099 | 0.197643041 | 0.849387834 | 0.964964904 | -7.460319811 |
| **P61289-2** | PSME3 | -0.018614088 | 3.180484718 | -0.916094676 | 0.392475662 | 0.759985165 | -7.460371383 |
| **Q96RL1** | UIMC1 | 0.000751932 | 3.18712407 | 0.025899271 | 0.98011806 | 0.994132259 | -7.46055324 |
| **Q9H267** | VPS33B | -0.001040253 | 3.163020677 | -0.073020774 | 0.943994794 | 0.98895887 | -7.460836818 |
| **Q6P996-1** | PDXDC1; LOC1027 | -0.003758881 | 3.182855322 | -0.249405877 | 0.810776393 | 0.955498404 | -7.460990197 |
| **Q8TEM1-1** | NUP210 | 0.019079291 | 3.218510545 | 0.798283332 | 0.452999128 | 0.796080086 | -7.461472641 |
| **Q6ZRS2** | SRCAP | -0.00284332 | 3.181097506 | -0.221197175 | 0.831754855 | 0.960474408 | -7.462509394 |

| | | | | | | | |
|---|---|---|---|---|---|---|---|
| **P61201-2** | COPS2 | 0.012515381 | 3.179455783 | 0.551597669 | 0.599741723 | 0.872097298 | -7.462744409 |
| **Q16401** | PSMD5 | 0.008650875 | 3.197187347 | 0.310740657 | 0.765765454 | 0.940484723 | -7.463357802 |
| **O43776** | NARS | -0.007067442 | 3.193895128 | -0.574661829 | 0.584932104 | 0.8653224 | -7.464520626 |
| **Q13217** | DNAJC3 | 0.005249643 | 3.197541776 | 0.486395099 | 0.642722863 | 0.891449441 | -7.464986093 |
| **O43823** | AKAP8 | 0.004293807 | 3.167769485 | 0.237636705 | 0.819509544 | 0.958064056 | -7.465591402 |
| **Q9UNM6** | PSMD13 | 0.017361794 | 3.185125054 | 0.942531754 | 0.379763842 | 0.750948043 | -7.466004287 |
| **Q9H814** | PHAX | -0.002427206 | 3.205996357 | -0.158082664 | 0.879210396 | 0.97338977 | -7.466544139 |
| **P16949-1** | STMN1 | -0.070086768 | 3.167139722 | -1.111403302 | 0.305973074 | 0.694598791 | -7.467358968 |
| **Q9Y3B8-1** | REXO2 | 0.009270187 | 3.173719713 | 0.204194854 | 0.844473114 | 0.962543524 | -7.467408883 |
| **O75122-3** | CLASP2 | 0.014670021 | 3.185689112 | 0.901689634 | 0.399536102 | 0.763210165 | -7.467640045 |
| **P35226** | BMI1; COMMD3-BI | -0.003814403 | 3.184458594 | -0.107923397 | 0.917325626 | 0.983235402 | -7.467818561 |
| **Q66K14-1** | TBC1D9B | -0.00348395 | 3.193795854 | -0.285292583 | 0.784333067 | 0.946321369 | -7.468810235 |
| **Q9Y617-1** | PSAT1 | -0.006293386 | 3.200257703 | -0.512098585 | 0.625587355 | 0.885387512 | -7.47004329 |
| **Q15334** | LLGL1 | 0.002977234 | 3.185590146 | 0.084304664 | 0.935362367 | 0.986793807 | -7.470598676 |
| **P39880-3** | CUX1 | 0.006449232 | 3.191528651 | 0.396773299 | 0.704291429 | 0.916268778 | -7.47117906 |
| **P46013** | MKI67 | -0.059303845 | 3.172665164 | -1.197842089 | 0.27300322 | 0.672884265 | -7.471853017 |
| **P35249** | RFC4 | -0.012497735 | 3.172042002 | -0.595367453 | 0.571819106 | 0.857347804 | -7.472016534 |
| **O15305** | PMM2 | -0.002170592 | 3.181888979 | -0.062855223 | 0.951778731 | 0.989472196 | -7.47226068 |
| **Q8N122** | RPTOR | 0.003137786 | 3.173511351 | 0.20410216 | 0.844542594 | 0.962543524 | -7.472497743 |
| **P07384** | CAPN1 | 0.011924578 | 3.167110618 | 0.65621821 | 0.534310028 | 0.840279726 | -7.473311132 |
| **Q9NTJ5** | SACM1L | 0.018834595 | 3.196757669 | 0.839327169 | 0.431195004 | 0.782692325 | -7.473843894 |
| **Q2TBE0** | CWF19L2 | -0.006893428 | 3.187852601 | -0.384207257 | 0.713135227 | 0.922407393 | -7.475126588 |
| **Q92747** | ARPC1A | -0.009199091 | 3.171900626 | -0.585283492 | 0.578183581 | 0.861366689 | -7.475364848 |
| **O43447** | PPIH | -0.01050441 | 3.160204497 | -0.366820719 | 0.725451046 | 0.926648409 | -7.476348821 |
| **P31150** | GDI1 | -0.009457337 | 3.187911346 | -0.227670758 | 0.82692658 | 0.959077751 | -7.478330769 |
| **Q8TBC4-1** | UBA3 | -0.007271193 | 3.17318024 | -0.376223402 | 0.718779334 | 0.925182096 | -7.479682064 |
| **O94903** | PROSC; PLPBP | 0.012942644 | 3.180904926 | 0.623553007 | 0.554252042 | 0.848970058 | -7.480193665 |
| **Q13263** | TRIM28 | 0.021579709 | 3.186473756 | 1.414965894 | 0.203349434 | 0.608695713 | -7.481464521 |
| **P35269** | GTF2F1 | -0.008514585 | 3.178954532 | -0.371174554 | 0.722358491 | 0.926635956 | -7.481571849 |
| **Q9UHB6-4** | LIMA1 | -0.003922324 | 3.185264635 | -0.236497083 | 0.820356692 | 0.958568719 | -7.48158972 |
| **O43765** | SGTA | -0.01765819 | 3.199083321 | -0.544501037 | 0.60434083 | 0.87347424 | -7.481751732 |
| **P52333-1** | JAK3 | 0.000905128 | 3.1867816 | 0.059659697 | 0.954226829 | 0.989472196 | -7.482293629 |
| **Q92888-3** | ARHGEF1 | 0.022463216 | 3.184159363 | 0.631022785 | 0.549651932 | 0.846638465 | -7.483221036 |
| **Q86V48-1** | LUZP1 | 0.004942486 | 3.211461624 | 0.160800148 | 0.877154347 | 0.973046824 | -7.484139873 |
| **Q14137** | BOP 1.00 | 0.003118577 | 3.167223585 | 0.152943817 | 0.883101211 | 0.974883858 | -7.484508558 |
| **O95639** | CPSF4 | -0.007157425 | 3.205006389 | -0.254155149 | 0.807260479 | 0.954765339 | -7.486353316 |
| **Q15369-1** | TCEB1; ELOC | -0.017589243 | 3.204258739 | -0.364293167 | 0.727248951 | 0.92710315 | -7.486761409 |
| **Q9Y4B4** | RAD54L2 | -0.001789011 | 3.176325407 | -0.095142329 | 0.927080207 | 0.985631941 | -7.487220667 |
| **Q96P48-6** | ARAP1 | 0.005655828 | 3.18396873 | 0.233435528 | 0.822633816 | 0.959068023 | -7.487244933 |

| | | | | | | | |
|---|---|---|---|---|---|---|---|
| **Q9Y5J1** | UTP18 | -0.005288979 | 3.199732217 | -0.332280991 | 0.750178494 | 0.935635612 | -7.487264066 |
| **P30419** | NMT1 | -0.017375343 | 3.192506902 | -0.238080834 | 0.819179468 | 0.957819113 | -7.487988277 |
| **P39019** | RPS19 | -0.02648215 | 3.195302231 | -0.800557914 | 0.451770734 | 0.79513757 | -7.488051203 |
| **P30085-1** | CMPK1 | 0.005016566 | 3.201360878 | 0.048185431 | 0.963021522 | 0.991283985 | -7.488880003 |
| **Q68CZ6** | HAUS3 | 0.005274885 | 3.197243065 | 0.244348648 | 0.814525528 | 0.95620584 | -7.489558539 |
| **O43815** | STRN | -0.008787885 | 3.170739935 | -0.638311034 | 0.545186282 | 0.84500683 | -7.489708794 |
| **Q0VDF9** | HSPA14 | -0.000867622 | 3.16288523 | -0.040720955 | 0.968745923 | 0.992306703 | -7.489988083 |
| **Q99459** | CDC5L | -0.015887449 | 3.168033107 | -0.711991011 | 0.501322662 | 0.826521799 | -7.48999429 |
| **Q12996-1** | CSTF3 | -0.0219601 | 3.161529907 | -0.509867839 | 0.627064774 | 0.885387512 | -7.490350912 |
| **Q9Y4X5** | ARIH1 | 0.008329152 | 3.16960766 | 0.463849349 | 0.657951315 | 0.896852471 | -7.491669153 |
| **Q14571-1** | ITPR2 | 0.014537246 | 3.178695384 | 0.330587307 | 0.751399594 | 0.935635612 | -7.492071309 |
| **P54277** | PMS1 | -0.011519638 | 3.203668091 | -0.444785278 | 0.670967979 | 0.901175787 | -7.492761714 |
| **Q9UGP8** | SEC63 | 0.012591922 | 3.18720248 | 0.552965351 | 0.598857642 | 0.871871293 | -7.493098991 |
| **Q9H3P2-1** | NELFA; WHSC2 | -0.005645082 | 3.200815825 | -0.244977343 | 0.814059159 | 0.95620584 | -7.494281236 |
| **Q15027** | ACAP1 | 0.000820124 | 3.189334871 | 0.052225736 | 0.959924037 | 0.990724405 | -7.494328114 |
| **Q7LBC6-1** | KDM3B | 0.019760024 | 3.176192928 | 1.062680279 | 0.325966348 | 0.710703247 | -7.495912284 |
| **Q13523** | PRPF4B | 0.008384668 | 3.185740135 | 0.700655991 | 0.507916857 | 0.82913633 | -7.496133825 |
| **O95861-1** | BPNT1 | 0.010741545 | 3.184789714 | 0.61981449 | 0.556563125 | 0.850691425 | -7.498878806 |
| **Q9Y3P9** | RABGAP1 | -0.004797486 | 3.191378728 | -0.258176028 | 0.804287585 | 0.953936916 | -7.499533171 |
| **P55786** | NPEPPS | -0.026173715 | 3.169679265 | -1.09499752 | 0.312589816 | 0.700451368 | -7.499795013 |
| **Q6UXN9** | WDR82 | 0.020237271 | 3.197992897 | 0.788152876 | 0.458498588 | 0.799639769 | -7.500920362 |
| **Q01433-1** | AMPD2 | -0.012574192 | 3.202335588 | -0.30307571 | 0.771341049 | 0.942381153 | -7.501206353 |
| **Q8N7H5** | PAF1 | -0.008567052 | 3.189854921 | -0.359653735 | 0.730553952 | 0.927984559 | -7.501317619 |
| **O14497** | ARID1A | -0.013560158 | 3.181016985 | -1.141855957 | 0.293997617 | 0.686599373 | -7.502136947 |
| **P19623** | SRM | -0.005757235 | 3.180634473 | -0.382412703 | 0.714402182 | 0.92336511 | -7.5037124 |
| **Q86Y56** | HEATR2; DNAAF5 | -0.000708247 | 3.18067318 | -0.042473474 | 0.967401747 | 0.992306703 | -7.504430478 |
| **Q9NZW5** | MPP6 | -0.00995685 | 3.189733605 | -0.300282654 | 0.773376446 | 0.943127531 | -7.504792074 |
| **Q96T51** | RUFY1 | -0.007477971 | 3.198982166 | -0.291211904 | 0.779999949 | 0.94472083 | -7.506175886 |
| **Q9H9A6** | LRRC40 | 0.013914938 | 3.1787464 | 0.505381575 | 0.630041667 | 0.88658892 | -7.506476716 |
| **Q5GLZ8-1** | HERC4 | 0.00076332 | 3.181438065 | 0.026672863 | 0.979524362 | 0.993783815 | -7.507683923 |
| **O75165** | DNAJC13 | 0.009870267 | 3.182760363 | 0.899641674 | 0.400547568 | 0.763806762 | -7.508037579 |
| **Q9Y4C1** | KDM3A | -0.039008566 | 3.154309392 | -0.72447143 | 0.494127758 | 0.821650279 | -7.508400191 |
| **Q6PJG2** | ELMSAN1 | 0.005614293 | 3.188060972 | 0.306293103 | 0.768998854 | 0.941649022 | -7.508536432 |
| **Q8IZL8** | PELP1 | 0.001616493 | 3.176246742 | 0.098220064 | 0.924729943 | 0.985113891 | -7.508964079 |
| **Q14677-3** | CLINT1 | -0.012141275 | 3.173150266 | -0.541369083 | 0.606376815 | 0.874996148 | -7.509156283 |
| **Q9NWY4** | C4orf27; HPF1 | 0.009347925 | 3.181953082 | 0.292590141 | 0.778992255 | 0.944053983 | -7.509802775 |
| **O95782** | AP2A1 | -0.011935898 | 3.173607069 | -0.612112865 | 0.561342527 | 0.852576894 | -7.509972371 |
| **Q9UL03** | INTS6 | -0.001495937 | 3.183398424 | -0.08469153 | 0.935066566 | 0.986793807 | -7.510491922 |
| **P06730-2** | EIF4E | 0.000284382 | 3.166847478 | 0.008400162 | 0.993550752 | 0.99756865 | -7.512251719 |

| | | | | | | | |
|---|---|---|---|---|---|---|---|
| **Q9HD45** | TM9SF3 | 0.010142855 | 3.170433608 | 0.151641499 | 0.884087809 | 0.975295424 | -7.512741704 |
| **P50579** | METAP2 | -0.015119728 | 3.160475313 | -0.368884066 | 0.723984738 | 0.926648409 | -7.514457214 |
| **O15084** | ANKRD28 | 0.010341921 | 3.196462543 | 0.589060638 | 0.575794782 | 0.860065716 | -7.515122201 |
| **Q7Z2Z2** | EFTUD1; EFL1 | 0.012854766 | 3.17856042 | 0.802838103 | 0.450541669 | 0.79513757 | -7.515459131 |
| **Q07666** | KHDRBS1 | -0.024299271 | 3.163757814 | -0.340225832 | 0.744460942 | 0.934206162 | -7.515615969 |
| **P50995** | ANXA11 | 0.01094824 | 3.182063586 | 0.591639894 | 0.574166911 | 0.85911412 | -7.515623371 |
| **Q5TBB1** | RNASEH2B | 0.012509771 | 3.165570906 | 0.663267995 | 0.53006584 | 0.839849063 | -7.515971957 |
| **O94762-1** | RECQL5 | 0.006294702 | 3.182149054 | 0.306635821 | 0.768749516 | 0.941649022 | -7.51656131 |
| **Q15751** | HERC1 | -0.001592963 | 3.186823573 | -0.12564306 | 0.903827896 | 0.980210603 | -7.517157303 |
| **Q9H6R4-1** | NOL6 | 0.005127532 | 3.195845708 | 0.372420151 | 0.721474771 | 0.926028604 | -7.517504236 |
| **Q96AG4** | LRRC59 | -0.002551006 | 3.212499523 | -0.058029067 | 0.955476268 | 0.989472196 | -7.518521748 |
| **Q8WUM4** | PDCD6IP | -0.018864867 | 3.167581809 | -1.114310414 | 0.304812702 | 0.693509999 | -7.519581532 |
| **Q8IV48** | ERI1 | 0.00089193 | 3.172865775 | 0.030430009 | 0.976641122 | 0.99316605 | -7.523077743 |
| **P61158** | ACTR3 | 0.021903626 | 3.191105271 | 0.630953878 | 0.549694259 | 0.846638465 | -7.523919554 |
| **Q8WYQ5-1** | DGCR8 | -0.001943588 | 3.186760915 | -0.090323618 | 0.930761524 | 0.986707862 | -7.523981394 |
| **Q9ULI0** | ATAD2B | 0.006306689 | 3.176909385 | 0.499866617 | 0.633711436 | 0.887509451 | -7.525745567 |
| **Q68CQ4** | DIEXF | -0.006009032 | 3.191119927 | -0.369031787 | 0.723879809 | 0.926648409 | -7.526862532 |
| **Q96PZ0** | PUS7 | 0.011023497 | 3.181261469 | 0.947666949 | 0.377331536 | 0.749474764 | -7.526869999 |
| **Q96CW5** | TUBGCP3 | -0.006366476 | 3.187684666 | -0.342271334 | 0.742991704 | 0.933447653 | -7.527020898 |
| **Q9H9T3-2** | ELP3 | -5.03E-05 | 3.186554127 | -0.003146335 | 0.997584365 | 0.998480498 | -7.528146194 |
| **P25788-1** | PSMA3 | -0.008559788 | 3.168771672 | -0.630892595 | 0.549731906 | 0.846638465 | -7.528386758 |
| **Q9UP83-2** | COG5 | -0.01741136 | 3.167942981 | -0.257124044 | 0.805065046 | 0.954249678 | -7.528598165 |
| **Q9Y4K1** | AIM1; CRYBG1 | 0.015794319 | 3.185960309 | 0.644206737 | 0.541590335 | 0.843570684 | -7.529241343 |
| **P28070** | PSMB4 | -0.002055417 | 3.176774161 | -0.102665779 | 0.921336496 | 0.984164603 | -7.529485807 |
| **Q92882** | OSTF1 | -0.004088682 | 3.212297603 | -0.236082251 | 0.820665126 | 0.958568719 | -7.529678248 |
| **P30086** | PEBP1 | 0.018670388 | 3.16101829 | 0.966552236 | 0.368489327 | 0.743611616 | -7.531978408 |
| **Q9Y530** | OARD1 | 0.002273951 | 3.159260753 | 0.156959064 | 0.88006081 | 0.973827734 | -7.532878248 |
| **P49454** | CENPF | -0.032937219 | 3.147891237 | -1.18325568 | 0.278347127 | 0.676427621 | -7.532991623 |
| **Q04760-1** | GLO1 | 0.009005556 | 3.181237098 | 0.337760728 | 0.746233112 | 0.934974996 | -7.533061129 |
| **Q14562** | DHX8 | -0.015021843 | 3.17341425 | -0.875145763 | 0.412794271 | 0.770775671 | -7.534196824 |
| **Q16630-2** | CPSF6 | 0.012983916 | 3.183010588 | 0.291051962 | 0.780116921 | 0.94472083 | -7.534414437 |
| **O60231** | DHX16 | -0.000976168 | 3.17924282 | -0.051320908 | 0.960617656 | 0.990984017 | -7.535386603 |
| **Q01082-1** | SPTBN1 | 0.010942294 | 3.166891165 | 0.483080826 | 0.644950025 | 0.891546795 | -7.535549082 |
| **O43143** | DHX15 | 0.010606141 | 3.188835647 | 1.175019641 | 0.281403441 | 0.678734164 | -7.537472989 |
| **Q14004-1** | CDK13 | 0.001159361 | 3.178907193 | 0.066092658 | 0.949299104 | 0.989472196 | -7.538677821 |
| **Q08378** | GOLGA3 | 0.015785924 | 3.180039127 | 1.089251606 | 0.314934819 | 0.701837878 | -7.539672692 |
| **Q8WVM8** | SCFD1 | 0.002102568 | 3.181693896 | 0.072034422 | 0.944749783 | 0.98904859 | -7.540221268 |
| **Q5VUA4** | ZNF318 | -0.00440671 | 3.178887332 | -0.334938096 | 0.748264366 | 0.935183323 | -7.541814536 |
| **Q9UJX3-1** | ANAPC7 | -0.006632422 | 3.193985792 | -0.288689417 | 0.781845454 | 0.945329802 | -7.543222203 |

| | | | | | | | |
|---|---|---|---|---|---|---|---|
| **Q6IA86-6** | ELP2 | -0.001567109 | 3.176577339 | -0.05876078 | 0.95491559 | 0.989472196 | -7.544609052 |
| **Q96KP4** | CNDP2 | -0.004521946 | 3.174075828 | -0.204847362 | 0.843984056 | 0.962543524 | -7.545431334 |
| **Q96J01** | THOC3 | 0.011793867 | 3.179485705 | 0.459769886 | 0.660726073 | 0.896934512 | -7.545495904 |
| **Q9NQW7-3** | XPNPEP1 | 0.001227118 | 3.185149211 | 0.028688195 | 0.977977748 | 0.993585863 | -7.545645018 |
| **P18031** | PTPN1 | 0.005107222 | 3.175039174 | 0.085110236 | 0.934746434 | 0.986793807 | -7.547519612 |
| **Q8NFD5-3** | ARID1B | -0.009063353 | 3.17584117 | -0.633979624 | 0.547837514 | 0.846115385 | -7.547721768 |
| **Q96F86** | EDC3 | 0.003081893 | 3.178213274 | 0.121207545 | 0.907203537 | 0.98075504 | -7.548769127 |
| **Q9Y5P4-3** | COL4A3BP | -0.005605099 | 3.175455124 | -0.392087302 | 0.707583637 | 0.918150554 | -7.548918162 |
| **Q9NWK9** | ZNHIT6 | 0.006778147 | 3.243190987 | 0.494920904 | 0.637012 | 0.888997073 | -7.551284435 |
| **P49591** | SARS | 0.03592838 | 3.176176304 | 0.66714214 | 0.52774259 | 0.839849063 | -7.551734824 |
| **P08238** | HSP90AB1 | 0.040620141 | 3.172745052 | 1.165224756 | 0.285075044 | 0.681061455 | -7.552001949 |
| **O14976** | GAK | -0.000878298 | 3.187867341 | -0.03099483 | 0.976207709 | 0.99316605 | -7.552013247 |
| **Q14151** | SAFB2 | 0.012167128 | 3.171782058 | 0.367676257 | 0.72484291 | 0.926648409 | -7.553470858 |
| **Q10570** | CPSF1 | 0.008626779 | 3.185665157 | 0.870346208 | 0.415225909 | 0.77203561 | -7.554459673 |
| **Q6DD88** | ATL3 | 0.008658451 | 3.197627259 | 0.282207585 | 0.786594695 | 0.946749713 | -7.555623654 |
| **P78362-2** | SRPK2 | 0.000502963 | 3.177657611 | 0.016240031 | 0.987532136 | 0.99582925 | -7.555841868 |
| **Q13242** | SRSF9 | -0.023881035 | 3.169448843 | -0.433410506 | 0.678793925 | 0.903807229 | -7.556071947 |
| **Q8WWY3** | PRPF31 | -0.023228418 | 3.173623667 | -0.506000701 | 0.629630394 | 0.886324146 | -7.558342073 |
| **Q8IY67-2** | RAVER1 | 0.006353082 | 3.19889867 | 0.302303204 | 0.771903805 | 0.942488971 | -7.559192211 |
| **Q9Y6K1** | DNMT3A | -0.004769439 | 3.195424585 | -0.228756272 | 0.826117748 | 0.959077751 | -7.559332532 |
| **Q9UL46** | PSME2 | 0.008532147 | 3.172451792 | 0.768158965 | 0.469488763 | 0.805952296 | -7.559477787 |
| **Q9UH65** | SWAP70 | -0.002203266 | 3.184279219 | -0.162006845 | 0.876241691 | 0.972759481 | -7.560323759 |
| **Q9BW27** | NUP85 | 0.003250486 | 3.173536396 | 0.134892756 | 0.896795486 | 0.978757548 | -7.561782552 |
| **P51812** | RPS6KA3 | -0.004982146 | 3.175506449 | -0.212438109 | 0.838300323 | 0.961230299 | -7.562042181 |
| **P05198** | EIF2S1 | -0.012802061 | 3.183746723 | -0.884398947 | 0.408135949 | 0.768299994 | -7.562430194 |
| **Q5VT52** | RPRD2 | 0.006071158 | 3.180498235 | 0.577370924 | 0.583206536 | 0.86495724 | -7.564355179 |
| **Q13126-1** | MTAP | -0.012958752 | 3.178752117 | -0.346811388 | 0.73973486 | 0.931261839 | -7.565471777 |
| **Q9HD20** | ATP13A1 | -0.004576234 | 3.192942194 | -0.268878122 | 0.796392147 | 0.950757673 | -7.567667992 |
| **Q9Y223-2** | GNE | -0.006414534 | 3.189525183 | -0.510802624 | 0.626445441 | 0.885387512 | -7.567768231 |
| **P11413-2** | G6PD | -0.006790187 | 3.17713033 | -0.320075782 | 0.758995354 | 0.9384213 | -7.568643362 |
| **P53618** | COPB1 | -0.00843503 | 3.190749979 | -0.847503825 | 0.426942904 | 0.779764001 | -7.568787987 |
| **O95071** | UBR5 | -0.01388661 | 3.16475204 | -0.960093972 | 0.371494943 | 0.744717822 | -7.570651497 |
| **Q9GZZ1-1** | NAA50 | -0.021354954 | 3.205593496 | -0.502289554 | 0.632097771 | 0.887490806 | -7.570853059 |
| **Q93074-2** | MED12 | -0.005833445 | 3.185600637 | -0.292752908 | 0.778873279 | 0.944053983 | -7.571285359 |
| **Q13107-1** | USP4 | -0.003965271 | 3.199179965 | -0.22062577 | 0.832181418 | 0.960474408 | -7.57262977 |
| **Q76FK4** | NOL8 | -0.010576076 | 3.190780957 | -0.634513082 | 0.547510558 | 0.846115385 | -7.573028586 |
| **Q12802-2** | AKAP13 | 0.002666639 | 3.181344897 | 0.187557995 | 0.856967149 | 0.968855822 | -7.573315258 |
| **Q92608-1** | DOCK2 | 0.009151771 | 3.16513775 | 1.126735098 | 0.299894305 | 0.690882899 | -7.573550046 |
| **Q9BS26** | ERP44 | -0.00262561 | 3.194985727 | -0.126635528 | 0.903072874 | 0.980210603 | -7.573623566 |

| | | | | | | | |
|---|---|---|---|---|---|---|---|
| **Q96K76** | USP47 | -0.012813321 | 3.183673993 | -0.780059524 | 0.46292556 | 0.802800142 | -7.575155574 |
| **P37198** | NUP62 | 0.006990533 | 3.183826173 | 0.304875286 | 0.770030673 | 0.941649022 | -7.577236087 |
| **P52756** | RBM5 | -0.005238618 | 3.159153645 | -0.184857086 | 0.858999822 | 0.968855822 | -7.577941017 |
| **P20618** | PSMB1 | -0.003063638 | 3.179484472 | -0.230010984 | 0.825183132 | 0.959077751 | -7.578025485 |
| **P62312** | LSM 6.00 | -0.003356386 | 3.188599557 | -0.064930655 | 0.95018904 | 0.989472196 | -7.579789737 |
| **P49848-3** | TAF6 | 0.009238475 | 3.175268536 | 0.404998166 | 0.698529645 | 0.913150177 | -7.58059398 |
| **Q12769** | NUP160 | 0.00874248 | 3.182191998 | 0.563961797 | 0.591776319 | 0.86906892 | -7.581773466 |
| **Q13347** | EIF3I | -0.025851732 | 3.19744775 | -0.960022433 | 0.371528342 | 0.744717822 | -7.581847019 |
| **P48637** | GSS | -0.011409314 | 3.170284239 | -0.598387651 | 0.569921009 | 0.855535705 | -7.581993852 |
| **Q9Y6I4-1** | USP3 | 0.00184726 | 3.17771615 | 0.110790462 | 0.91513953 | 0.9828683 | -7.582236016 |
| **P50502** | ST13 | -0.01966165 | 3.171402663 | -0.495052111 | 0.636924321 | 0.888997073 | -7.582410972 |
| **Q5UIP0** | RIF1 | -0.013771944 | 3.176897832 | -1.057505273 | 0.32815086 | 0.713117775 | -7.582850988 |
| **O43847-2** | NRD1; NRDC | -0.013353675 | 3.178067761 | -0.787399645 | 0.458909347 | 0.799919868 | -7.584295666 |
| **Q92667** | AKAP1 | 0.018890379 | 3.199040054 | 0.473586528 | 0.65135206 | 0.894398914 | -7.5848787 |
| **O95232-1** | LUC7L3 | -0.031534787 | 3.200839274 | -0.426583814 | 0.683511648 | 0.905838206 | -7.5851111 |
| **O94804** | STK10 | 0.000228565 | 3.175503589 | 0.018176491 | 0.986045647 | 0.995531556 | -7.585741148 |
| **O60502** | MGEA5 | 0.005014226 | 3.176260734 | 0.311686287 | 0.765078628 | 0.940369694 | -7.585931982 |
| **P16435** | POR | 0.003401986 | 3.18923751 | 0.176628108 | 0.865199976 | 0.9700838 | -7.586440929 |
| **P35250-1** | RFC2 | -0.01779519 | 3.193823041 | -0.430563159 | 0.680759752 | 0.903980599 | -7.58651359 |
| **Q92905** | COPS5 | -0.009952042 | 3.186594335 | -0.511382309 | 0.626061539 | 0.885387512 | -7.587000631 |
| **Q92688** | ANP32B | 0.017698509 | 3.183070987 | 0.82694838 | 0.437690353 | 0.786862931 | -7.587431888 |
| **Q15020** | SART3 | -0.01516988 | 3.17636207 | -0.872814208 | 0.413974211 | 0.771528292 | -7.587593258 |
| **P41240** | CSK | 0.003361544 | 3.18937248 | 0.22590119 | 0.8282456 | 0.959077751 | -7.589413941 |
| **Q86WB0** | ZC3HC1 | -0.007870528 | 3.163026763 | -0.556552984 | 0.596542073 | 0.871378088 | -7.589436857 |
| **Q9UJW0-1** | DCTN4 | 0.001544981 | 3.165114796 | 0.093991552 | 0.927959186 | 0.9859101 | -7.589760265 |
| **P42166** | TMPO | 0.018904344 | 3.190987866 | 0.643348369 | 0.542112958 | 0.843603519 | -7.589775961 |
| **Q9P2D3-1** | HEATR5B | 0.006884467 | 3.180329225 | 0.309391942 | 0.766745444 | 0.940879515 | -7.590638611 |
| **Q15428** | SF3A2 | -0.006808778 | 3.1864539 | -0.266443386 | 0.798186123 | 0.951650153 | -7.590773807 |
| **Q9UQR0** | SCML2 | 0.010098327 | 3.184261616 | 0.548181629 | 0.601953089 | 0.872497827 | -7.591215851 |
| **Q14165** | MLEC | 0.00275622 | 3.198136552 | 0.105791687 | 0.918951531 | 0.983742622 | -7.591334548 |
| **Q01844-5** | EWSR1 | 0.005810229 | 3.168115754 | 0.320936988 | 0.758371929 | 0.9384213 | -7.593567852 |
| **Q96RS6** | NUDCD1 | -0.00718237 | 3.191379017 | -0.544179027 | 0.604549983 | 0.873617609 | -7.595807114 |
| **P62888** | RPL30 | -0.017567815 | 3.187103275 | -0.429761793 | 0.681313509 | 0.903980599 | -7.595824169 |
| **Q8TDY2-1** | RB1CC1 | -0.000638385 | 3.170355049 | -0.048549169 | 0.962742634 | 0.991283985 | -7.599471616 |
| **Q99460** | PSMD1 | 0.01738905 | 3.176372476 | 0.968815175 | 0.367440641 | 0.74226186 | -7.599559324 |
| **Q9UNF0** | PACSIN2 | -0.021292519 | 3.186126093 | -0.60533306 | 0.565570284 | 0.853133834 | -7.599658304 |
| **Q15477** | SKIV2L | -0.009438065 | 3.188331308 | -0.573102325 | 0.585926775 | 0.8653224 | -7.600043205 |
| **P54920** | NAPA | 0.010121088 | 3.18522987 | 0.208366832 | 0.841347495 | 0.962112957 | -7.60019193 |
| **Q8N543** | OGFOD1 | -0.004110368 | 3.194615221 | -0.18499763 | 0.858894021 | 0.968855822 | -7.600529922 |

| | | | | | | | |
|---|---|---|---|---|---|---|---|
| **Q9Y6D5** | ARFGEF2 | -0.00025356 | 3.180905577 | -0.018769046 | 0.985590795 | 0.995531556 | -7.600707872 |
| **Q16222-1** | UAP1 | -0.006488964 | 3.191942697 | -0.351597661 | 0.736307678 | 0.929681266 | -7.601882028 |
| **P45973** | CBX5 | 0.008128686 | 3.165380105 | 0.421526367 | 0.687016673 | 0.907305098 | -7.60205818 |
| **Q92556-1** | ELMO1 | -0.005874611 | 3.166252402 | -0.194619148 | 0.851658631 | 0.966218452 | -7.602725705 |
| **O43660-1** | PLRG1 | 0.016323783 | 3.194969899 | 0.596436909 | 0.571146559 | 0.856724384 | -7.603412269 |
| **P43034** | PAFAH1B1 | 0.004773286 | 3.183948789 | 0.271591951 | 0.794394108 | 0.949806048 | -7.604096964 |
| **Q149N8** | SHPRH | -0.008784598 | 3.196971488 | -0.541187264 | 0.606495127 | 0.874996148 | -7.605989569 |
| **P49711-1** | CTCF | -0.000344492 | 3.215360465 | -0.022424322 | 0.982785098 | 0.994932161 | -7.606408169 |
| **P20839-6** | IMPDH1 | 0.002096827 | 3.189816837 | 0.106914229 | 0.918095289 | 0.983585096 | -7.606418872 |
| **Q96JB2-1** | COG3 | -0.001276621 | 3.173956947 | -0.057086773 | 0.956198345 | 0.989472196 | -7.607293911 |
| **Q5VZ89-7** | DENND4C | 0.001559002 | 3.176643994 | 0.082460097 | 0.936772888 | 0.987232823 | -7.607657601 |
| **Q70J99-3** | UNC13D | -0.004125677 | 3.195707565 | -0.212924568 | 0.837936431 | 0.96111408 | -7.607901588 |
| **Q7Z7A1** | CNTRL | 0.004143442 | 3.187894403 | 0.192425271 | 0.853307087 | 0.967121941 | -7.608003527 |
| **Q05086** | UBE3A | -0.005080454 | 3.185543987 | -0.301473515 | 0.772508386 | 0.942764023 | -7.608529997 |
| **P19525** | EIF2AK2 | -0.008061389 | 3.179416893 | -0.347337914 | 0.739357526 | 0.931081875 | -7.60858327 |
| **P52209** | PGD | -0.020532111 | 3.167988942 | -0.924792562 | 0.388258312 | 0.75788842 | -7.608763431 |
| **Q13596** | SNX1 | -0.005167077 | 3.164315515 | -0.306318637 | 0.768980276 | 0.941649022 | -7.610924673 |
| **P13693** | TPT1 | -0.018497693 | 3.225282575 | -0.892939201 | 0.403871235 | 0.766336431 | -7.611402309 |
| **Q92769** | HDAC2 | 0.007480809 | 3.191924764 | 0.428587074 | 0.682125645 | 0.90445398 | -7.611594657 |
| **Q08379** | GOLGA2 | -0.006155674 | 3.178902569 | -0.308833497 | 0.767151353 | 0.940879515 | -7.611902277 |
| **P52788-1** | SMS | -0.002195964 | 3.185602025 | -0.145754528 | 0.888550375 | 0.976290023 | -7.613107579 |
| **O96019-1** | ACTL6A | -0.003725508 | 3.181177185 | -0.14876761 | 0.886265773 | 0.975708125 | -7.613465298 |
| **Q7Z6E9** | RBBP6 | -0.007170018 | 3.182537269 | -0.661972906 | 0.530843922 | 0.839849063 | -7.614101928 |
| **P07686** | HEXB | 0.001334211 | 3.195104709 | 0.037875941 | 0.970928273 | 0.992540547 | -7.61423218 |
| **Q16666-1** | IFI16 | 0.025551897 | 3.173600212 | 1.093821254 | 0.313068702 | 0.701051533 | -7.617489673 |
| **O60488-1** | ACSL4 | -0.005798345 | 3.176805186 | -0.431422215 | 0.680166368 | 0.903980599 | -7.617923083 |
| **P07355** | ANXA2 | 0.016243561 | 3.178182837 | 0.624486548 | 0.553675858 | 0.848478354 | -7.617943033 |
| **Q9GZS3** | WDR61 | -0.001381344 | 3.164312692 | -0.041114837 | 0.968443807 | 0.992306703 | -7.619996589 |
| **P31943** | HNRNPH1 | 0.011482515 | 3.18638711 | 0.914265655 | 0.393366888 | 0.760194378 | -7.620276615 |
| **Q9H0J9** | PARP12 | 0.001729324 | 3.218698853 | 0.104033781 | 0.920292645 | 0.983949006 | -7.620485717 |
| **P08195-4** | SLC3A2 | -0.006281691 | 3.198972445 | -0.194334495 | 0.851872471 | 0.966280789 | -7.621605701 |
| **P48449-1** | LSS | 0.000667406 | 3.170137107 | 0.033281941 | 0.974452797 | 0.99316605 | -7.622163309 |
| **Q9BY44-1** | EIF2A | -0.012814303 | 3.176404295 | -0.708534147 | 0.503327708 | 0.827688992 | -7.62460332 |
| **Q9Y3R5-1** | DOPEY2 | -0.008246866 | 3.193744375 | -0.434857076 | 0.677796246 | 0.903662615 | -7.627025843 |
| **P54132** | BLM | -0.005168276 | 3.174813459 | -0.290089968 | 0.780820592 | 0.944918194 | -7.628032225 |
| **Q13131-2** | PRKAA1 | 0.019645206 | 3.193918652 | 0.440954781 | 0.673598496 | 0.902333511 | -7.628177415 |
| **Q14008** | CKAP5 | -0.015395996 | 3.176074469 | -1.180887817 | 0.27922293 | 0.676725357 | -7.629291868 |
| **O00487** | PSMD14 | -0.002959573 | 3.187510304 | -0.08595562 | 0.934100112 | 0.986793807 | -7.630122347 |
| **Q9H4E7** | DEF6 | 0.020808902 | 3.186118869 | 0.598413869 | 0.569904549 | 0.855535705 | -7.631114208 |

| | | | | | | | |
|---|---|---|---|---|---|---|---|
| **O43148-2** | RNMT | 0.016366049 | 3.182015339 | 0.40379544 | 0.699370855 | 0.913245694 | -7.631709544 |
| **P06239-3** | LCK | 0.001168585 | 3.166019754 | 0.080783187 | 0.938055418 | 0.987739899 | -7.631878549 |
| **P49750-4** | YLPM1 | -0.002661764 | 3.190042257 | -0.137594828 | 0.894743014 | 0.978625172 | -7.632345578 |
| **Q14258** | TRIM25 | -0.013272273 | 3.195777594 | -0.799620263 | 0.45227683 | 0.795197094 | -7.633213523 |
| **Q9NWT1** | PAK1IP1 | -0.016792741 | 3.191507674 | -0.517892744 | 0.621758648 | 0.883385606 | -7.63380963 |
| **P61204** | ARF3 | -0.020327609 | 3.149489289 | -0.543554554 | 0.604955706 | 0.874044933 | -7.634603764 |
| **P25786-2** | PSMA1 | 0.013014244 | 3.167597698 | 0.865688827 | 0.417595574 | 0.774100988 | -7.635384551 |
| **P11766** | ADH5 | 0.013835835 | 3.167404512 | 0.671530037 | 0.525119077 | 0.837933534 | -7.635395421 |
| **Q9Y266** | NUDC | -0.020856632 | 3.171444899 | -1.13817714 | 0.295423287 | 0.688880164 | -7.635562348 |
| **Q12907** | LMAN2 | 0.009246485 | 3.193674439 | 0.298542653 | 0.77464543 | 0.943227271 | -7.637707007 |
| **Q13619-1** | CUL4A | -0.005384928 | 3.189273617 | -0.403657111 | 0.699467634 | 0.913245694 | -7.637954241 |
| **P40938** | RFC3 | 0.006148419 | 3.18441611 | 0.353808364 | 0.734726916 | 0.92925216 | -7.638067244 |
| **P19367-3** | HK1 | 0.010205072 | 3.195590034 | 0.564455234 | 0.591459688 | 0.868764508 | -7.638106224 |
| **P18754-2** | RCC1 | 0.011204704 | 3.182154699 | 0.779059912 | 0.463474391 | 0.802800142 | -7.639040046 |
| **Q9Y6D6** | ARFGEF1 | -0.00786206 | 3.175149293 | -0.469124504 | 0.654371968 | 0.896376773 | -7.641403604 |
| **P68400** | CSNK2A1 | 0.008680803 | 3.186028935 | 0.43879131 | 0.675086422 | 0.90314827 | -7.64145779 |
| **Q96RU3-2** | FNBP1 | -0.02312535 | 3.181919435 | -0.566864752 | 0.589914933 | 0.868583051 | -7.641460695 |
| **O95155-4** | UBE4B | -0.003989033 | 3.174790359 | -0.273514464 | 0.792979685 | 0.949070183 | -7.64264168 |
| **P12956** | XRCC6 | 0.029299082 | 3.171476018 | 1.217688213 | 0.265872732 | 0.668906358 | -7.642795373 |
| **Q9UNS2** | COPS3 | 0.01158808 | 3.195129981 | 0.611745464 | 0.561571143 | 0.852576894 | -7.642909666 |
| **Q96A65** | EXOC4 | -0.002870487 | 3.201714509 | -0.221892265 | 0.831236044 | 0.960187608 | -7.642942726 |
| **Q9UKY7** | CDV3 | -0.003287968 | 3.18177672 | -0.138987332 | 0.893685622 | 0.978276697 | -7.643997137 |
| **P48506** | GCLC | 0.002550922 | 3.179370718 | 0.07246702 | 0.944418649 | 0.98904859 | -7.644811274 |
| **O00186** | STXBP3 | 0.009259284 | 3.149053496 | 0.24536682 | 0.813770285 | 0.95620584 | -7.645383701 |
| **Q14444-1** | CAPRIN1 | -0.019550465 | 3.191299402 | -0.847755382 | 0.42681257 | 0.779764001 | -7.645820876 |
| **Q12888-2** | TP53BP1 | 0.015201041 | 3.18031688 | 0.949360837 | 0.376531846 | 0.749049177 | -7.647467492 |
| **Q5T8P6** | RBM26 | -0.007734996 | 3.176611881 | -0.317352277 | 0.760968172 | 0.938803125 | -7.648128711 |
| **Q9NUQ9** | FAM49B | -0.000692484 | 3.176833822 | -0.03614044 | 0.972259674 | 0.992928787 | -7.649637788 |
| **P17174** | GOT1 | -0.001208212 | 3.163976224 | -0.076916679 | 0.941013348 | 0.988377491 | -7.651742923 |
| **P78347** | GTF2I | 0.029822535 | 3.176889882 | 0.893007471 | 0.403837277 | 0.766336431 | -7.652355508 |
| **Q9UKJ3-1** | GPATCH8 | 0.001815508 | 3.19215106 | 0.139262471 | 0.893476725 | 0.978276697 | -7.652600464 |
| **O14979-1** | HNRNPDL; HNRPD | -0.006132805 | 3.181992738 | -0.403088895 | 0.699865241 | 0.913245694 | -7.652821707 |
| **Q15648** | MED1 | 0.012221951 | 3.194255228 | 0.65115364 | 0.537372214 | 0.84325938 | -7.653604957 |
| **Q69YN4** | KIAA1429; VIRMA | -0.012525956 | 3.184807348 | -0.52108438 | 0.61965508 | 0.882603014 | -7.654515527 |
| **Q9BV73-1** | CEP250 | -0.007305286 | 3.193154001 | -0.449467265 | 0.667759532 | 0.90001396 | -7.655762499 |
| **P23526-1** | AHCY | -0.011492572 | 3.190996915 | -1.189594328 | 0.276014087 | 0.67488468 | -7.655921011 |
| **Q8NB90-1** | SPATA5 | -0.004135417 | 3.178200221 | -0.306373975 | 0.768940014 | 0.941649022 | -7.65673482 |
| **Q99570** | PIK3R4 | -0.0041099 | 3.173152807 | -0.285488447 | 0.784189554 | 0.946296623 | -7.65687286 |
| **Q9Y6K5** | OAS3 | -0.001764504 | 3.161507697 | -0.125373846 | 0.90403272 | 0.980210603 | -7.658048713 |

| | | | | | | | |
|---|---|---|---|---|---|---|---|
| **O75832** | PSMD10 | -0.000481186 | 3.195534747 | -0.011535063 | 0.99114403 | 0.996915979 | -7.658495948 |
| **Q9Y383** | LUC7L2 | 0.00892423 | 3.165185042 | 0.36518387 | 0.72661516 | 0.92710315 | -7.658594155 |
| **Q96GM5** | SMARCD1 | 0.000824115 | 3.169685804 | 0.050847443 | 0.960980618 | 0.991042447 | -7.661750124 |
| **Q09028** | RBBP4 | 0.008812488 | 3.190864765 | 0.366161717 | 0.725919629 | 0.926870088 | -7.661890767 |
| **Q04446** | GBE1 | -0.003415796 | 3.197512976 | -0.185993406 | 0.858144497 | 0.968855822 | -7.66339859 |
| **Q86WR0** | CCDC25 | -0.003464348 | 3.181182779 | -0.231849223 | 0.823814414 | 0.959077751 | -7.663471614 |
| **Q9NZB2-6** | FAM120A | -0.010671227 | 3.182728868 | -0.493166885 | 0.638184724 | 0.889065271 | -7.663762697 |
| **O94776** | MTA2 | -0.011439597 | 3.182452257 | -0.456987706 | 0.662621793 | 0.897184895 | -7.664499598 |
| **P63244** | GNB2L1; RACK1 | -0.027483111 | 3.194421848 | -1.162913569 | 0.285947246 | 0.681580409 | -7.66548846 |
| **Q9NTK5-1** | OLA1 | -0.007783203 | 3.174338645 | -0.86088617 | 0.42004953 | 0.776300813 | -7.666658787 |
| **Q12768** | KIAA0196; WASHC | 0.003643264 | 3.173009446 | 0.184227336 | 0.859473933 | 0.968855822 | -7.669772233 |
| **P38919** | EIF4A3 | 0.036746692 | 3.186026843 | 0.839570365 | 0.431068094 | 0.782692325 | -7.671931091 |
| **P48739-3** | PITPNB | -0.00273966 | 3.174414673 | -0.113803928 | 0.912842668 | 0.982195964 | -7.674066469 |
| **Q6PCE3** | PGM2L1 | 0.004798651 | 3.172671936 | 0.299061832 | 0.774266713 | 0.943158294 | -7.674360287 |
| **P43686** | PSMC4 | 0.028658866 | 3.186065857 | 0.541737718 | 0.606136979 | 0.874956086 | -7.674951006 |
| **Q9NW08** | POLR3B | 0.002618406 | 3.205902431 | 0.138197028 | 0.894285708 | 0.97851842 | -7.676527128 |
| **Q13561-2** | DCTN2 | 0.001324894 | 3.176686197 | 0.052967651 | 0.95935533 | 0.990513797 | -7.677269905 |
| **Q92900** | UPF1 | -0.006486061 | 3.190571435 | -0.856411468 | 0.422345393 | 0.777824327 | -7.679941289 |
| **P51003** | PAPOLA | -0.001064799 | 3.170822347 | -0.060596216 | 0.953509303 | 0.989472196 | -7.681068258 |
| **P49721** | PSMB2 | -0.006380826 | 3.161984475 | -0.305045697 | 0.769906629 | 0.941649022 | -7.682500275 |
| **P29350-3** | PTPN6 | 0.005056767 | 3.191344317 | 0.315758784 | 0.762123346 | 0.939498834 | -7.683029087 |
| **O95487-3** | SEC24B | -0.002201074 | 3.178977235 | -0.102407384 | 0.921533682 | 0.984164603 | -7.683316393 |
| **Q6ZNB6-1** | NFXL1 | 0.002441033 | 3.200993441 | 0.120219489 | 0.907955783 | 0.98075504 | -7.683577529 |
| **O43390-1** | HNRNPR | 0.014025566 | 3.183381604 | 0.792666117 | 0.456042769 | 0.798084321 | -7.684402441 |
| **P51572-2** | BCAP31 | 0.015413882 | 3.191744828 | 0.494638447 | 0.637200772 | 0.889065271 | -7.684440696 |
| **Q92576-1** | PHF3 | 0.010468855 | 3.175007806 | 0.535419742 | 0.610254794 | 0.877075676 | -7.685295951 |
| **Q13618** | CUL3 | -0.008743588 | 3.181605675 | -0.698619873 | 0.509107366 | 0.82913633 | -7.68595266 |
| **P08621-1** | SNRNP70 | 0.012425698 | 3.199573688 | 0.424812837 | 0.684738049 | 0.905978473 | -7.686233921 |
| **Q9HBL8** | NMRAL1 | 0.000961662 | 3.160034751 | 0.049109514 | 0.962313014 | 0.991283985 | -7.686313409 |
| **O60343-1** | TBC1D4 | -0.003414432 | 3.175628831 | -0.17240843 | 0.868383357 | 0.970949315 | -7.688003372 |
| **Q92619** | HMHA1; ARHGAP4 | 0.014901309 | 3.19540231 | 0.436487847 | 0.676672369 | 0.903662615 | -7.688502675 |
| **Q96QK1** | VPS35 | -0.010882239 | 3.180993753 | -0.546928908 | 0.602765182 | 0.872497827 | -7.689181848 |
| **Q00688** | FKBP3 | 0.038213508 | 3.190364683 | 0.638781038 | 0.544899073 | 0.84497922 | -7.690063107 |
| **Q06203** | PPAT | -0.012016649 | 3.191695571 | -0.446735732 | 0.669630468 | 0.900662615 | -7.691140861 |
| **Q15052** | ARHGEF6 | -0.003898467 | 3.162383104 | -0.225770721 | 0.828342874 | 0.959077751 | -7.693622044 |
| **Q12788** | TBL3 | 0.002212283 | 3.183708003 | 0.162409497 | 0.8759372 | 0.972759481 | -7.693677085 |
| **Q9UBW7** | ZMYM2 | 0.021270349 | 3.194627887 | 0.60045785 | 0.568622121 | 0.854816036 | -7.694336078 |
| **P25787** | PSMA2 | 0.006299347 | 3.187511365 | 0.220587927 | 0.83220967 | 0.960474408 | -7.695633386 |
| **P52597** | HNRNPF | 0.009755321 | 3.191041649 | 0.756132771 | 0.476186055 | 0.809713018 | -7.697811177 |

| | | | | | | | |
|---|---|---|---|---|---|---|---|
| **Q3V6T2-3** | CCDC88A | 0.008691795 | 3.171090142 | 0.363343084 | 0.727925252 | 0.92710315 | -7.698343118 |
| **Q8TD19** | NEK9 | -0.021026899 | 3.19399334 | -0.511679556 | 0.625864734 | 0.885387512 | -7.70005518 |
| **O75152** | ZC3H11A | 0.010268407 | 3.178121293 | 0.383277028 | 0.713791846 | 0.92287652 | -7.700108854 |
| **Q96KP1** | EXOC2 | 0.000361718 | 3.18892223 | 0.019156096 | 0.985293694 | 0.995531556 | -7.701300856 |
| **Q7KZF4** | SND1 | -0.007745008 | 3.181794367 | -1.058067804 | 0.327912829 | 0.713117775 | -7.701342881 |
| **Q86WJ1-1** | CHD1L | -0.004833522 | 3.181242654 | -0.467228635 | 0.655657237 | 0.896497734 | -7.70179317 |
| **O43264** | ZW10 | 0.00148119 | 3.187873405 | 0.036061514 | 0.972320225 | 0.992928787 | -7.702031014 |
| **Q9H3S7** | PTPN23 | -0.001285637 | 3.177075646 | -0.08533189 | 0.934576967 | 0.986793807 | -7.70217158 |
| **Q15185** | PTGES3 | 0.018819004 | 3.192605179 | 0.573601432 | 0.585608332 | 0.8653224 | -7.704401411 |
| **Q9BUJ2-1** | HNRNPUL1 | -0.007140759 | 3.175963378 | -0.500737319 | 0.633131302 | 0.887490806 | -7.706112251 |
| **Q8IYB7** | DIS3L2 | -0.009100673 | 3.194631096 | -0.333813969 | 0.74907393 | 0.935521689 | -7.707525521 |
| **Q5K651** | SAMD9 | 0.002907483 | 3.166625843 | 0.172837709 | 0.86805938 | 0.970868803 | -7.70764978 |
| **Q06124** | PTPN11 | 0.004682797 | 3.182447989 | 0.134246773 | 0.897286297 | 0.978757548 | -7.709015594 |
| **Q68CP9-1** | ARID2 | 0.008164118 | 3.178849975 | 0.575405331 | 0.584458233 | 0.8653224 | -7.709483294 |
| **P07437** | TUBB | 0.013704802 | 3.157961575 | 0.432270736 | 0.679580504 | 0.903893384 | -7.709735912 |
| **Q9H444** | CHMP4B | -0.021160909 | 3.204994656 | -0.283377709 | 0.785736606 | 0.946704945 | -7.711601568 |
| **P78316** | NOP14 | 0.002521249 | 3.202834206 | 0.121663326 | 0.906856568 | 0.98075504 | -7.711911339 |
| **Q9UKM9-1** | RALY | -0.011724209 | 3.173678682 | -0.339326584 | 0.745107219 | 0.934766597 | -7.712057167 |
| **P42345** | MTOR | 0.002858433 | 3.159507127 | 0.190765031 | 0.854555106 | 0.967777825 | -7.714881502 |
| **Q96BP3** | PPWD1 | 0.007784778 | 3.201768253 | 0.212020022 | 0.838613104 | 0.96130156 | -7.714925261 |
| **P00491** | PNP | 0.005287208 | 3.190072023 | 0.248867351 | 0.81117537 | 0.955498404 | -7.716738333 |
| **O15164** | TRIM24 | -0.008348556 | 3.178649492 | -0.363260313 | 0.727984183 | 0.92710315 | -7.718434462 |
| **Q460N5** | PARP14 | -0.028602339 | 3.176560524 | -0.649506366 | 0.538370567 | 0.843524953 | -7.721324813 |
| **P37802-2** | TAGLN2 | 0.020912771 | 3.169933665 | 0.431873618 | 0.679854665 | 0.903893384 | -7.721651015 |
| **Q9BTW9-4** | TBCD | -0.003830502 | 3.183153911 | -0.279778884 | 0.788376771 | 0.947805127 | -7.721692922 |
| **Q8IY21** | DDX60 | -0.008762547 | 3.172596105 | -0.299642313 | 0.77384336 | 0.943158294 | -7.724147546 |
| **P23743-1** | DGKA | -0.005985873 | 3.156834916 | -0.356711869 | 0.732652896 | 0.928164811 | -7.724387898 |
| **P20042** | EIF2S2 | 0.010328558 | 3.168013778 | 0.464931251 | 0.657216411 | 0.896852471 | -7.724757805 |
| **Q8N201** | INTS1 | -0.004013614 | 3.187816547 | -0.366970187 | 0.725344786 | 0.926648409 | -7.725529215 |
| **P38606** | ATP6V1A | 0.006524171 | 3.165137628 | 0.580861761 | 0.580987404 | 0.86380444 | -7.725626064 |
| **Q9Y2I7-1** | PIKFYVE | 0.000904668 | 3.183817035 | 0.064784797 | 0.950300753 | 0.989472196 | -7.726791882 |
| **Q15233** | NONO | -0.02942916 | 3.183112876 | -0.96057933 | 0.371268406 | 0.744717822 | -7.726831441 |
| **Q96I25** | RBM17 | 0.007316291 | 3.179537161 | 0.219126872 | 0.833300659 | 0.96068017 | -7.727693986 |
| **Q99497** | PARK7 | 0.01622391 | 3.197667849 | 0.706926124 | 0.504262187 | 0.828319674 | -7.728981532 |
| **P35606** | COPB2 | -0.019880465 | 3.17938621 | -0.80937136 | 0.447033191 | 0.792204208 | -7.730935523 |
| **P42858** | HTT | 0.006789595 | 3.167151997 | 0.402622962 | 0.700191352 | 0.913245694 | -7.732403817 |
| **P22059** | OSBP | 0.004422321 | 3.179699181 | 0.142859761 | 0.890746354 | 0.977483396 | -7.734063673 |
| **Q9HC38-2** | GLOD4 | -0.00551029 | 3.1673119 | -0.243830367 | 0.814910053 | 0.95620584 | -7.734643529 |
| **O00139-4** | KIF2A | -0.007523203 | 3.176809725 | -0.325575794 | 0.755017295 | 0.937112609 | -7.735884586 |

| | | | | | | | |
|---|---|---|---|---|---|---|---|
| **A2RRP1-1** | NBAS | -0.005754944 | 3.1906398 | -0.554581751 | 0.597813739 | 0.871651708 | -7.737084811 |
| **Q70CQ2** | USP34 | -0.005086696 | 3.174099046 | -0.479698389 | 0.647227093 | 0.89271426 | -7.739320825 |
| **O43149** | ZZEF1 | 0.003276512 | 3.191591207 | 0.275013568 | 0.79187736 | 0.948542293 | -7.740583931 |
| **P23284** | PPIB | 0.00890083 | 3.200677258 | 0.70469032 | 0.505563385 | 0.82913633 | -7.742169252 |
| **Q15437** | SEC23B | -0.002358221 | 3.182697245 | -0.090624256 | 0.930531793 | 0.986707862 | -7.743638662 |
| **P15170-3** | GSPT1 | 0.014430056 | 3.191559739 | 0.826150503 | 0.438111407 | 0.786897076 | -7.743960381 |
| **Q92620** | DHX38 | -0.000558162 | 3.178160953 | -0.049759619 | 0.96181459 | 0.991283985 | -7.744992622 |
| **O43290** | SART1 | -0.007580495 | 3.174374272 | -0.528361714 | 0.614873186 | 0.880352106 | -7.746641948 |
| **Q9NZM3** | ITSN2 | 0.000988016 | 3.19370928 | 0.06567195 | 0.9496213 | 0.989472196 | -7.748011668 |
| **P50395-1** | GDI2 | 0.011208738 | 3.167646528 | 0.759951621 | 0.474052322 | 0.808788313 | -7.748198883 |
| **P00492** | HPRT1 | 0.003482872 | 3.172629698 | 0.205515027 | 0.843483715 | 0.962543524 | -7.748740922 |
| **P40121** | CAPG | 0.006685187 | 3.17864823 | 0.314600184 | 0.762963668 | 0.939644172 | -7.749112103 |
| **P55010** | EIF5 | -0.01057279 | 3.192875827 | -0.507966248 | 0.62832567 | 0.885908173 | -7.750043997 |
| **O14981** | BTAF1 | 0.001733013 | 3.17997224 | 0.117242425 | 0.910222957 | 0.981900848 | -7.750647421 |
| **Q9H4A3-7** | WNK1 | -0.009375687 | 3.182984329 | -0.466967854 | 0.655834128 | 0.896497734 | -7.753055867 |
| **Q13123** | IK | -0.00375273 | 3.170931093 | -0.203640282 | 0.844888827 | 0.962551891 | -7.753466666 |
| **O43719** | HTATSF1 | -0.010695343 | 3.170189512 | -0.693144413 | 0.512317861 | 0.831227449 | -7.754838223 |
| **P22307-1** | SCP2 | -0.009987981 | 3.170023407 | -0.503924554 | 0.6310101 | 0.887123867 | -7.757523943 |
| **Q8NE71-1** | ABCF1 | -0.015553754 | 3.179894232 | -0.599291476 | 0.569353713 | 0.855106852 | -7.757640561 |
| **Q9BZF1-1** | OSBPL8 | -0.006552105 | 3.175205151 | -0.287534708 | 0.782690774 | 0.94563233 | -7.757738939 |
| **Q9HC35** | EML4 | -0.011559159 | 3.171415377 | -0.448945692 | 0.66811658 | 0.90001396 | -7.757878793 |
| **P41227-1** | NAA10 | -0.001744844 | 3.178200968 | -0.120128181 | 0.908025304 | 0.98075504 | -7.759221358 |
| **Q9P016** | THYN1 | 0.002936133 | 3.175901246 | 0.173246736 | 0.867750714 | 0.970868803 | -7.761660981 |
| **Q9P2J5** | LARS | 0.012459867 | 3.183119787 | 0.991273281 | 0.357158503 | 0.734600528 | -7.764472708 |
| **O60763-2** | USO1 | 0.007482351 | 3.176589551 | 0.481853171 | 0.645776008 | 0.892138826 | -7.767535341 |
| **Q13177** | PAK2 | 0.00981576 | 3.179319477 | 0.584214824 | 0.578860498 | 0.861567376 | -7.768309649 |
| **Q9UHX1-2** | PUF60 | 0.020819118 | 3.18556576 | 0.545216189 | 0.60387647 | 0.873076436 | -7.769593019 |
| **P46060** | RANGAP1 | -0.011854299 | 3.191127138 | -0.424258289 | 0.685122287 | 0.906008084 | -7.770706518 |
| **Q14498-1** | RBM39 | -0.008034959 | 3.196901426 | -0.408169387 | 0.696313846 | 0.912062672 | -7.77696619 |
| **P04406-1** | GAPDH | -0.008436331 | 3.177786091 | -0.64201769 | 0.54292377 | 0.843669625 | -7.778696553 |
| **Q9Y618-3** | NCOR2 | -0.003773488 | 3.193133923 | -0.265142153 | 0.799145452 | 0.952055634 | -7.779338823 |
| **Q86XP3-1** | DDX42 | 0.021441232 | 3.188299486 | 0.609205938 | 0.563152898 | 0.852578515 | -7.779509941 |
| **Q99536** | VAT1 | -0.000327921 | 3.190304833 | -0.023038713 | 0.982313532 | 0.994932161 | -7.78131718 |
| **Q96KR1** | ZFR | -0.006318944 | 3.188688211 | -0.391907985 | 0.707709754 | 0.918150554 | -7.782025545 |
| **Q9Y2L5-1** | TRAPPC8 | 0.000760861 | 3.177636124 | 0.056081037 | 0.956969086 | 0.989500907 | -7.784294066 |
| **P49773** | HINT1 | 0.007363045 | 3.178716486 | 0.467498228 | 0.655474394 | 0.896497734 | -7.784388817 |
| **P57772** | EEFSEC | 0.001337024 | 3.192373613 | 0.098307004 | 0.924663565 | 0.985113891 | -7.785643662 |
| **O15355** | PPM1G | -0.002928797 | 3.182044622 | -0.217079727 | 0.834829963 | 0.960912428 | -7.786589506 |
| **Q8N1G2** | CMTR1; FTSJD2 | 0.000810131 | 3.172290334 | 0.072983703 | 0.944023168 | 0.98895887 | -7.788199647 |

| | | | | | | |
|---|---|---|---|---|---|---|
| **Q9ULW0** | TPX2 | 0.003175226 | 3.116314359 | 0.185840528 | 0.858259559 | 0.968855822 | -7.7896622 |
| **O94874-1** | UFL1 | -0.003279868 | 3.16279498 | -0.090802497 | 0.930395594 | 0.986707862 | -7.790196467 |
| **P13796** | LCP1 | 0.026873865 | 3.187995782 | 1.235369867 | 0.259654778 | 0.663330292 | -7.791630517 |
| **P49756-1** | RBM25 | -0.014890153 | 3.179685521 | -0.509186632 | 0.627516309 | 0.885387512 | -7.792694903 |
| **Q9UGU0** | TCF20 | -0.001486623 | 3.186308684 | -0.136189519 | 0.895810367 | 0.978757548 | -7.793452588 |
| **Q9BTC0** | DIDO1 | -0.004613932 | 3.181307703 | -0.53152566 | 0.612800499 | 0.878766193 | -7.795594275 |
| **P07602-3** | PSAP | 0.002847081 | 3.225666151 | 0.104653136 | 0.919820104 | 0.983742622 | -7.796796521 |
| **P60900** | PSMA6 | -0.004411942 | 3.178135726 | -0.214929912 | 0.836436804 | 0.960912428 | -7.796810103 |
| **Q96FV9-1** | THOC1 | -0.00347849 | 3.187130662 | -0.240640693 | 0.817277765 | 0.956656724 | -7.797027273 |
| **Q96DI7** | SNRNP40 | -0.000552984 | 3.176233626 | -0.047973068 | 0.963184348 | 0.991283985 | -7.79898301 |
| **Q15365** | PCBP1 | 0.007880053 | 3.192074545 | 0.50582207 | 0.62974904 | 0.886334123 | -7.799496691 |
| **P52701** | MSH6 | -0.012914634 | 3.183242714 | -0.944658533 | 0.378755032 | 0.750267026 | -7.800486839 |
| **Q8IY18** | SMC5 | -0.0043625 | 3.183156633 | -0.275570267 | 0.791468139 | 0.948542293 | -7.801276702 |
| **P39748** | FEN1 | -1.6E-05 | 3.166224349 | -0.000681292 | 0.999476929 | 0.999602744 | -7.801386941 |
| **Q93009** | USP7 | 0.008797486 | 3.172271153 | 0.933790285 | 0.383931834 | 0.754456977 | -7.801395916 |
| **Q9UI08-2** | EVL | 0.014836745 | 3.170792526 | 0.696450194 | 0.510377969 | 0.829456831 | -7.80237915 |
| **P09972** | ALDOC | -0.017950377 | 3.190768002 | -0.514397678 | 0.624066627 | 0.885078428 | -7.805504939 |
| **P62191** | PSMC1 | -0.008828842 | 3.173887094 | -0.430557252 | 0.680763833 | 0.903980599 | -7.80938781 |
| **P55072** | VCP | 0.03775274 | 3.166774199 | 1.004628829 | 0.351151511 | 0.732449134 | -7.811569978 |
| **Q00534** | CDK6 | -0.011276231 | 3.184111123 | -0.415293908 | 0.691347587 | 0.909547372 | -7.816265149 |
| **Q8IX12** | CCAR1 | 0.015994235 | 3.17473869 | 0.556994212 | 0.596257641 | 0.871378088 | -7.816697889 |
| **Q9Y4E8** | USP15 | 0.003852258 | 3.189381438 | 0.22013811 | 0.832545512 | 0.960582935 | -7.819317315 |
| **Q9Y263** | PLAA | -0.004902974 | 3.179226852 | -0.158760968 | 0.878697095 | 0.97338977 | -7.82243074 |
| **Q9H4A4** | RNPEP | 0.001678217 | 3.174513949 | 0.118944144 | 0.908926905 | 0.981305104 | -7.823305857 |
| **P41250** | GARS | 0.001211152 | 3.181128898 | 0.064841949 | 0.95025698 | 0.989472196 | -7.82422399 |
| **Q9NZJ4-1** | SACS | 0.002673763 | 3.20044491 | 0.228508183 | 0.826302583 | 0.959077751 | -7.826239799 |
| **Q96SB8** | SMC6 | 0.001680944 | 3.168782607 | 0.141511402 | 0.891769582 | 0.977930894 | -7.827016083 |
| **O15294** | OGT | 0.004875806 | 3.185150523 | 0.189984187 | 0.855142232 | 0.96809704 | -7.82838327 |
| **Q14C86-6** | GAPVD1 | 0.00259361 | 3.184938448 | 0.149328693 | 0.885840474 | 0.975708125 | -7.828448564 |
| **O00471** | EXOC5 | -0.000144145 | 3.186503067 | -0.012734338 | 0.99022335 | 0.996494555 | -7.828631143 |
| **Q96F07-2** | CYFIP2 | -0.018796687 | 3.171502311 | -0.517176295 | 0.622231381 | 0.883385606 | -7.830894773 |
| **P29401-2** | TKT | 0.034105279 | 3.182857334 | 0.966221314 | 0.368642876 | 0.743611616 | -7.831252102 |
| **Q7Z3B4** | NUP54 | 0.000607675 | 3.162509774 | 0.021794643 | 0.983268406 | 0.995280405 | -7.831848351 |
| **Q9NYU2-1** | UGGT1 | 0.013463038 | 3.188565845 | 0.696026656 | 0.510626242 | 0.829637115 | -7.832189667 |
| **Q99986** | VRK1 | -0.00506569 | 3.172814311 | -0.23186411 | 0.823803331 | 0.959077751 | -7.832291763 |
| **Q9H2G2-2** | SLK | 0.005611564 | 3.175819763 | 0.557652235 | 0.5958336 | 0.871378088 | -7.833094148 |
| **Q15717** | ELAVL1 | -0.00294892 | 3.175092993 | -0.156103071 | 0.880708796 | 0.974002142 | -7.833126834 |
| **P25398** | RPS12 | 0.007912493 | 3.200703357 | 0.166141933 | 0.873115779 | 0.97237605 | -7.838080914 |
| **Q8IWX8** | CHERP | 0.002366568 | 3.162089087 | 0.050998133 | 0.960865096 | 0.991042447 | -7.840303774 |

| | | | | | | | |
|---|---|---|---|---|---|---|---|
| **P00338-3** | LDHA | 0.015779964 | 3.182797013 | 1.225026473 | 0.263276811 | 0.666686026 | -7.840518965 |
| **O75376-1** | NCOR1 | 0.003695264 | 3.188821184 | 0.245121781 | 0.813952027 | 0.95620584 | -7.84151895 |
| **P49915** | GMPS | -0.022914291 | 3.183797678 | -0.63793483 | 0.545416239 | 0.845033545 | -7.843273252 |
| **Q92945** | KHSRP | 0.009675696 | 3.184050759 | 0.660487139 | 0.53173745 | 0.839849063 | -7.846820991 |
| **P43490** | NAMPT | -0.017495032 | 3.186452833 | -0.577504212 | 0.583121715 | 0.86495724 | -7.850088951 |
| **Q92835** | INPP5D | 0.001381746 | 3.192551931 | 0.08248116 | 0.936756779 | 0.987232823 | -7.850255437 |
| **Q9NWH9-1** | SLTM | -0.003836903 | 3.179320375 | -0.153874871 | 0.882396011 | 0.97451158 | -7.853798013 |
| **P0DMV8** | HSPA1B; HSPA1A | -0.003156215 | 3.197522202 | -0.150282242 | 0.885117781 | 0.975708125 | -7.854740264 |
| **P42224-1** | STAT1 | -0.02113558 | 3.189983587 | -0.435407361 | 0.677416906 | 0.903662615 | -7.856615903 |
| **Q96L91** | EP400 | 0.004408095 | 3.187711737 | 0.298102829 | 0.774966313 | 0.943473392 | -7.858743699 |
| **P12268** | IMPDH2 | -0.015073259 | 3.183668127 | -0.759001642 | 0.474582499 | 0.809054593 | -7.859107054 |
| **Q9NY33** | DPP3 | -0.002929181 | 3.179388561 | -0.113087054 | 0.913388987 | 0.982219313 | -7.859252218 |
| **Q16531** | DDB1 | -0.011826731 | 3.188732604 | -0.910183584 | 0.395361456 | 0.761244496 | -7.863257065 |
| **P41091** | EIF2S3 | -0.017352186 | 3.192426765 | -0.462109428 | 0.659134055 | 0.896852471 | -7.867156434 |
| **Q15459** | SF3A1 | -0.015754263 | 3.169696286 | -0.696556788 | 0.510315498 | 0.829456831 | -7.867571245 |
| **Q7L1Q6-3** | BZW1 | -0.005768741 | 3.188185915 | -0.172917723 | 0.867998996 | 0.970868803 | -7.868704568 |
| **Q15021** | NCAPD2 | -0.005495755 | 3.187604076 | -0.521618157 | 0.619303652 | 0.882418852 | -7.8718109 |
| **O43172-1** | PRPF4 | -0.000680304 | 3.164267421 | -0.059796337 | 0.954122138 | 0.989472196 | -7.874222359 |
| **Q07955-1** | SRSF1 | 0.003912608 | 3.176215579 | 0.086998097 | 0.933303182 | 0.986793807 | -7.875146679 |
| **Q9Y5B9** | SUPT16H | -0.010394862 | 3.178337242 | -0.818617522 | 0.442100988 | 0.789324124 | -7.875476724 |
| **Q15643** | TRIP11 | -0.00175865 | 3.189205044 | -0.075822006 | 0.941850974 | 0.988377491 | -7.87629348 |
| **Q709C8** | VPS13C | -0.010844914 | 3.179428403 | -0.728608038 | 0.49175826 | 0.820113219 | -7.877042703 |
| **P35998** | PSMC2 | 0.011277368 | 3.17811547 | 0.398207554 | 0.703285149 | 0.915549813 | -7.878231202 |
| **Q9Y5K6** | CD2AP | -0.004149897 | 3.188344057 | -0.187403414 | 0.857083453 | 0.968855822 | -7.881782652 |
| **Q6P179** | ERAP2 | -0.007653126 | 3.19733632 | -0.374441869 | 0.720041401 | 0.925644897 | -7.885686883 |
| **O60884** | DNAJA2 | 0.006641204 | 3.209932998 | 0.16841565 | 0.871398015 | 0.971931713 | -7.88968326 |
| **Q15075** | EEA1 | 0.003628717 | 3.181473754 | 0.231576201 | 0.824017658 | 0.959077751 | -7.893338568 |
| **O15144** | ARPC2 | 0.007404588 | 3.194781397 | 0.284964377 | 0.784573568 | 0.946344035 | -7.893569029 |
| **P14317** | HCLS1 | 0.003518155 | 3.183241418 | 0.084979326 | 0.934846523 | 0.986793807 | -7.898022501 |
| **P60842** | EIF4A1 | -0.008098456 | 3.198276079 | -0.546413409 | 0.60309954 | 0.872497827 | -7.89907408 |
| **Q86U86-1** | PBRM1 | -0.007050659 | 3.181772858 | -0.721115638 | 0.496055575 | 0.822932915 | -7.900588404 |
| **Q08945** | SSRP1 | -0.003824912 | 3.174837272 | -0.364251916 | 0.72727831 | 0.92710315 | -7.903283026 |
| **P30101** | PDIA3 | 0.024300036 | 3.209582884 | 0.8172504 | 0.442827806 | 0.789629405 | -7.909914643 |
| **P50570-1** | DNM2 | 0.005477856 | 3.190295788 | 0.550030491 | 0.600755663 | 0.87241889 | -7.909928458 |
| **Q9UBL3-1** | ASH2L | -0.00075782 | 3.179012986 | -0.05354142 | 0.958915532 | 0.990326778 | -7.911474103 |
| **O00410-3** | IPO5 | -0.017895014 | 3.176730014 | -0.760982997 | 0.473477179 | 0.808788313 | -7.913954126 |
| **Q96T23** | RSF1 | -0.002228057 | 3.177616964 | -0.07204839 | 0.944739091 | 0.98904859 | -7.915281998 |
| **P18669** | PGAM1; LOC64357 | 0.03516125 | 3.192098126 | 0.660180425 | 0.531922024 | 0.839849063 | -7.919126858 |
| **Q9BYW2** | SETD2 | -0.001321353 | 3.181615642 | -0.153567147 | 0.882629075 | 0.974633495 | -7.923414794 |

| | | | | | | | |
|---|---|---|---|---|---|---|---|
| **P28838** | LAP3 | -0.001071696 | 3.180545139 | -0.074155816 | 0.943126069 | 0.988738927 | -7.92518199 |
| **Q14978-2** | NOLC1 | 0.013515363 | 3.179245744 | 0.673410665 | 0.523997203 | 0.837176492 | -7.925299011 |
| **P49790-3** | NUP153 | -0.000878397 | 3.187260313 | -0.060062675 | 0.953918077 | 0.989472196 | -7.927149127 |
| **Q8NFC6** | BOD1L1 | 0.008446269 | 3.163413544 | 0.491805943 | 0.639095418 | 0.88909352 | -7.928689067 |
| **Q8N3C0** | ASCC3 | -0.00910041 | 3.183237459 | -0.496790124 | 0.635763498 | 0.888277483 | -7.928727832 |
| **O95602** | POLR1A | -0.004315456 | 3.197592417 | -0.3575068 | 0.732085486 | 0.928164811 | -7.92911478 |
| **Q9Y520-4** | PRRC2C | -0.007313553 | 3.201874584 | -0.593608059 | 0.572926553 | 0.858139449 | -7.930792411 |
| **P14625** | HSP90B1 | 0.031382623 | 3.212740332 | 0.992037599 | 0.356812569 | 0.734600528 | -7.931489626 |
| **O95466-2** | FMNL1 | 0.003907061 | 3.170833266 | 0.19422222 | 0.851956819 | 0.966280789 | -7.931711295 |
| **Q15393-1** | SF3B3 | -0.011614701 | 3.170850772 | -0.845379789 | 0.428044532 | 0.780598411 | -7.932921122 |
| **P48147** | PREP | 0.004504025 | 3.1895186 | 0.243031543 | 0.815502827 | 0.95624343 | -7.933830533 |
| **P43487** | RANBP1 | -0.008437952 | 3.170857534 | -0.205871531 | 0.843216588 | 0.962543524 | -7.93409477 |
| **Q9NSD9** | FARSB | -0.001914807 | 3.171424768 | -0.167914549 | 0.871776526 | 0.971931713 | -7.938137844 |
| **P11216** | PYGB | -0.003378115 | 3.178555659 | -0.265847865 | 0.798625122 | 0.951999489 | -7.938403766 |
| **Q13435** | SF3B2 | 0.012785271 | 3.173827415 | 0.547389053 | 0.602466816 | 0.872497827 | -7.939695195 |
| **O43175** | PHGDH | -0.014779634 | 3.18716627 | -0.748310144 | 0.480577274 | 0.81235548 | -7.943169822 |
| **Q9Y4L1** | HYOU1 | -0.007965996 | 3.205478478 | -0.451886028 | 0.66610497 | 0.89988165 | -7.943574267 |
| **Q9BUQ8** | DDX23 | 0.005480127 | 3.173208836 | 0.283325773 | 0.785774685 | 0.946704945 | -7.944705035 |
| **P52948-1** | NUP98 | 0.005601424 | 3.180444171 | 0.447388169 | 0.669183355 | 0.90021364 | -7.946163247 |
| **Q9NTJ3-1** | SMC4 | -0.011602429 | 3.187089121 | -0.662957735 | 0.530252177 | 0.839849063 | -7.948239708 |
| **O15067** | PFAS | -0.001807511 | 3.185784963 | -0.121239516 | 0.907179198 | 0.98075504 | -7.949940221 |
| **O14646** | CHD1 | -0.010592497 | 3.180811526 | -0.424529576 | 0.684934303 | 0.905978473 | -7.950798095 |
| **Q7L014** | DDX46 | -0.015279449 | 3.160411921 | -0.725612517 | 0.49347337 | 0.821410327 | -7.959855042 |
| **Q9UIA9** | XPO7 | 0.005526636 | 3.16276249 | 0.218294814 | 0.833922147 | 0.96068017 | -7.963899802 |
| **O75592-1** | MYCBP2 | 0.010808141 | 3.164388498 | 0.724073233 | 0.494356251 | 0.821686278 | -7.964744042 |
| **P35659-1** | DEK | 0.000969656 | 3.173922105 | 0.039923988 | 0.969357232 | 0.992469575 | -7.965994371 |
| **Q14683** | SMC1A | 0.015588311 | 3.165865921 | 0.87895793 | 0.410870378 | 0.769906516 | -7.980916609 |
| **P62333** | PSMC6 | 0.000508181 | 3.176995927 | 0.025422539 | 0.980483939 | 0.994229224 | -7.981752731 |
| **Q16555-1** | DPYSL2 | 0.011462068 | 3.167768351 | 0.57448323 | 0.585045966 | 0.8653224 | -7.988726151 |
| **P23921** | RRM1 | -0.009637513 | 3.156778552 | -0.579865737 | 0.581620076 | 0.8642176 | -7.992346854 |
| **P49368-1** | CCT3 | 0.015662339 | 3.183345285 | 0.88075987 | 0.409963301 | 0.769831818 | -7.994273869 |
| **P15121** | AKR1B1 | -0.009392822 | 3.169982812 | -0.50072415 | 0.633140074 | 0.887490806 | -7.999661566 |
| **P08575-1** | PTPRC | -0.004680774 | 3.150290293 | -0.2102528 | 0.839935558 | 0.961392936 | -8.005269132 |
| **P54577** | YARS | 0.02871736 | 3.196729958 | 0.684412414 | 0.51746494 | 0.833280927 | -8.005920749 |
| **O14980** | XPO1 | 0.008555204 | 3.179940927 | 0.746214 | 0.481758599 | 0.81235548 | -8.008724946 |
| **Q86VP6-1** | CAND1 | -0.007515242 | 3.172926628 | -0.629933161 | 0.550321492 | 0.846853429 | -8.008915525 |
| **O75717** | WDHD1 | -0.004409096 | 3.176129055 | -0.314861692 | 0.762773968 | 0.939644172 | -8.009664526 |
| **Q9Y265** | RUVBL1 | 0.004630648 | 3.168708045 | 0.433732812 | 0.678571574 | 0.903662615 | -8.014523517 |
| **Q29RF7-1** | PDS5A | -0.00662138 | 3.180523437 | -0.412670599 | 0.693174298 | 0.909991705 | -8.017289207 |

| | | | | | | | |
|---|---|---|---|---|---|---|---|
| **A0AVT1-1** | UBA6 | -0.004323348 | 3.176703866 | -0.362085052 | 0.728821167 | 0.927586142 | -8.01986535 |
| **Q9Y3I0** | C22orf28; RTCB | -0.011634227 | 3.173494297 | -0.402786422 | 0.700076937 | 0.913245694 | -8.029228038 |
| **Q92841** | DDX17 | -0.00789331 | 3.203080347 | -0.359226952 | 0.730858295 | 0.92815165 | -8.030321497 |
| **P49792** | RANBP2 | -0.007553914 | 3.174467084 | -0.86522595 | 0.417831626 | 0.774192321 | -8.034969233 |
| **P22087** | FBL | 0.003098294 | 3.177450989 | 0.056408523 | 0.956718113 | 0.989472196 | -8.040067204 |
| **P31939** | ATIC | 0.005585412 | 3.174893096 | 0.55333455 | 0.598619115 | 0.871871293 | -8.048520462 |
| **Q9H2P0** | ADNP | -0.001765993 | 3.182228519 | -0.213301116 | 0.837654786 | 0.96111408 | -8.048869581 |
| **Q8N1F7-1** | NUP93 | -0.015941774 | 3.171109288 | -0.531768747 | 0.612641413 | 0.878766193 | -8.048989566 |
| **Q14152** | EIF3A | 0.024149214 | 3.19664209 | 0.775239321 | 0.465576222 | 0.803432467 | -8.051428504 |
| **Q99832** | CCT7 | 0.005561152 | 3.186647122 | 0.777650273 | 0.464249112 | 0.803074528 | -8.058055535 |
| **Q96PK6-1** | RBM14; RBM14-RE | 0.005819674 | 3.210263558 | 0.101748535 | 0.922036492 | 0.984164603 | -8.058138897 |
| **P43246-1** | MSH2 | -0.002175201 | 3.176349462 | -0.174660594 | 0.866683959 | 0.970722967 | -8.061565099 |
| **P35573-1** | AGL | -0.001094165 | 3.186081924 | -0.097000697 | 0.925660997 | 0.985476218 | -8.064575015 |
| **O43707** | ACTN4 | 0.002808047 | 3.182023152 | 0.288489615 | 0.781991698 | 0.945362757 | -8.071015232 |
| **Q9NR09** | BIRC6 | -0.005203829 | 3.191053281 | -0.324184621 | 0.756022746 | 0.937424764 | -8.071975296 |
| **P33176** | KIF5B | 0.002184672 | 3.175245783 | 0.154091592 | 0.882231878 | 0.97451158 | -8.076813379 |
| **O95347-1** | SMC2 | 0.008437931 | 3.186167251 | 0.443338886 | 0.671960672 | 0.901355316 | -8.077745414 |
| **P00813** | ADA | 0.020595751 | 3.177374985 | 0.786652612 | 0.459316978 | 0.800279253 | -8.077883932 |
| **P41252** | IARS | -0.006082713 | 3.179082412 | -0.573292466 | 0.585805448 | 0.8653224 | -8.088330175 |
| **P35579-1** | MYH9 | 0.032438116 | 3.187170287 | 1.210163001 | 0.268557512 | 0.67097152 | -8.098514995 |
| **Q92614-1** | MYO18A | -0.003444786 | 3.186887515 | -0.194653672 | 0.851632696 | 0.966218452 | -8.099870816 |
| **P17980** | PSMC3 | -0.002591822 | 3.177729731 | -0.193171004 | 0.852746661 | 0.96682982 | -8.101624997 |
| **Q15046-2** | KARS | -0.007130256 | 3.188091882 | -0.35008589 | 0.737389471 | 0.929681266 | -8.108063264 |
| **P53621-2** | COPA | -0.00996938 | 3.183150544 | -0.654650436 | 0.535256777 | 0.840770085 | -8.108267077 |
| **P22102-1** | GART | 0.01228839 | 3.174520365 | 0.838520012 | 0.431616403 | 0.783036167 | -8.11107954 |
| **P78527** | PRKDC | -0.015949077 | 3.177977068 | -1.301336047 | 0.237553447 | 0.644090219 | -8.116340275 |
| **A6NHR9-1** | SMCHD1 | 0.003908865 | 3.177199184 | 0.272889152 | 0.793439644 | 0.949122853 | -8.118825582 |
| **P17987** | TCP1 | 0.017260298 | 3.18540845 | 0.622938158 | 0.554631729 | 0.849209691 | -8.12065228 |
| **P14868** | DARS | 0.000406879 | 3.186150527 | 0.023112596 | 0.982256824 | 0.994932161 | -8.121132626 |
| **Q92621** | NUP205 | -0.009357762 | 3.200764253 | -0.281588408 | 0.787048889 | 0.947114226 | -8.123434046 |
| **Q9Y230** | RUVBL2 | -0.007414957 | 3.165467778 | -0.246159372 | 0.813182548 | 0.956167049 | -8.129264075 |
| **Q96T58** | SPEN | 0.001147445 | 3.187631397 | 0.137890009 | 0.894518851 | 0.97851842 | -8.139474666 |
| **P26639-2** | TARS | 0.000216212 | 3.172286413 | 0.013580839 | 0.989573503 | 0.996219143 | -8.141891036 |
| **Q01518-1** | CAP1 | -0.010935151 | 3.187322469 | -0.342261891 | 0.742998484 | 0.933447653 | -8.147349915 |
| **Q14566** | MCM6 | -0.010607992 | 3.167551017 | -0.460367735 | 0.660319065 | 0.896934512 | -8.165841136 |
| **P26640** | VARS; VARS2 | -0.003315116 | 3.173473796 | -0.091735794 | 0.929682478 | 0.986707862 | -8.174312914 |
| **P17844** | DDX5 | -0.011053564 | 3.202574131 | -0.467678627 | 0.655352058 | 0.896497734 | -8.174899031 |
| **Q7Z6Z7** | HUWE1 | 0.007857792 | 3.185984482 | 0.537492084 | 0.608902415 | 0.876559102 | -8.178441965 |
| **P22234** | PAICS | -0.000464325 | 3.194802932 | -0.013209977 | 0.989858207 | 0.996253288 | -8.184184144 |

| | | | | | | | |
|---|---|---|---|---|---|---|---|
| **Q02880-2** | TOP2B | 0.008520933 | 3.181887022 | 0.607845107 | 0.564001598 | 0.852578515 | -8.18999676 |
| **P24928** | POLR2A | 0.005434781 | 3.185705354 | 0.301342705 | 0.77260372 | 0.942764023 | -8.191339283 |
| **P29144** | TPP2 | 0.008298397 | 3.188262069 | 0.483902153 | 0.644397729 | 0.891474831 | -8.200030172 |
| **P11388-4** | TOP2A | 0.006289291 | 3.20826092 | 0.17687329 | 0.865015091 | 0.9700838 | -8.209948993 |
| **P49588** | AARS | -0.002686205 | 3.174614424 | -0.185577263 | 0.85845771 | 0.968855822 | -8.213436703 |
| **Q9UQE7** | SMC3 | 0.011575292 | 3.174080161 | 0.439934397 | 0.674300067 | 0.903121044 | -8.215871983 |
| **P22314** | UBA1 | 0.007565458 | 3.168606892 | 0.545094135 | 0.603955707 | 0.873076436 | -8.232683883 |
| **O75083** | WDR1 | -0.00616169 | 3.180980496 | -0.445189291 | 0.670690822 | 0.901175787 | -8.245939871 |
| **P07814** | EPRS | 0.00419929 | 3.171293918 | 0.533125954 | 0.61175362 | 0.878435299 | -8.248916217 |
| **Q02790** | FKBP4 | 0.000263151 | 3.194745445 | 0.010431345 | 0.991991366 | 0.997515682 | -8.259113347 |
| **O60841** | EIF5B | 0.00060484 | 3.17739378 | 0.055150767 | 0.957682039 | 0.989567408 | -8.264322239 |
| **P48643** | CCT5 | 0.018904161 | 3.190961792 | 0.614757963 | 0.55969827 | 0.851665881 | -8.269545929 |
| **P52566** | ARHGDIB | -0.003673925 | 3.187825827 | -0.273250802 | 0.793173615 | 0.949070183 | -8.270451407 |
| **P49736** | MCM2 | -0.000698479 | 3.175218512 | -0.032559901 | 0.975006806 | 0.99316605 | -8.303989742 |
| **O75533-1** | SF3B1 | -0.002501706 | 3.17105889 | -0.323411245 | 0.756581916 | 0.937613995 | -8.305167992 |
| **Q9Y4A5** | TRRAP | -0.003309167 | 3.177994569 | -0.243555356 | 0.815114112 | 0.956212145 | -8.308994781 |
| **P12270** | TPR | 0.005563826 | 3.167917908 | 0.478181949 | 0.648249305 | 0.893481222 | -8.309617607 |
| **Q5T4S7-1** | UBR4 | 0.002294292 | 3.174797752 | 0.305520063 | 0.769561372 | 0.941649022 | -8.311904934 |
| **P11586** | MTHFD1 | -0.012158465 | 3.18219444 | -0.484263597 | 0.644154756 | 0.891474831 | -8.312782435 |
| **P35580-4** | MYH10 | 0.001123973 | 3.181542763 | 0.044663966 | 0.96572181 | 0.992001409 | -8.340512514 |
| **P78371-1** | CCT2 | 0.005501037 | 3.185131754 | 0.305124252 | 0.769849451 | 0.941649022 | -8.3489031 |
| **P06744** | GPI | 0.011793657 | 3.195010455 | 0.595199286 | 0.571924904 | 0.857347804 | -8.375815801 |
| **P49321** | NASP | 0.004164103 | 3.185317219 | 0.121808149 | 0.906746324 | 0.98075504 | -8.481807752 |
| **P31948-2** | STIP1 | -8.55E-05 | 3.202846217 | -0.003851712 | 0.997042806 | 0.998480498 | -8.513020752 |
| **P08670** | VIM | -0.002723531 | 3.196188071 | -0.285986063 | 0.783824984 | 0.946296623 | -8.522945469 |
| **P12004** | PCNA | 0.004176672 | 3.16566523 | 0.356727246 | 0.732641918 | 0.928164811 | -8.536316581 |
| **Q8WXH0-2** | SYNE2 | 0.010204367 | 3.190925066 | 0.384120004 | 0.713196805 | 0.922407393 | -8.541262758 |
| **P21333** | FLNA | -0.00419268 | 3.182053283 | -0.501335207 | 0.632733102 | 0.887490806 | -8.575645255 |
| **Q9Y490** | TLN1 | -0.000176788 | 3.182479924 | -0.00965949 | 0.992583933 | 0.99756865 | -8.615347534 |
| **P11142-1** | HSPA8 | 0.009407834 | 3.209034449 | 0.847442181 | 0.426974846 | 0.779764001 | -8.703961396 |
